# Biosynthetic characterization and combinatorial biocatalysis of the cysteine protease inhibitor E-64

**DOI:** 10.1101/2024.10.09.617497

**Authors:** Mengting Liu, Xin Zang, Niko W. Vlahakis, Jose A. Rodriguez, Masao Ohashi, Yi Tang

## Abstract

E-64 is an irreversible and selective cysteine protease inhibitor prominently used in chemical biology and drug discovery. In this work, we uncovered and characterized the NRPS-independent pathway responsible for biosynthesis of E-64, which is widely conserved in fungi. Heterologous reconstitution and biochemical assays show the pathway starts with epoxidation of fumaric acid to the warhead (2*S*,3*S*)-*trans*-epoxysuccinic acid with an α-ketoglutarate (αKG)/Fe(II)-dependent oxygenase, followed by successive condensation with an l-amino acid by an ATP-grasp enzyme, and with an amine by the first characterized amide bond synthetase from fungi. Both amide bond-forming enzymes displayed significant biocatalytic potential, including scalability, stereoselectivity towards the warhead and broader substrate scopes in forming the amide bonds. Combinatorial biocatalysis with the two amide-bond forming enzymes generated a library of cysteine protease inhibitors and led to more potent analogs towards cathepsin B. In addition, preparative synthesis of clinically relevant cysteine protease inhibitors was accomplished from a single reaction mixture. Our work highlights the importance of biosynthetic investigation for enzyme discovery and the potential of amide bond-forming enzymes as biocatalysts for a library synthesis of small molecules.

## Main

E-64 (**1**) is a fungal natural product that has played prominent roles in drug discovery and chemical biology (**Fig. 1a**)^1–13^, as hundreds of studies have used **1** as a chemical probe (https://pubchem.ncbi.nlm.nih.gov/compound/123985#section=Literature). Isolated from *Aspergillus japonicus* TPR-64 in 1978, **1** is the first *trans*-epoxysuccinic acid (***t*-ES**)-based irreversible, potent, and selective inhibitor against cysteine proteases such as papain, calpain, and cysteine cathepsins (**Fig. 1a**)^5,14,15^. Cysteine proteases are ubiquitously conserved in all kingdoms of life, and are involved in multifaceted physiological roles such as apoptosis, inflammation, and immune responses^11,16,17^. As a result, cysteine proteases have been a potential drug target for multiple diseases including cancer, neurodegenerative disorders, muscular dystrophy, and infectious diseases^2,3,12,13,17–19^. Upon binding to a cysteine protease, the electrophilic ***t*-ES** warhead in **1** is covalently captured by the thiolate of the catalytic cysteine (**Fig. 1a**)^15,20^, a feature that led to the development of probes based on **1** for activity-based protein profiling (ABPP)^6–8,10^. Numerous biosynthetic variants of **1** have been isolated from many fungi such as *Aspergillus oryzae* and *Penicillium citrinum* (**Supplementary Fig. 1**)^6,21–24^. Extensive synthetic efforts have resulted in a variety of analogs such as CA-074^25^, CLIK-148^26^, and NYC-488^27^ that show selectivity towards cathepsin B, cathepsin L, and calpain, respectively (**Fig. 1b** and **Supplementary Fig. 1**). E-64d (Loxistatin), a prodrug for E-64c (**2**, Loxistatin acid) was in Phase III trial for treatment of muscular dystrophy (**Fig. 1b**)^2^, and was also repurposed for treating viral infections such as COVID caused by SARS-CoV-2^1,28^. Despite such decorated track record of **1**, the enzymes responsible for formation of **1** have remained elusive.

**Fig. 1.**
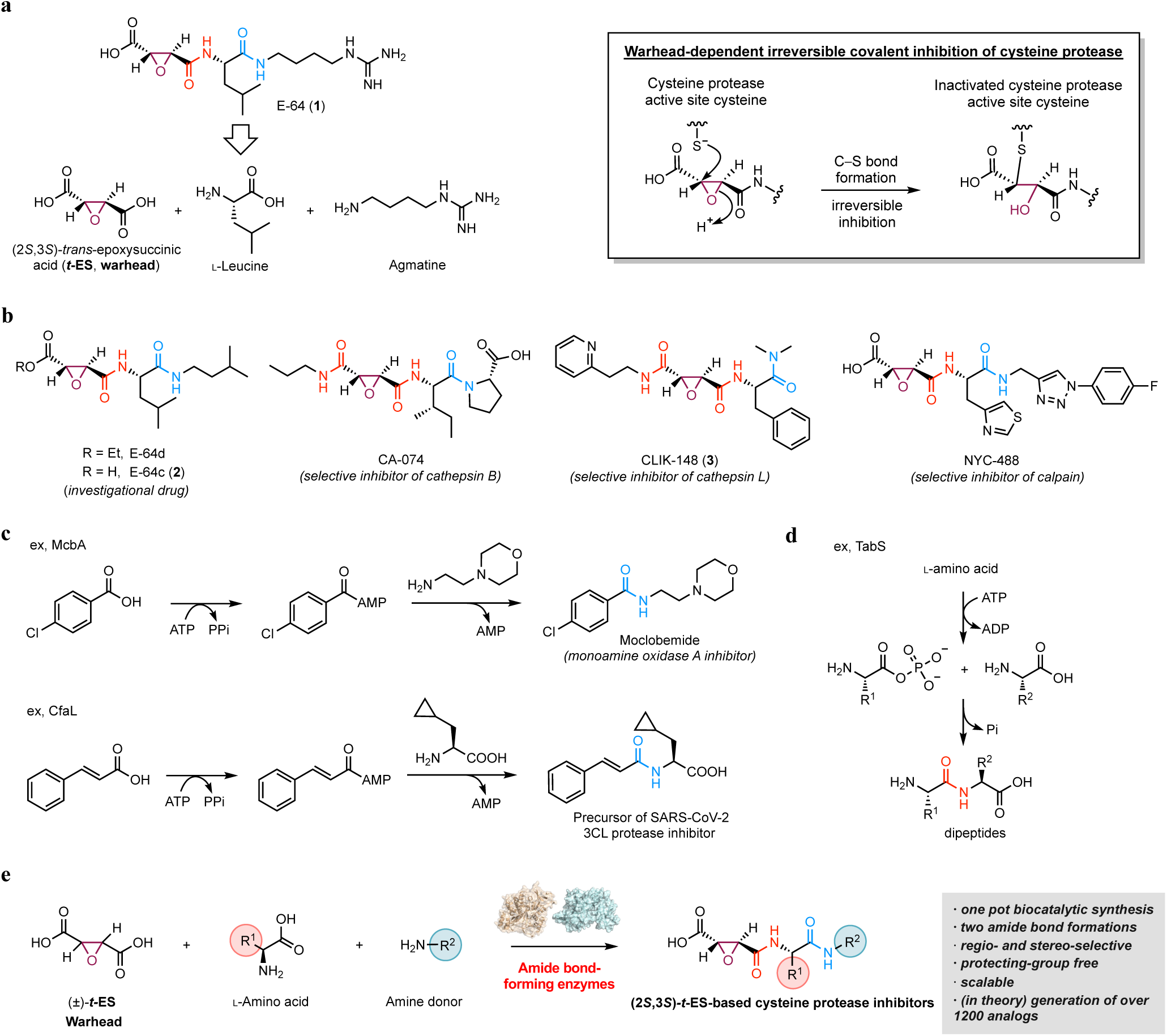
Amide bond-forming enzyme(s) are hypothesized to be involved in biosynthesis of E-64 (1). **a**, Structure, mode of action, and retrobiosynthesis of **1**. E-64 (**1**) contains the *trans*-epoxysuccinic acid (***t*-ES**) warhead that is the chemical basis for the covalent inhibition of cysteine proteases. **1** is proposed to form from the condensation of (2*S*,3*S*)-***t-*ES**, l-Leu, and agmatine. **b,** Structures of synthetic cysteine protease inhibitors based on **1**. **c**, Mechanism and application of amide bond synthetase (ABS) such as McbA and CfaL from the marinacarboline and coronatine biosynthetic pathways, respectively; **d,** Mechanism and applications of ATP-grasp enzyme such as TabS from tabtoxin biosynthetic pathway. **e**, In this work, we discovered an ATP-grasp enzyme and an ABS involved in the biosynthesis of **1**, and utilized them to realize combinatorial biocatalysis of E-64 analogs.

Biosynthetically, **1** is formed from the step-wise condensation of three distinct building blocks, the “warhead” (2*S*,3*S*)-***t*-ES** (a dicarboxylic acid), l-Leu (an amino acid), and agmatine (an amine) (**Fig. 1a, b**), using amide bond-forming enzyme(s) (**Fig. 1a**). Amide bond-forming enzymes have found wide-spread interests as biocatalysts for the synthesis of pharmaceuticals due to the ubiquity of the amide functionality (**Supplementary Fig. 2a-b**)^29,30^. Enzymatic synthesis of amides can obviate the protection-deprotection steps associated with functionally rich building blocks, and can achieve chemo- and regio-selectivity with atom-economy. Two classes of enzymes that are not associated with nonribosomal peptide synthetases (NRPSs) (**Supplementary Fig. 2c-d** and **Supplementary Fig. 3**), the ATP-grasp enzymes and amide-bond synthetases (ABSs) in particular, have attracted attention due to direct activation and amidation of carboxylic acids without the need for partnering enzymes^2,23,26,30–35^. ATP-grasp enzymes phosphorylate the carboxylic acid partner followed by condensation with the amine donor, while ABSs adenylate the carboxylic acid followed by amidation with amine nucleophiles (**Fig. 1c, d** and **Supplementary Fig. 2c**). Whereas bacterial examples, such as the ATP-grasp enzymes TabS^35^, PnaC^36^, and CysD^37^ as well as ABSs McbA^32,38^, CfaL^34^, and CysC^37^, have been demonstrated to be synthetically useful (**Fig. 1c, d** and **Supplementary Fig. 2c**), only a few putative ATP-grasp enzymes have been reported^39,40^ and no ABSs have been identified from fungi (**Supplementary Fig. 4a-b**). Given the tripartite structure of **1** and the analogs connected by two amide bonds, we hypothesized that amide bond-forming enzymes involved in the biosynthesis of **1** can be harnessed for the biocatalytic synthesis and structural diversification of both natural and synthetic E-64 analogs. Here, we described the discovery, characterization and application of a pair of ATP-grasp enzymes and ABSs from the E-64 biosynthetic pathway toward these applications (**Fig. 1e**).

## Results

### Discovery and characterization of the biosynthetic gene clusters of E-64 (1) and the analogs

A common strategy to identify biosynthetic gene clusters (BGCs) of amide-containing natural products, such as peptides and pseudopeptides, is to search for NRPS-anchored BGCs. This is especially suitable in fungi, as nearly all fungal peptidyl and pseudopeptidyl natural products are constructed by NRPS(s) (**Supplementary Fig. 2** and **Supplementary Fig. 3**)^41–44^. However, this strategy would not identify BGC candidates that encode an NRPS-independent pathway, as hypothesized for **1**. Direct searching using ATP-grasp enzymes and ABSs in fungi are challenging due to the lack of characterized examples. As a result, an alternative enzyme beacon that form a common structural feature, such as the polyamine fragment, was used in BGC identification (**Supplementary Fig. 1**). Polyamines, such as putrescine, agmatine, and cadaverine, are formed from the decarboxylation of l-Orn, l-Arg, and l-Lys, respectively, by PLP-dependent decarboxylases^44^. Although those polyamines are known primary metabolites in fungi, we reasoned that the BGCs for **1** and related compounds might encode a dedicated PLP-dependent lysine or ornithine decarboxylase to increase supplies of polyamines as building blocks.

Using *N*-dimethyllysine decarboxylase FlvG^45^ from the flavunoidine BGC as a beacon, we searched the genomes of fungal producers of **1** and analogs. Two putative BGCs (*acp1* and *acp2*) were identified from *A. oryzae* (**Fig. 2a**), which are also present in the closely related fungus *Aspergillus flavus* (*cp1* and *cp2*) (**Fig. 2a**). Each BGC encodes four proteins, including a putative αKG/Fe(II)-dependent oxygenase (*cp1A*/*cp2A*), a hypothetical protein (HP) (*cp1B*/*cp2B*) with no detectable Pfam domain, a PLP-dependent decarboxylase (*cp1C*/*cp2C*), and a protein (*cp1D*/*cp2D*) that is predicted to belong to the ANL-family^46^, of which ABSs are also members (**Supplementary Fig. 18**). Each protein in the *cp1* BGC shares ∼50% amino acid sequence identity to the corresponding homolog in the *cp2* BGC (**Fig. 2a**). A sequence similarity network (SSN)^47^ analysis of Cp1B and Cp2B revealed that there are many related proteins in Ascomycota, including the proposed ATP-grasp enzyme AnkG^40^ involved in NK13650C biosynthesis **Supplementary Fig. 4b-c**). Structural analysis of Cp1B and Cp2B by Foldseek^48^ identified homoglutathione (hGSH) synthetase as well as the recently characterized bacterial peptidylpolyamine synthetases (YgiC and YjfC)^49^ as the closest structure homolog (**Supplementary Fig. 7**), which is an ATP-grasp enzyme that catalyzes the condensation of γ-glutamylcysteine and β-alanine to form tripeptide hGSH^50^. The predicted functions of the four encoded proteins in these BGCs are therefore consistent with the required steps for biosynthesis of molecules related to **1** (**Fig. 1a** and **Supplementary Fig. 1**).

**Figure 2.**
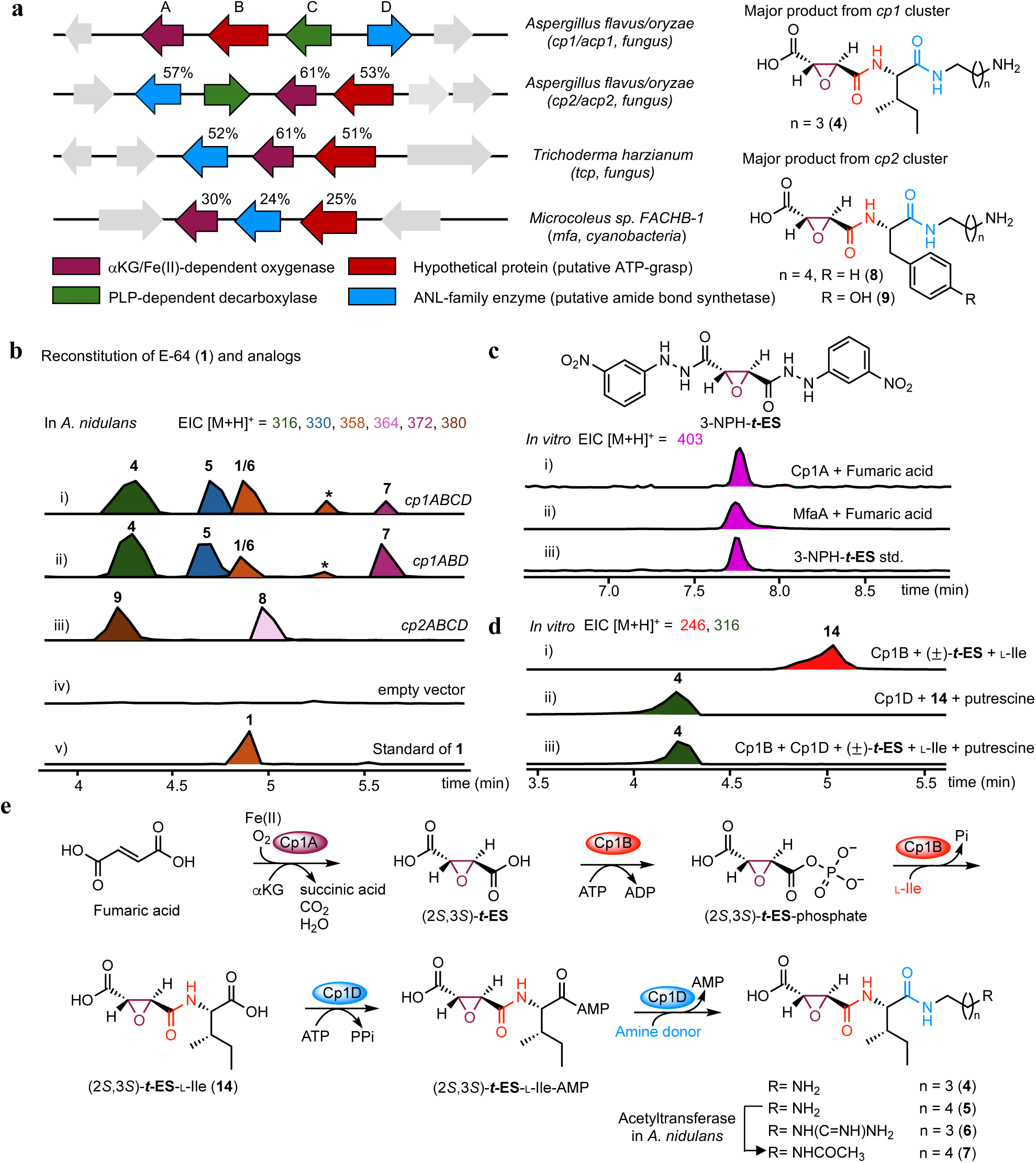
Biosynthetic pathway for E-64 (1) and identifying new amide-forming enzymes. **a**, BGCs of **1** from *A. flavus* and *A. oryzae,* and other homologous BGCs. The % amino acid sequence identity to each corresponding Cp1 enzyme is shown. **b**, LC/QTOF analysis of metabolites produced by different gene combinations from *cp1* and *cp2* clusters in the heterologous host *A. nidulans* are shown in i–v. Selected ion chromatography traces are shown, and the colors of the traces match to the indicated mass and compounds. *: not isolated. **c**, Cp1A catalyzed the epoxidation on fumaric acid to ***t*-ES**. Assays were carried out at 30 °C for 3 h in 100 μL of 50 mM sodium phosphate buffer with 0.2 mM FeSO_4_, 2 mM αKG, 2 mM ascorbate, 1 mM of fumaric acid, and 10 μM of Cp1A or MfaA The enzymatic products were derivatized with 3-nitrophenylhydrazine (3-NPH) to increase MS sensitivity. Selected ion monitoring of 3-NPH-***t*-ES** ([M + H]^+^ = 403) is shown. **d**, Enzyme assays with Cp1B and Cp1D. The reaction for i) was performed in 100 μL of 50 mM sodium phosphate buffer (pH8.0) containing 25 μM Cp1B, 5 mM *trans*-epoxy-succinic acid or analog, 2.5 mM l-Ile, 10 mM ATP, and 10 mM MgCl_2_ at 30 °C for 16 h. The reaction for ii) was performed in 25 μM Cp1B, 25 μM Cp1D, 5 mM (±)-*trans*-epoxy-succinic acid, 2.5 mM amino acid, 5 mM amine, 10 mM ATP cofactor, and 10 mM MgCl_2_ at 30 °C for 16 h. Selected ion monitoring of **4** ([M + H]^+^ = 316) and **14** ([M + H]^+^ = 246). **e**, Biosynthetic pathway of **1** and the analogs.

Heterologous expression of *cp1A-D* genes in *A. nidulans* ΛEMΛST^51^ (**Supplementary Fig. 6**) led to the production of multiple new hydrophilic metabolites with molecular weights (MWs) of 315 (**4**), 329 (**5**), 357 (**1** and **6**), and 371 (**7**) (**Fig. 2b**). Isolation and NMR analysis showed **4, 5** and **6** are analogs of **1** containing a ***t*-ES***-*l-Ile condensed with putrescine, cadaverine, and agmatine, respectively (**Supplementary Tables 6, 8-10** and **Supplementary Figs. 37-41, 49-63**)^21^. Compound **6** is the l-Ile variant of **1** and coelutes with **1** during analysis. NMR analysis showed **7** is a derivative of **5** that has been *N*-acetylated (**Supplementary Table 11** and **Supplementary Figs. 64-68**), a reaction that is likely catalyzed by an endogenous acyltransferase in *A. nidulans*. The absolute stereochemistry of ***t*-ES** moiety in those compounds was deduced to be (2*S*,3*S*) based on comparisons of NMR spectra and optical rotation with reported values^21^. In contrast, heterologous expression of the *cp2* BGC in *A. nidulans* afforded **8** and **9**, which were characterized to be ***t*-ES-**l-Phe-cadaverine and ***t*-ES-**l-Tyr-cadaverine, respectively (**Fig. 2a,b**, **Supplementary Tables 12-13**, and **Supplementary Figs. 69-78**). Two minor products **10** and **11** were also detected from this strain (**Supplementary Fig. 9**), of which **10** is characterized to be *N-*acetyl-**8** (**Supplementary Table 14** and **Supplementary Figs. 79-83**) and **11** is proposed to be *N-*acetyl-**9** based on MW.

These results confirmed that both *cp1* and *cp2* BGCs are responsible for the NRPS-independent biosynthesis of **1** and analogs. Removing the PLP-dependent decarboxylase *cp1C* from each *A. nidulans* transformant did not affect the metabolic profiles (**Fig. 2b** and **Supplementary Fig. 9**), suggesting the amine precursors are abundant in the heterologous host. Omitting any of *cp1A*, *cp1B* or *cp1D* from the transformant abolished the production of ***t*-ES** containing compounds related to **1** (**Supplementary Fig. 9**), thereby establishing these three genes constitute a minimum biosynthetic cassette. Indeed, such three-gene BGCs can be found in more than > 100 fungal species with > 40 different genera, as well as in cyanobacteria such as *Microcoleus spp.,* nearly all of which are not known to produce related compounds (**Fig. 2a**, and **Supplementary Figs. 1, 5a-b**). It is worth noting that the transformant lacking *cp1A* was able to produce the malic acid and fumaric acid analogs of **4**, which were characterized to be **12** and **13**, respectively (**Supplementary Tables 15-16** and **Supplementary Figs. 9, 75-84**), in agreement with the proposed role of Cp1A as the epoxidase.

### Cp1A and its homologs are a warhead (2*S*,3*S*)-*trans*-epoxysuccinic acid synthase

To confirm the function of Cp1A as an epoxide-forming enzyme, we overexpressed and purified the recombinant *C*-His-tagged Cp1A from *Escherichia coli* BL21 (DE3) (**Supplementary Fig. 10**). Enzyme assays were performed in the presence of fumaric acid and the products were derivatized with 3-nitrophenylhydrazine (3-NPH). LC/MS analysis showed that while no 3-NPH-***t*-ES** can be observed in the absence of Cp1A, fumaric acid was efficiently converted into ***t*-ES** by Cp1A in the presence of typical cofactors required for an αKG/Fe(II)-dependent oxygenase (**Fig. 2c**). Using a coupled assay with Cp1A and Cp1B to form ***t*-ES**-l-Ile (*vide infra*), the stereochemistry of the epoxide formed by Cp1A was assigned to be (2*S*,3*S*) based on comparison of retention time to those of synthetic (2*S*,3*S*)-***t*-ES**-l-Ile (**14**) and (2*R*,3*R*)-***t*-ES**-l-Ile (**Supplementary Fig. 11a**). Cp1A is not able to catalyze the epoxidation of the malyl-(**12**) and fumaryl-l-Ile-putrescine (**13**) (**Supplementary Fig. 11b**), demonstrating Cp1A stereoselectively epoxidizes free fumaric acid to give the warhead (2*S*,3*S*)-***t*-ES**. Interestingly, when we used succinic acid instead of fumaric acid as the substrate of Cp1A, 3-NPH-***t*-ES** and 3-NPH-fumaric acid were also produced, albeit with lower efficiency (**Supplementary Fig. 11c**). As similar to the reported multifunctional αKG/Fe(II)-dependent oxygenase AsqJ^52^, we reasoned that Cp1A is able to catalyze the desaturation of succinic acid to fumaric acid which is subsequently epoxidized into (2*S*,3*S*)-***t*-ES**.

An analog of **1**, circinamide^53^, was isolated from cyanobacteria *Anabena* spp. (**Supplementary Fig. 1**). A homologous BGC of **1** can be found in cyanobacteria genomes including *Microcoleus spp* (*mfa*), although the identities between Mfa and Cp1 enzymes are low (∼20 to 30%). Recombinant MfaA (**Supplementary Fig. 10**) is also able to catalyze the formation of (2*S*,3*S*)-***t*-ES** from fumaric acid (**Fig. 2c** and **Supplementary Fig. 11a**), representing the first bacterial example of an epoxysuccinic acid synthase.

### Cp1B and Cp2B are a novel family of pseudodipeptide-forming ATP-grasp enzyme

The putative ATP-grasp enzyme Cp1B and the ANL-family enzyme Cp1D are responsible for the amide bond-forming steps to give **1** and analogs. Both enzymes were purified from *E. coli* BL21 (DE3) (**Supplementary Fig. 10**). When assayed in the presence of (2*S*,3*S*)-***t*-ES**, l-Ile, putrescine, ATP and MgCl_2_, formation of **4** was observed (**Fig. 2d**). When putrescine or Cp1D was omitted from the reaction mixture, formation of (2*S*,3*S*)-***t*-ES**-l-Ile (**14**) was observed (**Fig. 2d**). This suggested formation of the first amide bond between (2*S*,3*S*)-***t*-ES** and l-Ile is catalyzed by Cp1B, while formation of the second amide bond between **14** and putrescine is catalyzed by Cp1D (**Fig. 2e**). Cp1B is confirmed to be an ATP-grasp enzyme from the formation of ADP in the reaction (**Supplementary Fig. 12a**)^50^. Replacing l-Ile with D-Ile did not lead to formation of (2*S*,3*S*)-***t*-ES**-D-Ile, demonstrating the stereospecificity of Cp1B towards l-amino acids (**Supplementary Fig. 13**). Cp1B is also enantiospecific for (2*S*,3*S*)-***t*-ES**, as steady-state kinetics showed Cp1B strongly phosphorylated (2*S*,3*S*)-***t*-ES** (*k*_cat_/*K*_M_ = 48.0 min^−1^mM^−1^) over (2*R*,3*R*)-***t*-ES** (not measurable) (**Supplementary Fig. 12a-b**). This property of Cp1B was demonstrated in a kinetic resolution of (±)-***t*-ES** through the amide-forming reaction, during which (±)-***t*-ES** was resolved through the formation of (2*S*,3*S*)-***t*-ES**-l-Ile (**14**) with diastereomeric ratio (d.r.) >98:2 (**Supplementary Fig. 14**). Cp1B is therefore a suitable biocatalyst to monoamidate (2*S*,3*S*)-***t*-ES** from racemic ***t*-ES**. In contrast, chemical synthesis of **1** and analogs requires asymmetric synthesis or resolution of stereochemically homogenous (2*S*,3*S*)-***t*-ES** monoester to prevent diamidation (**Supplementary Fig. 17**)^6^.

The X-ray crystal structure of Cp1B in a complex with ADP and Mg^2+^ ions was determined to 2.7-Å-resolution (**Fig. 3a**). Despite having no detectable sequence identity with other characterized ATP-grasp enzymes, the structure of Cp1B displays the three domains architecture (domains A, B, and C) found in ATP-grasp enzymes such as hGSH synthetases, and a conserved ATP binding site at the interface of domains A and C (**Fig. 3a** and **Supplementary Fig. 15**)^50,54^. Both open (PDB code: 3KAK) and closed (PDB code: 3KAL) active site forms of hGSH synthetases have been crystallized, with the phosphorylation and amide bond forming steps being catalyzed in the closed form^54^. The crystalized Cp1B structure is in the closed form, as the ATP binding site is covered by the lid-domain with the P-loop and the A-loop from the domain A (**Fig. 3a** and **Supplementary Fig. 15**)^54^. Similar to the ATP binding site of hGSH synthetases, the ADP-Mg^2+^ complex in the Cp1B structure is sandwiched between the strands of two anti-parallel β-sheets (**Fig. 3a, b** and **Supplementary Figs. 12c, 15**), and interacts with the conserved residues R139, D141, E163, and N165 (**Supplementary Figs. 12c**). Individual mutations of those residues to alanine either abolished or significantly attenuated enzymatic activity of Cp1B (**Supplementary Figs. 12d**). Hence, although annotated as a HP, our biochemical and structural characterization demonstrated Cp1B is the first biochemically validated case of an ATP-grasp enzyme in fungal natural product biosynthesis.

**Figure 3.**
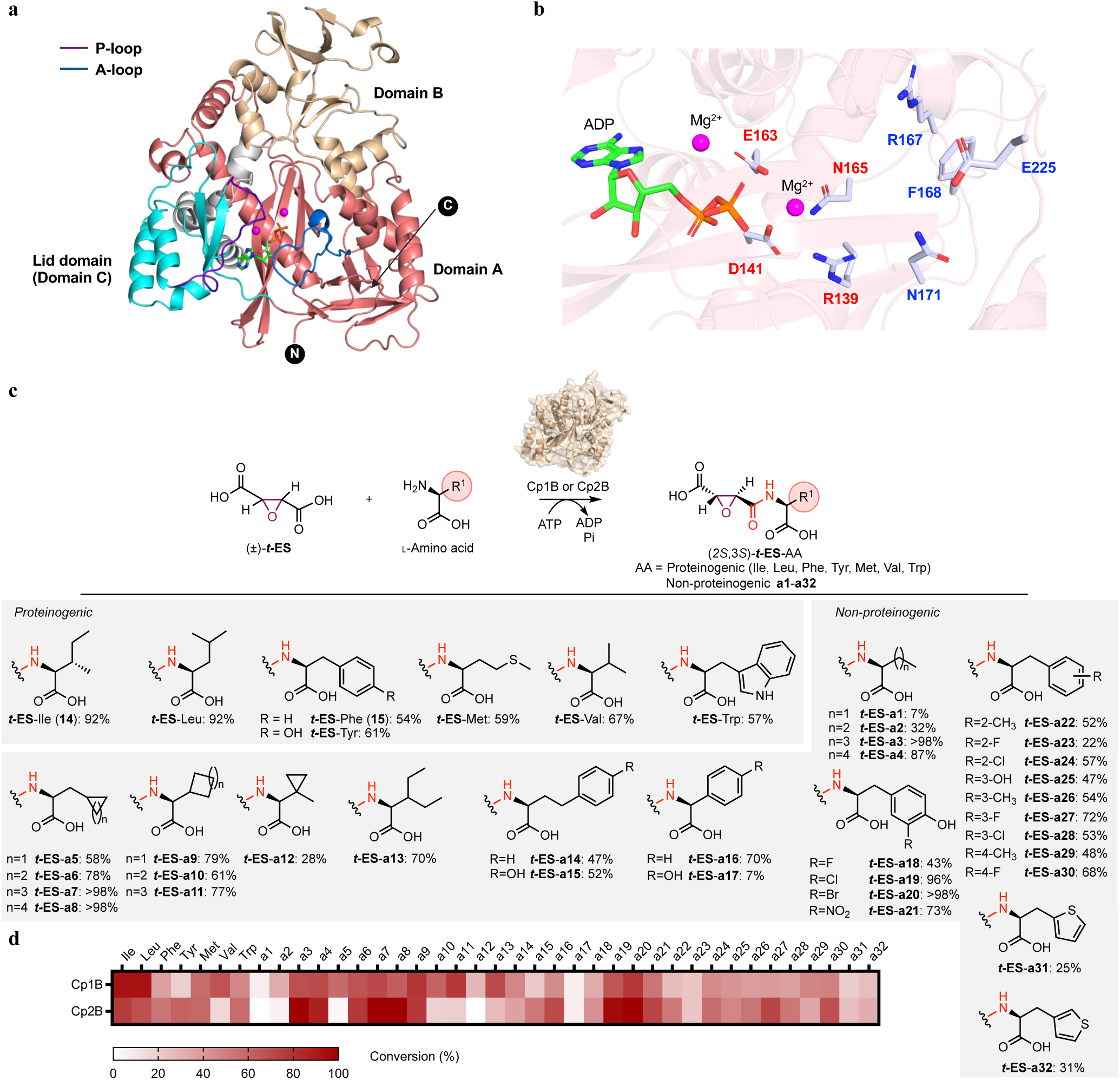
Stereoselective (2*S*,3*S*)-*t*-ES-l-amino acid synthesis by Cp1B and Cp2B. **a**, Crystal structure of Cp1B in a complex with ADP and two Mg^2+^ ions. Domains A (residues 5–200 and 444–502), B (residues 200–323), and C (residues 365–443) are shown as deepsalmon, wheat, and cyan, respectively. The helical connection (residues 324–364) between domains B and C is shown in grey. The P-loop (Gly-rich loop) and A-loop (Ala-rich loop) are highlighted in purple and blue, respectively. ADP is colored in light green, while Mg^2+^ ions are shown as magenta spheres. The Cp1B structure adopts a closed form (see **Supplementary Figs. 15**). **b**, Enlarged view of the ATP binding site. Note that clear electron density of MES was observed, but not shown in this version of figure. The clear electron density of adenosine was observed, and modeled ADP is shown in this version of figure. The conserved residues with hGSH synthetases such as R139, D141, E163, and N165 are labeled as red, which are potentially involved in the phosphorylation of the carboxylate in substrate. The other non-conserved residues with hGSH synthetases are shown in blue, which are likely involved in the substrate binding. **c**, Structure of proteinogenic and non-proteinogenic (**a1-a32**) amino acid substrate scope for Cp1B and Cp2B. The higher percent conversions shown represent the higher number between reactions catalyzed by either enzyme. As shown in the legend of Fig. 3d, the % conversion was estimated based on the calibration curves of enzymatically prepared standards. **d**, Heat map (estimated %conversion) comparing ligation of amino acid with ***t*-ES** by Cp1B and Cp2B, based on the calibration curves of enzymatically prepared standards. Assays were carried out with enzymes (25 μM), (±)-***t*-ES** (5 mM), amino acid (2.5 mM), 10 mM MgCl_2_, 10 mM ATP in 100 μL of 50 mM sodium phosphate buffer (pH 8.0) at 30 °C for 16 h.

We next explored the substrate scopes of Cp1B and Cp2B towards enzymatic synthesis of (2*S*,3*S*)-***t*-ES-AA**, where **AA** is any l-amino acid. Cp1B and Cp2B preferentially monoamidated (2*S*,3*S*)-***t*-ES** with hydrophobic amino acids including l-Ile, l-Leu, l-Val, l-Met, l-Phe, l-Trp, and l-Tyr, to varying conversions (**Fig. 3c,d**). A wide range of non-proteinogenic, nonpolar amino acids (**a1**-**a32**) were also ligated to (2*S*,3*S*)-***t*-ES**, including pharmaceutical building blocks such as cyclopropyl-l-alanine **a5** and cyclobutyl-l-alanine **a6** by Cp1B (**Fig. 3c,d** and **Supplementary Fig. 15**). Both enzymes, especially Cp2B, can also efficiently catalyze the mono-amidation of (2*S*,3*S*)-***t*-ES** with aromatic amino acids with diverse substitutions including phenolic alcohol, halide, nitro, methyl, and thiophenyl groups (**a18**-**a32**) (**Fig. 3c,d**). Nonaromatic amino acids with either nitrogen or oxygen-containing side chains, and α,α-disubstituted amino acids are not well-tolerated by Cp1B and Cp2B (**Supplementary Fig. 22**). Overall, Cp1B prefers aliphatic amino acids and has a broader substrate scope than that of Cp2B, which prefers aromatic and bulkier amino acids (**Fig. 3d**). The complementary substrate scopes could therefore be leveraged towards the biocatalytic synthesis of a diverse array of (2*S*,3*S*)-***t*-ES**-**AA** compounds. Indeed, thirty-eight 2,3-epoxyamides prepared with either Cp1B or Cp2B were isolated by a single-step purification from preparative enzymatic reactions and characterized by NMR (**Supplementary Tables 17-54 and Figs 94-283**).

### Cp1D and Cp2D are first fungal amide bond synthetases

To confirm the role of Cp1D in catalyzing the second amide bond-forming reaction, Cp1D was directly assayed with **14** and an amine nucleophile. As expected, in the presence of putrescine, cadaverine, or agmatine, Cp1D catalyzed the formation of **4**, **5** or **6**, respectively (**Fig. 2d** and **Supplementary Fig. 19**). Cp1D can efficiently amidate **14** to **6** with a *k*_cat_/*K*_M_ (apparent) of 79.8 min^−1^mM^−1^, much more efficient compared to that of (2*R*,3*R*)-***t*-ES**-l-Ile (**Supplementary Fig. 20**). The mechanism of Cp1D was confirmed to be that of an ABS, as formation of **14**-AMP in the presence of ATP was detected by LC/MS (**Supplementary Fig. 20**). This establishes Cp1D to be the first characterized fungal ABS.

The substrate scope of Cp1D was comprehensively explored using a two-stage screen. First, twenty *N*-succinyl proteinogenic l-amino acids (**suc-AA**) were synthesized (**Supplementary Figs. 26a, 314-330**) and assayed in the presence of ATP and isopentylamine. Parallel to the substrate preference of Cp1B, Cp1D preferentially amidates seven **suc-AA** substrates containing hydrophobic amino acids (l-Ile, l-Leu, l-Val, l-Met, l-Phe, l-Trp, and l-Tyr) (**Supplementary Figs. 26a**). The **suc-AA** substrates containing l-Ala and l-Pro can also be amidated by Cp1D, albeit with low efficiency. Adenylation of the **suc-AA** by Cp1D and Cp2D were also assayed by the hydroxylamine-based colorimetric assay^55^ (**Supplementary Figs. 26b**). Cp2D adenylated **suc**-l-Trp and **suc**-l-Tyr more efficiently compared to Cp1D, in agreement with the preference for aromatic amino acids displayed by Cp2B over Cp1B. Using the thirty-nine (2*S*,3*S*)-***t*-ES**-**AA** prepared by Cp1B (**Fig.3c**), Cp1D can amidate all with agmatine as judged by LC/MS analysis (**Supplementary Fig. 24**). Interestingly, Cp1D can also amidate the terminal carboxylate of l-Asp-l-Phe, l-Asp-l-Leu, and glutaryl-l-Leu with moderate activity to form isopentyl amidated dipeptides (**Supplementary Fig. 32**). As Cp1D/Cp2D do not have any sequence similarity to characterized bacterial ABSs such as McbA^32,38^, DdaF^56^, CfaL^34^, and CysC^37^, Cp1D/Cp2D can be grouped into a new family of ABSs that activates *N*-acyl amino acids and dipeptides, instead of simple organic acids shown by the bacterial ABSs (**Fig. 1c** and **Supplementary Fig. 8**).

The second stage assays assessed the scope of nucleophiles that can be used by Cp1D, during which in vitro reaction of Cp1D with (2*S*,3*S*)-***t*-ES**-l-Phe (**15**) and diverse amines were performed (**Fig. 4** and **Supplementary Fig. 27**). Remarkably, Cp1D can accept forty-one of the selected amine nucleophiles to form analogs of **1** with excellent conversions (**Fig. 4**). These include primary amines that are: diamines with different chain lengths (**b1**-**b7**); alkanoamines (**b8**-**b9**), aliphatic amines with the different chain lengths (**b10**-**b17**), arylamines (**b18**-**b20**), alkynyl amines (**b21-b24**), azidoamines (**b25-b27**), heteroarylamines (**b29**-**b32**) and alkyl hydrazine (**b33**). Alkylamines with carboxylic acids at one end were not tolerated by Cp1D (**Supplementary Fig. 25**), although alkylamines capped with carboxylic acid methyl ester (**b34**-**b37**) can be efficiently incorporated into diamide products. Among secondary amine nucleophiles, dimethylamine (**b28**) is an excellent substrate for Cp1D, while bulkier amines such as diethylamine and dipropylamine were not accepted (**Supplementary Fig. 25**). Most amino acids and their methyl esters, except for Gly-OMe (**b34**), were not accepted as an amine donor by Cp1D. The dipeptide Gly-l-Tyr-OMe (**b38**) was well-accepted by Cp1D as a nucleophile, suggesting that the substrate scope could be further explored to towards synthesis of pseudotetrapeptides.

**Figure 4.**
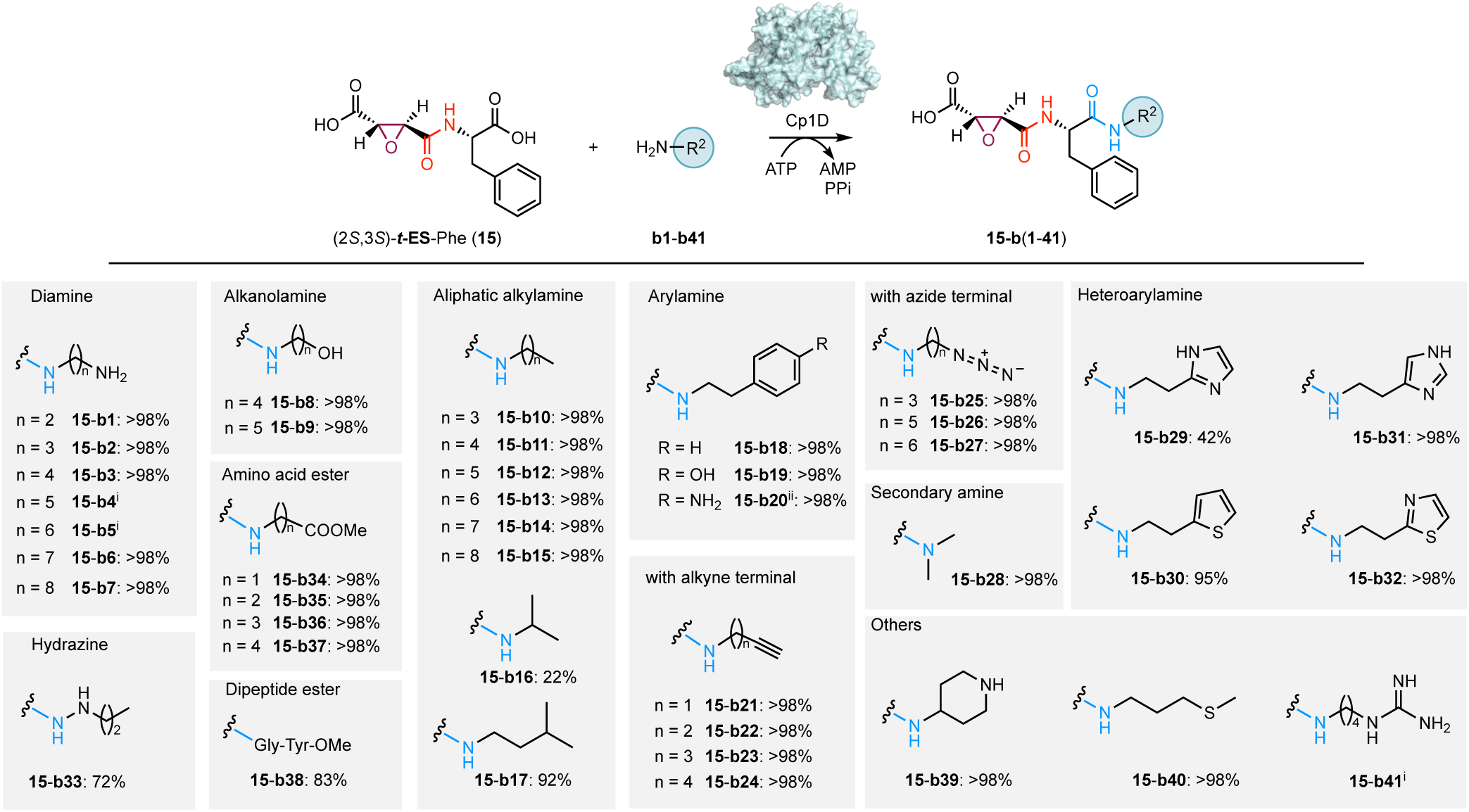
A broad range of amines is accepted by Cp1D to form the warhead-containing diamides. Amine substrates (**b1-b41**) were converted into the corresponding diamides (**15-b1** to **15-b41**) assayed with **15** and Cp1D. Assays were carried out with Cp1D (25 μM), (2*S*,3*S*)-***t*-ES**-l-Phe (2 mM), an amine (5 mM), 10 mM ATP, and 10 mM MgCl_2_ in 100 μ L of 50 mM sodium phosphate buffer (pH 8.0) at 30 °C for 16 h. The reactions were analyzed by LC/MS. Conversion (%) of **15** to the corresponding amide product was calculated based on HPLC peak area ratios between product and **15** at 204 nm (% Conversion = (peak area of product / (peak area of **15** + peak area of product)) × 100%). i: product peak was detected but overlap with **15** on the LC/MS chromatogram. ii: While **b20** has two nucleophilic amines (aromatic and aliphatic amines) in the structure, only single major product peak was detected on the HPLC trace. One of the proposed structures is shown.

### Combinatorial biocatalytic synthesis and screening of cysteine protease inhibitors

Combinatorial biocatalysis is the enzymatic version of combinatorial synthesis, and uses cascaded enzymatic transformations for library construction^57,58^. Applications of combinatorial biocatalysis are limited due to the lack of examples in which two or more enzymes in the same cascade are both highly promiscuous^59^. Given the broad substrate scope established for Cp1B and Cp1D, a library of more than 1200 E-64 analogs can be generated and screened as ***t*-ES** based cysteine protease inhibitors. To access a fraction of that library in a proof of concept study, a 96-well assay containing the warhead (±)-***t*-ES**, two enzymes (Cp1B and Cp1D), eight proteinogenic amino acids (l-Ile, l-Leu, l-Val, l-Met, l-Phe, l-Trp and l-Tyr, with l-His as negative control) and twelve amine donors including newly tested amines such as histamine (**b42**), 4-(2-aminoethyl)-pyridine (**b43**), and tryptamine (**b44**) were performed. Masses correspond to the expected products were detected in all samples except those with l-His as negative controls (**Supplementary Fig. 28**).

By coupling this combinatorial biocatalytic platform with a fluorescence-based assay (**Fig. 5a**), we set out to identify highly potent human cathepsin B inhibitors. A library of thirty-nine analogs of (2*S*,3*S*)-***t*-ES-AA-** agmatine was generated by varying the amino acid (**Supplementary Fig. 24**). The crude reaction mixtures in 96-wells format were directly screened for cathepsin B inhibitory activity in a fluorometric endpoint assay^60^. In agreement with reported structure-activity-relationship (SAR) for cathepsin B inhibitors^9,61^, analogs having an aliphatic amino acid such as l-cyclobutylalanine (**a6**), l-cyclopentylglycine (**a10**), l-cyclohexylglycine (**a11**), and 3-ethyl-l-norvaline (**a13**) exhibited the strongest cathepsin B inhibition (**Fig. 5b**). We then enzymatically prepared a library with l-cyclopentylglycine (**a10**) as the amino acid building block and forty-one different amine donors (**Fig. 5c** and **Supplementary Fig. 30**). While arylamines (**b18**-**b19**) and heteroarylamines (**b29**-**b32**) decreased the inhibitory activity compared to agmatine (**b41**), alkyl amines with terminal amine (**b7**), alcohol (**b9**), methyl (**b13**), alkyne (**b24**), and azide (**b26**) all displayed comparable or more potent inhibition (**Fig. 5c, d**). Therefore, these results support the utility of this combinatorial biocatalysis approach for cysteine protease inhibitor screening, which can be easily extended to other disease-relevant enzymes such as cathepsin C^62^.

**Figure 5.**
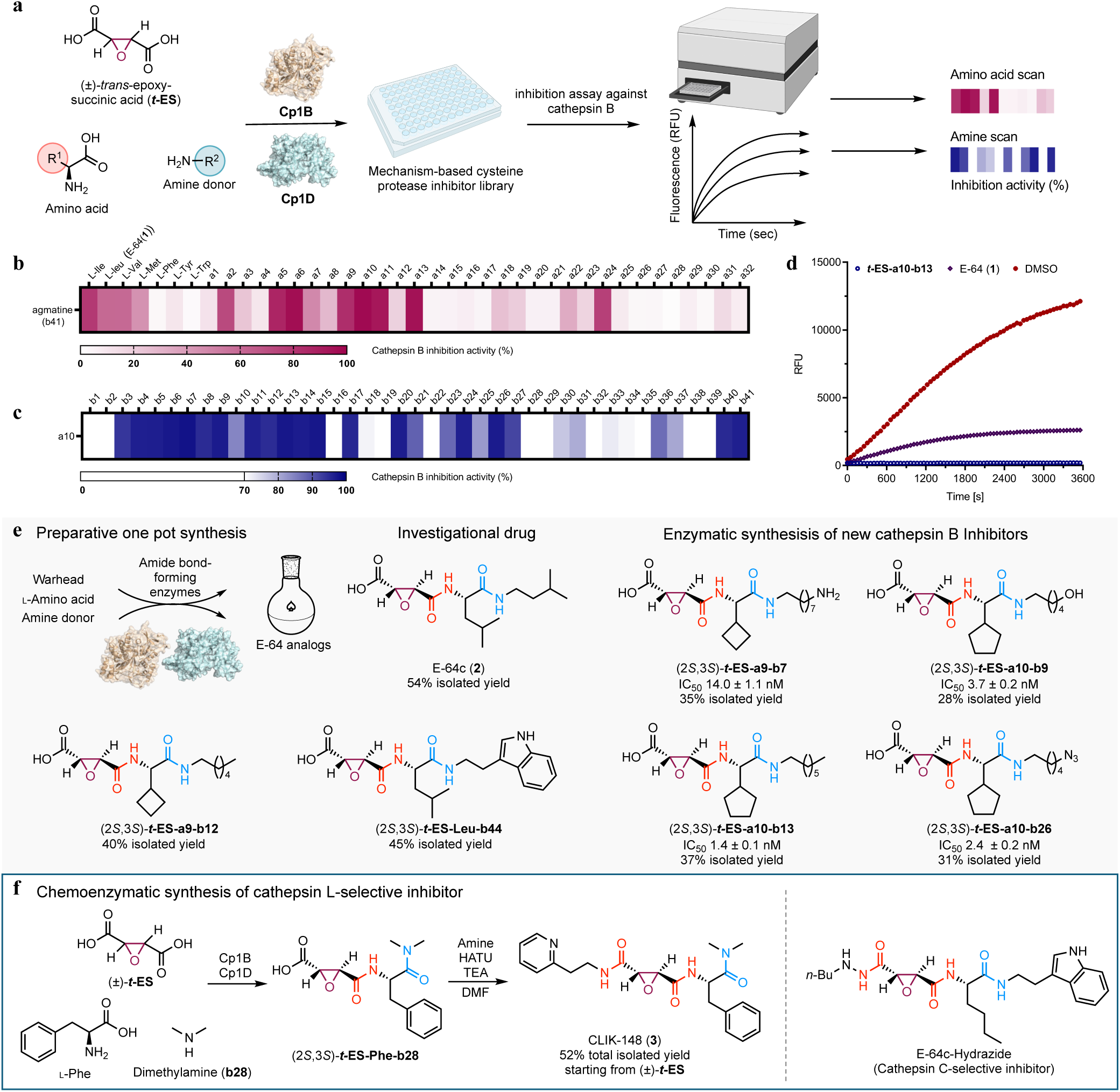
Biocatalytic platform for screening and synthesis of *t-*ES-based cysteine protease inhibitors. **a**, General workflow. **b**, Cathepsin B inhibitory activity of the crude reaction containing (2*S*,3*S*)-***t*-ES**-**AA**-agmatine where **AA** is an amino acid (**a1**-**a32**), and **c**, Cathepsin B inhibitory activity of the crude reaction containing (2*S*,3*S*)-***t*-ES**-**a10**-**B** where **B** is an amine (**b1**-**b41**). The average inhibition activity is presented as a percentage of the positive control (n=2, where n represents the number of independent experiments). **d**, Selected data output for fluorescence-based cathepsin B inhibition assay using a fluorogenic substrate *Z*-Phe-Arg-AMC. RFU represents relative fluorescence unit. **e**, Biocatalytic one-pot synthesis of **2** and newly identified cathepsin B inhibitors. The % isolated yields are shown for all synthesized compounds. IC_50_ of selected new inhibitors against cathepsin B are also shown. Reaction conditions for Fig. 5b and 5c; the assays were carried out with Cp1B (25 μM), Cp1D (25 μM), 2 mM (±)-***t*-ES**, 1 mM l-amino acid, 1 mM amine, 10 mM ATP, and 10 mM MgCl_2_ in 100 μL of 50 mM sodium phosphate buffer (pH 8.0) at 30 °C for 16 h. Reaction condition for Fig. 5e; large scale 20 mL in vitro reactions with 5 mM (±)-***t*-ES**, 2.5 mM l-amino acid, 2.5 mM amine donor, Cp1B (2.5 μM), and Cp1D (2.5 μM) in 50 mM sodium phosphate (pH 8.0) was carried out at 30 °C for 16 h. **f**, Chemoenzymatic synthesis of CLIK-148 (**3**) in preparative scale. After the biocatalytic synthesis of (2*S*,3*S*)-***t*-ES-Phe-b28** by following the same method described in the reaction condition for Fig. 5e using l-Phe as amino acid and dimethylamine (**b28**) as amine, the crude mixture can be directly used for the subsequent chemical condensation. The chemoenzymatic approach can also be applied to synthesis of the cathepsin C-selective inhibitor.

### Preparative synthesis of E-64 analogs

To assess the detailed inhibitory activity of hit compounds identified from the combinatorial biocatalytic platform, preparative scale reactions with Cp1B and Cp1D in a single reaction vessel were performed. The targeted compounds include (2*S*,3*S*)-***t*-ES-a9-b7**, (2*S*,3*S*)-***t*-ES-a9-b12**, (2*S*,3*S*)-***t*-ES**-**a10-b9**, (2*S*,3*S*)-***t*-ES-a10-b13**, (2*S*,3*S*)-***t*-ES-a10-b26**, (2*S*,3*S*)-***t*-ES-Leu-b44**, and the investigational drug E-64c (**2**) (**Fig. 5e**). Based on the reported stereoselective synthesis of **1**, chemical synthesis of those compounds would require not only the preparation of the (2*S*,3*S*)-***t*-ES** monoester, but also subsequent multistep synthesis with protection-deprotection steps (**Supplementary Fig. 29**)^6^. In contrast, biocatalytic synthesis readily afforded the targeted compounds with isolated yield of 28% to 54% (**Fig. 5e**), which were structurally confirmed with NMR (**Supplementary Tables 55-60** and **Supplementary Figs. 42-43, 284-313)**. It is worth noting that the preparative synthesis of **2** coupled with the ATP-recycling system could also afford the nearly identical conversion with the standard conditions (**Supplementary Fig. 36**), further highlighting the cost-effectiveness of the biocatalytic cascade synthesis^63^. With those pure compounds in hand, we performed detailed kinetic characterization and confirmed the new analogs (2*S*,3*S*)-***t*-ES**-**a10-b9**, (2*S*,3*S*)-***t*-ES-a10-b13**, and (2*S*,3*S*)-***t*-ES-a10-b26** all exhibited lower IC_50_ values toward cathepsin B than that of **1** (**Fig. 5d-e** and **Supplementary Fig. 33**). As those newly generated inhibitors bear the warhead (2*S*,3*S*)-***t*-ES,** the covalent adduct between the (2*S*,3*S*)-***t*-ES** moiety and active site cysteine in cysteine proteases is expected to remain as the mechanism of irreversible inhibition^20^. Indeed, when crystallized with papain, (2*S*,3*S*)-***t*-ES-a9-b7** prepared from the enzymatic reaction was bound covalently to the active site cysteine in the same conformation as **1** and E-64d (**Supplementary Fig. 34**).

The biocatalytic platform with Cp1B and Cp1D cannot yet directly synthesize cathepsin l-selective inhibitors frequently used in biological studies such as CLIK-148 and CAA0225, which have an additional amide bond to the ***t*-ES** moiety (**Fig. 5f**)^26,64–66^. A facile combination of biocatalytic and chemical syntheses was explored to access these compounds. For example, CLIK-148 can be easily obtained with ∼50% total isolated yield from the biocatalytic synthesis of (2*S*,3*S*)-***t*-ES-Phe-b29**, followed by chemical condensation with 2-pyridylethylamine (**Fig. 5f**, **Supplementary Table 7** and **Supplementary Figs. 44-48**). Compared to the reported chemical synthesis of CLIK-148 that requires more than six steps, purification of all intermediates, and expensive starting material of (2*S*,3*S*)-***t*-ES** diethyl ester (**Supplementary Fig. 29**), the chemoenzymatic synthesis is considerably simpler, greener and more cost-effective^6,67^. Therefore, such integration of chemical synthesis into the biocatalytic platform can further expand the structure diversity of ***t*-ES**-based inhibitor library that can be screened for desired properties including potency and selectivity (**Fig. 5f**).

## Discussion

Biosynthetic investigation of natural products have contributed significantly to the identification of new classes of enzymes and useful biocatalysts for synthesis and modification of bioactive compounds^44^. In this work, we discovered two families of amide bond-forming enzymes that have not been previously characterized from fungi: The ATP grasp enzymes Cp1B (Cp2B) were initially annotated as HPs and the ABSs Cp1D (Cp2D) annotated as ANL-family proteins, both from the well-explored genomes of *A. oryzae* and *A. flavus*^44,68^. Heterologous expression, biochemical assays, and structural characterizations uncovered the roles of these enzyme in catalyzing amide bond formations during the biosynthesis of **1** and analogs. Taking the biosynthetic pathway of **1** as an example, αKG/Fe(II)-dependent oxygenase Cp1A stereoselectively forms a warhead (2*S*,3*S*)-***t*-ES** from fumaric acid, which is condensed with l-Leu by the ATP-grasp enzyme Cp1B. The ABS Cp1D then catalyzes the condensation of the resultant (2*S*,3*S*)-***t*-ES**-l-Leu with agmatine to complete the pathway (**Fig. 2e**).

It is somewhat surprising at the onset of this study that the biosynthesis of **1** has remained elusive, despite the important role of **1** and related compounds in chemical biology and drug discovery. The difficulties in the identification of the correct BGC could stem from our findings that the biosynthetic pathway does not involve NRPS(s) for the amide-forming steps. Particularly in fungi, NRPS-independent amide bond formation steps in the biosynthesis of peptidyl and pseudopeptidyl natural products are exceptionally rare^41,44^. Our discovery and characterization of Cp1B (and Cp2B) and Cp1D (and Cp2D) hence established this mode of amide bond formation in fungal secondary metabolism. Homologous BGCs are found to be widely conserved in hundreds of different Ascomycetes species most of which are not known to produce E-64 analogs (**Supplementary Figs. 4c, 5**). Many homologs of Cp1B and/or Cp1D with >30% amino acid identity can be readily found in the fungal genomes deposited to the NCBI genome database. Furthermore, homologs to both enzymes can also be found in BGCs that are not homologous to that of **1** (**Supplementary Figs. 4c, 8**). Genome mining efforts using Cp1B and Cp1D as enzymatic beacons would lead to the discovery of new amide-bond containing natural products.

The concept of combinatorial biocatalysis for the generation of compound libraries appeared in the early 2000s as an emerging technology in drug discovery^57,58^. Through biosynthetic studies, several ATP-grasp enzymes and ABSs have been identified from the biosynthetic pathways of bacterial bioactive oligopeptides^31,32,34^, including warhead-armed pseudopeptides dapdiamides^56^, rhizocticins^69^ and cystargolides^37,70^. While these enzymes have been explored for generation of natural product analogs or applications in biocatalysis, the broad utility in combinatorial biocatalysis is limited due to the unwanted cross-reactivity^71^ of the enzymes toward substrates^37^ and/or due to the lack of pairing with additional enzymes having similar or great substrate promiscuity^59^. Here, by taking advantage of the broad substrate scope of both Cp1B and Cp1D, we demonstrated combinatorial biocatalysis to create a library of E-64 analogs using a 96-well format (**Fig. 5a**). The high-throughput enzymatic library generation was further coupled to direct fluorescence-based screening to rapidly identify new and more potent cathepsin B inhibitors (**Fig. 5**). By simply changing the protease target or the assay, this panel of epoxy-diamides can be screened for other purposes including the design of new ABPP probes.

Compared to chemical synthesis of E-64 and related compounds (**Supplementary Fig. 29**)^6^, the combined Cp1B and Cp1D enzymatic synthesis has several key advantages. First, the enantio-specificity displayed by Cp1B towards (2*S,*3*S*)-***t*-ES** enables the use of (±)-***t*-ES** as a starting material, which is significantly more cost-effective and does not require stereoselective synthesis. Secondly, the lack of cross-reactivity between Cp1B and Cp1D towards the amino acid and amine substrates, respectively, enable a one-pot reaction and obviates the need for protection and deprotections steps that are a stable in chemical synthesis of amides from functionally rich starting materials. Thirdly, the ATP required for amide-bond formation can be easily regenerated using cofactor recycling methods that are well-documented^63^, further lowering the cost of the biocatalytic synthesis as AMP is in general more cost-effective than ATP (**Supplementary Fig. 36**). The scalable one-pot enzymatic synthesis was demonstrated in the preparation of a number of novel analogs of E-64 at reasonable isolated yield, which enabled the determination of IC_50_ values, and use in cocrystallization studies. We also demonstrated the facile coupling of enzymatic and chemical syntheses in the preparative synthesis of cathepsin-L selective inhibitors CLIK-148 with >50% yield. Likewise, the chemoenzymatic cascade reactions can be simply adapted to synthesize the related inhibitor CAA0225, as well as the recently developed cathepsin C-selective inhibitor that is the hydrazide analog of **1**^72^ (**Fig. 5f** and **Supplementary Fig. 1**). Furthermore, Cp1B was also shown to use malic acid and fumaric acid as dicarboxylic acid substrates in the synthesis of **12** and **13**. Although not exhaustively explored here, the enzymes can be explored to synthesize non-epoxide containing pseudopeptides that have different applications (**Supplementary Fig. 31**-32). Lastly, given the high expression and robustness of both enzymes, one can envision that with directed evolution approaches, the substrate scopes can be further expanded to generate much more diverse amide-containing molecules^73^.

In conclusion, nearly four decades after its initial discovery^14^, a family of BGCs involved in the biosynthesis of **1** and related compounds has been characterized. The use of amide-bond forming enzymes as versatile biocatalysts illustrates the power of repurposing biosynthetic enzymes for catalysis. We anticipate that continuous discovery of amide-forming enzymes from natural product biosynthesis will expand the toolbox available to accelerate drug discovery and development of greener chemical synthesis.

## Methods

### Strains and culture conditions

*Aspergillus flavus* NRRL3357 was obtained from the Agricultural Research Service Culture Collection (NRRL). It was maintained in liquid PDB medium (PDA medium without agar) at 28 °C for isolation of genomic DNA. *Aspergillus nidulans* A1145 ΛEMΛST host was previously developed in our lab.^51^ The *A. nidulans* strain was grown at 28 °C in CD media (1 L: 10 g Glucose, 50 mL 20 × Nitrate salts, 1 mL Trace elements, pH 6.5, and 20 g/L Agar for solid cultivation) for sporulation or in CD-ST media (1L: 20 g Starch, 20 g Casamino acids, 50 mL 20 × Nitrate salts, 1 mL Trace elements, pH 6.5)^51^ for heterologous expression of gene cluster, compounds production and RNA extraction. *Escherichia coli* strains were cultivated either on lysogeny broth (LB) agar plates or in LB liquid medium. Growth media were supplied with antibiotics as required at the following concentrations: kanamycin (50 μg mL^−1^), and ampicillin (100 μg mL^−1^). *Saccharomyces cerevisiae* strain BJ5464-NpgA (*MATα ura3-52 his3-Δ200 leu2-Δ1 trp1 pep4::HIS3 prb1 Δ1.6R can1 GAL*) was used as the yeast host for *in vivo* homologous recombination to construct the *A. nidulans* plasmids.

### General molecular biology techniques

*E. coli* TOP10 cells were used for cloning, following standard recombinant DNA techniques. *E. coli* BL21(DE3) (Novagen) was used as the *E. coli* host for protein expression. DNA restriction enzymes were used as recommended by the manufacturer (New England Biolabs, NEB). Genomic DNA from all fungal strains was prepared using LETS isolation buffer (10 mM Tris-HCI, pH 8.0, 20 mM EDTA, 0.5% SDS, 0.1 M LiCl). PCR was performed using Q5 High-Fidelity DNA Polymerase (NEB). The gene-specific primers are listed in Supplementary Table 2. PCR products were confirmed by DNA sequencing. For isolation of RNA from *A. nidulans* transformants, the strains were grown on CD-ST liquid for 3 days at 28 °C. The RNA extraction steps were performed using RiboPure™ Yeast RNA Isolation Kit (Ambion) following the manufacturer’s instructions. Residual genomic DNA in the extracts was digested by DNase I (2 U/mL) (Invitrogen) at 37 °C for 4 hours. SuperScript III First-Strand Synthesis System (Invitrogen) was used for cDNA synthesis with Oligo-dT primers following directions from the user manual.

### Heterologous expression in *A. nidulans*

For heterologous expression in *A. nidulans*, three plasmid vectors, pYTU, pYTP, and pYTR containing auxotrophic markers for uracil (*pyrG*), pyridoxine (*pyroA*), and riboflavin (*riboB*), respectively, were used to construct plasmids for *A. nidulans* heterologous expression. Genes in the *cp1* and *cp2* cluster were amplified with PCR from the genomic DNA of *A. flavus*. The *gpdA* promoters from *Penicillium oxalicum (*constitutive *POgpdA), A. niger* (constitutive *gpdA*, *glaA* induced by starch) and *Penicillium expansum (*constitutive *PEgpdA)* were amplified by PCR. pYTP and pYTR were digested with *PacI*/*NotI*. pYTU was digested with *PacI*/*NotI* (keep *glaA* on vector). The amplified gene fragments and the corresponding vectors were co-transformed into *S. cerevisiae* strain BJ5464-NpgA for homologous recombination. The yeast plasmids were extracted using Zymoprep^TM^ Yeast Plasmid Miniprep I (Zymo Inc. USA), and then electrically transformed into *E. coli* TOP10 to isolate single plasmids. The plasmids were extracted from *E. coli* using the Zyppy^TM^ Plasmid Miniprep Kit (Zymo Research) and confirmed with sequencing by Largen and Primordium Lab. The preparation of the protoplasts of *A. nidulans* and transformation was described in the supplementary methods in Supplementary Information.

### Proteins expression and purification

The intron free ORFs of Cp1A, Cp1B, Cp1D, Cp2B, Cp2C, and Cp2D were amplified by PCR using cDNA from the corresponding *A. nidulans* transformant as a template, and ligated to linear expression vector pET28a via Gibson Assembly (New England Biolabs, NEB), according to the manufacturer’s protocol. *mfaA* gene obtained from the NCBI database, were codon-optimized based on the codon preference of *E. coli* by Integrated DNA Technologies (IDT), Inc. The gene for CHU (Polyphosphate kinase) was also synthesized by IDT. The plasmids were then transformed into *E. coli* BL21(DE3) individually and grown overnight in 5 mL of LB medium with 50 μg/mL kanamycin at 37 °C. The overnight cultures were used as seed cultures for 1 L fresh LB media containing 50 μg/mL kanamycin and incubated at 37 °C until the OD_600_ reached 0.8. The cultures were cooled on ice, before addition of 0.1 mM isopropyl-β-D-thiogalactopyranoside (IPTG, GoldBio, USA) to induce protein expression. The expression was performed at 16 °C for 20 h, 220 rpm. *E. coli* cells were harvested by centrifugation at 5300 rpm for 15 min and resuspended in 30 mL A10 buffer (50 mM sodium phosphate buffer, 150 mM NaCl, 10 mM imidazole, pH 8.0) containing 1 tablet of PierceTM protease inhibitor (Thermo Scientific). The cell suspension was lysed on ice by sonication and the lysate was centrifuged at 17,000 g for 30 min at 4°C to remove the insoluble cellular debris. The recombinant hexa-His-tagged proteins were purified individually from corresponding soluble fractions by affinity chromatography with Ni-NTA agarose resin (Qiagen) according to the manufacturer’s instructions. As for preparing the protein sample for the crystallization, the protein was further loaded onto size exclusion chromatography (GE healthcare, Superdex 200) for further purification with the buffer of 50 mM Tris pH 8.0 with 100 mM NaCl.

The purified proteins were concentrated and exchanged into storage buffer (50 mM sodium phosphate buffer, 200 mM NaCl, 10% glycerol, pH 8.0) with Amicon Ultra-30K concentrator (Merck Millipore). SDS-PAGE was performed to check the protein purity and Bradford Protein Assay (Bio-Rad) was used to calculate protein concentration with bovine serum albumin (BSA, Sigma-Aldrich) as standard. The proteins were used for the crystallization (Cp1B) or aliquoted and stored at −80 °C until used in in vitro assays. The plasmids used for protein purification are listed in Supplementary Table 3. See Supplementary Fig. 6, and Fig. 9 for SDS–PAGE analysis.

### In vitro assays

For *in vitro* assay for Cp1A and MfaA, 100 μL reactions were performed at 30 °C for 3 h, in 50 mM sodium phosphate buffer (pH 8.0) containing 0.2 mM FeSO_4_, 2 mM αKG, 2 mM ascorbate, 1 mM of substrate, and 10 μM of Cp1A or MfaA. The reaction mixture in the absence of protein was prepared as the negative control. Enzyme reactions were quenched by adding 100 µL acetonitrile and centrifuged at 17,000 *g* for 5 min, which was subjected to 3-nitrophenylhydrazine (3-NPH) derivatization.^74^ As for 3-NPH derivatization to directly detect 1,2-dicarboxylic acid: all reagents for derivatization were freshly prepared prior to use. 50 μL volumes of reaction mixture were sequentially treated with 50 μL of 50 mM 3-NPH (Sigma-Aldrich) in MeOH–H_2_O (70:30, v/v), 50 μL of 50 mM EDC (Oakwood Chemical) in MeOH–H_2_O (70:30, v/v), and 50 μL of 7% *v/v* pyridine (Acros Organics) in MeOH– H_2_O (70:30, v/v) and mixed thoroughly. Derivatization mixtures were incubated at 37 °C for 30 min and centrifuged at 17,000 *g* for 10 minutes. The supernatant was subjected to Quadrupole time-of-flight (QTOF) analysis with an Agilent Quadrupole Time of Flight LC/MS (6545 LC/Q-TOF) with a reverse-phase column (Agilent Poroshell, 120 EC-C18, 2.7 μm, 3.0 × 50 mm) using positive-mode electrospray ionization with 1% MeCN-H_2_O (containing 0.1% formic acid) for first 2 minutes, then a linear gradient of 1–95% for 9 minutes, followed by 95% MeCN for 3 minutes with a flow rate of 0.6 ml min^−1^.

For *in vitro* assay for Cp1B and Cp2B, assays were carried out in 50 mM sodium phosphate buffer (pH 8.0), containing 25 μM enzyme, 5 mM *trans*-epoxy-succinic acid or analog, 2.5 mM amino acid, ATP cofactor (10 mM), MgCl_2_ (10 mM) in 100 μL of solution at 30 °C for 16 h. The reaction was then quenched with 120 μL MeCN and centrifuged at 17,000 *g* for 5 min. The supernatant was subjected to QTOF analysis with the same conditions as mentioned above, or analyzed by a Agilent LC/MSD iQ with a reverse-phase column (AgilentTM InfinityLab Poroshell 120 Aq-C18, 2.7 μm, 100 Å, 2.1 × 100 mm) using positive and negative mode electrospray ionization with a linear gradient of 1-99% MeCN-H_2_O supplemented with 0.1% (v/v) formic acid with a flow rate of 0.6 mL/min. The detailed gradient conditions are shown in each legend of Supplementary Figures. The % conversion shown in Fig. 3 was estimated based on the calibration curves of enzymatically prepared standards that was generated from peak areas at 204 nm by HPLC. As for the determination of d.r values, the sample was analyzed by chiral analytical HPLC with a CHIRALPAK^®^ IA-3 column (150 x 4.6 mm, 3 μm) at room temperature (flow rate 1 mL/min, 40% MeCN–H_2_O with 0.1% trifluoroacetic acid).

Owing to the insolubility of Cp1C in *E. coli* BL21(DE3), the decarboxylase Cp2C from *cp2* cluster was expressed for characterization instead. *In vitro* assays of Cp2C were performed in 50 μL of 50 mM sodium phosphate buffer (pH 8.0), containing 50 μM of Cp2C, 100 μM PLP, 2 mM l-amino acid such as l-ornithine, l-lysine, and l-arginine at 30 °C. After 1 hours at 30 °C, all reactions were quenched with 50 μL MeCN, centrifuged at 17,000 *g* for 5 min, which was subjected to LC-MS analysis after dansyl derivatization with dansyl chloride (TCI). As for dansyl derivatization to detect decarboxylated product, 1 M borate buffer (pH 8.0) and dansyl chloride solution (10 mM final concentration) were added to the mixture. After incubation at 30 °C for 1 hour, the resultant reaction mixture was centrifuged at 17,000 *g* for 5 min. The resultant supernatant was subjected to QTOF analysis using the same gradient methods described above.

*In vitro* assay of Cp1D was performed in 50 mM sodium phosphate buffer (pH 8.0). The typical reaction contains 25 μM of Cp1D, 2.5 mM mono-amide such as **14** or **15**, 5 mM amine, 10 mM ATP, and 10 mM MgCl_2_ in 100 μL of solution. After incubation at 30 °C for 16 h, the subsequent sample preparation and QTOF or LC/MS (Agilent LC/MSD iQ) analysis followed the same methods as described for the Cp1B reaction. The detailed gradient conditions for LC/MS analysis are shown in each legend in Supplementary Figures. The % conversion shown in Fig. 4 was estimated based on the HPLC peak area ratio of product versus substrate **15** at UV 204 nm (% Conversion = (peak area of product / (peak area of **15** + peak area of product)) × 100%).

As for Cp1B mutants, the plasmids pML8010 containing the wild-type *cp1B* gene, was used as the template for PCR-based site-directed mutagenesis. DNA sequencing was used to confirm the identities including the mutated positions of the expression plasmids. Following expression and purification, Cp1B mutants were subjected to activity assays as described above.

### Stepwise enzymatic assay with Cp1A/MfaA and Cp1B for synthesis of (2*S*,3*S*)-*t*-ES-l-isoleucine (14)

Purified Cp1A/MfaA were added to 50 mM sodium phosphate buffer (pH 8.0) in 50 mL containing 0.2 mM FeSO_4_, 2 mM αKG, 2 mM ascorbate, 1 mM of fumaric acid, and 10 μM of Cp1A or MfaA at 30 °C for 16 h. Then the protein was removed by Amicon concentrators (Millipore). Subsequently 10 μM enzyme, 2.5 mM l-isoleucine, ATP cofactor (10 mM), MgCl_2_ (10 mM) were added followed by incubation at 30°C for 16 h. The enzyme in reaction solution was removed by ultrafiltration. The obtained filtrate was adjusted to pH around 2∼3 with 5 M H_2_SO_4_ solution and extracted with ethyl acetate. The solvent was subsequently removed under reduced pressure and further purified via semi-preparative HPLC. The enzymatically prepared pure product was subjected to NMR analysis as well as analyzed by chiral analytical HPLC with a CHIRALPAK^®^ IA-3 column (150 x 4.6 mm, 3 μm) at room temperature (flow rate 1 mL/min, 40% MeCN–H_2_O with 0.1% trifluoroacetic acid).

### Coupled activity assay with Cp1B and Cp1D

Coupled *in vitro* activity assay for Cp1B with Cp1D was performed in 100 μL of 50 mM sodium phosphate buffer (pH 8.0) which contains 25 μM Cp1B, 25 μM Cp1D, 5 mM (±)-*trans*-epoxy-succinic acid, 2.5 mM amino acid, 5 mM amine, 10 mM ATP cofactor, and 10 mM MgCl_2_. The reaction mixture was incubated at 30 °C for 16 h before quenching the reaction.. The subsequent sample preparation and LC/MS analysis followed the same methods as described for the Cp1B reaction.

### Steady-state kinetic analysis of Cp1D and Cp2D

100-μl reaction mixtures containing 5 μM Cp1D or Cp2D, 10 mM agmatine, 10 mM ATP, and 10 mM MgCl_2_ and the different concentrations of (2*S,*3*S*)-*t*-ES-Ile (**14**) (0.2mM to 5.0 mM) or (2*R,*3*R*)-*t*-ES-Ile (0.2 mM to 3.0 mM) in 50 mM sodium phosphate buffer (pH8.0) were incubated at 30 °C for 5 min. Then, the reaction was quenched with an equal volume of cold acetonitrile. Protein was precipitated and removed by centrifugation and the supernatant analyzed by HPLC. The formation of **6** was estimated by a standard curve of **6** that was generated from peak areas at 204 nm by HPLC. Data fitting with nonlinear regression was performed using GraphPad Prism 9, and apparent *K*_M_ and *k*_cat_ values represent mean ± s.d. of three independent replicates.

### ADP detection from Cp1B assay

The reaction mixtures (200 μL) contained 0.25 μM Cp1B, 10 mM ATP, 12 mM MgCl_2_, 300 *µ*M NADH, 500 *µ*M phosphoenolpyruvic acid (PEP), 41 units/mL pyruvate kinase (PK, Sigma), 59 units/mL lactate dehydrogenase (LDH, Sigma), 10 mM KCl and 100 mM Tris-HCl (pH 8.0) with 1 mM epoxy-succinic acid or other acid donors and 5 mM l-Phe. PK and LDH were stored in 10 mM HEPES (pH 7.0) with 100 mM KCl and 0.1 mM EDTA with 50% glycerol. PK requires potassium ion as an essential cofactor for its activity. The reaction mixture was incubated at 30 °C, and the consumption of NADH was monitored continuously for 60 min with a TECAN M200 plate reader by measuring the absorbance at 340 nm. The consumption of NADH reflects the formation of ADP on the incubation of Cp1B with tested 1,2-dicarboxylic acid substrates with excess amount of l-Phe. Therefore, the phosphorylation activity of Cp1B toward each substrate was derived from the consumption of NADH (the formation of ADP). The phosphorylation activity of Cp1B for (2*S*,3*S*)-***t*-ES** was set as 100% activity to calculate the relative activity for the other substrate.

For the steady-state kinetics analysis of Cp1B for the phosphorylation, the same procedure mentioned above was used except for various concentrations of (2*S*,3*S*)-***t*-ES** being used. Briefly, The reaction mixtures (100 μL) contained 1.0 μM Cp1B, 10 mM ATP, 12 mM MgCl_2_, 300 *µ*M NADH, 500 *µ*M phosphoenolpyruvic acid (PEP), 41 units/mL pyruvate kinase (PK, Sigma), 59 units/mL lactate dehydrogenase (LDH, Sigma), 10 mM KCl and 100 mM Tris-HCl (pH 8.0) with various concentration (0.04 mM to 1 mM) of (2*S*,3*S*)-***t*-ES** and 5 mM l-Phe. The reaction mixture was incubated at 30 °C, and the consumption of NADH at 10 min was used to derive the reaction rate (velocity) for the phosphorylation. Kinetic constants were derived from velocity versus substrate concentration data using a nonlinear regression fitting method with GraphPad Prism 9.

### Synthesis of *N*-succinyl proteinogenic amino acids and (2*R*,3*R*)-*t*-ES-Ile

All *N*-succinyl amino acids except for *N*-succinyl-Gly (Sigma-Aldrich) were synthesized following the protocol reported by Sumida et al^75^. A stirring solution of amino acid (2.0 mmol) and 20% NaOH (0.4 mL) in H_2_O (1.7 mL), succinic anhydride (0.21 g, 2.1 mmol) and 20% NaOH (4.2 mL) were separately added at room temperature. The reaction temperature was strictly maintained under 50 °C to prevent undesired racemization. After 2 h, 1 M HCl was added to adjust the pH of the reaction mixture to become pH around 3.0 and then extracted with equal volume ethyl acetate for two times. The combined organic layer was washed with brine (15 mL), dried over MgSO_4_, and filtered. The filtrate was evaporated to obtain semi-purified product. The solid residue was purified by HPLC using water (0.1% trifluoroacetic acid) and acetonitrile as mobile phase. (2*R*,3*R*)-*t*-ES-Ile was synthesized according to the reported procedure^76^ starting from (2*R*,3*R*)-3-(Ethoxycarbonyl)oxirane-2-carboxylic acid (Enamine Ltd). ^1^H NMR (500 MHz, DMSO): 8.68 ppm (1H, d, *J*=8.4 Hz), 4.21 ppm (1H, dd, *J*=5.7, 8.3 Hz), 3.76 ppm (1H, d, *J*=1.8 Hz), 3.46 ppm (1H, d, *J*=1.8 Hz), 1.81 ppm (1H, m), 1.19 ppm (1H, m), 1.41 ppm (1H, m), 0.86 ppm (6H, m). ^13^C NMR (125 MHz, DMSO) *δ* 172.3, 168.8, 165.4, 56.6, 52.3, 51.1, 36.3, 15.5, 24.7, 11.3. HRMS (ESI, M+H^+^) calculated for C_10_H_16_NO_6_^+^ 246.0972; found 246.0993. 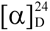 + 20° (*c* 0.1, MeOH).

### Hydroxamate-based colorimetric assay to test substrate specificity for Cp1D

Hydroxamate-based colorimetric assay was used to test substrate specificity toward *N*-succinyl amino acids for amide bond synthase Cp1D/Cp2D.^77^ The reaction was performed in 150 μL of 50 mM Tris buffer (pH 8.0) containing 20 μM of Cp1D or Cp2D, 15 mM of ATP, 5 mM of *N*-succinyl amino acid substrate, 200 mM hydroxylamine, and 10 mM MgCl_2_. The reaction is quenched after the incubation for 8 h at 30 °C by addition of equivalent volume (150 μL)of stopping solution (10% (w/v) FeCl_3_ and 3.3% (w/v) trichloroacetic acid dissolved in 0.7 M HCl). The precipitated enzyme was removed by centrifugation and 200 μL of the supernatant was transferred to a 96-well plate and the absorbance of the ferric-hydroxamate complex at 540 nm was measured by a Tecan M200 plate reader. The absorbance at 540 nm was used to calculate the relative activity, and the absorbance *N*-succinyl-l-Leu and *N*-succinyl-l-Tyr after the subtraction of that from each negative control (without Cp1D or Cp2D) were set as 100% activity for Cp1D and Cp2D, respectively.

### Enzymatic synthesis of *trans-*epoxysuccinyl amino acids

To obtain (2*S*,3*S*)-*trans-*epoxysuccinyl amino acids, large scale 20 mL in vitro reaction containing purified Cp1B or Cp2B (2.5 μM, 0.1 mol%) with 5 mM (±)-*trans*-epoxy-succinic acid, 2.5 mM amino acid, 10 mM ATP, 10 mM MgCl_2_ in 50 mM sodium phosphate buffer (pH 8.0) was performed at room temperature for 16 h. The protein was removed by Amicon concentrators (Millipore), and the concentrate was washed two times with three volumes of water. The filtrate was combined and was carefully adjusted to pH 2∼3 with 5 M H_2_SO_4_ solution. The acidified filtrate was further extracted with equal volume ethyl acetate for two times. The combined organic layer was washed with brine, dried over MgSO_4_, and filtered. The solvent was evaporated in vacuo to give a crude mixture that was further purified by HPLC (water and acetonitrile, both supplemented with 0.1% trifluoroacetic acid) on a Cosmosil, C18 AR-II column (5.0 µm, 10 ID X 250 mm, Nacalai Tesque) to afford at least >1.0 mg of the corresponding product with varying isolated yields. Thirty-eight examples were characterized by NMR (**Supplementary Tables 17-54** and **Supplementary Figs. 94-283**).

### Chemical analysis and compound isolation

For small scale metabolite analysis in *A. nidulans*, transformants were selected for on CD sorbitol agar (2% glucose as carbon source) appropriately supplemented with riboflavin, uracil, and/or pyridoxine for the set of plasmids introduced. The CD-ST agar was inoculated with spores and incubated at 28 °C for 4 days. The agar was then collected and extracted with acetone for 30 min with sonication. After centrifugation, the supernatant (200 μL) was concentrated and then re-suspended in methanol (100 μL). QTOF analyses were performed with the same method described in in vitro reaction of Cp1B. Some LC/MS analyses were performed on a Agilent LC/MSD iQ (AgilentTM InfinityLab Poroshell 120 Aq-C18, 2.7 μm, 100 Å, 2.1 × 100 mm) using positive and negative mode electrospray ionization with a linear gradient of 1-99% CH_3_CN-H_2_O supplemented with 0.1% (v/v) formic acid in 13.25 min followed by 99% CH_3_CN for 3 min with a flow rate of 0.6 mL/min. For large-scale analysis, *A. nidulans* transformants were inoculated to 40 plates each of which contains 50 mL CDST agar plates, which were placed in a 28 °C incubator for 3–4 days. After 4 day, the solid agar cultures were cut into small pieces and was extracted extensively with acetone. The residual was loaded on a normal-phase CombiFlash^®^ system and was subjected to flash chromatography with a gradient of CH_2_Cl_2_/MeOH for initial separation. Metabolites of interest, tracked by analytical HPLC and LC/MS, were purified from the corresponding fractions by reverse-phase semipreparative HPLC with a COSMOSIL column with flow rate of 4 mL/min of solvents A (H_2_O with 0.1% trifluoroacetic acid) and B (acetonitrile). NMR spectra were obtained with a Bruker AV500 spectrometer with a 5 mm dual cryoprobe at the UCLA Molecular Instrumentation Center. (^1^HNMR 500 MHz, ^13^CNMR 125 MHz). High resolution mass spectra were also recorded on a 6545 QTOF high resolution mass spectrometer (UCLA Molecular Instrumentation Center).

### Enzymatic or chemoenzymatic synthesis of cysteine protease inhibitor at preparative scale

Large scale 20 mL in vitro reaction containing 5 mM (±)-*trans*-epoxy-succinate, 2.5 mM l-amino acid, 2.5 mM amine donor, Cp1B (2.5 μM, 0.1 mol%), and Cp1D (2.5 μM, 0.1 mol%) was carried out at 30 °C for 16 h. The protein was removed by Amicon concentrators (Millipore), and the concentrate was washed two times with three volumes of water. For diamine or alkanoamine as amine donor, the filtrate was evaporated in vacuo and then directly subject to reverse-phase CombiFlash system (Teledyne) with a gradient of MeCN–H_2_O (0-5 min 0%-5% MeCN; 5-10 min 5%-20% MeCN, 10-20 min 20%-60% MeCN, 20-25 min 60% B, 25-35 min 100% B). Fractions containing target compound were combined and further purified with HPLC with a COSMOSIL column (Nacalai Tesque Inc., 5C18-AR-II, 10 ID x 250mm) with flow rate of 4 mL/min of solvents A (H_2_O with 0.1% trifluoroacetic acid) and B (acetonitrile). For other amine donors with hydrophobic terminal, the filtrate was combined, and pH was adjusted to around 2∼3 with 5 M H_2_SO_4_ solution. The acidified filtrate was further extracted with equal volume ethyl acetate for two times. The combined organic layer was washed with brine, dried over MgSO_4_, and filtered. The solvent was evaporated in vacuo to give a crude mixture that was further purified by HPLC. The isolated yield for each compound is shown in **Fig. 5e**. For example, E-64c was obtained 9 mg with 54% isolated yield from one-pot reaction. Spectroscopic and physical properties of E-64c were identical to those reported in the literature^78^: ^1^H NMR (DMSO-*d*_6_, 500 MHz) *δ* 0.83–0.89 ppm (12H, m), 1.27 (2H, q, *J* = 7.1 Hz), 1.45–1.58 (4H, m, CH_2_, 2 × CH), 3.04 (2H, m), 3.45 (1H, d, *J* = 1.8 Hz), 3.66 (1H, d, *J* = 1.9 Hz), 4.30 (1H, m), 8.03 (1H, t, *J* = 5.4 Hz), 8.56 (1H, d, *J* = 8.4 Hz); ^13^C NMR (DMSO-*d*_6_, 125 MHz) *δ* 171.0, 168.8, 164.9, 52.7, 51.2, 51.2, 41.1, 38.0, 36.8, 25.1, 24.3, 22.9, 22.4, 22.4, 21.7. HRMS (ESI, M+H^+^) calculated for C_15_H_27_N_2_O_5_^+^ 315.1914; found 315.1921.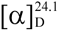 + 43° (*c* 0.1, MeOH).

To chemoenzymatically synthesize selective cathepsin L inhibitor CLIK-148, large scale 20 mL in vitro reaction containing 5 mM (±)-*trans*-epoxy-succinate, 2.5 mM l-Phe, 2.5 mM dimethylamine, Cp1B (2.5 μM, 0.1 mol%), and Cp1D (2.5 μM, 0.1 mol%) was carried out at 30 °C for 16 h. The protein was removed by following the same procedure as mentioned above. The subsequent filtrate acidification, and extraction, followed the same methods as described for amine donors with hydrophobic terminal. The solvent was evaporated in vacuo to give a crude mixture. The crude mixture was then dissolved in DMF. 2-(2-aminoethyl) pyridine (Combi-Blocks; 1.2 eq), HATU (Combi-Blocks; 1.2 eq), Et_3_N (Sigma-Aldrich; 3 eq), were added to the above DMF solution at 0 °C, the resulting mixture was stirred at room temperature until all the substrate was consumed. The reaction mixture was applied to reverse-phase HPLC chromatography using a COSMOSIL column (Nacalai Tesque Inc., 5C18 MS-II, 10 ID x 250mm, flow rate 4 mL/min, MeCN–H_2_O with 0.1% trifluoroacetic acid), to yield 11 mg of CLIK-148 (52% isolated yield).

### Cathepsin B inhibition assay

Cathepsin B Inhibitor Screening assay^60^ utilizes the ability of cathepsin B to cleave the synthetic AMC (7-Amino-4-methylcoumarin) based peptide substrate to release AMC, which can be easily quantified using a fluorometer or fluorescence microplate reader. In the presence of a cathepsin B inhibitor, the cleavage of the substrate is reduced/abolished resulting in decrease or total loss of the AMC fluorescence. Recombinant human procathepsin B (R&D systems) was activated to mature cathepsin B by incubation at 37 °C for 20 min in activation buffer (20 mM Na-acetate pH 5.5, 1 mM EDTA, 5 mM DTT, and 100 mM NaCl). Cathepsin B activity was then assayed in final buffer conditions of 0.04 ng/μL cathepsin B, 40 μM *Z*-Phe-Arg-AMC (Sigma-Aldrich), 40 mM citrate phosphate (pH 5.5), 1 mM EDTA, 100 mM NaCl, 5 mM DTT, and 0.01% Brij at 37 °C in triplicate. Cleavage of *Z*-Phe-Arg-AMC to generate fluorescent AMC was monitored at relative fluorescence unit (RFU) (excitation 360 nm, emission 460 nm) were recorded over a period of 30 min in infinite M200 PRO multimode microplate reader (Tecan). Choose two time points (T_1_ & T_2_) in the linear range of the plot and obtain the corresponding values for the fluorescence (RFU_1_ and RFU_2_). Calculate the slope for all test Inhibitor Samples and Enzyme Control by dividing the net ΔRFU (RFU_2_-RFU_1_) values with the time ΔT (T_2_-T_1_). Calculate the relative inhibition as follows:

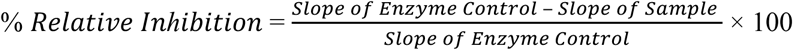

To test the effect of amino acid donor toward the inhibitory activity, the ***t*-ES** based compounds were obtained from coupled *in vitro* activity assay with 39 different amino acid donors while the amine donor was kept as agmatine. The assay contained 20 μM Cp1B, 20 μM Cp1D, 2 mM (±)-*trans*-epoxy-succinate, 39 different amino acid donor (1 mM), 1 mM agmatine as amine donor, 10 mM ATP, 10 mM MgCl_2_ in 100 μL of 50 mM sodium phosphate buffer (pH 8.0) at 30°C for overnight. The reaction was stopped by heat inactivation and centrifuged at 17,000 *g* for 5 mins. The supernatant was serially diluted and used (the final concentration is < 100 nM) for cathepsin B inhibition assay. The unnatural amino acid l-cyclopropylalanine (**a5**), l-cyclobutylalanine (**a6**), (*S*)-2-amino-2-cyclobutylacetic acid (**a9**), (*S*)-2-amino-2-cyclopentylacetic acid (**a10**), (*S*)-2-amino-2-cyclohexylacetic acid (**a11**), l-2-amino-3-ethylpentanoic acid (**a13**) were found to have the stronger inhibitory activity toward cathepsin B.

To test the effect of amine donor toward the inhibitory activity, the ***t*-ES** based compounds were obtained from coupled *in vitro* activity assay with 41 different amine donors while the amino acid was kept as **a10**. The assay contained 20 μM Cp1B, 20 μM Cp1D, 2 mM (±)-*trans*-epoxy-succinic acid, 1 mM **a10**, different amine donor (1 mM), 10 mM ATP cofactor, 10 mM MgCl_2_ in 100 μL of solution at 30°C for overnight. The reaction was treated as mentioned above, and the diluted sample was used for cathepsin B inhibition assay. Our screening assay found that **b4−b7**, **b13−b15**, **b17**, **b20**, **b24**, and **b26** as the amine donor exhibited potent inhibition.

Kinetic analyses of enzymatic synthesized inhibitions of cathepsin B were conducted to determine IC_50_. The concentrations of selected compounds and E-64 ranged from 2174 nM to 0.27 nM (2-fold serial dilution). IC_50_ values were calculated as the inhibitor concentration that reduced cathepsin B activity by 50%. Kinetic assays were run in a 96-well plate format. Data analysis was conducted using Prism GraphPad software.

### Crystallization of Cp1B, papain and papain-E64 analog complexes

The protein concentration used for crystallization of Cp1B was 15 mg/mL. The protein was incubated with ATP, MgCl_2_, and (±)-*trans*-epoxy-succinic acid in a ratio of 1:1:1:5 for 30 minutes. Then the protein mixed with the crystallization mother liquid in a ratio of 1:1 with a sitting drop vapor diffusion method. The crystals were observed after one week at the condition of 0.1 M MES/imidazole pH 6.5, 10% w/v PEG 20000, 20% v/v PEG MME 550, 0.02 M sodium l-glutamate, 0.02 M dl-alanine, 0.02 M glycine, 0.02 M dl-lysine HCl, 0.02 M dl-serine.

As for the crystallization of papain, twice-crystallized papain from papaya latex was purchased from Sigma as a buffered aqueous suspension approximately 25 mg/mL in protein concentration. Aliquots of this suspension were mixed with methanol, at a 1:2 volume ratio of papain suspension to methanol, in the sample wells of sitting drop crystallization trays allowing up to 30 μL sample volume per well. For all crystallization experiments, a total volume of 15 μL was targeted, though crystals were successfully grown in up to 30 μL volumes. These drops were incubated against a reservoir solution containing 59% methanol and 889 mM NaCl. The crystal used for determination of the unliganded papain structure was grown as above; all others were grown from seeds. For seeding, crystals were propagated by crushing up previously grown papain crystals by repeated pipetting in their mother liquor and transferring small fragments to freshly prepared sitting drops using a strand of horsehair. Papain crystals would typically appear in this condition between 48 and 72 hours without seeding, but formed in 24 hours if seeded. All crystals adopted a prismatic, diamond-shaped morphology.

Papain was also co-crystallized with E-64 (**1**), E-64d, and (2*S*,3*S*)-***t*-ES-a9-b7** using the same protocol as above, with the addition of the chosen inhibitor compound dissolved in solution to the crystallization well. **1** was purchased from Sigma and dissolved to 1.25 mg/mL (3.5 mM) in 66% methanol, E-64d was purchased from Selleck Chemicals and dissolved to 1.25 mg/mL (3.7 mM) in 66% methanol, and approximately 1 mg purified (2*S*,3*S*)-***t*-ES-a9-b7** was dissolved to 2.5 mg/mL (6.8 mM) in 66% methanol. For co-crystallization experiments with each compound, 2.4 μL of each compound solution was added to the crystallization well and mixed with protein solution. For these trials, the concentration of papain in each well was 0.35 mM, such that the estimated molar excess of inhibitor was 1.4 x for **1**, 1.5 x for E-64d, and 2.7 x for (2*S*,3*S*)-***t*-ES-a9-b7**. For all co-crystallization experiments, crystals were seeded with fragments of unliganded papain crystals as described above and appeared after approximately 24 hours. Crystal development in the presence of **1** or analogs was inconsistent without seeding.

### Diffraction data collection

The crystals of Cp1B were flash-cooled and stored in liquid nitrogen. The data were collected at beamline 17-ID-2 at the National Synchrotron Light Source II. Diffraction data were collected at the wavelength of 0.97933 Å.

The crystals of papain and papain with the inhibitor including E-64 (**1**), E-64d, and (2*S*,3*S*)-***t*-ES-a9-b7** were mounted on Mitegen loops and flash frozen under a 100 K nitrogen stream. Full X-ray diffraction datasets were acquired using a Rigaku FRE+ rotating anode X-ray diffractometer using a Cu Kα source (emitting X-ray photons 1.54 Å in wavelength) and equipped with a Rigaku HTC detector. All rotation datasets were collected taking 2-minute integrated exposures over oscillations of 0.5 degrees per exposure totaling approximately 26 hours of data collection per crystal, at a detector distance of either 78 or 74 mm. This configuration enabled visualization of reflections up to 1.4 Å in resolution at the detector edge.

### Data processing, structure determination, and refinement

The diffraction data for the Cp1B crystals was indexed, integrated, and scaled using the XDS package^79^. The structure of Cp1B was solved by molecular replacement. The Alphafold3^80^ predicted structure was used as the searching model by the Phaser embedded in the PHENIX suite^81^. Refinement was done by the refinement embedded in the PHENIX suite. The statistics are summarized in Table S4. The coordinate of the model was validated by Coot^82^. The picture is drawn by Pymol.

Data frames in OSC file format for papain and papain with the inhibitor were reduced in XDS, reflection intensities were scaled in XSCALE, and converted to MTZ format using XDSCONV.^79,83^ Reflections extending to a resolution of 1.4 Å for each inhibitor-bound structure, and to 1.5 Å for the unbound papain structure, were included for integration, phasing, and refinement. Phases were retrieved by molecular replacement using Phaser-MR through the PHENIX GUI with a known X-ray diffraction structure of papain determined to 1.65 Å resolution (PDB ID: 9PAP).^84–86^ For residue discrepancies between the protein sequence in this PDB entry and the sequence of papain reported in Uniprot (Accession code: P00784), the sequence from Uniprot was adopted into the model for refinement.

Refinement of each structure was carried out in PHENIX adopting the following strategy: PHENIX was instructed to refine XYZ coordinates against both reciprocal space data and real space maps, occupancies, and individual B-factors. All protein and ligand atoms were treated as anisotropic in B-factor refinement, though solvent atoms remained isotropic. No hydrogen atoms were modeled on any molecule. Three cycles of such refinement were run prior to inspecting the agreement between the atomic coordinates and electron density map in COOT, and building solvent molecules or adjusting side-chain positions to satisfy disagreements revealed in the difference Fourier map.^82^ Water molecules were built in positions marked by positive difference Fourier peaks exceeding 3 sigma levels where hydrogen-bonding partners were present within 2.5 – 3.3 Å proximity. Methanol molecules were only built in sites where a water molecule did not fully abolish the difference Fourier density upon subsequent refinement cycles, and where the carbon atom of the methanol molecule would not exist within 3.3 Å of any other non-bonded atoms. Following model adjustments, the coordinates were saved and the same refinement protocol in PHENIX repeated.

For structures of papain complexed with **1** or its analogs, prominent positive difference Fourier density indicating presence of a molecule bound to the active cysteine 25 typically developed after one or two of these iterations. The respective ligand was modeled into this density using pre-existing monomers in the CCP4 library for **1** and E-64d, and a custom ligand built using COOT’s ligand builder for (2*S*,3*S*)-***t*-ES-a9-b7**. Restraints for (2*S*,3*S*)-***t*-ES-a9-b7** were generated using eLBOW in the PHENIX GUI, with the PDB file for the custom ligand modeled in COOT as input.^87^ CIF files for each ligand were input as restraints for subsequent refinement of each respective complex structure, a bond length of 1.8 Å, with an allowed standard deviation of 0.1 Å, was enforced for the covalent linkage between the sulfur atom of Cys 25 and the ligand’s carbon C2, and the occupancy of all atoms in the ligand was refined in PHENIX as a single group. As additional validation, ligand-omit maps were generated for each ligand-bound structure by deleting the ligand from the model and refining the original MTZ file naïve to the presence of the ligand against it.

## Data availability

The data that support the findings of this study are available within the paper and its Supplementary Information, or are available from the corresponding author upon reasonable request. The atomic coordinates of Cp1B, apo-papain, papain with **1**, papain with E-64d, and papain with (2*S*,3*S*)-***t*-ES-a9-b7** have been deposited in the Protein Data Bank (http://www.rcsb.org) under the accession code 9CJN, 9CLH, 9CKT, 9CKW, and 9CKY respectively.

## Acknowledgement

This work was supported by the Emerging Pathogen Initiative from the Howard Hughes Medical Institute to Y.T to J.A.R. The authors thank the staff of beamline 17-ID-2 at the National Synchrotron Light Source II for access and help with the X-ray data collection. We thank Chuhang Luo for help with cysteine protease inhibition assays.

## Author contributions

M.L.,M.O., and Y.T. developed the hypothesis and conceived the idea for the study. M.L., M.O., N.V., J.A.R., and Y.T. designed the experiments. M.L. performed all *in vivo*, *in vitro* experiments, and cysteine protease inhibition assays as well as compound synthesis, isolation and characterization. M.L. and M.O. performed bioinformatic analysis and identified the biosynthetic gene clusters. X. Z. and N.W.V. preformed structural biology experiments. M. O., X. Z., N.W.V., and J.A.R. analyzed the crystal structures. All authors analyzed and discussed the results. M.L., M.O., and Y.T. prepared the manuscript.

## Additional information

Supplementary Information is available in the online version of the paper.

Correspondence and requests for materials should be addressed to Y. T. and M.O.

