## Supporting Information for "Biosynthetic characterization and combinatorial biocatalysis of the cysteine protease inhibitor E-64"

### Table of Contents

**Complemented Experimental Procedures**
1. Preparation of protoplast of *A. nidulans* and transformations **S10**
2. Spectroscopic data of *N*-succinyl proteinogenic amino acids **S10**
3. Preparation of (2*S*,3*S*)-*t*-ES or (2*R*,3*R*)-*t*-ES standard **S12**
4. Amino acid sequences of recombinant proteins used in this study **S13**

**Supplementary Tables**
**Table 1**. Bioinformatic analysis of *cp1* gene cluster **S15**
**Table 2.** Primers used in this study **S16**
**Table 3.** Plasmids used in this study **S18**
**Table 4.** X-ray data collection and refinement statistics of Cp1B **S19**
**Table 5.** Statistics from crystallographic data reduction and refinement for papain with/without inhibitor **S20**
**Table 6.** Spectroscopic data of compound **E-64 (1)** **S21**
**Table 7.** Spectroscopic data of compound **CLIK-148 (3)** **S22**
**Table 8.** Spectroscopic data of compound **CPI-2 (4)** **S23**
**Table 9.** Spectroscopic data of compound **CPI-3 (5)** **S24**
**Table 10.** Spectroscopic data of compound **6** **S25**
**Table 11.** Spectroscopic data of compound **7** **S26**
**Table 12.** Spectroscopic data of compound **8** **S27**
**Table 13.** Spectroscopic data of compound **9** **S28**
**Table 14.** Spectroscopic data of compound **10** **S29**
**Table 15.** Spectroscopic data of compound **12** **S30**
**Table 16.** Spectroscopic data of compound **13** **S31**
**Table 17.** Spectroscopic data of compound **14** **S32**
**Table 18.** Spectroscopic data of compound **15** **S33**
**Table 19.** Spectroscopic data of compound (2*S*,3*S*)-*t*-ES-Leu **S34**
**Table 20.** Spectroscopic data of compound (2*S*,3*S*)-*t*-ES-Val **S35**
**Table 21.** Spectroscopic data of compound (2*S*,3*S*)-*t*-ES-Tyr **S36**
**Table 22.** Spectroscopic data of compound (2*S*,3*S*)-*t*-ES-Trp **S37**
**Table 23.** Spectroscopic data of compound (2*S*,3*S*)-*t*-ES-a1 **S38**
**Table 24.** Spectroscopic data of compound (2*S*,3*S*)-*t*-ES-a2 **S39**
**Table 25.** Spectroscopic data of compound (2*S*,3*S*)-*t*-ES-a3 **S40**
**Table 26.** Spectroscopic data of compound (2*S*,3*S*)-*t*-ES-a4 **S41**
**Table 27.** Spectroscopic data of compound (2*S*,3*S*)-*t*-ES-a5 **S42**
**Table 28.** Spectroscopic data of compound (2*S*,3*S*)-*t*-ES-a6 **S43**
**Table 29.** Spectroscopic data of compound (2*S*,3*S*)-*t*-ES-a7 **S44**
**Table 30.** Spectroscopic data of compound (2*S*,3*S*)-*t*-ES-a8 **S45**
**Table 31.** Spectroscopic data of compound (2*S*,3*S*)-*t*-ES-a9 **S46**
**Table 32.** Spectroscopic data of compound (2*S*,3*S*)-*t*-ES-a10 **S47**
**Table 33.** Spectroscopic data of compound (2*S*,3*S*)-*t*-ES-a11 **S48**
**Table 34.** Spectroscopic data of compound (2*S*,3*S*)-*t*-ES-a12 **S49**
**Table 35.** Spectroscopic data of compound (2*S*,3*S*)-*t*-ES-a13 **S50**
**Table 36.** Spectroscopic data of compound (2*S*,3*S*)-*t*-ES-a14 **S51**
**Table 37.** Spectroscopic data of compound (2*S*,3*S*)-*t*-ES-a15 **S52**
**Table 38.** Spectroscopic data of compound (2*S*,3*S*)-*t*-ES-a16 **S53**
**Table 39.** Spectroscopic data of compound (2*S*,3*S*)-*t*-ES-a17 **S54**
**Table 40.** Spectroscopic data of compound (2*S*,3*S*)-*t*-ES-a18 **S55**

**Table 41.** Spectroscopic data of compound (2*S*,3*S*)-*t*-ES-a19 **S56**
**Table 42.** Spectroscopic data of compound (2*S*,3*S*)-*t*-ES-a20 **S57**
**Table 43.** Spectroscopic data of compound (2*S*,3*S*)-*t*-ES-a21 **S58**
**Table 44.** Spectroscopic data of compound (2*S*,3*S*)-*t*-ES-a22 **S59**
**Table 45.** Spectroscopic data of compound (2*S*,3*S*)-*t*-ES-a23 **S60**
**Table 46.** Spectroscopic data of compound (2*S*,3*S*)-*t*-ES-a24 **S61**
**Table 47.** Spectroscopic data of compound (2*S*,3*S*)-*t*-ES-a25 **S62**
**Table 48.** Spectroscopic data of compound (2*S*,3*S*)-*t*-ES-a26 **S63**
**Table 49.** Spectroscopic data of compound (2*S*,3*S*)-*t*-ES-a27 **S64**
**Table 50.** Spectroscopic data of compound (2*S*,3*S*)-*t*-ES-a28 **S65**
**Table 51.** Spectroscopic data of compound (2*S*,3*S*)-*t*-ES-a29 **S66**
**Table 52.** Spectroscopic data of compound (2*S*,3*S*)-*t*-ES-a30 **S67**
**Table 53.** Spectroscopic data of compound (2*S*,3*S*)-*t*-ES-a31 **S68**
**Table 54.** Spectroscopic data of compound (2*S*,3*S*)-*t*-ES-a32 **S69**
**Table 55.** Spectroscopic data of compound (2*S*,3*S*)-*t*-ES-a9-b7 **S70**
**Table 56.** Spectroscopic data of compound (2*S*,3*S*)-*t*-ES-a9-b12 **S71**
**Table 57.** Spectroscopic data of compound (2*S*,3*S*)-*t*-ES-a10-b9 **S72**
**Table 58.** Spectroscopic data of compound (2*S*,3*S*)-*t*-ES-a10-b13 **S73**
**Table 59.** Spectroscopic data of compound (2*S*,3*S*)-*t*-ES-a10-b26 **S74**
**Table 60.** Spectroscopic data of compound (2*S*,3*S*)-*t*-ES-Leu-b43 **S75**

**Supplementary Figures**

**Figure 1.** Other reported E-64 analogs and E-64 derived cysteine protease inhibitors **S76**
**Figure 2.** Significance and biosynthetic machinery of amide functionality **S77**
**Figure 3.** Amide bond formation in fumaryl dipeptides and penilumamide from NRPS **S78**
**Figure 4.** Limited examples of fungal ATP-grasp enzyme involved in natural product biosynthesis **S79**
**Figure 5.** E-64-like biosynthetic gene clusters are widely conserved in a plethora of fungi **S81**
**Figure 6.** Plasmids used for heterologous expression **S82**
**Figure 7.** Bioinformatic analysis of Cp1B **S83**
**Figure 8.** Non-E-64 biosynthetic gene clusters containing homologues of Cp1B and Cp1D **S84**
**Figure 9.** LC/MS analysis of extracts from the heterologous expression of *cp1* and *cp2* in *A. nidulans*. **S85**
**Figure 10.** SDS-PAGE gels of purified proteins used in this study **S86**
**Figure 11.** Absolute configuration of *t*-ES and substrate for Cp1A **S87**
**Figure 12.** Cp1B is a new ATP-grasp enzyme **S88**
**Figure 13.** D-amino acids are not substrate for Cp1B **S90**
**Figure 14.** Examination of the kinetic resolution potential for Cp1B **S91**
**Figure 15.** The crystal structure of Cp1B adopts a closed active site form **S92**
**Figure 16.** SDS-PAGE gels of Cp1B mutant **S93**
**Figure 17.** Previous work for the preparation of (2*S*, 3*S*)-*trans*-epoxy-succinate **S94**
**Figure 18.** Bioinformatic analysis of Cp1D **S95**
**Figure 19.** In vitro reaction of Cp1D with 14 and amines **S96**
**Figure 20.** Cp1D is a new family of diamide-forming amide bond synthetase not CoA ligase **S97**
**Figure 21.** Structure of unnatural amino acid tested but could not be accepted by Cp1B/Cp2B **S98**
**Figure 22.** HPLC traces of (2*S*,3*S*)-*t*-ES-amino acids formed by the Cp1B reaction **S100**
**Figure 23.** Substrate scope test for Cp1D against coupling of isopentylamine with synthesized 20 *N*-succinyl proteinogenic amino acids **S101**
**Figure 24.** HPLC traces of combinatorial biocatalysis of Cp1B and Cp1D with agmatine and different L-amino acids **S103**
**Figure 25.** Structures of amine nucleophiles which could not be accepted by Cp1D or Cp2D **S104**
**Figure 26.** Exploration of substrate scope of Cp1D/Cp2D with synthesized twenty *N*-succinyl proteinogenic amino acids **S105**

**Figure 27.** HPLC traces from the in vitro reaction of Cp1D with (2*S*,3*S*)-*t*-ES-L-Phe (**15**) and different amines **S106**
**Figure 28.** LC-MS analysis of one-pot synthesis E-64 analogs by Cp1B and Cp1D in 96-well plate **S110**
**Figure 29.** Reported chemical synthesis for E-64 (**1**) and CLIK-148 (**3**) **S111**
**Figure 30.** HPLC traces of combinatorial biocatalysis with a10 and different amines **S112**
**Figure 31.** Dicarboxylic acids scope of Cp1B **S114**
**Figure 32.** Structures of (pseudo)dipeptides tested for the acid acceptor of Cp1D **S115**
**Figure 33.** In vitro cathepsin B inhibition assay of enzymatic synthesized inhibitors **S116**
**Figure 34.** Structures of the active site of papain (no inhibitor bound, panels a-b) and papain bound to E-64 analogs from X-ray diffraction (panels c-h). **S117**
**Figure 35.** In vitro assay of Cp2C **S119**
**Figure 36.** Preparative-scale synthesis of E-64c using ATP Regeneration system **S120**
**Figure 37.** $^{1}$H NMR spectrum of E-64 (**1**) in DMSO-$d_6$ **S121**
**Figure 38.** $^{13}$C NMR spectrum of E-64 (**1**) in DMSO-$d_6$ **S121**
**Figure 39.** HSQC spectrum of E-64 (**1**) in DMSO-$d_6$ **S122**
**Figure 40.** HMBC spectrum of E-64 (**1**) in DMSO-$d_6$ **S122**
**Figure 41.** $^{1}$H-$^{1}$H COSY spectrum of E-64 (**1**) in DMSO-$d_6$ **S123**
**Figure 42.** $^{1}$H NMR spectrum of E-64c (**2**) in DMSO-$d_6$ **S123**
**Figure 43.** $^{13}$C NMR spectrum of E-64c (**2**) in DMSO-$d_6$ **S124**
**Figure 44.** $^{1}$H NMR spectrum of CLIK148 (**3**) in $CDCl_3$ **S124**
**Figure 45.** $^{13}$C NMR spectrum of CLIK148 (**3**) in $CDCl_3$ **S125**
**Figure 46.** HSQC spectrum of CLIK148 (**3**) in $CDCl_3$ **S125**
**Figure 47.** HMBC spectrum of CLIK148 (**3**) in $CDCl_3$ **S126**
**Figure 48.** $^{1}$H-$^{1}$H COSY spectrum of CLIK148 (**3**) in $CDCl_3$ **S126**
**Figure 49.** $^{1}$H NMR spectrum of CPI-2 (**4**) in DMSO-$d_6$ **S127**
**Figure 50.** $^{13}$C NMR spectrum of CPI-2 (**4**) in DMSO-$d_6$ **S127**
**Figure 51.** HSQC spectrum of CPI-2 (**4**) in DMSO-$d_6$ **S128**
**Figure 52.** HMBC spectrum of CPI-2 (**4**) in DMSO-$d_6$ **S128**
**Figure 53.** $^{1}$H-$^{1}$H COSY spectrum of CPI-2 (**4**) in DMSO-$d_6$ **S129**
**Figure 54.** $^{1}$H NMR spectrum of CPI-3 (**5**) in DMSO-$d_6$ **S129**
**Figure 55.** $^{13}$C NMR spectrum of CPI-3 (**5**) in DMSO-$d_6$ **S130**
**Figure 56.** HSQC spectrum of CPI-3 (**5**) in DMSO-$d_6$ **S130**
**Figure 57.** HMBC spectrum of CPI-3 (**5**) in DMSO-$d_6$ **S131**
**Figure 58.** $^{1}$H-$^{1}$H COSY spectrum of CPI-3 (**5**) in DMSO-$d_6$ **S131**
**Figure 59.** $^{1}$H NMR spectrum of compound **6** in DMSO-$d_6$ **S132**
**Figure 60.** $^{13}$C NMR spectrum of compound **6** in DMSO-$d_6$ **S132**
**Figure 61.** HSQC spectrum of compound **6** in DMSO-$d_6$ **S133**
**Figure 62.** HMBC spectrum of compound **6** in DMSO-$d_6$ **S133**
**Figure 63.** $^{1}$H-$^{1}$H COSY spectrum of compound **6** in DMSO-$d_6$ **S134**
**Figure 64.** $^{1}$H NMR spectrum of compound **7** in DMSO-$d_6$ **S134**
**Figure 65.** $^{13}$C NMR spectrum of compound **7** in DMSO-$d_6$ **S135**
**Figure 66.** HSQC spectrum of compound **7** in DMSO-$d_6$ **S135**
**Figure 67.** HMBC spectrum of compound **7** in DMSO-$d_6$ **S136**
**Figure 68.** $^{1}$H-$^{1}$H COSY spectrum of compound **7** in DMSO-$d_6$ **S136**
**Figure 69.** $^{1}$H NMR spectrum of compound **8** in DMSO-$d_6$ **S137**
**Figure 70.** $^{13}$C NMR spectrum of compound **8** in DMSO-$d_6$ **S137**
**Figure 71.** HSQC spectrum of compound **8** in DMSO-$d_6$ **S138**
**Figure 72.** HMBC spectrum of compound **8** in DMSO-$d_6$ **S138**
**Figure 73.** $^{1}$H-$^{1}$H COSY spectrum of compound **8** in DMSO-$d_6$ **S139**
**Figure 74.** $^{1}$H NMR spectrum of compound **9** in DMSO-$d_6$ **S139**
**Figure 75.** $^{13}$C NMR spectrum of compound **9** in DMSO-$d_6$ **S140**

**Figure 76.** HSQC spectrum of compound **9** in DMSO-$d_6$ **S140**
**Figure 77.** HMBC spectrum of compound **9** in DMSO-$d_6$ **S141**
**Figure 78.** $^1$H-$^1$H COSY spectrum of compound **9** in DMSO-$d_6$ **S141**
**Figure 79.** $^1$H NMR spectrum of compound **10** in DMSO-$d_6$ **S142**
**Figure 80.** $^{13}$C NMR spectrum of compound **10** in DMSO-$d_6$ **S142**
**Figure 81.** HSQC spectrum of compound **10** in DMSO-$d_6$ **S143**
**Figure 82.** HMBC spectrum of compound **10** in DMSO-$d_6$ **S143**
**Figure 83.** $^1$H-$^1$H COSY spectrum of compound **10** in DMSO-$d_6$ **S144**
**Figure 84.** $^1$H NMR spectrum of compound **12** in DMSO-$d_6$ **S144**
**Figure 85.** $^{13}$C NMR spectrum of compound **12** in DMSO-$d_6$ **S145**
**Figure 86.** HSQC spectrum of compound **12** in DMSO-$d_6$ **S145**
**Figure 87.** HMBC spectrum of compound **12** in DMSO-$d_6$ **S146**
**Figure 88.** $^1$H-$^1$H COSY spectrum of compound **12** in DMSO-$d_6$ **S146**
**Figure 89.** $^1$H NMR spectrum of compound **13** in DMSO-$d_6$ **S147**
**Figure 90.** $^{13}$C NMR spectrum of compound **13** in DMSO-$d_6$ **S147**
**Figure 91.** HSQC spectrum of compound **13** in DMSO-$d_6$ **S148**
**Figure 92.** HMBC spectrum of compound **13** in DMSO-$d_6$ **S148**
**Figure 93.** $^1$H-$^1$H COSY spectrum of compound **13** in DMSO-$d_6$ **S149**
**Figure 94.** $^1$H NMR spectrum of compound **14** in DMSO-$d_6$ **S149**
**Figure 95.** $^{13}$C NMR spectrum of compound **14** in DMSO-$d_6$ **S150**
**Figure 96.** HSQC spectrum of compound **14** in DMSO-$d_6$ **S150**
**Figure 97.** HMBC spectrum of compound **14** in DMSO-$d_6$ **S151**
**Figure 98.** $^1$H-$^1$H COSY spectrum of compound **14** in DMSO-$d_6$ **S151**
**Figure 99.** $^1$H NMR spectrum of compound **15** in DMSO-$d_6$ **S152**
**Figure 100.** $^{13}$C NMR spectrum of compound **15** in DMSO-$d_6$ **S152**
**Figure 101.** HSQC spectrum of compound **15** in DMSO-$d_6$ **S153**
**Figure 102.** HMBC spectrum of compound **15** in DMSO-$d_6$ **S153**
**Figure 103.** $^1$H-$^1$H COSY spectrum of compound **15** in DMSO-$d_6$ **S154**
**Figure 104.** $^1$H NMR spectrum of compound **(2*S*,3*S*)-*t*-ES-Leu** in DMSO-$d_6$ **S154**
**Figure 105.** $^{13}$C NMR spectrum of compound **(2*S*,3*S*)-*t*-ES-Leu** in DMSO-$d_6$ **S155**
**Figure 106.** HSQC spectrum of compound **(2*S*,3*S*)-*t*-ES-Leu** in DMSO-$d_6$ **S155**
**Figure 107.** HMBC spectrum of compound **(2*S*,3*S*)-*t*-ES-Leu** in DMSO-$d_6$ **S156**
**Figure 108.** $^1$H-$^1$H COSY spectrum of compound **(2*S*,3*S*)-*t*-ES-Leu** in DMSO-$d_6$ **S156**
**Figure 109**. $^1$H NMR spectrum of compound **(2*S*,3*S*)-*t*-ES-Val** in DMSO-$d_6$ **S157**
**Figure 110.** $^{13}$C NMR spectrum of compound **(2*S*,3*S*)-*t*-ES-Val** in DMSO-$d_6$ **S157**
**Figure 111.** HSQC spectrum of compound **(2*S*,3*S*)-*t*-ES-Val** in DMSO-$d_6$ **S158**
**Figure 112.** HMBC spectrum of compound **(2*S*,3*S*)-*t*-ES-Val** in DMSO-$d_6$ **S158**
**Figure 113.** $^1$H-$^1$H COSY spectrum of compound **(2*S*,3*S*)-*t*-ES-Val** in DMSO-$d_6$ **S159**
**Figure 114.** $^1$H NMR spectrum of compound **(2*S*,3*S*)-*t*-ES-Tyr** in DMSO-$d_6$ **S159**
**Figure 115.** $^{13}$C NMR spectrum of compound **(2*S*,3*S*)-*t*-ES-Tyr** in DMSO-$d_6$ **S160**
**Figure 116.** HSQC spectrum of compound **(2*S*,3*S*)-*t*-ES-Tyr** in DMSO-$d_6$ **S160**
**Figure 117.** HMBC spectrum of compound **(2*S*,3*S*)-*t*-ES-Tyr** in DMSO-$d_6$ **S161**
**Figure 118.** $^1$H-$^1$H COSY spectrum of compound **(2*S*,3*S*)-*t*-ES-Tyr** in DMSO-$d_6$ **S161**
**Figure 119.** $^1$H NMR spectrum of compound **(2*S*,3*S*)-*t*-ES-Trp** in DMSO-$d_6$ **S162**
**Figure 120.** $^{13}$C NMR spectrum of compound **(2*S*,3*S*)-*t*-ES-Trp** in DMSO-$d_6$ **S162**
**Figure 121.** HSQC spectrum of compound **(2*S*,3*S*)-*t*-ES-Trp** in DMSO-$d_6$ **S163**
**Figure 122.** HMBC spectrum of compound **(2*S*,3*S*)-*t*-ES-Trp** in DMSO-$d_6$ **S163**
**Figure 123.** $^1$H-$^1$H COSY spectrum of compound **(2*S*,3*S*)-*t*-ES-Trp** in DMSO-$d_6$ **S164**
**Figure 124.** $^1$H NMR spectrum of compound **(2*S*,3*S*)-*t*-ES-a1** in DMSO-$d_6$ **S164**
**Figure 125.** $^{13}$C NMR spectrum of compound **(2*S*,3*S*)-*t*-ES-a1** in DMSO-$d_6$ **S165**
**Figure 126.** HSQC spectrum of compound **(2*S*,3*S*)-*t*-ES-a1** in DMSO-$d_6$ **S165**

**Figure 127.** HMBC spectrum of compound **(2*S*,3*S*)-*t*-ES-a1** in DMSO-$d_6$ **S166**
**Figure 128.** $^1$H-$^1$H COSY spectrum of compound **(2*S*,3*S*)-*t*-ES-a1** in DMSO-$d_6$ **S166**
**Figure 129.** $^1$H NMR spectrum of compound **(2*S*,3*S*)-*t*-ES-a2** in DMSO-$d_6$ **S167**
**Figure 130.** $^{13}$C NMR spectrum of compound **(2*S*,3*S*)-*t*-ES-a2** in DMSO-$d_6$ **S167**
**Figure 131.** HSQC spectrum of compound **(2*S*,3*S*)-*t*-ES-a2** in DMSO-$d_6$ **S168**
**Figure 132.** HMBC spectrum of compound **(2*S*,3*S*)-*t*-ES-a2** in DMSO-$d_6$ **S168**
**Figure 133.** $^1$H-$^1$H COSY spectrum of compound **(2*S*,3*S*)-*t*-ES-a2** in DMSO-$d_6$ **S169**
**Figure 134.** $^1$H NMR spectrum of compound **(2*S*,3*S*)-*t*-ES-a3** in DMSO-$d_6$ **S169**
**Figure 135.** $^{13}$C NMR spectrum of compound **(2*S*,3*S*)-*t*-ES-a3** in DMSO-$d_6$ **S170**
**Figure 136.** HSQC spectrum of compound **(2*S*,3*S*)-*t*-ES-a3** in DMSO-$d_6$ **S170**
**Figure 137.** HMBC spectrum of compound **(2*S*,3*S*)-*t*-ES-a3** in DMSO-$d_6$ **S171**
**Figure 138.** $^1$H-$^1$H COSY spectrum of compound **(2*S*,3*S*)-*t*-ES-a3** in DMSO-$d_6$ **S171**
**Figure 139.** $^1$H NMR spectrum of compound **(2*S*,3*S*)-*t*-ES-a4** in DMSO-$d_6$ **S172**
**Figure 140.** $^{13}$C NMR spectrum of compound **(2*S*,3*S*)-*t*-ES-a4** in DMSO-$d_6$ **S172**
**Figure 141.** HSQC spectrum of compound **(2*S*,3*S*)-*t*-ES-a4** in DMSO-$d_6$ **S173**
**Figure 142.** HMBC spectrum of compound **(2*S*,3*S*)-*t*-ES-a4** in DMSO-$d_6$ **S173**
**Figure 143.** $^1$H-$^1$H COSY spectrum of compound **(2*S*,3*S*)-*t*-ES-a4** in DMSO-$d_6$ **S174**
**Figure 144.** $^1$H NMR spectrum of compound **(2*S*,3*S*)-*t*-ES-a5** in DMSO-$d_6$ **S174**
**Figure 145.** $^{13}$C NMR spectrum of compound **(2*S*,3*S*)-*t*-ES-a5** in DMSO-$d_6$ **S175**
**Figure 146.** HSQC spectrum of compound **(2*S*,3*S*)-*t*-ES-a5** in DMSO-$d_6$ **S175**
**Figure 147.** HMBC spectrum of compound **(2*S*,3*S*)-*t*-ES-a5** in DMSO-$d_6$ **S176**
**Figure 148.** $^1$H-$^1$H COSY spectrum of compound **(2*S*,3*S*)-*t*-ES-a5** in DMSO-$d_6$ **S176**
**Figure 149.** $^1$H NMR spectrum of compound **(2*S*,3*S*)-*t*-ES-a6** in DMSO-$d_6$ **S177**
**Figure 150.** $^{13}$C NMR spectrum of compound **(2*S*,3*S*)-*t*-ES-a6** in DMSO-$d_6$ **S177**
**Figure 151.** HSQC spectrum of compound **(2*S*,3*S*)-*t*-ES-a6** in DMSO-$d_6$ **S178**
**Figure 152.** HMBC spectrum of compound **(2*S*,3*S*)-*t*-ES-a6** in DMSO-$d_6$ **S178**
**Figure 153.** $^1$H-$^1$H COSY spectrum of compound **(2*S*,3*S*)-*t*-ES-a6** in DMSO-$d_6$ **S179**
**Figure 154.** $^1$H NMR spectrum of compound **(2*S*,3*S*)-*t*-ES-a7** in DMSO-$d_6$ **S179**
**Figure 155.** $^{13}$C NMR spectrum of compound **(2*S*,3*S*)-*t*-ES-a7** in DMSO-$d_6$ **S180**
**Figure 156.** HSQC spectrum of compound **(2*S*,3*S*)-*t*-ES-a7** in DMSO-$d_6$ **S180**
**Figure 157.** HMBC spectrum of compound **(2*S*,3*S*)-*t*-ES-a7** in DMSO-$d_6$ **S181**
**Figure 158.** $^1$H-$^1$H COSY spectrum of compound **(2*S*,3*S*)-*t*-ES-a7** in DMSO-$d_6$ **S181**
**Figure 159.** $^1$H NMR spectrum of compound **(2*S*,3*S*)-*t*-ES-a8** in DMSO-$d_6$ **S182**
**Figure 160.** $^{13}$C NMR spectrum of compound **(2*S*,3*S*)-*t*-ES-a8** in DMSO-$d_6$ **S182**
**Figure 161.** HSQC spectrum of compound **(2*S*,3*S*)-*t*-ES-a8** in DMSO-$d_6$ **S183**
**Figure 162.** HMBC spectrum of compound **(2*S*,3*S*)-*t*-ES-a8** in DMSO-$d_6$ **S183**
**Figure 163.** $^1$H-$^1$H COSY spectrum of compound **(2*S*,3*S*)-*t*-ES-a8** in DMSO-$d_6$ **S184**
**Figure 164.** $^1$H NMR spectrum of compound **(2*S*,3*S*)-*t*-ES-a9** in DMSO-$d_6$ **S184**
**Figure 165.** $^{13}$C NMR spectrum of compound **(2*S*,3*S*)-*t*-ES-a9** in DMSO-$d_6$ **S185**
**Figure 166.** HSQC spectrum of compound **(2*S*,3*S*)-*t*-ES-a9** in DMSO-$d_6$ **S185**
**Figure 167.** HMBC spectrum of compound **(2*S*,3*S*)-*t*-ES-a9** in DMSO-$d_6$ **S186**
**Figure 168.** $^1$H-$^1$H COSY spectrum of compound **(2*S*,3*S*)-*t*-ES-a9** in DMSO-$d_6$ **S186**
**Figure 169.** $^1$H NMR spectrum of compound **(2*S*,3*S*)-*t*-ES-a10** in DMSO-$d_6$ **S187**
**Figure 170.** $^{13}$C NMR spectrum of compound **(2*S*,3*S*)-*t*-ES-a10** in DMSO-$d_6$ **S187**
**Figure 171.** HSQC spectrum of compound **(2*S*,3*S*)-*t*-ES-a10** in DMSO-$d_6$ **S188**
**Figure 172.** HMBC spectrum of compound **(2*S*,3*S*)-*t*-ES-a10** in DMSO-$d_6$ **S188**
**Figure 173.** $^1$H-$^1$H COSY spectrum of compound **(2*S*,3*S*)-*t*-ES-a10** in DMSO-$d_6$ **S189**
**Figure 174.** $^1$H NMR spectrum of compound **(2*S*,3*S*)-*t*-ES-a11** in DMSO-$d_6$ **S189**
**Figure 175.** $^{13}$C NMR spectrum of compound **(2*S*,3*S*)-*t*-ES-a11** in DMSO-$d_6$ **S190**
**Figure 176.** HSQC spectrum of compound **(2*S*,3*S*)-*t*-ES-a11** in DMSO-$d_6$ **S190**
**Figure 177.** HMBC spectrum of compound **(2*S*,3*S*)-*t*-ES-a11** in DMSO-$d_6$ **S191**

**Figure 178.** $^1$H-$^1$H COSY spectrum of compound **(2*S*,3*S*)-*t*-ES-a11** in DMSO-$d_6$ **S191**
**Figure 179.** $^1$H NMR spectrum of compound **(2*S*,3*S*)-*t*-ES-a12** in DMSO-$d_6$ **S192**
**Figure 180.** $^{13}$C NMR spectrum of compound **(2*S*,3*S*)-*t*-ES-a12** in DMSO-$d_6$ **S192**
**Figure 181.** HSQC spectrum of compound **(2*S*,3*S*)-*t*-ES-a12** in DMSO-$d_6$ **S193**
**Figure 182.** HMBC spectrum of compound **(2*S*,3*S*)-*t*-ES-a12** in DMSO-$d_6$ **S193**
**Figure 183.** $^1$H-$^1$H COSY spectrum of compound **(2*S*,3*S*)-*t*-ES-a12** in DMSO-$d_6$ **S194**
**Figure 184.** $^1$H NMR spectrum of compound **(2*S*,3*S*)-*t*-ES-a13** in DMSO-$d_6$ **S194**
**Figure 185.** $^{13}$C NMR spectrum of compound **(2*S*,3*S*)-*t*-ES-a13** in DMSO-$d_6$ **S195**
**Figure 186.** HSQC spectrum of compound **(2*S*,3*S*)-*t*-ES-a13** in DMSO-$d_6$ **S195**
**Figure 187.** HMBC spectrum of compound **(2*S*,3*S*)-*t*-ES-a13** in DMSO-$d_6$ **S196**
**Figure 188.** $^1$H-$^1$H COSY spectrum of compound **(2*S*,3*S*)-*t*-ES-a13** in DMSO-$d_6$ **S196**
**Figure 189.** $^1$H NMR spectrum of compound **(2*S*,3*S*)-*t*-ES-a14** in DMSO-$d_6$ **S197**
**Figure 190.** $^{13}$C NMR spectrum of compound **(2*S*,3*S*)-*t*-ES-a14** in DMSO-$d_6$ **S197**
**Figure 191.** HSQC spectrum of compound **(2*S*,3*S*)-*t*-ES-a14** in DMSO-$d_6$ **S198**
**Figure 192.** HMBC spectrum of compound **(2*S*,3*S*)-*t*-ES-a14** in DMSO-$d_6$ **S198**
**Figure 193.** $^1$H-$^1$H COSY spectrum of compound **(2*S*,3*S*)-*t*-ES-a14** in DMSO-$d_6$ **S199**
**Figure 194.** $^1$H NMR spectrum of compound **(2*S*,3*S*)-*t*-ES-a15** in DMSO-$d_6$ **S199**
**Figure 195.** $^{13}$C NMR spectrum of compound **(2*S*,3*S*)-*t*-ES-a15** in DMSO-$d_6$ **S200**
**Figure 196.** HSQC spectrum of compound **(2*S*,3*S*)-*t*-ES-a15** in DMSO-$d_6$ **S200**
**Figure 197.** HMBC spectrum of compound **(2*S*,3*S*)-*t*-ES-a15** in DMSO-$d_6$ **S201**
**Figure 198.** $^1$H-$^1$H COSY spectrum of compound **(2*S*,3*S*)-*t*-ES-a15** in DMSO-$d_6$ **S201**
**Figure 199.** $^1$H NMR spectrum of compound **(2*S*,3*S*)-*t*-ES-a16** in DMSO-$d_6$ **S202**
**Figure 200.** $^{13}$C NMR spectrum of compound **(2*S*,3*S*)-*t*-ES-a16** in DMSO-$d_6$ **S202**
**Figure 201.** HSQC spectrum of compound **(2*S*,3*S*)-*t*-ES-a16** in DMSO-$d_6$ **S203**
**Figure 202.** HMBC spectrum of compound **(2*S*,3*S*)-*t*-ES-a16** in DMSO-$d_6$ **S203**
**Figure 203.** $^1$H-$^1$H COSY spectrum of compound **(2*S*,3*S*)-*t*-ES-a16** in DMSO-$d_6$ **S204**
**Figure 204.** $^1$H-$^1$H COSY spectrum of compound **(2*S*,3*S*)-*t*-ES-a16** in DMSO-$d_6$ **S204**
**Figure 205.** $^{13}$C NMR spectrum of compound **(2*S*,3*S*)-*t*-ES-a17** in DMSO-$d_6$ **S205**
**Figure 206.** HSQC spectrum of compound **(2*S*,3*S*)-*t*-ES-a17** in DMSO-$d_6$ **S205**
**Figure 207.** HMBC spectrum of compound **(2*S*,3*S*)-*t*-ES-a17** in DMSO-$d_6$ **S206**
**Figure 208.** $^1$H-$^1$H COSY spectrum of compound **(2*S*,3*S*)-*t*-ES-a17** in DMSO-$d_6$ **S206**
**Figure 209.** $^1$H NMR spectrum of compound **(2*S*,3*S*)-*t*-ES-a18** in DMSO-$d_6$ **S207**
**Figure 210.** $^{13}$C NMR spectrum of compound **(2*S*,3*S*)-*t*-ES-a18** in DMSO-$d_6$ **S207**
**Figure 211.** HSQC spectrum of compound **(2*S*,3*S*)-*t*-ES-a18** in DMSO-$d_6$ **S208**
**Figure 212.** HMBC spectrum of compound **(2*S*,3*S*)-*t*-ES-a18** in DMSO-$d_6$ **S208**
**Figure 213.** $^1$H-$^1$H COSY spectrum of compound **(2*S*,3*S*)-*t*-ES-a18** in DMSO-$d_6$ **S209**
**Figure 214.** $^1$H NMR spectrum of compound **(2*S*,3*S*)-*t*-ES-a19** in DMSO-$d_6$ **S209**
**Figure 215.** $^{13}$C NMR spectrum of compound **(2*S*,3*S*)-*t*-ES-a19** in DMSO-$d_6$ **S210**
**Figure 216.** HSQC spectrum of compound **(2*S*,3*S*)-*t*-ES-a19** in DMSO-$d_6$ **S210**
**Figure 217.** HMBC spectrum of compound **(2*S*,3*S*)-*t*-ES-a19** in DMSO-$d_6$ **S211**
**Figure 218.** $^1$H-$^1$H COSY spectrum of compound **(2*S*,3*S*)-*t*-ES-a19** in DMSO-$d_6$ **S211**
**Figure 219.** $^1$H NMR spectrum of compound **(2*S*,3*S*)-*t*-ES-a20** in DMSO-$d_6$ **S212**
**Figure 220.** $^{13}$C NMR spectrum of compound **(2*S*,3*S*)-*t*-ES-a20** in DMSO-$d_6$ **S212**
**Figure 221.** HSQC spectrum of compound **(2*S*,3*S*)-*t*-ES-a20** in DMSO-$d_6$ **S213**
**Figure 222.** HMBC spectrum of compound **(2*S*,3*S*)-*t*-ES-a20** in DMSO-$d_6$ **S213**
**Figure 223.** $^1$H-$^1$H COSY spectrum of compound **(2*S*,3*S*)-*t*-ES-a20** in DMSO-$d_6$ **S214**
**Figure 224.** $^1$H NMR spectrum of compound **(2*S*,3*S*)-*t*-ES-a21** in DMSO-$d_6$ **S214**
**Figure 225.** $^{13}$C NMR spectrum of compound **(2*S*,3*S*)-*t*-ES-a21** in DMSO-$d_6$ **S215**
**Figure 226.** HSQC spectrum of compound **(2*S*,3*S*)-*t*-ES-a21** in DMSO-$d_6$ **S215**
**Figure 227.** HMBC spectrum of compound **(2*S*,3*S*)-*t*-ES-a21** in DMSO-$d_6$ **S216**
**Figure 228.** $^1$H-$^1$H COSY spectrum of compound **(2*S*,3*S*)-*t*-ES-a21** in DMSO-$d_6$ **S216**

**Figure 229.** $^{1}$H NMR spectrum of compound **(2*S*,3*S*)-*t*-ES-a22** in DMSO-$d_6$ **S217**
**Figure 230.** $^{13}$C NMR spectrum of compound **(2*S*,3*S*)-*t*-ES-a22** in DMSO-$d_6$ **S217**
**Figure 231.** HSQC spectrum of compound **(2*S*,3*S*)-*t*-ES-a22** in DMSO-$d_6$ **S218**
**Figure 232.** HMBC spectrum of compound **(2*S*,3*S*)-*t*-ES-a22** in DMSO-$d_6$ **S218**
**Figure 233.** $^{1}$H-$^{1}$H COSY spectrum of compound **(2*S*,3*S*)-*t*-ES-a22** in DMSO-$d_6$ **S219**
**Figure 234.** $^{1}$H NMR spectrum of compound **(2*S*,3*S*)-*t*-ES-a23** in DMSO-$d_6$ **S219**
**Figure 235.** $^{13}$C NMR spectrum of compound **(2*S*,3*S*)-*t*-ES-a23** in DMSO-$d_6$ **S220**
**Figure 236.** HSQC spectrum of compound **(2*S*,3*S*)-*t*-ES-a23** in DMSO-$d_6$ **S220**
**Figure 237.** HMBC spectrum of compound **(2*S*,3*S*)-*t*-ES-a23** in DMSO-$d_6$ **S221**
**Figure 238.** $^{1}$H-$^{1}$H COSY spectrum of compound **(2*S*,3*S*)-*t*-ES-a23** in DMSO-$d_6$ **S221**
**Figure 239.** $^{1}$H NMR spectrum of compound **(2*S*,3*S*)-*t*-ES-a24** in DMSO-$d_6$ **S222**
**Figure 240.** $^{13}$C NMR spectrum of compound **(2*S*,3*S*)-*t*-ES-a24** in DMSO-$d_6$ **S222**
**Figure 241.** HSQC spectrum of compound **(2*S*,3*S*)-*t*-ES-a24** in DMSO-$d_6$ **S223**
**Figure 242.** HMBC spectrum of compound **(2*S*,3*S*)-*t*-ES-a24** in DMSO-$d_6$ **S223**
**Figure 243.** $^{1}$H-$^{1}$H COSY spectrum of compound **(2*S*,3*S*)-*t*-ES-a24** in DMSO-$d_6$ **S224**
**Figure 244.** $^{1}$H NMR spectrum of compound **(2*S*,3*S*)-*t*-ES-a25** in DMSO-$d_6$ **S224**
**Figure 245.** $^{13}$C NMR spectrum of compound **(2*S*,3*S*)-*t*-ES-a25** in DMSO-$d_6$ **S225**
**Figure 246.** HSQC spectrum of compound **(2*S*,3*S*)-*t*-ES-a25** in DMSO-$d_6$ **S225**
**Figure 247.** HMBC spectrum of compound **(2*S*,3*S*)-*t*-ES-a25** in DMSO-$d_6$ **S226**
**Figure 248.** $^{1}$H-$^{1}$H COSY spectrum of compound **(2*S*,3*S*)-*t*-ES-a25** in DMSO-$d_6$ **S226**
**Figure 249.** $^{1}$H NMR spectrum of compound **(2*S*,3*S*)-*t*-ES-a26** in DMSO-$d_6$ **S227**
**Figure 250.** $^{13}$C NMR spectrum of compound **(2*S*,3*S*)-*t*-ES-a26** in DMSO-$d_6$ **S227**
**Figure 251.** HSQC spectrum of compound **(2*S*,3*S*)-*t*-ES-a26** in DMSO-$d_6$ **S228**
**Figure 252.** HMBC spectrum of compound **(2*S*,3*S*)-*t*-ES-a26** in DMSO-$d_6$ **S228**
**Figure 253.** $^{1}$H-$^{1}$H COSY spectrum of compound **(2*S*,3*S*)-*t*-ES-a26** in DMSO-$d_6$ **S229**
**Figure 254.** $^{1}$H NMR spectrum of compound **(2*S*,3*S*)-*t*-ES-a27** in DMSO-$d_6$ **S229**
**Figure 255.** $^{13}$C NMR spectrum of compound **(2*S*,3*S*)-*t*-ES-a27** in DMSO-$d_6$ **S230**
**Figure 256.** HSQC spectrum of compound **(2*S*,3*S*)-*t*-ES-a27** in DMSO-$d_6$ **S230**
**Figure 257.** HMBC spectrum of compound **(2*S*,3*S*)-*t*-ES-a27** in DMSO-$d_6$ **S231**
**Figure 258.** $^{1}$H-$^{1}$H COSY spectrum of compound **(2*S*,3*S*)-*t*-ES-a27** in DMSO-$d_6$ **S231**
**Figure 259.** $^{1}$H NMR spectrum of compound **(2*S*,3*S*)-*t*-ES-a28** in DMSO-$d_6$ **S232**
**Figure 260.** $^{13}$C NMR spectrum of compound **(2*S*,3*S*)-*t*-ES-a28** in DMSO-$d_6$ **S232**
**Figure 261.** HSQC spectrum of compound **(2*S*,3*S*)-*t*-ES-a28** in DMSO-$d_6$ **S233**
**Figure 262.** HMBC spectrum of compound **(2*S*,3*S*)-*t*-ES-a28** in DMSO-$d_6$ **S233**
**Figure 263.** $^{1}$H-$^{1}$H COSY spectrum of compound **(2*S*,3*S*)-*t*-ES-a28** in DMSO-$d_6$ **S234**
**Figure 264.** $^{1}$H NMR spectrum of compound **(2*S*,3*S*)-*t*-ES-a29** in DMSO-$d_6$ **S234**
**Figure 265.** $^{13}$C NMR spectrum of compound **(2*S*,3*S*)-*t*-ES-a29** in DMSO-$d_6$ **S235**
**Figure 266.** HSQC spectrum of compound **(2*S*,3*S*)-*t*-ES-a29** in DMSO-$d_6$ **S235**
**Figure 267.** HMBC spectrum of compound **(2*S*,3*S*)-*t*-ES-a29** in DMSO-$d_6$ **S236**
**Figure 268.** $^{1}$H-$^{1}$H COSY spectrum of compound **(2*S*,3*S*)-*t*-ES-a29** in DMSO-$d_6$ **S236**
**Figure 269.** $^{1}$H NMR spectrum of compound **(2*S*,3*S*)-*t*-ES-a30** in DMSO-$d_6$ **S237**
**Figure 270.** $^{1}$H NMR spectrum of compound **(2*S*,3*S*)-*t*-ES-a30** in DMSO-$d_6$ **S237**
**Figure 271.** HSQC spectrum of compound **(2*S*,3*S*)-*t*-ES-a30** in DMSO-$d_6$ **S238**
**Figure 272.** HMBC spectrum of compound **(2*S*,3*S*)-*t*-ES-a30** in DMSO-$d_6$ **S238**
**Figure 273.** $^{1}$H-$^{1}$H COSY spectrum of compound **(2*S*,3*S*)-*t*-ES-a30** in DMSO-$d_6$ **S239**
**Figure 274.** $^{1}$H NMR spectrum of compound **(2*S*,3*S*)-*t*-ES-a31** in DMSO-$d_6$ **S239**
**Figure 275.** $^{13}$C NMR spectrum of compound **(2*S*,3*S*)-*t*-ES-a31** in DMSO-$d_6$ **S240**
**Figure 276.** HSQC spectrum of compound **(2*S*,3*S*)-*t*-ES-a31** in DMSO-$d_6$ **S240**
**Figure 277.** HMBC spectrum of compound **(2*S*,3*S*)-*t*-ES-a31** in DMSO-$d_6$ **S241**
**Figure 278.** $^{1}$H-$^{1}$H COSY spectrum of compound **(2*S*,3*S*)-*t*-ES-a31** in DMSO-$d_6$ **S241**
**Figure 279.** $^{1}$H NMR spectrum of compound **(2*S*,3*S*)-*t*-ES-a32** in DMSO-$d_6$ **S242**

**Figure 280.** $^{13}C$ NMR spectrum of compound **(2*S*,3*S*)-*t*-ES-a32** in DMSO-$d_6$ **S242**
**Figure 281.** HSQC spectrum of compound **(2*S*,3*S*)-*t*-ES-a32** in DMSO-$d_6$ **S243**
**Figure 282.** HMBC spectrum of compound **(2*S*,3*S*)-*t*-ES-a32** in DMSO-$d_6$ **S243**
**Figure 283.** $^{1}H$-$^{1}H$ COSY spectrum of compound **(2*S*,3*S*)-*t*-ES-a32** in DMSO-$d_6$ **S244**
**Figure 284.** $^{1}H$ NMR spectrum of compound **(2*S*,3*S*)-*t*-ES-a9-b7** in DMSO-$d_6$ **S244**
**Figure 285.** $^{13}C$ NMR spectrum of compound **(2*S*,3*S*)-*t*-ES-a9-b7** in DMSO-$d_6$ **S245**
**Figure 286.** HSQC spectrum of compound **(2*S*,3*S*)-*t*-ES-a9-b7** in DMSO-$d_6$ **S245**
**Figure 287.** HMBC spectrum of compound **(2*S*,3*S*)-*t*-ES-a9-b7** in DMSO-$d_6$ **S246**
**Figure 288.** $^{1}H$-$^{1}H$ COSY spectrum of compound **(2*S*,3*S*)-*t*-ES-a9-b7** in DMSO-$d_6$ **S246**
**Figure 289.** $^{1}H$ NMR spectrum of compound **(2*S*,3*S*)-*t*-ES-a9-b12** in DMSO-$d_6$ **S247**
**Figure 290.** $^{13}C$ NMR spectrum of compound **(2*S*,3*S*)-*t*-ES-a9-b12** in DMSO-$d_6$ **S247**
**Figure 291.** HSQC spectrum of compound **(2*S*,3*S*)-*t*-ES-a9-b12** in DMSO-$d_6$ **S248**
**Figure 292.** HMBC spectrum of compound **(2*S*,3*S*)-*t*-ES-a9-b12** in DMSO-$d_6$ **S248**
**Figure 293.** $^{1}H$-$^{1}H$ COSY spectrum of compound **(2*S*,3*S*)-*t*-ES-a9-b12** in DMSO-$d_6$ **S249**
**Figure 294.** $^{1}H$ NMR spectrum of compound **(2*S*,3*S*)-*t*-ES-a10-b9** in DMSO-$d_6$ **S249**
**Figure 295.** $^{13}C$ NMR spectrum of compound **(2*S*,3*S*)-*t*-ES-a10-b9** in DMSO-$d_6$ **S250**
**Figure 296.** HSQC spectrum of compound **(2*S*,3*S*)-*t*-ES-a10-b9** in DMSO-$d_6$ **S250**
**Figure 297.** HMBC spectrum of compound **(2*S*,3*S*)-*t*-ES-a10-b9** in DMSO-$d_6$ **S251**
**Figure 298.** $^{1}H$-$^{1}H$ COSY spectrum of compound **(2*S*,3*S*)-*t*-ES-a10-b9** in DMSO-$d_6$ **S251**
**Figure 299.** $^{1}H$ NMR spectrum of compound **(2*S*,3*S*)-*t*-ES-a10-b13** in acetone-$d_6$ **S252**
**Figure 300.** $^{13}C$ NMR spectrum of compound **(2*S*,3*S*)-*t*-ES-a10-b13** in acetone-$d_6$ **S252**
**Figure 301.** HSQC spectrum of compound **(2*S*,3*S*)-*t*-ES-a10-b13** in acetone-$d_6$ **S253**
**Figure 302.** HMBC spectrum of compound **(2*S*,3*S*)-*t*-ES-a10-b13** in acetone-$d_6$ **S253**
**Figure 303.** $^{1}H$-$^{1}H$ COSY spectrum of compound **(2*S*,3*S*)-*t*-ES-a10-b13** in acetone-$d_6$ **S254**
**Figure 304.** $^{1}H$ NMR spectrum of compound **(2*S*,3*S*)-*t*-ES-a10-b26** in acetone-$d_6$ **S254**
**Figure 305.** $^{13}C$ NMR spectrum of compound **(2*S*,3*S*)-*t*-ES-a10-b26** in acetone-$d_6$ **S255**
**Figure 306.** HSQC spectrum of compound **(2*S*,3*S*)-*t*-ES-a10-b26** in acetone-$d_6$ **S255**
**Figure 307.** HMBC spectrum of compound **(2*S*,3*S*)-*t*-ES-a10-b26** in acetone-$d_6$ **S256**
**Figure 308.** $^{1}H$-$^{1}H$ COSY spectrum of compound **(2*S*,3*S*)-*t*-ES-a10-b26** in acetone-$d_6$ **S256**
**Figure 309.** $^{1}H$ NMR spectrum of compound **(2*S*,3*S*)-*t*-ES-Leu-b43** in DMSO-$d_6$ **S257**
**Figure 310.** $^{13}C$ NMR spectrum of compound **(2*S*,3*S*)-*t*-ES-Leu-b43** in DMSO-$d_6$ **S257**
**Figure 311.** HSQC spectrum of compound **(2*S*,3*S*)-*t*-ES-Leu-b43** in DMSO-$d_6$ **S258**
**Figure 312.** HMBC spectrum of compound **(2*S*,3*S*)-*t*-ES-Leu-b43** in DMSO-$d_6$ **S258**
**Figure 313.** $^{1}H$-$^{1}H$ COSY spectrum of compound **(2*S*,3*S*)-*t*-ES-Leu-b43** in DMSO-$d_6$ **S259**
**Figure 314.** $^{1}H$ NMR spectrum of compound ***N*-succinyl-L-alanine** in DMSO-$d_6$ **S259**
**Figure 315.** $^{13}C$ NMR spectrum of compound ***N*-succinyl-L-alanine** in DMSO-$d_6$ **S260**
**Figure 316.** $^{1}H$ NMR spectrum of compound ***N*-succinyl-L-valine** in $CD_3OD$ **S260**
**Figure 317.** $^{13}C$ NMR spectrum of compound ***N*-succinyl-L-valine** in $CD_3OD$ **S261**
**Figure 318.** $^{1}H$ NMR spectrum of compound ***N*-succinyl-L-leucine** in $D_2O$ **S261**
**Figure 319.** $^{13}C$ NMR spectrum of compound ***N*-succinyl-L-leucine** in $D_2O$ **S262**
**Figure 320.** $^{1}H$ NMR spectrum of compound ***N*-succinyl-L-isoleucine** in $CD_3OD$ **S262**
**Figure 321.** $^{13}C$ NMR spectrum of compound ***N*-succinyl-L-isoleucine** in $CD_3OD$ **S263**
**Figure 322.** $^{1}H$ NMR spectrum of compound ***N*-succinyl-L-methionine** in DMSO-$d_6$ **S263**
**Figure 323.** $^{13}C$ NMR spectrum of compound ***N*-succinyl-L-methionine** in DMSO-$d_6$ **S264**
**Figure 324.** $^{1}H$ NMR spectrum of compound ***N*-succinyl-L-proline** in DMSO-$d_6$ **S264**
**Figure 325.** $^{1}H$ NMR spectrum of compound ***N*-succinyl-L-phenylalanine** in $CD_3OD$ **S265**
**Figure 326.** $^{13}C$ NMR spectrum of compound ***N*-succinyl-L-phenylalanine** in $CD_3OD$ **S265**
**Figure 327.** $^{1}H$ NMR spectrum of compound ***N*-succinyl-L-tyrosine** in DMSO-$d_6$ **S266**
**Figure 328.** $^{13}C$ NMR spectrum of compound ***N*-succinyl-L-tyrosine** in DMSO-$d_6$ **S266**
**Figure 329.** $^{1}H$ NMR spectrum of compound ***N*-succinyl-L-tryptophan** in DMSO-$d_6$ **S267**
**Figure 330.** $^{13}C$ NMR spectrum of compound ***N*-succinyl-L-tryptophan** in DMSO-$d_6$ **S267**

#### Complemented Experimental procedures

##### 1. Preparation of protoplast of *A. nidulans* and transformation

The transformation of *Aspergillus nidulans* A1145 ΔEMΔST[1], spores were inoculated into 50 mL liquid CD media in a 125-mL flask and germinated at 30 °C shaking at 250 rpm for ~9 h. The 20 X Nitrate salts solution was prepared by dissolving 120 g $NaNO_3$, 10.4 g KCl, 10.4 g $MgSO_4$•$7H_2O$, and 30.4 g $KH_2PO_4$ in 1 L double distilled water. The trace elements solution (100 mL) contained 2.20 g $ZnSO_4$•$7H_2O$, 1.10 g $H_3BO_3$, 0.50 g $MnCl_2$•$4H_2O$, 0.16 g $FeSO_4$•$7H_2O$, 0.16 g $CoCl_2$•$5H_2O$, 0.16 g $CuSO_4$•$5H_2O$, and 0.11 g $(NH_4)_6Mo_7O_{24}$•$4H_2O$. The dropout components for selection for the three expression vectors were uracil/uridine, pyridoxine and riboflavin. *A. nidulans* A1145 ΔEM was initially grown on CD agar plates containing 10 mM uridine, 5 mM uracil, 0.5 µg/mL pyridoxine HCl and 2.5 µg/mL riboflavin at 37°C for 5 days. The germinated spores were harvested by centrifugation at 3,500 rpm for 10 min, and washed with Osmotic buffer (10 mL, 1.2 M $MgSO_4$, 10 mM sodium phosphate buffer, pH 5.8). The mycelia were then mixed with Osmotic buffer (10 mL, 30 mg lysing enzymes from *Trichoderma*, 20 mg Yatalase) in a 125-mL flask. Protoplasts were prepared by incubating the mixture overnight at 30 °C with gentle shaking at 80 rpm. Cells were collected in a 30-mL Corex tube and overlaid gently by 10 mL of Trapping buffer (0.6 M sorbitol, 0.1 M Tris HCI, pH 7.0). Centrifugation at 3,500 rpm for 15 min at 4 °C layered the protoplasts at the interface of the two buffers. The protoplasts were then pipetted to a sterile 15-mL falcon tube and washed with STC buffer (10 mL, 1.2 M sorbitol, 10 mM $CaCl_2$, 10 mM Tris-HCI pH 7.5). The protoplasts were resuspended in STC buffer (1 mL).

For each transformation, 3 µL of each plasmid (>100 ng/µL) was added to 60 µL of the *A. nidulans* A1145 ΔSTΔEM protoplast suspension prepared as above, and the mixture was incubated for 1 h on ice. 600 µL PEG solution (60% PEG, 50 mM of $CaCl_2$, and 50 mM of Tris-HCl, pH 7.5) was added to the protoplast mixture, followed by additional incubation at room temperature for 20 min. The mixture was spread on the CD sorbitol plate (CD solid medium with 1.2 M sorbitol and the appropriate supplements: 10 mM of uridine, 5 mM of uracil, 0.5 µg/mL of pyridoxine HCl, and/or 2.5 µg/mL of riboflavin according to the markers in the transformed plasmids) and incubated at 37 °C for 3-4 days.

##### 2. Spectroscopic data of *N*-succinyl proteinogenic amino acids

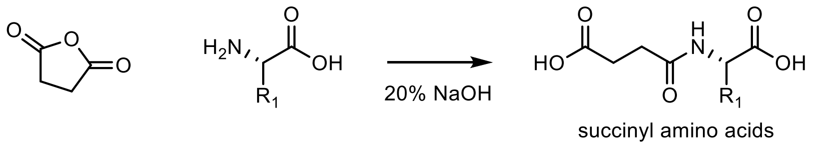

Spectroscopic data of *N*-succinyl-L-alanine matched literature spectra[2].

$^1$H NMR (500 MHz, DMSO) $\delta$ 8.15 ppm (1H, t, $J$ = 7.0 Hz), 4.17 ppm (1H, m), 2.37 ppm (4H, m), 1.24 ppm (3H, d, $J$ = 7.3 Hz). $^{13}$C NMR (125 MHz, DMSO) $\delta$ 174.3, 173.8, 170.8, 47.5, 29.8, 29.1, 17.3. HRMS (ESI, M+H$^+$) calculated for $C_7H_{12}NO_5^+$ 190.0710; found 190.0700.

Spectroscopic data of *N*-succinyl-L-valine agree with the reported literature value[3].

$^1$H NMR (500 MHz, $CD_3OD$): $\delta$ 4.35 ppm (1H, dd, $J$ = 2.9, 5.6 Hz), 2.59 ppm (4H, m), 2.15 ppm (1H, m), 0.95 ppm (6H, m). $^{13}$C NMR (125 MHz, $CD_3OD$) $\delta$ 176.3, 174.9, 174.7, 58.8, 31.6, 31.2, 30.2, 19.5, 18.2. HRMS (ESI, M+H$^+$) calculated for $C_9H_{16}NO_5^+$ 218.1023; found 218.1021.

Spectroscopic data of *N*-succinyl-L-leucine agree with the reported literature value[4].

$^1$H NMR (500 MHz, $D_2O$): $\delta$ 4.36 ppm (1H, dd, $J$ = 5.6, 9.1Hz), 2.61 ppm (2H, m), 2.55 ppm (2H, m), 1.61 ppm (3H, m), 0.88 ppm (3H, d, $J$ = 6.5 Hz), 0.84 ppm (3H, d, $J$ = 6.5 Hz). $^{13}$C NMR (125 MHz, $CD_3OD$) $\delta$ 178.1, 177.9, 176.0, 53.0, 41.5, 32.1, 31.8, 26.3, 24.2, 22.7. HRMS (ESI, M+H$^+$) calculated for $C_{10}H_{18}NO_5^+$ 232.1179; found 232.1173.

Spectroscopic data of *N*-succinyl-L-isoleucine agree with the reported literature value[2].

$^1$H NMR (500 MHz, $CD_3OD$): $\delta$ 4.37 (1H, t, $J$ = 5.4 Hz), 2.59 (4H, m), 1.88 (1H, m), 1.52 (1H, m), 1.25 (1H, m), 0.93 (6H, m). $^{13}$C NMR (125 MHz, $CD_3OD$) $\delta$ 176.3, 174.9, 174.6, 58.1, 38.4, 31.3, 30.3, 26.2, 16.0, 11.8. HRMS (ESI, M+H$^+$) calculated for $C_{10}H_{18}NO_5^+$ 232.1179; found 218.1180.

Spectroscopic data of *N*-succinyl-L-methionine agree with the reported literature value[2].

$^1$H NMR (500 MHz, DMSO): $\delta$ 8.12 ppm (1H, d, $J$ = 7.9), 4.37 ppm (1H, m), 2.47 ppm (6H, m), 2.05 ppm (3H, s), 1.97 ppm (1H, m), 1.87 ppm (1H, m). $^{13}$C NMR (125 MHz, DMSO) $\delta$ 174.5, 174.1, 172.2, 51.6, 31.5, 30.4, 30.3, 29.6, 15.1.

$^1$H NMR data of *N*-succinyl-L-proline agree with the reported literature value[2].

$^1$H NMR (500 MHz, DMSO): $^1$H NMR (500 MHz, DMSO) $\delta$ 4.48 ppm (0.5H, dd, $J$ = 8.5 Hz), 4.20 ppm (1H, dd, $J$ = 3.6, 8.9 Hz), 3.51 ppm (2H, m), 2.42 ppm (4H, m), 2.10 ppm (1H, m), 1.90-1.75 ppm (3H, m). HRMS (ESI, M+H$^+$) calculated for $C_9H_{14}NO_5^+$ 216.0866; found 216.0854.

Spectroscopic data of *N*-succinyl-L-serine agree with the reported literature value[2].

$^1$H NMR (500 MHz, DMSO): $\delta$ 8.00 ppm (1H, d, $J$ = 8.0 Hz), 4.24 ppm (1H, m), 3.60 ppm (3H, m), 2.38 ppm (4H, m). $^{13}$C NMR (125 MHz, DMSO) $\delta$ 173.9, 172.2, 171.2, 61.5, 54.7, 29.8, 29.1. HRMS (ESI, M+H$^+$) calculated for $C_7H_{12}NO_6^+$ 206.0659; found 206.0665.

Spectroscopic data of *N*-succinyl-L-theronine agree with the reported literature value[2].

$^1$H NMR (500 MHz, DMSO): $\delta$ 7.77 ppm (1H, m), 4.19 ppm (1H, dd, $J$ = 3.3, 8.8 Hz), 4.07 ppm (1H, m, $J$ = 3.3, 6.4 Hz), 2.48 ppm (4H, m), 1.02 (3H, d, $J$ = 6.4 Hz). $^{13}$C NMR (125 MHz, DMSO) $\delta$ 173.8, 172.2, 171.5, 66.4, 57.6, 29.8, 29.2, 20.3. HRMS (ESI, M+H$^+$) calculated for $C_8H_{14}NO_6^+$ 220.0816; found 220.0811.

Spectroscopic data of *N*-succinyl-L-cysteine agree with the reported literature value[2].

H NMR (500 MHz, DMSO): $\delta$ 8.21 ppm (1H, d, $J$ = 7.5 Hz), 4.36 ppm (1H, dt, $J$ = 6.3, 13.3 Hz), 2.90 ppm (1H, d, $J$ = 5.7, 13.1 Hz), 2.78 ppm (1H, $J$ = 7.9, 13.1 Hz), 2.37 ppm (4H, m). HRMS (ESI, M+H$^+$) calculated for $C_7H_{12}SNO_5^+$ 222.0431; found 222.0425.

Spectroscopic data of *N*-succinyl-L-asparagine agree with the reported literature value[2].

H NMR (500 MHz, DMSO): 8.06 ppm (1H, d, $J$ = 7.9 Hz), 7.31 ppm (1H, s), 6.86 ppm (1H, s), 4.46 ppm (1H, m), 2.48 ppm (6H, m). HRMS (ESI, M+H$^+$) calculated for $C_8H_{13}N_2O_6^+$ 233.0768; found 233.0752.

Spectroscopic data of *N*-succinyl-L-glutamine agree with the reported literature value[2].

$^1$H NMR (500 MHz, DMSO): $\delta$ 8.14 ppm (1H, d, $J$ = 7.8 Hz), 7.26 ppm (1H, s), 6.77 ppm (1H, s), 4.14 ppm (1H, m), 2.39 ppm (4H, m), 2.10 ppm (2H, m), 1.92 ppm (1H, m), 1.73 ppm (1H, m). $^{13}$C NMR (125 MHz, DMSO) $\delta$ 173.8,

173.5, 173.5, 171.2, 51.6, 30.8, 29.8, 29.1, 27.0. HRMS (ESI, M+H$^+$) calculated for $C_9H_{15}N_2O_6^+$ 247.0925; found 247.0925.

Spectroscopic data of *N*-succinyl-L-phenylalanine agree with the reported literature value[3].

$^1$H NMR (500 MHz, $CD_3OD$): $\delta$ 7.23 ppm (5H, m), 4.66 ppm (1H, dd, *J* = 5.2, 8.5 Hz), 3.18 ppm (1H, dd, *J* = 5.3, 13.9 Hz), 2.96 ppm (1H, dd, *J* = 8.6, 13.9), 2.48 ppm (4H, m). $^{13}$C NMR (125 MHz, $CD_3OD$) $\delta$ 176.1, 174.6, 174.3, 138.3, 130.3, 129.4, 127.8, 55.0, 38.4, 31.3, 30.2. HRMS (ESI, M+H$^+$) calculated for $C_9H_{14}NO_5^+$ 216.0866; found 216.0854.

Spectroscopic data of *N*-succinyl-L-tyrosine agree with the reported literature value[2].

$^1$H NMR (500 MHz, DMSO): 8.10 ppm (1H, d, *J* = 8.0 Hz), 6.98 ppm (2H, m), 6.64 (2H, m), 4.32 ppm (1H, dd, *J* = 4.4, 8.8 Hz), 2.89 ppm (1H, dt, *J* = 3.7, 8.9 Hz), 2.73 ppm (1H, dt, *J* = 5.3, 8.8 Hz), 2.34 ppm (4H, m). $^{13}$C NMR (125 MHz, $CD_3OD$) $\delta$ 174.2, 173.6, 171.4, 156.3, 130.5, 128.1, 115.4, 54.3, 36.6, 30.3, 29.5. HRMS (ESI, M+H$^+$) calculated for $C_{13}H_{16}NO_6^+$ 282.0972; found 282.0967.

Spectroscopic data of *N*-succinyl-L-tryptophan agree with the reported literature value[2].

$^1$H NMR (500 MHz, DMSO): 8.16 ppm (1H, d, *J* = 7.8 Hz), 7.55 ppm (1H, t, *J* = 7.3 Hz), 7.35 (1H, dd, *J* = 5.6, 8.1Hz), 7.17 ppm (1H, m), 7.07 ppm (1H, m), 6.99 ppm (1H, m), 4.52 ppm (1H, m), 3.19 ppm (1H, m), 3.05 ppm (1H, m), 2.39 ppm (4H, m). $^{13}$C NMR (125 MHz, $CD_3OD$) $\delta$ 174.2, 173.8, 171.5, 136.4, 127.6, 123.9, 121.2, 118.7, 118.5, 111.7, 110.2, 53.4, 30.2, 29.3, 27.5. HRMS (ESI, M+H$^+$) calculated for $C_{15}H_{17}N_2O_5^+$ 305.1132; found 305.1130.

Spectroscopic data of *N*-succinyl-L-aspartate agree with the reported literature value[2].

$^1$H NMR (500 MHz, DMSO): 8.19 ppm (1H, d, *J* = 8.0 Hz), 4.49 ppm (1H, m), 2.64 ppm (1H, m), 2.52 ppm (1H, m), 2.36 ppm (4H, m). HRMS (ESI, M+H$^+$) calculated for $C_8H_{12}NO_7^+$ 234.0608; found 234.0600.

Spectroscopic data of *N*-succinyl-L-glutamic acid agree with the reported literature value[3].

$^1$H NMR (500 MHz, $D_2O$): 4.40 ppm (1H, dd, *J* = 5.1, 9.3), 2.58 ppm (4H, m), 2.47 ppm (2H, m), 2.18 ppm (1H, m), 1.98 ppm (1H, m). $^{13}$C NMR (125 MHz, $D_2O$) $\delta$ 180.3, 180.1, 178.3, 178.2, 55.2, 33.2, 32.3, 32.0, 29.0. HRMS (ESI, M+H$^+$) calculated for $C_9H_{14}NO_7^+$ 248.0765; found 248.0760.

Spectroscopic data of *N*-succinyl-L-histidine agree with the reported literature value[2].

$^1$H NMR (500 MHz, $D_2O$): 8.47 ppm (1H, d, *J* = 5.7), 7.17 ppm (1H, d, *J* = 5.7), 4.65 ppm (1H, m), 3.22 ppm (1H, m), 3.05 ppm (1H, m), 2.48 ppm (4H, m). HRMS (ESI, M+H$^+$) calculated for $C_{10}H_{14}N_3O_5^+$ 256.0928; found 256.0927.

Spectroscopic data of *N*-succinyl-L-lysine agree with the reported literature value[2].

$^1$H NMR (500 MHz, $D_2O$): 4.25 ppm (1H, m), 2.86 ppm (1H, m), 2.52 ppm (4H, m), 1.79 ppm (1H, m), 1.65 ppm (1H, m), 1.55 ppm (2H, m), 1.34 ppm (2H, m). $^{13}$C NMR (125 MHz, $D_2O$) $\delta$ 176.8, 175.7, 174.9, 52.3, 39.1, 29.9, 29.8, 29.0, 26.1, 21.9. HRMS (ESI, M+H$^+$) calculated for $C_{10}H_{19}N_2O_5^+$ 247.1288; found 247.1290.

Spectroscopic data of *N*-succinyl-L-arginine agree with the reported literature value[2].

$^1$H NMR (500 MHz, $D_2O$): 4.26 ppm (1H, m), 3.10 ppm (2H, t, *J* = 6.9 Hz), 2.55 ppm (4H, m), 1.81 ppm (1H, m), 1.65 ppm (1H, m), 1.54 ppm (2H, m). $^{13}$C NMR (125 MHz, $D_2O$) $\delta$ 176.9, 175.6, 174.8, 156.7, 52.4, 30.2, 29.9, 29.0, 27.7, 24.2. HRMS (ESI, M+H$^+$) calculated for $C_{10}H_{19}N_4O_5^+$ 275.1350; found 275.1347.

#### 3. Preparation of (2*S*,3*S*)-*t*-ES or (2*R*,3*R*)-*t*-ES standard

(2*S*,3*S*)-*t*-ES or (2*R*,3*R*)-*t*-ES was obtained by the hydrolysis from corresponding (2*S*,3*S*)-diethyl-2,3-epoxysuccinate and (2*R*,3*R*)-diethyl-2,3-epoxysuccinate, respectively. A solution of aqueous NaOH (40 mg, 2 eq.)

was added to (2*S*,3*S*)-diethyl-2,3-epoxysuccinate (94 mg, 1 eq.) in an ice bath. The resulting solution was stirred for 2 h at 0°C, then for 30 min at room temperature, after which the solution was neutralized and lyophilized to yield (2*S*,3*S*)-*t*-ES. $^{1}$H NMR (400 MHz, $D_2O$): 3.39 (s, 2H); $^{13}$C NMR (100 MHz, $D_2O$): 175.1, 53.9. Similarly (2*R*,3*R*)-diethyl-2,3-epoxysuccinate was treated as described above to yield (2*R*,3*R*)-*t*-ES. $^{1}$H NMR (400 MHz, $D_2O$): 3.45 (s, 2H); $^{13}$C NMR (100 MHz, $D_2O$): 174.6, 53.9.

**4. Sequence information**

Cp1A amino acid sequence

MQQFVRNVNPARIGDITTQVSRVVPHLIASDLSCATRSSHVTEVSNALRKSGILKVSLQFKDDASKYLQNL
ILGLHKHHGHGLPITHSASRGWFWDIRPNSTTFQTPSHQARSETMQEFPWHTDCSYEEAPPKYFALQVLR
EDRCGGGTLSVMNVGKLSSMLSPSTCAALLRPQFRIDVPPEFVKNDASRHIIGSLMAADSSGAPNMLRFR
EDIMTPLNVEAAAALVELKDRLLGLEVQAETLHLTPDCLPRGSVVLMDNRRWLHARNEVMDPERHLRR
VRWDARPFPAMTM

Cp1B amino acid sequence

MKIPAPQQLQQLHVSLDGGHYEPVTTFDPAKATYLQDQEALQENLLRLCSVNGWHKSSRAACSPRPVLV
SSEHQRRWRELHEALVLAITDIVERWLTDPEARFPERMPLEPEEEDLLRWIDEQVPHNLPQYRDCRGSWR
PDFLVEEENSEDGSGPVENFRISEINARFSFNGFMFATCGQQAIHDMGICDNGNGLVGATDPAKILKGLLR
LFQPGLPLHLLKGDEAGVDIHMLVDFLDRYLGITPRFIMPADLRLLHEPQAKGGYKLCCVVKNPDSCDPA
TLIYHDGDILEEIHQVGLELHQREIRALEPEMLRQISLRCFNDMRTILLVHDKRMLGIVRQELENLVARNV
LTLSQAKILDKGIPETILPGSLDLDQAIARCKEMPELKDEYILKPIRSGKGDGIVFGEDLNSEEWISRLEGLR
SAQLIPGGGTCIVQRKVKQLLYDVVLRPTGVKTRYPLIGTYHSINGEFLGVGVWRSSPDRICAISHGGAWT
VSVMRDE

Cp1D amino acid sequence

MAPTTFSLKEVLAVAEIHPFYNPAVEYPPTPETIKSAIELADKRSTDIDLSSLPLVSKKDLYKAIARLTDDTSP
QNEYRRSSYVSITGGGSGGLPLMFVTDTKENRNQRAVFGEFLSTCGVVEPHDWILTTHTSGYFYRSLDLL
SEILENAGATVLSAGNYMTPAEVVHALAHYHVNVITGDGSQVVQVVHHISTLPAEEKAKIKLTKVLYTSE
PLTETQQIHIRATLGPVKICSVWGSAEAGPCALSDPDLTSPERPPGTMDFIFDTRQVVIEILPHSASEGDSSA
GVKSVPDGEEGIIVQTSLVRRRNPLVRYITGDVGSLQPLPEKARAIIPESELEHLRVLRLRGRDRRFSFKWF
GIYFEFENIVSFMQGDKTGVLQWQVILATLESSPQTKLEIRLLRQANNEHIMTKEELLNKLEKYFFILPENE
HLFQVTFLDDLSGFEKSSTGNKVMKFVDKVH

Cp2B amino acid sequence

MKYPTTGQLQQVHLGIGPKGYEPVASYQGDKQLYTQEHEILQASILGFCPEHLWHHGSNKASCPRPILVT
AKHQEQLEQLHNALVTAIVDIVKRWWTDLDARFPERMPLTRDEEDLLRWLEHQHSHNGVPYEARLGSW
RPDFLVGDYSGGPSTETYRLTEINARFCFNGFMHQAYGQEGLSDLGAGRNGLIHATDSSKILDGLLSLFNP
DRPLHLLKGEEPGIDIHMFIDFVYRHIGIKPRLITPADLRLIPDPQKKDGSKLCCLVKDQQNASLINESRLLV
TSKGEVVEEVHQVGLELHQHELFGLSREMLREISLRCFNDMRTILLVHDKRMLGIIKQEMPTLVARKVLT

HDQGEALERGISDSFIPGSSELNELIQTLTDSPELRKEYLLKPIRGGKGAGIIFGDEVGPDEWLSTLERLRNP
HFVSGNTMYVVQRRIWPRLYEVILNSSGDRGNYPLIGTYHTTNGQLLGLGTWRSSPDRICAVSHGGGWIC
SVLDEYAESSE

Cp2D amino acid sequence
MTTKSFSLSEVLAVAKRHPFYNPEIQYPLDETALQAVRDWAVKNQTEVDLRFQPLLHKNDIYKTVERLTH
DASPENVYRESSYMSITGGGSGGVPMMFAVDVHENRQQRAQMGKLLRNCGVIRRKDWVLSVHISGGFY
RSLDLTTETMENAGATVLSAGNYMEPEEVVQALAHYHVNVLTGDASQIVQLACYISTLPLERQRQIQINK
IIYTSEPLTGAQRAFLRATLGDVKICSVMGSSEAGPWALSNPDLVGEENLNSSSMDFVFDTRDMIIEILSPA
GLDDGKPPSDIDPLPLGETGIIVQTSLRRLRNPLVRYITGDLGSLHPLPEIASAVVPESERQYLRVLRMQGR
DRRFSFKWYGAYFEFEKMKALLQAEECGVLQWQVILDQLESSGLPTLQVRLLRAPSRADVLSEEQLVKR
VRTFFLVLPENEDVFSIVFVKNLDGFERSSTAGKVISFVDRLH

Epoxysuccinate synthase MfaA from *Microcoleus sp. FACHB-1* amino acid sequence
MIAAKKTDLLAIEENPLILPASCLFQIDTKNDIDFDAYASALFEAGIILLDLGFDNPDASIMTTIVEHLGTIDT
HDGKGMVIWDVKYDANVDQDKGTRSLTTKKFPIHTDASFEEPPPQYVALYVVAEDSLGGGITQLIDGRQI
LQHLSREAISVLQTKAFKFRVPQEFIKNKAYIEASILNGEGNFRYRQEVLILDDCTPQELQAIGELELLLAN
KSLIKSIFLKTGTIIIFDNGRFLHGRTKVRDKNRHLKRLRFQAKQTRFGVDCETYVYRKESGCLG

Polyphosphate kinase (CHU)
MATDFSKLSKYVETLRVKPKQSIDLKKDFDTDYDHKMLTKEEGEELLNLGISKLSEIQEKLYASGTKSVLI
VFQAMDAAGKDGTVKHIMTGLNPQGVKVTSFKVPSKIELSHDYLWRHYVALPATGEIGIFNRSHYENVL
VTRVHPEYLLSEQTSGVTAIEQVNQKFWDKRFQQINNFEQHISENGTIVLKFFLHVSKKEQKKRFIERIELD
TKNWKFSTGDLKERAHWKDYRNAYEDMLANTSTKQAPWFVIPADDKWFTRLLIAEIICTELEKLNLTFP
TVSLEQKAELEKAKAELVAEKSSD

**Supplementary Tables**

**Supplementary Table 1.** Comparative BLASTP analysis of the *cp1* gene cluster with homologous cluster from ascomycetes *Aspergillus flavus*, *Trichoderma harzianum*, and cyanobacterium *Microcoleus sp. FACHB-1*.

| *Aspergillus flavus* *cp1* | *Aspergillus flavus* (% identity / % similarity) *cp2* | *Trichoderma harzianum* (% identity / % similarity) *tcp* | *Microcoleus sp. FACHB-1* (% identity / % similarity) *mfa* | Homolog in Swiss-Prot database (% identity) |
|---|---|---|---|---|
| Cp1A XP_041142152.1 | Cp2A (61/76) XP_041145391.1 | TcpA (61/76) KKO98516.1 | MfaA (30/52) MBD2130326.1 | L-asparagine oxygenase (29%) Q9Z4Z5.1 |
| Cp1B XP_041142153.1 | Cp2B (53 / 69) XP_041145392.1 | TcpB (51/66) KKO98515.1 | MfaB (25/41) MBD2130328.1 | No hits |
| Cp1C XP_041142154.1 | Cp2C (58/74) XP_041145390.1 | *Not conserved* | *Not conserved* | FlvG (42%) PLP-dependent decarboxylase B8NHE2.1 |
| Cp1D XP_041142155.1 | Cp2D (57/74) XP_041145389.1 | TcpD (52/70) KKO98517.1 | MfaD (24/43) MBD2130327.1 | No hits |

**Supplementary Table 2.** Primers used in this study

| Primers | Sequence (5'-3') |
|---|---|
| pML 8001 F1 | CTGAGCTTCATCCCCAGCATCATTACACCTCAGCAATGCAACAATTCGTACGCAATGTC |
| pML 8001 R1 | AGAATCAGTAAGCTCACATGTATTCCTGGAGCAAACTTCCTTGTGGACGAAATGCAGC |
| pML 8001 F2 | TTTGCTCCAGGAATACATGTGAGCTTAC |
| pML 8001 R2 | TTTTGCGATTGTTTGAAGTGTTCTGTATGC |
| pML 8001 F3 | CAAGTGCATACAGAACACTTCAAACAATCGCAAAAATGAAGATTCCAGCTCCCCAAC |
| pML 8001 R3 | ACACAGTGGAGGACATACCCGTAATTTTCTGATTTAAATGTGGTTGCCACTTAGTCAGGG |
| pML 8002 F1 | CCTTCTCTGAACAATAAACCCCACAGAAGGCATTTATGATGGTGACGGACATGGC |
| pML 8002 R1 | TCAGTAAGCTCACATGTATTCCTGGAGCAAAGCCGTGAACCCTAATATAAACATCAAAGC |
| pML 8002 F2 | TTTGCTCCAGGAATACATGTGAGCTTAC |
| pML 8002 R2 | TTTTGCGATTGTTTGAAGTGTTCTGTATGC |
| pML 8002 F3 | CAAGTGCATACAGAACACTTCAAACAATCGCAAAAATGGCTCCTACCACATTTTCCC |
| pML 8002 R3 | TGATGAGACCCAACAACCATGATACCAGGGGATTTAAATGTTCTTCTGCCGTTGTCGACG |
| pML 8003 F1 | CTCTCCCTTCTCTGAACAATAAACCCCACAGAAGGCATTTATGATGGTGACGGACATGGC |
| pML 8003 R1 | ATAGGTCGCCAGGTACGACCAGTTCGGAAGATCAGGAAATGTTCTTCTGCCGTTGTCGAC |
| pML 8003 F2 | CGACATACGCTGCGTCGACAACGGCAGAAGAACATTTCCTGATCTTCCGAACTGGTCG |
| pML 8003 R2 | TGCTGAGGTGTAATGATGCTGGGGATGA |
| pML 8003 F3 | CTGAGCTTCATCCCCAGCATCATTACACCTCAGCAATGCAACAATTCGTACGCAATGTC |
| pML 8003 R3 | AGTGATGAGACCCAACAACCATGATACCAGGGGATTTCTTCCTTGTGGACGAAATGCAGC |
| pML 8004 F1 | CTCCCTTCTCTGAACAATAAACCCCACAGAAGGCATTTATGATGGTGACGGACATGGC |
| pML 8004 R1 | GGTCTCTCCCGTCACCCAAATCAATTCACCGGAGTAAATGTTCTTCTGCCGTTGTCGAC |
| pML 8004 F2 | ACGCTGCGTCGACAACGGCAGAAGAACATTTACTCCGGTGAATTGATTTGGGTG |
| pML 8004 R2 | GTTGTTGGAGTTGTTGGGGAGCTGGAATCTTCATTGTTTAGATGTGTCTATGTGGCGGG |
| pML 8004 F3 | CTAACCATTACCCCGCCACATAGACACATCTAAACAATGAAGATTCCAGCTCCCCAAC |
| pML 8004 R3 | GGAGTGATGAGACCCAACAACCATGATACCAGGGGATTTGTGGTTGCCACTTAGTCAGGG |
| pML 8005 F1 | GAGCTTCATCCCCAGCATCATTACACCTCAGCAATGCAACAATTCGTACGCAATGTC |
| pML 8005 R1 | TAGCTCGTTCGGCACCTTTAATCAAATGTGGTTGCCACTTAGTCAGGGCT |
| pML 8005 F2 | ATGGGCTTAAAGCCCTGACTAAGTGGCAACCACATTTGATTAAAGGTGCCGAACGAGCTA |
| pML 8005 R2 | CACAGTGGAGGACATACCCGTAATTTTCTGATTTCGGAATGTCTCGGATTAATATCACCG |
| pML 8006 F1 | GCTTCATCCCCAGCATCATTACACCTCAGCAATGCAACAATTCGTACGCAATGTC |
| pML 8006 R1 | GGGTCTCTCCCGTCACCCAAATCAATTCACCGGAGTAAATGTGGTTGCCACTTAGTCAGG |
| pML 8006 F2 | ATGGGCTTAAAGCCCTGACTAAGTGGCAACCACATTTACTCCGGTGAATTGATTTGGGTG |
| pML 8006 R2 | GAACCTCCTTGAGGGAAAATGTGGTAGGAGCCATTGTTTAGATGTGTCTATGTGGCGGG |
| pML 8006 F3 | CTAACCATTACCCCGCCACATAGACACATCTAAACAATGGCTCCTACCACATTTTCCC |
| pML 8006 R3 | TCAACACAGTGGAGGACATACCCGTAATTTTCTGATTTAGTTCTTCTGCCGTTGTCGACG |
| pML 8007 F1 | TTCTCTGAACAATAAACCCCACAGAAGGCATTTATGCAACATATAGGACGTCATTTCCAC |
| pML 8007 R1 | AATCAGTAAGCTCACATGTATTCCTGGAGCAAACGGTTCTTGCGATTTGTTTTCTTTTCC |
| pML 8007 F2 | TTTGCTCCAGGAATACATGTGAGCTTAC |
| pML 8007 R2 | TTTTGCGATTGTTTGAAGTGTTCTGTATGC |
| pML 8007 F3 | ACAAGTGCATACAGAACACTTCAAACAATCGCAAAAATGAAGTACCCTACTACCGGACAG |
| pML 8007 R3 | GATGAGACCCAACAACCATGATACCAGGGGATTTAAATCTAGCTGCAGCTGAAGTACCAG |
| pML 8008 F1 | CTGAGCTTCATCCCCAGCATCATTACACCTCAGCAATGGGAAGTGTGGGCATCTCT |
| pML 8008 R1 | TAGAATCAGTAAGCTCACATGTATTCCTGGAGCAAATACTCTCTGCCAAGAGCAATAGGC |
| pML 8008 F2 | TTTTCCTCGCCTATTGCTCTTGGCAGAGAGTATTTGCTCCAGGAATACATGTGAGCTTAC |
| pML 8008 R2 | CCTCCGACAACGAGAAGCTTTTCGTGGTCATTTTTGCGATTGTTTGAAGTGTTCTGTATG |
| pML 8008 F3 | TACAAGTGCATACAGAACACTTCAAACAATCGCAAAAATGACCACGAAAAGCTTCTCGT |
| pML 8008 R3 | CAGTGGAGGACATACCCGTAATTTTCTGATTTAAATGAACACCAGGAATCCAGCATTAGC |
| pML 8009 F1 | CATCATCACAGCAGCGGCCTGGTGCCGCGCGGCAGCATGCAACAATTCGTACGCAATGTC |
| pML 8009 R1 | GGATCTCAGTGGTGGTGGTGGTGGTGCTCGAGTCACTACATAGTCATGGCAGGAAATGGC |
| pML 8010 F1 | ATCATCATCACAGCAGCGGCCTGGTGCCGCGCGGCAGCATGAAGATTCCAGCTCCCCAAC |
| pML 8010 R1 | GATCTCAGTGGTGGTGGTGGTGGTGCTCGAGTCACTATTCATCACGCATAACAGAAACCG |
| pML 8011 F1 | ATCATCATCACAGCAGCGGCCTGGTGCCGCGCGGCAGCATGGCTCCTACCACATTTTCCC |
| pML 8011 R1 | TCAGTGGTGGTGGTGGTGGTGCTCGAGTCATCAATGTACCTTGTCTACAAACTTCATGAC |

**Supplementary Table 2 (continued).** Primers used in this study.

| Primers | Sequence (5'-3') |
|---|---|
| pML 8012 F1 | ATCATCATCACAGCAGCGGCCTGGTGCCGCGCGGCAGCATGGGAAGTGTGGGCATC |
| pML 8012 R1 | GTGGTGGTGGTGGTGGTGCTCGAGTCATTAGAAATCTTGGTAAATTACAGTGTAGCTATT |
| pML 8013 F1 | CATCATCACAGCAGCGGCCTGGTGCCGCGCGGCAGCATGAAGTACCCTACTACCGGACAG |
| pML 8013 R1 | CCGGATCTCAGTGGTGGTGGTGGTGGTGCTCGAGTCATCACTCTGAGCTCTCCGCG |
| pML 8014 F1 | CATCATCACAGCAGCGGCCTGGTGCCGCGCGGCAGCATGACCACGAAAAGCTTCTCGTTG |
| pML 8014 R1 | AGTGGTGGTGGTGGTGGTGCTCGAGTCATCAATGCAAACGATCTACAAAGCTAATAACTT |
| pML 8015 F1 | CATCACAGCAGCGGCCTGGTGCCGCGCGGCAGCATGATCGCCGCTAAAAAAACGGACT |
| pML 8015 R1 | GCAGCCGGATCTCAGTGGTGGTGGTGGTGGTGCTCGAGTCACCCAAGGCACCCGGACTC |
| pML 8016 F1 | AAATAATTTTGTTTAACTTTAAGAAGGAGATATACCATGGCAACCGATTTTAGCAAACTG |
| pML 8016 R1 | GTTAGCAGCCGGATCTCAGTGGTGGTGGTGGTGGTGATCGCTTGATTTTTCTGCAACCAG |

**Supplementary Table 3.** Plasmids used in this study

| Plasmids | Vector | Genes |
|---|---|---|
| pML 8001 | pYTU | *cp1A* (oxygenase)-*cp1B* (HP) |
| pML 8002 | pYTP | *cp1C* (decarboxylase)-*cp1D* (AMP-binding) |
| pML 8003 | pYTP | *cp1C* (decarboxylase)-*cp1D* (AMP-binding)-*cp1A* (oxygenase) |
| pML 8004 | pYTP | *cp1C* (decarboxylase)-*cp1D* (AMP-binding)-*cp1B* (HP) |
| pML 8005 | pYTU | *cp1A* (oxygenase)-*cp1B* (HP)-*cp1C* (decarboxylase) |
| pML 8006 | pYTU | *cp1A* (oxygenase)-*cp1B* (HP)-*cp1D* (AMP-binding) |
| pML 8007 | pYTP | *cp2A* (oxygenase)-*cp2D* (AMP-binding) |
| pML 8008 | pYTU | *cp2C* (decarboxylase)-*cp2B* (HP) |
| pML 8009 | pET28a | *cp1A* (oxygenase) |
| pML 8010 | pET28a | *cp1B* (HP) |
| pML 8011 | pET28a | *cp1D* (AMP-binding) |
| pML 8012 | pET28a | *cp2C* (decarboxylase) |
| pML 8013 | pET28a | *cp2B* (HP) |
| pML 8014 | pET28a | *cp2D* (AMP-binding) |
| pML 8015 | pET28a | *mfaA* (oxygenase) |
| pML 8016 | pET28a | Polyphosphate kinase |

**Supplementary Table 4.** X-ray data collection and refinement statistics of Cp1B.

| | Cp1B (PDB 9CJN) |
|---|---|
| **Data collection** | |
| Space group | $P2_12_12_1$ |
| Cell dimensions | |
| *a*, *b*, *c* (Å) | 69.85, 96.23, 129.0 |
| α, β, γ (°) | 90.00, 90.00, 90.00 |
| Resolution (Å) | 77.14-2.70 (2.80-2.70) |
| $R_{sym}$ or $R_{merge}$ | 0.130 (1.22) |
| $I$ / σ$I$ | 16.0 (2.2) |
| *CC(1/2)* | 0.999 (0.774) |
| Completeness (%) | 99.9 (99.1) |
| Redundancy | 13.1 (13.0) |
| **Refinement** | |
| Resolution (Å) | 77.14-2.70 (2.79-2.70) |
| No. reflections | 24508 (2358) |
| $R_{work}$ / $R_{free}$ | 0.1932/0.2346 |
| No. atoms | 4214 |
| Protein | 3967 |
| Ligand/ion | 162 |
| Water | 85 |
| *B*-factors (Å$^2$) | 66.78 |
| Protein | 65.82 |
| Ligand/ion | 96.33 |
| Water | 55.31 |
| R.m.s. deviations | |
| Bond lengths (Å) | 0.003 |
| Bond angles (°) | 0.57 |
| Ramachandran plot (%) | |
| Outliers | 0.00 |
| Favored | 96.37 |
| Allowed | 3.63 |

The structure was obtained using diffraction data from a single crystal. Values in parentheses are for highest-resolution shell.

**Supplementary Table 5.** Table of statistics from crystallographic data reduction and refinement for structure of papain without inhibitor bound, or co-crystallized with E64, E64-d or the biosynthetic variant, and (2*S*,3*S*)-***t*-ES-a9-b7**.

| | Apo Papain (PDB, 9CLH) | Papain E-64 (**1**) (PDB, 9CKT) | Papain E-64d (PDB, 9CKW) | Papain (2*S*,3*S*)-***t*-ES-a9-b7** (PDB, 9CKY) |
|---|---|---|---|---|
| Temperature | 100 K | 100 K | 100 K | 100 K |
| Wavelength | 1.54 Å | 1.54 Å | 1.54 Å | 1.54 Å |
| Data processing | | | | |
| Crystal system | Orthorhombic | Orthorhombic | Orthorhombic | Orthorhombic |
| Space group | $P2_12_12_1$ | $P2_12_12_1$ | $P2_12_12_1$ | $P2_12_12_1$ |
| Unit cell constants: | | | | |
| a, b, c (Å) | 42.45, 49,23, 101.66 | 42.41, 48.83, 101.73 | 43.88, 49.30, 90.69 | 42.47, 48.95, 101.58 |
| α, β, γ (°) | 90.00, 90.00, 90.00 | 90.00, 90.00, 90.00 | 90.00, 90.00, 90.00 | 90.00, 90.00, 90.00 |
| Resolution (Å) | 50.83 – 1.50 (1.60 – 1.50) | 39.14 – 1.40 (1.50 – 1.40) | 26.56 – 1.40 (1.50 – 1.40) | 35.25 – 1.40 (1.50 – 1.40) |
| No. unique reflections | 62159 (10442) | 41260 (6615) | 39328 (7086) | 42180 (7511) |
| $R_{merge}$ | 0.065 (0.143) | 0.065 (0.507) | 0.086 (0.186) | 0.065 (0.653) |
| $R_{meas}$ | 0.07 (0.154) | 0.067 (0.531) | 0.09 (0.196) | 0.068 (0.687) |
| Completeness (%) | 88.1 (61.9) | 97.1 (84.8) | 99.5 (97.5) | 99.0 (95.9) |
| Redundancy | 7.30 (7.56) | 12.66 (11.37) | 12.70 (10.09) | 12.56 (10.22) |
| I/σ | 19.50 (10.75) | 21.48 (5.03) | 20.47 (9.76) | 20.70 (4.07) |
| $CC_{1/2}$ (%) | 99.8 (98.9) | 99.9 (95.7) | 99.8 (98.5) | 99.9 (91.7) |
| Refinement program | *PHENIX* | *PHENIX* | *PHENIX* | *PHENIX* |
| $R_{work}$, $R_{free}$ | 0.1387, 0.1678 | 0.1901, 0.2211 | 0.1366, 0.1715 | 0.1597, 0.1988 |
| B-factors ($Å^2$) | | | | |
| Protein | 13.01 | 18.69 | 11.63 | 18.99 |
| Ligand/ion/other | 9.46 | 27.61 | 29.46 | 28.16 |
| Water | 25.38 | 25.25 | 28.55 | 34.50 |
| RMSD Bonds (Å) | 0.005 | 0.006 | 0.006 | 0.004 |
| RMSD Angles (°) | 0.722 | 0.950 | 0.840 | 0.749 |
| Ramachandran statistics | | | | |
| Outliers (%) | 0.00 | 0.00 | 0.00 | 0.55 |
| Favored (%) | 98.07 | 98.10 | 98.10 | 98.35 |
| Allowed (%) | 1.93 | 1.90 | 1.90 | 1.10 |
| Number of protein atoms | 1821 | 1707 | 1804 | 1765 |
| Number of water atoms | 251 | 182 | 294 | 319 |
| Number of ligand/ion atoms | 2 | 27 | 26 | 28 |
| Occupancy of inhibitor atoms | N/A | 80% | 75% | 81% |

**Supplementary Table 6.** Spectroscopic data of compound **E-64 (1)**

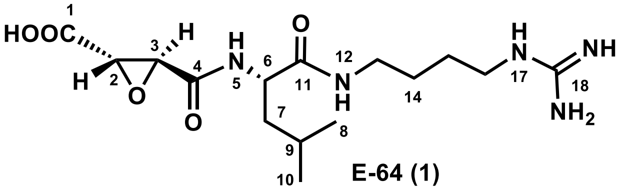

**E-64 (1)**

| Position | E-64 (**1**) in DMSO-$d_6$ $\delta_H$ ($J$ in Hz) | Position |
|---|---|---|
| 1 | | 170.8, C |
| 2 | 3.02, d (1.9) | 54.3, CH |
| 3 | 3.35, d (1.8) | 52.1, CH |
| 4 | | 167.2, C |
| 5 | 8.75, d (8.4) | |
| 6 | 4.25, t (5.5, 8.9) | 51.5, CH |
| 7 | 1.43, m | 40.7, $CH_2$ |
| 8 | 1.56, m | 24.3, CH |
| 9 | 0.83, m | 21.6, $CH_3$ |
| 10 | 0.86, m | 22.9, $CH_3$ |
| 11 | | 171.8, C |
| 12 | 8.22, t (5.8) | |
| 13 | 2.98, m<br>3.09, m | 37.7, $CH_2$ |
| 14 | 1.39, m | 26.1, $CH_2$ |
| 15 | 1.42, m | 25.7, $CH_2$ |
| 16 | 3.08, m | 40.3, $CH_2$ |
| 17 | 8.46, t (5.8) | |
| 18 | | 157.1, C |

NMR spectrum (500 MHz) for $^1H$, NMR spectrum (125 MHz) for $^{13}C$, DMSO-$d_6$, "m" means overlapped or multiple with other signals. Chemical shifts are reported in ppm.

HRMS (ESI, M+H$^+$) calculated for $C_{15}H_{28}N_5O_5^+$ 358.2085; found 358.2079.

$[\alpha]_D^{24.1}$ + 56° ($c$ 0.1, MeOH). Compound **1** showed the same positive optical rotation as reported E-64[5].

**Supplementary Table 7.** Spectroscopic data of compound **CLIK-148 (3)**

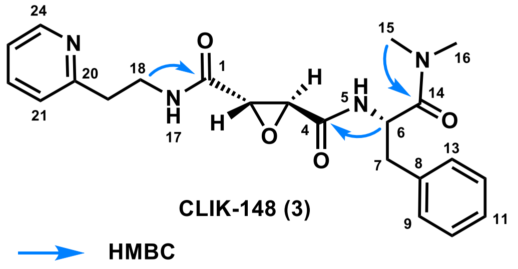

CLIK-148 (3)

| Position | **CLIK-148 (3)** in $CDCl_3$ $\delta_H$ (*J* in Hz) | $\delta_C$, type |
|---|---|---|
| 1 | | 166.8, C |
| 2 | 3.44, s | 54.1, CH |
| 3 | 3.32, m | 54.0, CH |
| 4 | | 165.7, C |
| 5 | 7.44, d (8.3) | |
| 6 | 5.07, m | 50.1, CH |
| 7 | 2.86, m | 39.3, $CH_2$ |
| | 2.96, m | |
| 8 | | 135.8, C |
| 9 | 7.15, m | 129.5, CH |
| 10 | 7.26, m | 128.7, CH |
| 11 | 7.23, m | 127.4, CH |
| 12 | 7.26, m | 128.7, CH |
| 13 | 7.15, m | 129.5, CH |
| 14 | | 170.8, C |
| 15 | 2.88, s | 35.9, $CH_3$ |
| 16 | 2.68, s | 37.1, $CH_3$ |
| 17 | 7.97, d (6.3) | |
| 18 | 3.73, m | 38.5, $CH_2$ |
| 19 | 3.31, m | 33.7, $CH_2$ |
| 20 | | 155.4, C |
| 21 | 7.79, d (8.0) | 127.7, CH |
| 22 | 8.28, t (7.8) | 145.1, CH |
| 23 | 7.74, t (6.8) | 124.9, CH |
| 24 | 8.72, d (5.7) | 141.9, CH |

NMR spectrum (500 MHz) for $^1H$, NMR spectrum (125 MHz) for $^{13}C$, $CDCl_3$, "m" means overlapped or multiple with other signals. Chemical shifts are reported in ppm.

HRMS (ESI, $M+H^+$) calculated for $C_{22}H_{27}N_4O_4^+$ 411.2027; found 411.2014.

$[\alpha]_D^{24.1}$ + 66° (*c* 0.1, MeOH). Compound **3** showed the same positive optical rotation as reported CLIK-148[6].

**Supplementary Table 8.** Spectroscopic data of compound **CPI-2 (4)**

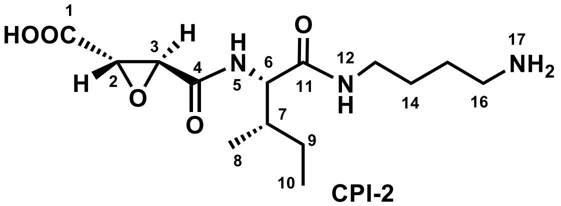

**CPI-2**

$^{1}$H NMR spectrum (500 MHz), $^{13}$C NMR spectrum (125 MHz), DMSO-$d_6$

| Position | CPI-2 (**4**) in DMSO-$d_6$ $\delta_H$ ($J$ in Hz) | $\delta_C$, type | Reported CPI-2 (**4**) in DMSO-$d_6$[7] $\delta_C$, type |
|---|---|---|---|
| 1 | | 168.8, C | 168.4, C |
| 2 | 3.75, d (1.7) | 52.5, CH | 51.2, CH |
| 3 | 3.46, d (1.7) | 51.3, CH | 52.5, CH |
| 4 | | 165.0, C | 164.6, C |
| 5 | 8.52, d (8.8) | | |
| 6 | 4.15 t (8.3) | 57.1, CH | 57.0, CH |
| 7 | 1.72, d (9.8) | 36.6, CH | 36.6, CH |
| 8 | 0.82 m | 15.4, $CH_3$ | 15.4, $CH_3$ |
| 9 | 1.08, m; 1.43, m | 24.4, $CH_2$ | 24.4, $CH_2$ |
| 10 | 0.82, m | 10.9, $CH_3$ | 11.0, $CH_3$ |
| 11 | | 170.3, C | 169.9, C |
| 12 | 8.17, t (5.8) | | |
| 13 | 3.01, m; 3.09, m | 38.0, $CH_2$ | 37.9, $CH_2$ |
| 14 | 1.42, m | 25.9, $CH_2$ | 25.9, $CH_2$ |
| 15 | 1.52, m | 24.6, $CH_2$ | 24.6, $CH_2$ |
| 16 | 2.78, d (6.8) | 38.5, $CH_2$ | 38.5, $CH_2$ |

NMR spectrum (500 MHz) for $^{1}$H, NMR spectrum (125 MHz) for $^{13}$C, DMSO-$d_6$, "m" means overlapped or multiple with other signals. Chemical shifts are reported in ppm.

HRMS (ESI, M+H$^+$) calculated for $C_{14}H_{26}N_3O_5^+$ 316.1867; found 316.1878.

$[\alpha]_D^{24.1}$ + 46° ($c$ 0.1, MeOH). Compound **4** showed the same positive optical rotation as reported CPI-2[7].

**Supplementary Table 9.** Spectroscopic data of compound **CPI-3 (5)**

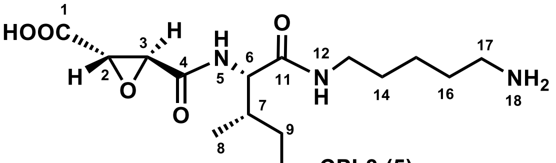

| Position | CPI-3 (**5**) in DMSO-$d_6$ $\delta_H$ (*J* in Hz) | $\delta_C$, type | Reported CPI-3 (**5**) in DMSO-$d_6$[7] $\delta_C$, type |
|---|---|---|---|
| 1 | | 168.8, C | 168.4, C |
| 2 | 3.75, d (1.7) | 52.5, CH | 52.5, CH |
| 3 | 3.46, d (1.7) | 51.2, CH | 51.2, CH |
| 4 | | 165.0, C | 164.6, C |
| 5 | 8.51, d (8.8) | | |
| 6 | 4.14 t (8.3) | 57.1, CH | 57.0, CH |
| 7 | 1.73, m | 36.6, CH | 36.6, CH |
| 8 | 0.82, m | 15.3, $CH_3$ | 15.4, $CH_3$ |
| 9 | 1.08, m; 1.39, m | 24.4, $CH_2$ | 24.4, $CH_2$ |
| 10 | 0.82, m | 11.0, $CH_3$ | 11.0, $CH_3$ |
| 11 | | 170.2, C | 169.8, C |
| 12 | 8.11, t (5.7) | | |
| 13 | 2.98, m; 3.10, m | 38.2, $CH_2$ | 38.1, $CH_2$ |
| 14 | 1.38, m | 28.3, $CH_2$ | 28.3, $CH_2$ |
| 15 | 1.26, m | 23.1, $CH_2$ | 23.1, $CH_2$ |
| 16 | 1.52, m | 26.6, $CH_2$ | 26.6, $CH_2$ |
| 17 | 2.74, d (6.8) | 38.7, $CH_2$ | 38.7, $CH_2$ |

NMR spectrum (500 MHz) for $^1$H, NMR spectrum (125 MHz) for $^{13}$C, DMSO-$d_6$, "m" means overlapped or multiple with other signals. Chemical shifts are reported in ppm.

HRMS (ESI, M+H$^+$) calculated for $C_{15}H_{28}N_3O_5^+$ 330.2023; found 330.2009.

$[\alpha]_D^{24.1}$ + 42° (*c* 0.1, MeOH).

Compound **5** showed the same positive optical rotation as reported CPI-3[7].

**Supplementary Table 10.** Spectroscopic data of compound **6**

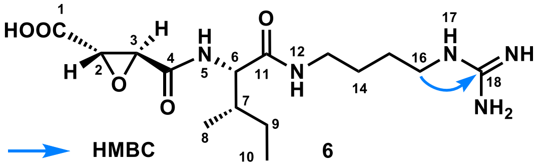

| Position | **6** in DMSO-$d_6$ (500MHz) $\delta_H$ ($J$ in Hz) | $\delta_C$, type |
|---|---|---|
| 1 | | 168.9, C |
| 2 | 3.74, d (1.9) | 52.5, CH |
| 3 | 3.44, d (1.8) | 51.3, CH |
| 4 | | 165.1, C |
| 5 | 8.51, d (8.8) | |
| 6 | 4.15, t (8.2) | 57.1, CH |
| 7 | 1.71, m | 36.6, CH |
| 8 | 0.82, m | 15.3, $CH_3$ |
| 9 | 1.07, m; 1.42, m | 24.4, $CH_2$ |
| 10 | 0.82, m | 10.9, $CH_3$ |
| 11 | | 170.2, C |
| 12 | 8.15, t (5.6) | |
| 13 | 3.00, m; 3.08, m | 38.0, $CH_2$ |
| 14 | 1.42, m | 26.0, $CH_2$ |
| 15 | 1.42, m | 26.1, $CH_2$ |
| 16 | 3.08, m | 40.4, $CH_2$ |
| 17 | 7.62, brs | |
| 18 | | 156.7, C |

NMR spectrum (500 MHz) for $^1$H, NMR spectrum (125 MHz) for $^{13}$C, DMSO-$d_6$, "m" means overlapped or multiple with other signals. Chemical shifts are reported in ppm.

HRMS (ESI, M+H$^+$) calculated for $C_{15}H_{28}N_5O_5^+$ 358.2085; found 358.2060.

$[\alpha]_D^{24.1}$ + 80° ($c$ 0.1, MeOH).

**Supplementary Table 11.** Spectroscopic data of compound **7**

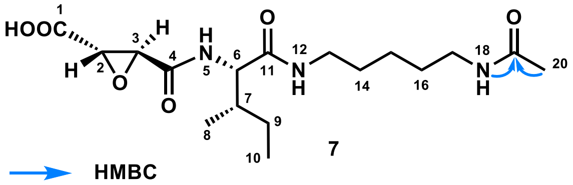

| Position | **7** in DMSO-$d_6$ (500MHz) $\delta_H$ ($J$ in Hz) | $\delta_C$, type |
|---|---|---|
| 1 | | 168.8, C |
| 2 | 3.75, d (1.7) | 52.5, CH |
| 3 | 3.46, d (1.7) | 51.2, CH |
| 4 | | 164.9, C |
| 5 | 8.51, d (8.9) | |
| 6 | 4.16 t (8.2) | 57.0, CH |
| 7 | 1.71, m | 36.7, CH |
| 8 | 0.81, m | 15.3, $CH_3$ |
| 9 | 1.07, m; 1.38, m | 24.3, $CH_2$ |
| 10 | 0.81, m | 11.0, $CH_3$ |
| 11 | | 170.1, C |
| 12 | 8.09, t (5.6) | |
| 13 | 2.98, m; 3.09, m | 38.4, $CH_2$ |
| 14 | 1.38, m | 28.8, $CH_2$ |
| 15 | 1.23, m | 23.8, $CH_2$ |
| 16 | 1.38, m | 28.6, $CH_2$ |
| 17 | 2.98, m | 38.4, $CH_2$ |
| 18 | 7.77 t (5.5) | |
| 19 | | 168.9, C |
| 20 | 1.77, s | 22.6, $CH_3$ |

NMR spectrum (500 MHz) for $^1$H, NMR spectrum (125 MHz) for $^{13}$C, DMSO-$d_6$, "m" means overlapped or multiple with other signals. Chemical shifts are reported in ppm.

HRMS (ESI, M+H$^+$) calculated for $C_{15}H_{28}N_5O_5^+$ 372.2129; found 372.2112.

$[\alpha]_D^{24.1}$ + 40° ($c$ 0.1, MeOH).

**Supplementary Table 12.** Spectroscopic data of compound **8**

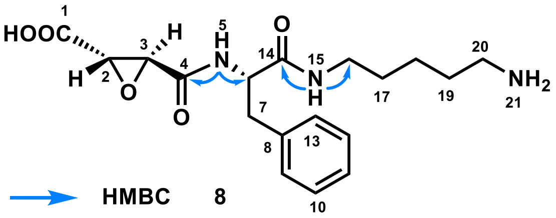

| Position | **8** in DMSO-$d_6$ $\delta_H$ ($J$ in Hz) | $\delta_C$, type |
|---|---|---|
| 1 | | 168.7, C |
| 2 | 3.59, s | 52.6, CH |
| 3 | 3.30, s | 51.4, CH |
| 4 | | 165.0, C |
| 5 | 8.63, d (8.5) | |
| 6 | 4.49, td (5.4, 8.9) | 54.2, CH |
| 7 | 2.81, dd (9.4, 13.6); 3.05, m | 37.8, $CH_2$ |
| 8 | | 137.6, C |
| 9 | 7.22, m | 129.2, CH |
| 10 | 7.27, m | 128.1, CH |
| 11 | 7.19, m | 126.4, CH |
| 12 | 7.27, m | 128.1, CH |
| 13 | 7.22, m | 129.2, CH |
| 14 | | 170.2, C |
| 15 | 8.10, t (5.6) | |
| 16 | 2.96, m | 38.3, $CH_2$ |
| 17 | 1.34, m | 28.4, $CH_2$ |
| 18 | 1.22, m | 23.1, $CH_2$ |
| 19 | 1.50, m | 26.7, $CH_2$ |
| 20 | 2.73, m | 38.7, $CH_2$ |

NMR spectrum (500 MHz) for $^1$H, NMR spectrum (125 MHz) for $^{13}$C, DMSO-$d_6$, "m" means overlapped or multiple with other signals. Chemical shifts are reported in ppm.

HRMS (ESI, M+H$^+$) calculated for $C_{18}H_{26}N_3O_5^+$ 364.1867; found 364.1860.

$[\alpha]_D^{24.1}$ + 136° ($c$ 0.1, MeOH).

**Supplementary Table 13.** Spectroscopic data of compound **9**

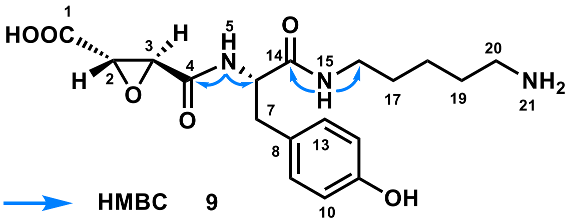

| Position | **9** in DMSO-$d_6$ $\delta_H$ (*J* in Hz) | $\delta_C$, type |
|---|---|---|
| 1 | | 168.7, C |
| 2 | 3.59, s | 52.6, CH |
| 3 | 3.30, s | 51.3, CH |
| 4 | | 164.9, C |
| 5 | 8.54, d (8.5) | |
| 6 | 4.39, td (5.3, 8.9) | 54.5, CH |
| 7 | 2.85, dd (5.3, 13.7); 3.05, m | 37.1, $CH_2$ |
| 8 | | 127.5, C |
| 9 | 6.99, m | 130.1, CH |
| 10 | 6.64, m | 114.9, CH |
| 11 | | 155.9, C |
| 12 | 6.64, m | 114.9, CH |
| 13 | 6.99, m | 130.1, CH |
| 14 | | 170.3, C |
| 15 | 8.05, t (5.6) | |
| 16 | 2.96, m | 38.3, $CH_2$ |
| 17 | 1.35, m | 28.4, $CH_2$ |
| 18 | 1.23, m | 23.1, $CH_2$ |
| 19 | 1.50, m | 26.7, $CH_2$ |
| 20 | 2.75, m | 38.7, $CH_2$ |

NMR spectrum (500 MHz) for $^1$H, NMR spectrum (125 MHz) for $^{13}$C, DMSO-$d_6$, "m" means overlapped or multiple with other signals. Chemical shifts are reported in ppm.

HRMS (ESI, M+H$^+$) calculated for $C_{18}H_{26}N_3O_6^+$ 380.1816; found 380.1793.

$[\alpha]_D^{24.1}$ + 86° (*c* 0.1, MeOH).

**Supplementary Table 14.** Spectroscopic data of **10**

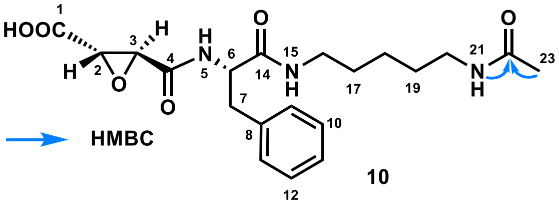

**10**

| Position | **10** in DMSO-$d_6$ $\delta_H$ ($J$ in Hz) | $\delta_C$, type |
|---|---|---|
| 1 | | 168.9, C |
| 2 | 3.59, s | 52.6, CH |
| 3 | 3.30, s | 51.3, CH |
| 4 | | 164.9, C |
| 5 | 8.61, d (8.5) | |
| 6 | 4.50, td (5.3, 9.0) | 54.1, CH |
| 7 | 2.80, dd (9.4, 13.6); 3.05, m | 37.9, $CH_2$ |
| 8 | | 137.5, C |
| 9 | 7.22, m | 129.2, CH |
| 10 | 7.26, m | 128.4, CH |
| 11 | 7.19, m | 126.4, CH |
| 12 | 7.26, m | 128.4, CH |
| 13 | 7.22, m | 129.2, CH |
| 14 | | 170.1, C |
| 15 | 8.07, t (5.6) | |
| 16 | 2.98, m | 38.4, $CH_2$ |
| 17 | 1.35, m | 28.8, $CH_2$ |
| 18 | 1.19, m | 23.7, $CH_2$ |
| 19 | 1.35, m | 28.6, $CH_2$ |
| 20 | 2.98, m | 38.5, $CH_2$ |
| 21 | 7.77, t (5.6) | |
| 22 | | 168.7, C |
| 23 | 1.77, s | 22.6, $CH_3$ |

NMR spectrum (500 MHz) for $^1$H, NMR spectrum (125 MHz) for $^{13}$C, DMSO-$d_6$, "m" means overlapped or multiple with other signals. Chemical shifts are reported in ppm.

HRMS (ESI, M+H$^+$) calculated for $C_{20}H_{28}N_3O_6^+$ 406.1973; found 406.1949.

$[\alpha]_D^{24.1}$ + 84° ($c$ 0.1, MeOH).

**Supplementary Table 15.** Spectroscopic data of compound **12**

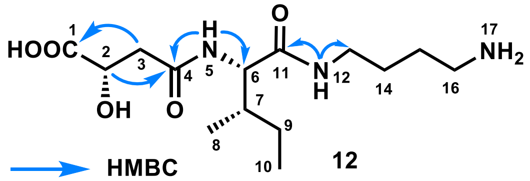

**12**

| Position | **12** in DMSO-$d_6$ (500MHz) $\delta_H$ ($J$ in Hz) | $\delta_C$, type |
|---|---|---|
| 1 | | 172.1, C |
| 2 | 4.22, d (1.7) | 68.3, CH |
| 3 | 2.36, m; 2.64, m | 40.0, $CH_2$ |
| 4 | | 172.1, C |
| 5 | 7.49, d (6.0) | |
| 6 | 4.14 t (8.3) | 56.2, CH |
| 7 | 1.67, d (9.8) | 37.4, CH |
| 8 | 0.82 m | 15.3, $CH_3$ |
| 9 | 1.08, m; 1.43, m | 24.2, $CH_2$ |
| 10 | 0.82, m | 11.1, $CH_3$ |
| 11 | | 170.5, C |
| 12 | 8.15, m | |
| 13 | 3.01, m; 3.09, m | 37.9, $CH_2$ |
| 14 | 1.42, m | 26.0, $CH_2$ |
| 15 | 1.52, m | 24.6, $CH_2$ |
| 16 | 2.78, d (6.8) | 38.6, $CH_2$ |

NMR spectrum (500 MHz) for $^1H$, NMR spectrum (125 MHz) for $^{13}C$, DMSO-$d_6$, "m" means overlapped or multiple with other signals. Chemical shifts are reported in ppm.

HRMS (ESI, M+H$^+$) calculated for $C_{14}H_{28}N_3O_5^+$ 318.2023; found 318.2000.

$[\alpha]_D^{24.1}$ + 56° ($c$ 0.1, MeOH).

**Supplementary Table 16.** Spectroscopic data of compound **13**

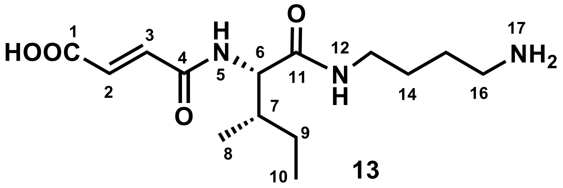

| Position | **13** in DMSO-$d_6$ (500MHz) $\delta_H$ ($J$ in Hz) | $\delta_C$, type |
|---|---|---|
| 1 | | 166.5, C |
| 2 | 7.12, d (15.5) | 137.0, CH |
| 3 | 6.51, d (15.5) | 129.9, CH |
| 4 | | 163.0, C |
| 5 | 8.58, d (8.8) | |
| 6 | 4.19 t (8.3) | 57.3, CH |
| 7 | 1.74, m | 36.5, CH |
| 8 | 0.79 m | 15.4, $CH_3$ |
| 9 | 1.09, m; 1.43, m | 24.4, $CH_2$ |
| 10 | 0.79, m | 10.9, $CH_3$ |
| 11 | | 170.5, C |
| 12 | 8.14, t (5.7) | |
| 13 | 3.01, m; 3.09, m | 37.9, $CH_2$ |
| 14 | 1.42, m | 26.0, $CH_2$ |
| 15 | 1.52, m | 24.6, $CH_2$ |
| 16 | 2.78, t (7.3) | 38.5, $CH_2$ |

NMR spectrum (500 MHz) for $^1$H, NMR spectrum (125 MHz) for $^{13}$C, DMSO-$d_6$, "m" means overlapped or multiple with other signals. Chemical shifts are reported in ppm.

HRMS (ESI, M+H$^+$) calculated for $C_{14}H_{26}N_3O_4^+$ 300.1918; found 300.1898.

$[\alpha]_D^{24.1}$ + 98° ($c$ 0.1, MeOH).

**Supplementary Table 17.** Spectroscopic data of (2*S*,3*S*)-*t*-ES-Ile (**14**)

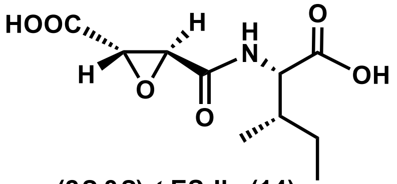

**(2*S*,3*S*)-*t*-ES-Ile (14)**

| Position | (2*S*,3*S*)-*t*-ES-Ile (**14**) in DMSO-$d_6$ $\delta_H$ (*J* in Hz) | $\delta_C$, type |
|---|---|---|
| 1 | | 169.0, C |
| 2 | 3.67, d (1.9) | 52.3, CH |
| 3 | 3.35, d (1.8) | 52.1, CH |
| 4 | | 165.9, C |
| 5 | 8.54, d (8.3) | |
| 6 | 4.22, dd (5.7, 8.4) | 56.5, CH |
| 7 | 1.81, m | 36.4, CH |
| 8 | 0.86, m | 15.6, $CH_3$ |
| 9 | 1.19, m; 1.39, m | 24.6, $CH_2$ |
| 10 | 0.86, m | 11.3, $CH_3$ |
| 11 | | 172.4, C |

NMR spectrum (500 MHz) for $^1$H, NMR spectrum (125 MHz) for $^{13}$C, DMSO-$d_6$, "m" means overlapped or multiple with other signals. Chemical shifts are reported in ppm.

HRMS (ESI, M+H$^+$) calculated for $C_{10}H_{16}NO_6^+$ 246.0972; found 246.0946.

$[\alpha]_D^{24.1}$ + 104° (*c* 0.1, MeOH).

**Supplementary Table 18.** Spectroscopic data of (2*S*,3*S*)-*t*-ES-Phe (**15**)

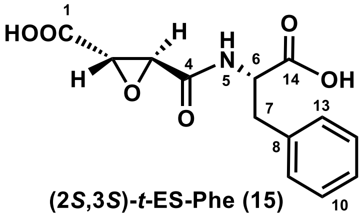

(2*S*,3*S*)-*t*-ES-Phe (15)

| | (2*S*,3*S*)-*t*-ES-Phe (**15**) in DMSO-$d_6$ | |
|---|---|---|
| Position | $\delta_H$ (*J* in Hz) | $\delta_C$, type |
| 1 | | 168.6, C |
| 2 | 3.59, d (1.8) | 52.5, CH |
| 3 | 3.30, d (1.8) | 51.3, CH |
| 4 | | 165.3, C |
| 5 | 8.67, d (8.3) | |
| 6 | 4.49, m | 53.4, CH |
| 7 | 2.92, m; 3.10, m | 36.5, $CH_2$ |
| 8 | | 137.3, C |
| 9 | 7.29, m | 129.2, CH |
| 10 | 7.22, m | 128.3, CH |
| 11 | 7.22, m | 126.6, CH |
| 12 | 7.29, m | 128.3, CH |
| 13 | 7.22, m | 129.2, CH |
| 14 | | 172.3, C |

NMR spectrum (500 MHz) for $^{1}H$, NMR spectrum (125 MHz) for $^{13}C$, DMSO-$d_6$, "m" means overlapped or multiple with other signals. Chemical shifts are reported in ppm.

HRMS (ESI, M+H$^+$) calculated for $C_{13}H_{14}NO_6^+$ 280.0816; found 280.0807.

$[\alpha]_D^{24.1}$ + 72° (*c* 0.1, MeOH).

**Supplementary Table 19.** Spectroscopic data of (2*S*,3*S*)-*t*-ES-Leu

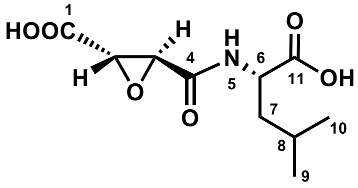

**(2*S*,3*S*)-*t*-ES-Leu**

| | (2*S*,3*S*)-*t*-ES-Leu in DMSO-$d_6$ (500MHz) | |
|---|---|---|
| Position | $\delta_H$ (*J* in Hz) | $\delta_C$, type |
| 1 | | 168.7, C |
| 2 | 3.63, d (1.9) | 52.5, CH |
| 3 | 3.45, d (1.8) | 51.3, CH |
| 4 | | 165.3, C |
| 5 | 8.69, d (8.0) | |
| 6 | 4.26, m | 50.4, CH |
| 7 | 1.54, m | 39.8, $CH_2$ |
| 8 | 1.62, m | 24.3, CH |
| 9 | 0.90, d (6.4) | 22.8, $CH_3$ |
| 10 | 0.85, d (6.4) | 21.2, $CH_3$ |
| 11 | | 173.4, C |

NMR spectrum (500 MHz) for $^1H$, NMR spectrum (125 MHz) for $^{13}C$, DMSO-$d_6$, "m" means overlapped or multiple with other signals. Chemical shifts are reported in ppm.

HRMS (ESI, M+H$^+$) calculated for $C_{10}H_{16}NO_6^+$ 246.0972; found 246.0965.

$[\alpha]_D^{24.1}$ + 90° (*c* 0.1, MeOH).

**Supplementary Table 20.** Spectroscopic data of (2*S*,3*S*)-*t*-ES-Val

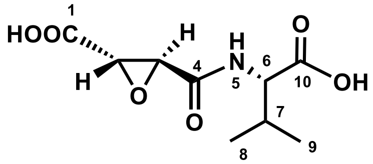

**(2*S*,3*S*)-*t*-ES-Val**

| Position | (2*S*,3*S*)-*t*-ES-Val in DMSO-$d_6$ (500MHz) $\delta_H$ (*J* in Hz) | $\delta_C$, type |
|---|---|---|
| 1 | | 168.8, C |
| 2 | 3.79, d (1.9) | 52.4, CH |
| 3 | 3.47, d (1.8) | 51.2, CH |
| 4 | | 165.4, C |
| 5 | 8.64, d (8.4) | |
| 6 | 4.20, dd (5.6, 8.5) | 57.4, CH |
| 7 | 2.09, m | 29.9, CH |
| 8 | 0.89, m | 17.8, $CH_3$ |
| 9 | 0.89, m | 19.1, $CH_3$ |
| 10 | | 172.4, C |

NMR spectrum (500 MHz) for $^1H$, NMR spectrum (125 MHz) for $^{13}C$, DMSO-$d_6$, "m" means overlapped or multiple with other signals. Chemical shifts are reported in ppm.

HRMS (ESI, M+H$^+$) calculated for $C_{10}H_{16}NO_6^+$ 232.0816; found 232.0820.

$[\alpha]_D^{24.1}$ + 152° (*c* 0.1, MeOH).

**Supplementary Table 21.** Spectroscopic data of (2*S*,3*S*)-*t*-ES-Tyr

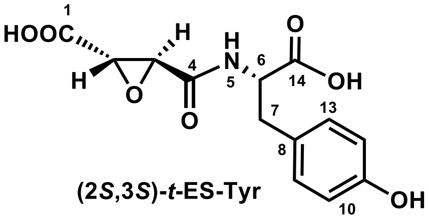

(2*S*,3*S*)-*t*-ES-Tyr

| Position | (2*S*,3*S*)-*t*-ES-Tyr in DMSO-$d_6$ (500MHz) $\delta_H$ (*J* in Hz) | $\delta_C$, type |
|---|---|---|
| 1 | | 168.6, C |
| 2 | 3.61, d (1.8) | 52.5, CH |
| 3 | 3.33, d (1.8) | 51.3, CH |
| 4 | | 165.1, C |
| 5 | 8.60, d (8.2) | |
| 6 | 4.40, m | 53.7, CH |
| 7 | 2.79, dd (9.4, 13.9); 2.97, dd (4.8, 13.9) | 35.8, $CH_2$ |
| 8 | | 127.0, C |
| 9 | 7.00, m | 130.1, CH |
| 10 | 6.66, m | 115.0, CH |
| 11 | | 156.0, C |
| 12 | 6.66, m | 115.0, CH |
| 13 | 7.00, m | 130.1, CH |
| 14 | | 172.4, C |

NMR spectrum (500 MHz) for $^1H$, NMR spectrum (125 MHz) for $^{13}C$, DMSO-$d_6$, "m" means overlapped or multiple with other signals. Chemical shifts are reported in ppm.

HRMS (ESI, M+H$^+$) calculated for $C_{13}H_{14}NO_7^+$ 296.0765; found 296.0777.

$[\alpha]_D^{24.1}$ + 72° (*c* 0.1, MeOH).

**Supplementary Table 22.** Spectroscopic data of (2*S*,3*S*)-*t*-ES-Trp

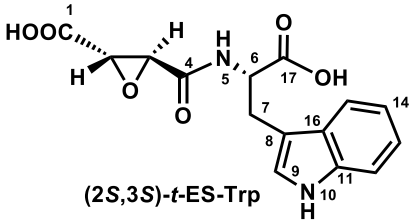

**(2*S*,3*S*)-*t*-ES-Trp**

| Position | (2*S*,3*S*)-*t*-ES-Trp in DMSO-$d_6$ (500MHz) $\delta_H$ (*J* in Hz) | $\delta_C$, type |
|---|---|---|
| 1 | | 168.6, C |
| 2 | 3.63, d (1.8) | 52.5, CH |
| 3 | 3.34, d (1.9) | 51.3, CH |
| 4 | | 165.2, C |
| 5 | 8.64, d (8.0) | |
| 6 | 4.52, td (4.9, 8.2) | 53.1, CH |
| 7 | 3.08, dd (4.0, 14.7); 3.20, dd (8.5, 14.7) | 26.9, $CH_2$ |
| 8 | | 136.1, C |
| 9 | 7.15, d (2.3) | 123.7, CH |
| 10 | 10.88, d (2.4) | |
| 11 | | 136.1, C |
| 12 | 7.34, d (8.1) | 111.5, CH |
| 13 | 7.07, m | 121.0, CH |
| 14 | 6.99, m | 118.5, CH |
| 15 | 7.53, d (7.9) | 118.2, CH |
| 16 | | 127.2, C |
| 17 | | 172.7, C |

NMR spectrum (500 MHz) for $^1$H, NMR spectrum (125 MHz) for $^{13}$C, DMSO-$d_6$, "m" means overlapped or multiple with other signals. Chemical shifts are reported in ppm.

HRMS (ESI, M+H$^+$) calculated for $C_{15}H_{15}N_2O_6^+$ 319.0925; found 319.0931.

$[\alpha]_D^{24.1}$ + 52° (*c* 0.1, MeOH).

**Supplementary Table 23.** Spectroscopic data of (2*S*,3*S*)-*t*-ES-a1

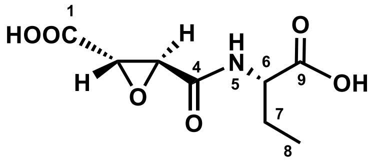

**(2*S*,3*S*)-*t*-ES-a1**

| Position | (2*S*,3*S*)-*t*-ES-a1 in DMSO-$d_6$ (500MHz) $\delta_H$ (*J* in Hz) | $\delta_C$, type |
|---|---|---|
| 1 | | 168.7, C |
| 2 | 3.69, d (1.8) | 52.5, CH |
| 3 | 3.47, d (1.8) | 51.2, CH |
| 4 | | 165.3, C |
| 5 | 8.68, d (7.7) | |
| 6 | 4.18, dd (5.1, 8.1) | 53.3, CH |
| 7 | 1.65, m; 1.75, m | 24.2, $CH_2$ |
| 8 | 0.88, m | 10.2, $CH_3$ |
| 9 | | 172.8, C |

NMR spectrum (500 MHz) for $^1$H, NMR spectrum (125 MHz) for $^{13}$C, DMSO-$d_6$, "m" means overlapped or multiple with other signals. Chemical shifts are reported in ppm.

HRMS (ESI, M+H$^+$) calculated for $C_8H_{12}NO_6^+$ 218.0659; found 218.0645.

$[\alpha]_D^{24.1}$ + 56° (*c* 0.1, MeOH).

**Supplementary Table 24.** Spectroscopic data of (2*S*,3*S*)-*t*-ES-a2

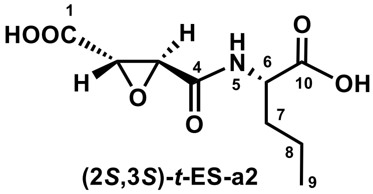

**(2*S*,3*S*)-*t*-ES-a2**

| Position | (2*S*,3*S*)-*t*-ES-a2 in DMSO-$d_6$ (500MHz) $\delta_H$ (*J* in Hz) | $\delta_C$, type |
|---|---|---|
| 1 | | 168.7, C |
| 2 | 3.67, d (1.8) | 52.5, CH |
| 3 | 3.47, d (1.8) | 51.2, CH |
| 4 | | 165.3, C |
| 5 | 8.70, d (7.8) | |
| 6 | 4.23, dd (5.0, 8.5) | 51.7, CH |
| 7 | 1.62, m; 1.69, m | 32.9, $CH_2$ |
| 8 | 1.32, m | 18.5, $CH_2$ |
| 9 | 0.87, m | 13.5, $CH_3$ |
| 10 | | 173.0, C |

NMR spectrum (500 MHz) for $^1$H, NMR spectrum (125 MHz) for $^{13}$C, DMSO-$d_6$, “m” means overlapped or multiple with other signals. Chemical shifts are reported in ppm.

HRMS (ESI, M+H$^+$) calculated for $C_9H_{14}NO_6^+$ 232.0816; found 232.0819.

$[\alpha]_D^{24.1}$ + 48° (*c* 0.1, MeOH).

**Supplementary Table 25.** Spectroscopic data of (2*S*,3*S*)-*t*-ES-a3

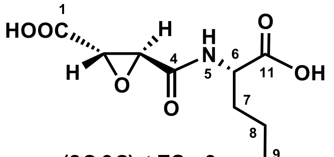

| Position | (2*S*,3*S*)-*t*-ES-a3 in DMSO-$d_6$ (500MHz) $\delta_H$ (*J* in Hz) | $\delta_C$, type |
|---|---|---|
| 1 | | 168.7, C |
| 2 | 3.67, d (1.8) | 52.5, CH |
| 3 | 3.46, d (1.8) | 51.4, CH |
| 4 | | 165.3, C |
| 5 | 8.69, d (7.8) | |
| 6 | 4.22, dd (4.9, 8.3) | 52.0, CH |
| 7 | 1.63, m; 1.72, m | 30.6, $CH_2$ |
| 8 | 1.27, m | 27.4, $CH_2$ |
| 9 | 1.27, m | 21.7, $CH_2$ |
| 10 | 0.86, m | 13.8, $CH_3$ |
| 11 | | 173.0, C |

NMR spectrum (500 MHz) for $^1H$, NMR spectrum (125 MHz) for $^{13}C$, DMSO-$d_6$, "m" means overlapped or multiple with other signals. Chemical shifts are reported in ppm.

HRMS (ESI, M+H$^+$) calculated for $C_{10}H_{16}NO_6^+$ 246.0972; found 246.0967.

$[\alpha]_D^{24.1}$ + 70 (*c* 0.1, MeOH).

**Supplementary Table 26.** Spectroscopic data of (2*S*,3*S*)-*t*-ES-a4

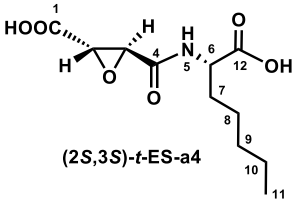

| Position | (2*S*,3*S*)-*t*-ES-a4 in DMSO-$d_6$ (500MHz) $\delta_H$ (*J* in Hz) | $\delta_C$, type |
|---|---|---|
| 1 | | 168.7, C |
| 2 | 3.67, d (1.8) | 52.5, CH |
| 3 | 3.45, d (1.8) | 51.3, CH |
| 4 | | 165.3, C |
| 5 | 8.69, d (7.8) | |
| 6 | 4.22, dd (4.9, 8.3) | 52.0, CH |
| 7 | 1.62, m; 1.71, m | 30.8, $CH_2$ |
| 8 | 1.27, m | 30.7, $CH_2$ |
| 9 | 1.27, m | 24.9, $CH_2$ |
| 10 | 1.27, m | 21.9, $CH_2$ |
| 11 | 0.86, t (6.8) | 13.9, $CH_3$ |
| 12 | | 173.0, C |

NMR spectrum (500 MHz) for $^1$H, NMR spectrum (125 MHz) for $^{13}$C, DMSO-$d_6$, "m" means overlapped or multiple with other signals. Chemical shifts are reported in ppm.

HRMS (ESI, M+H$^+$) calculated for $C_{11}H_{18}NO_6^+$ 260.1129; found 260.1133.

$[\alpha]_D^{24.1}$ + 64° (*c* 0.1, MeOH).

**Supplementary Table 27.** Spectroscopic data of (2*S*,3*S*)-*t*-ES-a5

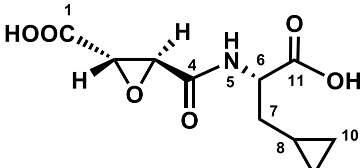

**(2*S*,3*S*)-*t*-ES-a5**

| | (2*S*,3*S*)-*t*-ES-a5 in DMSO-$d_6$ (500MHz) | |
|---|---|---|
| Position | $\delta_H$ (*J* in Hz) | $\delta_C$, type |
| 1 | | 168.7, C |
| 2 | 3.69, d (1.8) | 52.6, CH |
| 3 | 3.48, d (1.8) | 51.3, CH |
| 4 | | 165.2, C |
| 5 | 8.72, d (7.9) | |
| 6 | 4.30, dd (5.4, 8.0) | 52.5, CH |
| 7 | 1.60, m | 35.7, $CH_2$ |
| 8 | 0.75, m | 7.7, CH |
| 9 | 0.40, m | 4.5, $CH_2$ |
| 10 | 0.04, m; 0.14, m | 3.9, $CH_2$ |
| 11 | | 172.9, C |

NMR spectrum (500 MHz) for $^1$H, NMR spectrum (125 MHz) for $^{13}$C, DMSO-$d_6$, "m" means overlapped or multiple with other signals. Chemical shifts are reported in ppm.

HRMS (ESI, M+H$^+$) calculated for $C_{10}H_{14}NO_6^+$ 244.0816; found 244.0788.

$[\alpha]_D^{24.1}$ + 120º (*c* 0.1, MeOH).

**Supplementary Table 28.** Spectroscopic data of (2*S*,3*S*)-*t*-ES-a6

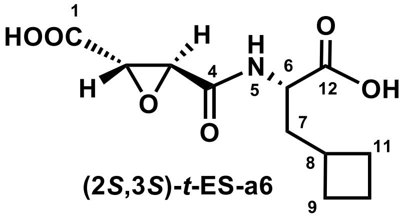

**(2*S*,3*S*)-*t*-ES-a6**

| Position | (2*S*,3*S*)-*t*-ES-a6 in DMSO-$d_6$ (500MHz) $\delta_H$ (*J* in Hz) | $\delta_C$, type |
|---|---|---|
| 1 | | 169.1, C |
| 2 | 3.65, d (2.0) | 52.9, CH |
| 3 | 3.46, d (1.9) | 51.7, CH |
| 4 | | 165.6, C |
| 5 | 8.65, d (8.2) | |
| 6 | 4.14, m | 51.1, CH |
| 7 | 1.75, m; 1.81, m | 38.3, $CH_2$ |
| 8 | 2.32, m | 32.8, CH |
| 9 | 1.62, m; 1.97, m | 28.0, $CH_2$ |
| 10 | 1.76, m | 18.5, $CH_2$ |
| 11 | 1.62, m; 1.97, m | 28.2, $CH_2$ |
| 12 | | 173.5, C |

NMR spectrum (500 MHz) for $^1$H, NMR spectrum (125 MHz) for $^{13}$C, DMSO-$d_6$, "m" means overlapped or multiple with other signals. Chemical shifts are reported in ppm.

HRMS (ESI, M+H$^+$) calculated for $C_{11}H_{16}NO_6^+$ 258.0972; found 258.0965.

$[\alpha]_D^{24.1}$ + 100° (*c* 0.1, MeOH).

**Supplementary Table 29.** Spectroscopic data of (2*S*,3*S*)-*t*-ES-a7

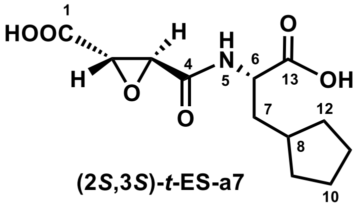

| Position | (2*S*,3*S*)-*t*-ES-a7 in DMSO-$d_6$ (500MHz) $\delta_H$ (*J* in Hz) | $\delta_C$, type |
|---|---|---|
| 1 | | 168.7, C |
| 2 | 3.64, d (1.8) | 52.5, CH |
| 3 | 3.45, d (1.8) | 51.6, CH |
| 4 | | 165.3, C |
| 5 | 8.70, d (7.9) | |
| 6 | 4.22, m | 51.3, CH |
| 7 | 1.69, m | 37.1, $CH_2$ |
| 8 | 1.82, m | 36.3, CH |
| 9 | 1.72, m | 32.2, $CH_2$ |
| 10 | 1.08, m | 31.7, $CH_2$ |
| 11 | 1.56, m | 24.7, $CH_2$ |
| 12 | 1.47, m | 24.5, $CH_2$ |
| 13 | | 173.3, C |

NMR spectrum (500 MHz) for $^1$H, NMR spectrum (125 MHz) for $^{13}$C, DMSO-$d_6$, "m" means overlapped or multiple with other signals. Chemical shifts are reported in ppm.

HRMS (ESI, M+H$^+$) calculated for $C_{12}H_{17}NO_6^+$ 272.1129; found 272.1134.

$[\alpha]_D^{24.1}$ + 44° (*c* 0.1, MeOH).

**Supplementary Table 30.** Spectroscopic data of (2*S*,3*S*)-*t*-ES-a8

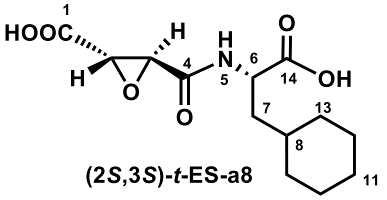

**(2*S*,3*S*)-*t*-ES-a8**

| Position | (2*S*,3*S*)-*t*-ES-a8 in DMSO-$d_6$ (500MHz) $\delta_H$ (*J* in Hz) | $\delta_C$, type |
|---|---|---|
| 1 | | 168.7, C |
| 2 | 3.64, d (1.8) | 52.5, CH |
| 3 | 3.44, d (1.8) | 51.3, CH |
| 4 | | 165.3, C |
| 5 | 8.70, d (7.9) | |
| 6 | 4.28, m | 49.7, CH |
| 7 | 1.57, m | 38.3, $CH_2$ |
| 8 | 1.31, m | 33.6, CH |
| 9 | 0.93, m; 1.64, m | 33.1, $CH_2$ |
| 10 | 1.60, m | 26.0, $CH_2$ |
| 11 | 1.13, m | 25.7, $CH_2$ |
| 12 | 0.83, m; 1.69, m | 31.5, $CH_2$ |
| 13 | 1.65, m | 25.5, $CH_2$ |
| 14 | | 173.5, C |

NMR spectrum (500 MHz) for $^1$H, NMR spectrum (125 MHz) for $^{13}$C, DMSO-$d_6$, "m" means overlapped or multiple with other signals. Chemical shifts are reported in ppm.

HRMS (ESI, M+H$^+$) calculated for $C_{13}H_{20}NO_6^+$ 286.1285; found 286.1283.

$[\alpha]_D^{24.1}$ + 48° (*c* 0.1, MeOH).

**Supplementary Table 31.** Spectroscopic data of (2*S*,3*S*)-*t*-ES-a9

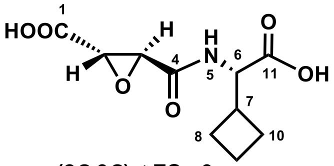

(2*S*,3*S*)-*t*-ES-a9

| Position | (2*S*,3*S*)-*t*-ES-a9 in DMSO-$d_6$ (500MHz) $\delta_H$ (*J* in Hz) | $\delta_C$, type |
|---|---|---|
| 1 | | 168.7, C |
| 2 | 3.72, d (1.8) | 52.4, CH |
| 3 | 3.46, d (1.8) | 51.2, CH |
| 4 | | 165.4, C |
| 5 | 8.70, d (8.0) | |
| 6 | 4.22, t (8.2) | 56.0, CH |
| 7 | 2.63, m | 36.3, CH |
| 8 | 1.82, m | 24.8, $CH_2$ |
| 9 | 1.94, m | 24.5, $CH_2$ |
| 10 | 1.74, m | 17.4, $CH_2$ |
| 11 | | 172.0, C |

NMR spectrum (500 MHz) for $^1$H, NMR spectrum (125 MHz) for $^{13}$C, DMSO-$d_6$, "m" means overlapped or multiple with other signals. Chemical shifts are reported in ppm.

HRMS (ESI, M+H$^+$) calculated for $C_{10}H_{14}NO_6^+$ 244.0816; found 244.0815.

$[\alpha]_D^{24.1}$ + 68° (*c* 0.1, MeOH).

**Supplementary Table 32.** Spectroscopic data of (2*S*,3*S*)-*t*-ES-a10

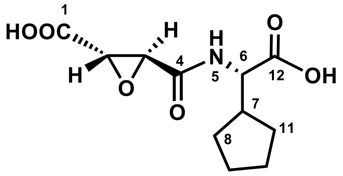

(2*S*,3*S*)-*t*-ES-a10

| | (2*S*,3*S*)-*t*-ES-a10 in DMSO-$d_6$ (500MHz) | |
|---|---|---|
| Position | $\delta_H$ (*J* in Hz) | $\delta_C$, type |
| 1 | | 168.8, C |
| 2 | 3.73, d (1.8) | 52.4, CH |
| 3 | 3.46, d (1.8) | 51.2, CH |
| 4 | | 165.3, C |
| 5 | 8.73, d (8.2) | |
| 6 | 4.18, m | 55.5, CH |
| 7 | 2.21, m | 41.1, CH |
| 8 | 1.66, m | 28.3, $CH_2$ |
| 9 | 1.49, m | 24.8, $CH_2$ |
| 10 | 1.56, m | 24.5, $CH_2$ |
| 11 | 1.29, m | 28.6, $CH_2$ |
| 12 | | 172.7, C |

NMR spectrum (500 MHz) for $^1$H, NMR spectrum (125 MHz) for $^{13}$C, DMSO-$d_6$, "m" means overlapped or multiple with other signals. Chemical shifts are reported in ppm.

HRMS (ESI, M+H$^+$) calculated for $C_{11}H_{16}NO_6^+$ 258.0972; found 258.0964.

$[\alpha]_D^{24.1}$ + 104° (*c* 0.1, MeOH).

**Supplementary Table 33.** Spectroscopic data of (2*S*,3*S*)-*t*-ES-a11

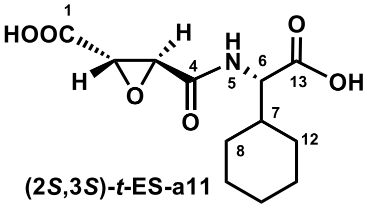

**(2*S*,3*S*)-*t*-ES-a11**

| | (2*S*,3*S*)-*t*-ES-a11 in DMSO-$d_6$ (500MHz) | |
|---|---|---|
| Position | $\delta_H$ (*J* in Hz) | $\delta_C$, type |
| 1 | | 168.8, C |
| 2 | 3.77, d (1.8) | 52.4, CH |
| 3 | 3.46, d (1.8) | 51.1, CH |
| 4 | | 165.3, C |
| 5 | 8.65, d (8.4) | |
| 6 | 4.18, m | 57.0, CH |
| 7 | 1.71, m | 39.0, CH |
| 8 | 1.01, m; 1.59, m | 29.1, $CH_2$ |
| 9 | 1.59, m | 25.5, $CH_2$ |
| 10 | 1.66, m | 25.6, $CH_2$ |
| 11 | 1.12, m | 25.5, $CH_2$ |
| 12 | 1.10, m; 1.56, m | 27.9, $CH_2$ |
| 13 | | 172.3, C |

NMR spectrum (500 MHz) for $^1$H, NMR spectrum (125 MHz) for $^{13}$C, DMSO-$d_6$, “m” means overlapped or multiple with other signals. Chemical shifts are reported in ppm.

HRMS (ESI, M+H$^+$) calculated for $C_{123}H_{18}NO_6^+$ 272.1129; found 272.1123.

$[\alpha]_D^{24.1}$ + 76° (*c* 0.1, MeOH).

**Supplementary Table 34.** Spectroscopic data of (2*S*,3*S*)-*t*-ES-a12

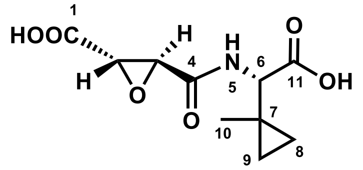

**(2*S*,3*S*)-*t*-ES-a12**

| | (2*S*,3*S*)-*t*-ES-a12 in DMSO-$d_6$ (500MHz) | |
|---|---|---|
| Position | $\delta_H$ (*J* in Hz) | $\delta_C$, type |
| 1 | | 168.8, C |
| 2 | 3.82, d (1.8) | 52.3, CH |
| 3 | 3.46, d (1.8) | 51.1, CH |
| 4 | | 165.1, C |
| 5 | 8.65, d (8.2) | |
| 6 | 3.73, m | 59.1, CH |
| 7 | | 17.7, C |
| 8 | 0.52, m; 0.69, m | 11.6, $CH_2$ |
| 9 | 0.32, m; 0.41, m | 12.0, $CH_2$ |
| 10 | 1.04, s | 19.2, $CH_3$ |
| 11 | | 171.6, C |

NMR spectrum (500 MHz) for $^1$H, NMR spectrum (125 MHz) for $^{13}$C, DMSO-$d_6$, "m" means overlapped or multiple with other signals. Chemical shifts are reported in ppm.

HRMS (ESI, M+H$^+$) calculated for $C_{10}H_{14}NO_6^+$ 244.0816; found 244.0810.

$[\alpha]_D^{24.1}$ + 132° (*c* 0.1, MeOH).

**Supplementary Table 35.** Spectroscopic data of (2*S*,3*S*)-*t*-ES-a13

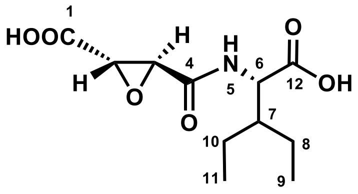

**(2*S*,3*S*)-*t*-ES-a13**

| | (2*S*,3*S*)-*t*-ES-a13 in DMSO-$d_6$ (500MHz) | |
|---|---|---|
| Position | $\delta_H$ (*J* in Hz) | $\delta_C$, type |
| 1 | | 168.8, C |
| 2 | 3.81, d (1.8) | 53.4, CH |
| 3 | 3.46, d (1.8) | 52.3, CH |
| 4 | | 165.5, C |
| 5 | 8.58, d (8.2) | |
| 6 | 4.44, m | 51.2, CH |
| 7 | 1.67, m | 43.1, CH |
| 8 | 1.18, m; 1.32, m | 22.0, $CH_2$ |
| 9 | 0.85, t (7.3) | 11.4, $CH_3$ |
| 10 | 1.32, m | 22.2, $CH_2$ |
| 11 | 0.85, t (7.3) | 11.5, $CH_3$ |
| 12 | | 173.0, C |

NMR spectrum (500 MHz) for $^1$H, NMR spectrum (125 MHz) for $^{13}$C, DMSO-$d_6$, "m" means overlapped or multiple with other signals. Chemical shifts are reported in ppm.

HRMS (ESI, M+H$^+$) calculated for $C_{11}H_{18}NO_6^+$ 260.1129; found 260.1105.

$[\alpha]_D^{24.1}$ + 138° (*c* 0.1, MeOH).

**Supplementary Table 36.** Spectroscopic data of (2*S*,3*S*)-*t*-ES-a14

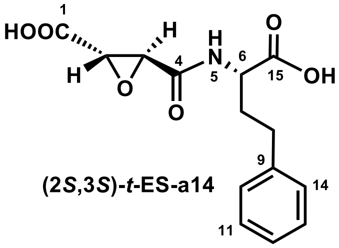

| | (2*S*,3*S*)-*t*-ES-a14 in DMSO-$d_6$ (500MHz) | |
|---|---|---|
| Position | $\delta_H$ (*J* in Hz) | $\delta_C$, type |
| 1 | | 169.2, C |
| 2 | 3.65, d (1.9) | 53.0, CH |
| 3 | 3.45, m | 52.0, CH |
| 4 | | 166.2, C |
| 5 | 8.73, d (7.9) | |
| 6 | 4.20, m | 52.2, CH |
| 7 | 1.93, m; 2.01, m | 33.1, $CH_2$ |
| 8 | 2.61, m | 31.8, $CH_2$ |
| 9 | | 141.3, C |
| 10 | 7.28, m | 128.8, CH |
| 11 | 7.19, m | 128.8, CH |
| 12 | 7.19, m | 126.4, CH |
| 13 | 7.28, m | 128.8, CH |
| 14 | 7.19, m | 128.8, CH |
| 15 | | 173.4, C |

NMR spectrum (500 MHz) for $^1$H, NMR spectrum (125 MHz) for $^{13}$C, DMSO-$d_6$, "m" means overlapped or multiple with other signals. Chemical shifts are reported in ppm.

HRMS (ESI, M+H$^+$) calculated for $C_{14}H_{16}NO_6^+$ 294.0972; found 294.0957.

$[\alpha]_D^{24.1}$ + 84° (*c* 0.1, MeOH).

**Supplementary Table 37.** Spectroscopic data of (2*S*,3*S*)-*t*-ES-a15

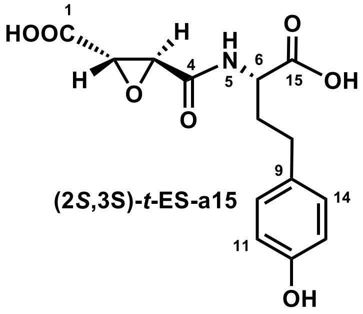

(2*S*,3S)-*t*-ES-a15

| Position | (2*S*,3*S*)-*t*-ES-a15 in DMSO-$d_6$ (500MHz) $\delta_H$ (*J* in Hz) | $\delta_C$, type |
|---|---|---|
| 1 | | 168.7, C |
| 2 | 3.67, d (1.9) | 52.6, CH |
| 3 | 3.48, m | 51.4, CH |
| 4 | | 165.5, C |
| 5 | 8.74, d (7.9) | |
| 6 | 4.17, m | 51.4, CH |
| 7 | 1.83, m; 1.95, m | 33.0, $CH_2$ |
| 8 | 2.44, m | 30.5, $CH_2$ |
| 9 | | 130.8, C |
| 10 | 6.97, m | 129.2, CH |
| 11 | 6.66, m | 115.1, CH |
| 12 | | 155.5, C |
| 13 | 6.66, m | 115.1, CH |
| 14 | 6.97, m | 129.2, CH |
| 15 | | 173.0, C |

NMR spectrum (500 MHz) for $^1$H, NMR spectrum (125 MHz) for $^{13}$C, DMSO-$d_6$, "m" means overlapped or multiple with other signals. Chemical shifts are reported in ppm.

HRMS (ESI, M+H$^+$) calculated for $C_{14}H_{16}NO_7^+$ 310.0921; found 310.0924.

$[\alpha]_D^{24.1}$ + 56° (*c* 0.1, MeOH)

**Supplementary Table 38.** Spectroscopic data of (2*S*,3*S*)-*t*-ES-a16

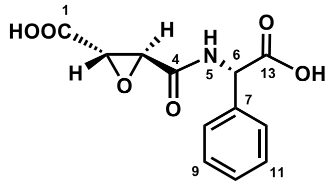

**(2*S*,3*S*)-*t*-ES-a16**

| Position | (2*S*,3*S*)-*t*-ES-a16 in DMSO-$d_6$ (500MHz) $\delta_H$ (*J* in Hz) | $\delta_C$, type |
|---|---|---|
| 1 | | 168.7, C |
| 2 | 3.79, d (1.9) | 52.3, CH |
| 3 | 3.50, d (1.9) | 51.3, CH |
| 4 | | 165.0, C |
| 5 | 9.26, d (7.3) | |
| 6 | 5.37, d (7.3) | 56.5, CH |
| 7 | | 136.7, C |
| 8 | 7.39, m | 127.6, CH |
| 9 | 7.39, m | 128.7, CH |
| 10 | 7.35, m | 128.2, CH |
| 11 | 7.39, m | 128.7, CH |
| 12 | 7.39, m | 127.6, CH |
| 13 | | 171.3, C |

NMR spectrum (500 MHz) for $^1$H, NMR spectrum (125 MHz) for $^{13}$C, DMSO-$d_6$, "m" means overlapped or multiple with other signals. Chemical shifts are reported in ppm.

HRMS (ESI, M+H$^+$) calculated for $C_{12}H_{12}NO_6^+$ 266.0659; found 266.0658.

$[\alpha]_D^{24.1}$ + 114° (*c* 0.1, MeOH)

**Supplementary Table 39.** Spectroscopic data of (2*S*,3*S*)-*t*-ES-a17

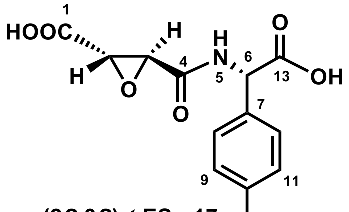

**(2*S*,3*S*)-*t*-ES-a17**

| | (2*S*,3*S*)-*t*-ES-a17 in DMSO-$d_6$ (500MHz) | |
|---|---|---|
| Position | $\delta_H$ (*J* in Hz) | $\delta_C$, type |
| 1 | | 168.8, C |
| 2 | 3.77, d (1.9) | 52.3, CH |
| 3 | 3.48, d (1.9) | 51.2, CH |
| 4 | | 164.8, C |
| 5 | 9.13, d (7.3) | |
| 6 | 5.21, d (7.3) | 56.0, CH |
| 7 | | 126.6, C |
| 8 | 7.19, m | 128.9, CH |
| 9 | 6.76, m | 115.4, CH |
| 10 | | 157.4, C |
| 11 | 6.76, m | 115.4, CH |
| 12 | 7.19, m | 128.9, CH |
| 13 | | 171.7, C |

NMR spectrum (500 MHz) for $^1$H, NMR spectrum (125 MHz) for $^{13}$C, DMSO-$d_6$, "m" means overlapped or multiple with other signals. Chemical shifts are reported in ppm.

HRMS (ESI, M+H$^+$) calculated for $C_{12}H_{12}NO_7^+$ 282.0608; found 282.0605.

$[\alpha]_D^{24.1}$ + 97° (*c* 0.1, MeOH)

**Supplementary Table 40.** Spectroscopic data of (2*S*,3*S*)-*t*-ES-a18

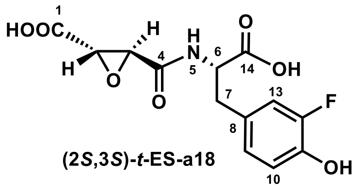

(2*S*,3*S*)-*t*-ES-a18

| | (2*S*,3*S*)-*t*-ES-a18 in DMSO-$d_6$ (500MHz) | |
|---|---|---|
| Position | $\delta_H$ (*J* in Hz) | $\delta_C$, type |
| 1 | | 168.6, C |
| 2 | 3.58, d (1.9) | 52.4, CH |
| 3 | 3.30, d (1.9) | 51.5, CH |
| 4 | | 165.3, C |
| 5 | 8.60, d (7.3) | |
| 6 | 4.42, m | 53.4, CH |
| 7 | 2.80, dd (9.5, 13.9); 2.98, dd (4.9, 13.9) | 35.5, $CH_2$ |
| 8 | | 128.5 d (6.07), C |
| 9 | 6.98, dd (1.8, 12.4) | 116.7 d (18.14), CH |
| 10 | 6.84, m | 117.4 d (3.22), CH |
| 11 | | 143.4 d (12.17), C |
| 12 | | 150.6 d (240.17), C |
| 13 | 6.84, m | 125.2 d (2.99), CH |
| 14 | | 172.3, C |

NMR spectrum (500 MHz) for $^1$H, NMR spectrum (125 MHz) for $^{13}$C, DMSO-$d_6$, "m" means overlapped or multiple with other signals. Chemical shifts are reported in ppm.

HRMS (ESI, M+H$^+$) calculated for $C_{13}H_{13}NO_7F^+$ 314.0671; found 314.0668.

$[\alpha]_D^{24.1}$ + 138° (*c* 0.1, MeOH)

**Supplementary Table 41.** Spectroscopic data of (2*S*,3*S*)-*t*-ES-a19

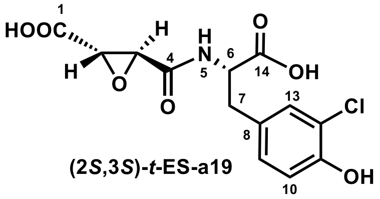

| | (2*S*,3*S*)-*t*-ES-a19 in DMSO-$d_6$ (500MHz) | |
|---|---|---|
| Position | $\delta_H$ (*J* in Hz) | $\delta_C$, type |
| 1 | | 169.0, C |
| 2 | 3.59, d (1.9) | 52.9, CH |
| 3 | 3.32, d (1.9) | 51.9, CH |
| 4 | | 165.7, C |
| 5 | 8.62, d (8.3) | |
| 6 | 4.41, m | 53.9, CH |
| 7 | 2.80, dd (9.4, 13.9); 2.98, dd (4.9, 13.9) | 35.7, $CH_2$ |
| 8 | | 129.4, C |
| 9 | 6.97, dd (2.1, 8.3) | 129.2, CH |
| 10 | 6.86, d (8.2) | 116.8, CH |
| 11 | | 152.1, C |
| 12 | | 119.6, C |
| 13 | 7.18, d (2.1) | 130.8, CH |
| 14 | | 172.7, C |

NMR spectrum (125 MHz) for $^{13}$C, DMSO-$d_6$, “m” means overlapped or multiple with other signals. Chemical shifts are reported in ppm.

HRMS (ESI, M+H$^+$) calculated for $C_{13}H_{13}NO_7Cl^+$ 330.0375; found 330.0376.

$[\alpha]_D^{24.1}$ + 130° (*c* 0.1, MeOH)

**Supplementary Table 42.** Spectroscopic data of (2*S*,3*S*)-*t*-ES-a20

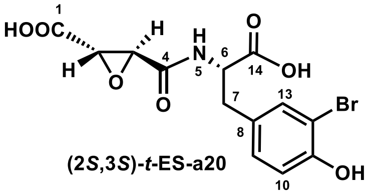

| Position | (2*S*,3*S*)-*t*-ES-a20 in DMSO-$d_6$ (500MHz) $\delta_H$ (*J* in Hz) | $\delta_C$, type |
|---|---|---|
| 1 | | 168.6, C |
| 2 | 3.59, d (1.8) | 52.5, CH |
| 3 | 3.31, d (1.8) | 51.5, CH |
| 4 | | 165.3, C |
| 5 | 8.61, d (8.3) | |
| 6 | 4.41, m | 53.5, CH |
| 7 | 2.80, dd (9.4, 13.9); 2.98, dd (4.9, 13.9) | 35.2, $CH_2$ |
| 8 | | 129.4, C |
| 9 | 7.02, dd (2.1, 8.2) | 129.4, CH |
| 10 | 6.85, d (8.2) | 116.1, CH |
| 11 | | 152.7, C |
| 12 | | 108.9, C |
| 13 | 7.33, d (2.1) | 133.3, CH |
| 14 | | 172.2, C |

NMR spectrum (500 MHz) for $^1H$, NMR spectrum (125 MHz) for $^{13}C$, DMSO-$d_6$, "m" means overlapped or multiple with other signals. Chemical shifts are reported in ppm.

HRMS (ESI, M+H$^+$) calculated for $C_{13}H_{13}NO_7Br^+$ 373.9870; found 373.9874.

$[\alpha]_D^{24.1}$ + 132 (*c* 0.1, MeOH)

**Supplementary Table 43.** Spectroscopic data of (2*S*,3*S*)-*t*-ES-a21

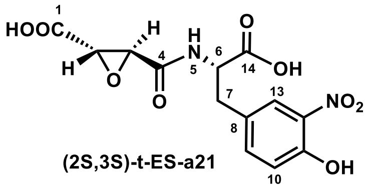

| Position | (2*S*,3*S*)-*t*-ES-a21 in DMSO-$d_6$ (500MHz) $\delta_H$ (*J* in Hz) | $\delta_C$, type |
|---|---|---|
| 1 | | 168.5, C |
| 2 | 3.58, d (1.9) | 52.5, CH |
| 3 | 3.32, d (1.9) | 51.4, CH |
| 4 | | 165.4, C |
| 5 | 8.66, d (8.4) | |
| 6 | 4.48, m | 53.1, CH |
| 7 | 2.89, dd (9.4, 13.9); 3.08, dd (4.9, 13.9) | 35.1, $CH_2$ |
| 8 | | 128.5, C |
| 9 | 7.41, dd (2.2, 8.6) | 136.2, CH |
| 10 | 7.06, d (8.5) | 119.0, CH |
| 11 | | 150.9, C |
| 12 | | 136.3, C |
| 13 | 7.76, d (2.2) | 125.5, CH |
| 14 | | 172.0, C |

NMR spectrum (500 MHz) for $^1$H, NMR spectrum (125 MHz) for $^{13}$C, DMSO-$d_6$, "m" means overlapped or multiple with other signals. Chemical shifts are reported in ppm.

HRMS (ESI, M+H$^+$) calculated for $C_{13}H_{13}N_2O_9^+$ 341.0616; found 341.0619.

$[\alpha]_D^{24.1}$ + 138 (*c* 0.1, MeOH)

**Supplementary Table 44.** Spectroscopic data of (2*S*,3*S*)-*t*-ES-a22

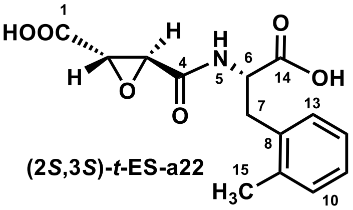

**(2*S*,3*S*)-*t*-ES-a22**

| | (2*S*,3*S*)-*t*-ES-a22 in DMSO-$d_6$ (500MHz) | |
|---|---|---|
| Position | $\delta_H$ (*J* in Hz) | $\delta_C$, type |
| 1 | | 168.5, C |
| 2 | 3.57, d (1.8) | 52.5, CH |
| 3 | 3.30, d (1.8) | 51.3, CH |
| 4 | | 165.3, C |
| 5 | 8.75, d (8.4) | |
| 6 | 4.48, m | 52.0, CH |
| 7 | 2.88, dd (9.9, 14.1);<br>3.14, dd (4.9, 14.2) | 34.2, $CH_2$ |
| 8 | | 136.1, C |
| 9 | | 135.5, C |
| 10 | 7.11, m | 130.1, CH |
| 11 | 7.11, m | 126.7, CH |
| 12 | 7.11, m | 125.6, CH |
| 13 | 7.12, m | 129.7, CH |
| 14 | | 172.5, C |
| 15 | 2.29, s | 18.9, $CH_3$ |

NMR spectrum (500 MHz) for $^1H$, NMR spectrum (125 MHz) for $^{13}C$, DMSO-$d_6$, "m" means overlapped or multiple with other signals. Chemical shifts are reported in ppm.

HRMS (ESI, M+H$^+$) calculated for $C_{14}H_{16}NO_6^+$ 294.0972; found 294.09812.

$[\alpha]_D^{24.1}$ + 90° (*c* 0.1, MeOH).

**Supplementary Table 45.** Spectroscopic data of (2*S*,3*S*)-*t*-ES-a23

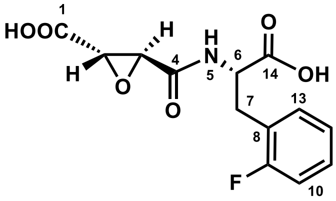

**(2*S*,3*S*)-*t*-ES-a23**

| Position | (2*S*,3*S*)-*t*-ES-a23 in DMSO-$d_6$ (500MHz) $\delta_H$ ($J$ in Hz) | $\delta_C$, type |
|---|---|---|
| 1 | | 168.5, C |
| 2 | 3.58, d (1.8) | 52.5, CH |
| 3 | 3.30, d (1.8) | 51.3, CH |
| 4 | | 165.3, C |
| 5 | 8.70, d (8.4) | |
| 6 | 4.53, m | 51.9, CH |
| 7 | 2.92, dd (9.5, 13.9); 3.19, m | 30.2, $CH_2$ |
| 8 | | 124.0, C, d (15.4) |
| 9 | | 160.8, C, d (244.2) |
| 10 | 7.14, m | 115.1, CH, d (21.6) |
| 11 | 7.29, m | 128.9, CH, d (8.1) |
| 12 | 7.13, m | 124.2, CH, d (3.3) |
| 13 | 7.29, m | 131.8, CH, d (4.5) |
| 14 | | 172.0, C |

NMR spectrum (500 MHz) for $^1H$, NMR spectrum (125 MHz) for $^{13}C$, DMSO-$d_6$, "m" means overlapped or multiple with other signals. Chemical shifts are reported in ppm.

HRMS (ESI, M+H$^+$) calculated for $C_{13}H_{13}NO_6F^+$ 298.0721; found 298.0723.

$[\alpha]_D^{24.1}$ + 74° ($c$ 0.1, MeOH).

**Supplementary Table 46.** Spectroscopic data of (2*S*,3*S*)-*t*-ES-a24

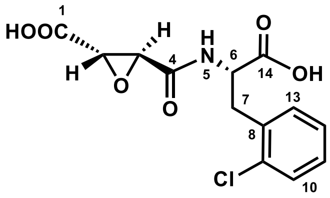

**(2*S*,3*S*)-*t*-ES-a24**

| | (2*S*,3*S*)-*t*-ES-a24 in DMSO-$d_6$ (500MHz) | |
|---|---|---|
| Position | $\delta_H$ (*J* in Hz) | $\delta_C$, type |
| 1 | | 168.5, C |
| 2 | 3.54, d (1.8) | 52.5, CH |
| 3 | 3.29, m | 51.3, CH |
| 4 | | 165.3, C |
| 5 | 8.73, d (8.6) | |
| 6 | 4.59, m | 51.4, CH |
| 7 | 2.98, dd (10.3, 13.9);<br>3.32, m | 34.5, $CH_2$ |
| 8 | | 134.9, C |
| 9 | | 133.3, C |
| 10 | 7.27, m | 127.0, CH |
| 11 | 7.27, m | 128.7, CH |
| 12 | 7.42, m | 129.3, CH |
| 13 | 7.33, m | 131.8, CH |
| 14 | | 172.1, C |

NMR spectrum (500 MHz) for $^1$H, NMR spectrum (125 MHz) for $^{13}$C, DMSO-$d_6$, "m" means overlapped or multiple with other signals. Chemical shifts are reported in ppm.

HRMS (ESI, M+H$^+$) calculated for $C_{13}H_{13}NO_6Cl^+$ 314.0426; found 314.0434.

$[\alpha]_D^{24.1}$ + 64 (*c* 0.1, MeOH).

**Supplementary Table 47.** Spectroscopic data of (2*S*,3*S*)-*t*-ES-a25

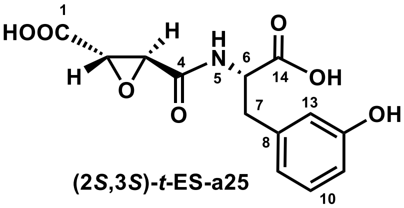

| Position | (2*S*,3*S*)-*t*-ES-a25 in DMSO-$d_6$ (500MHz) $\delta_H$ (*J* in Hz) | $\delta_C$, type |
|---|---|---|
| 1 | | 168.6, C |
| 2 | 3.61, d (1.8) | 52.5, CH |
| 3 | 3.32, d (1.8) | 51.3, CH |
| 4 | | 165.2, C |
| 5 | 8.65, d (8.2) | |
| 6 | 4.44, m | 53.4, CH |
| 7 | 2.83, dd (9.4, 13.9); 3.00, dd (4.8, 13.9) | 36.5, $CH_2$ |
| 8 | | 138.6, C |
| 9 | 7.00, m | 116.0, CH |
| 10 | 6.66, m | 129.2, CH |
| 11 | | 113.6, CH |
| 12 | 6.66, m | 157.2, C |
| 13 | 7.00, m | 119.7, CH |
| 14 | | 172.3, C |

NMR spectrum (500 MHz) for $^{1}$H, NMR spectrum (125 MHz) for $^{13}$C, DMSO-$d_6$, "m" means overlapped or multiple with other signals. Chemical shifts are reported in ppm.

HRMS (ESI, M+H$^+$) calculated for $C_{13}H_{14}NO_7^+$ 296.0765; found 296.0767.

$[\alpha]_D^{24.1}$ + 144° (*c* 0.1, MeOH).

**Supplementary Table 48.** Spectroscopic data of (2*S*,3*S*)-*t*-ES-a26

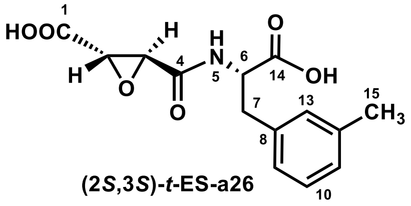

**(2*S*,3*S*)-*t*-ES-a26**

| | (2*S*,3*S*)-*t*-ES-a26 in DMSO-$d_6$ (500MHz) | |
|---|---|---|
| Position | $\delta_H$ (*J* in Hz) | $\delta_C$, type |
| 1 | | 168.6, C |
| 2 | 3.60, d (1.8) | 52.5, CH |
| 3 | 3.31, d (1.8) | 51.3, CH |
| 4 | | 165.2, C |
| 5 | 8.64, d (8.2) | |
| 6 | 4.46, m | 53.4, CH |
| 7 | 2.87, dd (9.5, 13.7); 3.06, dd (4.8, 13.8) | 36.4, $CH_2$ |
| 8 | | 137.2, C |
| 9 | 7.02, m | 126.2, CH |
| 10 | 7.17, t (7.7) | 128.2, CH |
| 11 | 7.02, m | 127.2, CH |
| 12 | | 137.2, C |
| 13 | 7.02, m | 129.8, CH |
| 14 | | 172.3, C |
| 15 | 2.27, s | 21.0, $CH_3$ |

NMR spectrum (500 MHz) for $^1$H, NMR spectrum (125 MHz) for $^{13}$C, DMSO-$d_6$, "m" means overlapped or multiple with other signals. Chemical shifts are reported in ppm.

HRMS (ESI, M+H$^+$) calculated for $C_{14}H_{16}NO_6^+$ 294.0972; found 294.0977.

$[\alpha]_D^{24.1}$ + 90° (*c* 0.1, MeOH).

**Supplementary Table 49.** Spectroscopic data of (2*S*,3*S*)-*t*-ES-a27

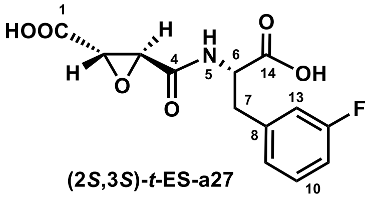

(2*S*,3*S*)-*t*-ES-a27

| | (2*S*,3*S*)-*t*-ES-a27 in DMSO-$d_6$ (500MHz) | |
|---|---|---|
| Position | $\delta_H$ (*J* in Hz) | $\delta_C$, type |
| 1 | | 168.5, C |
| 2 | 3.59, d (1.8) | 52.4, CH |
| 3 | 3.30, d (1.8) | 51.3, CH |
| 4 | | 165.2, C |
| 5 | 8.69, d (8.3) | |
| 6 | 4.51, m | 53.0, CH |
| 7 | 2.93, dd (9.7, 13.8); 3.13, dd (4.9, 13.8) | 36.1, $CH_2$ |
| 8 | | 140.2, C |
| 9 | 7.03, m | 125.3, CH |
| 10 | 7.32, m | 130.1, CH |
| 11 | 7.04, m | 113.4, CH |
| 12 | | 161.2, C |
| 13 | 7.03, m | 115.9, CH |
| 14 | | 172.1, C |

NMR spectrum (500 MHz) for $^1$H, NMR spectrum (125 MHz) for $^{13}$C, DMSO-$d_6$, "m" means overlapped or multiple with other signals. Chemical shifts are reported in ppm.

HRMS (ESI, M+H$^+$) calculated for $C_{13}H_{13}NO_6F^+$ 298.0721; found 298.0723.

$[\alpha]_D^{24.1}$ + 92° (*c* 0.1, MeOH).

**Supplementary Table 50.** Spectroscopic data of (2*S*,3*S*)-*t*-ES-a28

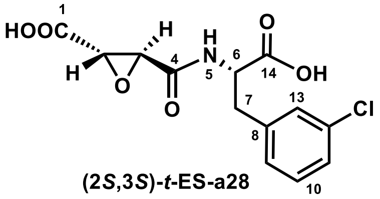

(2*S*,3*S*)-*t*-ES-a28

| | (2*S*,3*S*)-*t*-ES-a28 in DMSO-$d_6$ (500MHz) | |
|---|---|---|
| Position | $\delta_H$ (*J* in Hz) | $\delta_C$, type |
| 1 | | 168.5, C |
| 2 | 3.59, d (1.8) | 52.5, CH |
| 3 | 3.30, d (1.8) | 51.4, CH |
| 4 | | 165.3, C |
| 5 | 8.68, d (8.2) | |
| 6 | 4.46, m | 53.0, CH |
| 7 | 2.92, m; 3.11, m | 36.0, $CH_2$ |
| 8 | | 139.9, C |
| 9 | 7.30, m | 129.1, CH |
| 10 | 7.32, m | 130.1, CH |
| 11 | 7.29, m | 126.6, CH |
| 12 | | 132.8, C |
| 13 | 7.19, m | 128.0, CH |
| 14 | | 172.1, C |

NMR spectrum (500 MHz) for $^1H$, NMR spectrum (125 MHz) for $^{13}C$, DMSO-$d_6$, "m" means overlapped or multiple with other signals. Chemical shifts are reported in ppm.

HRMS (ESI, M+H$^+$) calculated for $C_{13}H_{13}NO_6Cl^+$ 314.0426; found 314.0434.

$[\alpha]_D^{24.1}$ + 70° (*c* 0.1, MeOH).

**Supplementary Table 51.** Spectroscopic data of (2*S*,3*S*)-*t*-ES-a29

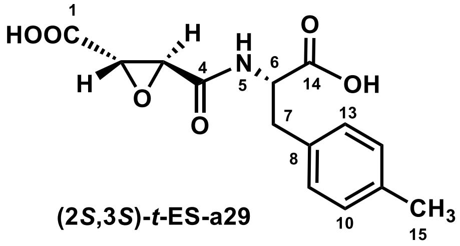

**(2*S*,3*S*)-*t*-ES-a29**

| | (2*S*,3*S*)-*t*-ES-a29 in DMSO-$d_6$ | |
|---|---|---|
| Position | $\delta_H$ (*J* in Hz) | $\delta_C$, type |
| 1 | | 168.6, C |
| 2 | 3.60, d (1.9) | 52.5, CH |
| 3 | 3.30, d (1.9) | 51.3, CH |
| 4 | | 165.2, C |
| 5 | 8.64, d (8.2) | |
| 6 | 4.46, m | 53.5, CH |
| 7 | 2.87, m; 3.05, m | 36.1, $CH_2$ |
| 8 | | 134, C |
| 9 | 7.10, m | 129, CH |
| 10 | 7.10, m | 129, CH |
| 11 | | 134.4, C |
| 12 | 7.10, m | 128.9, CH |
| 13 | 7.10, m | 128.9, CH |
| 14 | | 172.4, C |
| 15 | 2.26, s | 20.7, $CH_3$ |

NMR spectrum (500 MHz) for $^1$H, NMR spectrum (125 MHz) for $^{13}$C, DMSO-$d_6$, "m" means overlapped or multiple with other signals. Chemical shifts are reported in ppm.

HRMS (ESI, M+H$^+$) calculated for $C_{14}H_{16}NO_6^+$ 294.0972; found 294.0981.

$[\alpha]_D^{24.1}$ + 109° (*c* 0.1, MeOH)

**Supplementary Table 52.** Spectroscopic data of (2*S*,3*S*)-*t*-ES-a30

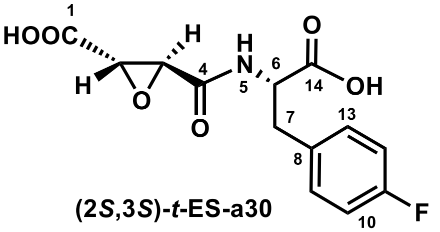

(2*S*,3*S*)-*t*-ES-a30

| | (2*S*,3*S*)-*t*-ES-a30 in DMSO-$d_6$ (500MHz) | |
|---|---|---|
| Position | $\delta_H$ (*J* in Hz) | $\delta_C$, type |
| 1 | | 168.5, C |
| 2 | 3.59, d (1.8) | 52.5, CH |
| 3 | 3.30, d (1.8) | 51.3, CH |
| 4 | | 165.2, C |
| 5 | 8.67, d (8.3) | |
| 6 | 4.48, m | 53.7, CH |
| 7 | 2.90, dd (9.6, 13.8); 3.09, dd (4.9, 13.9) | 35.8, $CH_2$ |
| 8 | | 133.5, C |
| 9 | 7.26, m | 131.0 d (8.06), CH |
| 10 | 7.11, m | 115.0 d (21.06), CH |
| 11 | | 161.1 d (242.13), C |
| 12 | 7.11, m | 115.0 d (21.06), CH |
| 13 | 7.26, m | 131.0 d (8.06), CH |
| 14 | | 172.2, C |

NMR spectrum (500 MHz) for $^1$H, NMR spectrum (125 MHz) for $^{13}$C, DMSO-$d_6$, "m" means overlapped or multiple with other signals. Chemical shifts are reported in ppm.

HRMS (ESI, M+H$^+$) calculated for $C_{13}H_{13}NO_6F^+$ 298.0721; found 298.0723.

$[\alpha]_D^{24.1}$ + 112° (*c* 0.1, MeOH).

**Supplementary Table 53.** Spectroscopic data of (2*S*,3*S*)-*t*-ES-a31

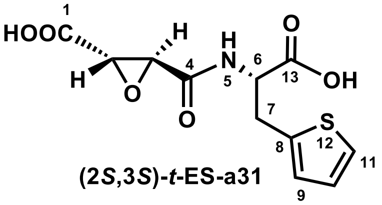

**(2*S*,3*S*)-*t*-ES-a31**

| Position | (2*S*,3*S*)-*t*-ES-a31 in DMSO-$d_6$ (500MHz) $\delta_H$ (*J* in Hz) | $\delta_C$, type |
|---|---|---|
| 1 | | 168.6, C |
| 2 | 3.66, d (1.9) | 52.5, CH |
| 3 | 3.38, d (1.9) | 51.7, CH |
| 4 | | 165.6, C |
| 5 | 8.70, d (8.1) | |
| 6 | 4.48, dd (4.6, 8.5) | 53.4, CH |
| 7 | 3.19, m; 3.32, m | 30.8, $CH_2$ |
| 8 | | 139.0, C |
| 9 | 6.90, d (3.4) | 126.5, CH |
| 10 | 6.95, dd (3.4, 5.2) | 126.8, CH |
| 11 | 7.36, d (5.1) | 124.9, CH |
| 13 | | 171.8, C |

NMR spectrum (500 MHz) for $^1$H, NMR spectrum (125 MHz) for $^{13}$C, DMSO-$d_6$, “m” means overlapped or multiple with other signals. Chemical shifts are reported in ppm.

HRMS (ESI, M+H$^+$) calculated for $C_{11}H_{12}NO_6S^+$ 286.0380; found 286.0382.

$[\alpha]_D^{24.1}$ + 50° (*c* 0.1, MeOH).

**Supplementary Table 54.** Spectroscopic data of (2*S*,3*S*)-*t*-ES-a32

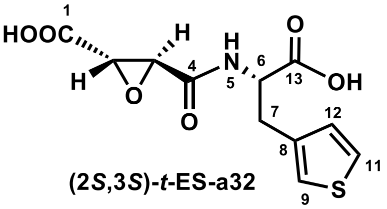

(2*S*,3*S*)-*t*-ES-a32

| | (2*S*,3*S*)-*t*-ES-a32 in DMSO-$d_6$ (500MHz) | |
|---|---|---|
| Position | $\delta_H$ (*J* in Hz) | $\delta_C$, type |
| 1 | | 168.6, C |
| 2 | 3.62, d (1.8) | 52.5, CH |
| 3 | 3.32, d (1.8) | 51.4, CH |
| 4 | | 165.3, C |
| 5 | 8.65, d (8.1) | |
| 6 | 4.48, dd (4.7, 8.1, 9.1) | 52.9, CH |
| 7 | 2.98, dd (4.7, 14.5); 3.10, dd (9.3, 14.4) | 31.1, $CH_2$ |
| 8 | | 137.4, C |
| 9 | 7.23, dd (1.2, 3.0) | 122.6, CH |
| 11 | 7.45, dd (3.0, 4.9) | 125.9, CH |
| 12 | 6.99, dd (1.3, 4.9) | 128.6, CH |
| 13 | | 172.3, C |

NMR spectrum (500 MHz) for $^1$H, NMR spectrum (125 MHz) for $^{13}$C, DMSO-$d_6$, "m" means overlapped or multiple with other signals. Chemical shifts are reported in ppm.

HRMS (ESI, M+H$^+$) calculated for $C_{11}H_{12}NO_6S^+$ 286.0380; found 286.0378.

$[\alpha]_D^{24.1}$ + 54° (*c* 0.1, MeOH).

**Supplementary Table 55.** Spectroscopic data of **(2*S*,3*S*)-*t*-ES-a9-b7**

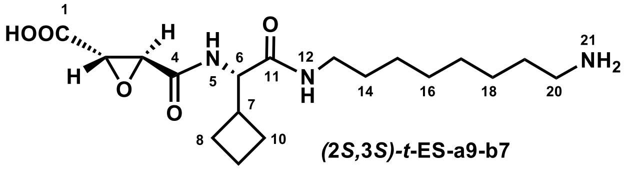

| | **(2*S*,3*S*)-*t*-ES-a9-b7** in DMSO-$d_6$ (500MHz) | |
|---|---|---|
| Position | $\delta_H$ ($J$ in Hz) | $\delta_C$, type |
| 1 | | 168.8, C |
| 2 | 3.73, d (1.8) | 52.5, CH |
| 3 | 3.44, d (1.8) | 51.3, CH |
| 4 | | 165.1, C |
| 5 | 8.53, d (8.5) | |
| 6 | 4.27, t (8.5) | 56.8, CH |
| 7 | 2.53, m | 37.4, CH |
| 8 | 1.73, m; 1.88, m | 24.7, $CH_2$ |
| 9 | 1.75, m | 17.5, $CH_2$ |
| 10 | 1.82, m | 24.3, $CH_2$ |
| 11 | | 169.5, C |
| 12 | 8.04, t (5.7) | |
| 13 | 2.96, m; 3.08, m | 38.3, $CH_2$ |
| 14 | 1.35, m | 29.0, $CH_2$ |
| 15 | 1.23, m | 26.1, $CH_2$ |
| 16 | 1.23, m | 28.5, $CH_2$ |
| 17 | 1.23, m | 25.7, $CH_2$ |
| 18 | 1.23, m | 28.5, $CH_2$ |
| 19 | 1.49, m | 27.0, $CH_2$ |
| 20 | 2.75, m | 38.8, $CH_2$ |

NMR spectrum (500 MHz) for $^1$H, NMR spectrum (125 MHz) for $^{13}$C, DMSO-$d_6$, "m" means overlapped or multiple with other signals. Chemical shifts are reported in ppm.

HRMS (ESI, M+H$^+$) calculated for $C_{18}H_{32}N_3O_5^+$ 370.2336; found 370.2332.

$[\alpha]_D^{24.1}$ + 36° (*c* 0.1, MeOH).

**Supplementary Table 56.** Spectroscopic data of **(2S,3S)-*t*-ES-a9-b12**

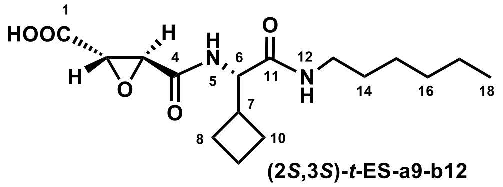

| | **(2*S*,3*S*)-*t*-ES-a9-b12** in DMSO-$d_6$ (500MHz) | |
|---|---|---|
| Position | $\delta_H$ ($J$ in Hz) | $\delta_C$, type |
| 1 | | 168.8, C |
| 2 | 3.74, d (1.8) | 52.5, CH |
| 3 | 3.45, d (1.8) | 51.2, CH |
| 4 | | 165.0, C |
| 5 | 8.53, d (8.4) | |
| 6 | 4.28, t (8.5) | 56.7, CH |
| 7 | 2.54, m | 37.5, CH |
| 8 | 1.75, m; 1.83, m | 24.7, $CH_2$ |
| 9 | 1.73, m | 17.4, $CH_2$ |
| 10 | 1.82, m | 24.3, $CH_2$ |
| 11 | | 169.5, C |
| 12 | 8.04, t (5.7) | |
| 13 | 2.95, m; 3.09, m | 38.3, $CH_2$ |
| 14 | 1.35, m | 29.0, $CH_2$ |
| 15 | 1.23, m | 25.9, $CH_2$ |
| 16 | 1.23, m | 30.9, $CH_2$ |
| 17 | 1.23, m | 22.1, $CH_2$ |
| 18 | 0.85, t (6.8) | 13.9, $CH_3$ |

NMR spectrum (500 MHz) for $^1H$, NMR spectrum (125 MHz) for $^{13}C$, DMSO-$d_6$, "m" means overlapped or multiple with other signals. Chemical shifts are reported in ppm.

HRMS (ESI, M+H$^+$) calculated for $C_{16}H_{27}N_2O_5^+$ 327.1914; found 327.1912.

$[\alpha]_D^{24.1}$ + 36° ($c$ 0.1, MeOH).

**Supplementary Table 57.** Spectroscopic data of **(2*S*,3*S*)-*t*-ES-a10-b9**

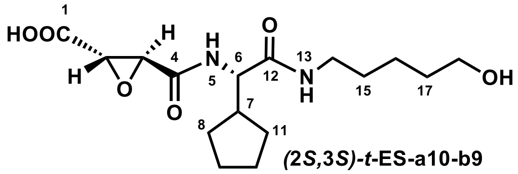

| | **(2*S*,3*S*)-*t*-ES-a10-b9** in DMSO-$d_6$ (500MHz) | |
|---|---|---|
| Position | $\delta_H$ (*J* in Hz) | $\delta_C$, type |
| 1 | | 168.8, C |
| 2 | 3.70, d (1.8) | 52.1, CH |
| 3 | 3.42, d (1.8) | 51.2, CH |
| 4 | | 164.9, C |
| 6 | 4.16, t (8.7) | 56.1, CH |
| 7 | 2.12, m | 42.2, CH |
| 8 | 1.26, m | 28.6, $CH_2$ |
| 9 | 1.54, m | 24.8, $CH_2$ |
| 10 | 1.45, m | 24.5, $CH_2$ |
| 11 | 1.51, m; 1.61, m | 28.6, $CH_2$ |
| 12 | | 170.3, C |
| 14 | 2.95, dt (6.9, 13.5); 3.09, dt (6.9, 13.5) | 38.3, $CH_2$ |
| 15 | 1.36, m | 28.8, $CH_2$ |
| 16 | 1.23, m | 22.9, $CH_2$ |
| 17 | 1.39, m | 32.1, $CH_2$ |
| 18 | 1.39, dd (8.7, 15.2) | 60.6, $CH_2$ |

NMR spectrum (500 MHz) for $^1$H, NMR spectrum (125 MHz) for $^{13}$C, DMSO-$d_6$, “m” means overlapped or multiple with other signals. Chemical shifts are reported in ppm.

HRMS (ESI, M+H$^+$) calculated for $C_{16}H_{27}N_2O_6^+$ 343.1864; found 343.1807.

$[\alpha]_D^{24.1}$ + 56° (*c* 0.1, MeOH).

**Supplementary Table 58.** Spectroscopic data of **(2*S*,3*S*)-*t*-ES-a10-b13**

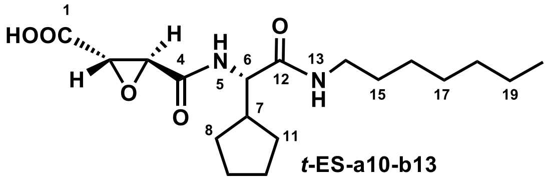

***t*-ES-a10-b13**

| | **(2*S*,3*S*)-*t*-ES-a10-b13** in acetone-$d_6$ (500MHz) | |
|---|---|---|
| Position | $\delta_H$ (*J* in Hz) | $\delta_C$, type |
| 1 | | 168.8, C |
| 2 | 3.69, d (1.8) | 54.2, CH |
| 3 | 3.59, d (1.8) | 52.6, CH |
| 4 | | 166.0, C |
| 5 | 7.47, d (8.7) | |
| 6 | 4.30, t (8.5) | 57.2, CH |
| 7 | 2.25, m | 43.9, CH |
| 8 | 1.37, m | 29.9, $CH_2$ |
| 9 | 1.50, m | 25.6, $CH_2$ |
| 10 | 1.60, m | 25.8 $CH_2$ |
| 11 | 1.63, m | 30.1, $CH_2$ |
| 12 | | 171.4, C |
| 13 | 7.37, t (5.8) | |
| 14 | 3.13, dt (6.4, 13.0); 3.25, dt (6.6, 13.2) | 39.8, $CH_2$ |
| 15 | 1.49, m | 30.2, $CH_2$ |
| 16 | 1.27, m | 32.5, $CH_2$ |
| 17 | 1.70, m | 29.3, $CH_2$ |
| 18 | 1.30, m | 27.5, $CH_2$ |
| 19 | 1.28, m | 23.2, $CH_2$ |
| 20 | 0.89, m | 14.3, $CH_3$ |

NMR spectrum (500 MHz) for $^1$H, NMR spectrum (125 MHz) for $^{13}$C, acetone-$d_6$, "m" means overlapped or multiple with other signals. Chemical shifts are reported in ppm.

HRMS (ESI, M+H$^+$) calculated for $C_{18}H_{31}N_2O_5^+$ 355.2227; found 355.2226.

$[\alpha]_D^{24.1}$ + 56° (*c* 0.1, MeOH).

**Supplementary Table 59.** Spectroscopic data of **(2*S*,3*S*)-*t*-ES-a10-b26**

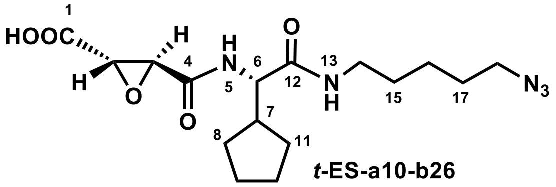

| | **(2*S*,3*S*)-*t*-ES-a10-b26** in acetone-$d_6$ (500MHz) | |
|---|---|---|
| Position | $\delta_H$ ($J$ in Hz) | $\delta_C$, type |
| 1 | | 168.7, C |
| 2 | 3.69, d (1.8) | 54.2, CH |
| 3 | 3.59, d (1.8) | 52.8, CH |
| 4 | | 166.1, C |
| 5 | 7.45, m | |
| 6 | 4.29, t (8.5) | 57.2, CH |
| 7 | 2.26, m | 43.8, CH |
| 8 | 1.32, m; 1.52, m | 30.2, $CH_2$ |
| 9 | 1.49, m | 25.6, $CH_2$ |
| 10 | 1.56, m | 25.6, $CH_2$ |
| 11 | 1.69, m | 29.3, $CH_2$ |
| 12 | | 171.4, C |
| 13 | 7.45, m | |
| 14 | 3.16, m; 3.26, m | 39.5, $CH_2$ |
| 15 | 1.51, m | 25.9, $CH_2$ |
| 16 | 1.40, m | 24.7, $CH_2$ |
| 17 | 1.51, m; 1.60, m | 29.1, $CH_2$ |
| 18 | 3.33, m | 51.9, $CH_2$ |

NMR spectrum (500 MHz) for $^1$H, NMR spectrum (125 MHz) for $^{13}$C, acetone-$d_6$, "m" means overlapped or multiple with other signals. Chemical shifts are reported in ppm.

HRMS (ESI, M+H$^+$) calculated for $C_{16}H_{26}N_5O_5^+$ 368.1928; found 368.1929.

$[\alpha]_D^{24.1}$ + 42° ($c$ 0.1, MeOH).

**Supplementary Table 60.** Spectroscopic data of **(2*S*,3*S*)-*t*-ES-Leu-b43**

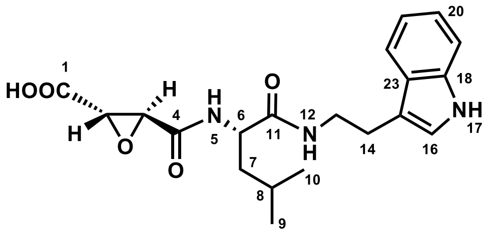

**(2S,3S)-*t*-ES-Leu-b43**

| Position | **(2*S*,3*S*)-*t*-ES-Leu-b43** in DMSO-$d_6$ (500MHz) | |
|---|---|---|
| | $\delta_H$ (*J* in Hz) | $\delta_C$, type |
| 1 | | 168.8, C |
| 2 | 3.67, d (1.8) | 52.7, CH |
| 3 | 3.48, d (1.8) | 51.2, CH |
| 4 | | 164.9, C |
| 5 | 8.58, d (8.4) | |
| 6 | 4.32, m | 51.2, CH |
| 7 | 1.44, m | 41.1, $CH_2$ |
| 8 | 1.52, m | 24.2, CH |
| 9 | 0.83, d (6.5) | 21.6, $CH_3$ |
| 10 | 0.87, d (6.5) | 22.9, $CH_3$ |
| 11 | | 171.1, C |
| 12 | 8.20, t (5.7) | |
| 13 | 3.31, m | 40.1, $CH_2$ |
| 14 | 2.81, m | 25.0, $CH_2$ |
| 15 | | 111.6, C |
| 16 | 7.13, d (2.3) | 122.7, CH |
| 17 | 10.81, m | |
| 18 | | 136.2, C |
| 19 | 7.32, d (8.1) | 111.3, CH |
| 20 | 7.05, t (7.5) | 120.9, CH |
| 21 | 6.97, t (7.4) | 118.2, CH |
| 22 | 7.53, d (7.8) | 118.2, CH |
| 23 | | 127.2, C |

NMR spectrum (500 MHz) for $^1H$, NMR spectrum (125 MHz) for $^{13}C$, DMSO-$d_6$, "m" means overlapped or multiple with other signals. Chemical shifts are reported in ppm.

HRMS (ESI, M+H$^+$) calculated for $C_{20}H_{26}N_3O_5^+$ 388.1867; found 388.1864.

$[\alpha]_D^{24.1}$ + 48° (*c* 0.1, MeOH).

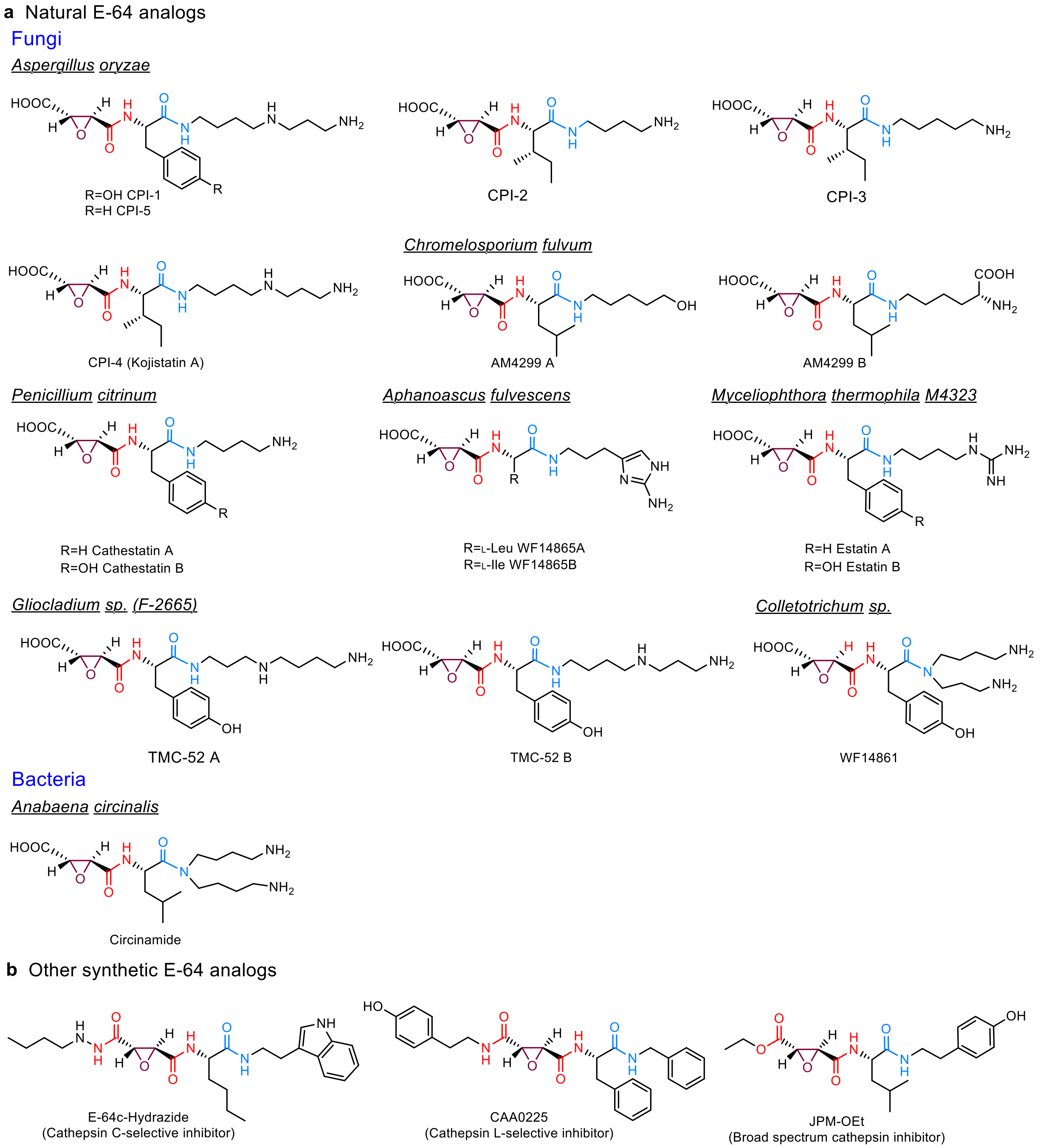

**Supplementary Fig. 1 | Natural and synthetic E-64 analogs**. **a**, Natural E-64 analogs, including CP1-1 to CPI-5[7], AM4299A and B[8], cathestatin A and B[9], estatin A and B[10], WF14865A and B[11], TMC-52 A and B[12], were isolated, from different fungal species, such as *Aspergillus*, *Penicillium*, *Aphanoascus*, *Myceliophthora*, *Gliocladium*, *Colletotrichum spp*. E-64 analog circinamide[13] was isolated from cyanobacteria *Anabaena circinalis*. Most of E-64 analogs contain putrescine and cadaverine or its derivatives at C-terminus. **b**, Other synthetic cysteine protease inhibitors derived from E-64[14].

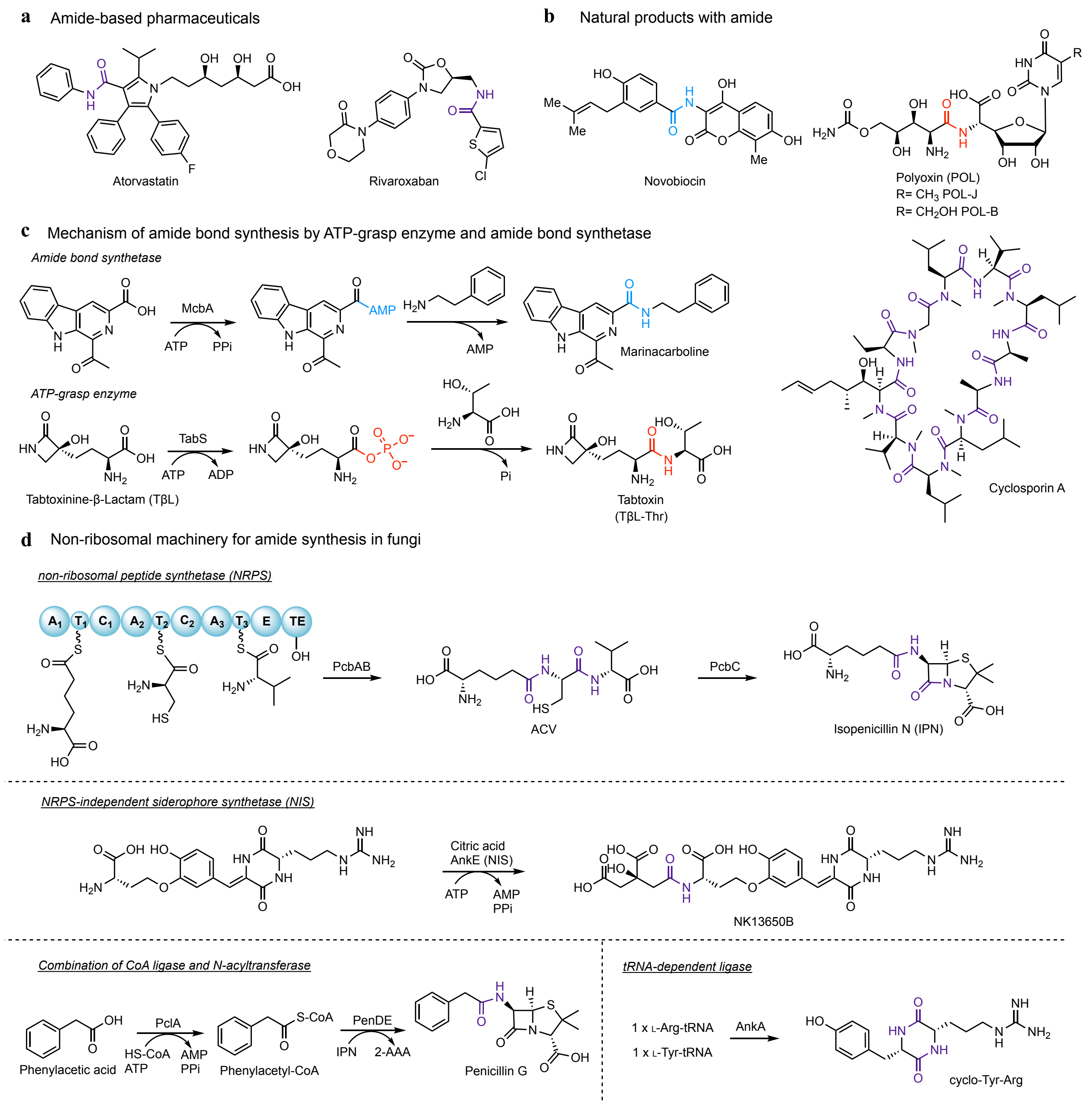

**Supplementary Fig. 2 | Significance and biosynthetic machinery of amide functionality**. **a**, Selected top-selling drugs containing amide bond. **b**, Representative amide-containing natural products used in clinic and agriculture. **c**, The mechanisms of the biocatalytically competent amide bond synthetase McbA and the ATP-grasp enzyme TabS. **d**, Examples of non-ribosomal machinery for amide synthesis in fungi. The amide bond formations in isopenicillin N (IPN) is catalyzed by three modules NRPS PcbAB. NRPS-independent siderophore synthetase (NIS) AnkE is proposed to catalyze the amide bond in NK13650B. The pair of CoA ligase PclA and *N*-acyltransferase PenDE forms the amide bond in the production of penicillin G and 2-aminoadipic acid (2-AAA). AnkA catalyzed the tRNAdependent amidation to form cyclo-Tyr-Arg. Domain abbreviations: A: adenylation; T: peptidyl-carrier protein; C: condensation; E: epimerization; TE: thioesterase. While a few ATP-grasp enzymes were proposed to involve the biosynthesis of the fungal peptides (also see **Extended Data Fig. 2**), ABSs have not been identified in the fungal natural product biosynthesis.

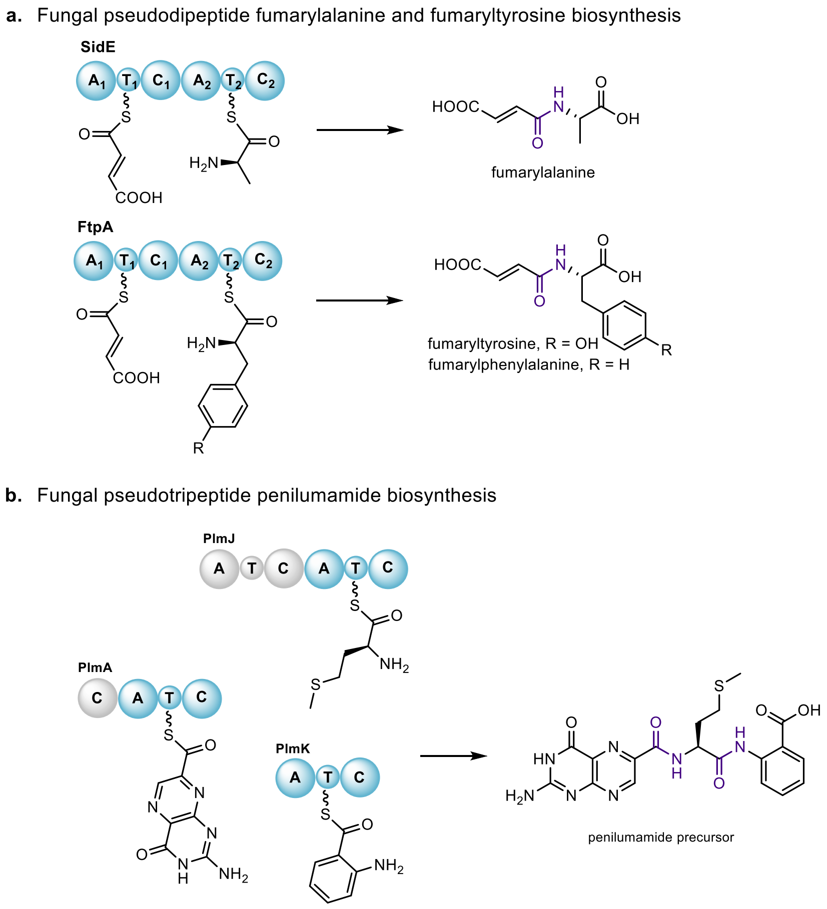

**Supplementary Fig. 3 | Amide bond formation in fumaryl dipeptides[15-16] and penilumamide[17] is derived from NRPS. a**, Dimodule NRPS, SidE and FtpA were responsible for biosynthesis of fumaryl pseudodipeptide fumarylalanine and fumaryltyrosine, respectively. **b**, In penilumamide precursor biosynthesis, NRPS, PlmA and PlmJ and PlmK were required for penilumamide biosynthesis. Domain highlighted in grey was proposed to be either skipped or inactive.

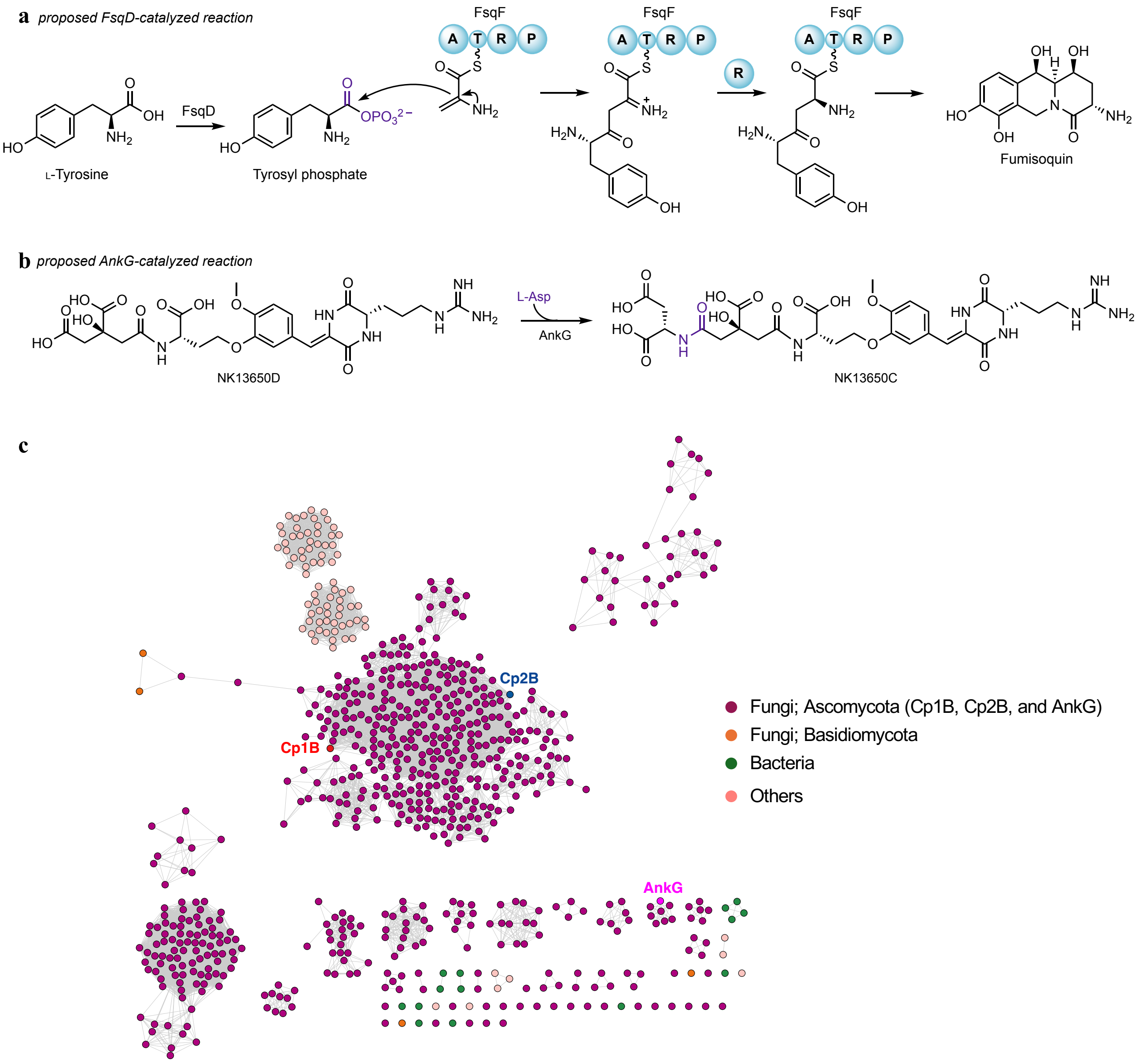

**Supplementary Fig. 4 | Limited examples of fungal ATP-grasp enzyme involved in natural product biosynthesis**. **a**, FsqD from fumisoquin biosynthesis was proposed to activate L-tyrosine to form tyrosyl phosphate. However, the functions of FsqD have not been biochemically characterized. **b**, AnkG was proposed to be an ATP-grasp enzyme that catalyzes the amide bond formation between L-aspartic acid and NK13650D to form NK13650C based on the in vivo experiments. However, no biochemical characterization was reported. **c**, Sequence similarity network analysis of Cp1B homologs from the UniProt database. Note that 5,000 maximum number target sequences from a blastp search with Cp1B as a query (expect threshold value: 5) were retrieved and subjected to SSN construction, with an alignment score threshold of 7. Cp1B and Cp2B in this study and the proposed ATP-grasp enzyme AnkG are highlighted. While Cp1B and Cp2B were located at a separate clade from AnkG, the SSN and the amino acid sequence identity between them (~30%) suggest these enzymes are distinct but distantly related, suggesting that AnkG also functions as a ATP-grasp enzyme as previously proposed. While no characterized enzymes were found in this SSN, interestingly, the SSN showed that putative Cp1B-like ATP-grasp enzymes are conserved in not only many fungi (Ascomycota and Basidiomycota) but also a few bacteria. Domain abbreviations: A: adenylation; T: peptidyl-carrier protein; R: reductase domain; P: pyridoxal phosphate binding domain.

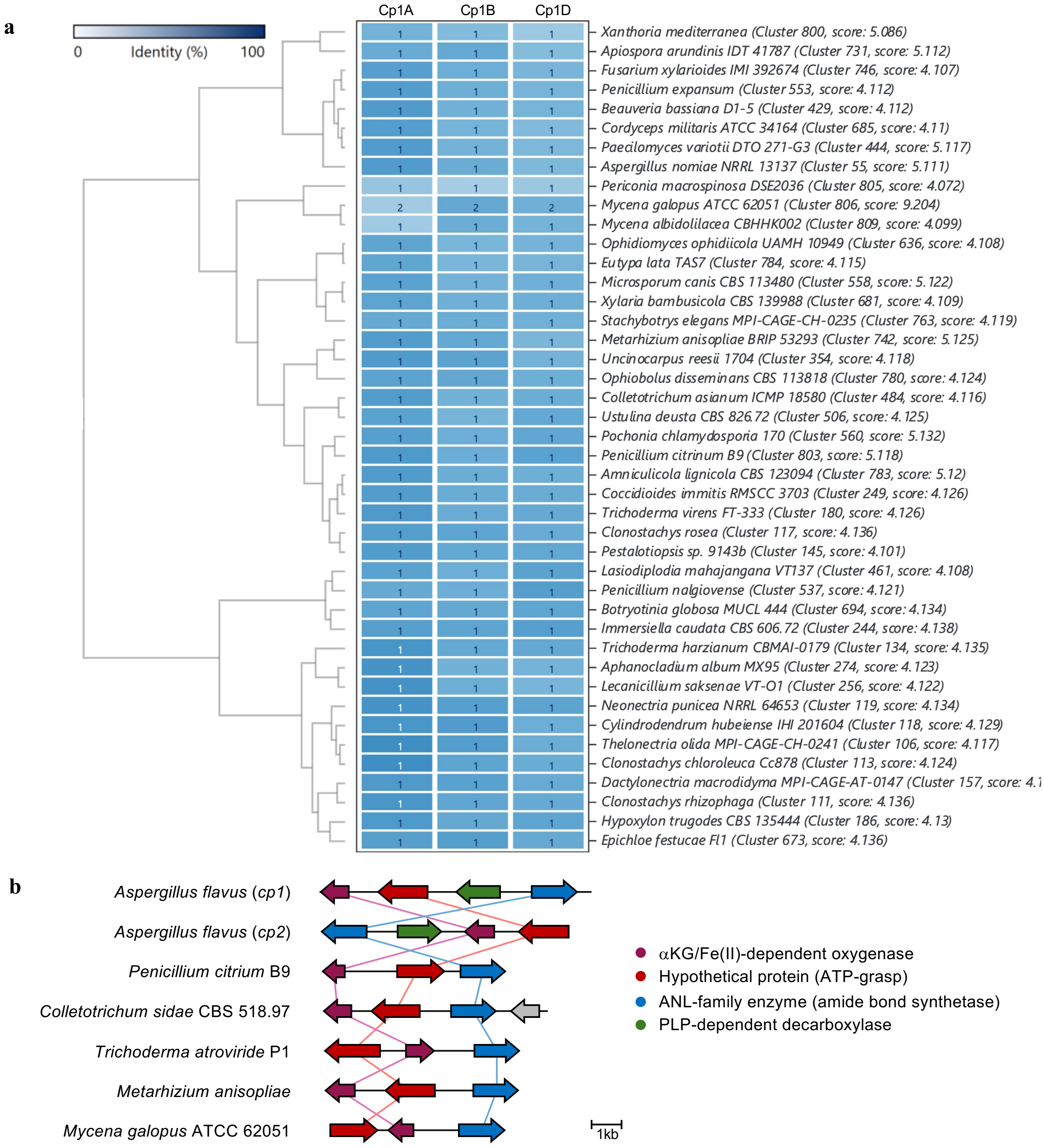

**Supplementary Fig. 5 | E-64-like biosynthetic gene clusters are widely conserved in a plethora of fungi. a**, Selected hit clusters are shown in a dendrogram (based on identity to query sequences) from cblaster search. A darker tint of blue resembles a higher percentage identity of the query in the output cluster. Three gene cassettes (*cpA*, *cpB*, and *cpD*) is highly conserved in more than > 40 different fungal genera such as *Aspergillus spp*, *Penicillium spp*, *Metarhizium spp*, *Trichoderma spp*, and *Mycena* spp (Basidiomycota). Two copies of E-64 like cluster are also present in *Mycena galopus*. **b**. Selected E-64 honologous biosynthetic gene clusters from clinker visualization, including reported E-64 analog producing fungi (*Aspergillus oryzae*, *Penicillium citrinum*, and *Colletotrichium spp*) and fungi not known to produce E-64 (*Trichoderma atroviride*, *Metarhizium anisopliae*, and *Mycena galopus* ATCC 62051). Nearly all the genes in those clusters are conserved except for PLP-dependent decarboxylase (the homolog of Cp1C).

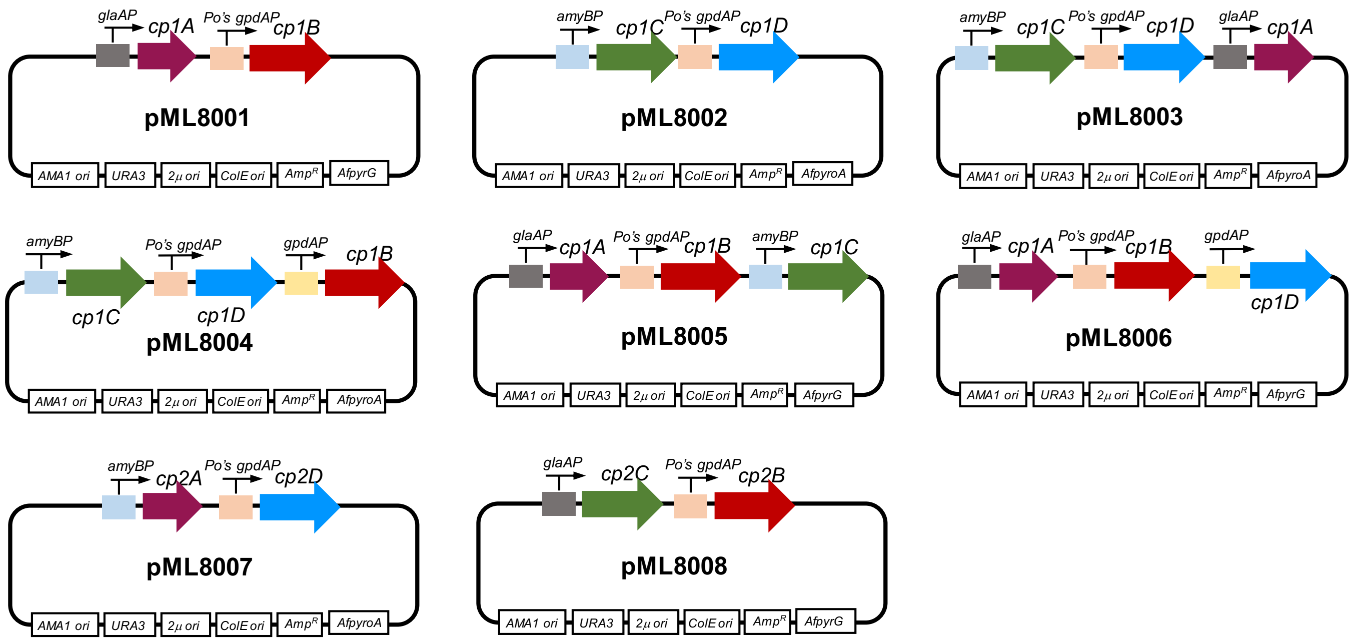

**Supplementary Fig. 6 | Plasmids used for heterologous expression of *cp1* and *cp2* genes in *A. nidulans* heterologous host**.

**a** domain search from Pfam database

**b** domain search from COG database

**c** BLASTP against SwissProt database

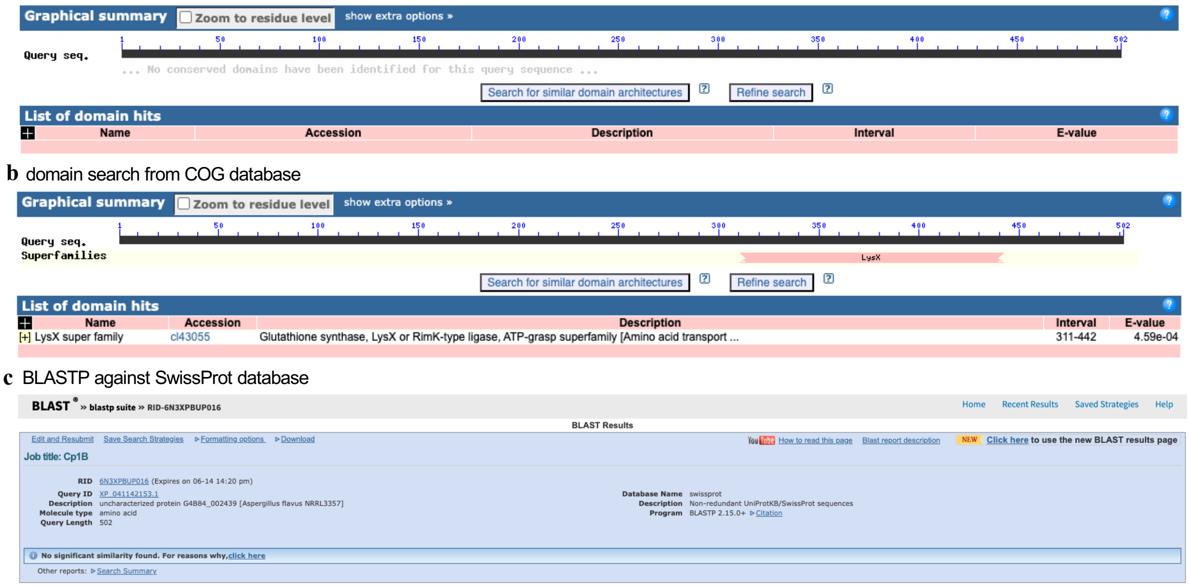

**d** TOP PDB hit from Protein structure search by Foldseek

| Target | Description | Scientific Name | Prob. | Seq. Id. | E-Value |
|---|---|---|---|---|---|
| 3kal-assembly1_B | Structure of homoglutathione synthetase from Gl… | Glycine max | 1.00 | 12.4 | 9.20e-12 |
| 2hgs-assembly1_A-2 | HUMAN GLUTATHIONE SYNTHETASE | Homo sapiens | 1.00 | 15.8 | 2.67e-12 |
| 1m0w-assembly1_A | Yeast Glutathione Synthase Bound to gamma-gl… | Saccharomyces cerevisiae | 1.00 | 13.4 | 3.45e-12 |
| 5oes-assembly3_E | The structure of a glutathione synthetase (StGS… | Solanum tuberosum | 1.00 | 12.8 | 7.11e-12 |
| 3kal-assembly1_A | Structure of homoglutathione synthetase from Gl… | Glycine max | 1.00 | 12.8 | 1.54e-11 |
| 5oeu-assembly1_A | The structure of a glutathione synthetase like-eff… | Globodera pallida | 1.00 | 11.9 | 4.20e-10 |
| 5oet-assembly1_B | The structure of a glutathione synthetase like-eff… | Globodera pallida | 1.00 | 13.3 | 3.79e-10 |
| 5oev-assembly1_A | The structure of a glutathione synthetase like-eff… | Globodera pallida | 1.00 | 11.3 | 6.04e-10 |
| 7uka-assembly1_A | YgiC from Escherichia coli K-12 in complex with … | Escherichia coli K-12 | 1.00 | 13.2 | 5.99e-8 |

**e** Predicted structure of Cp1B

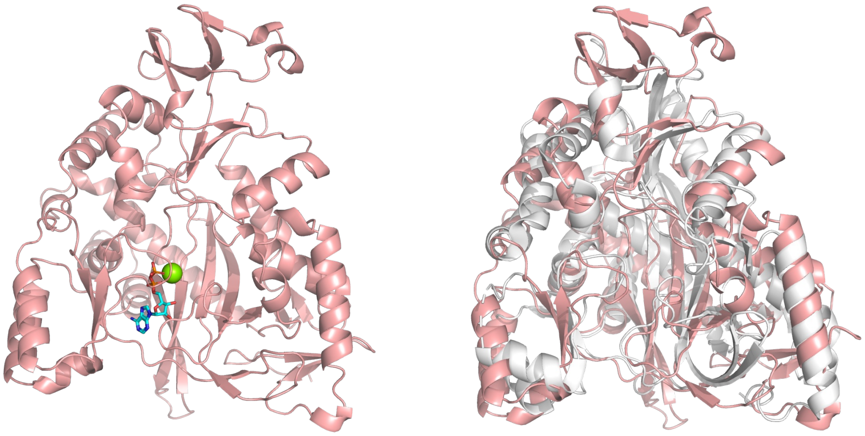

**Supplementary Fig. 7 | Bioinformatic analysis of Cp1B. a**, Pfam domain search for Cp1B on NCBI website. **b**, COG domain search for Cp1B on NCBI website. The small portion of Cp1B has a LysX superfamily domain with high E-value ($4.59 \times 10^{-4}$). **c**, BLASTP search using Cp1B as the query against SwissProt database. No characterized homolog of Cp1B was identified from the database. **d**, Structural analysis of Cp1B/Cp2B by Foldseek[18] identified homoglutathione synthase (3KAL, PDB number)[19] from *Glycine max* as closet structure homolog. **e**. Left, AF3[20] predicted Cp1B with ADP and $Mg^{2+}$. Right, overlay of Cp1B and 3KAL, shown in cartoon representation, RMSD = 5.697 (2147 to 2147 atoms by using "super" align method in pymol ).

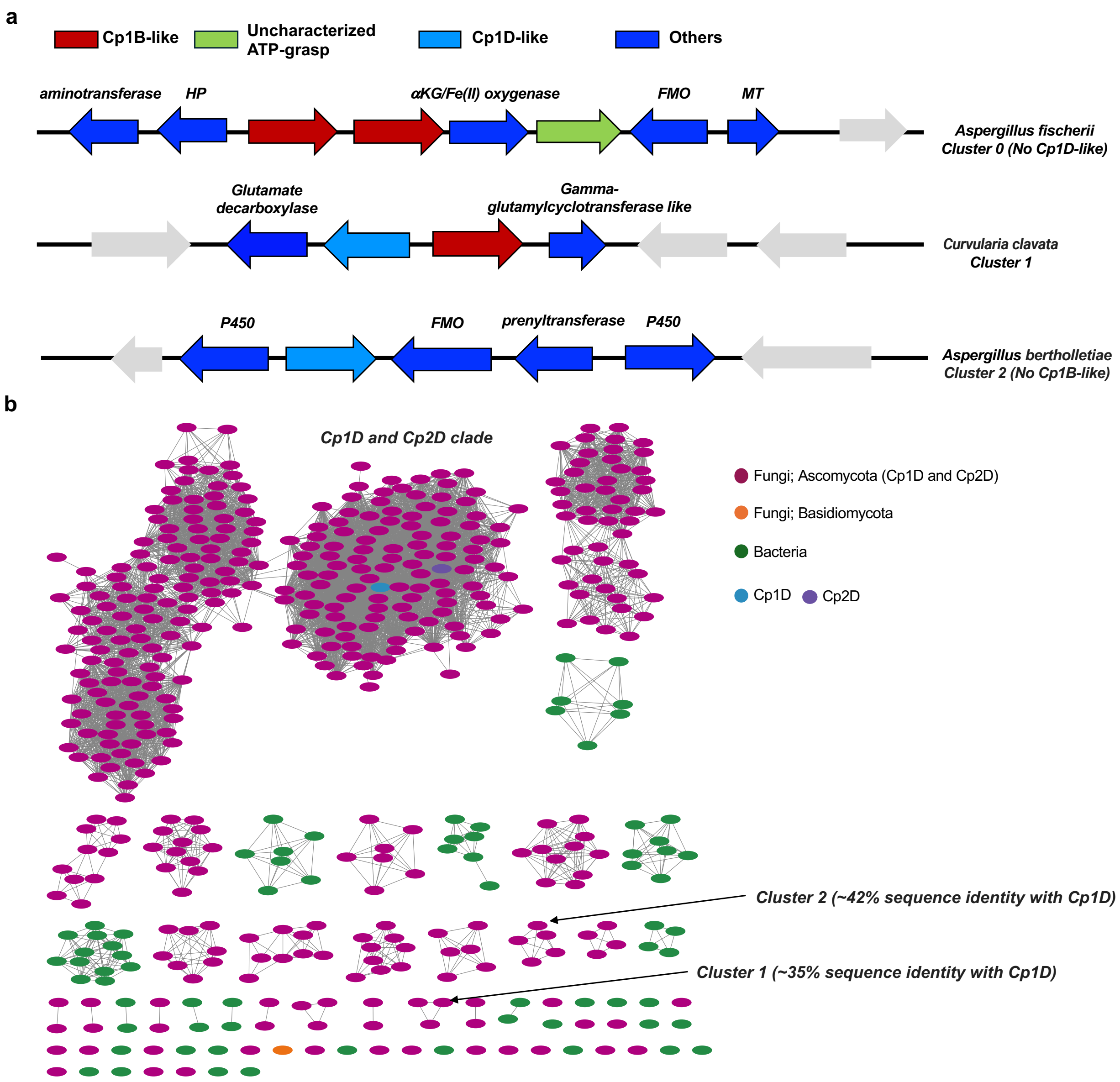

**Supplementary Fig. 8 | Non-E-64 biosynthetic gene clusters containing homologs of Cp1B and Cp1D. a**, The BGCs with Cp1B-like and/or Cp1D-like enzymes could be found in fungi such as *Aspergillus fischerii* (Cluster 0), *Curvularia clavata* (Cluster 1), *Aspergillus bertholletiae* (Cluster 2). These cryptic clusters such as Cluster 0 encode one or multiple ATP-grasp enzyme homologous to Cp1B, and the presence of multiple amide bond forming enzyme or oxidation enzyme implies that these clusters may code new amide-containing compounds with oxidative modifications. **b**, Sequence similarity network (SSN) of Cp1D and homologues. Protein sequences (maximum 1000) were obtained by performing a BLASTp with query e-value of 5 on the EFI-EST website. The SSN of Cp1D was generated with alignment score threshold of 160. The analysis of SSN led to identify new BGCs such as Cluster 1 and Cluster 2 which potentially code new amide-containing natural products. As similar to the SSN of Cp1B (**Extended Data Fig. 2c**), Cp1D-like enzymes could also be found in the bacterial genomes.

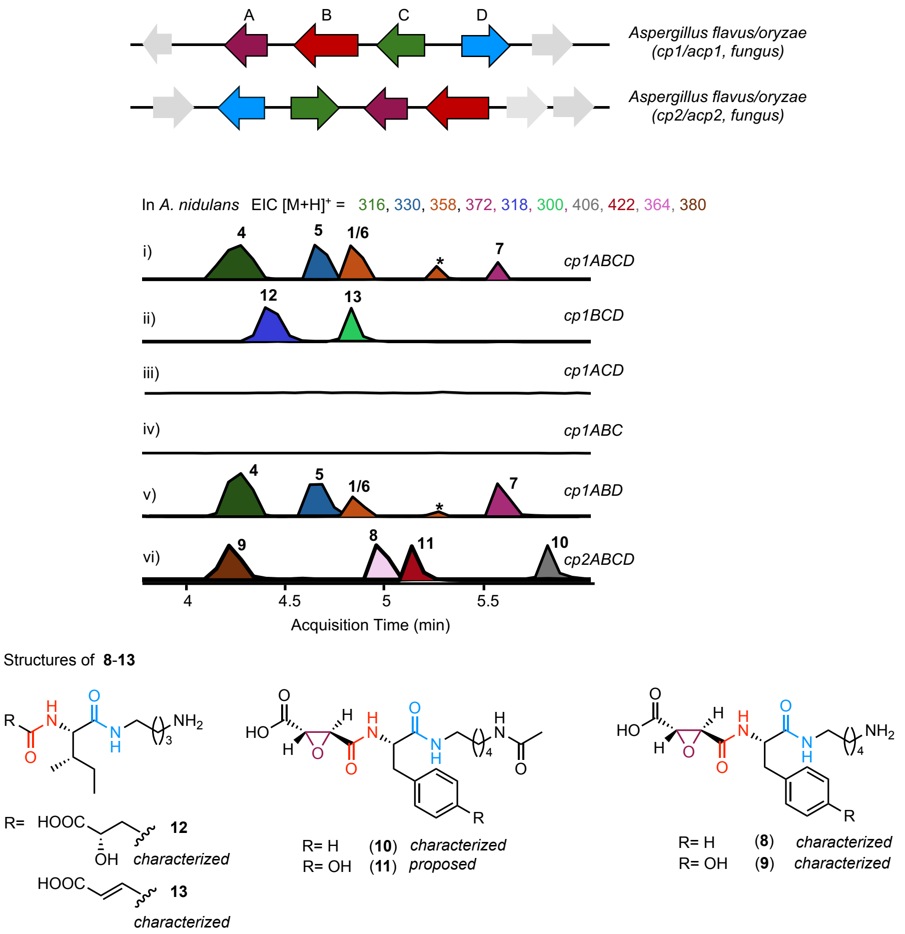

**Supplementary Fig. 9 | LC/MS analysis of extracts from the heterologous expression of *cp1* and *cp2* in *A. nidulans*.** LC/MS analyses include *cp1ABCD* (i), *cp1BCD* (ii), *cp1ACD* (iii), *cp1ABC* (iv), *cp1ABD* (v), and *cp2ABCD* (vi). Selected ion chromatograms correspond to the $[M+H]^+$ for **1** ($[M+H]^+$ = 358), **4** ($[M+H]^+$ = 316), **5** ($[M+H]^+$ = 330), **6** ($[M+H]^+$ = 358), **7** ($[M+H]^+$ = 372), **8** ($[M+H]^+$ = 364), **9** ($[M+H]^+$ = 380), **10** ($[M+H]^+$ = 406), **11** ($[M+H]^+$ = 422), **12** ($[M+H]^+$ = 318), and **13** ($[M+H]^+$ = 300). The y-axis represents the mass intensity and is presented on the same scale. Heterologous expression of three gene cassette *cp1ABD* is sufficient for the biosynthesis of **1** and the analogs in *A. nidulans*. As polyamines are abundant primary metabolites in fungi, the PLP-dependent decarboxylase Cp1C is not essential for the biosynthesis of **1** and the analogs in heterologous host *A. nidulans*. Interestingly, the heterologous expression of *cp1BCD* led to the formation of malic acid (**12**) and fumaric acid (**13**) derivatives, suggesting the role of Cp1A as an epoxidase. This result further supported the highly promiscuous characters of both Cp1B and Cp1D as a biocatalyst. The structures of all compounds except for **11** were determined by NMR.

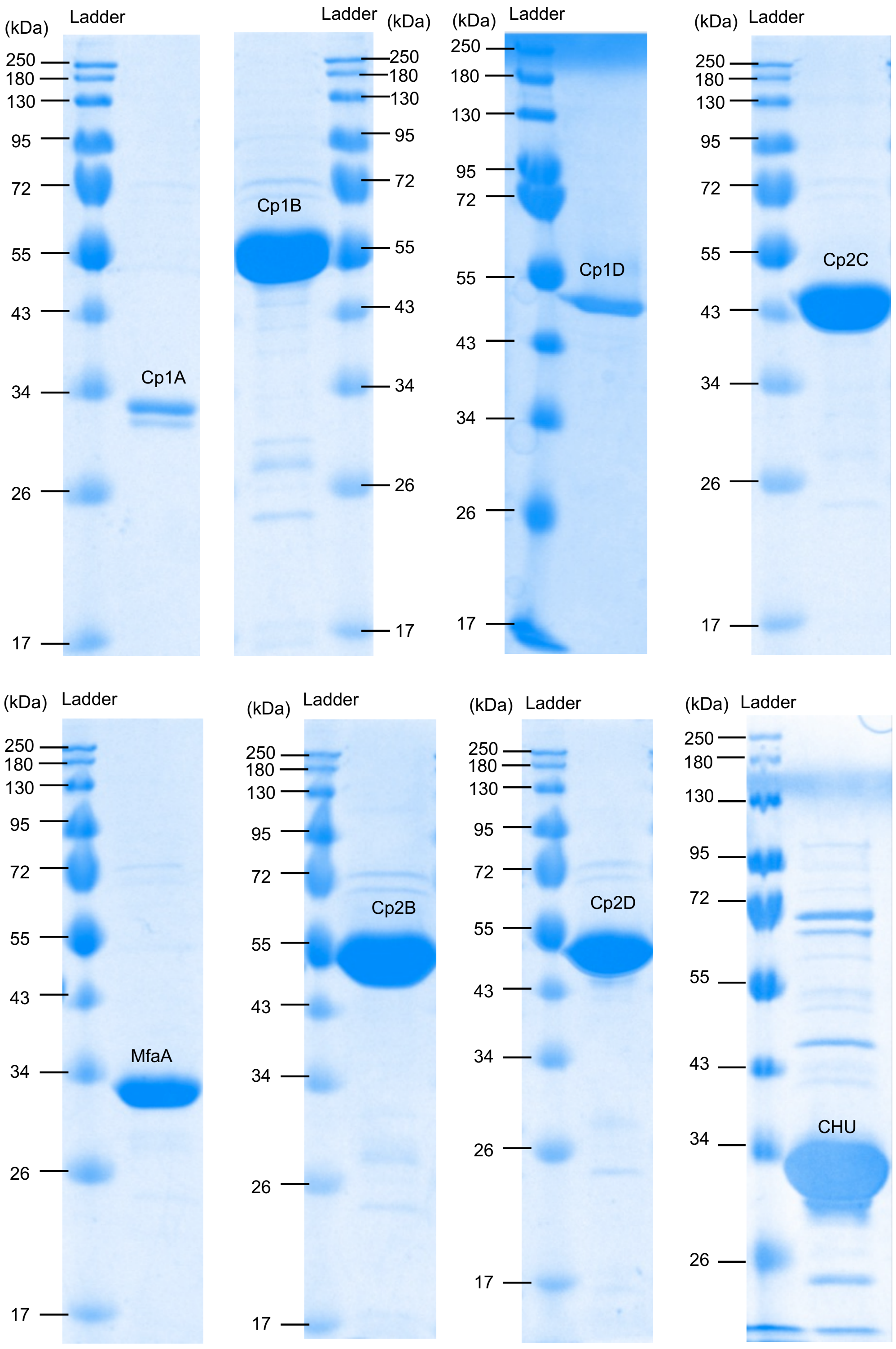

**Supplementary Fig. 10 | SDS-PAGE gels of purified proteins used in this study.** Expected molecular weights of Cp1A (oxygenase), Cp1B (HP), Cp1D (ABS), MfaA (oxygenase), Cp2C (decarboxylase), Cp2B (HP), Cp2D (ABS), CHU (Polyphosphate kinase) are 33 kDa, 57 kDa, 52 kDa, 32 kDa, 45 kDa, 57 kDa, 52 kDa and 36 kDa, respectively. These experiments were repeated three times independently and representative results are shown.

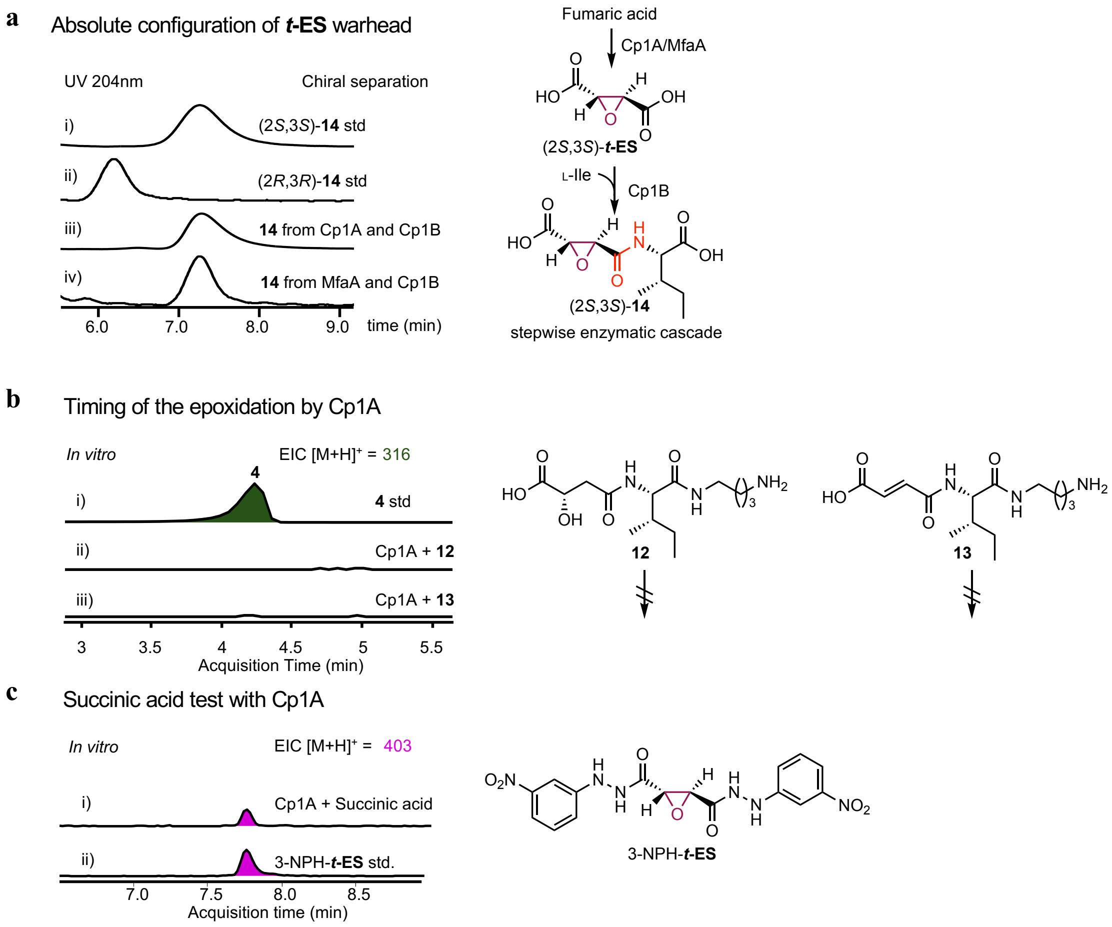

**Supplementary Fig. 11 | Absolute configuration of *t*-ES and substrate for Cp1A. a**, Enzymatic synthesis of (2*S*,3*S*)-***t*-ES** from Cp1A or MfaA, which was subjected to the subsequent enzymatic transformation with Cp1B. Briefly, the reaction was performed in 50 mM sodium phosphate buffer (pH 8.0) containing 0.2 mM $FeSO_4$, 2 mM αKG, 2 mM ascorbate, 1 mM of substrate, and 10 µM of Cp1A or MfaA at 30 °C for 16 h. Then the protein was removed by Amicon concentrators (Millipore). Subsequently 10 µM enzyme, 2.5 mM L-isoleucine, ATP cofactor (10 mM), $MgCl_2$ (10 mM) were added followed by incubation at 30°C for 16 h. The formation of (2*S*,3*S*)-**14** established the absolute configuration of the enzymatically synthesized warhead to be 2*S*, 3*S* in comparison to the HPLC retention time of standards. The HPLC analysis was performed with a CHIRALPAK® IA-3 column (150 x 4.6 mm, 3 µm) at room temperature (flow rate 1 mL/min, 40% MeCN–$H_2O$ with 0.1% trifluoroacetic acid). **b**, LC/MS analysis of in vitro reaction of Cp1A with **12** or **13**. 100 µL reactions were performed at 30 °C for 3 h, in 50 mM sodium phosphate buffer (pH 8.0) containing 0.2 mM $FeSO_4$, 2 mM αKG, 2 mM ascorbate, 1 mM of substrate **12** or **13**, and 10 µM of Cp1A. **4** was not observed in enzymatic assay of Cp1A with **12** (ii) and **13** (iii) in the presence of αKG, ascorbate, and $Fe^{2+}$. The traces show selected ion monitoring of **4** ($[M+H]^+$ = 316) after addition of fumaryl- (**13**) or L-malyl-L-Ile-putrescine (**12**) to the enzymatic reaction mixtures. **c**, In vitro reaction of Cp1A with succinic acid. The same reaction condition with Extended Data Fig. 5b was used except for succinic acid being used as the substrate. After overnight incubation at 30 °C, the product was derivatized with 3-NPH. Selected ion monitoring of 3-NPH-***t*-ES** ($[M+H]^+$ = 403) is shown. Note that 3-NPH-fumaric acid was also observed when succinic acid was used as the substrate.

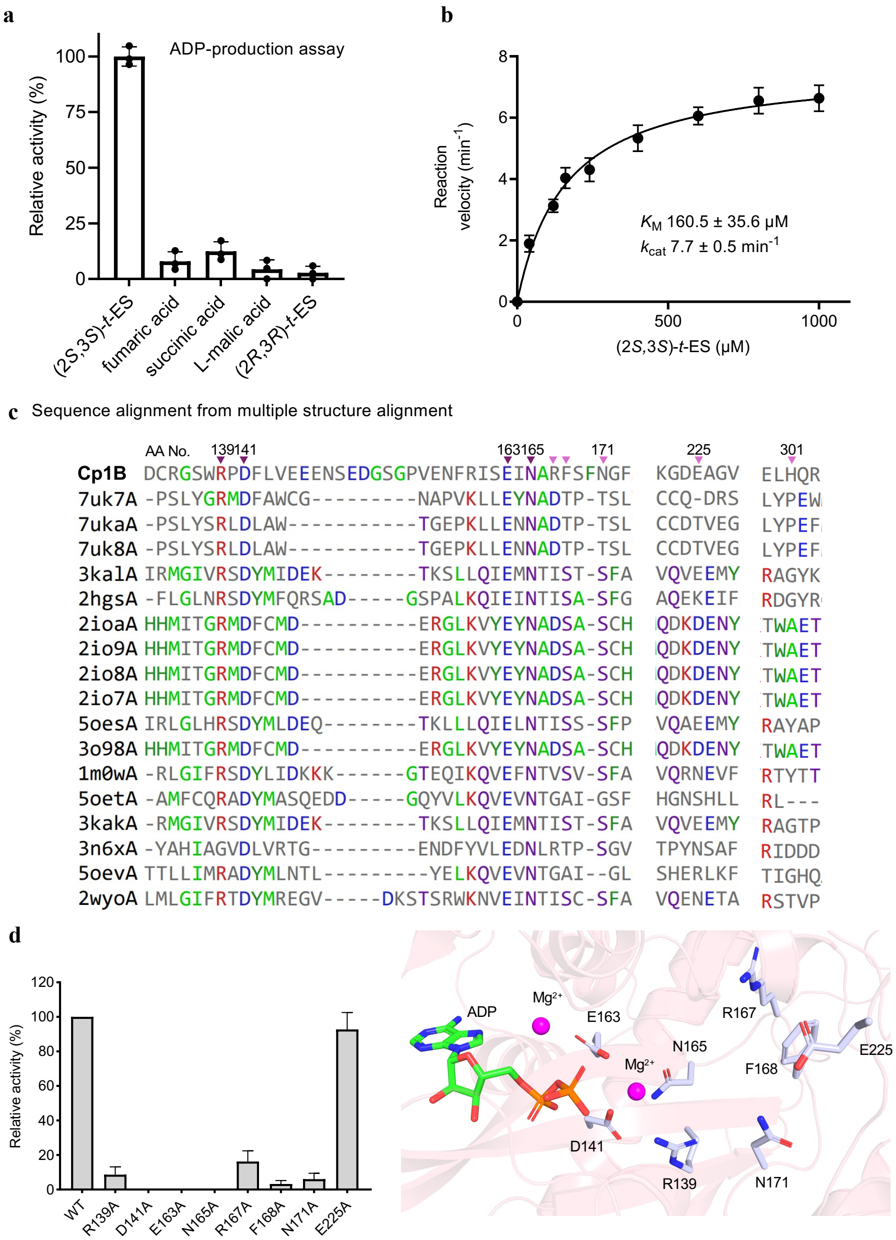

**Supplementary Fig. 12 | Cp1B is a new ATP-grasp enzyme. a**, Relative rates of ADP formation on incubation of Cp1B with potential dicarboxylic acid substrates. Cp1B strongly prefers (2*S*,3*S*)-***t*-ES** as the acid acceptor. The reaction was performed in 200 μL of 100 mM Tris-HCl (pH 8.0) containing 0.25 μM Cp1B, 10 mM ATP, 12 mM $MgCl_2$, 300 $\mu$M NADH, 500 $\mu$M phosphoenolpyruvic acid (PEP), 41 units/mL pyruvate kinase (PK, Sigma), 59 units/mL lactate dehydrogenase (LDH, Sigma), 10 mM KCl with 1 mM epoxy-succinic acid or other acid donors and 5 mM L-Phe. The relative phosphorylation activities of Cp1B toward each substrate were derived by the consumption of NADH at the time point where each reaction mixture was incubated at 30 °C for 30 min. Error bars indicate s.d. of three independent replicates. **b**, Apparent Michaelis-Menten plots for the Cp1B catalyzed phosphorylation of (2*S*,3*S*)-***t*-ES**. The values represent means ± s.d., and error bars indicate s.d. of more than three independent replicates. The reaction mixtures (100 μL) contained 1.0 μM Cp1B, 10 mM ATP, 12 mM $MgCl_2$, 300

$\mu$M NADH, 500 $\mu$M phosphoenolpyruvic acid (PEP), 41 units/mL pyruvate kinase (PK, Sigma), 59 units/mL lactate dehydrogenase (LDH, Sigma), 10 mM KCl and 100 mM Tris-HCl (pH 8.0) with various concentration (0.04 mM to 1 mM) of (2*S*,3*S*)-*t*-ES and 5 mM L-Phe. The reaction mixture was incubated at 30 °C, and the consumption of NADH at 10 min was used to derive the reaction rate for enzyme kinetics. Kinetic constants were derived from velocity versus substrate concentration data using a nonlinear regression fitting method with GraphPad Prism 9. **c**, Structure-based multiple sequence alignment of Cp1B with other characterized ATP-grasp enzymes including hGSH synthetases. **d**, The activity of each mutant is compared to that of wild-type Cp1B which was quantified by the formation of (2*S*,3*S*)-**14**. Error bars indicate s.d. of three independent replicates. The mutation of conserved residues such as R139, D141, E163, and N165, which are expected to form the interaction with ATP and/or $Mg^{2+}$, significantly decreased or abolished the activity. Since the mutation of R167, which is not conserved in other characterized ATP-grasp enzyme, also significantly decreased the activity, we reasoned that R167 likely anchors the carboxylic acid of (2*S*,3*S*)-**t-ES** or amino acid donor through hydrogen bond formation. Other residues unique to Cp1B such as F168 and N171 diminished the activity, implying that these residues might have important role in forming the active site pocket.

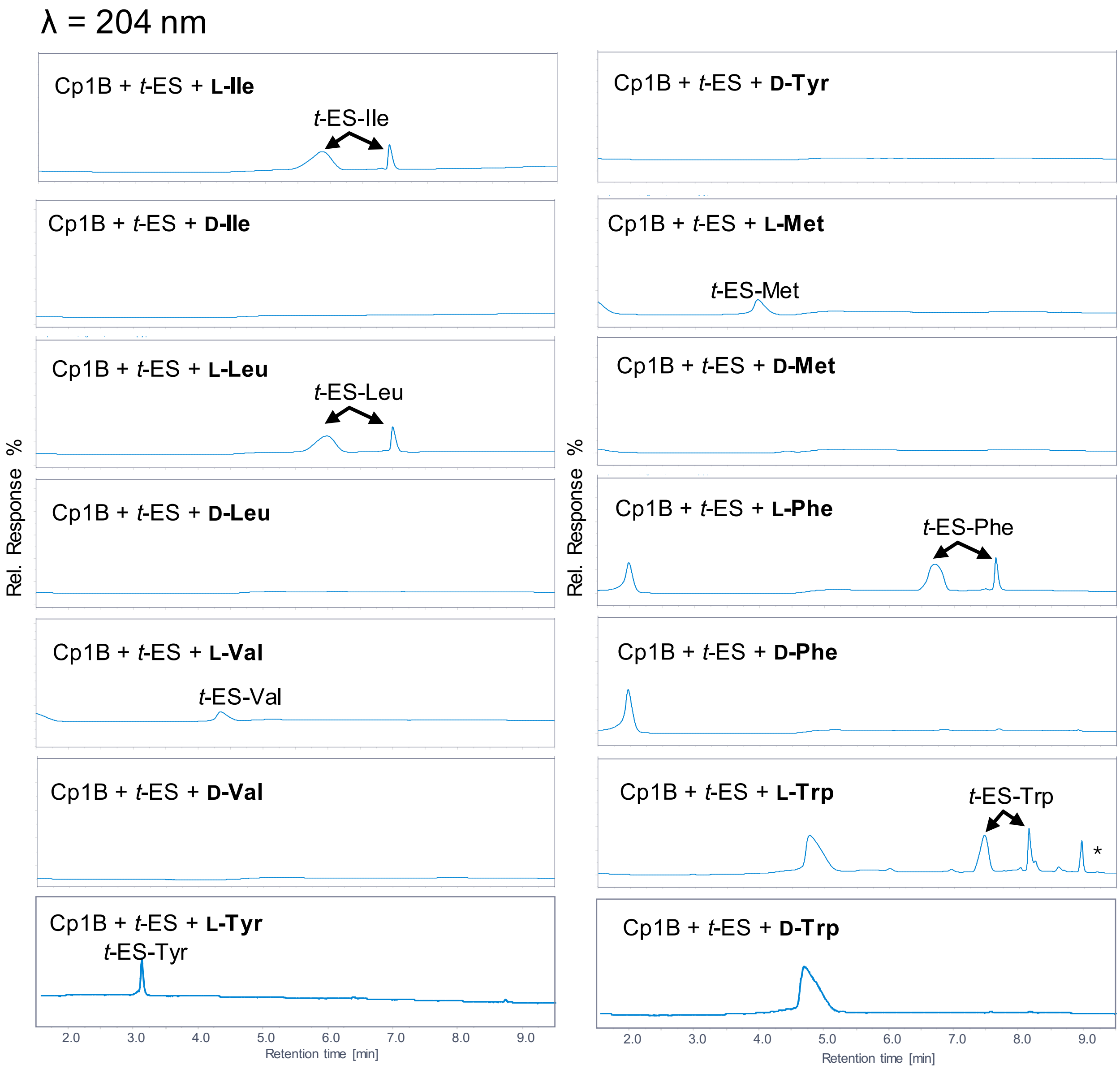

**Supplementary Fig. 13 | D-amino acids are not substrate for Cp1B.** LC/MS analysis of *in vitro* assay of Cp1B with hydrophobic D-amino acids. The corresponding products were not detected when D-amino acid was used. Assays were carried out with enzymes (25 μM), (±)-*trans*-epoxy-succinate (5 mM) and amino acid (2.5 mM), 10 mM $MgCl_2$, 10 mM ATP, 50 mM sodium phosphate. Note: Epoxy-succinyl L-amino acids were present as two split peaks which result from different charge state (0.1% formic acid was used as the additive for LC/MS solvents). The LC-MS elution gradients method: 0-0.25 min, 1% eluent B; 0.25-13.0 min, 1-99% eluent B; 13.0-16.0 min, 99% eluent B; 16.0-18.0 min, 1% eluent B using Agilent LC/MSD iQ (AgilentTM InfinityLab Poroshell 120 Aq-C18, 2.7 μm, 100 Å, 2.1 × 100 mm).

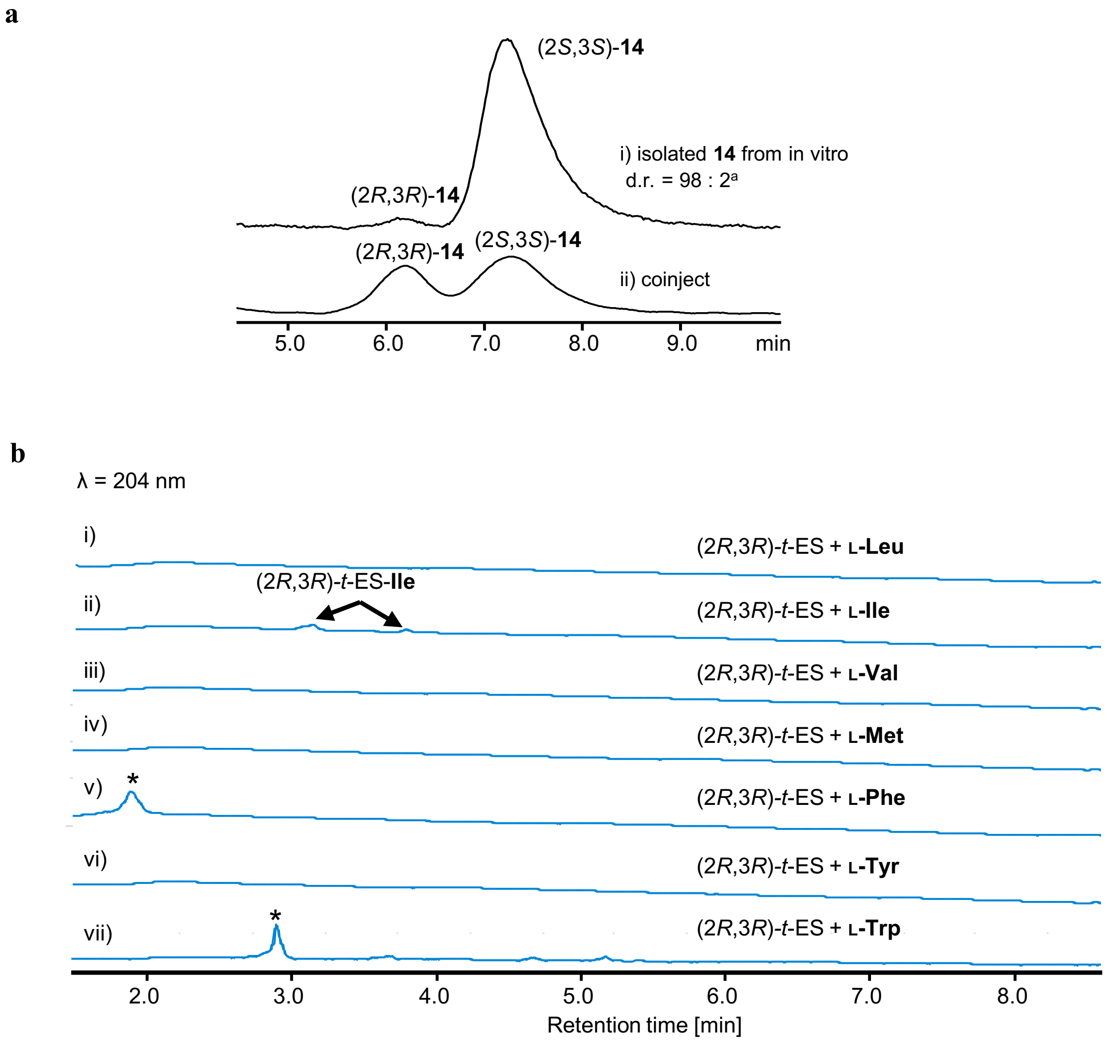

**Supplementary Fig. 14 | Examination of the kinetic resolution potential for Cp1B. a**. i) The HPLC chromatogram for ***t*-ES**-L-Ile **(14)** that was enzymatically prepared from racemic *t*-ES with L-Ile. d.r. value was calculated from HPLC peak area ratio using chiral analytical HPLC with a CHIRALPAK® IA-3 (150 x 4.6 mm, 3 µm) at room temperature (flow rate 1 mL/min, 40% MeCN–$H_2O$ with 0.1% trifluoroacetic acid). ii) the coinjection of (2*S*,3*S*)-*t*-ES-L-Ile (**14)** and (2*R*,3*R*)-**14**. **b**. (2*R*,3*R*)-*t*-ES test with proteinogenic amino acids. While only in vitro assay with L-Ile gave trace amount pseudopeptides, no product can be observed when tested with other L-amino acids. The LC-MS elution gradients method: 0-0.25 min, 1% eluent B; 0.25-13.0 min, 1-99% eluent B; 13.0-16.0 min, 99% eluent B; 16.0-18.0 min, 1% eluent B using Agilent LC/MSD iQ (AgilentTM InfinityLab Poroshell 120 Aq-C18, 2.7 µm, 100 Å, 2.1 × 100 mm). *: amino acid substrate.

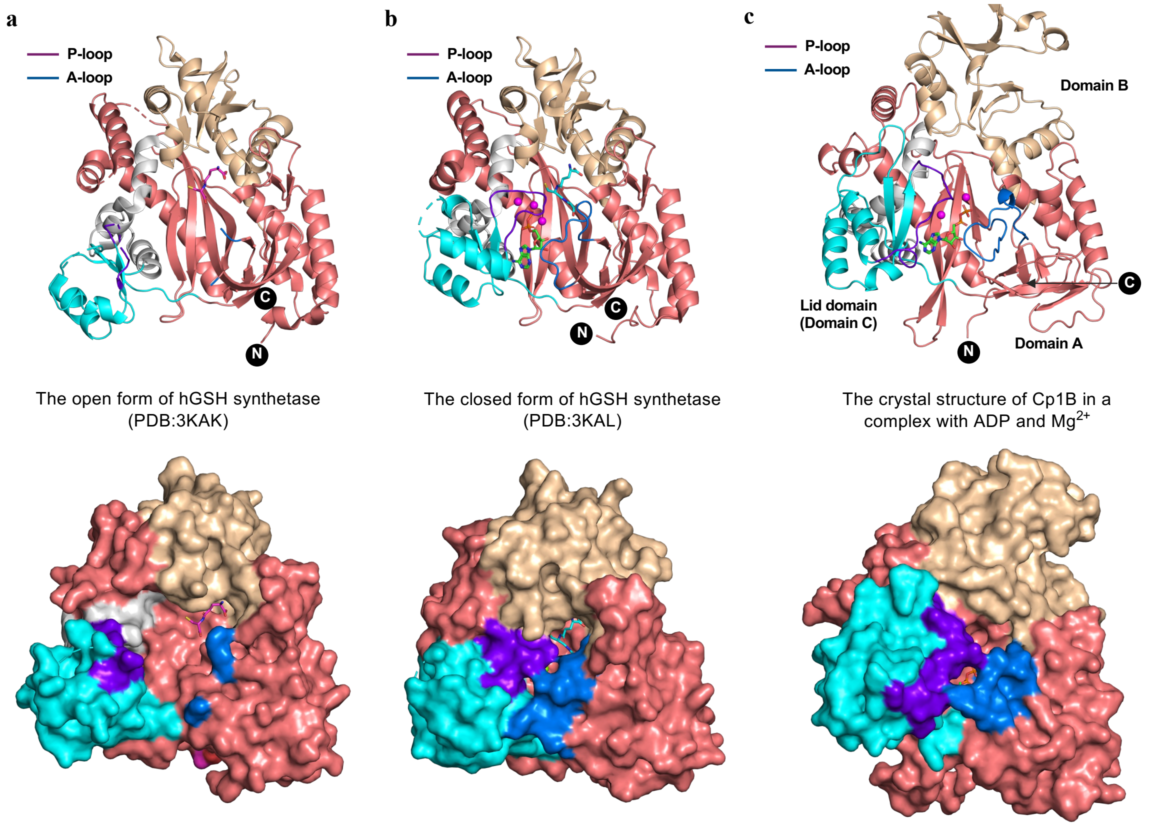

**Supplementary Fig. 15 | The crystal structure of Cp1B adopts a closed active site form.** Comparisons of overall structure folds of hGSH synthetase with the open active site form (3KAK) (**a**), with the closed active site form (3KAL) (**b**), and Cp1B in a complex with ADP and $Mg^{2+}$ (**c**) are shown. All structures have three characteristic domains, Domain A (deepsalmon), Domain B (wheat), and lid domain (Domain C, cyan), which is a typical fold of ATP-grasp enzymes. Surface representation of the crystal structures is shown below of each cartoon representation. Stick drawings show γ-glutamylcysteine (magenta) in the open form, ADP (green) and hGSH (cyan) in the closed form, and ADP (green) in the Cp1B structure. $Mg^{2+}$ ions in the closed form and Cp1B are shown as magenta spheres. P-loop (Gly-rich loop) and A-loop (Ala-rich loop) are shown in purple and blue, respectively. In the open form, the P-loop and A-loop are disordered in contrast to those in the closed form and the Cp1B structure. Consequently, in the open form (3KAK), the lid domain and A-loop leave the nucleotide binding site open. In contrast, the lid domain with P-loop and A-loop enclose the active site in the closed form (3KAL) and the Cp1B structure. These structural comparisons therefore suggested that the crystal structure of Cp1B adopts a closed active site for the catalysis.

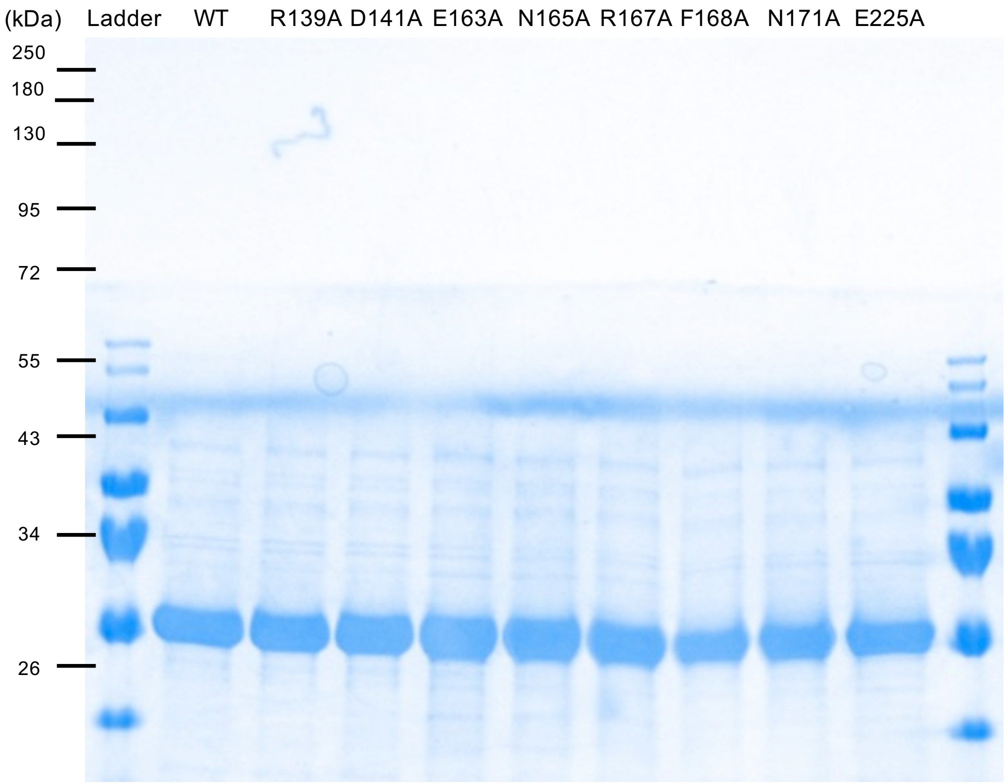

**Supplementary Fig. 16 | SDS-PAGE gels of Cp1B mutant.**

a. chiral resolution by L-arginine (Kanaoka, Hesse)

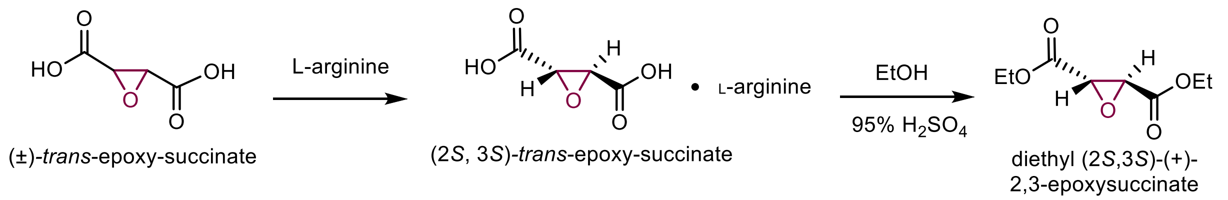

b. stereoselective synthesis (Bogyo)

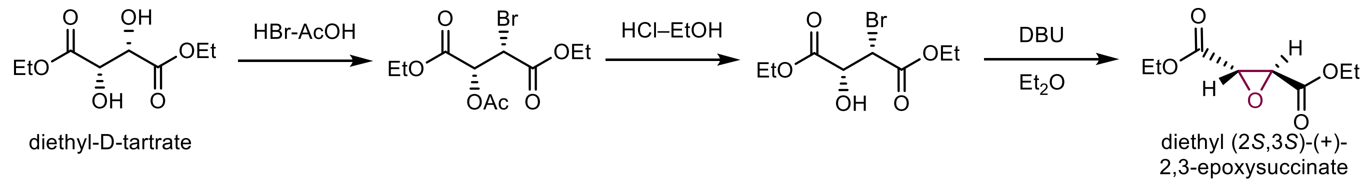

**Supplementary Fig. 17 | Previous work for the preparation of (2*S*,3*S*)-*trans*-epoxy-succinic acid**[21-23].

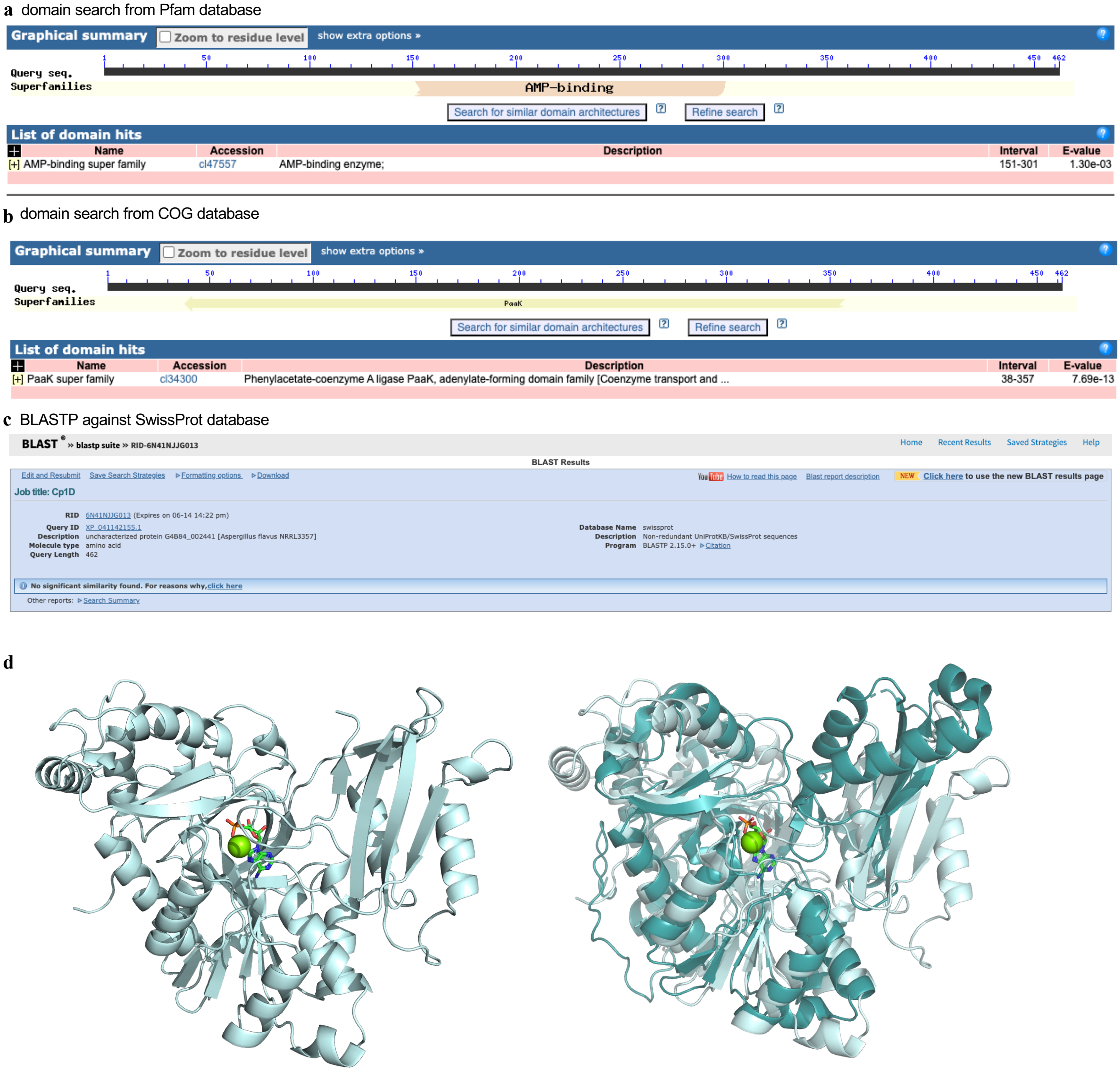

**Supplementary Fig. 18 | Bioinformatic analysis of Cp1D. a**. Domain search of Cp1D from Pfam database. Cp1D has the weak conservation of AMP-binding superfamily domain. **b**. Domain search of Cp1D from COG database. Cp1D belongs to the family of phenylacetate-CoA ligase, suggesting Cp1D is also an ANL-family enzyme. **c**. BLASTP search using Cp1D as the query against SwissProt database. No characterized homolog of Cp1D was identified from the database. **d**. Left, AF3[20] predicted Cp1D with AMP and $Mg^{2+}$. Right, overlay of Cp1D and PDB 6he0[24], shown in cartoon representation, RMSD = 5.367, (2096 to 2096 atoms by using “super” alignment method in pymol).

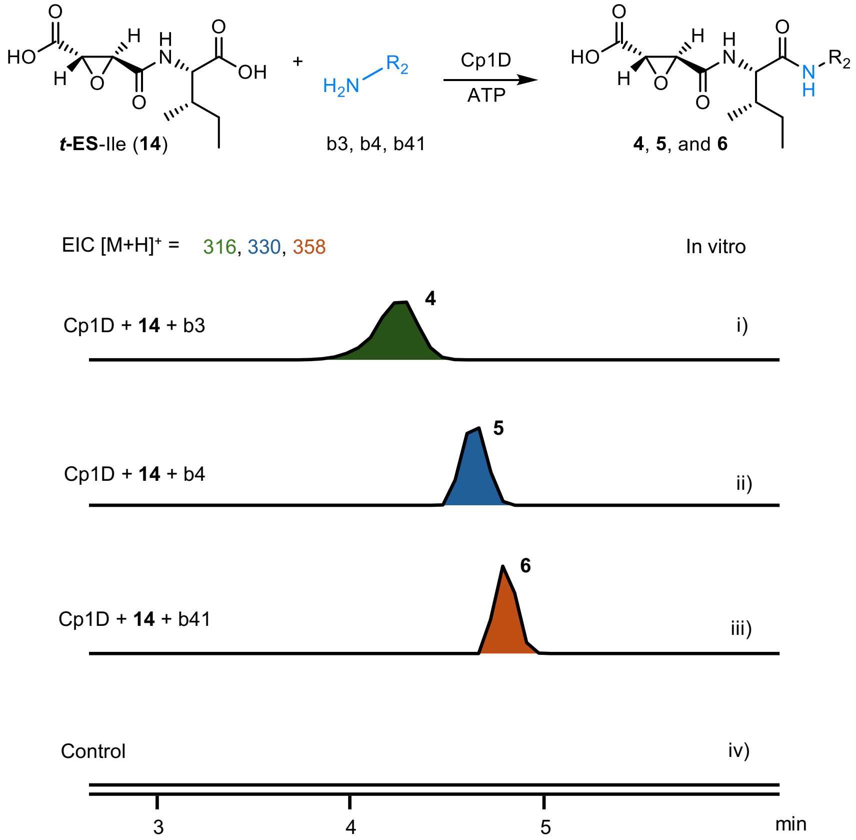

**Supplementary Fig. 19 | In vitro reaction of Cp1D with 14 and amines**. LC/MS analysis of in vitro assay of Cp1D with **14** and putrescine (b3, i), cadaverine (b4, ii), and agmatine (b41, iii). The exclusive formation of **4** ($[M+H]^+$ = 316), **5** ($[M+H]^+$ = 330) and **6** ($[M+H]^+$ = 358) was detected.

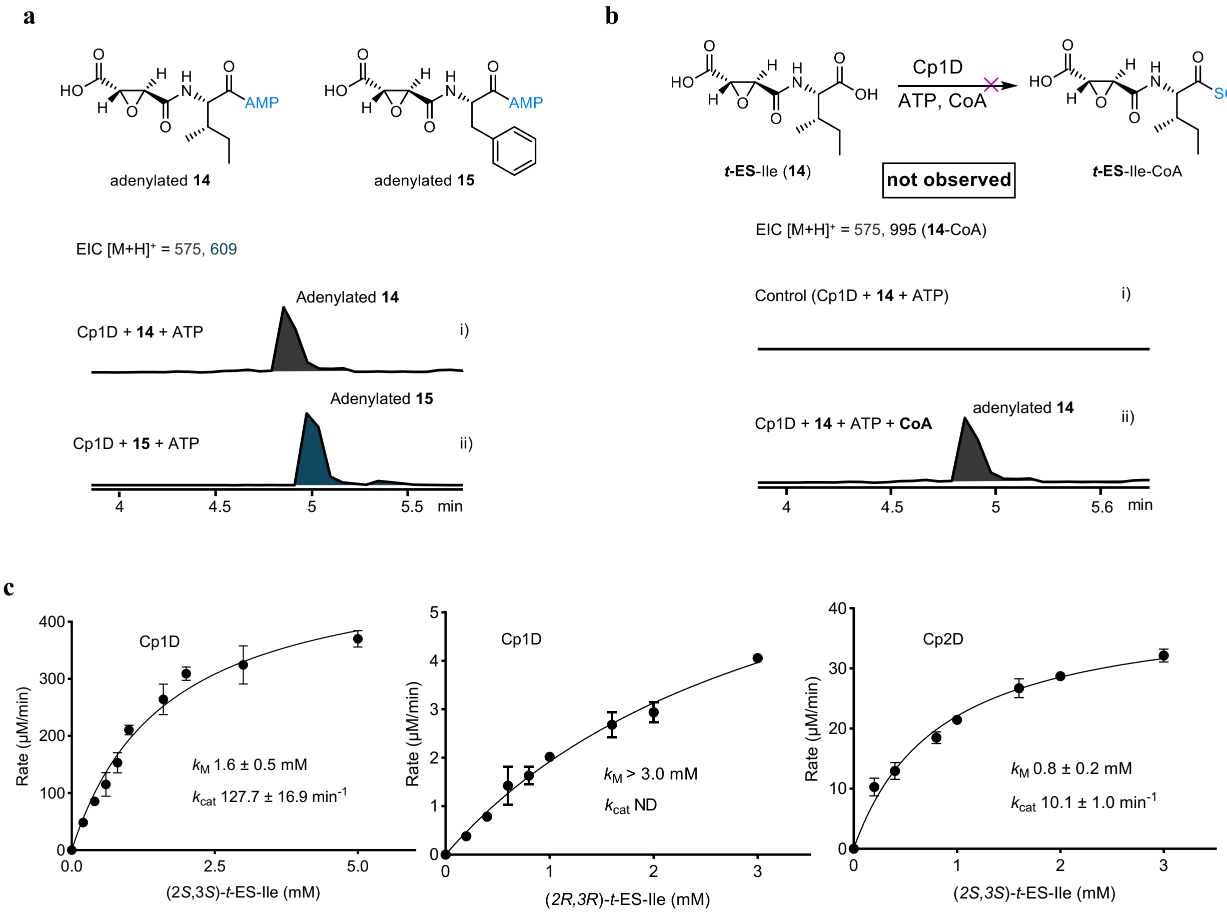

**Supplementary Fig. 20 | Cp1D is a new family of diamide-forming amide bond synthetase not CoA ligase. a**. Cp1D activity assay with (2*S*,3*S*)-*t*-ES-Ile (**14**), and (2*S*,3*S*)-*t*-ES-Phe (**15**) as substrate, the MS peak of adenylated **14** (*m/z* 575 for $[M+H]^+$) and adenylated **15** (*m/z* 609 for $[M+H]^+$) was observed. Assays were carried out with enzymes (25 μM), **14** or **15** (0.4 mM), 10 mM $MgCl_2$, 10 mM ATP in 100 μL of 50 mM sodium phosphate (pH 8.0). Reactions were analyzed by UPLC-MS after incubation at 30 °C for 10 min. **b**. Assays were carried out with enzymes (25 μM), **14** (0.4 mM), 1 mM CoA-SH, 10 mM $MgCl_2$, 10 mM ATP in 100 μL of 50 mM sodium phosphate (pH 8.0). Reactions were analyzed by UPLC-MS after incubation at 30 °C for 1 h. Selected ion chromatography of *t*-ES-Ile-CoA (*m/z* 995 for $[M+H]^+$) and adenylated **14** (*m/z* 575 for $[M+H]^+$) is shown. The MS peak of *t*-ES-Ile-CoA was not observed. **c**. Apparent Michaelis-Menten plots for the Cp1D and Cp2D catalyzed amidation with single diastereomer (2*S*,3*S*)-*t*-ES-Ile or (2*R*,3*R*)-*t*-ES-Ile. The kinetic constants are shown as an apparent value.

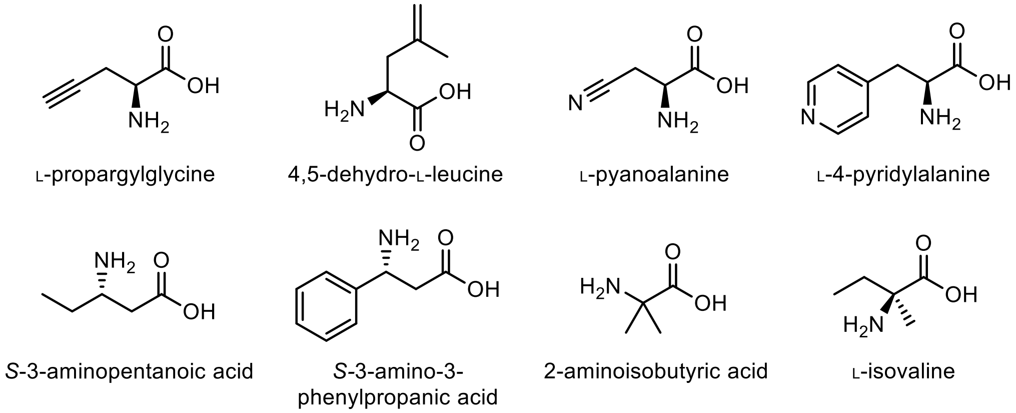

**Supplementary Fig. 21 | Structure of unnatural amino acid tested but could not be accepted by Cp1B/Cp2B.** Amino acids with double bond, triple bond, nitrile, pyridine are disfavored. α, α-disubstituted amino acid, *N*-methylated amino acid was also not preferred.

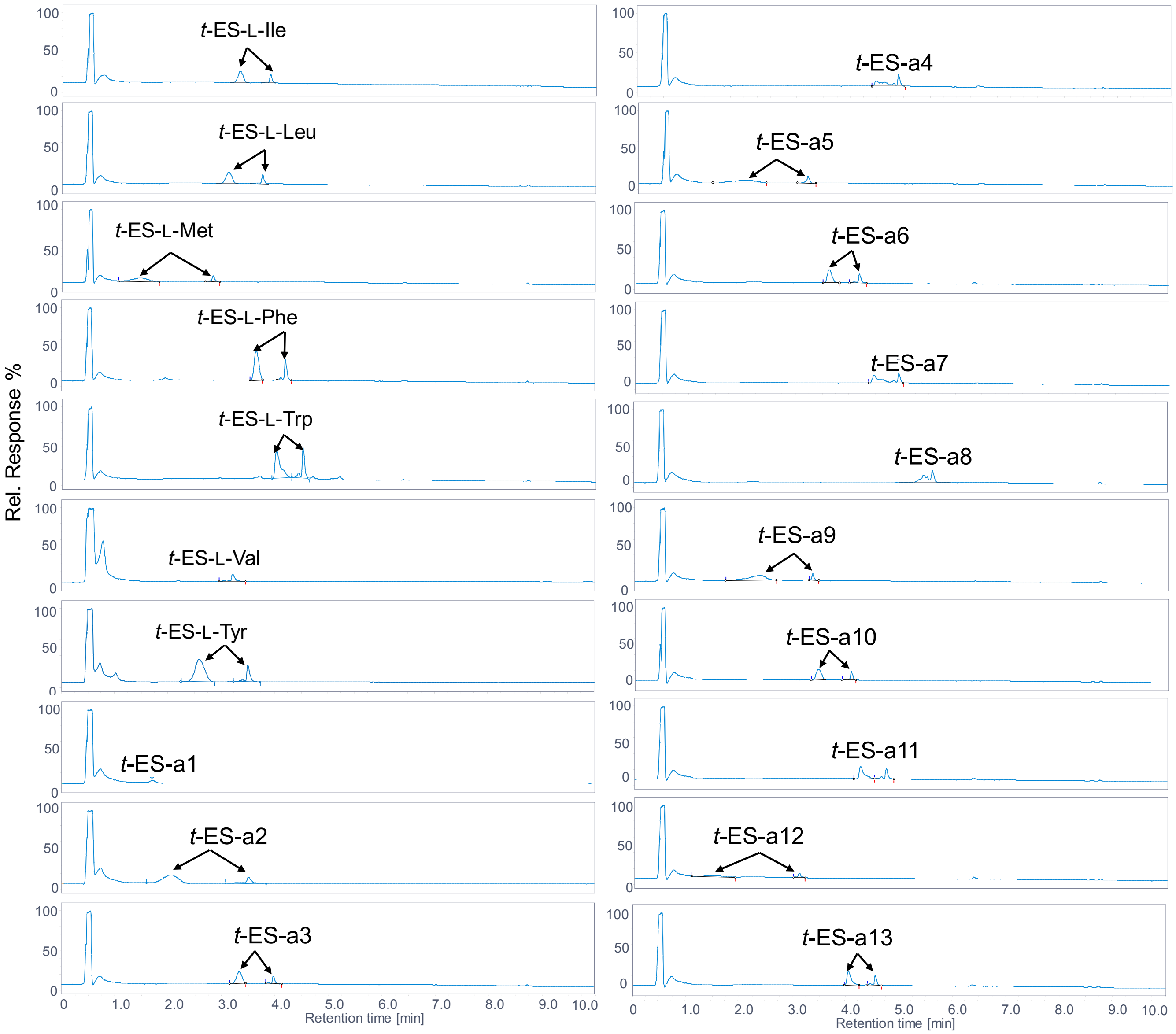

*t*-ES-L-Ile
*t*-ES-L-Leu
*t*-ES-L-Met
*t*-ES-L-Phe
*t*-ES-L-Trp
*t*-ES-L-Val
*t*-ES-L-Tyr
*t*-ES-a1
*t*-ES-a2
*t*-ES-a3
*t*-ES-a4
*t*-ES-a5
*t*-ES-a6
*t*-ES-a7
*t*-ES-a8
*t*-ES-a9
*t*-ES-a10
*t*-ES-a11
*t*-ES-a12
*t*-ES-a13
Rel. Response %
100
50
0
0
1.0
2.0
3.0
4.0
5.0
6.0
7.0
8.0
9.0
10.0
Retention time [min]

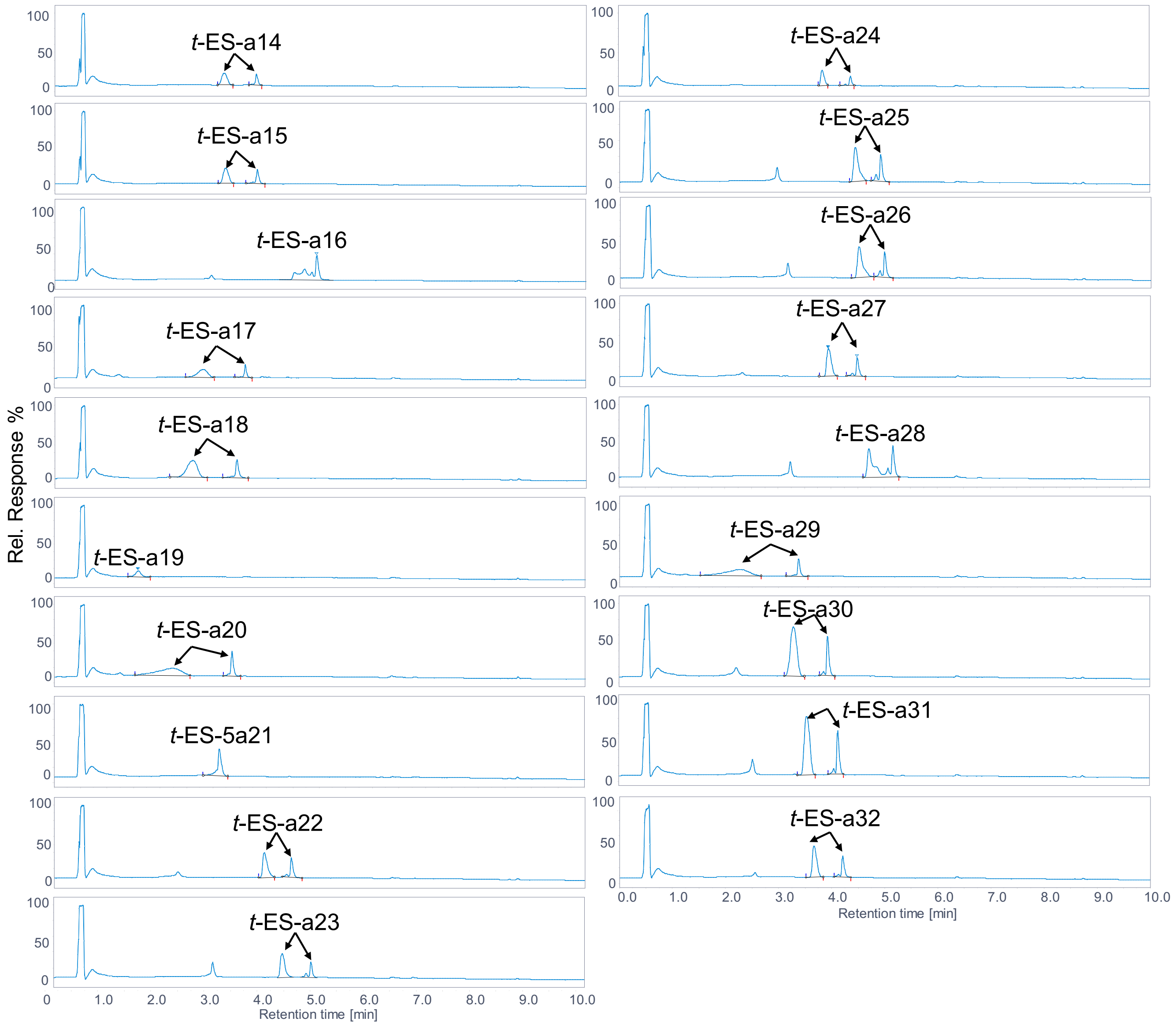

**Supplementary Fig. 22 | HPLC traces of (2*S*,3*S*)-*t*-ES-amino acids formed by the Cp1B reaction.** Substrate scope test for Cp1B against coupling of (±)-***t*-ES** with different proteinogenic and non-proteinogenic amino acids. Cp1B was found to accept hydrophobic L-amino acids (I, L, M, F, W, V, Y, a1-a32). Assays were carried out with enzymes (25 µM), (±)-*trans*-epoxy-succinate (5 mM) and amino acid (2.5 mM), 10 mM $MgCl_2$, 10 mM ATP in 50 mM sodium phosphate buffer (pH 8.0). Note: the corresponding products were present as two split peaks which result from different charge state of the product (0.1% formic acid was used as the additive for LC/MS solvents). Only one peak was observed during compound isolation by HPLC as 0.1% trifluoroacetic acid was added to eluents. Percentage yield was quantified based on the standard curve of **14** or the enzymatically synthesized standard at λ 204 nm. The LC/MS elution gradients method: 0-0.25 min, 1% eluent B; 0.25-13.0 min, 1-99% eluent B; 13.0-16.0 min, 99% eluent B; 16.0-18.0 min, 1% eluent B using Agilent LC/MSD iQ (AgilentTM InfinityLab Poroshell 120 Aq-C18, 2.7 µm, 100 Å, 2.1 × 100 mm).

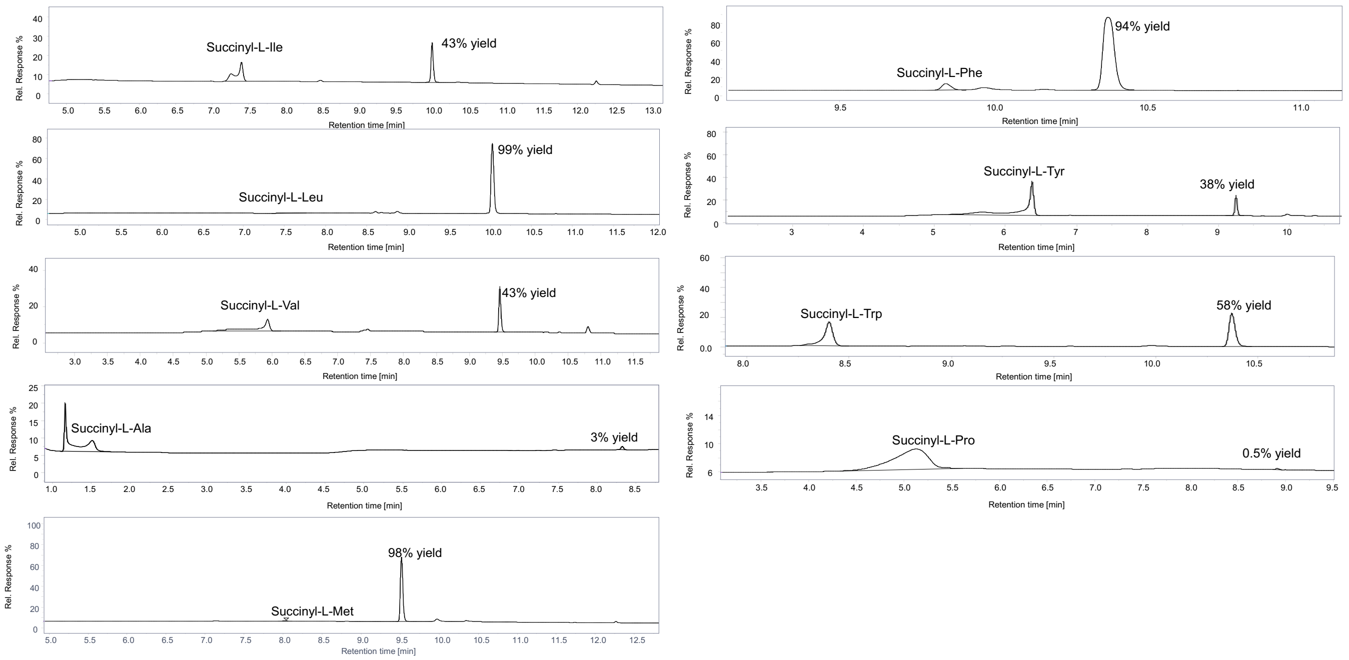

**Supplementary Fig. 23 | Substrate scope test for Cp1D against coupling of isopentylamine with synthesized 20 *N*-succinyl proteinogenic amino acids**. Cp1D was found to accept *N*-succinyl proteinogenic amino acids with hydrophobic amino acids. Assays were carried out with enzyme (25 µM) of Cp1D, *N*-succinyl proteinogenic amino acid (2 mM) and isopentylamine (5 mM), 10 mM $MgCl_2$, 10mM ATP, in 50 mM sodium phosphate buffer (pH 8.0). Reactions were analyzed by LC/MS after incubation at 30°C for 16 h. Conversion of *N*-succinyl proteinogenic amino acid to the amide products was calculated using HPLC peak area ratios at λ 204 nm of product and substrate. The LC-MS elution gradients method: 0-0.25 min, 1% eluent B; 0.25-13.0 min, 1-99% eluent B; 13.0-16.0 min, 99% eluent B; 16.0-18.0 min, 1% eluent B using Agilent LC/MSD iQ (AgilentTM InfinityLab Poroshell 120 Aq-C18, 2.7 µm, 100 Å, 2.1 × 100 mm).

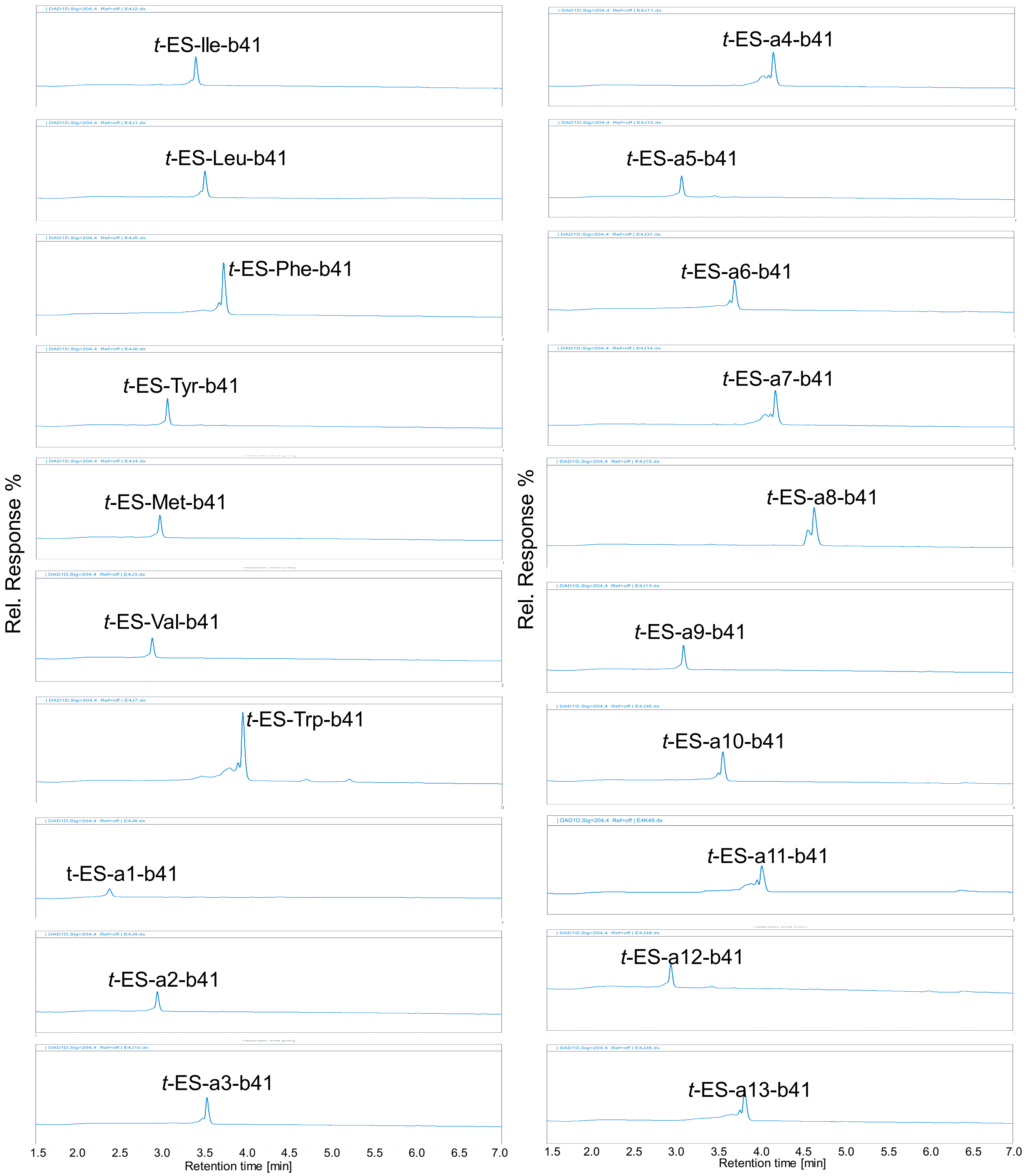
t-ES-Ile-b41
t-ES-Leu-b41
t-ES-Phe-b41
t-ES-Tyr-b41
t-ES-Met-b41
t-ES-Val-b41
t-ES-Trp-b41
t-ES-a1-b41
t-ES-a2-b41
t-ES-a3-b41
t-ES-a4-b41
t-ES-a5-b41
t-ES-a6-b41
t-ES-a7-b41
t-ES-a8-b41
t-ES-a9-b41
t-ES-a10-b41
t-ES-a11-b41
t-ES-a12-b41
t-ES-a13-b41
Rel. Response %
Rel. Response %
1.5 2.0 2.5 3.0 3.5 4.0 4.5 5.0 5.5 6.0 6.5 7.0
Retention time [min]
1.5 2.0 2.5 3.0 3.5 4.0 4.5 5.0 5.5 6.0 6.5 7.0
Retention time [min]

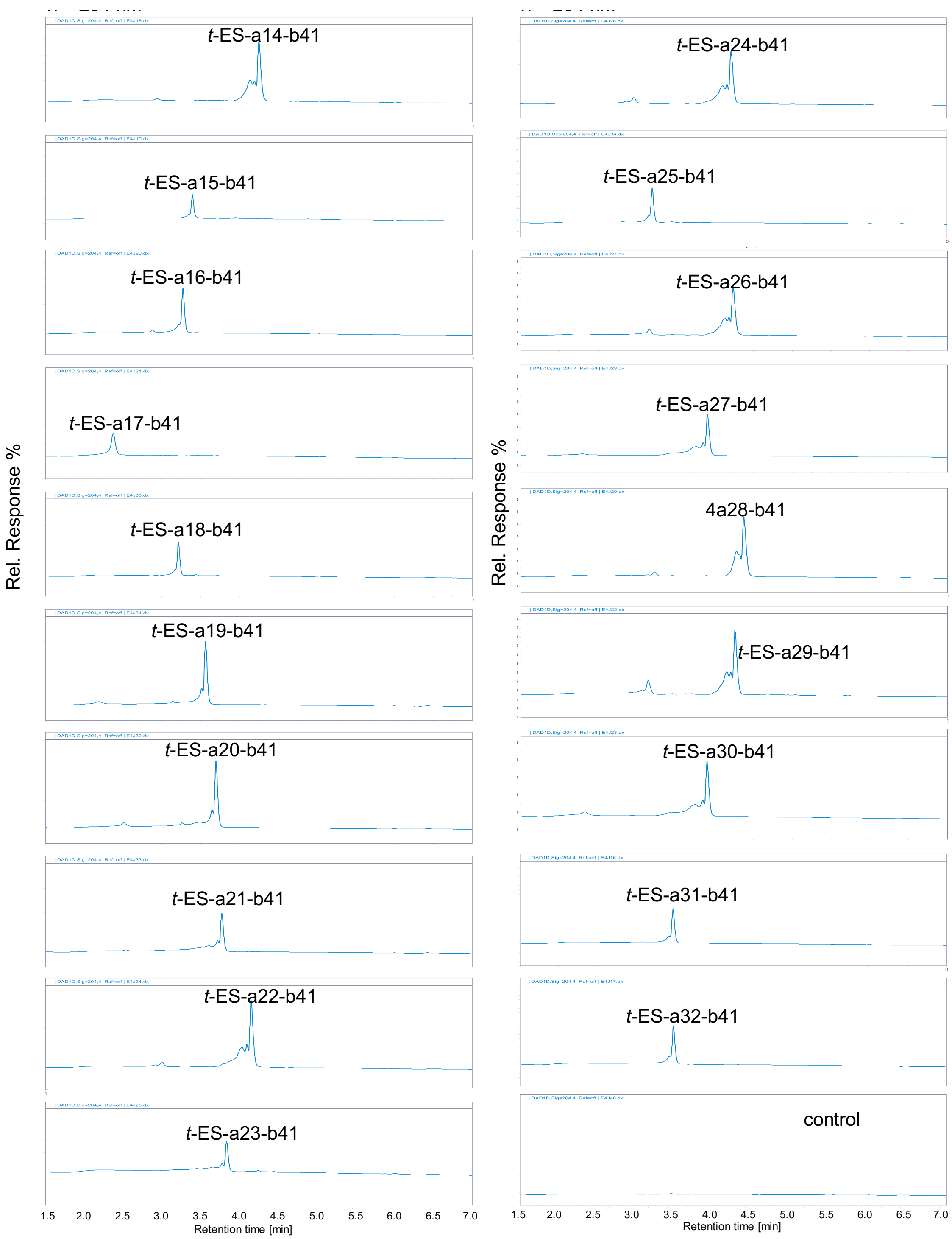

**Supplementary Fig. 24 | HPLC traces of combinatorial biocatalysis with agmatine and different L-amino acids.** Assays were carried out with Cp1B (25 µM), and Cp1D (25 µM), 2 mM (±)-*trans*-epoxy-succinate, 1 mM amino acid, 1 mM agmatine (b41) at 30 °C for 16 h. The LC-MS elution gradients method: 0-0.25 min, 1% eluent B; 0.25-13.0 min, 1-99% eluent B; 13.0-16.0 min, 99% eluent B; 16.0-18.0 min, 1% eluent B using Agilent LC/MSD iQ (AgilentTM InfinityLab Poroshell 120 Aq-C18, 2.7 µm, 100 Å, 2.1 × 100 mm).

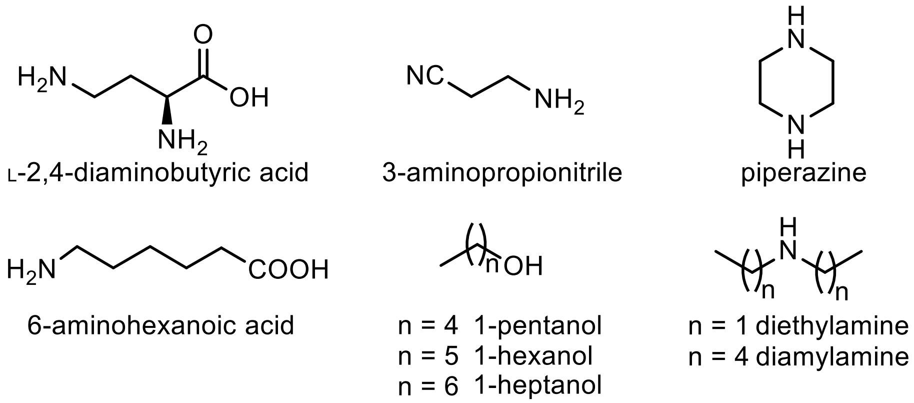

**Supplementary Fig. 25 | Structures of amine nucleophiles which could not be accepted by Cp1D or Cp2D.** Amines with negative charge functional groups were disfavored, such as carboxylic acid. On the other hand, amines with carboxylate ester can be used as amine nucleophiles. Secondary amines were also not preferred except for dimethylamine.

**a**

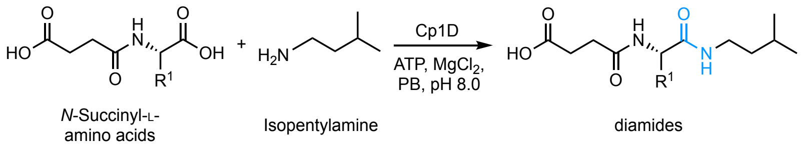

| Substrate | Percentage yield (Cp1D) % | Substrate | Percentage yield (Cp1D) % |
|---|---|---|---|
| Succinyl-L-Ile | 43 | Succinyl-L-Pro | 0.5 |
| Succinyl-L-Leu | >99 | Succinyl-L-Ser | ND |
| Succinyl-L-Val | 43 | Succinyl-L-Thr | ND |
| Succinyl-L-Met | 98 | Succinyl-L-Asn | ND |
| Succinyl-L-Ala | 3 | Succinyl-L-Gln | ND |
| Succinyl-L-Phe | 92 | Succinyl-L-Arg | ND |
| Succinyl-L-Tyr | 38 | Succinyl-L-His | ND |
| Succinyl-L-Trp | 58 | Succinyl-L-Lys | ND |
| Succinyl-L-Cys | ND | Succinyl-L-Asp | ND |
| Succinyl-L-Gly | ND | Succinyl-L-Glu | ND |

**b** Adenylation domain assay for Cp1D/Cp2D

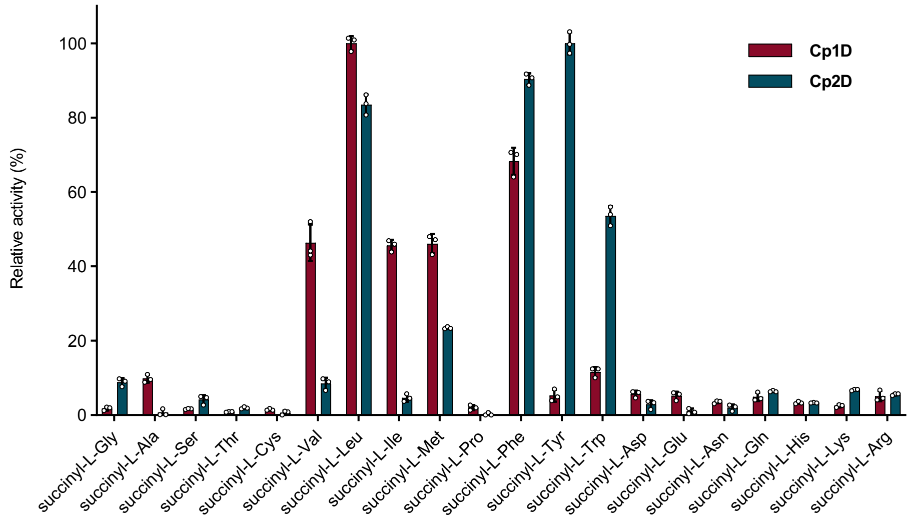

**Supplementary Fig. 26 | Exploration of substrate scope of Cp1D/Cp2D with synthesized twenty *N*-succinyl proteinogenic amino acids. a**, Substrate scope test for Cp1D with 20 synthesized *N*-succinyl-L-amino acids coupling with isopentylamine. Cp1D was found to accept *N*-succinyl-L-amino acids with hydrophobic amino acids (L, I, V, M, F, Y, and W). Assays were carried out with enzymes (25 µM), *N*-succinyl-L-amino acids (2 mM) and isoamylamine (5 mM), 10 mM $MgCl_2$, 10 mM ATP in 100 µL of 50 mM sodium phosphate buffer (pH 8.0). Reactions were analyzed by LC/MS after incubation at 30 °C for 16 h. Conversion (%) of *N*-succinyl-L-amino acid to the corresponding amide product was calculated based on HPLC peak area (UV = 204 nm) ratios between product and starting material. ND means the product was not detected. **b,** Hydroxamate-based colorimetric assay was employed to assess adenylation specificity toward *N*-succinyl-L-amino acids for Cp1D/Cp2D. The reaction was performed in 150 μL of Tris buffer (pH 8.0) containing 20 μM of Cp1D or Cp2D, 15 mM of ATP, 5 mM of *N*-succinyl-L-amino acid substrate, 200 mM hydroxylamine, and 10 mM $MgCl_2$. After the incubation for 8 h at 30 ºC, the reaction was quenched by addition of equivalent volume of stopping solution (10% (w/v) $FeCl_3$ and 3.3% (w/v) trichloroacetic acid dissolved in 0.7 M HCl). The precipitated enzyme was removed by centrifugation and the supernatant was measured for its absorbance at 540 nm by a TECAN M200 plate reader. The assays confirmed that both Cp1D and Cp2D prefer hydrophobic L-amino acids in *N*-succinyl-L-amino acid, while Cp2D preferred aromatic amino acids compared to Cp1D, which is consistent with their native substrate.

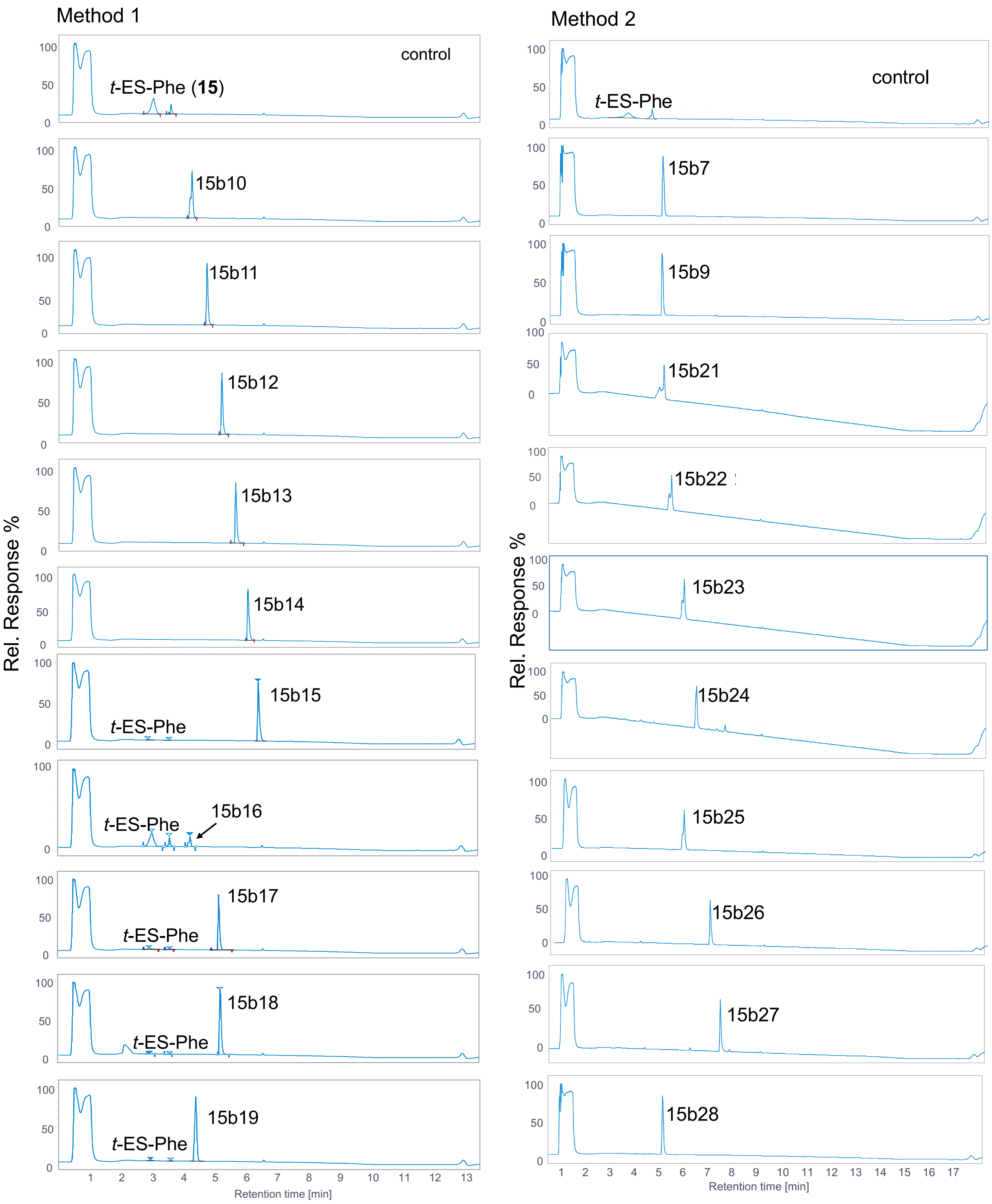

Method 1
control
t-ES-Phe (15)
15b10
15b11
15b12
15b13
15b14
15b15
t-ES-Phe
15b16
t-ES-Phe
15b17
t-ES-Phe
15b18
t-ES-Phe
15b19
t-ES-Phe
Rel. Response %
Retention time [min]
Method 2
control
t-ES-Phe
15b7
15b9
15b21
15b22
15b23
15b24
15b25
15b26
15b27
15b28
Rel. Response %
Retention time [min]

Method 2

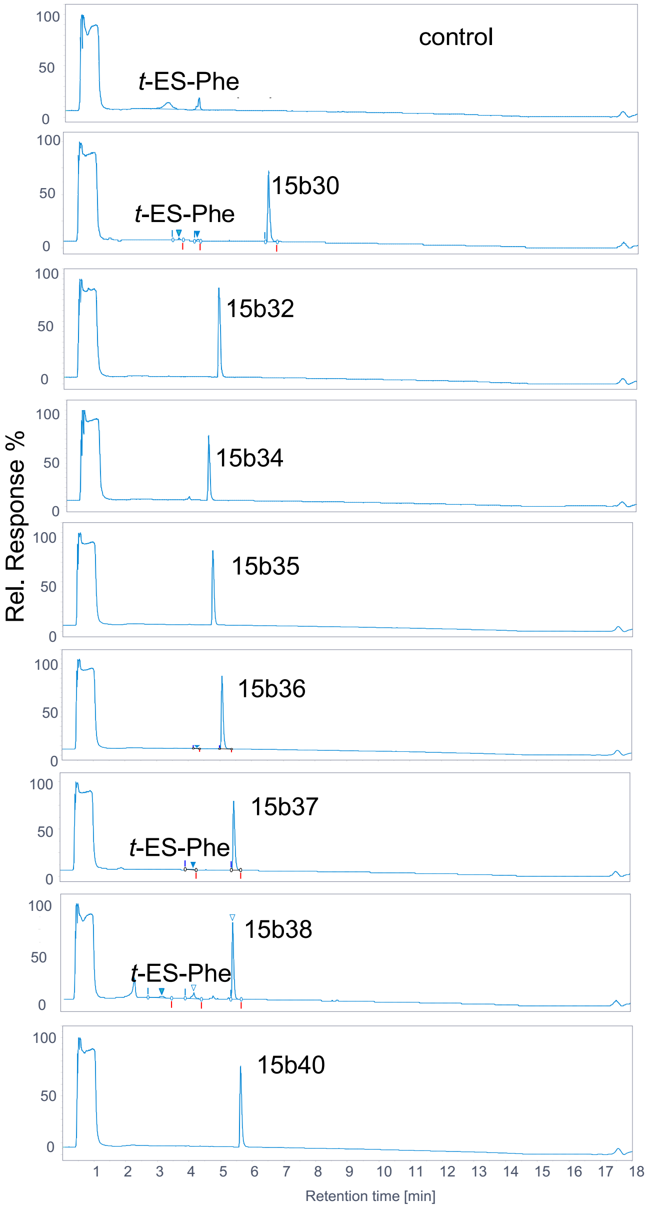

Method 3

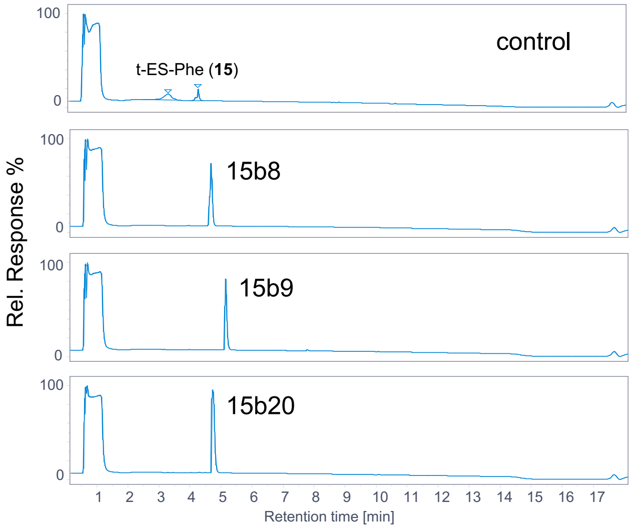

Method 4

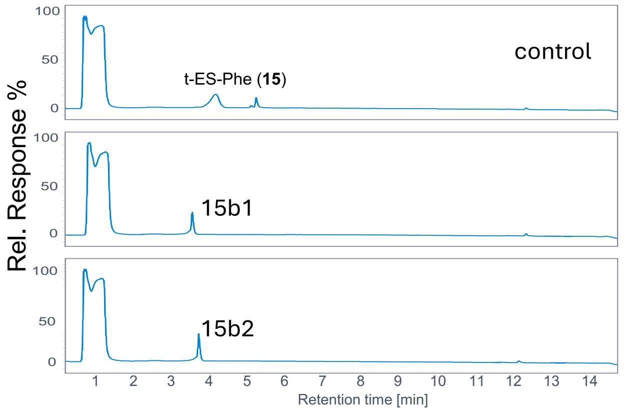

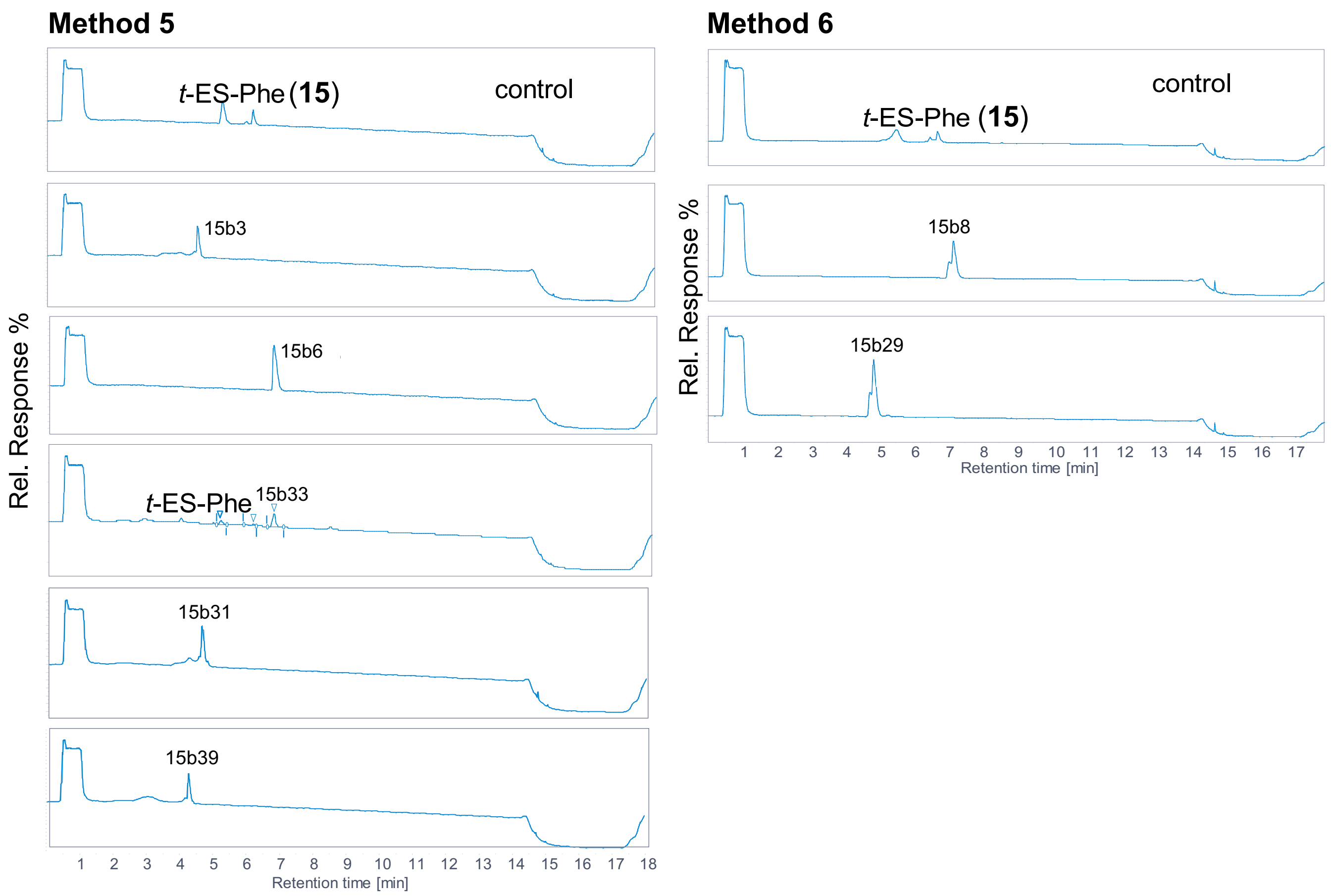

**Supplementary Fig. 27 | HPLC traces from the in vitro reaction of Cp1D with (2*S*,3*S*)-*t*-ES-L-Phe (15) and different amines.** Assays were carried out with enzyme (25 µM) of Cp1D, (2*S*,3*S*)-*t*-ES-L-Phe (**15**) (2 mM) and amine (5 mM). The traces were recorded at λ 204 nm. Cp1D was found to accept a variety of primary amines. The LC/MS analysis using Agilent LC/MSD iQ (AgilentTM InfinityLab Poroshell 120 Aq-C18, 2.7 µm, 100 Å, 2.1 × 100 mm) with elution gradients method, **Method 1**: 0-0.25 min, 1% eluent B; 0.25-8.5 min, 1-99% eluent B; 8.5-11.5 min, 99% eluent B; 11.5-13.5 min, 1% eluent B. **Method 2**: 0-0.25 min, 1% eluent B; 0.25-13.0 min, 1-99% eluent B; 13.0-16.0 min, 99% eluent B; 16.0-18.0 min, 1% eluent B. **Method 3**: 0-0.25 min, 1% eluent B; 0.25-13.0 min, 1-80% eluent B; 13.0-16.0 min, 99% eluent B; 16.0-18.0 min, 1% eluent B. **Method 4**: 0-0.25 min, 1% eluent B; 0.25-13.0 min, 1-60% eluent B; 13.0-16.0 min, 99% eluent B; 16.0-18.0 min, 1% eluent B. **Method 5**: 0-0.25 min, 1% eluent B; 0.25-13.0 min, 1-40% eluent B; 13.0-16.0 min, 99% eluent B; 16.0-18.0 min, 1% eluent B. **Method 6**: 0-0.25 min, 1% eluent B; 0.25-13.0 min, 1-30% eluent B; 13.0-16.0 min, 99% eluent B; 16.0-18.0 min, 1% eluent B.

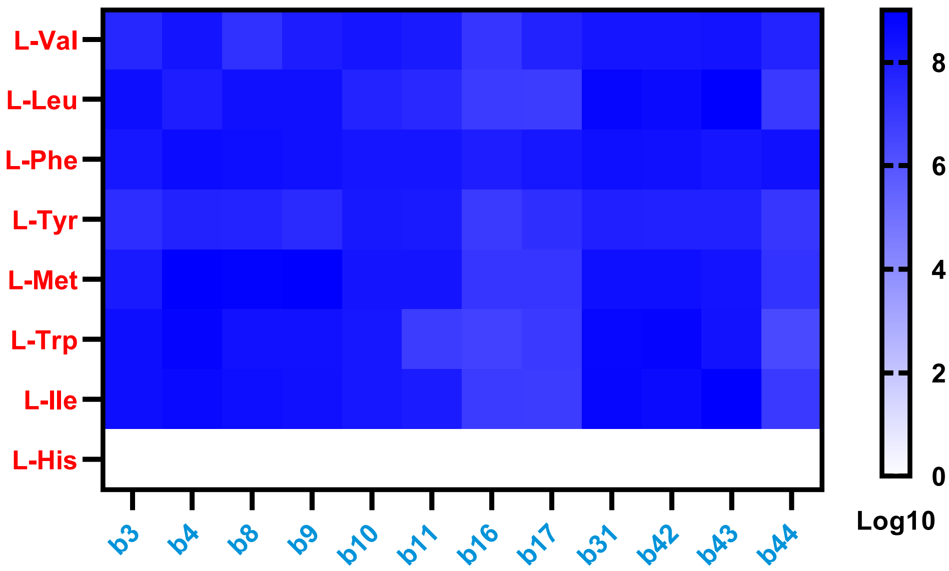

**Supplementary Fig. 28 | LC/MS analysis of one-pot synthesis E-64 analogs by Cp1B and Cp1D in 96-well plate.** Mass area for the corresponding products from ligation of proteinogenic amino acids (Y-axis: V, L, F, Y, M, W, I) with amine donors (X-axis: b3-4, b8-11, b16-17, b31, b42 (histamine), b43 (4-(2-aminoethyl)-pyridine), and b44 (tryptamine)) is shown as the heat map. Assays were carried out with enzymes Cp1B and Cp1D (25 μM), 2 mM (±)-*trans*-epoxy-succinate, 1 mM L-amino acid, and 1 mM amine at 30 °C. The production of the corresponding products was determined by LC/MS analysis following 16 h incubation.

**a**) E-64 synthesis

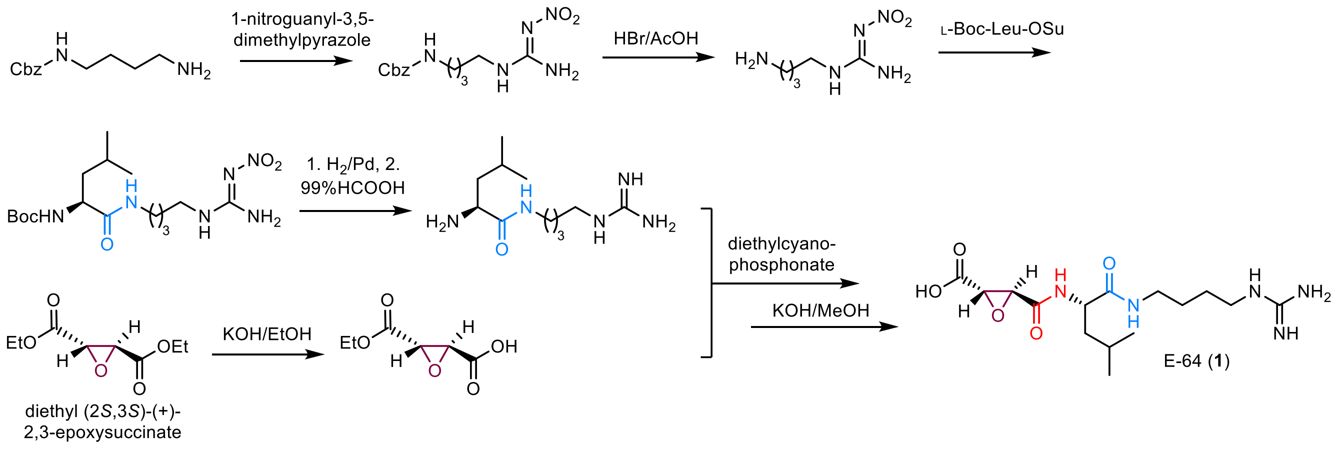

**b**) CLIK-148 synthesis

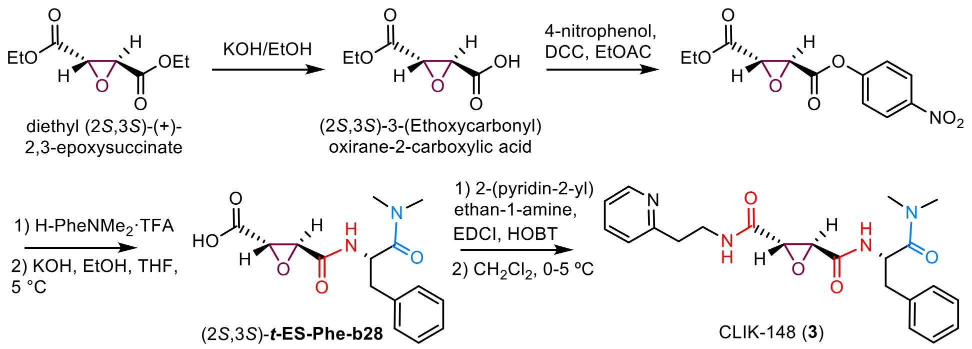

**Supplementary Fig. 29 | Reported chemical synthesis for E-64 (a)[25] and CLIK-148 (b)[6].**

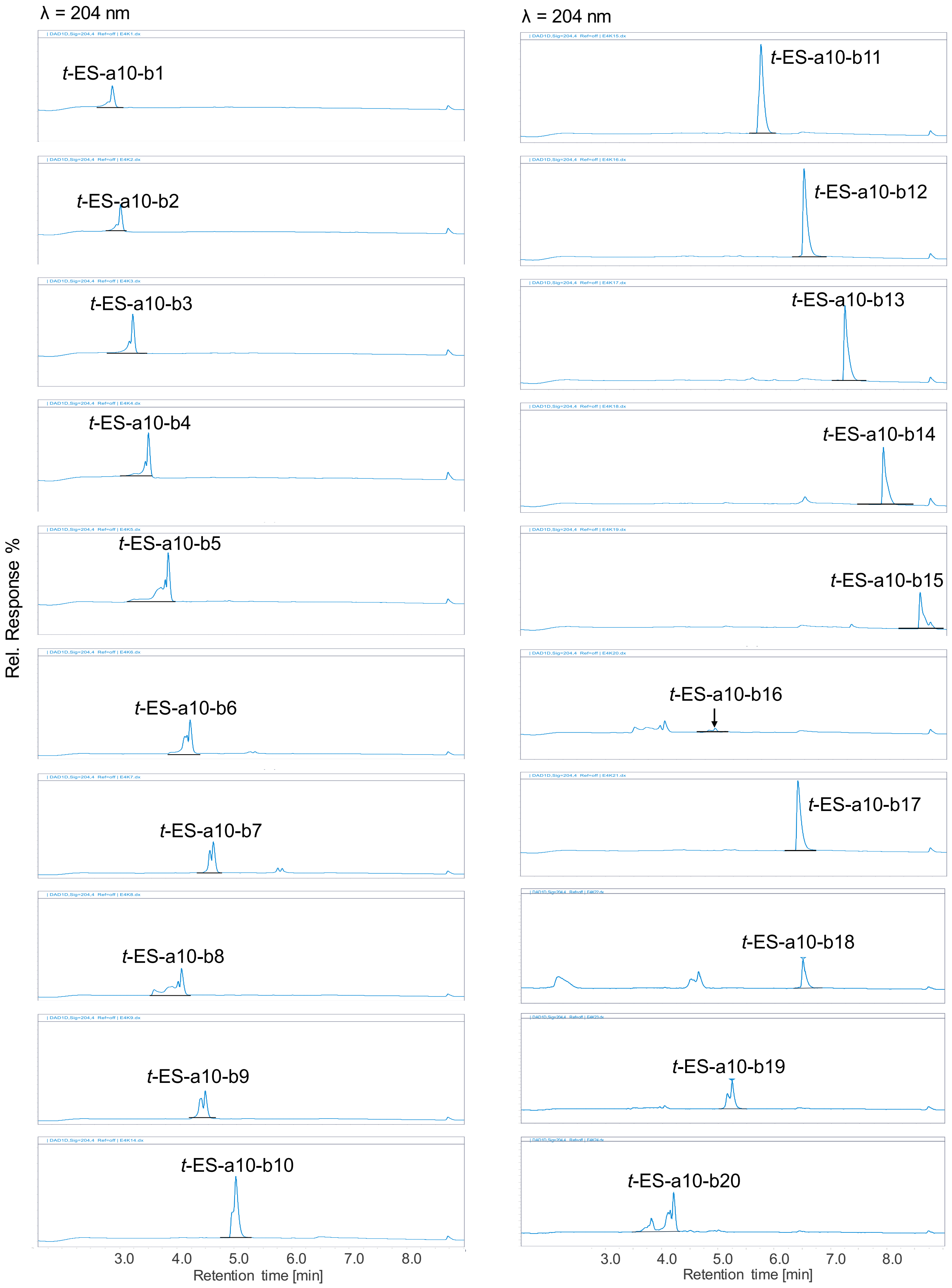

λ = 204 nm
λ = 204 nm
t-ES-a10-b1
t-ES-a10-b2
t-ES-a10-b3
t-ES-a10-b4
t-ES-a10-b5
t-ES-a10-b6
t-ES-a10-b7
t-ES-a10-b8
t-ES-a10-b9
t-ES-a10-b10
t-ES-a10-b11
t-ES-a10-b12
t-ES-a10-b13
t-ES-a10-b14
t-ES-a10-b15
t-ES-a10-b16
t-ES-a10-b17
t-ES-a10-b18
t-ES-a10-b19
t-ES-a10-b20
Rel. Response %
3.0 4.0 5.0 6.0 7.0 8.0
Retention time [min]
3.0 4.0 5.0 6.0 7.0 8.0
Retention time [min]

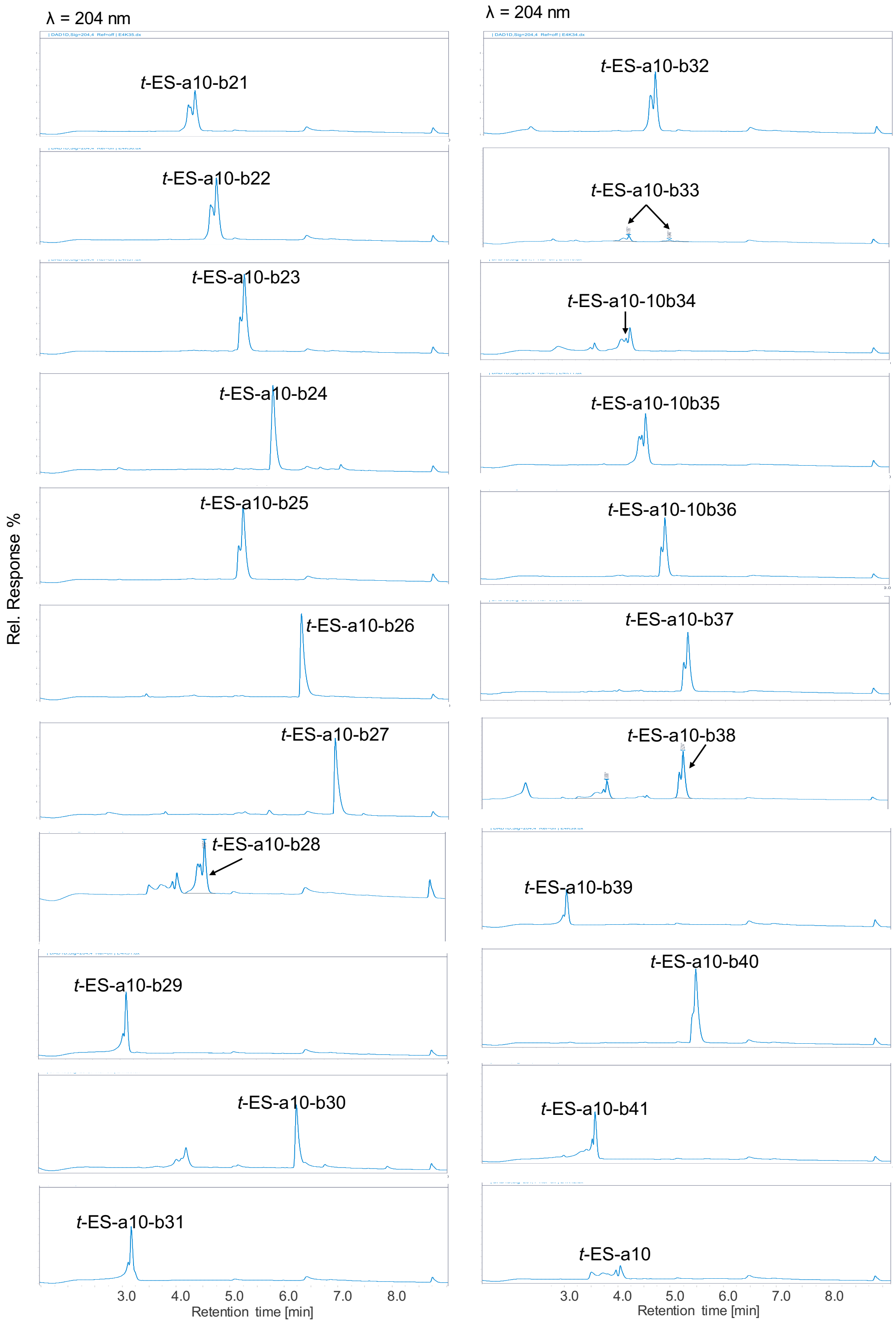

**Supplementary Fig. 30 | HPLC traces of combinatorial biocatalysis with a10 and different amines.** Assays were carried out with Cp1B (25 µM), and Cp1D (25 µM), 2 mM (±)-*trans*-epoxy-succinate, 1 mM **a10**, 1 mM amine at 30 ºC for 16 h. The LC/MS elution gradients method: 0-0.25 min, 1% eluent B; 0.25-13.0 min, 1-99% eluent B; 13.0-16.0 min, 99% eluent B; 16.0-18.0 min, 1% eluent B using Agilent LC/MSD iQ (AgilentTM InfinityLab Poroshell 120 Aq-C18, 2.7 µm, 100 Å, 2.1 × 100 mm).

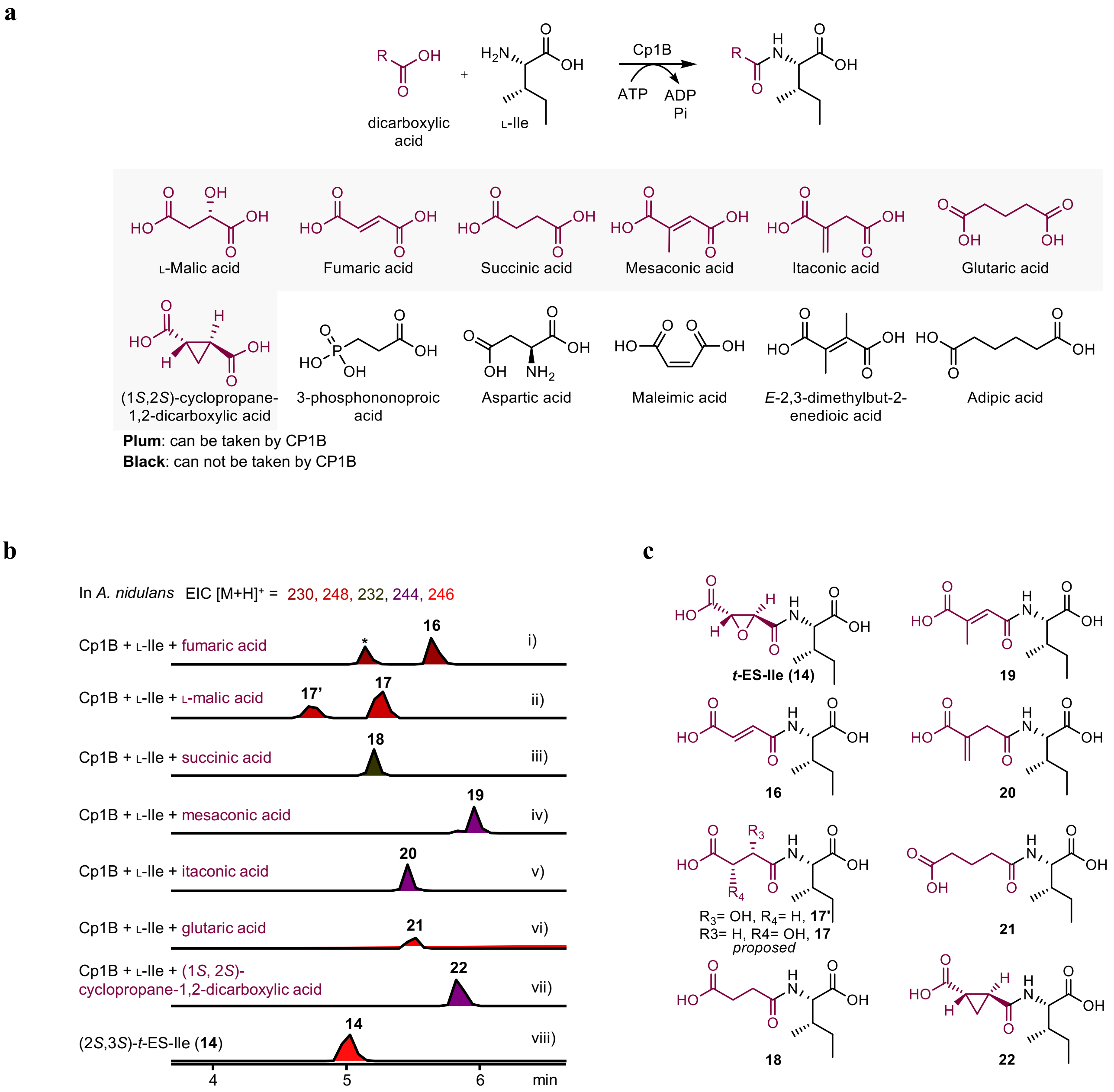

**Supplementary Fig. 31 | Dicarboxylic acid scope for Cp1B. a**. Structure of dicarboxylic acids tested with Cp1B. Note: dicarboxylic acids which can be taken by Cp1B are highlighted in plum while dicarboxylic acids which cannot be taken by Cp1B are colored in black. **b**. LC/MS analysis of the predicted product from the ligation of various dicarboxylic acids with L-Ile. The products were detected when L-malic acid, succinic acid, fumaric acid, mesaconic acid, itaconic acid, (1*S*,2*S*)-*trans*-cyclopropane-1,2-dicarboxylic acid, and glutaric acid were used as the acid acceptor. *the parent MS is different with **16**. **c**. Predicted structures of corresponding products.

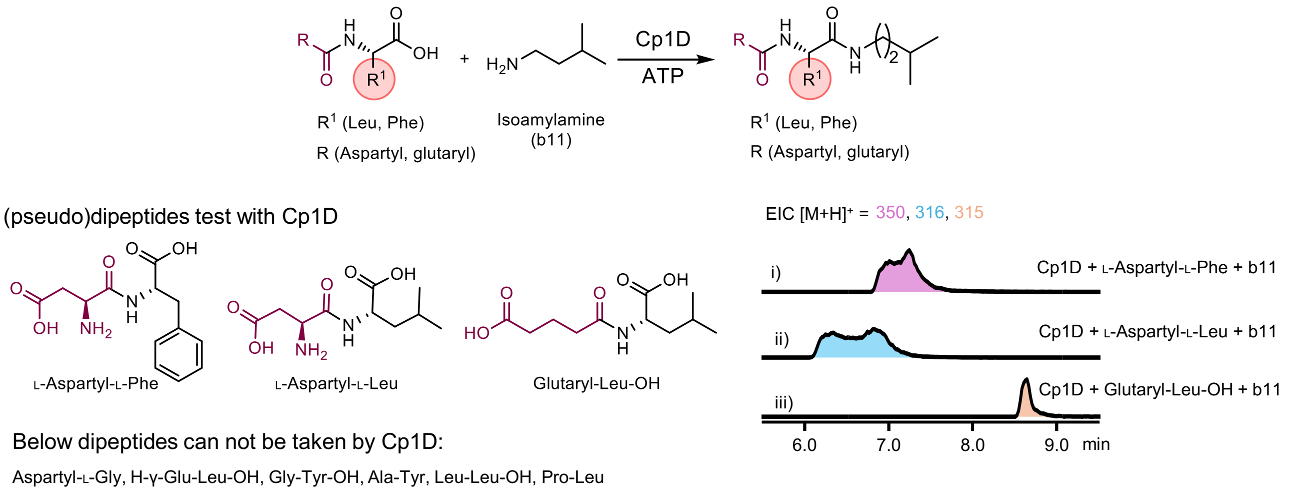

**Supplementary Fig. 32 | Structures of (pseudo)dipeptides tested for the acid acceptor of Cp1D**. LC/MS analysis of in vitro reaction of Cp1D with the (pseudo)dipeptides and isoamylamine (isopentylamine). EICs for L-Aspartyl-L-Phe, L-Aspartyl-L-Leu, and Glutaryl-L-Leu are $[M+H]^+ = 350$, $[M+H]^+ = 316$, and $[M+H]^+ = 315$, respectively.

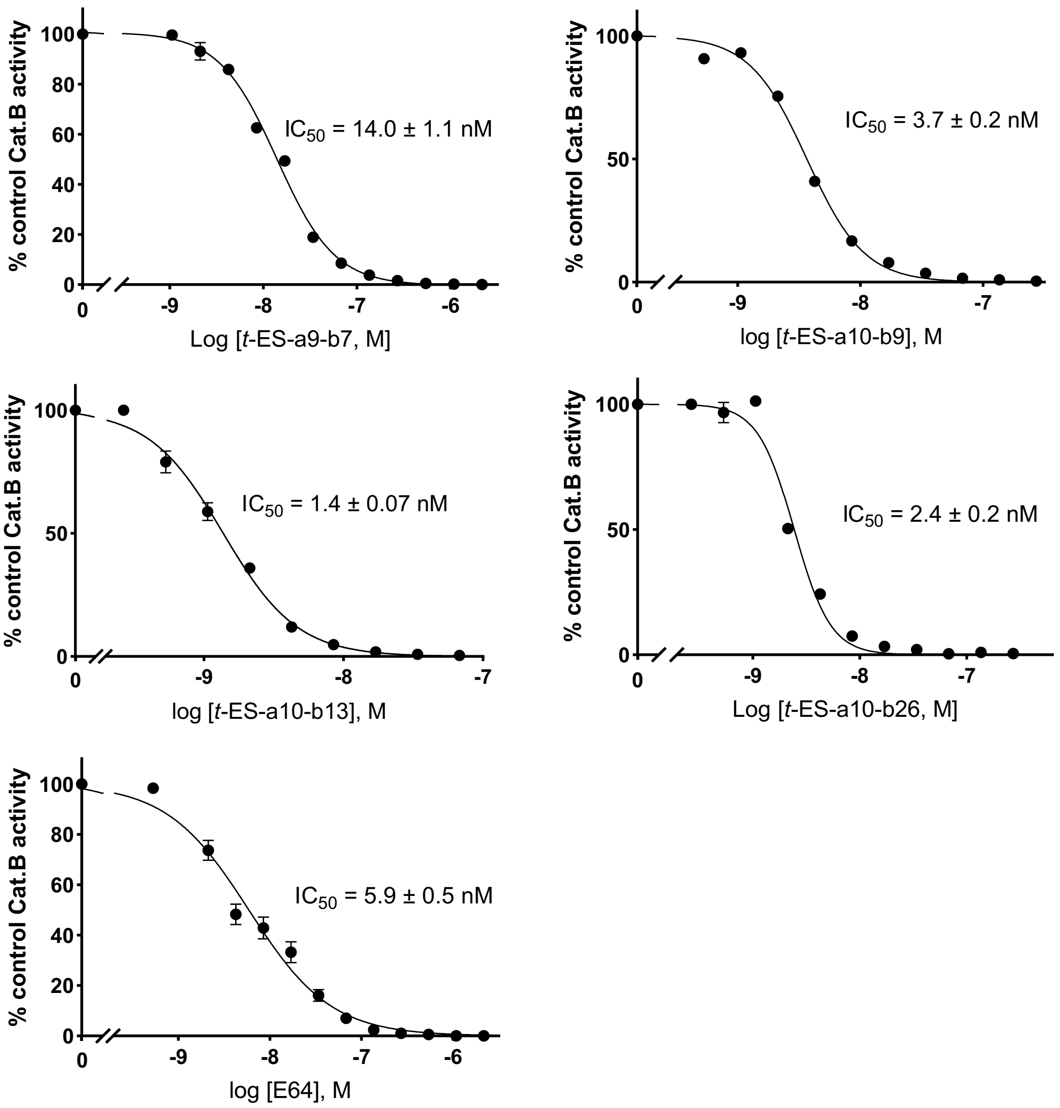

**Supplementary Fig**. **33 |** In vitro cathepsin B inhibition assay of enzymatic synthesized inhibitors.

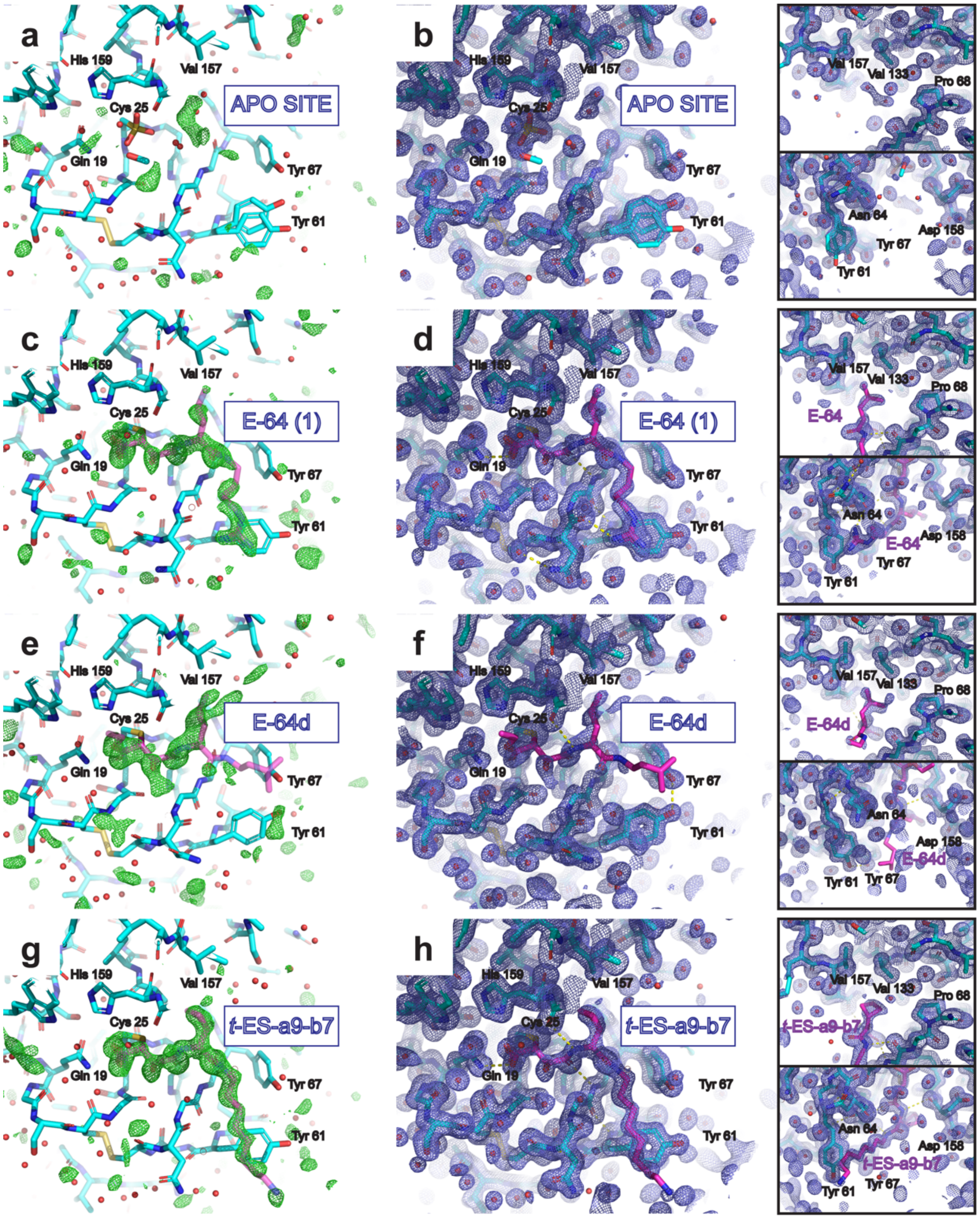

**Supplementary Fig. 34 | Structures of the active site of papain (no inhibitor bound, panels a-b) and papain bound to E-64 analogs from X-ray diffraction (panels c-h)**. For the unliganded active site, atomic coordinates are superimposed on a difference Fourier map at 3σ following complete modeling and refinement (**a**), and the electron density map generated following refinement at 1σ (**b**). For the E-64-bound active site, atomic coordinates are superimposed on a ligand-omit difference Fourier map at 3σ, revealing positive density that E-64 (**1**) (translucent magenta model) was modeled into (**c**), and the electron density map for the refined structure with the ligand included at 1σ (**d**). The same is shown for the papain-E64-d active site (**e-f**) and the papain-(2*S*,3*S*)-***t*-ES-a9-b7** active site (**g-h**). Insets for each structure highlight a hydrophobic pocket adjacent to the active site occupied by hydrophobic side-chains of each ligand (top inset), and the solvent-facing region adjacent to the active site occupied by each inhibitor's tail (bottom inset). Yellow dashed lines indicate potential hydrogen-bonding interactions.

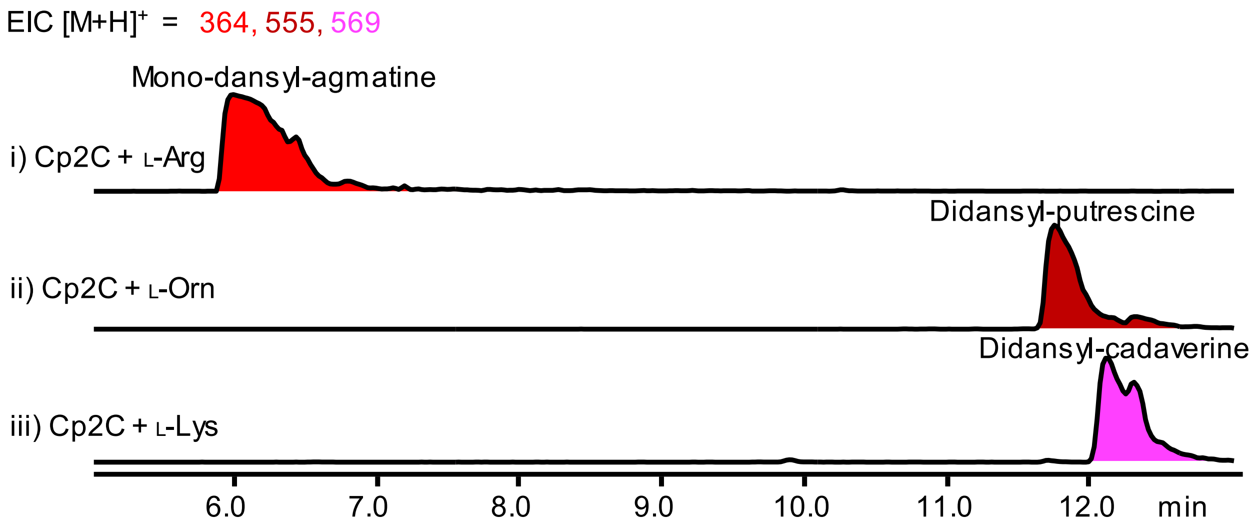

**Supplementary Fig. 35 | In vitro assay of Cp2C with L-Ornithine, L-Arginine and L-Lysine.** The formation of corresponding decarboxylated product agmatine (mono-dansyl-agmatine, $[M+H]^+ = 364$), putrescine (didansyl-putrescine, $[M+H]^+ = 555$), and cadaverine (didansyl-cadaverine, $[M+H]^+ = 569$) was observed.

**a**

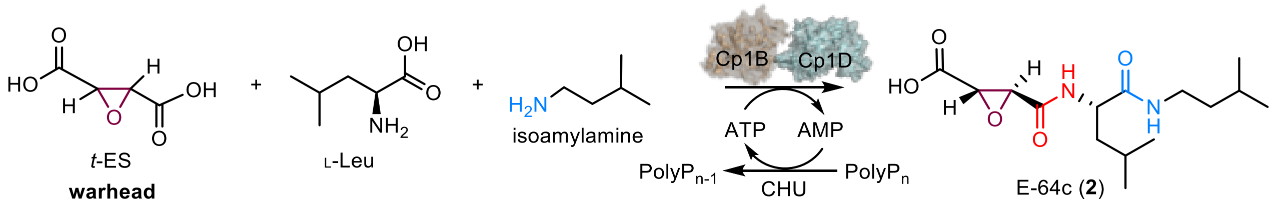

**b**

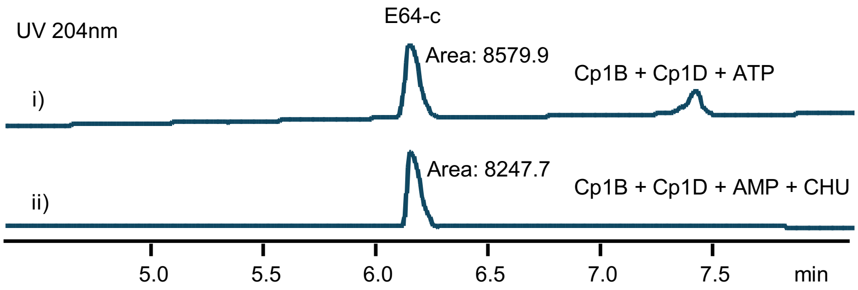

**Supplementary Fig. 36. Preparative-scale synthesis of E-64c using ATP Regeneration system**. **a**, Scheme for E-64c (**2**) from (±)-*t*-ES, L-Leu, and isoamylamine (isopentylamine) catalyzed by Cp1B and Cp1D, with recycling the ATP using the Polyphosphate kinase (CHU)[26] and polyphosphate ($PolyP_n$). **b**, Preparative-scale synthesis of E-64c with addition of 10 mM ATP (i) or using ATP Regeneration system (ii). Reaction conditions for i): 5 mM (±)-*t*-ES, 2.5 mM L-Leu, 5 mM isoamylamine, 10 mM ATP, 2.5 µM Cp1B, and 2.5 µM Cp1D. Reaction conditions for ii): 5 mM (±)-*t*-ES, 2.5 mM L-Leu, 5 mM isoamylamine, 10 mM AMP, 20 mg $mL^{-1}$ $PolyP_n$, 2.5 µM Cp1B, 2.5 µM Cp1D, 50 µM CHU. Incubation at 30°C for 48 h. A comparable production of E-64c formation was observed.

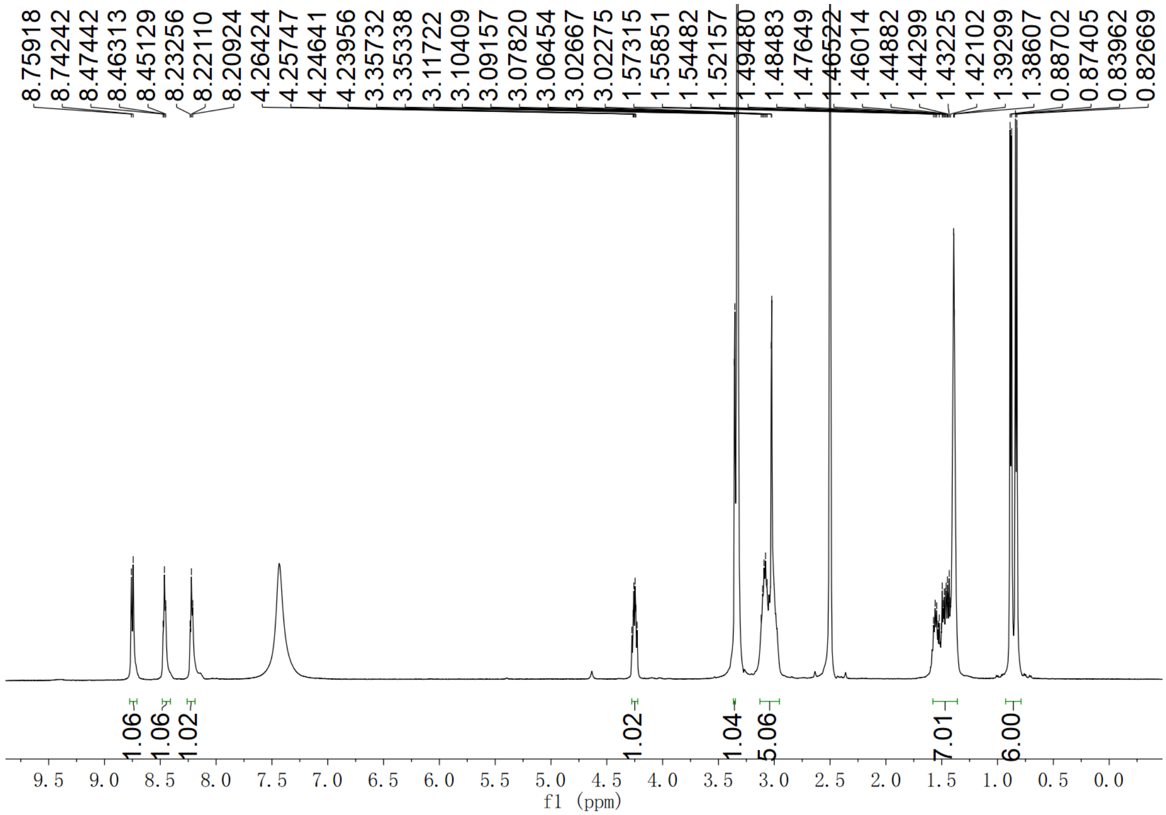

**Supplementary Fig. 37.** $^{1}$**H NMR spectrum of E-64 (1) in DMSO-$d_6$**

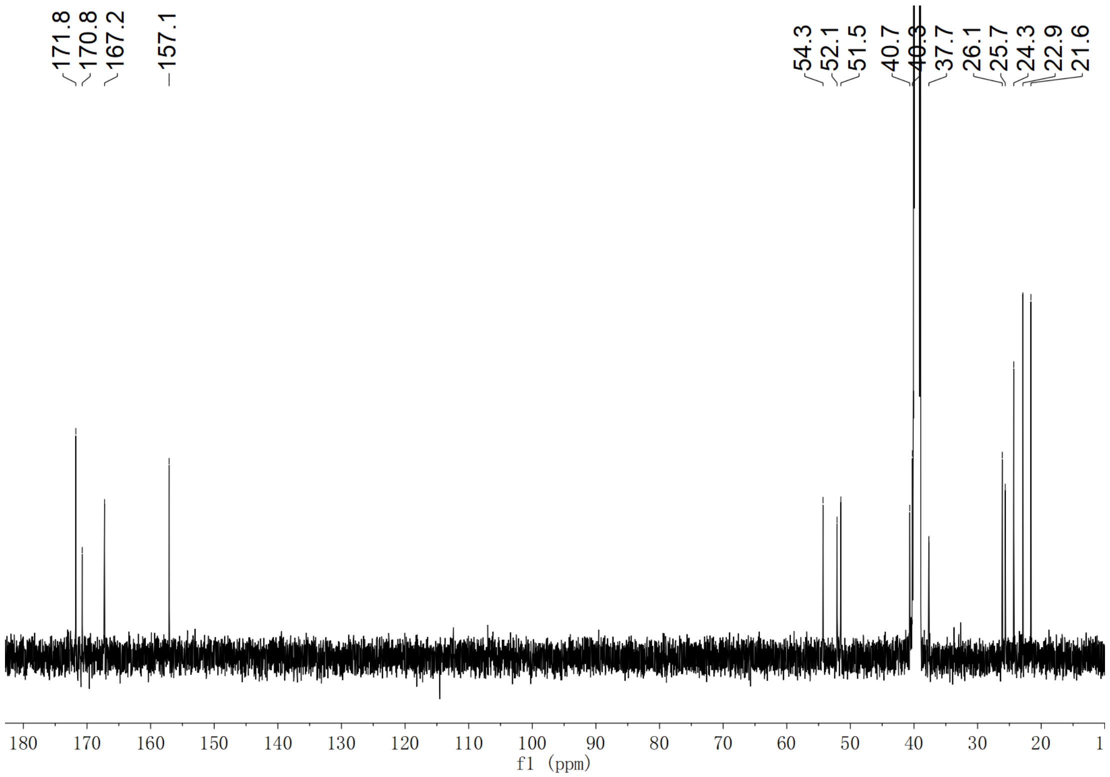

**Supplementary Fig. 38.** $^{13}$**C NMR spectrum of E-64 (1) in DMSO-$d_6$**

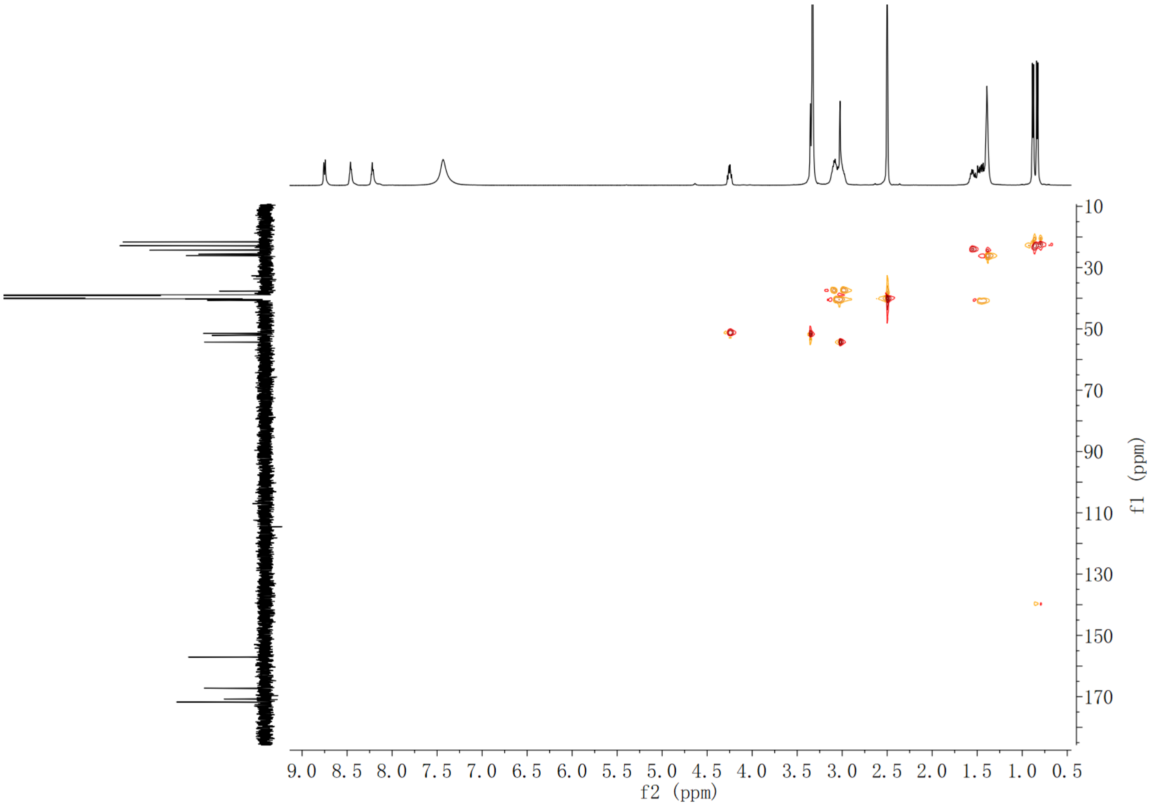

**Supplementary Fig. 39. HSQC spectrum of E-64 (1) in DMSO-$d_6$**

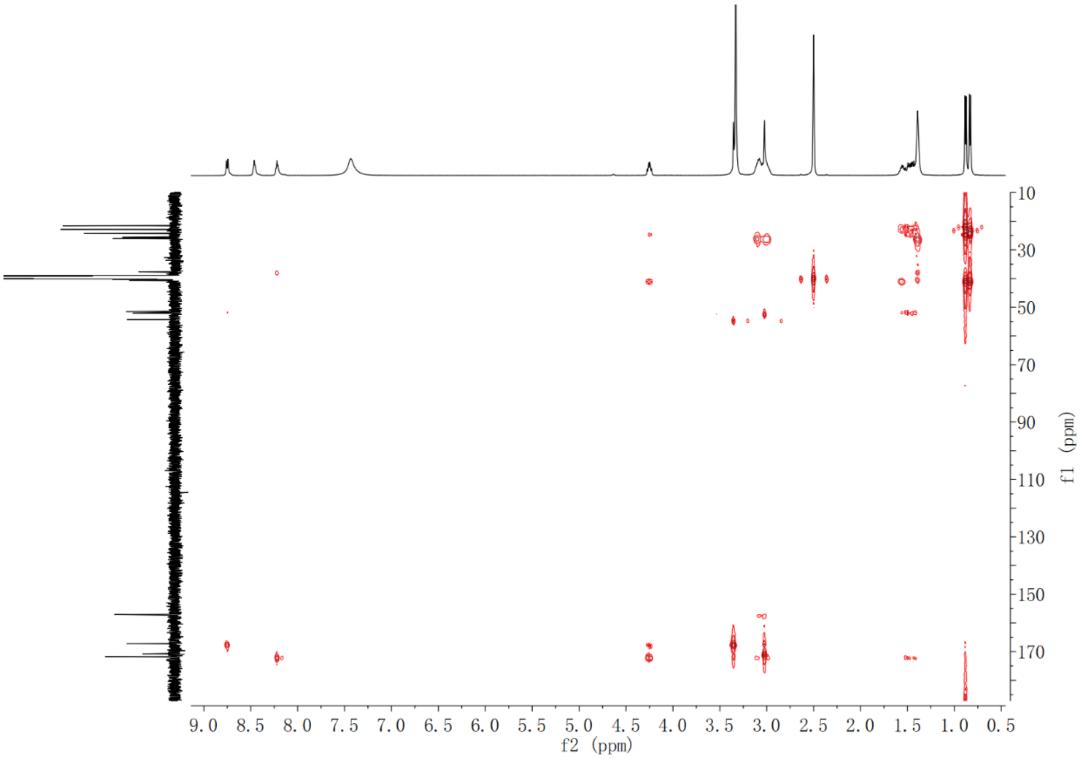

**Supplementary Fig. 40. HMBC spectrum of E-64 (1) in DMSO-$d_6$**

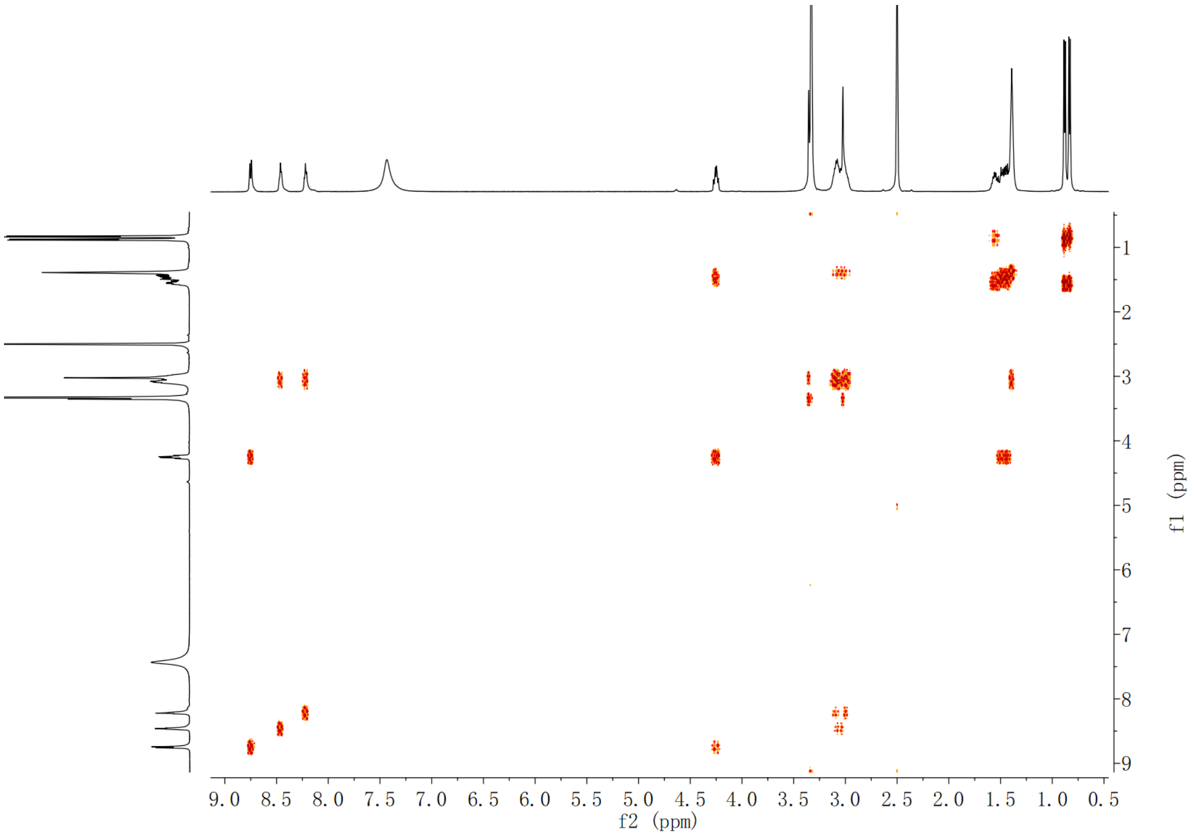

**Supplementary Fig. 41.** $^{1}$**H-**$^{1}$**H COSY spectrum of E-64 (1) in DMSO-*d*$_6$**

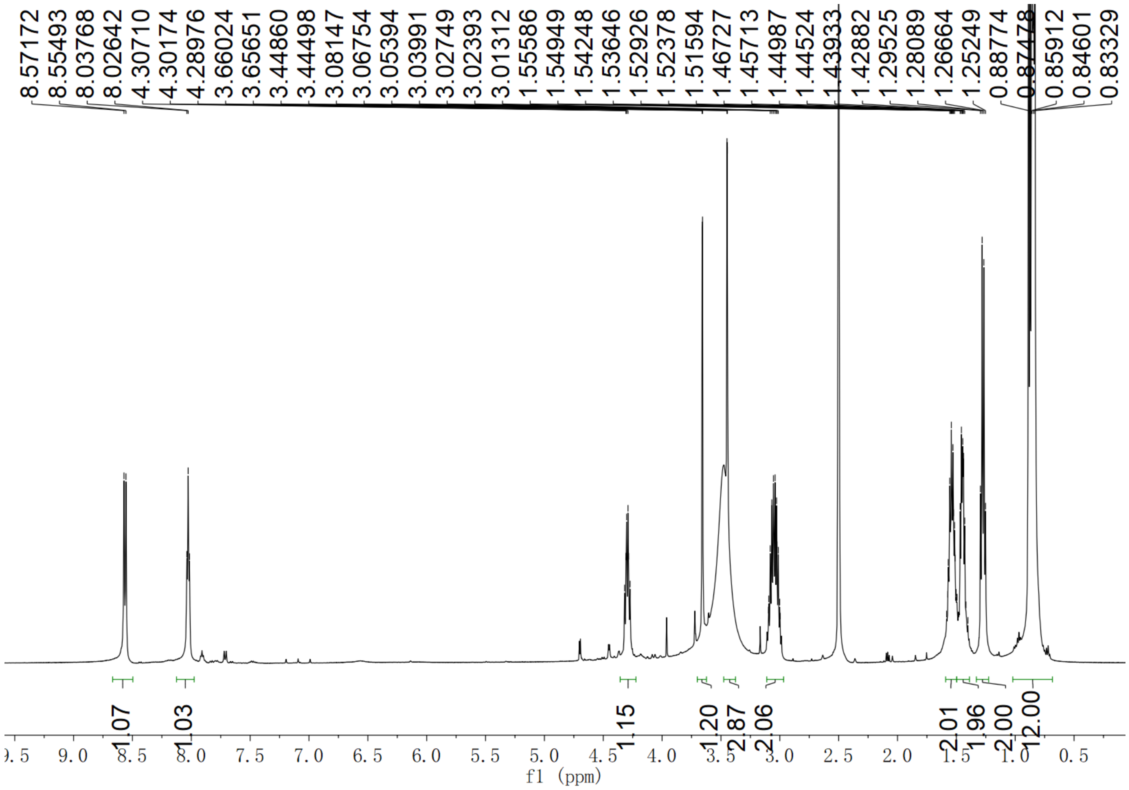

**Supplementary Fig. 42.** $^{1}$**H NMR spectrum of E-64c (2) in DMSO-*d*$_6$**

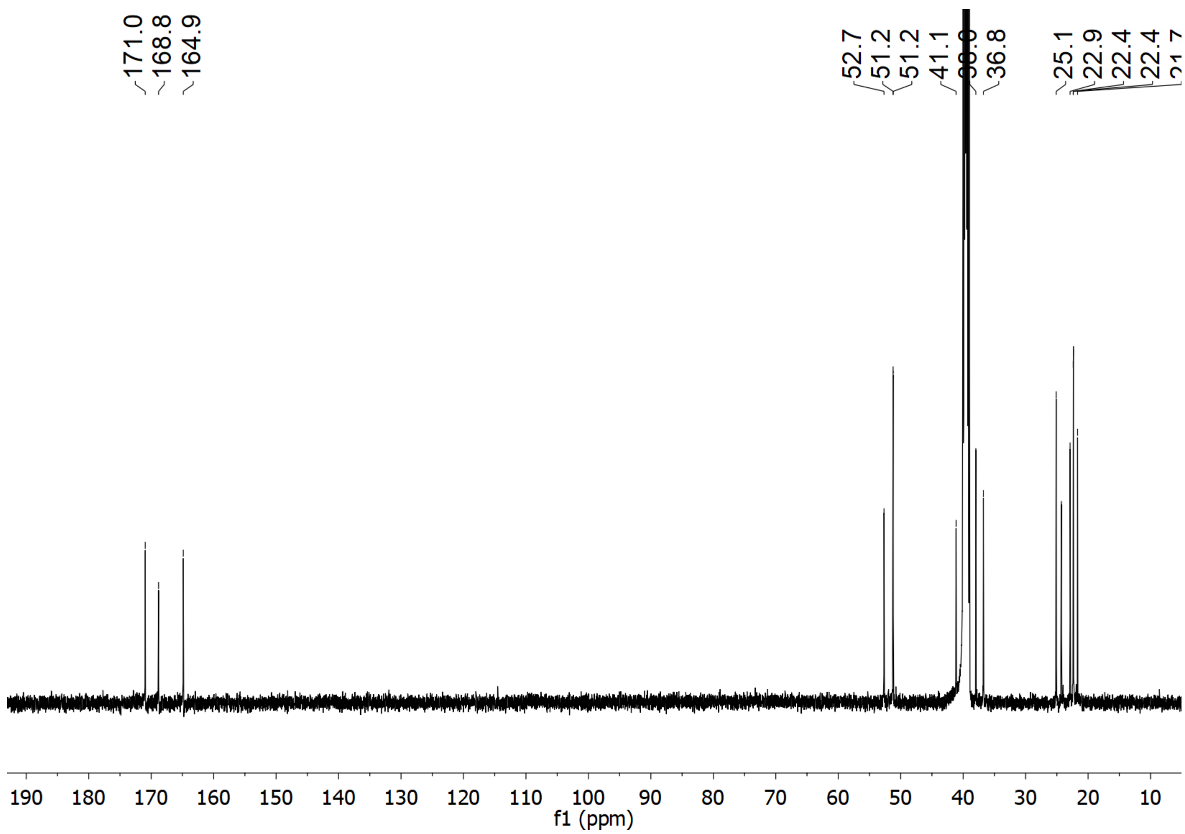

Supplementary Fig. 43. $^{13}C$ NMR spectrum of E-64c (2) in DMSO-$d_6$

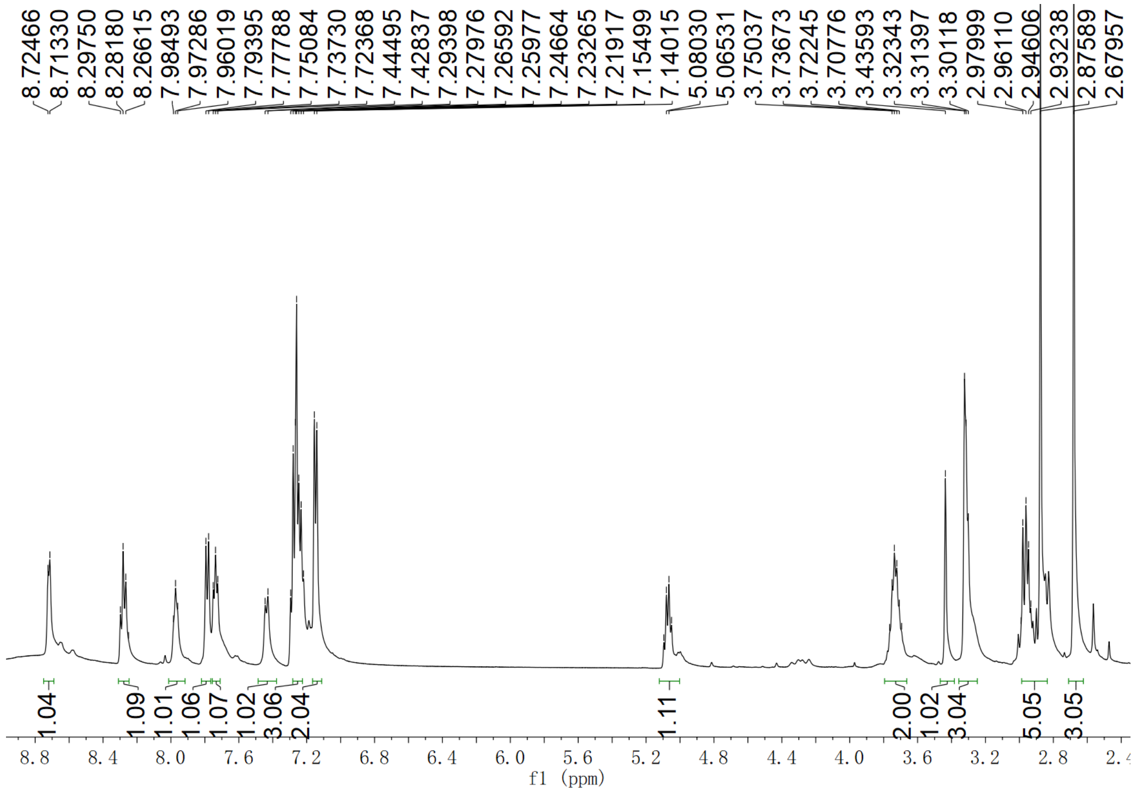

Supplementary Fig. 44. $^{1}H$ NMR spectrum of CLIK148 (3) in $CDCl_3$

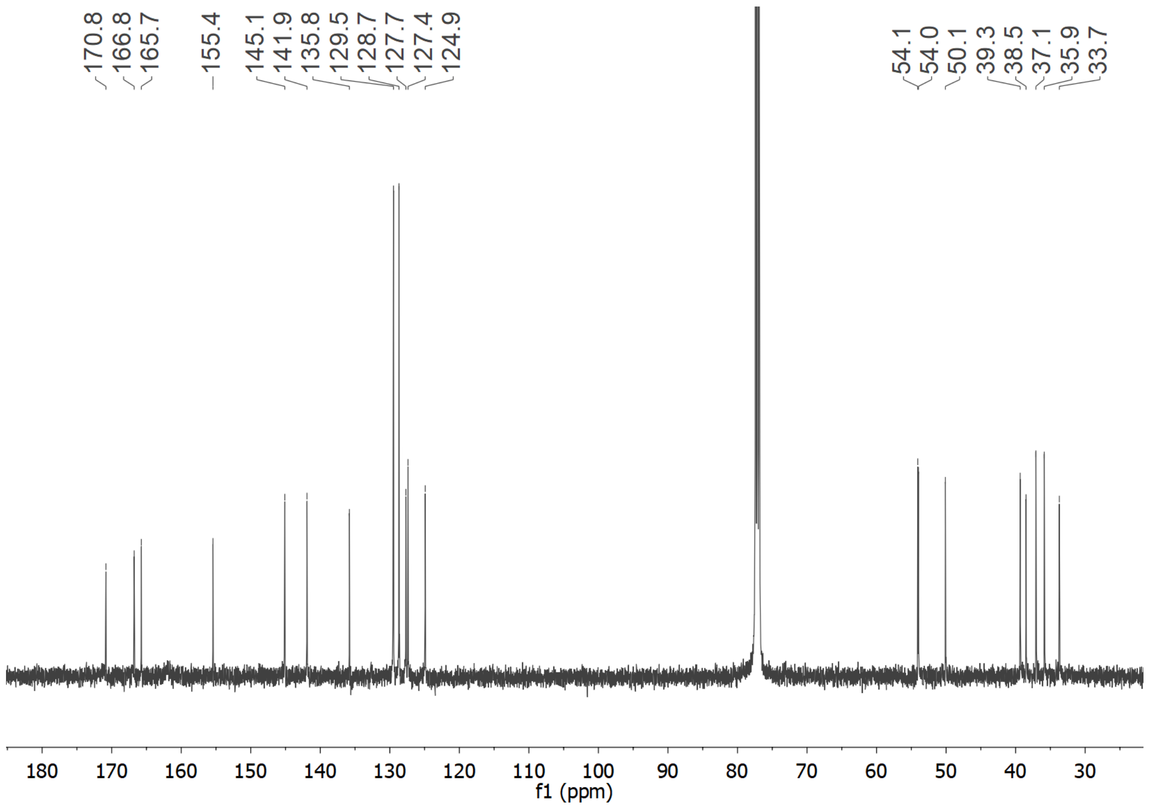

**Supplementary Fig. 45. $^{13}$C NMR spectrum of CLIK148 (3) in $CDCl_3$**

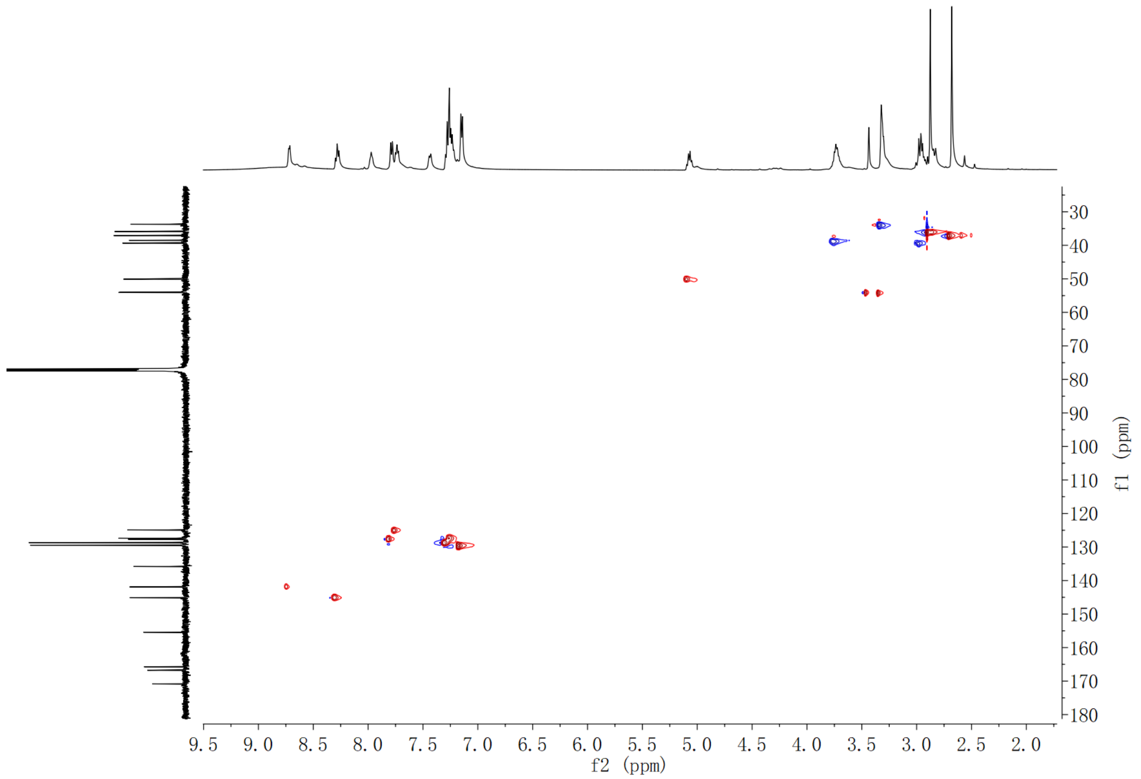

**Supplementary Fig. 46. HSQC spectrum of CLIK148 (3) in $CDCl_3$**

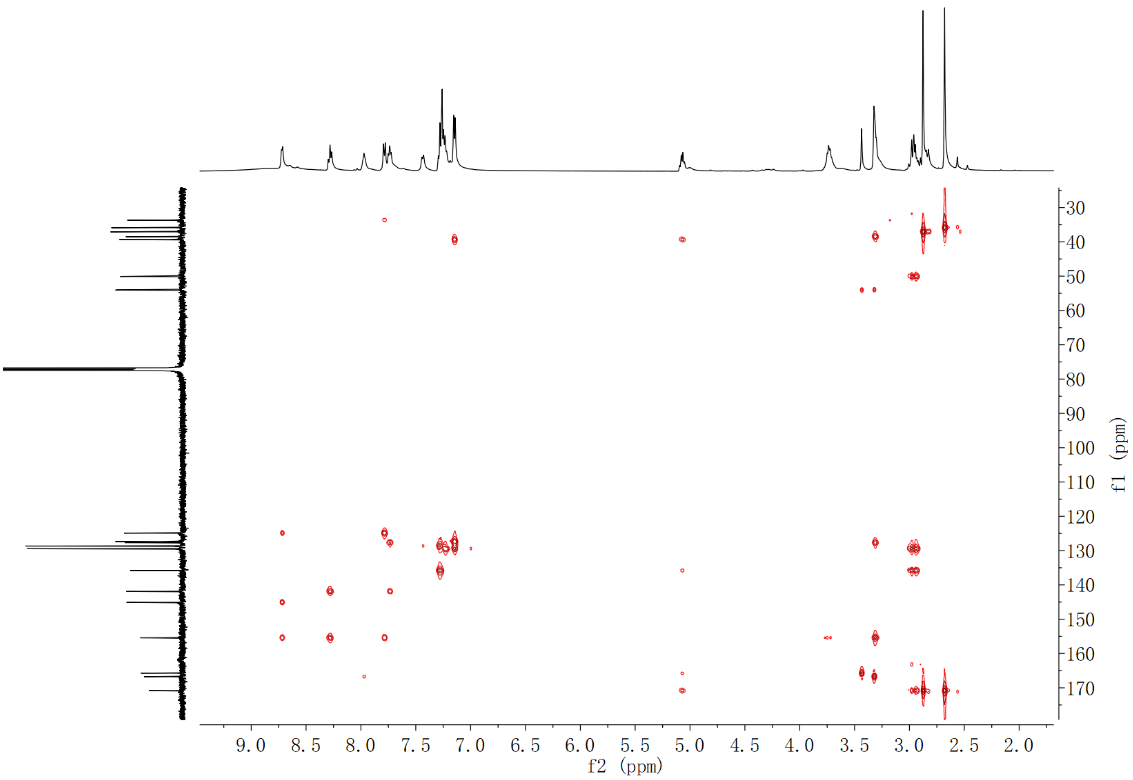

**Supplementary Fig. 47. HMBC spectrum of CLIK148 (3) in $CDCl_3$**

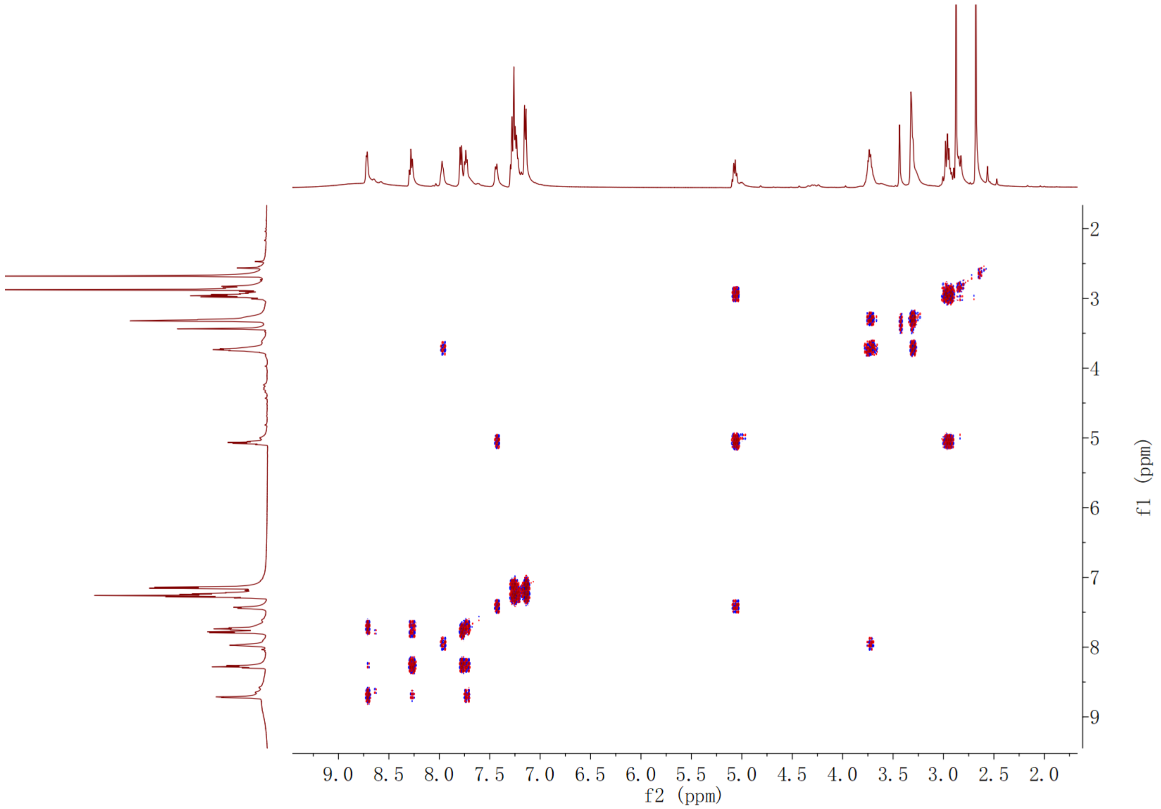

**Supplementary Fig. 48. $^1H$-$^1H$ COSY spectrum of CLIK148 (3) in $CDCl_3$**

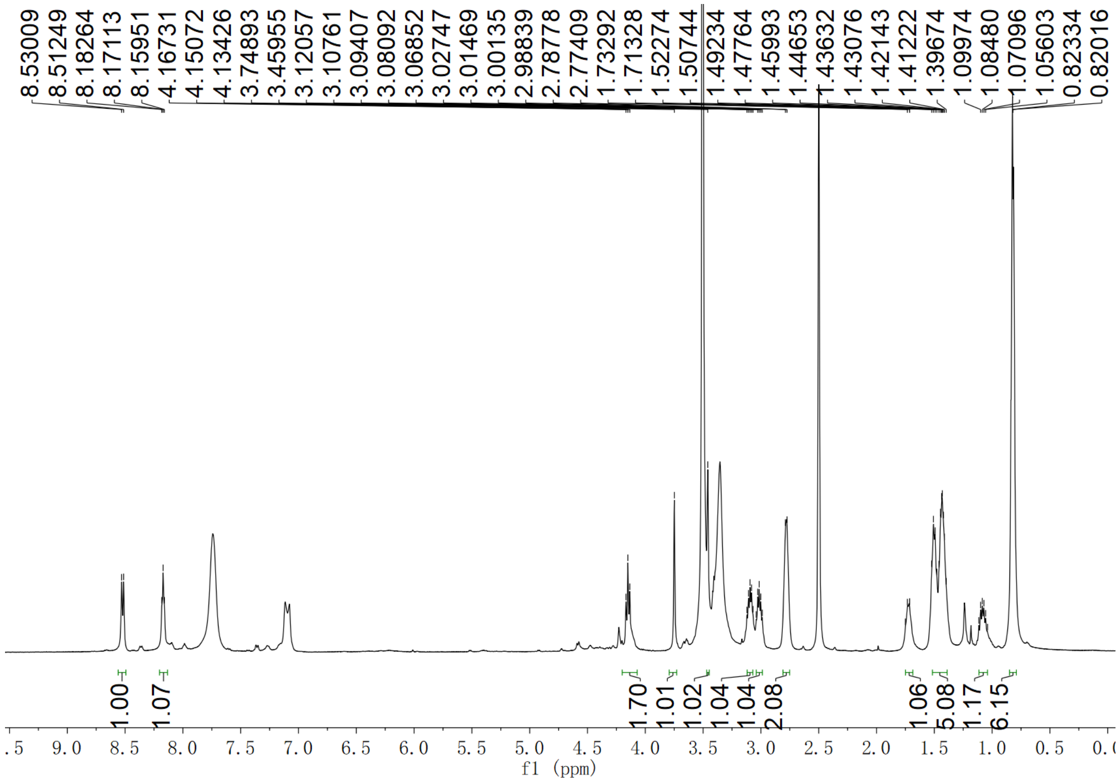

**Supplementary Fig. 49. $^{1}$H NMR spectrum of CPI-2 (4) in DMSO-$d_6$**

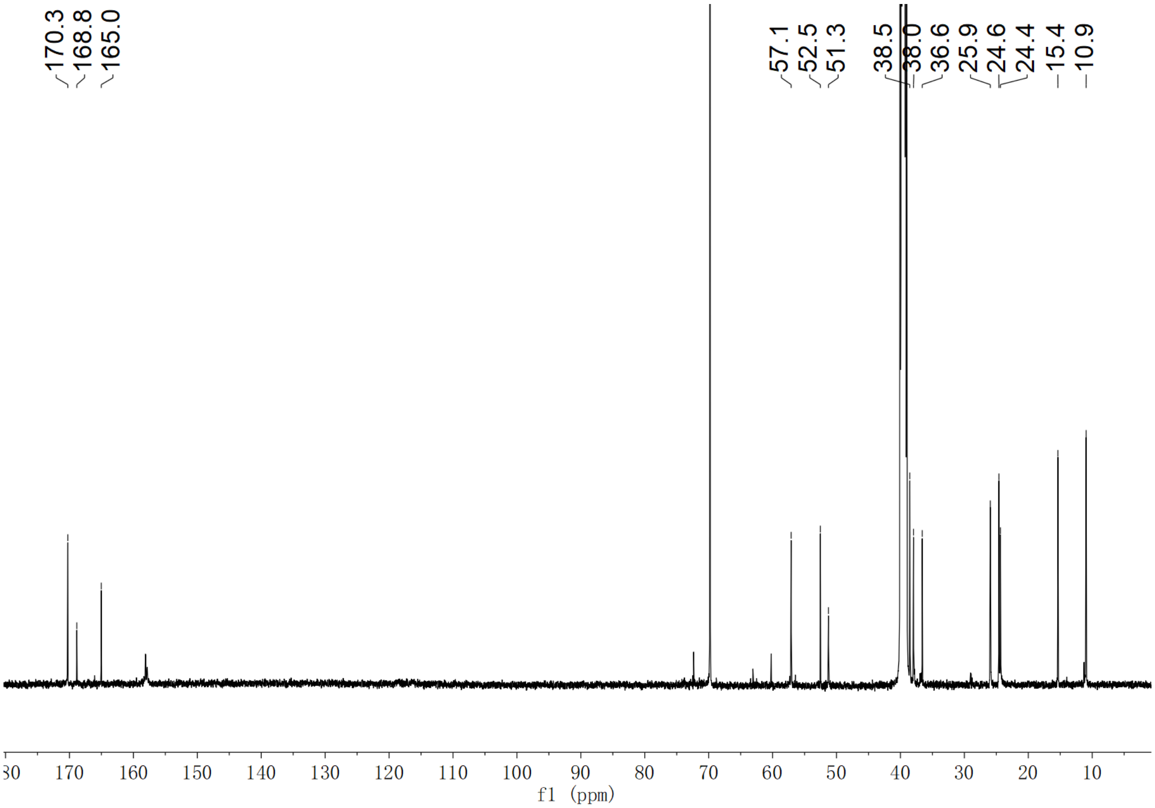

**Supplementary Fig. 50. $^{13}$C NMR spectrum of CPI-2 (4) in DMSO-$d_6$**

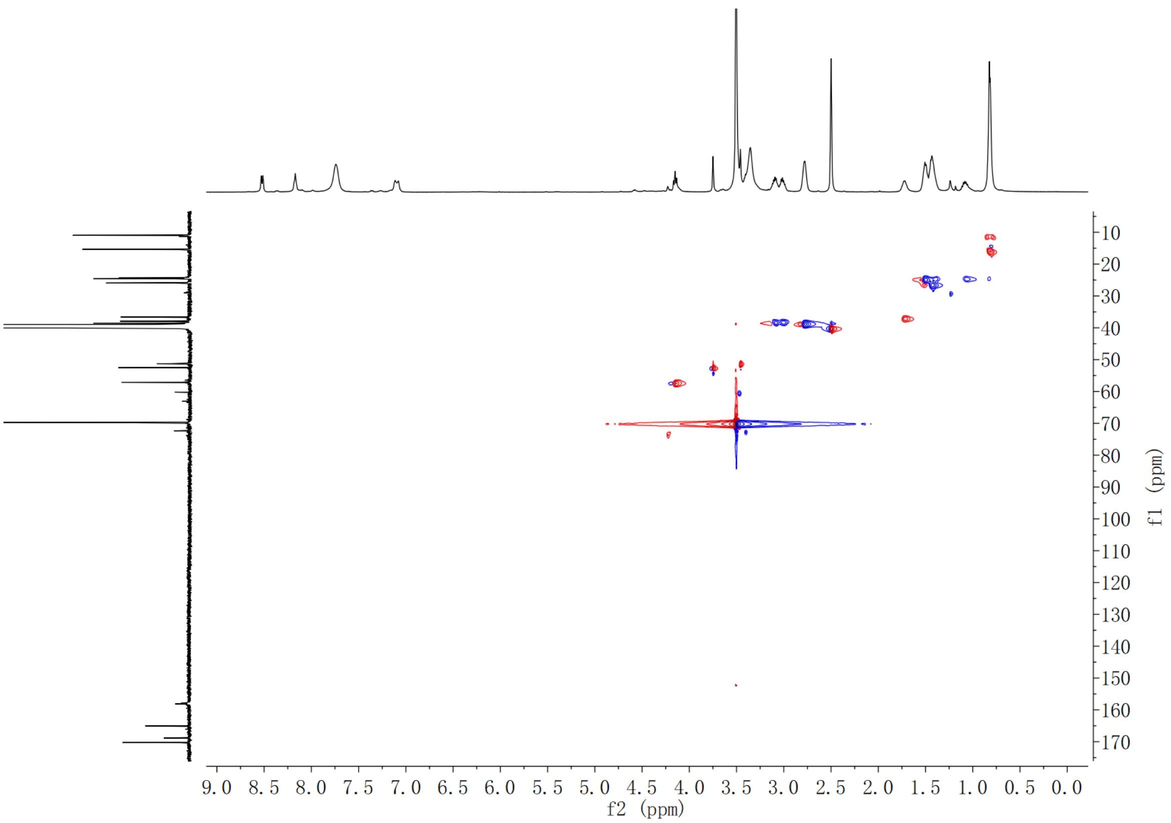

**Supplementary Fig. 51. HSQC spectrum of CPI-2 (4) in DMSO-*d*$_6$**

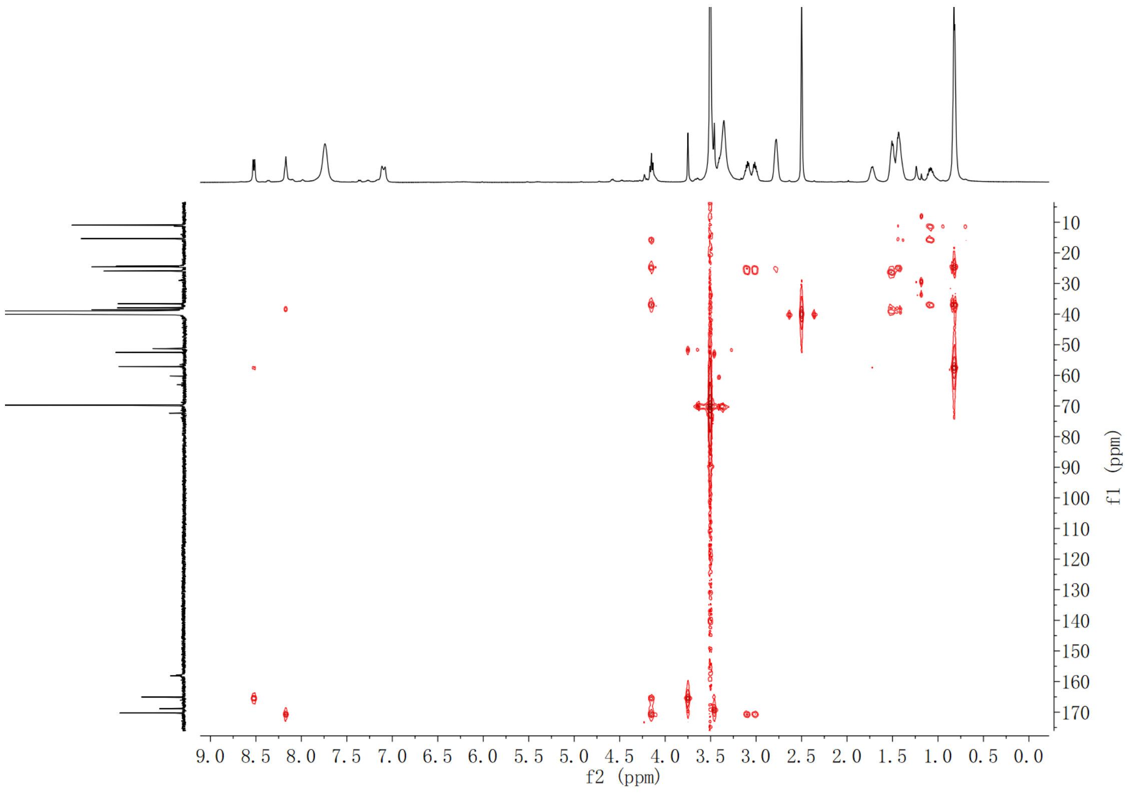

**Supplementary Fig. 52. HMBC spectrum of CPI-2 (4) in DMSO-*d*$_6$**

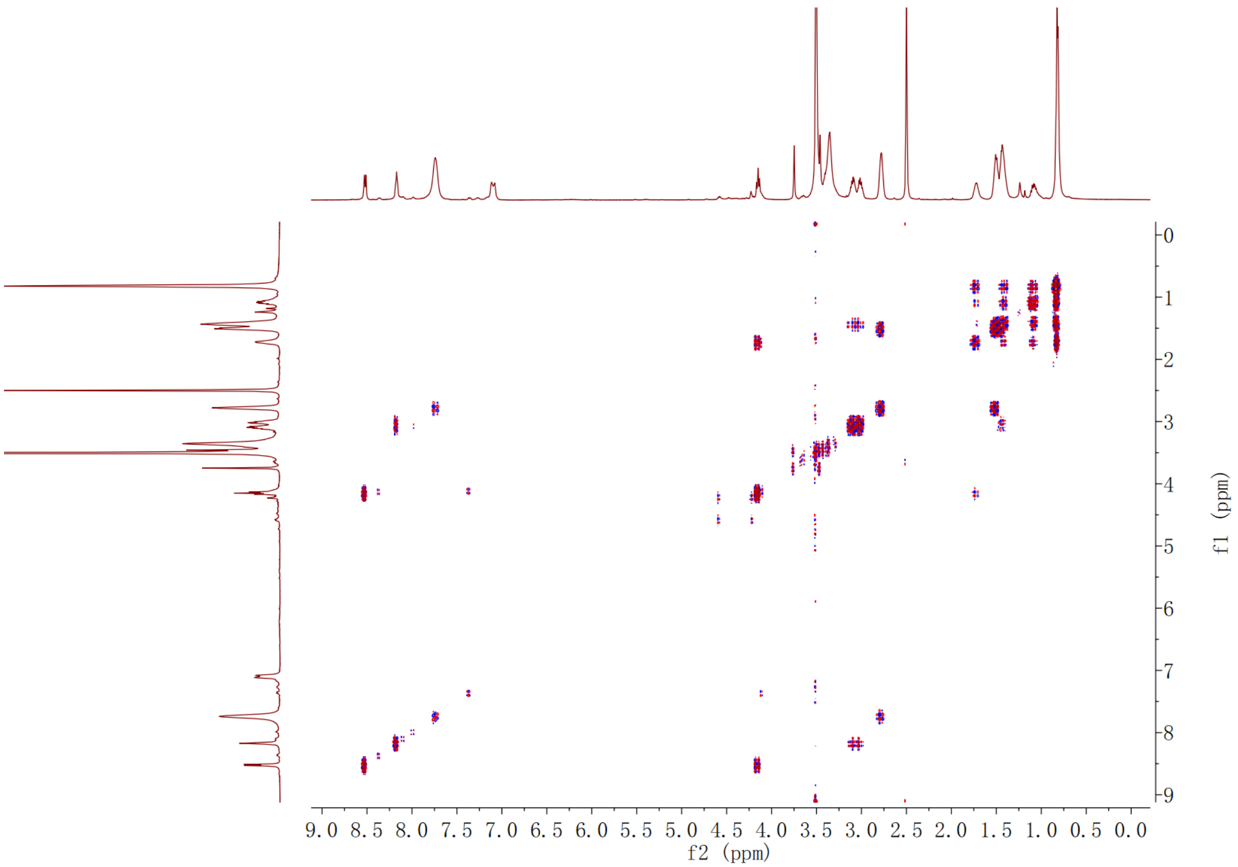

**Supplementary Fig. 53. $^{1}$H-$^{1}$H COSY spectrum of CPI-2 (4) in DMSO-$d_6$**

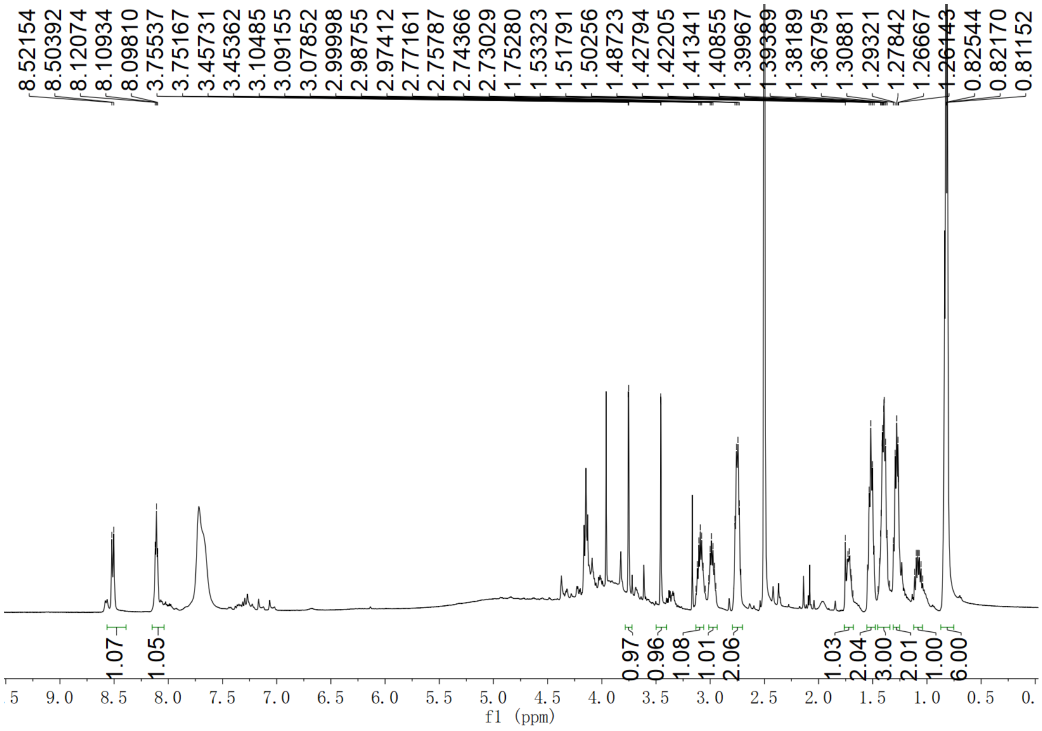

**Supplementary Fig. 54. $^{1}$H NMR spectrum of CPI-3 (5) in DMSO-$d_6$**

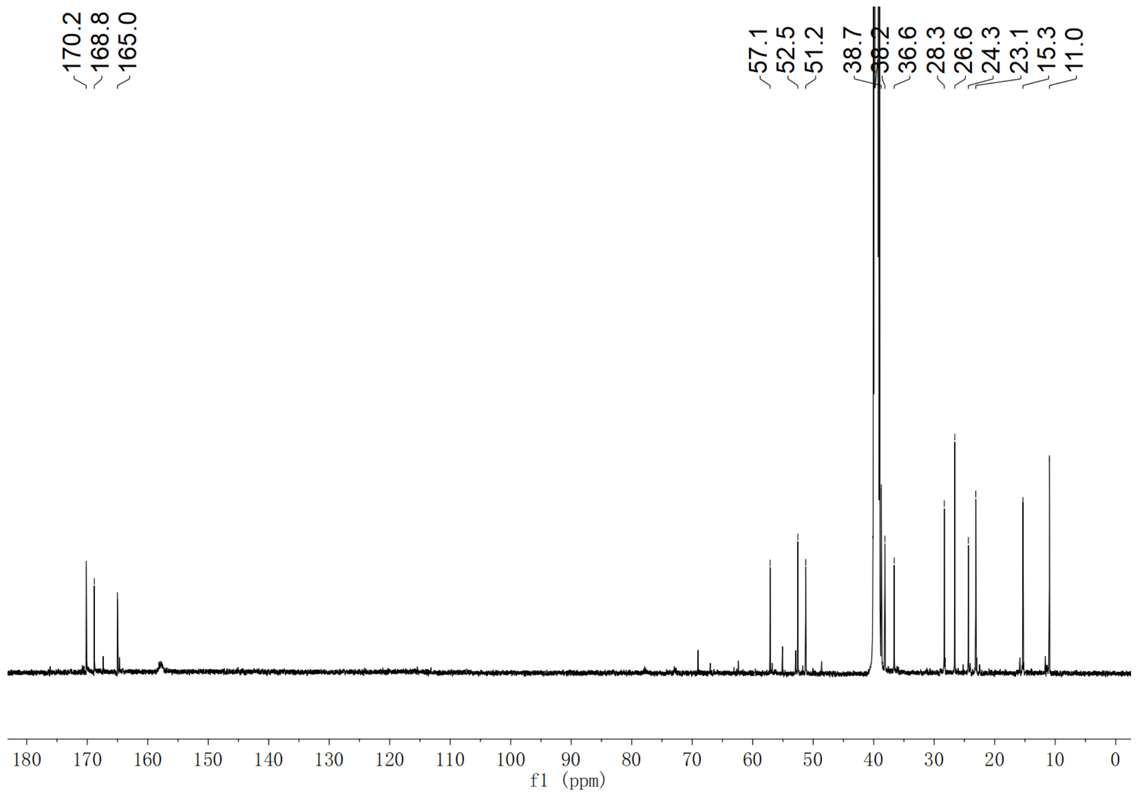

**Supplementary Fig. 55. $^{13}$C NMR spectrum of CPI-3 (5) in DMSO-*d*$_6$**

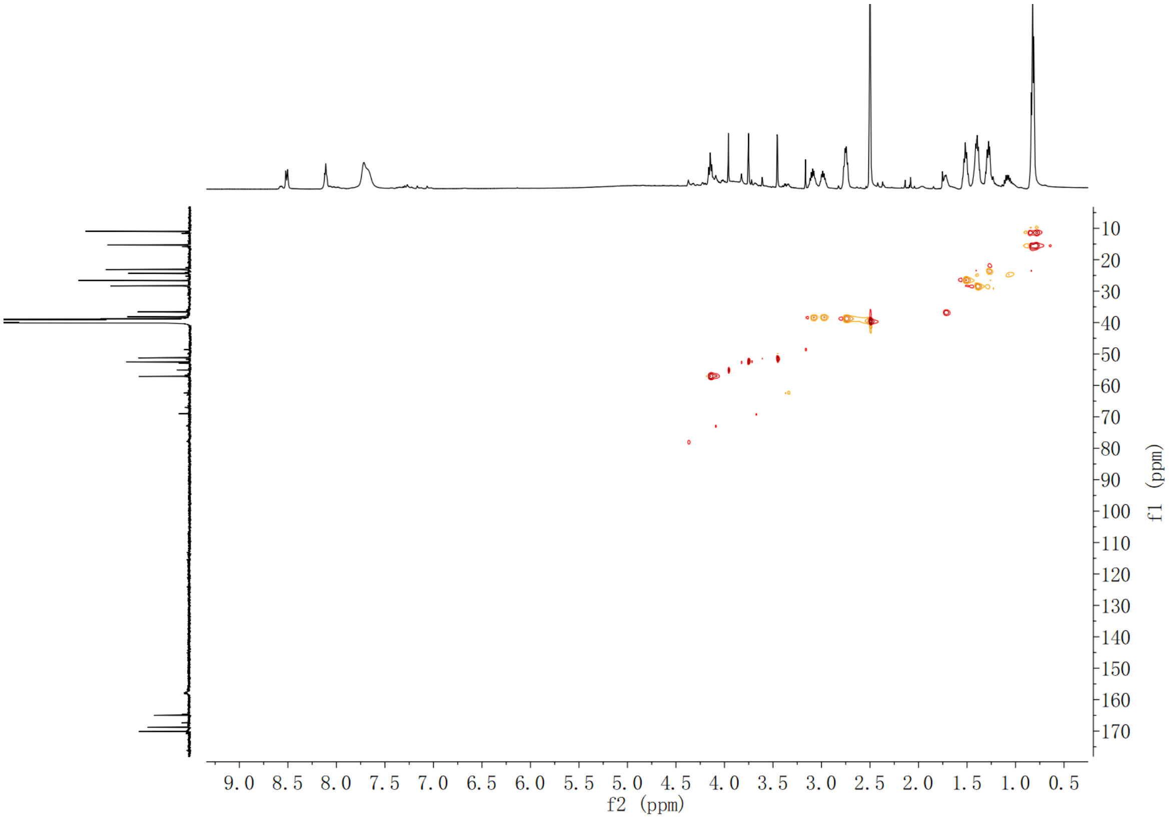

**Supplementary Fig. 56. HSQC spectrum of CPI-3 (5) in DMSO-*d*$_6$**

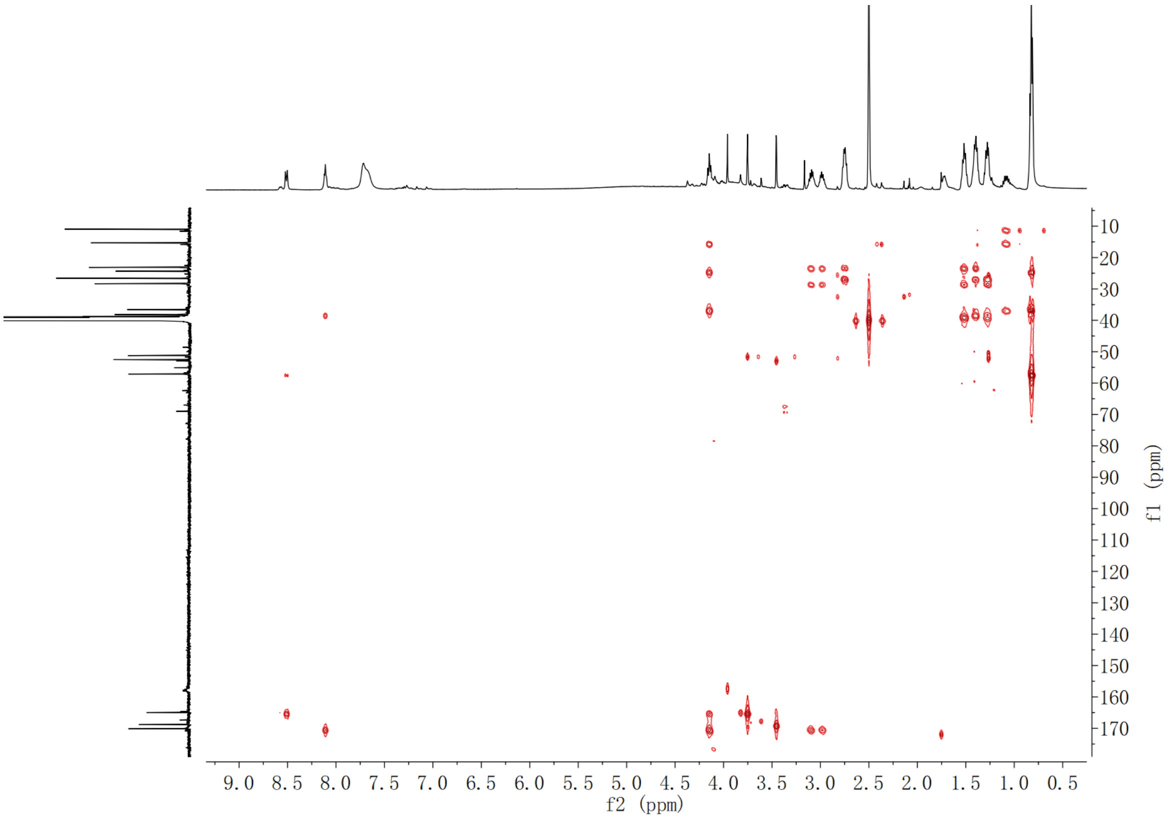

**Supplementary Fig. 57. HMBC spectrum of CPI-3 (5) in DMSO-*$d_6$***

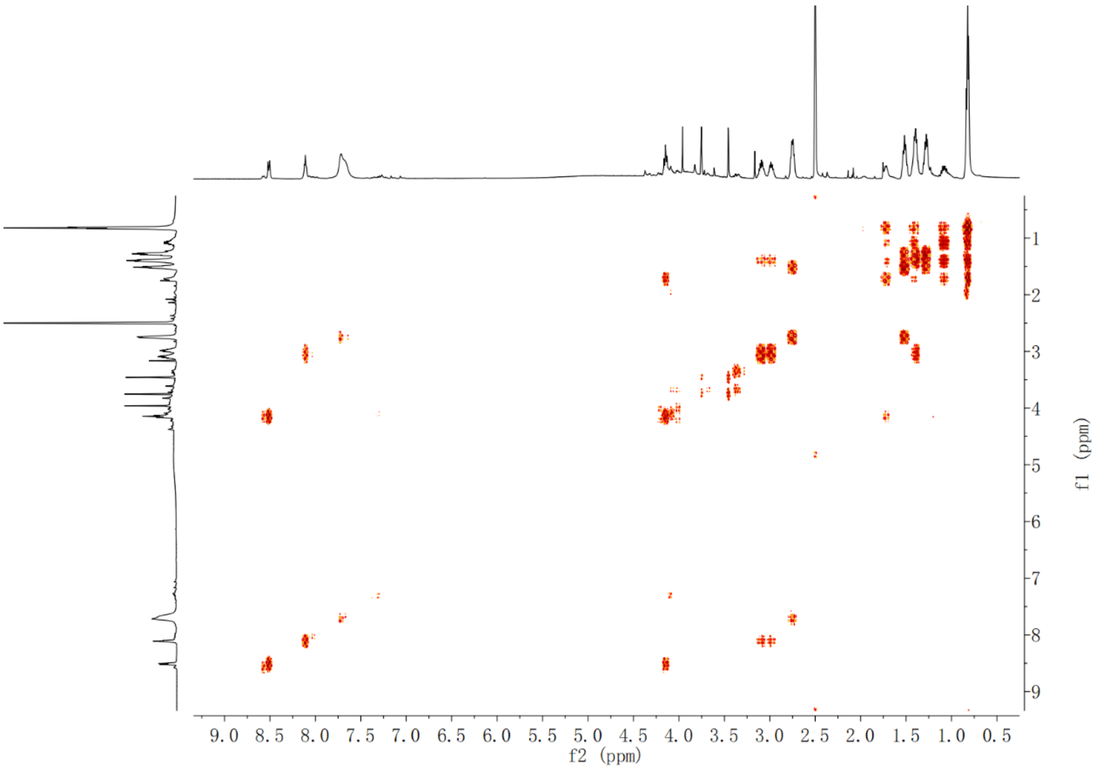

**Supplementary Fig. 58. $^1$H-$^1$H COSY spectrum of CPI-3 (5) in DMSO-*$d_6$***

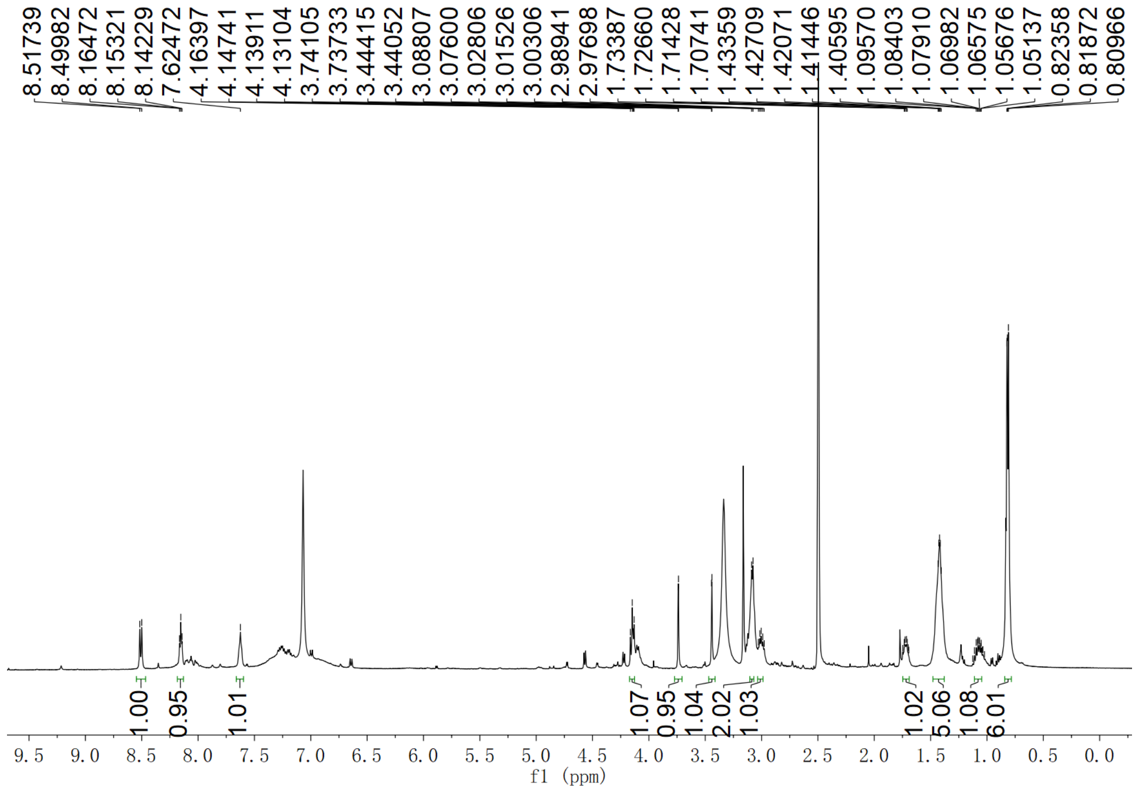

**Supplementary Fig. 59. $^{1}$H NMR spectrum of compound 6 in DMSO-$d_6$**

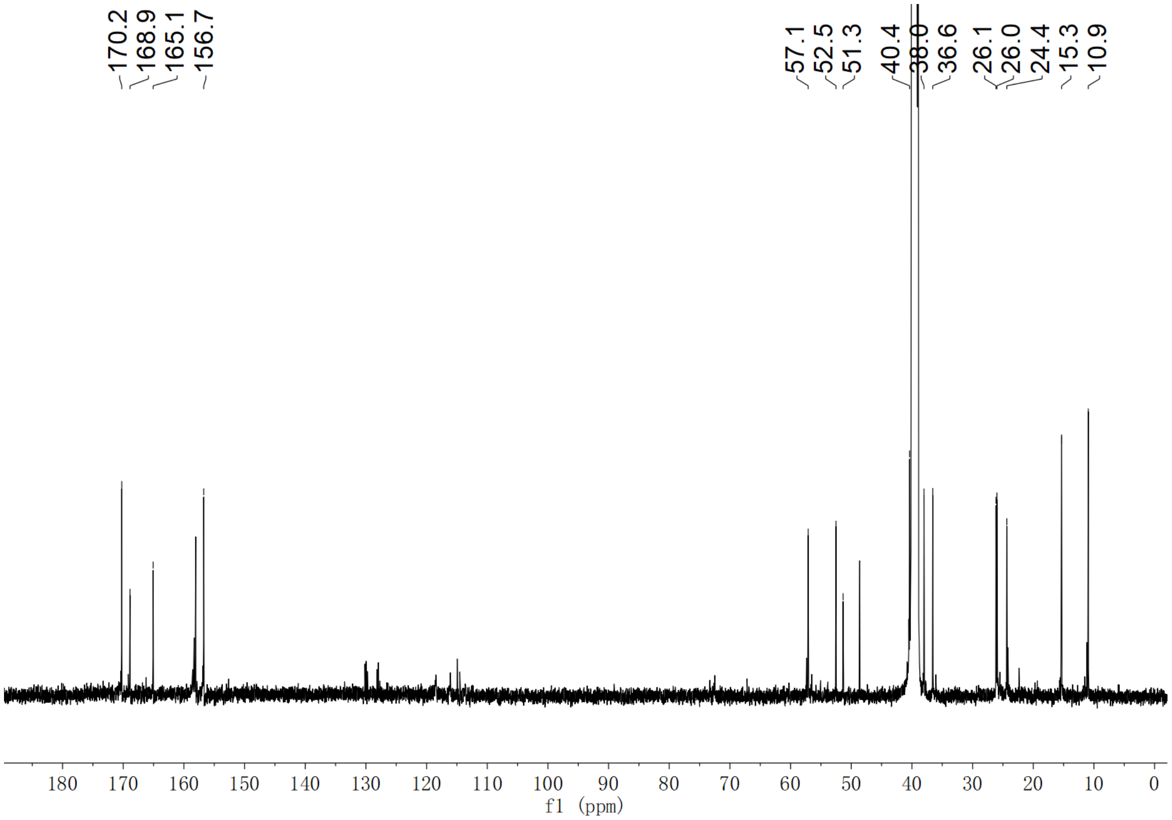

**Supplementary Fig. 60. $^{13}$C NMR spectrum of compound 6 in DMSO-$d_6$**

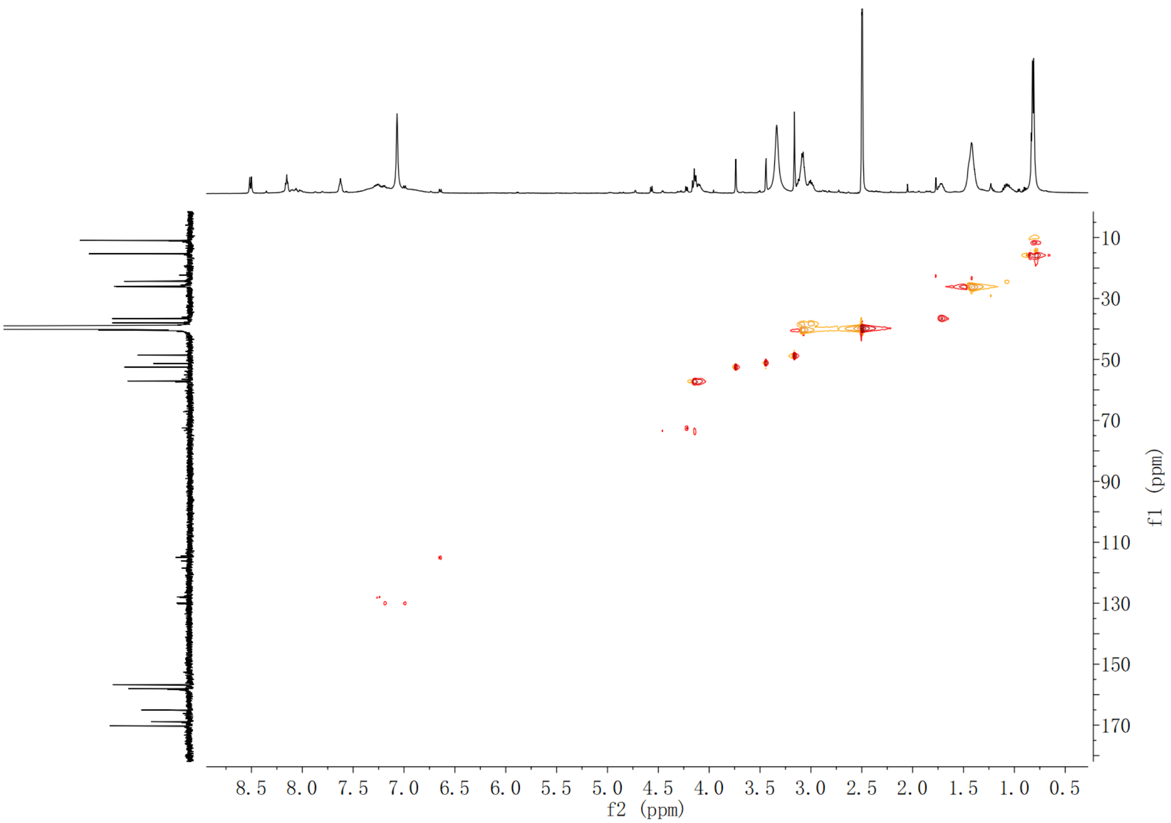

**Supplementary Fig. 61. HSQC spectrum of compound 6 in DMSO-*d*$_6$**

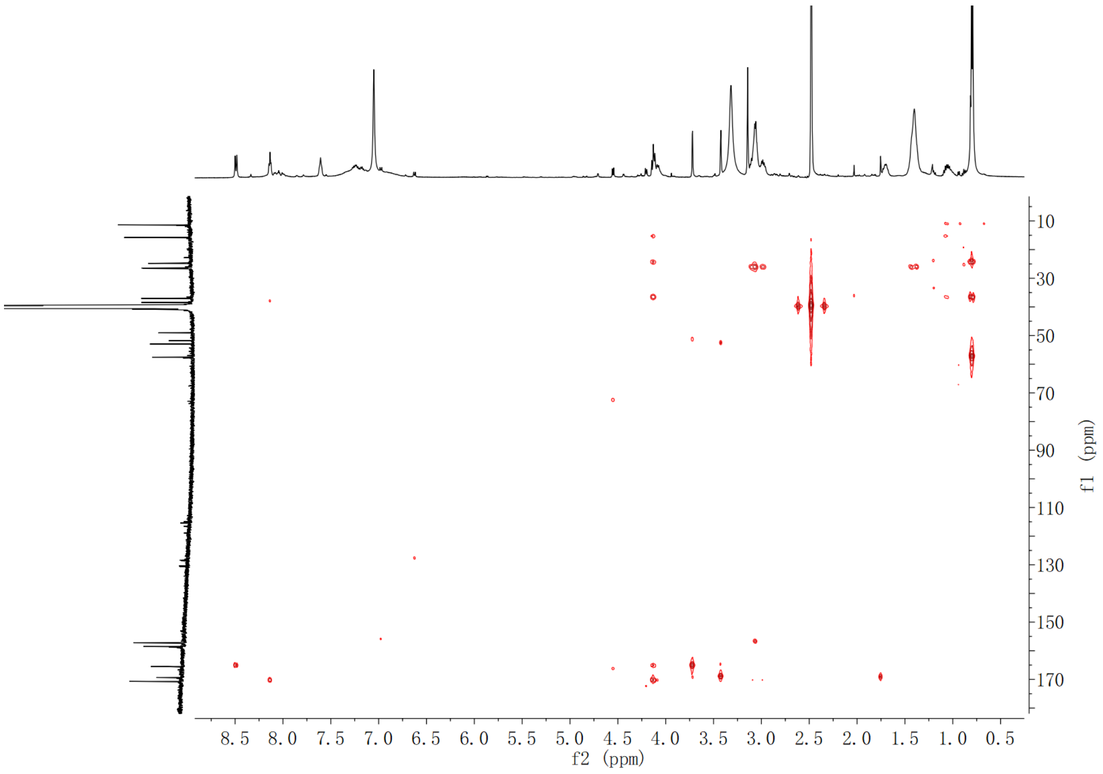

**Supplementary Fig. 62. HMBC spectrum of compound 6 in DMSO-*d*$_6$**

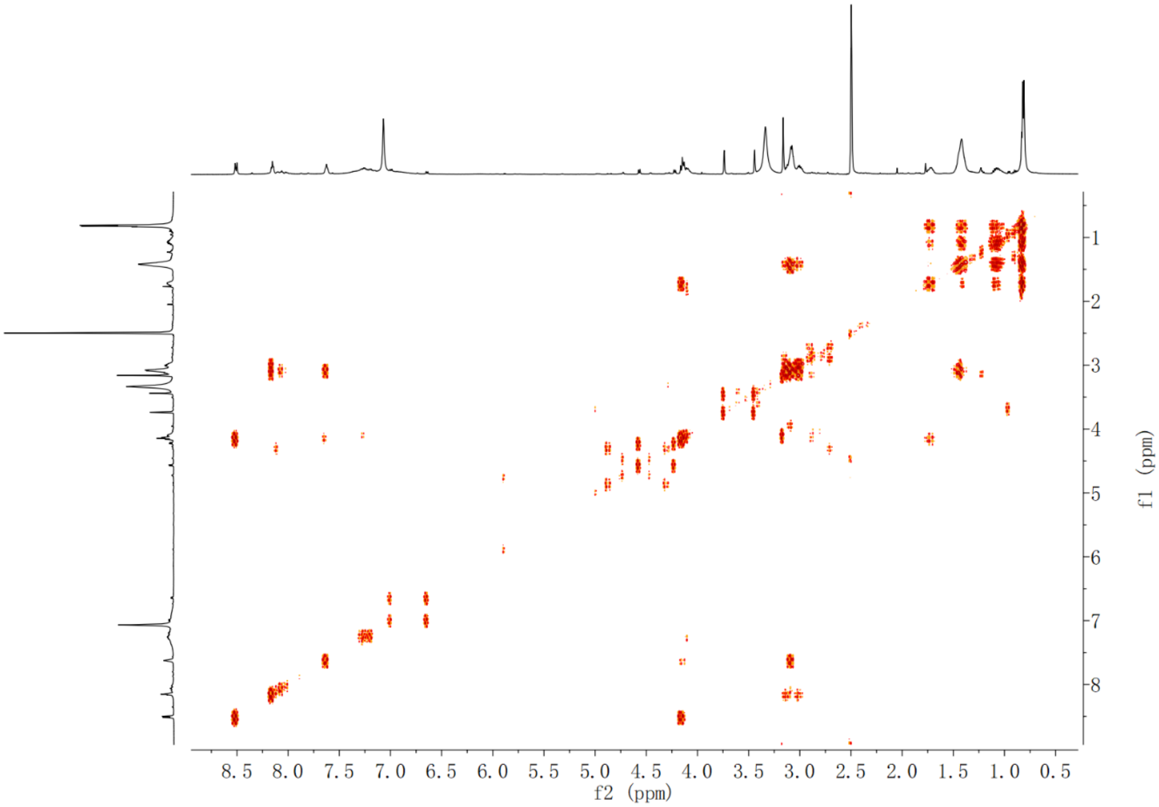

**Supplementary Fig. 63. $^{1}$H-$^{1}$H COSY spectrum of compound 6 in DMSO-*d*$_6$**

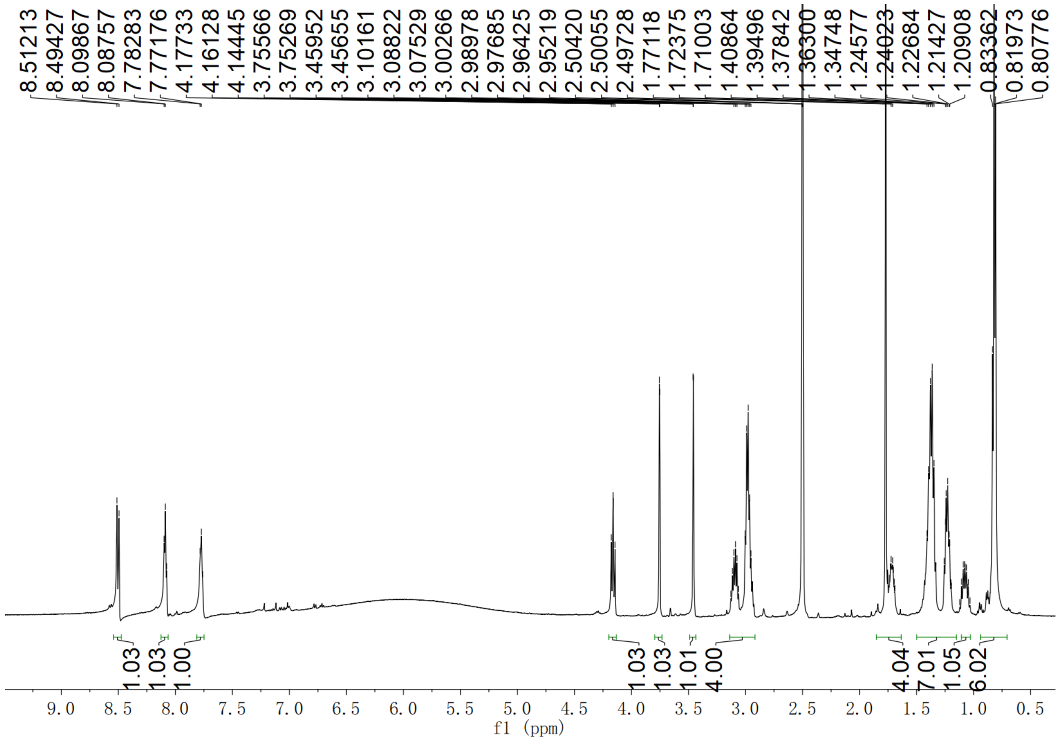

**Supplementary Fig. 64 $^{1}$H NMR spectrum of compound 7 in DMSO-*d*$_6$**

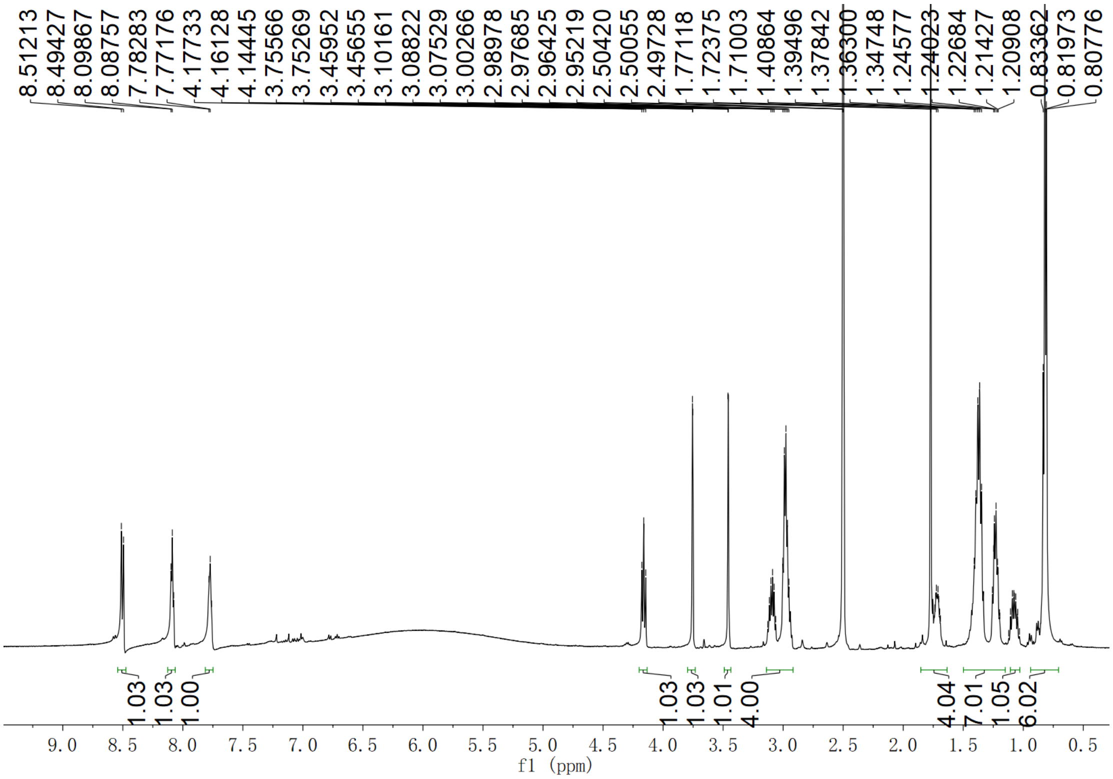

**Supplementary Fig. 65. $^{13}$C NMR spectrum of compound 7 in DMSO-$d_6$**

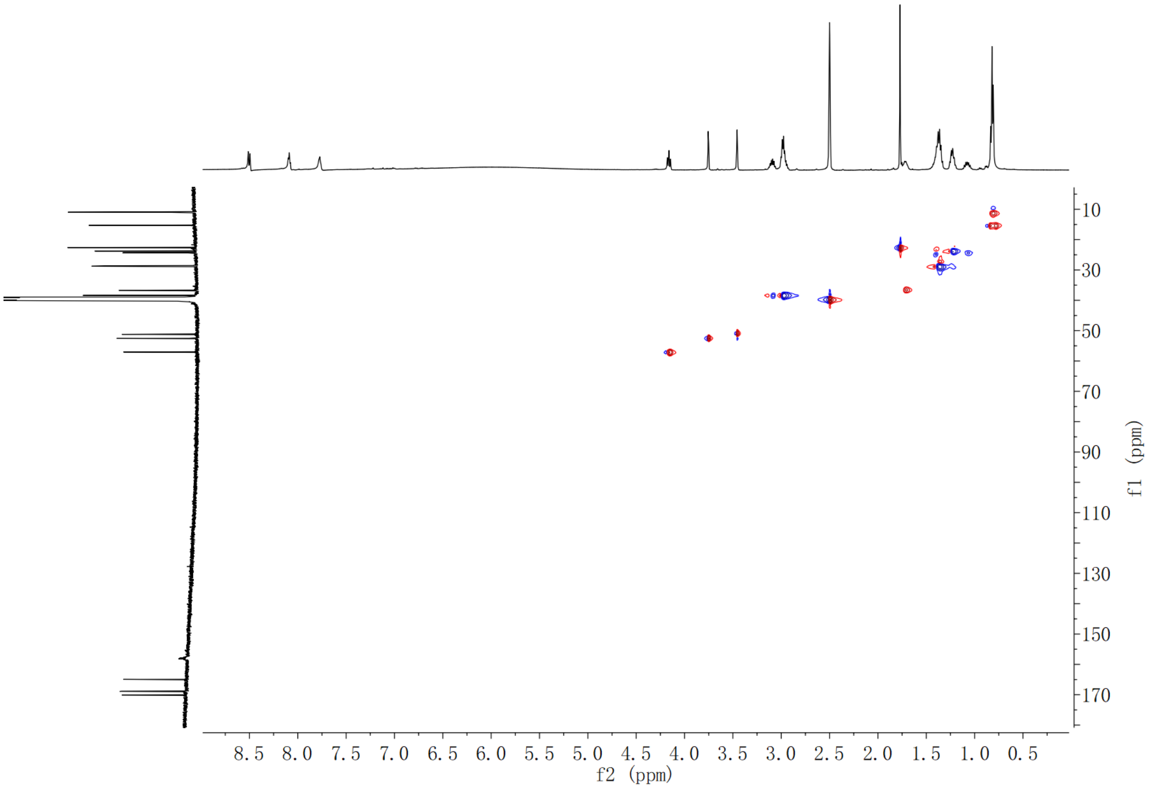

**Supplementary Fig. 66. HSQC spectrum of compound 7 in DMSO-$d_6$**

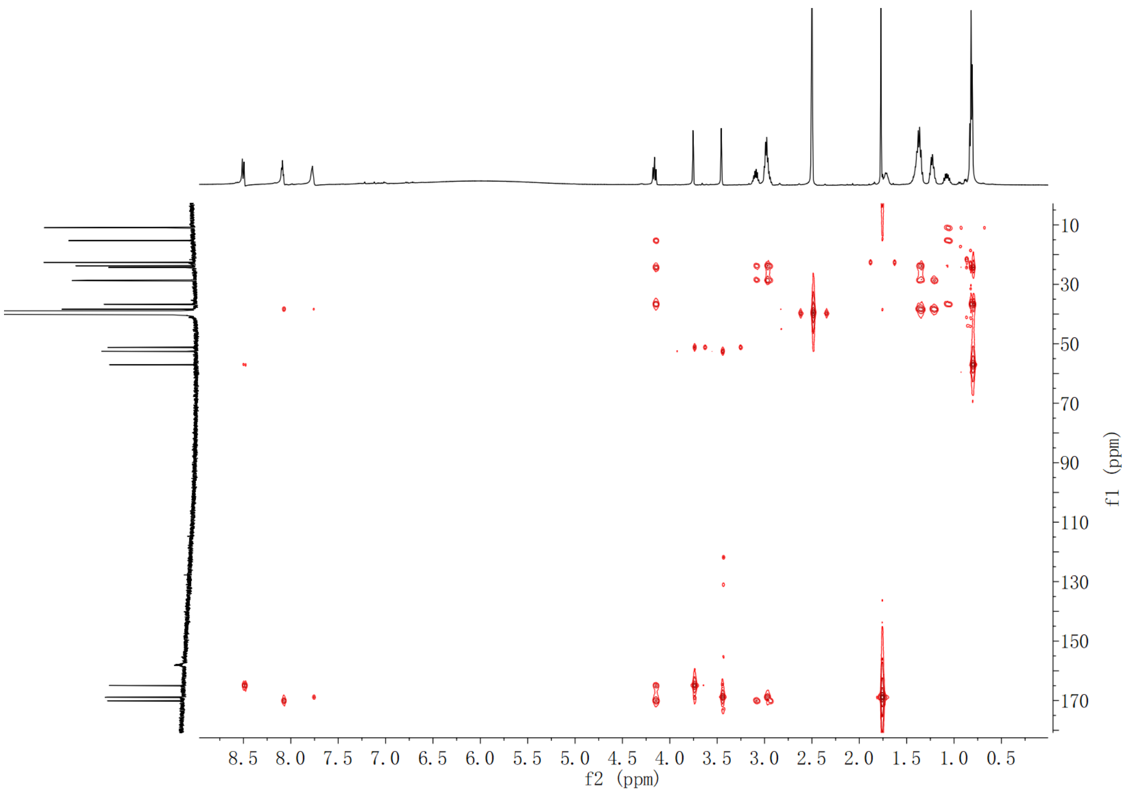

**Supplementary Fig. 67. HMBC spectrum of compound 7 in DMSO-$d_6$**

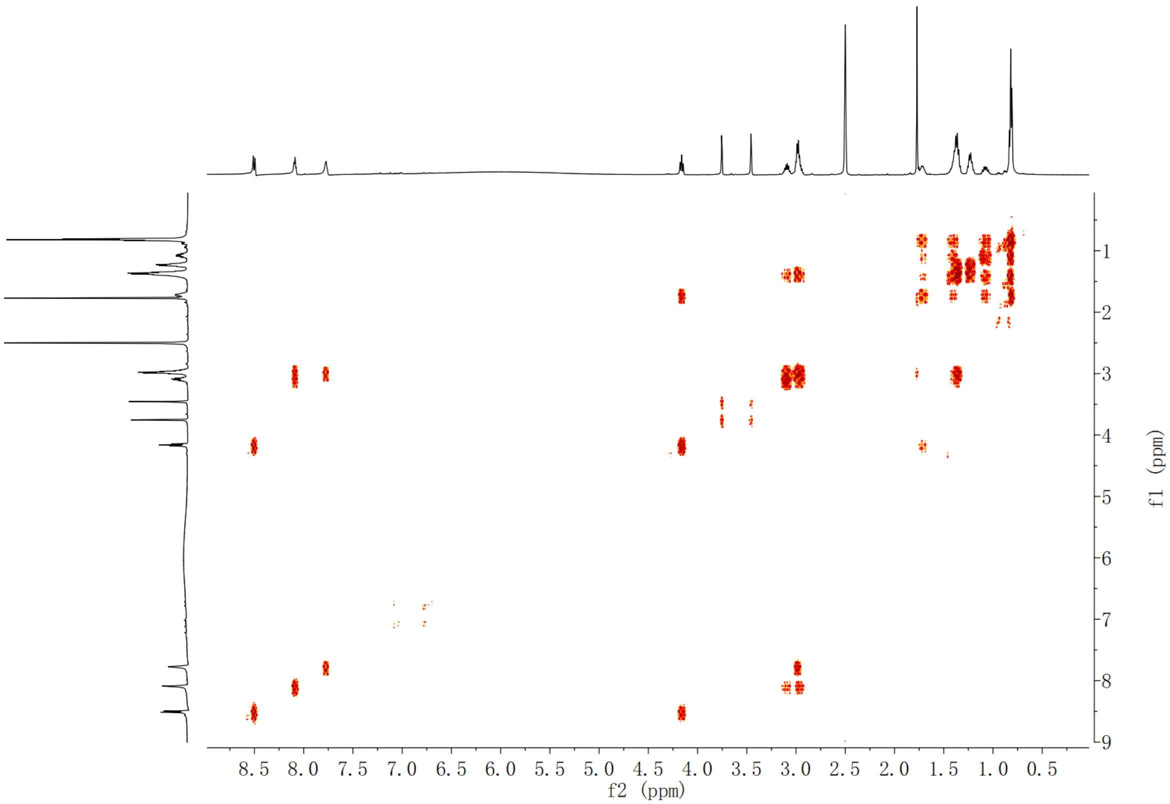

**Supplementary Fig. 68. $^{1}$H-$^{1}$H COSY spectrum of compound 7 in DMSO-$d_6$**

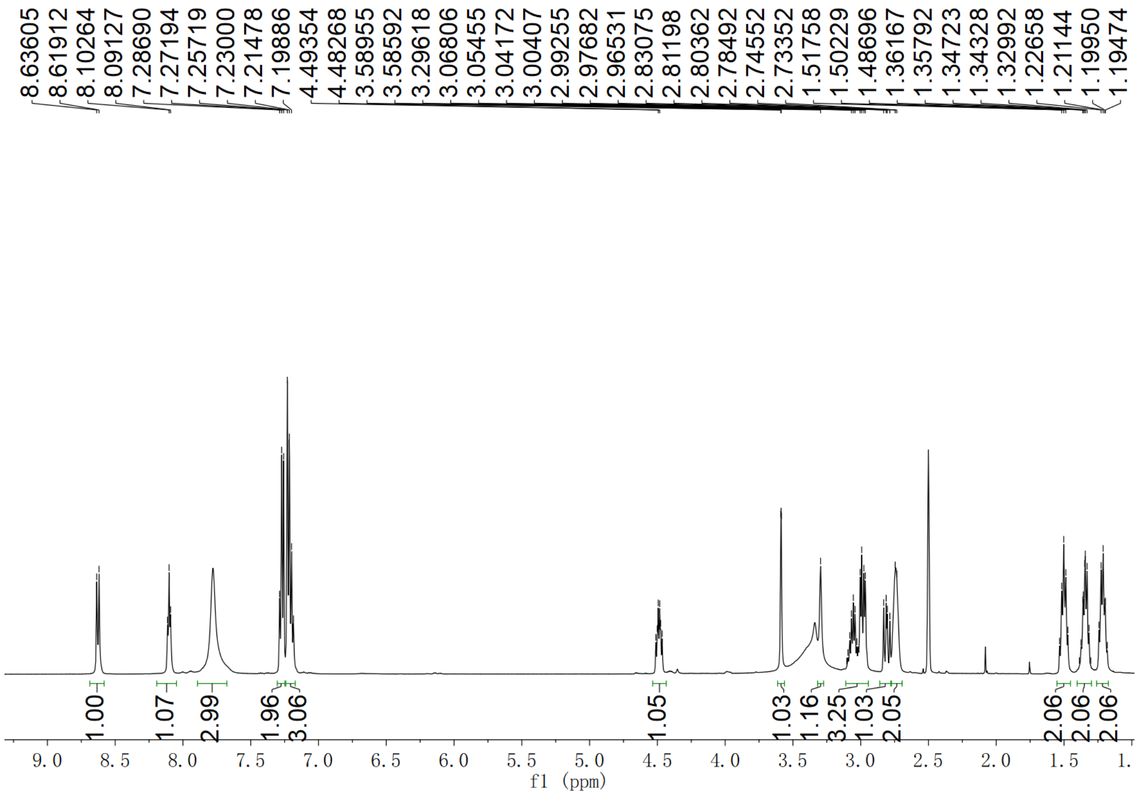

**Supplementary Fig. 69. $^{1}$H NMR spectrum of compound 8 in DMSO-$d_6$**

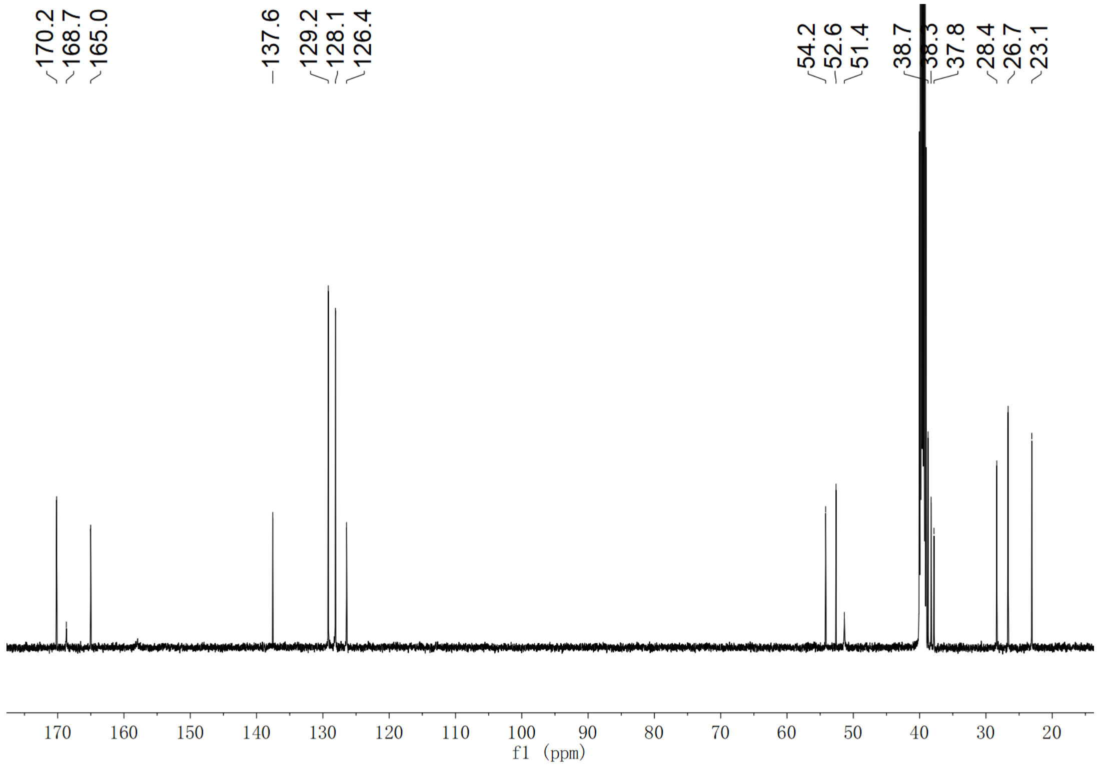

**Supplementary Fig. 70. $^{13}$C NMR spectrum of compound 8 in DMSO-$d_6$**

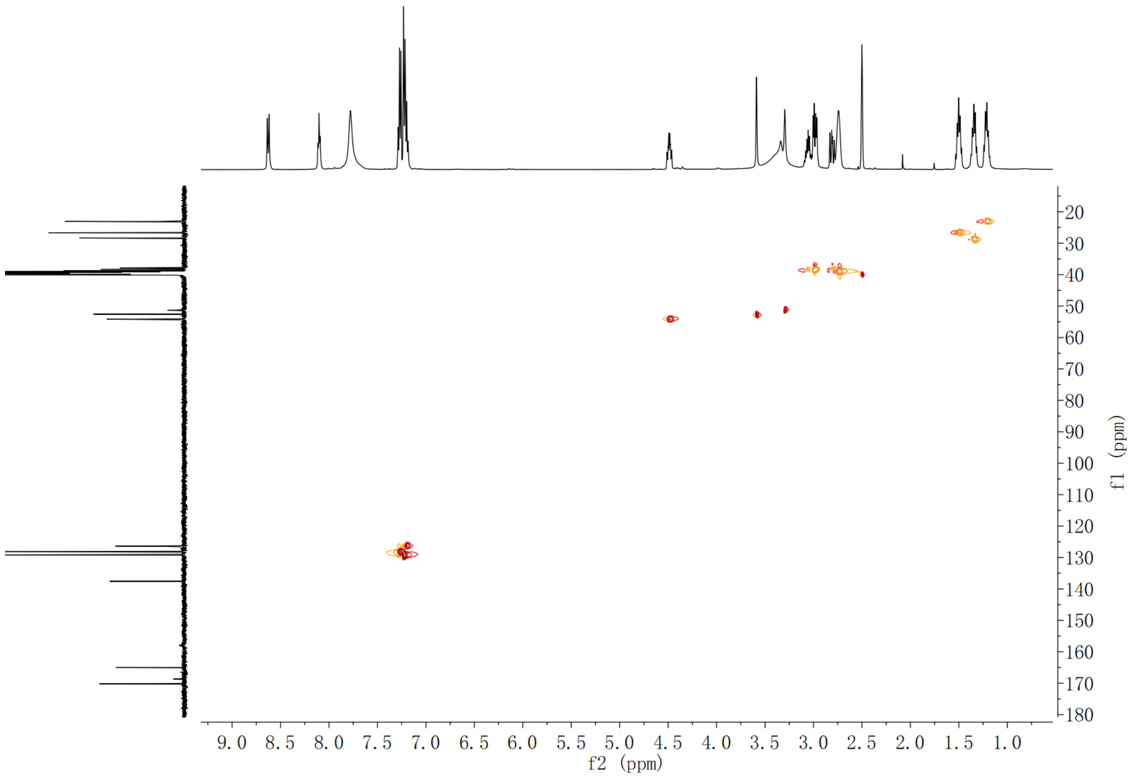

**Supplementary Fig. 71. HSQC spectrum of compound 8 in DMSO-$d_6$**

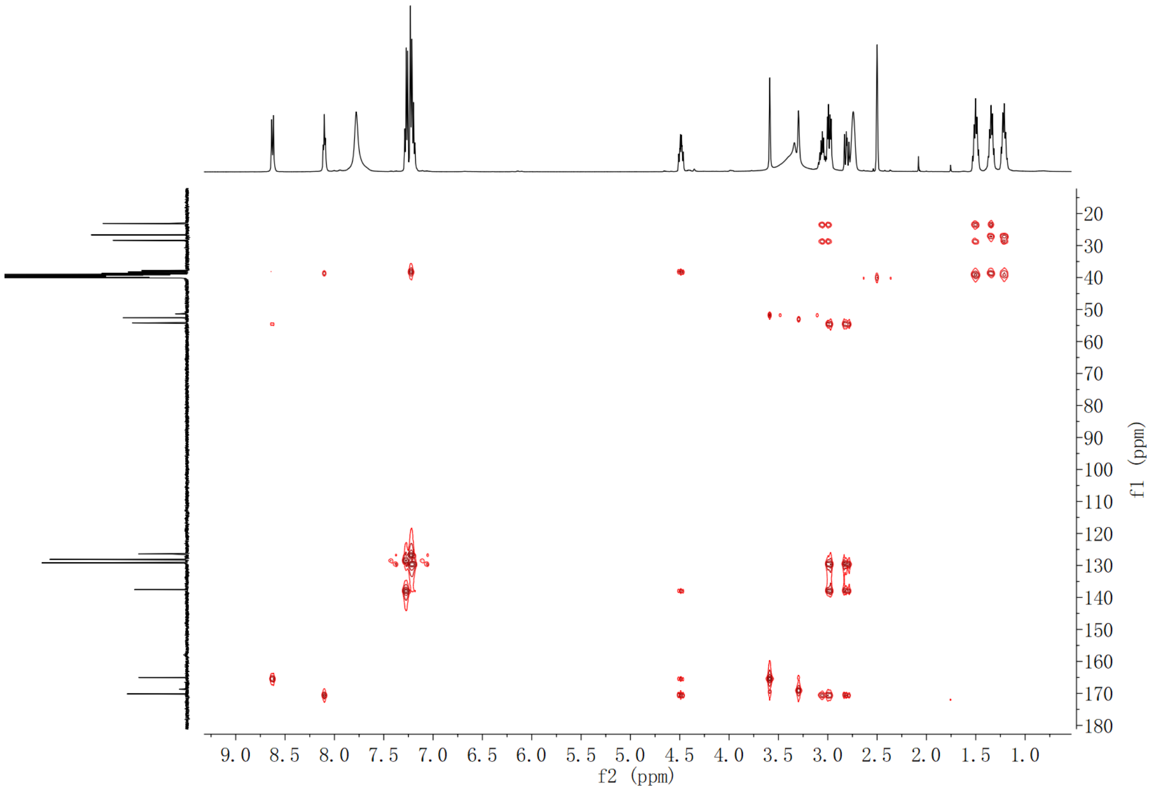

**Supplementary Fig. 72. HMBC spectrum of compound 8 in DMSO-$d_6$**

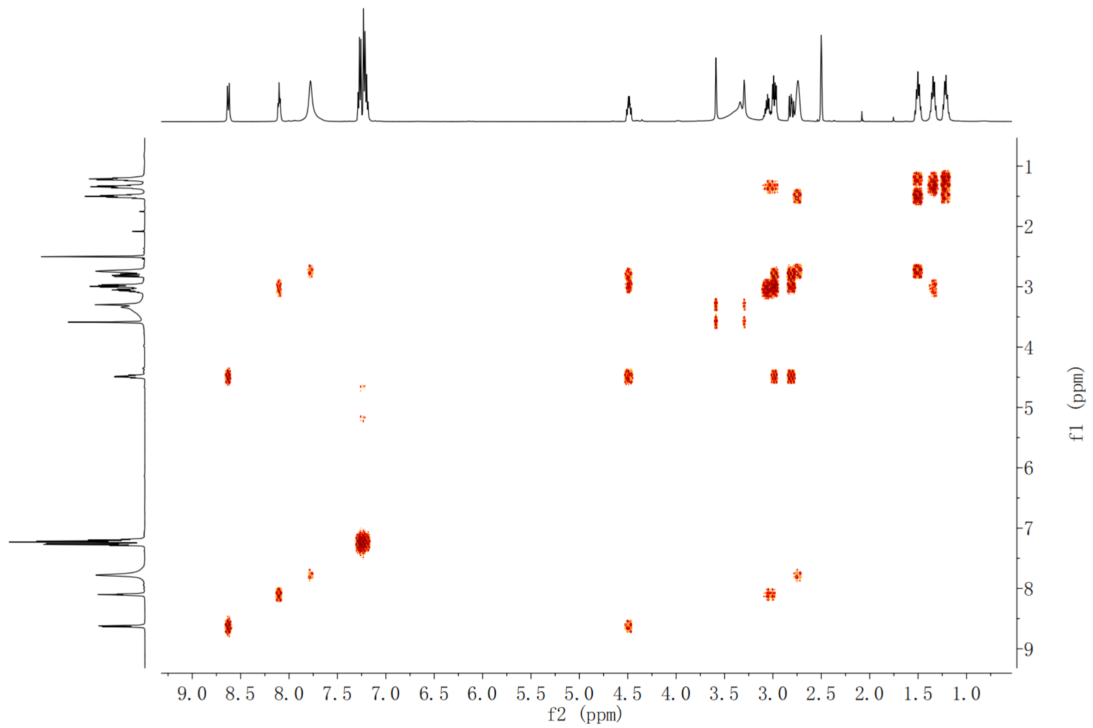

**Supplementary Fig. 73. $^{1}H$-$^{1}H$ COSY spectrum of compound 8 in DMSO-$d_6$**

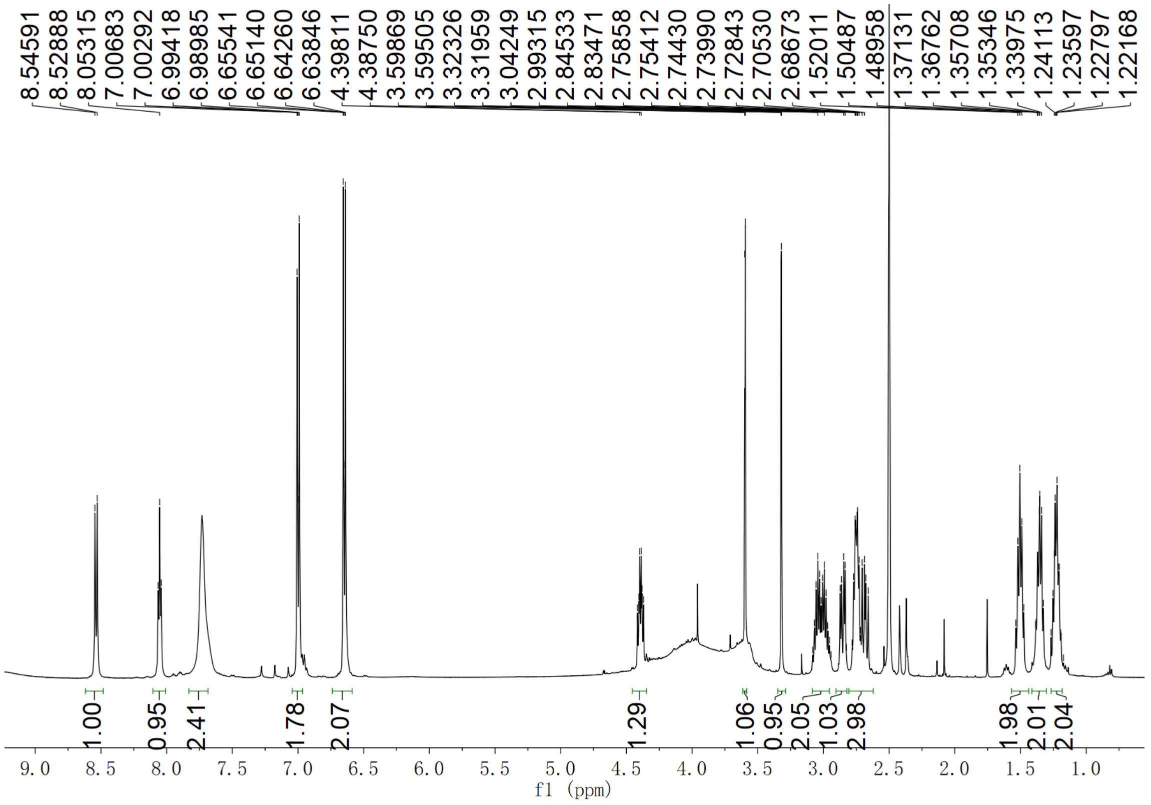

**Supplementary Fig. 74. $^{1}H$ NMR spectrum of compound 9 in DMSO-$d_6$**

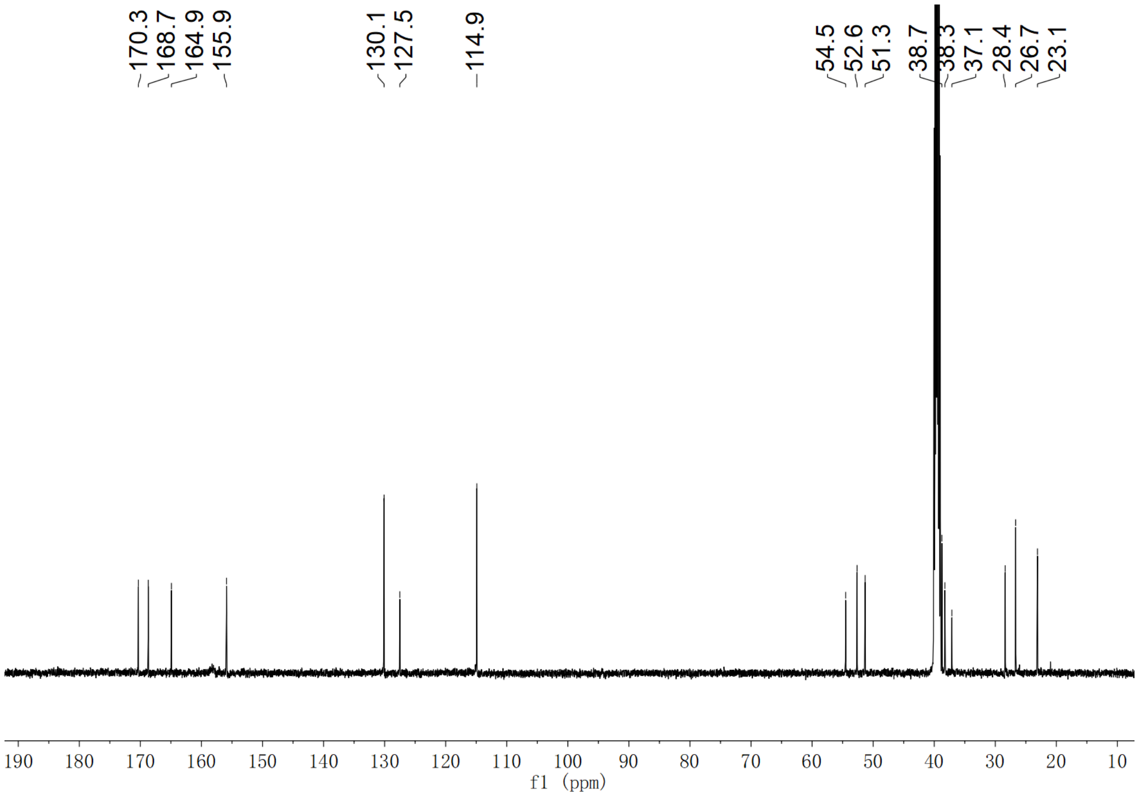

**Supplementary Fig. 75. $^{13}$C NMR spectrum of compound 9 in DMSO-$d_6$**

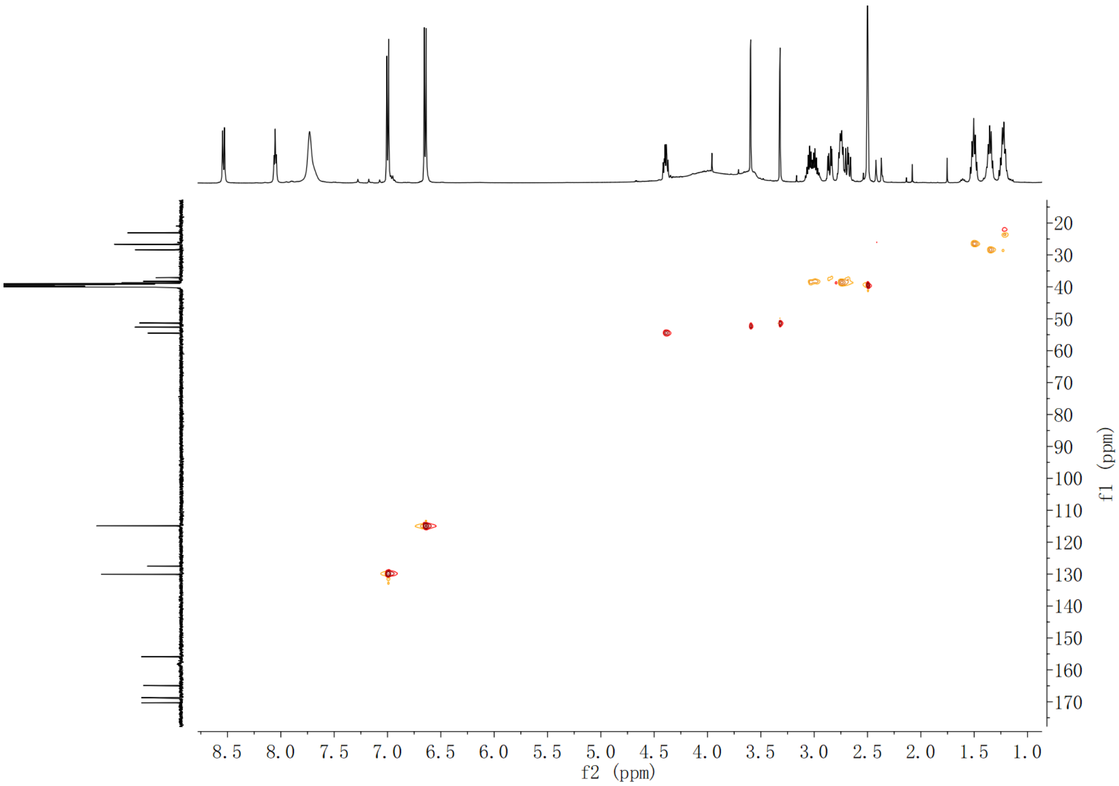

**Supplementary Fig. 76. HSQC spectrum of compound 9 in DMSO-$d_6$**

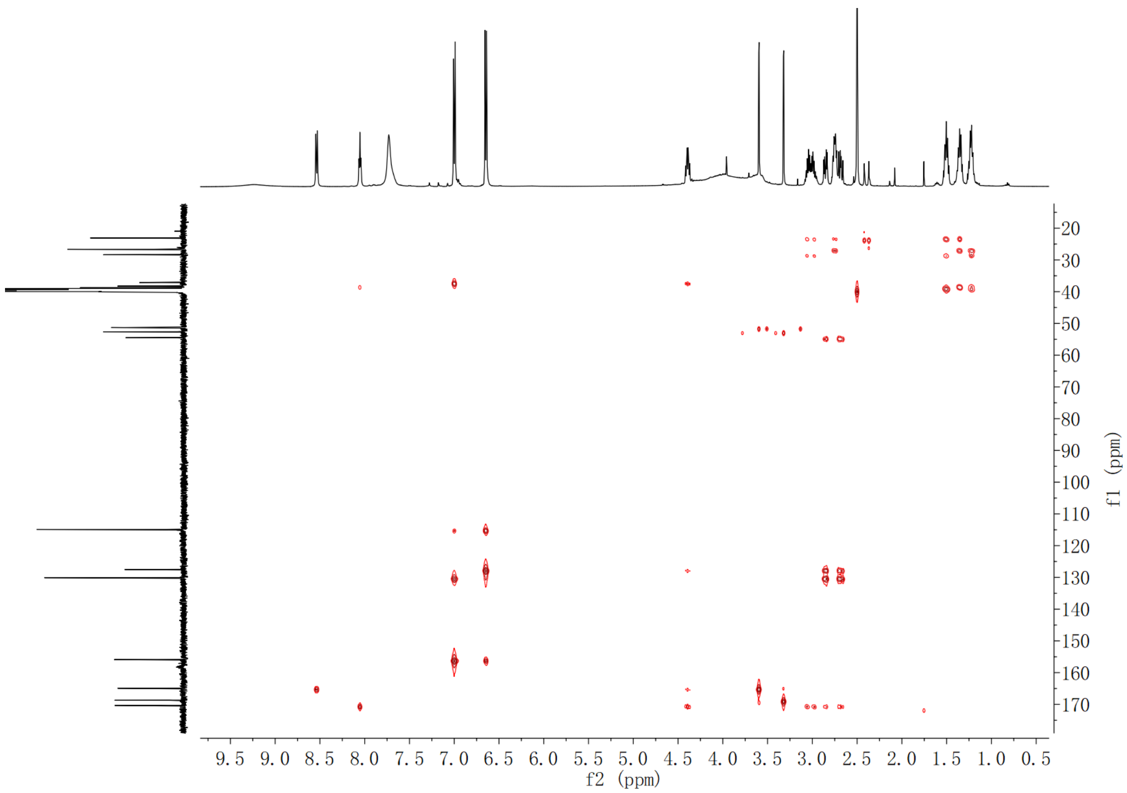

**Supplementary Fig. 77. HMBC spectrum of compound 9 in DMSO-$d_6$**

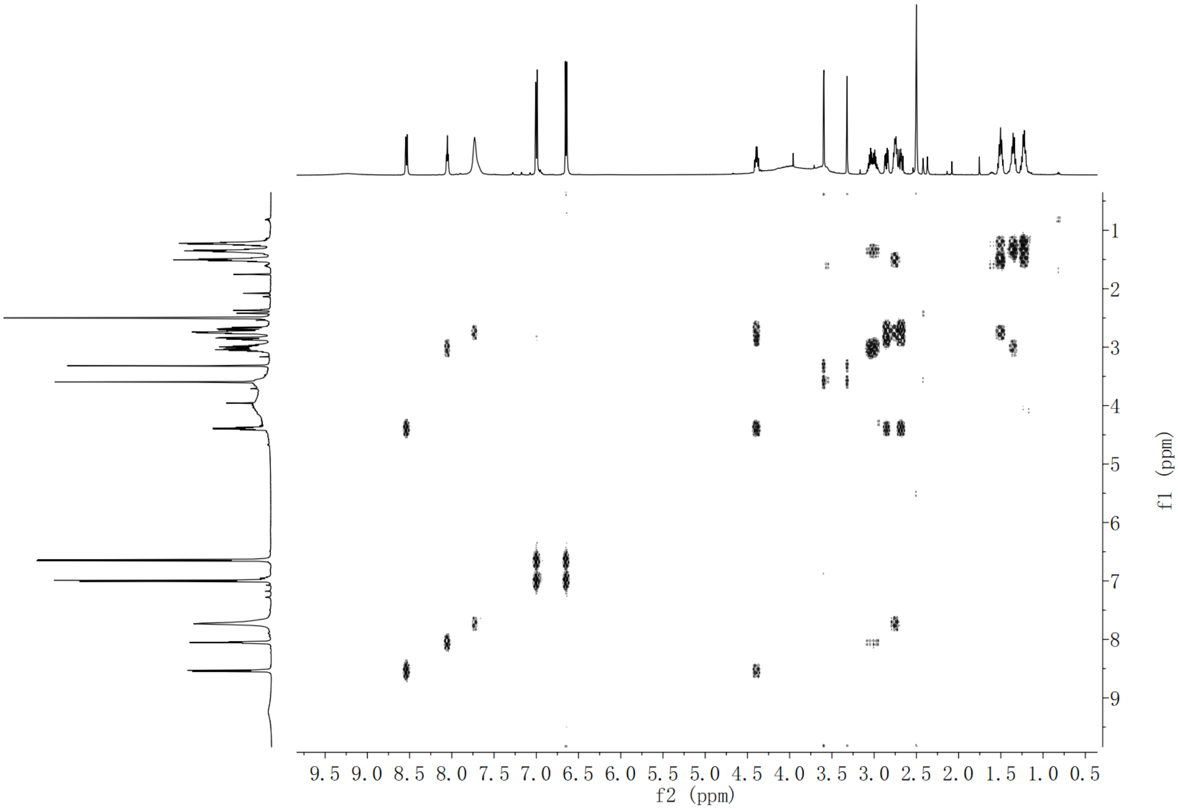

**Supplementary Fig. 78. $^1$H-$^1$H COSY spectrum of compound 9 in DMSO-$d_6$**

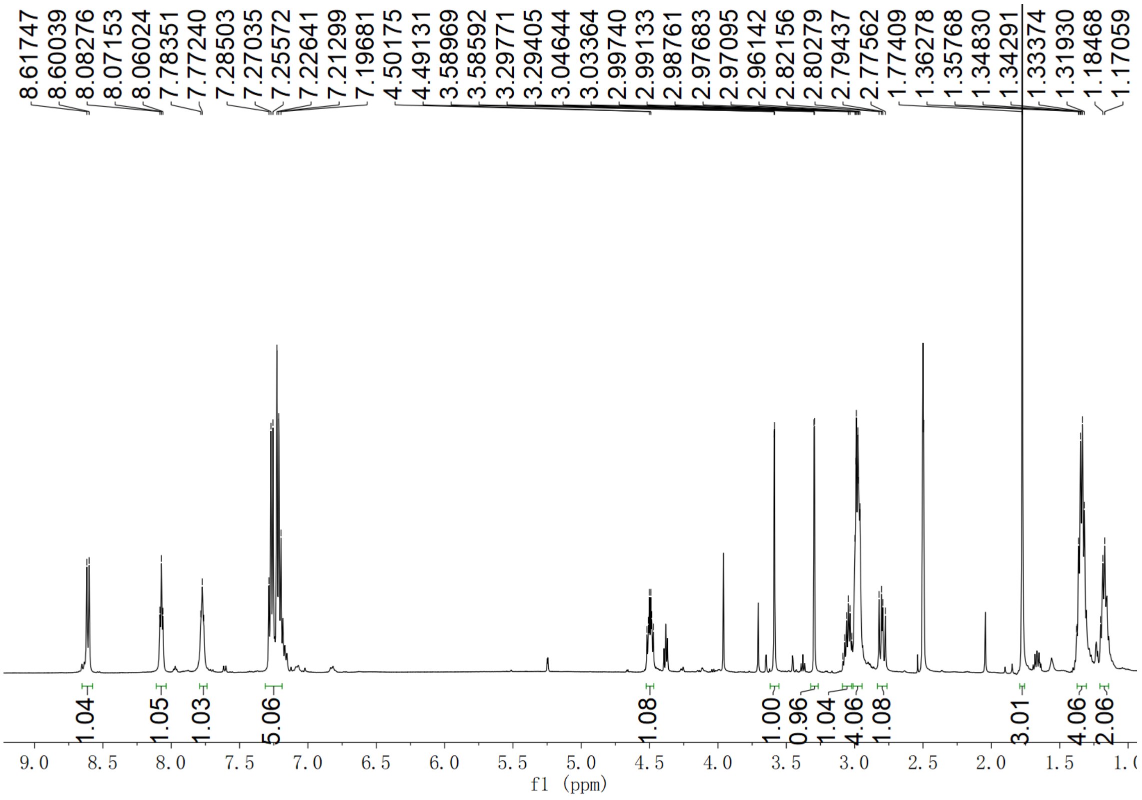

**Supplementary Fig. 79. $^{1}$H NMR spectrum of compound 10 in DMSO-$d_6$**

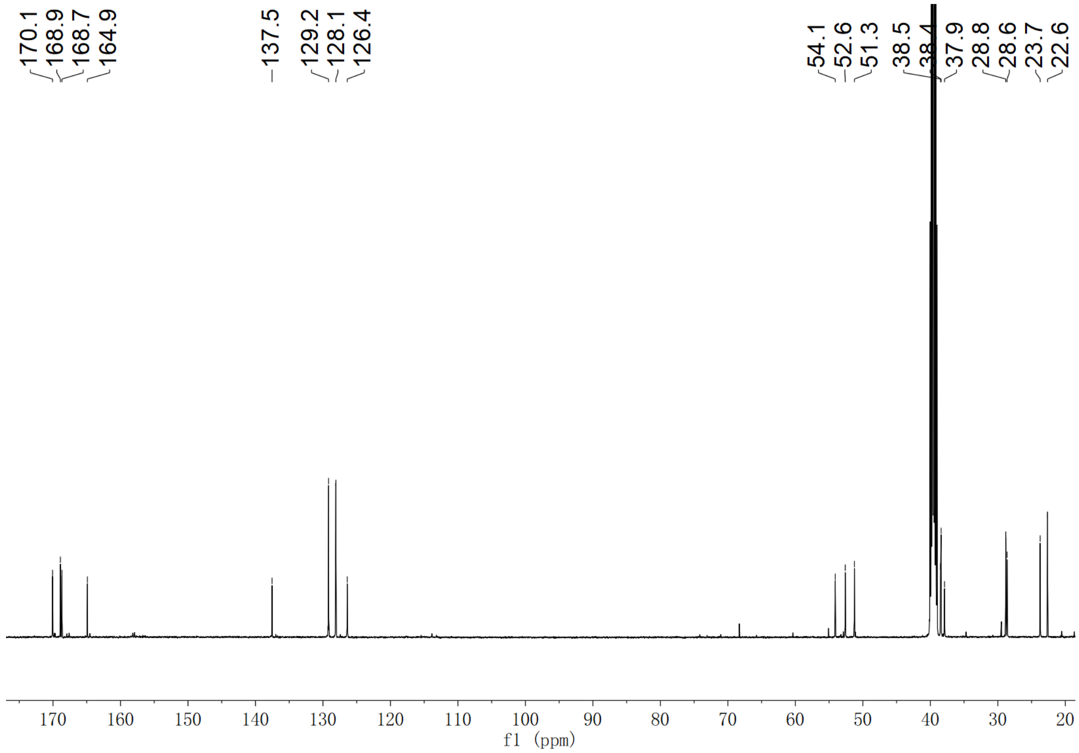

**Supplementary Fig. 80. $^{13}$C NMR spectrum of compound 10 in DMSO-$d_6$**

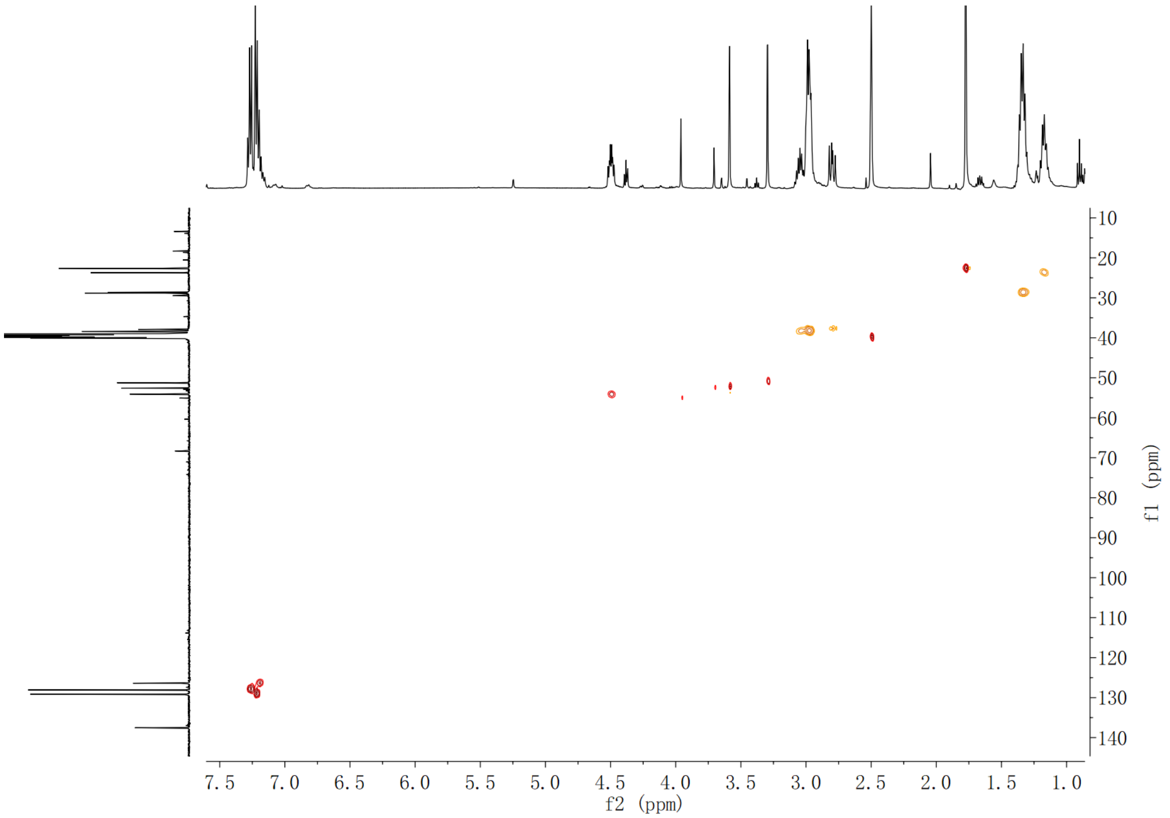

**Supplementary Fig. 81. HSQC spectrum of compound 10 in DMSO-*d*$_6$**

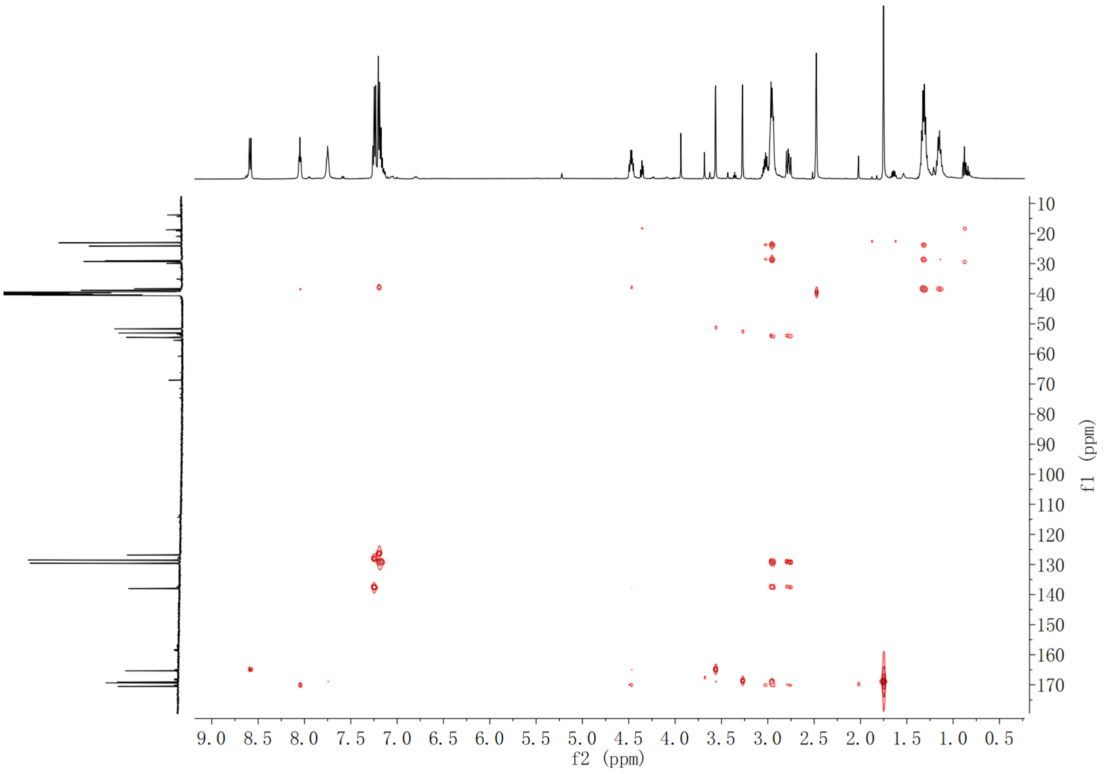

**Supplementary Fig. 82. HMBC spectrum of compound 10 in DMSO-*d*$_6$**

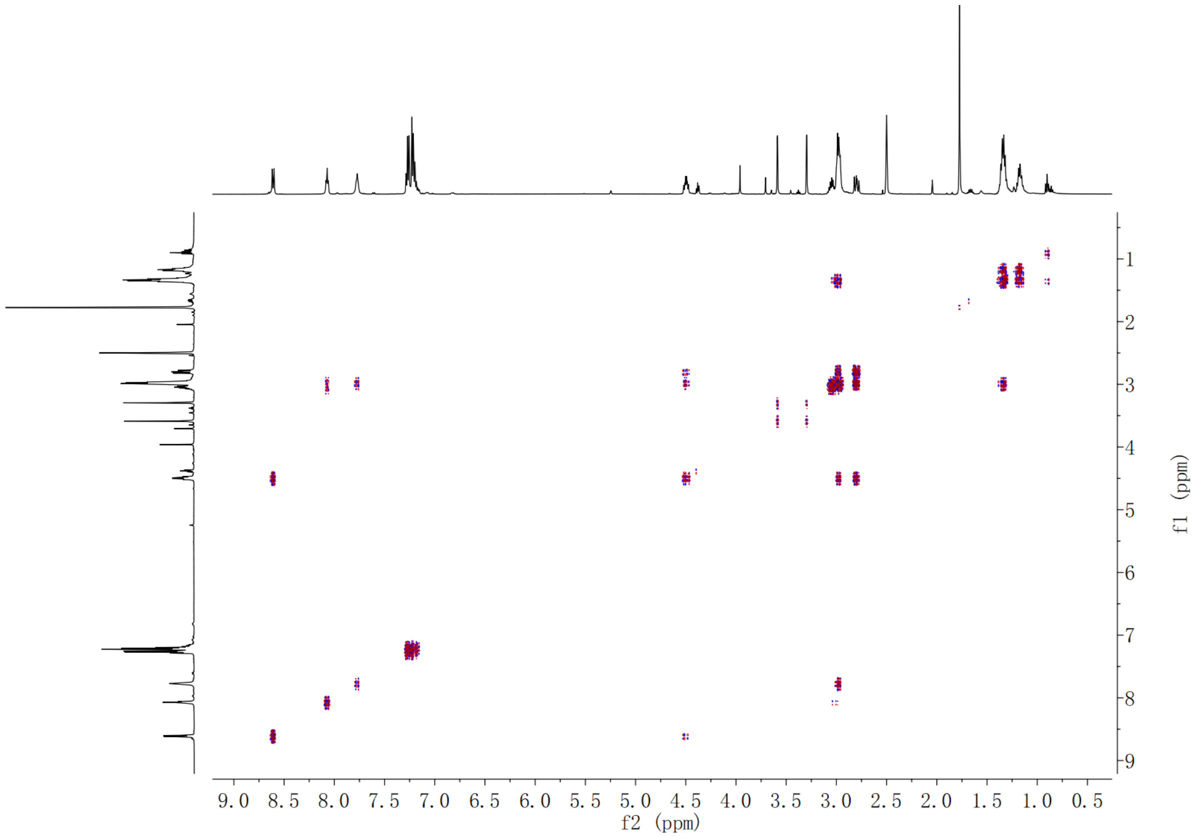

**Supplementary Fig. 83. $^1$H-$^1$H COSY spectrum of compound 10 in DMSO-$d_6$**

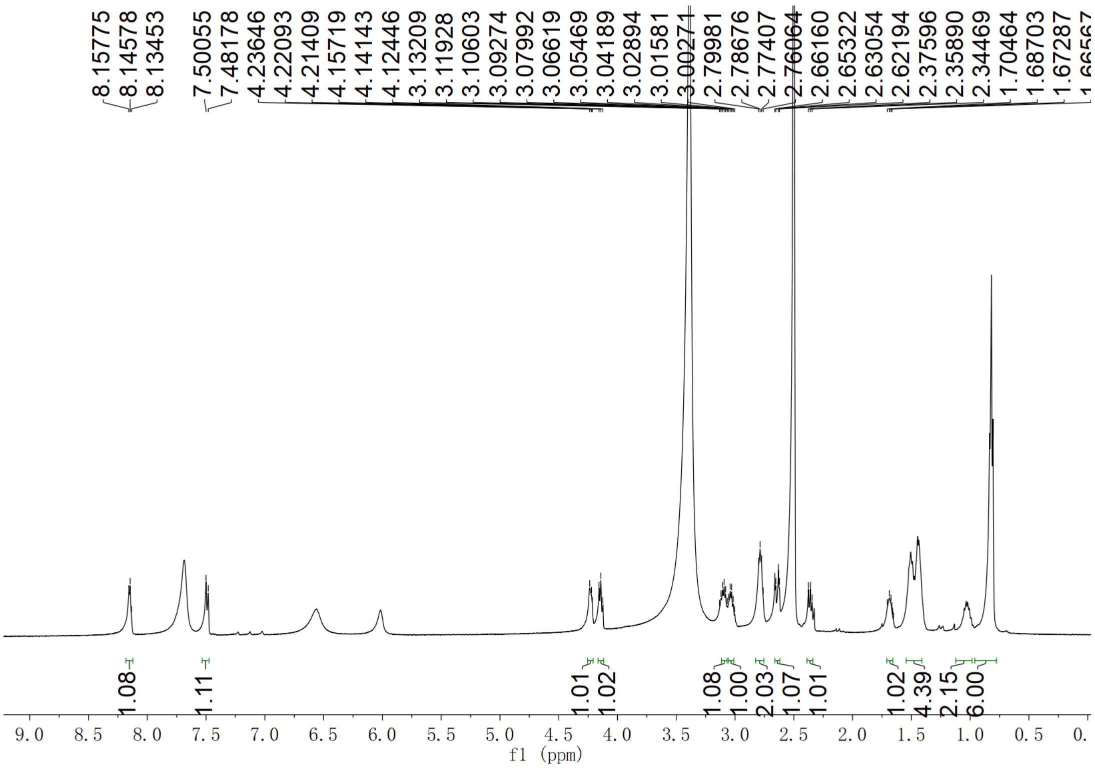

**Supplementary Fig. 84. $^1$H NMR spectrum of compound 12 in DMSO-$d_6$**

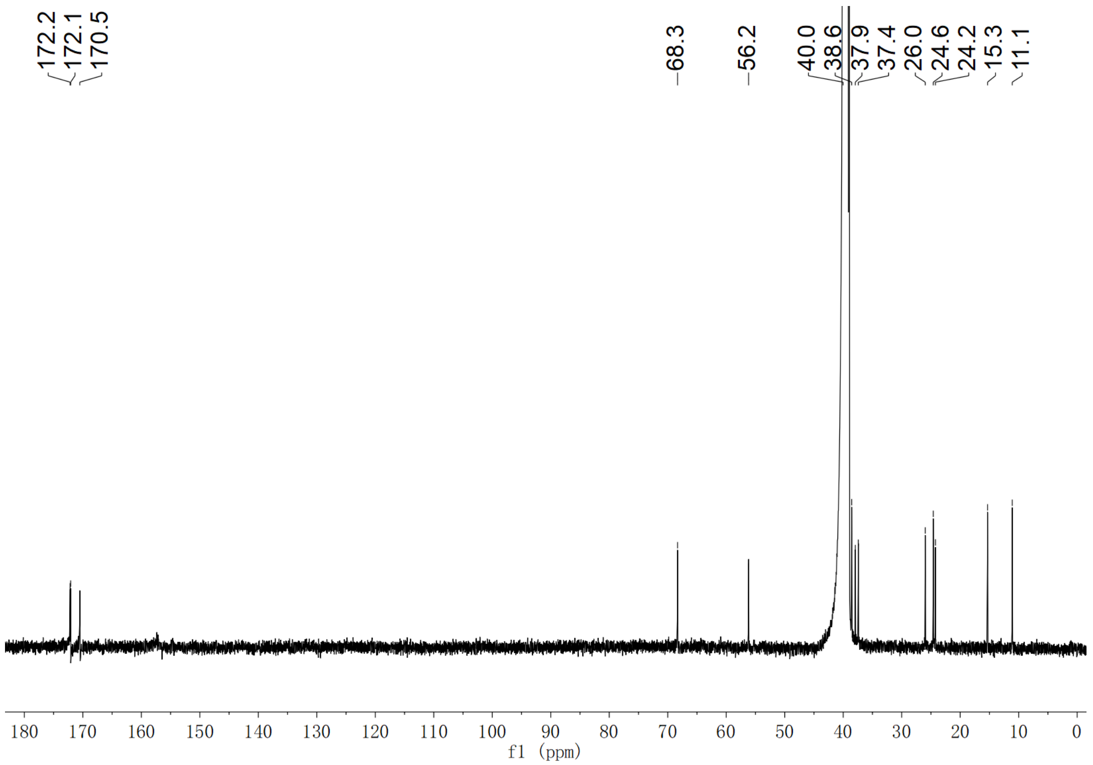

**Supplementary Fig. 85. $^{13}$C NMR spectrum of compound 12 in DMSO-$d_6$**

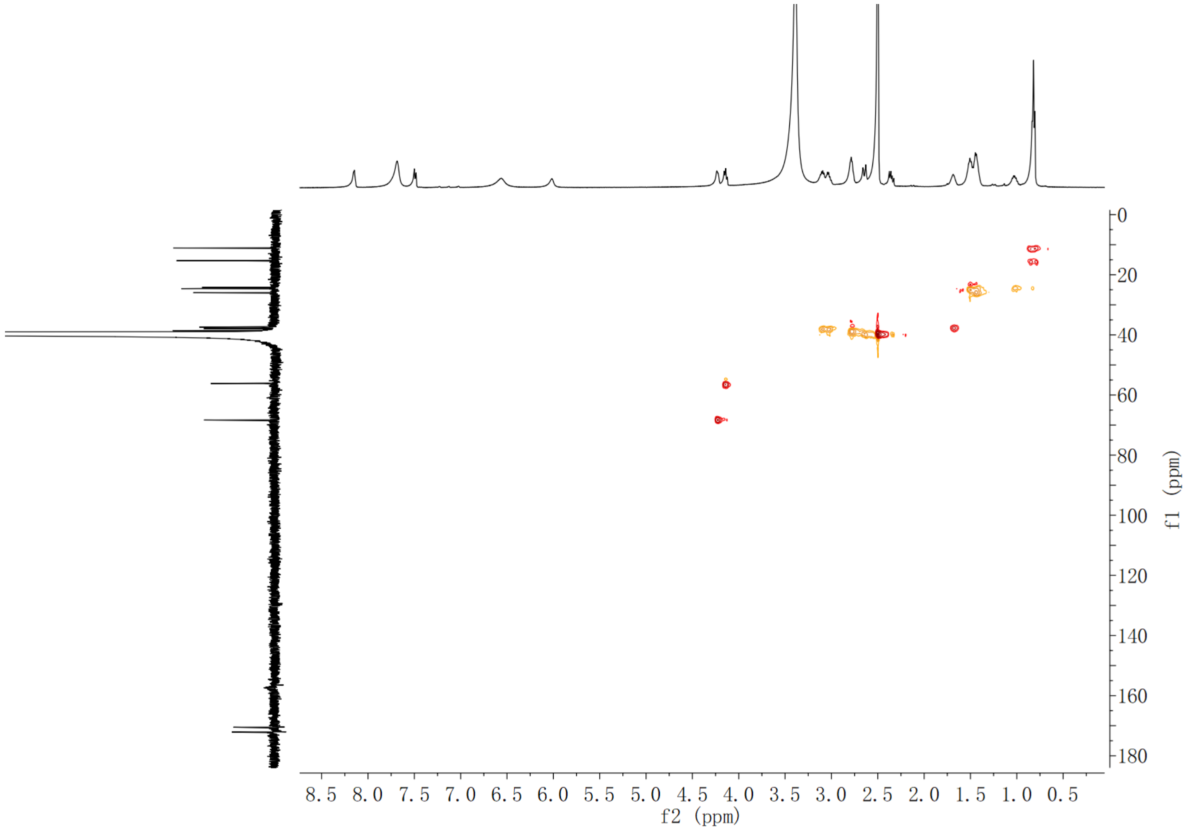

**Supplementary Fig. 86. HSQC spectrum of compound 12 in DMSO-$d_6$**

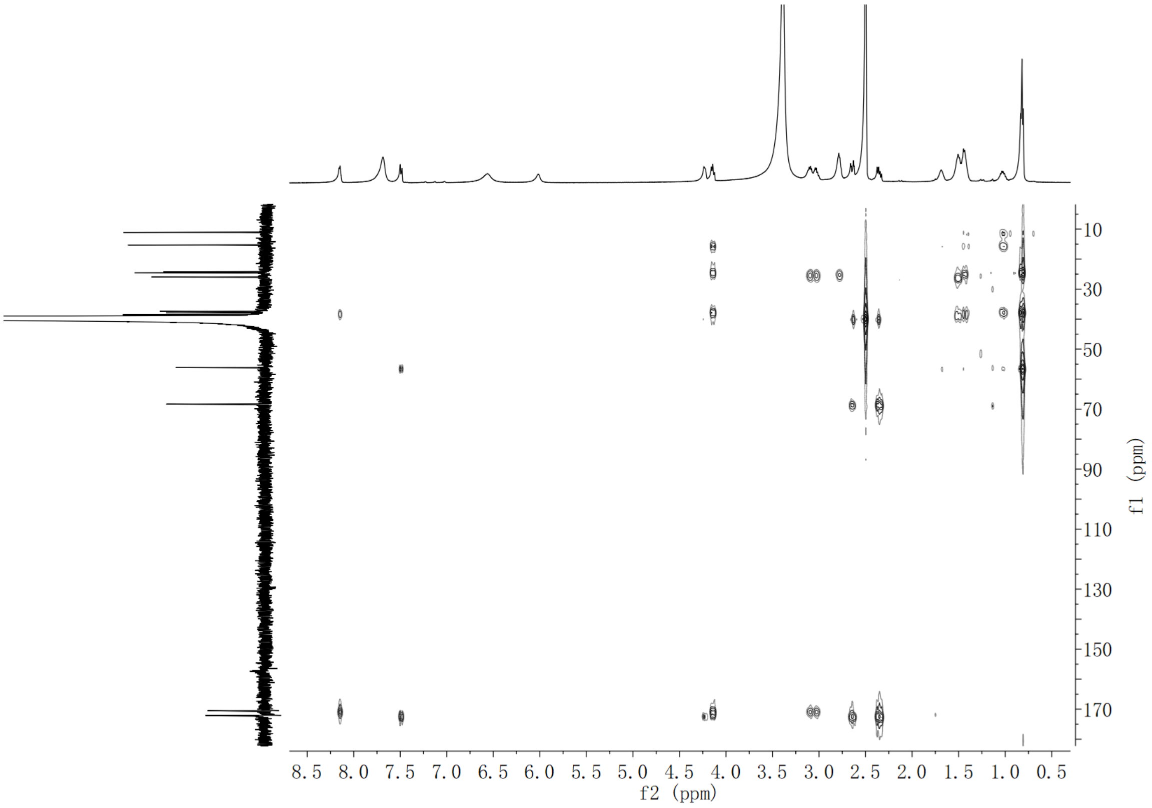

**Supplementary Fig. 87. HMBC spectrum of compound 12 in DMSO-*d*$_6$**

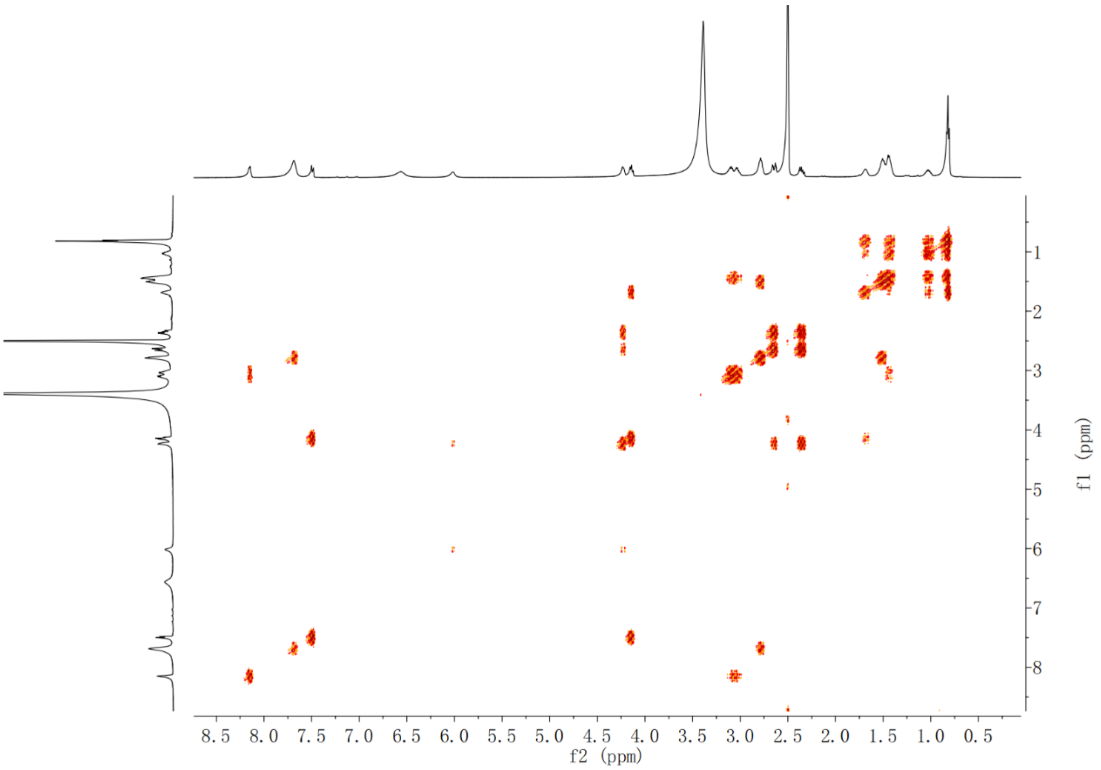

**Supplementary Fig. 88. $^1$H-$^1$H COSY spectrum of compound 12 in DMSO-*d*$_6$**

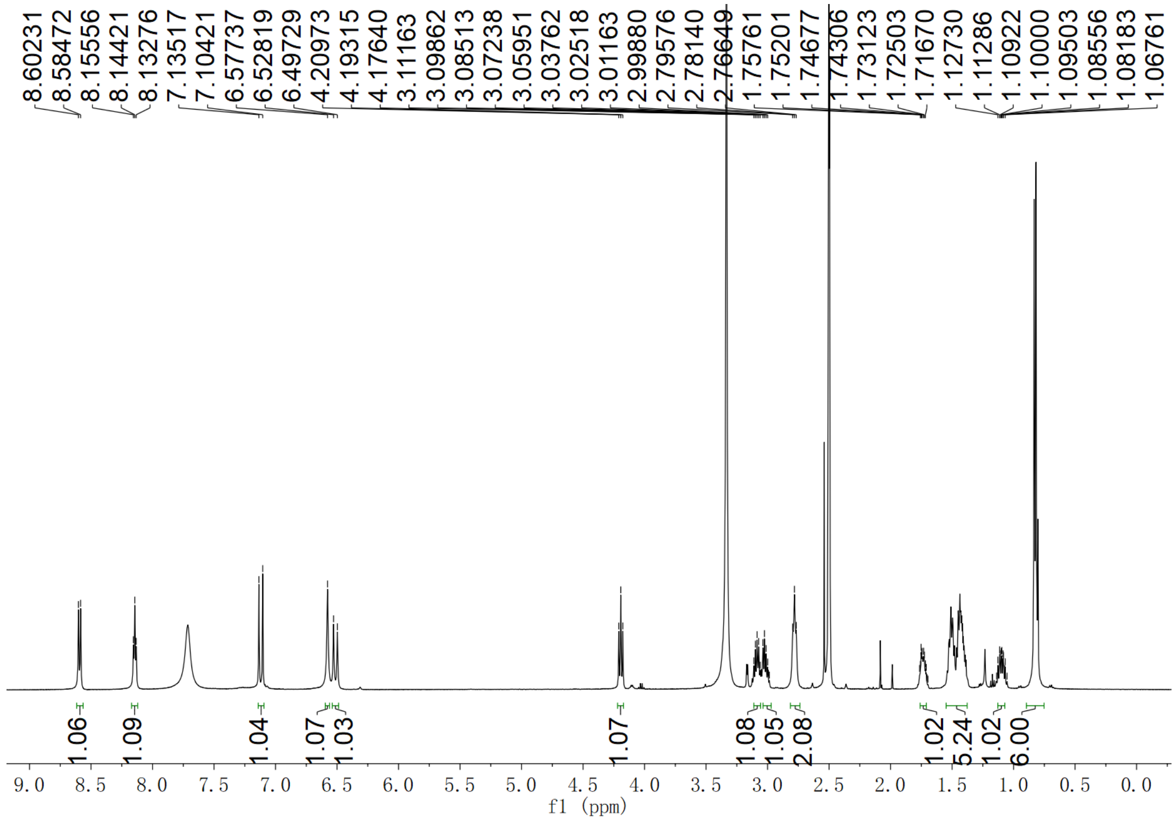

Supplementary Fig. 89. $^{1}H$ NMR spectrum of compound 13 in DMSO-$d_6$

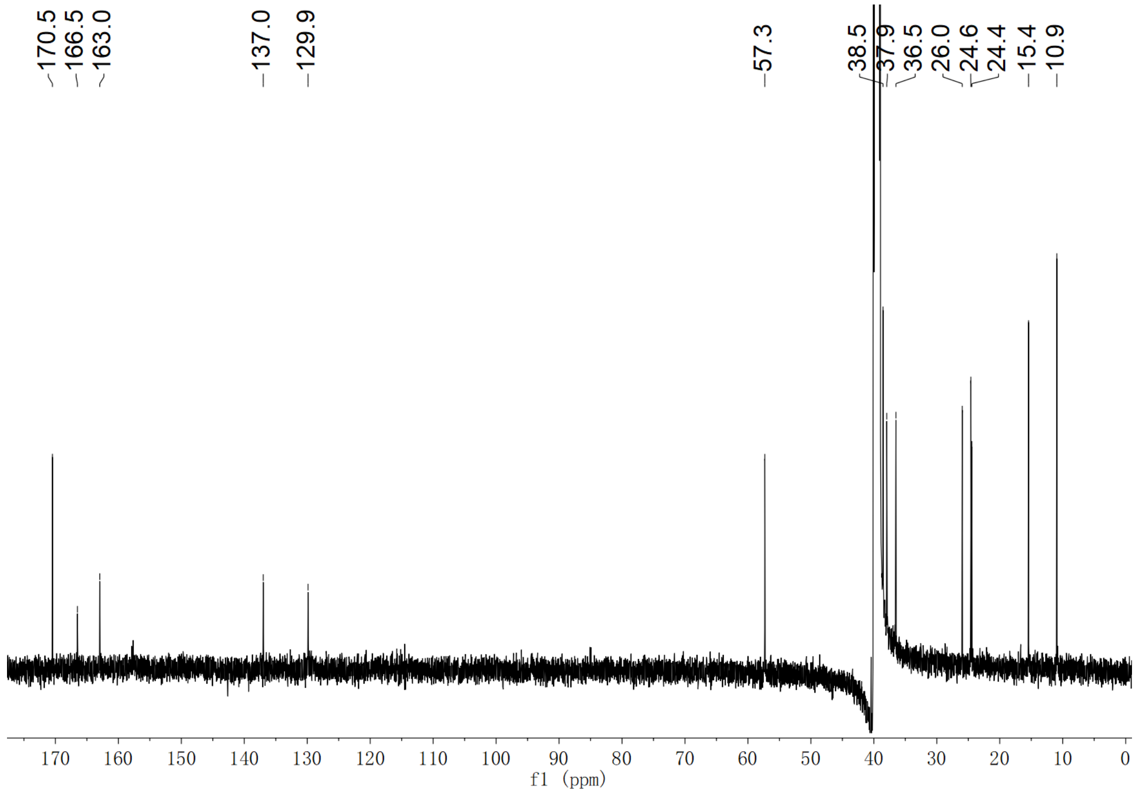

Supplementary Fig. 90. $^{13}C$ NMR spectrum of compound 13 in DMSO-$d_6$

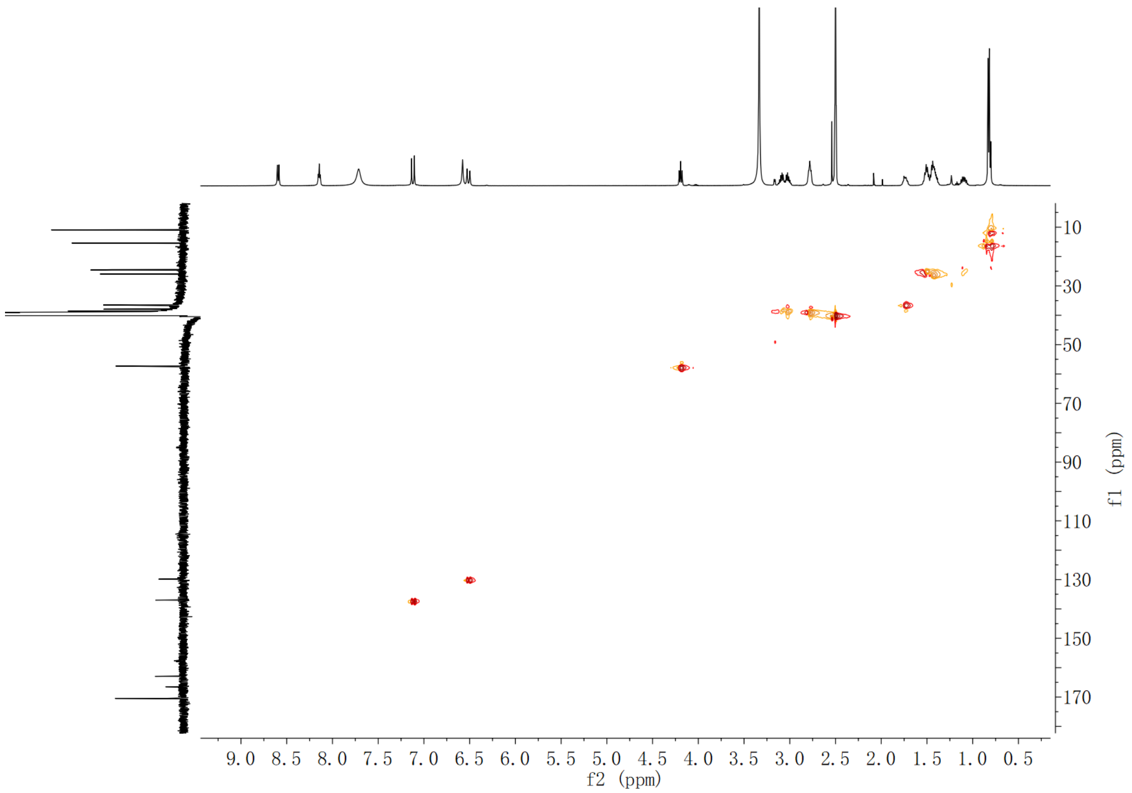

**Supplementary Fig. 91. HSQC spectrum of compound 13 in DMSO-$d_6$**

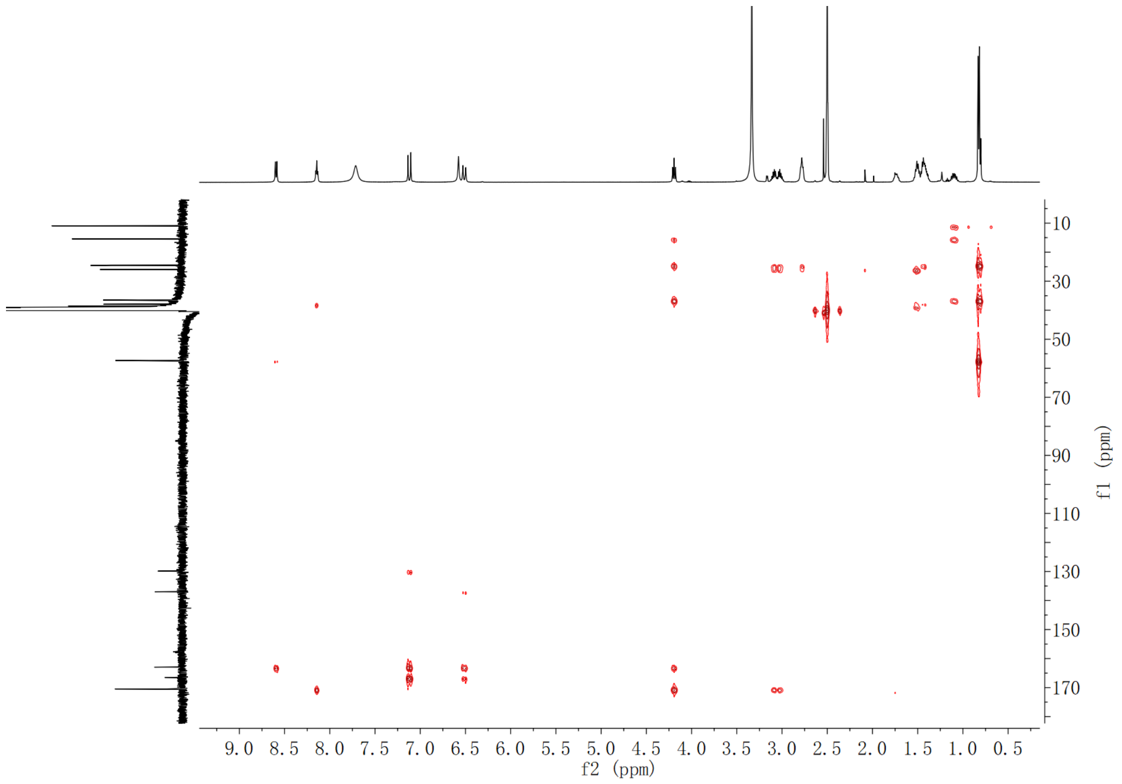

**Supplementary Fig. 92. HMBC spectrum of compound 13 in DMSO-$d_6$**

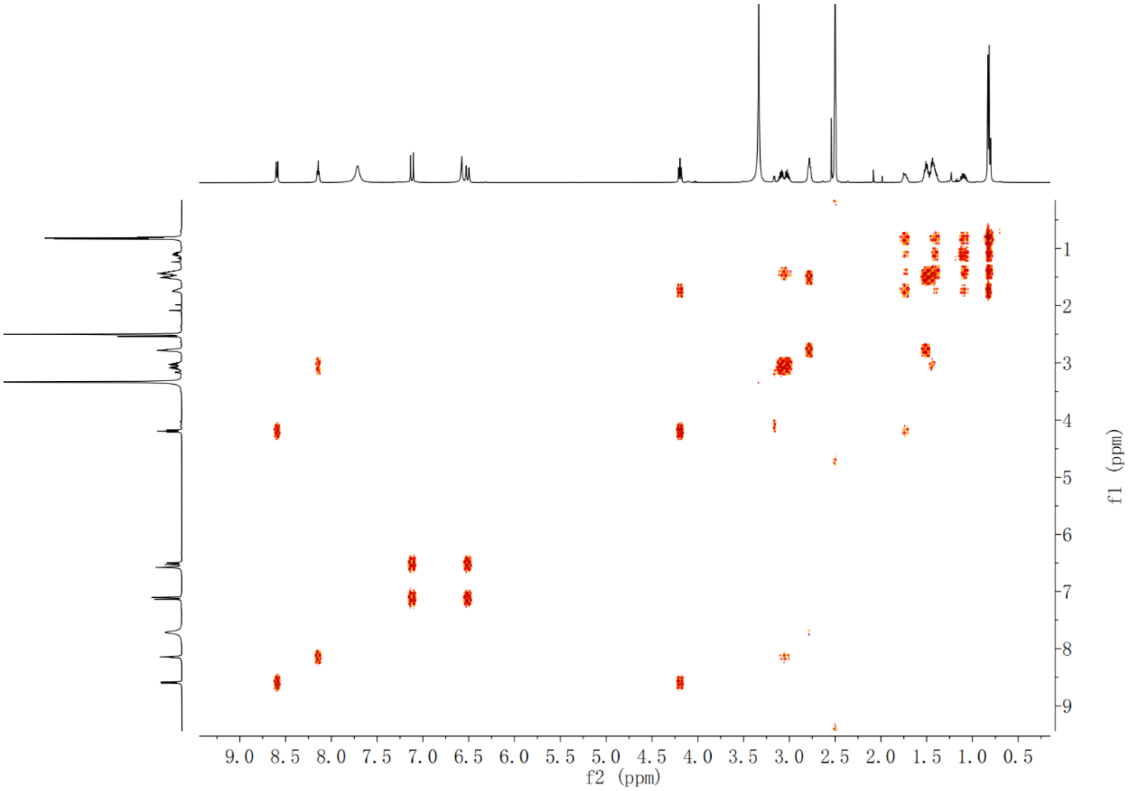

**Supplementary Fig. 93. $^{1}$H-$^{1}$H COSY spectrum of compound 13 in DMSO-$d_6$**

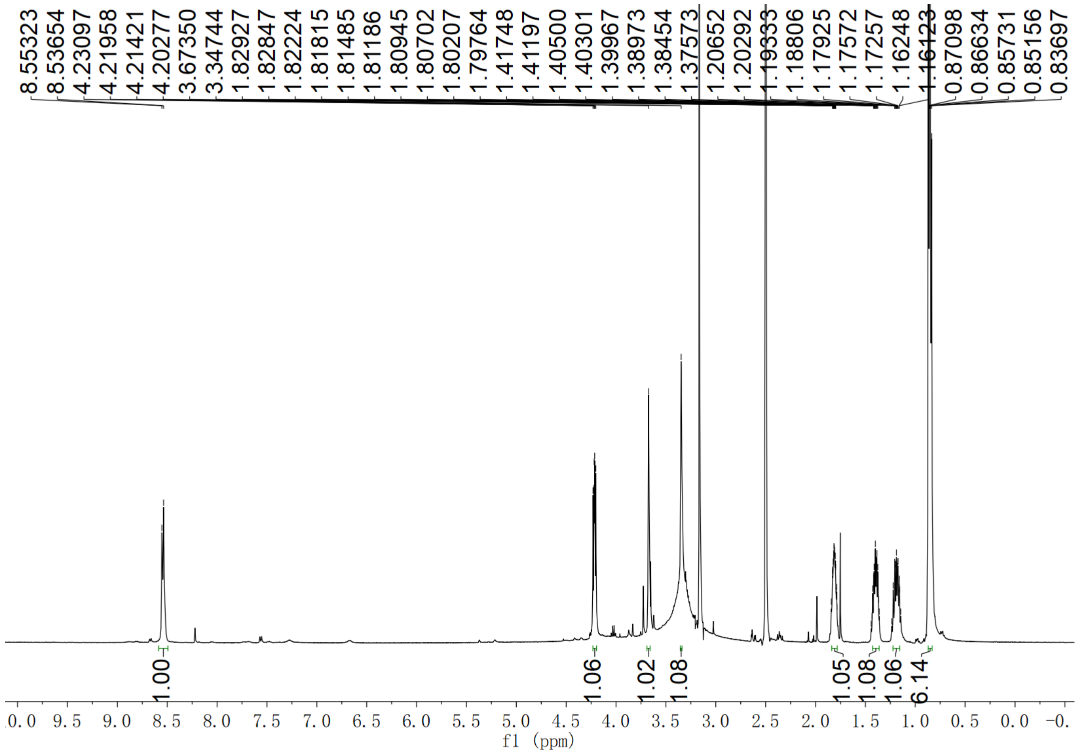

**Supplementary Fig. 94. $^{1}$H NMR spectrum of compound 14 in DMSO-$d_6$**

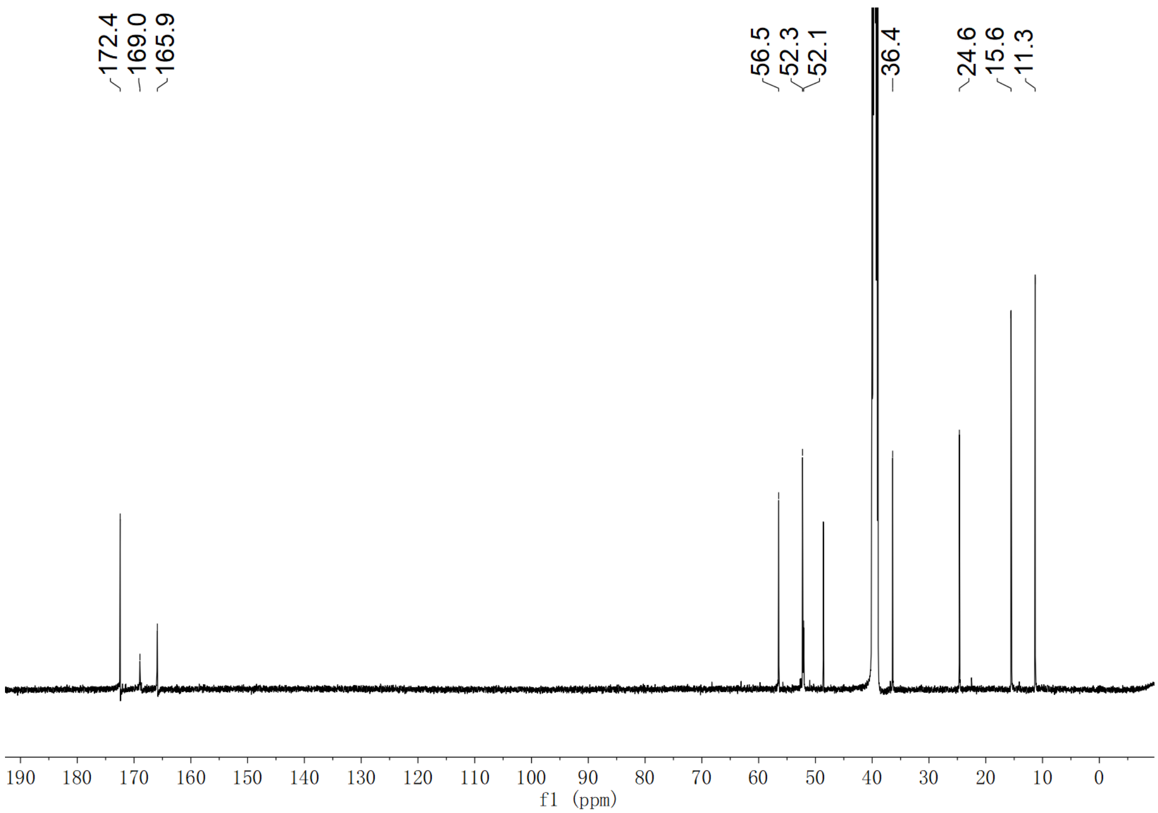

**Supplementary Fig. 95. $^{13}$C NMR spectrum of compound 14 in DMSO-*d*$_6$**

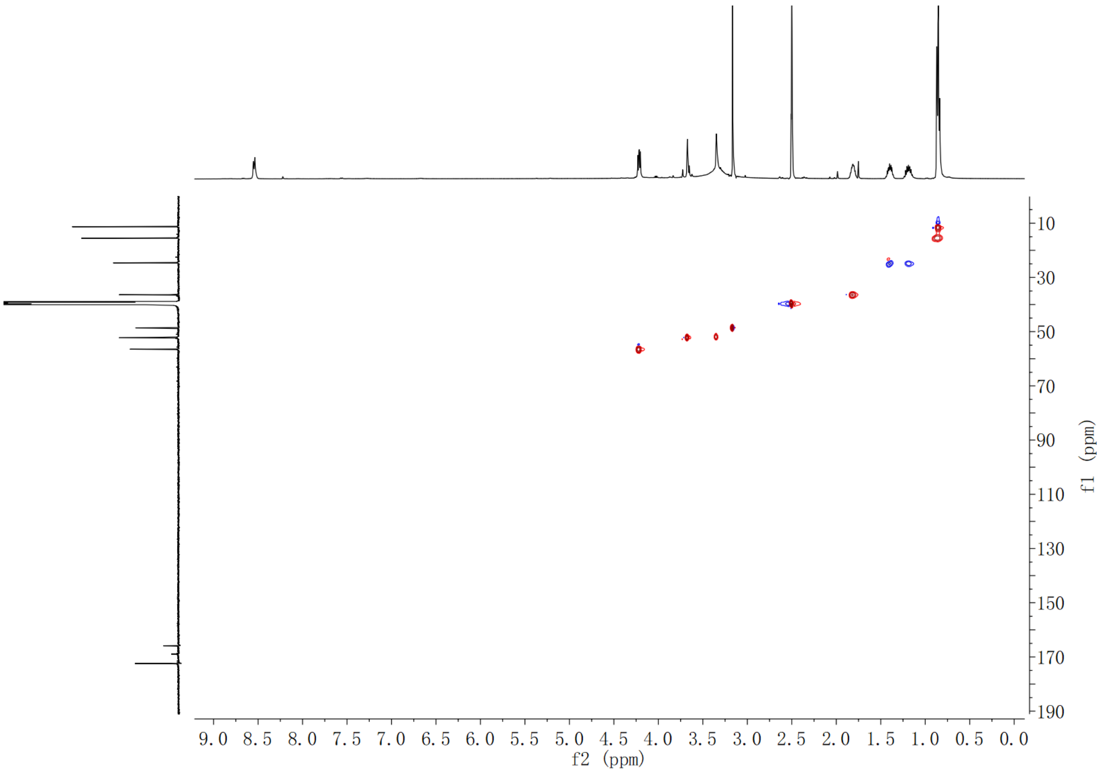

**Supplementary Fig. 96. HSQC spectrum of compound 14 in DMSO-*d*$_6$**

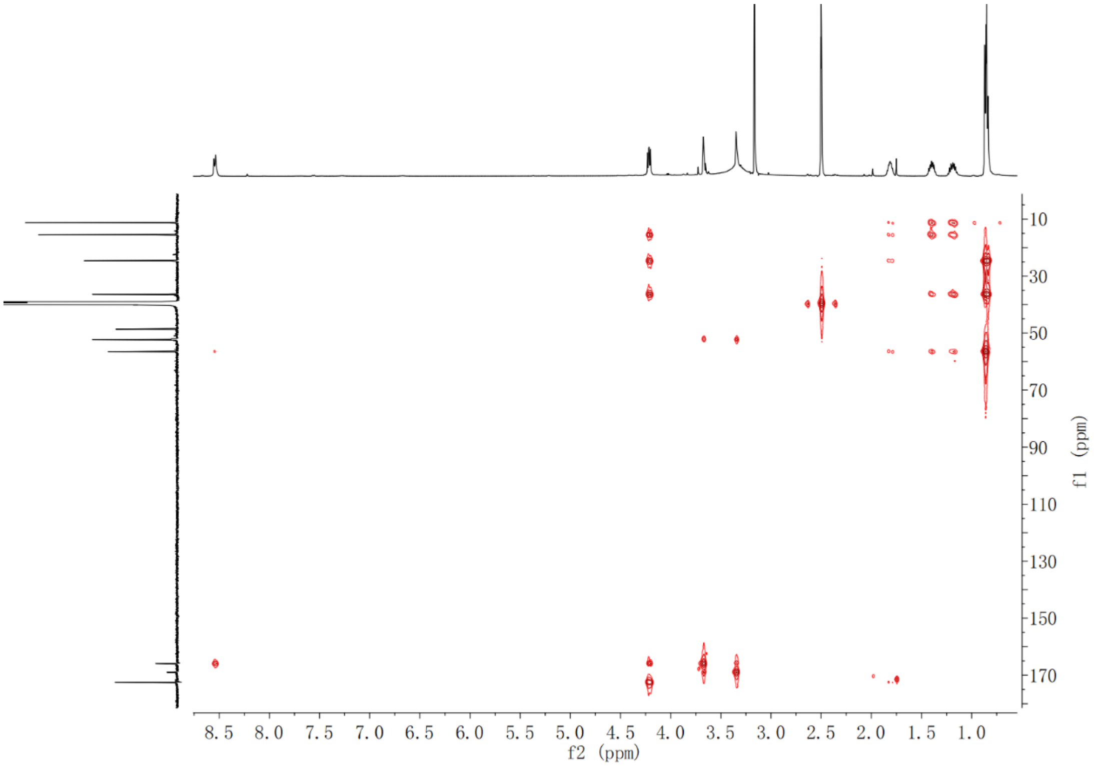

**Supplementary Fig. 97. HMBC spectrum of compound 14 in DMSO-$d_6$**

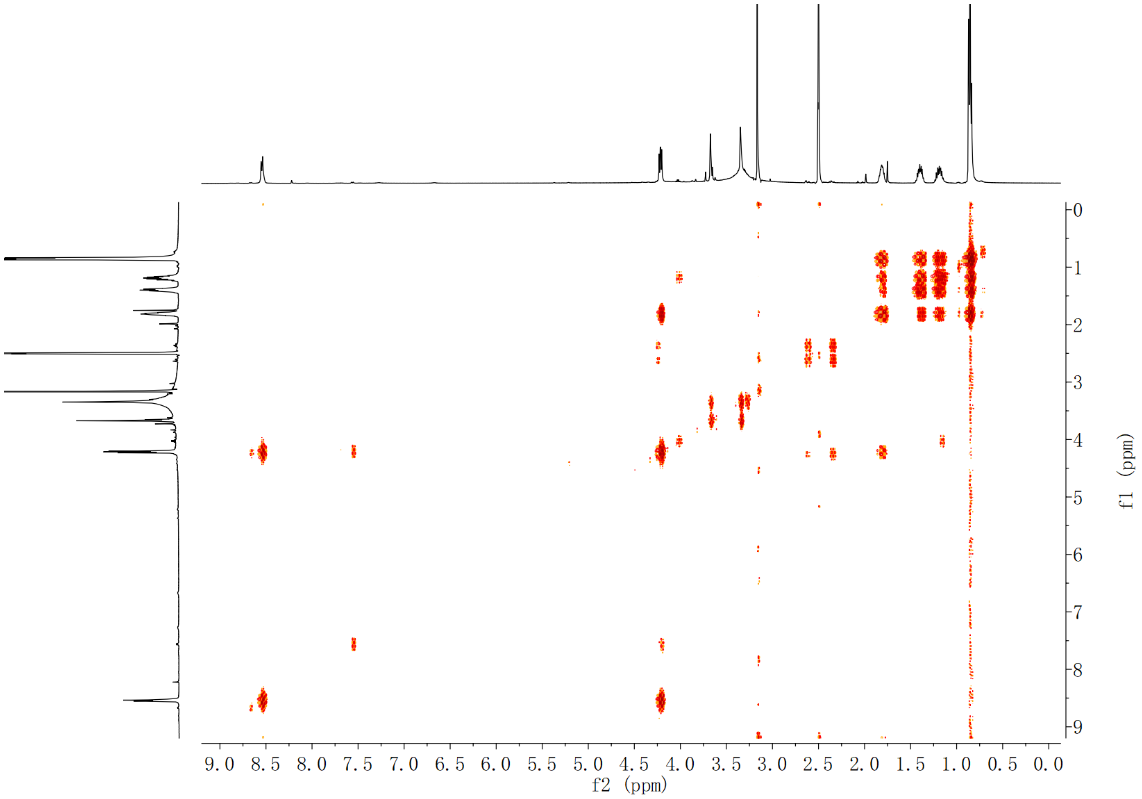

**Supplementary Fig. 98. $^1$H-$^1$H COSY spectrum of compound 14 in DMSO-$d_6$**

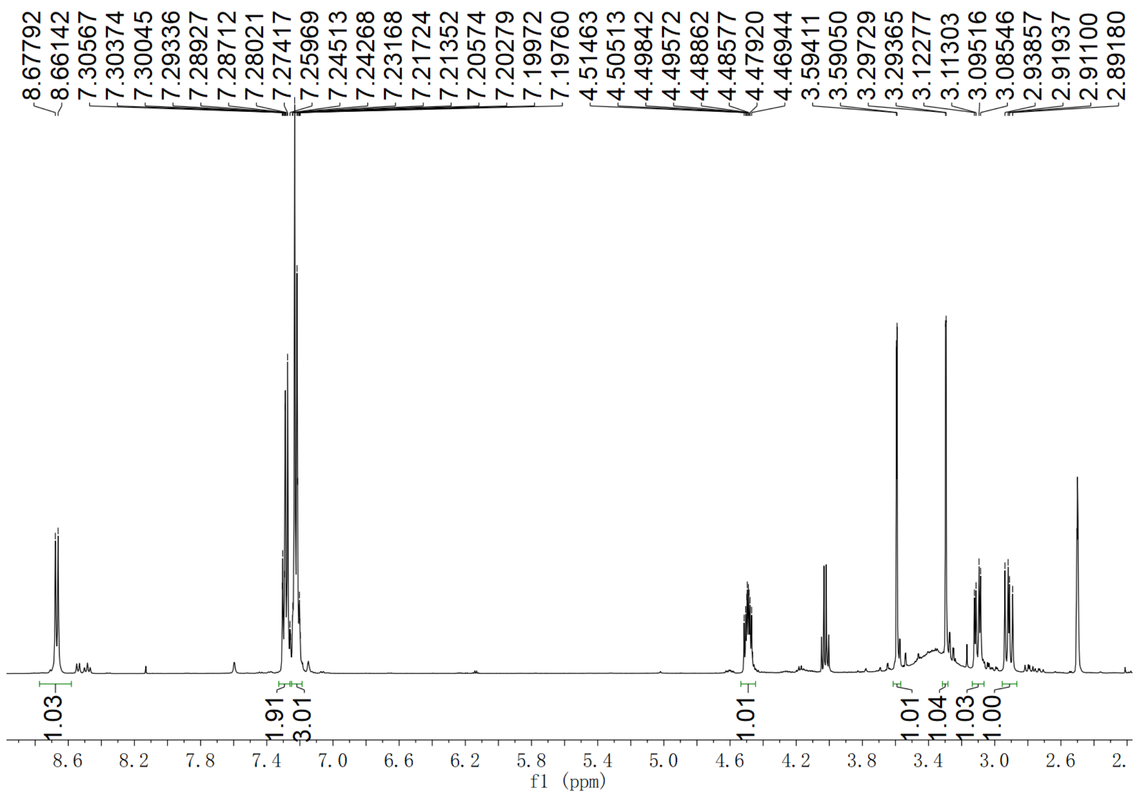

**Supplementary Fig. 99.** $^{1}H$ **NMR spectrum of compound 15 in DMSO-$d_6$**

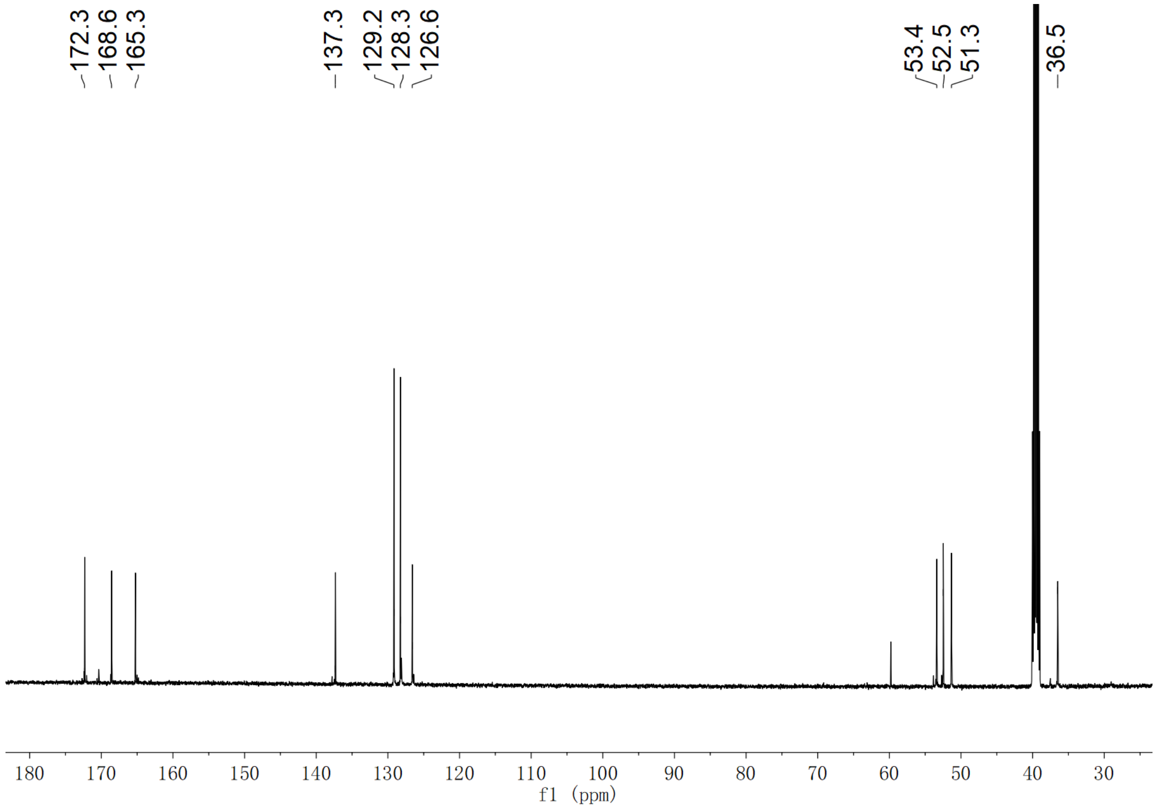

**Supplementary Fig. 100.** $^{13}C$ **NMR spectrum of compound 15 in DMSO-$d_6$**

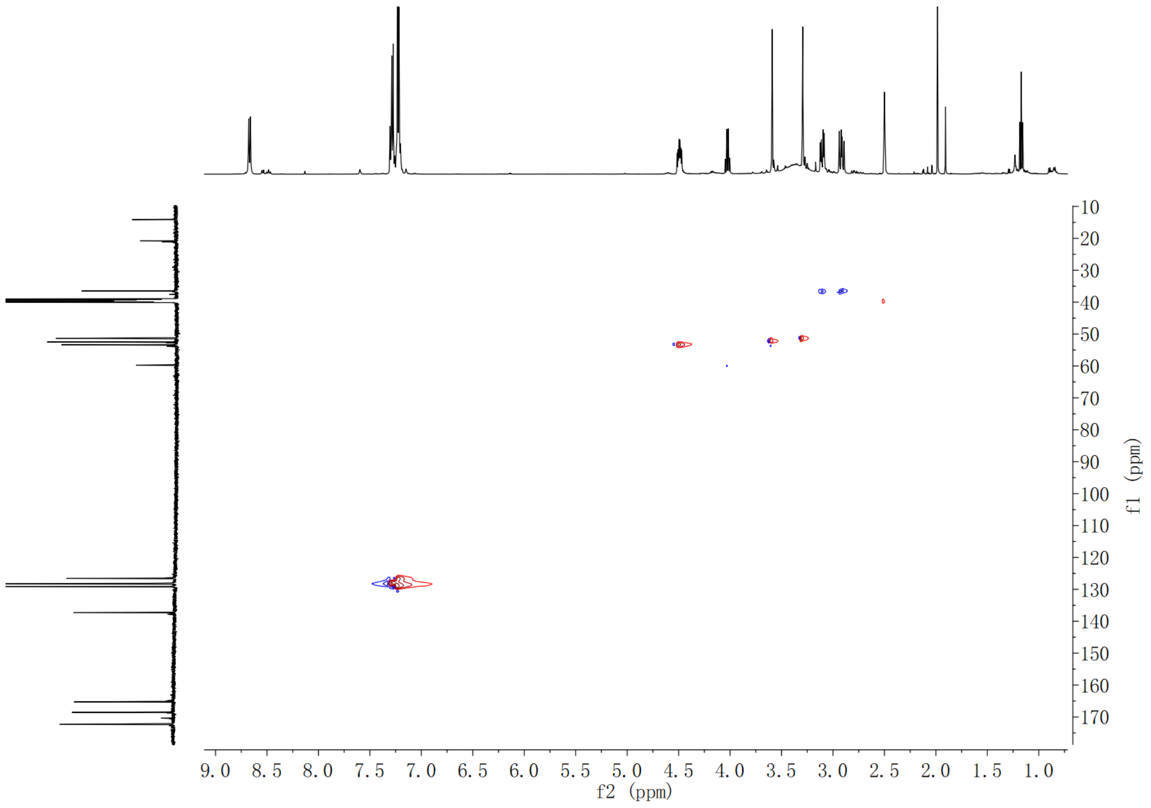

**Supplementary Fig. 101. HSQC spectrum of compound 15 in DMSO-*d*$_6$**

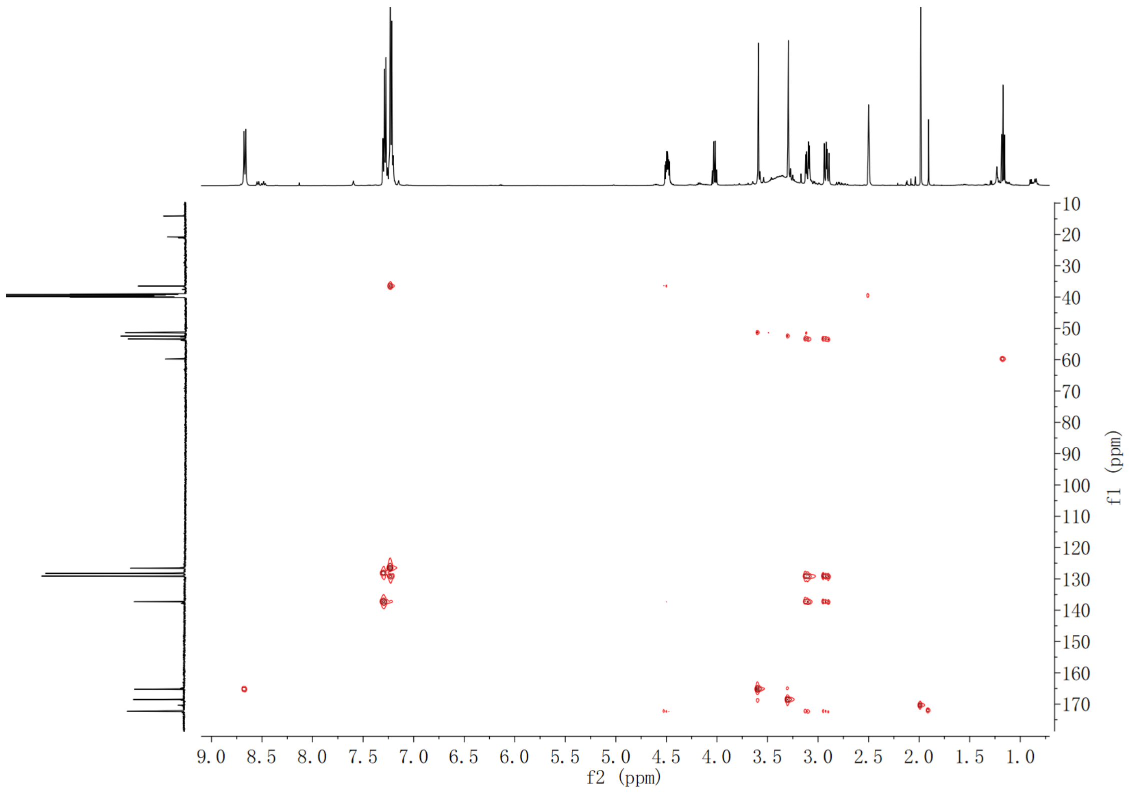

**Supplementary Fig. 102. HMBC spectrum of compound 15 in DMSO-*d*$_6$**

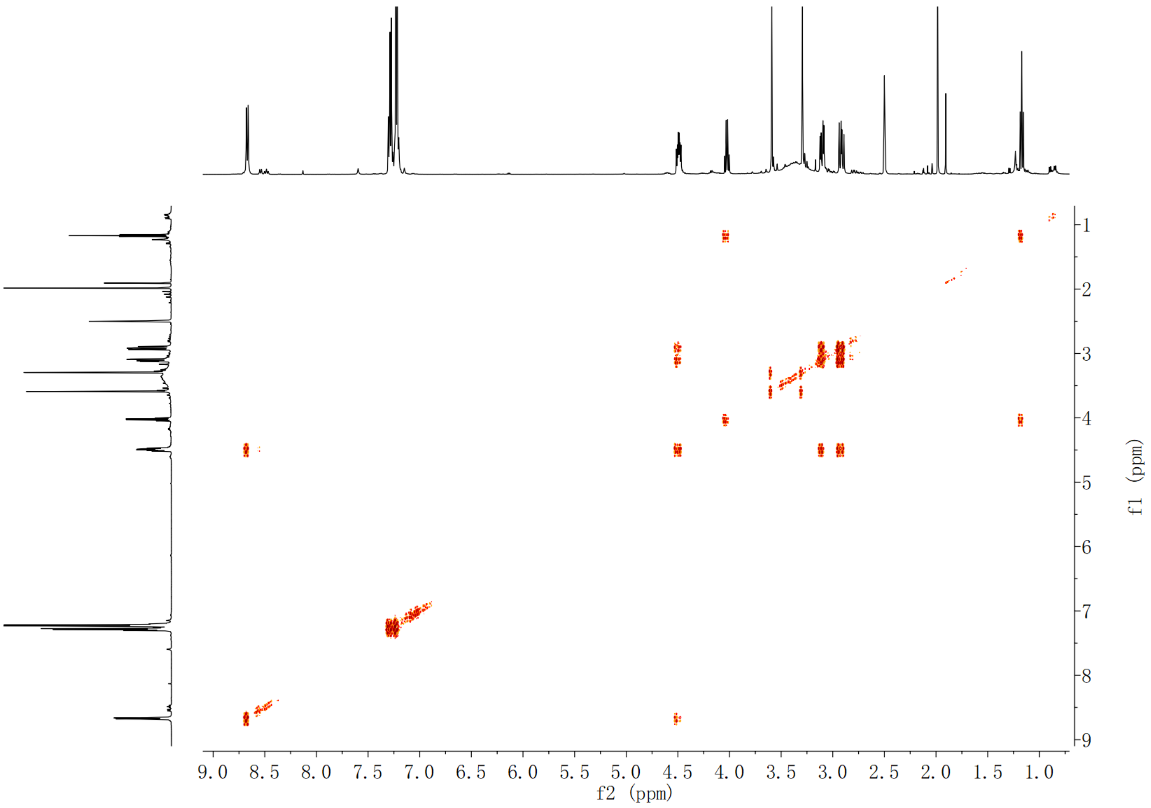

**Supplementary Fig. 103. $^{1}$H-$^{1}$H COSY spectrum of compound 15 in DMSO-*d*$_6$**

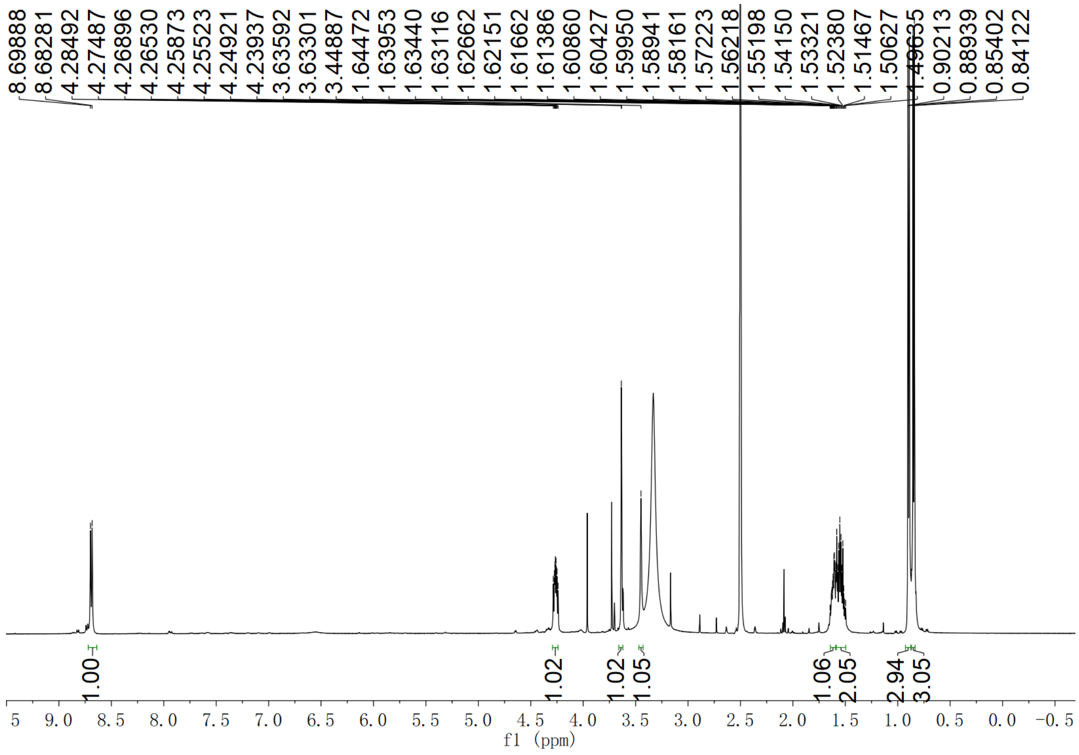

**Supplementary Fig. 104. $^{1}$H NMR spectrum of compound (2*S*,3*S*)-*t*-ES-Leu in DMSO-*d*$_6$**

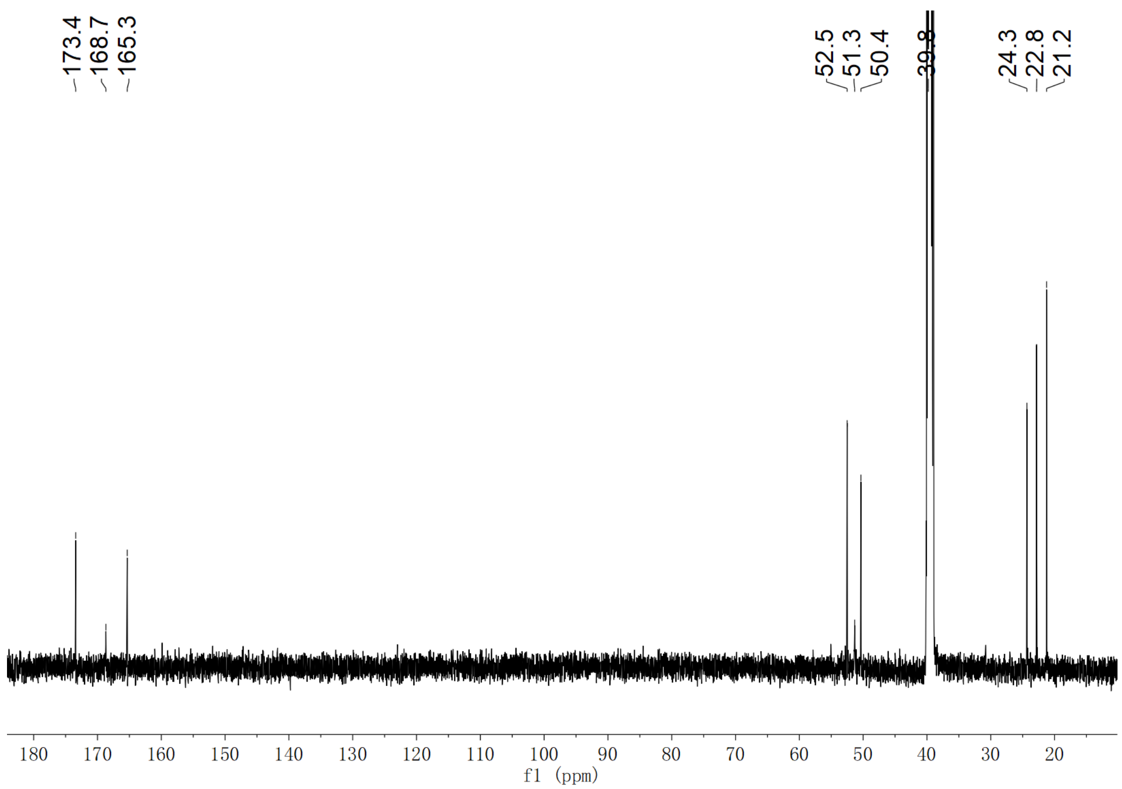

**Supplementary Fig. 105.** $^{13}$**C NMR spectrum of compound (2*S*,3*S*)-*t*-ES-Leu in DMSO-*d*$_6$**

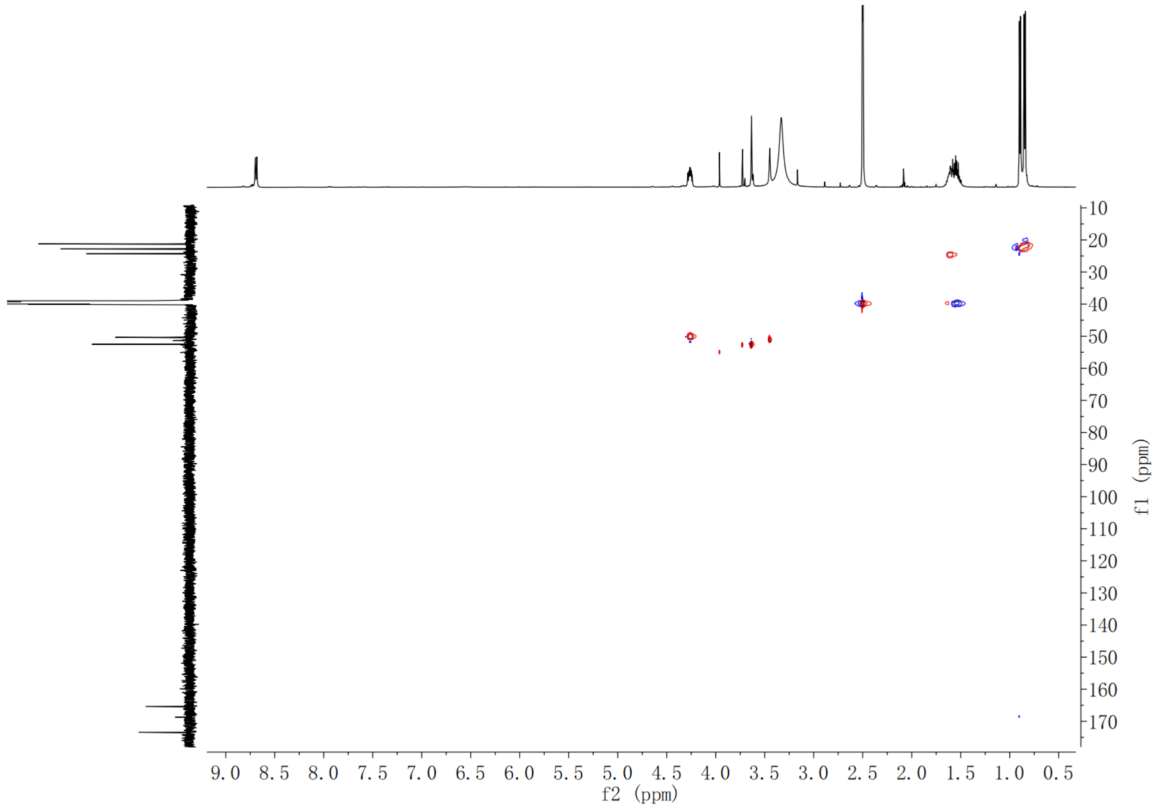

**Supplementary Fig. 106. HSQC spectrum of compound (2*S*,3*S*)-*t*-ES-Leu in DMSO-*d*$_6$**

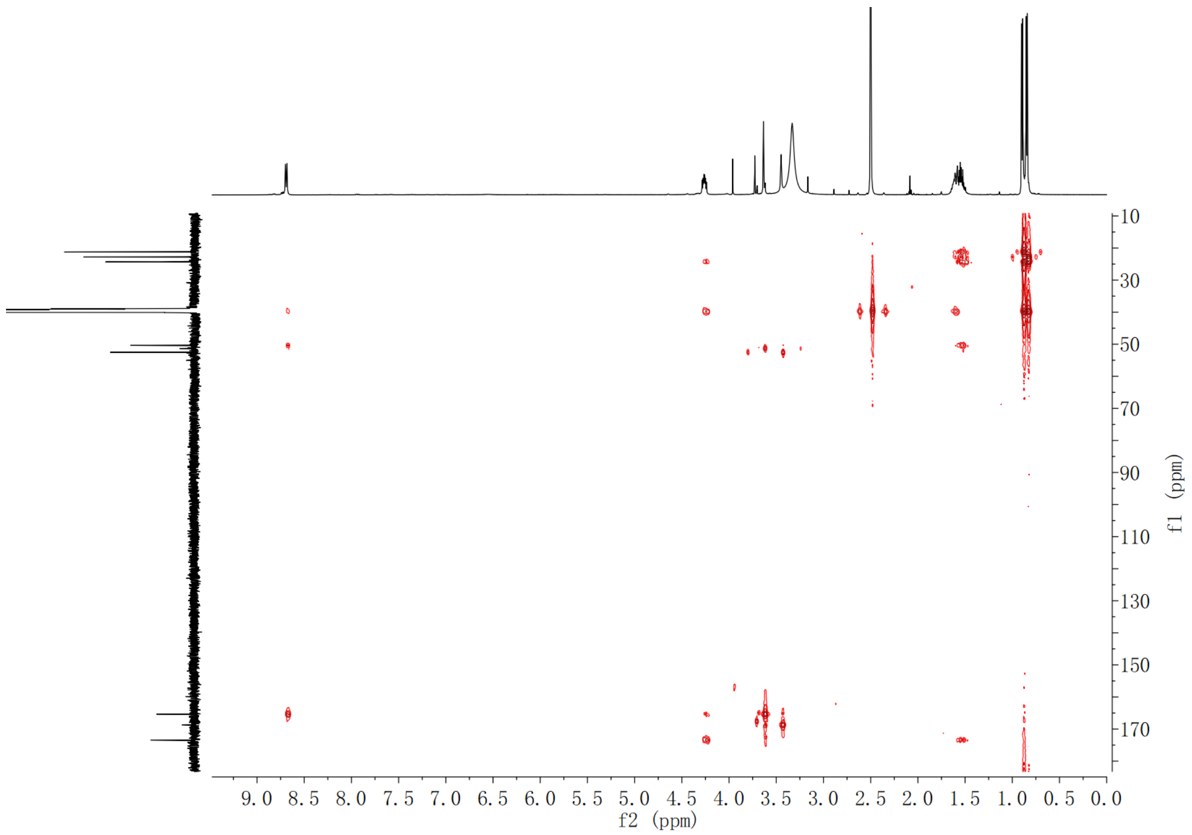

**Supplementary Fig. 107. HMBC spectrum of compound (2*S*,3*S*)-*t*-ES-Leu in DMSO-*d*$_6$**

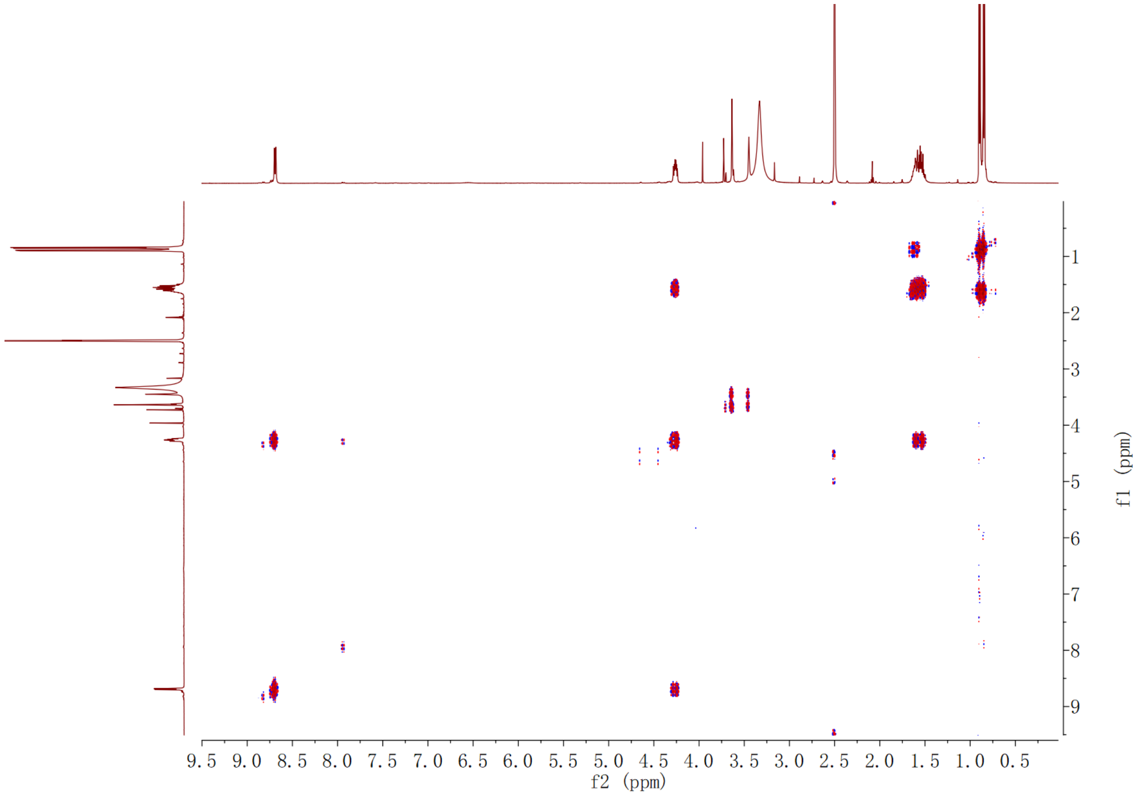

**Supplementary Fig. 108. $^1$H-$^1$H COSY spectrum of compound (2*S*,3*S*)-*t*-ES-Leu in DMSO-*d*$_6$**

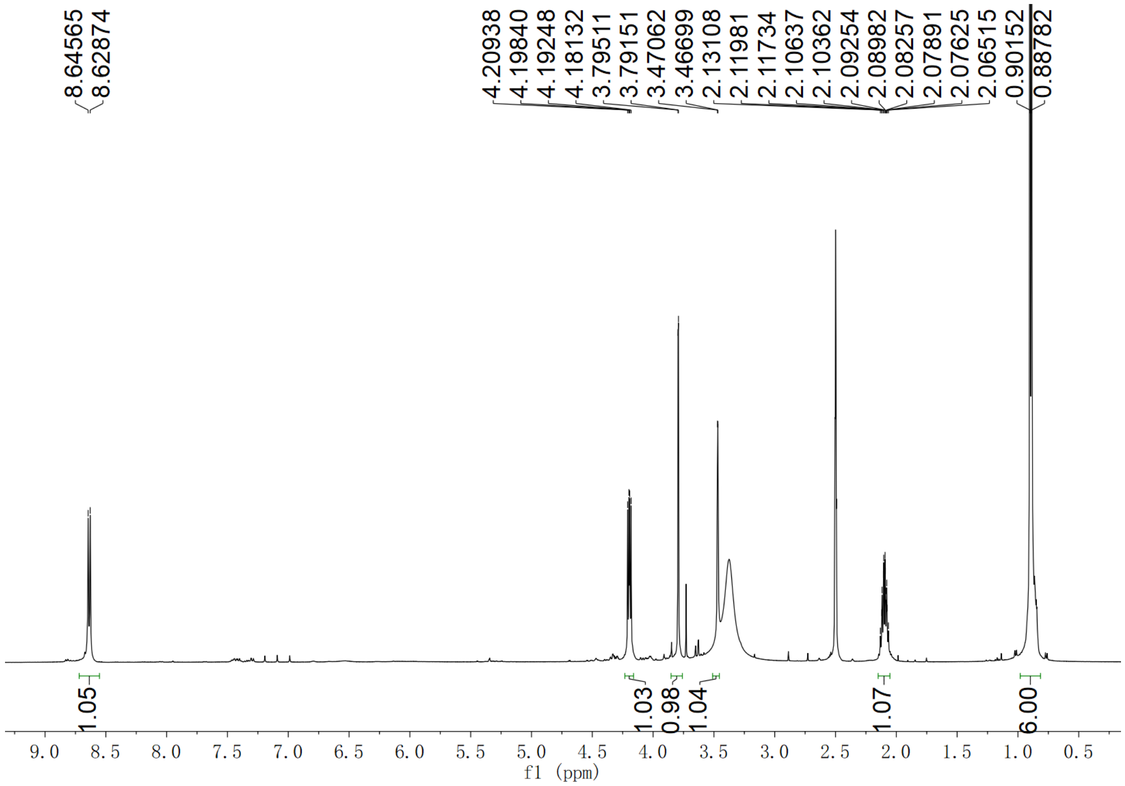

**Supplementary Fig. 109. $^{1}$H NMR spectrum of compound (2*S*,3*S*)-*t*-ES-Val in DMSO-$d_6$**

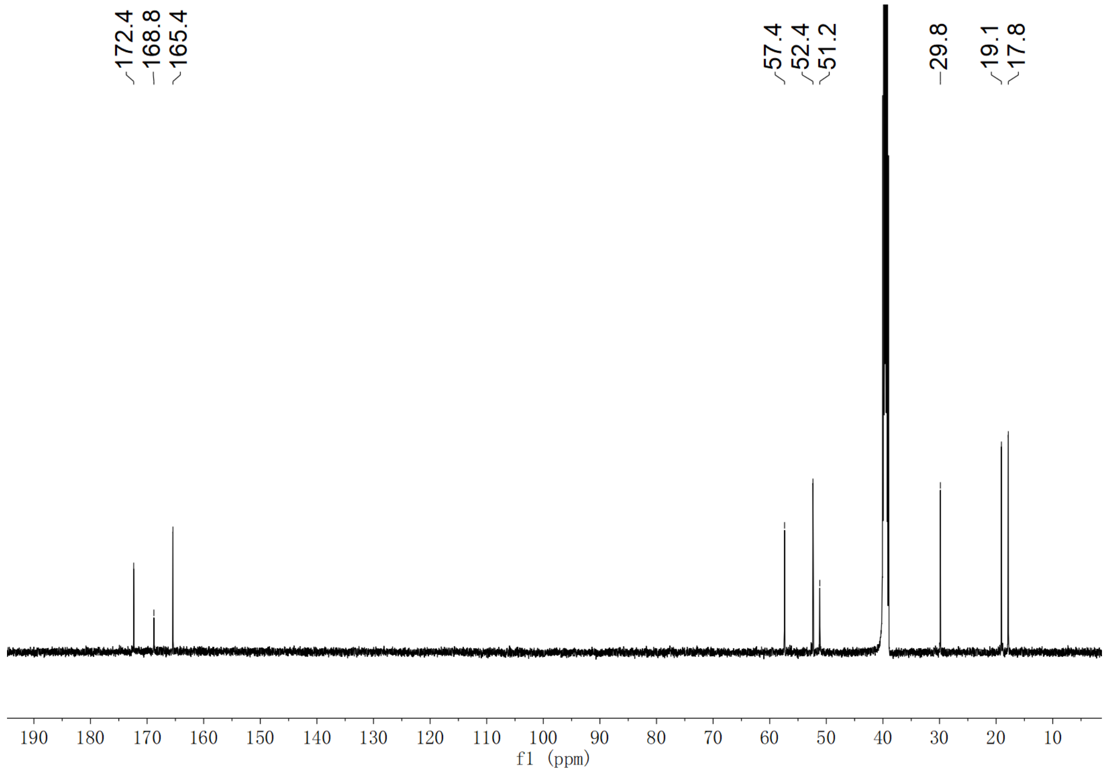

**Supplementary Fig. 110. $^{13}$C NMR spectrum of compound (2*S*,3*S*)-*t*-ES-Val in DMSO-$d_6$**

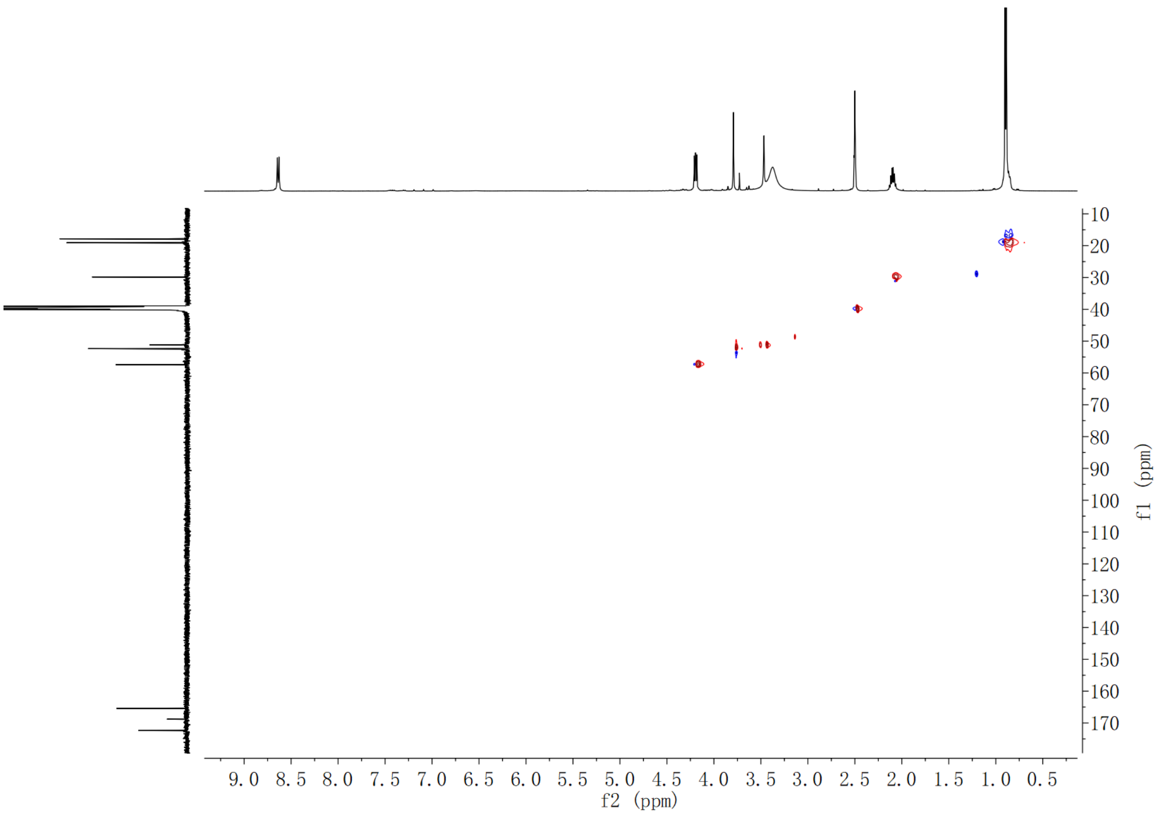

**Supplementary Fig. 111. HSQC spectrum of compound (2*S*,3*S*)-*t*-ES-Val in DMSO-*d*$_6$**

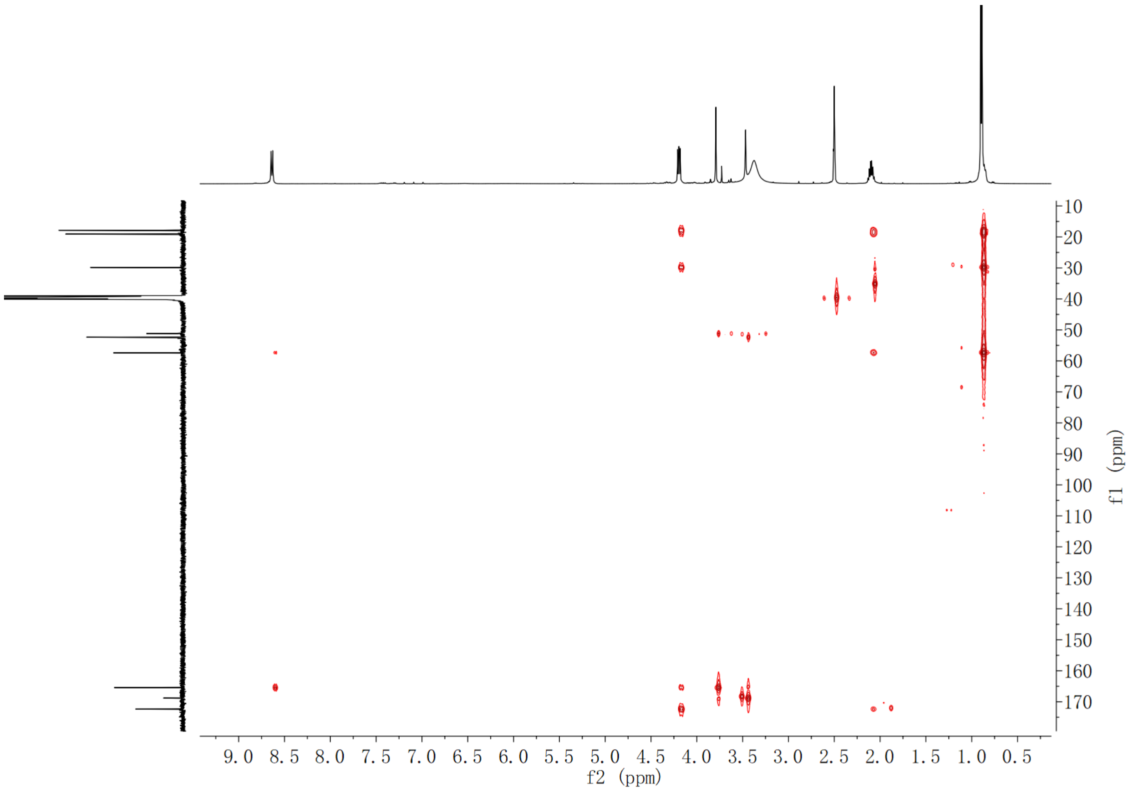

**Supplementary Fig. 112. HMBC spectrum of compound (2*S*,3*S*)-*t*-ES-Val in DMSO-*d*$_6$**

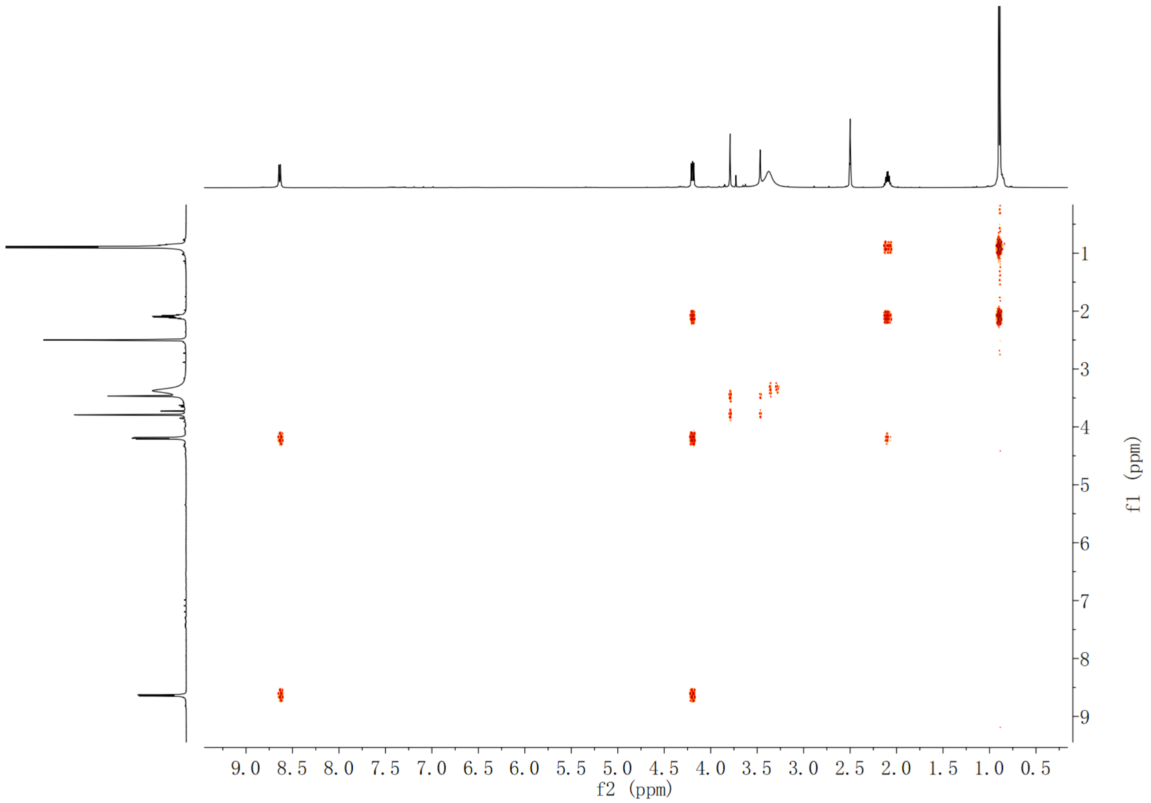

**Supplementary Fig. 113.** $^{1}$**H-**$^{1}$**H COSY spectrum of compound (2*S*,3*S*)-*t*-ES-Val in DMSO-*d*$_6$**

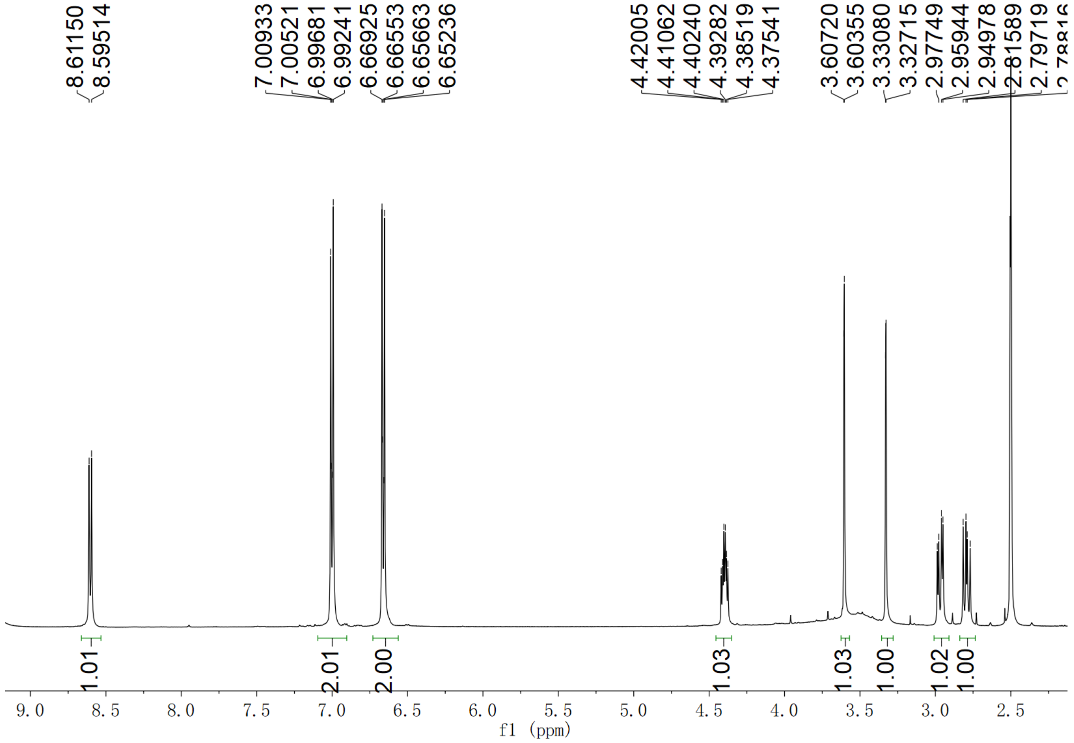

**Supplementary Fig. 114.** $^{1}$**H NMR spectrum of compound (2*S*,3*S*)-*t*-ES-Tyr in DMSO-*d*$_6$**

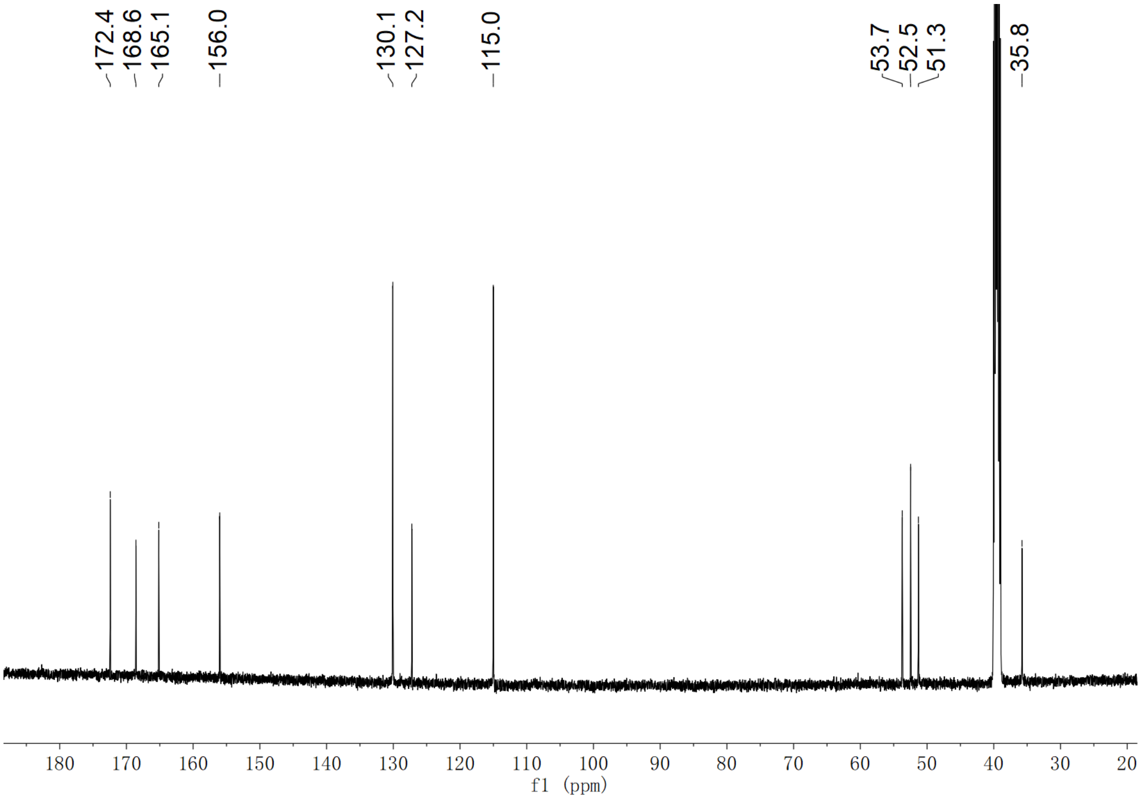

**Supplementary Fig. 115. $^{13}$C NMR spectrum of compound (2*S*,3*S*)-*t*-ES-Tyr in DMSO-$d_6$**

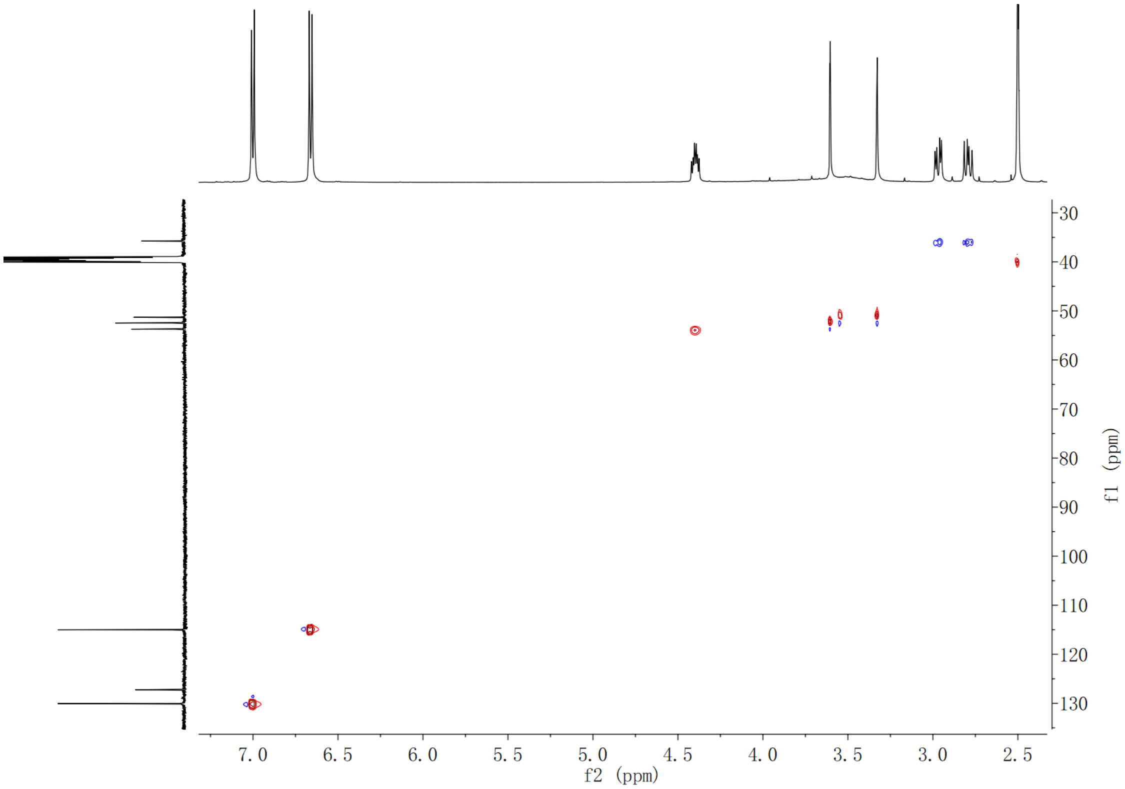

**Supplementary Fig. 116. HSQC spectrum of compound (2*S*,3*S*)-*t*-ES-Tyr in DMSO-$d_6$**

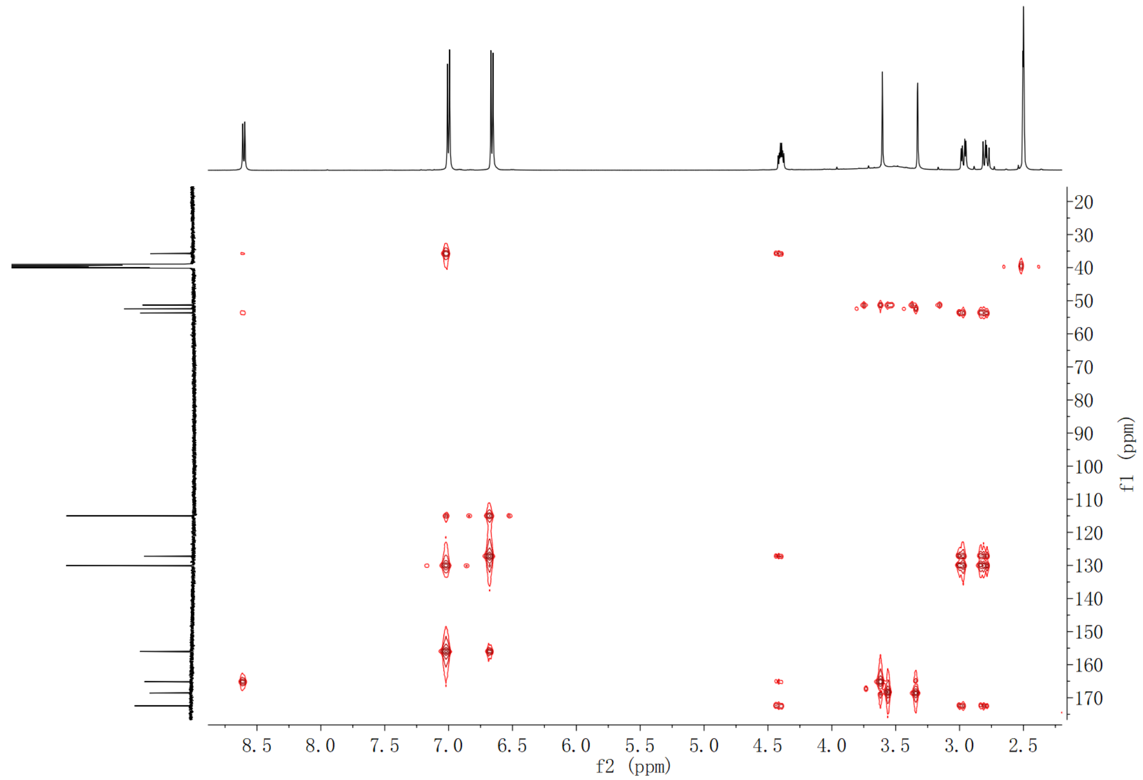

**Supplementary Fig. 117. HMBC spectrum of compound (2*S*,3*S*)-*t*-ES-Tyr in DMSO-$d_6$**

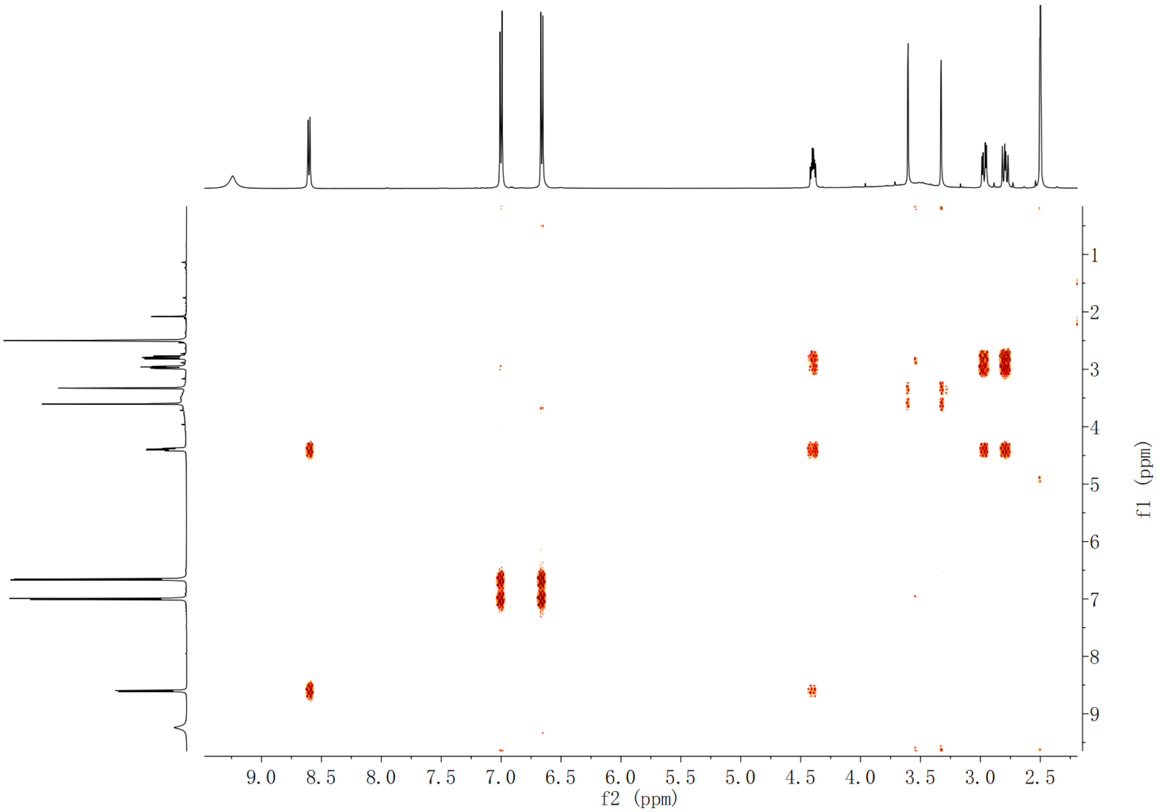

**Supplementary Fig. 118. $^{1}$H-$^{1}$H COSY spectrum of compound (2*S*,3*S*)-*t*-ES-Tyr in DMSO-$d_6$**

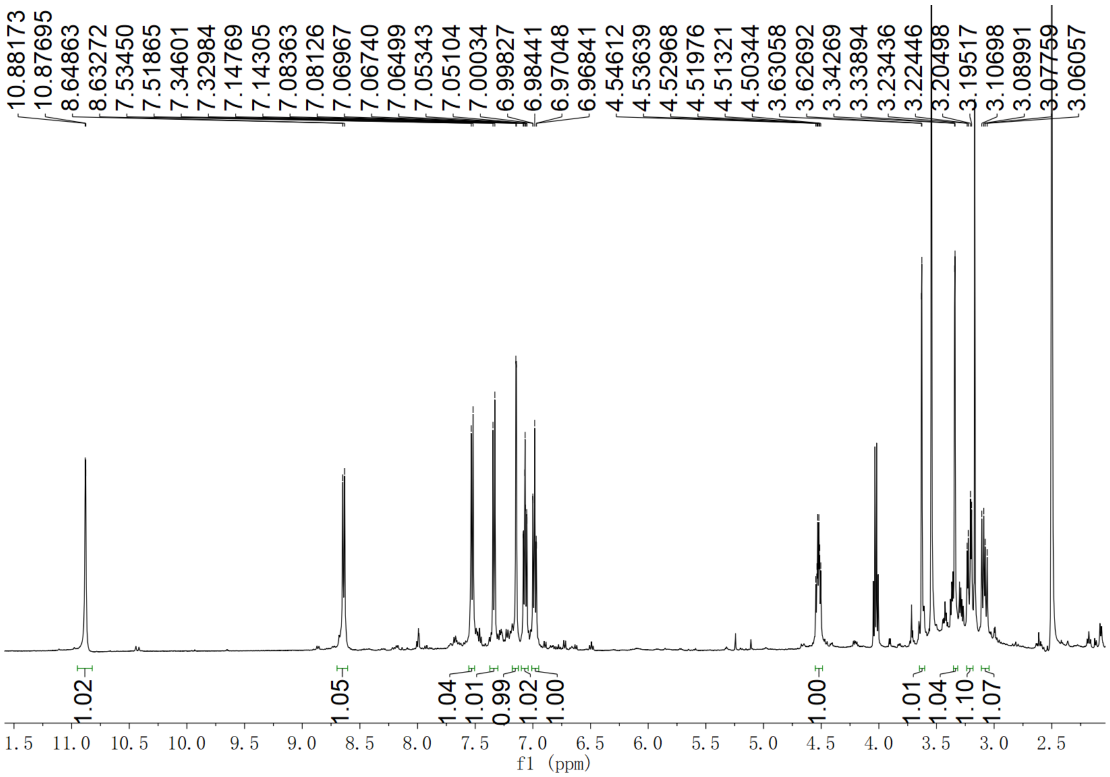

Supplementary Fig. 119. $^{1}$H NMR spectrum of compound (2*S*,3*S*)-*t*-ES-Trp in DMSO-$d_6$

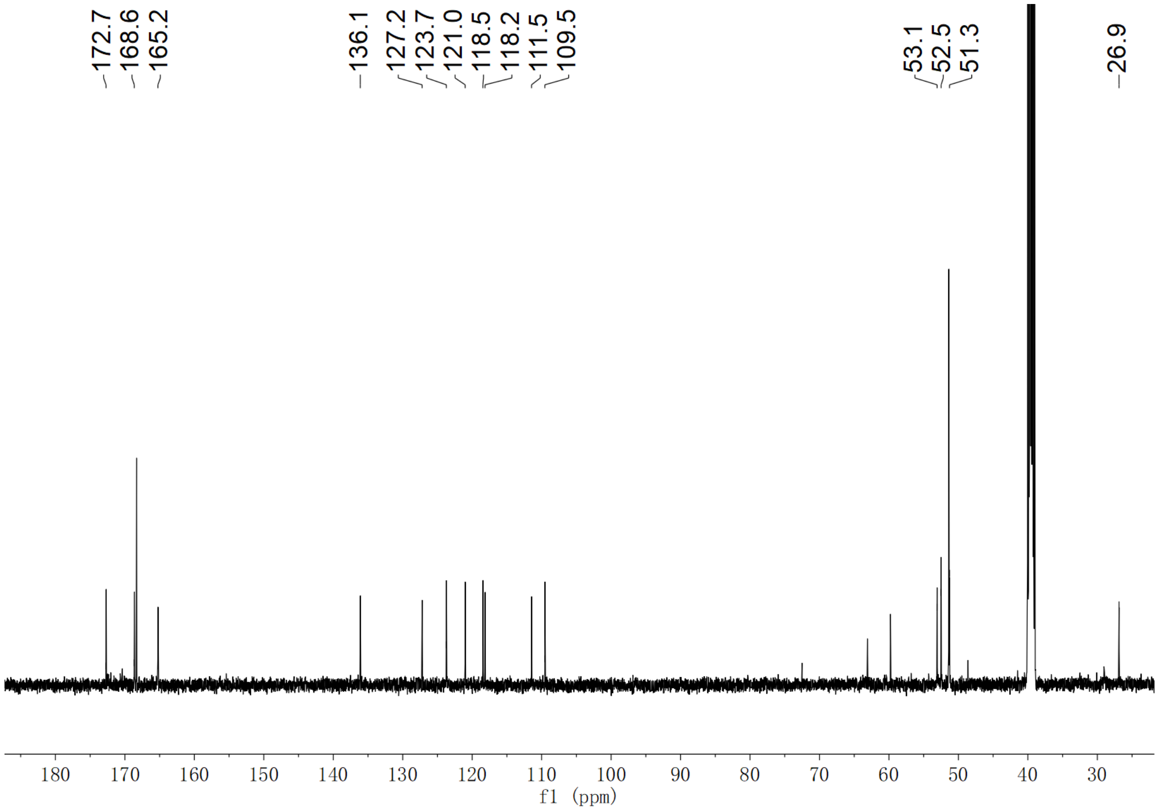

Supplementary Fig. 120. $^{13}$C NMR spectrum of compound (2*S*,3*S*)-*t*-ES-Trp in DMSO-$d_6$

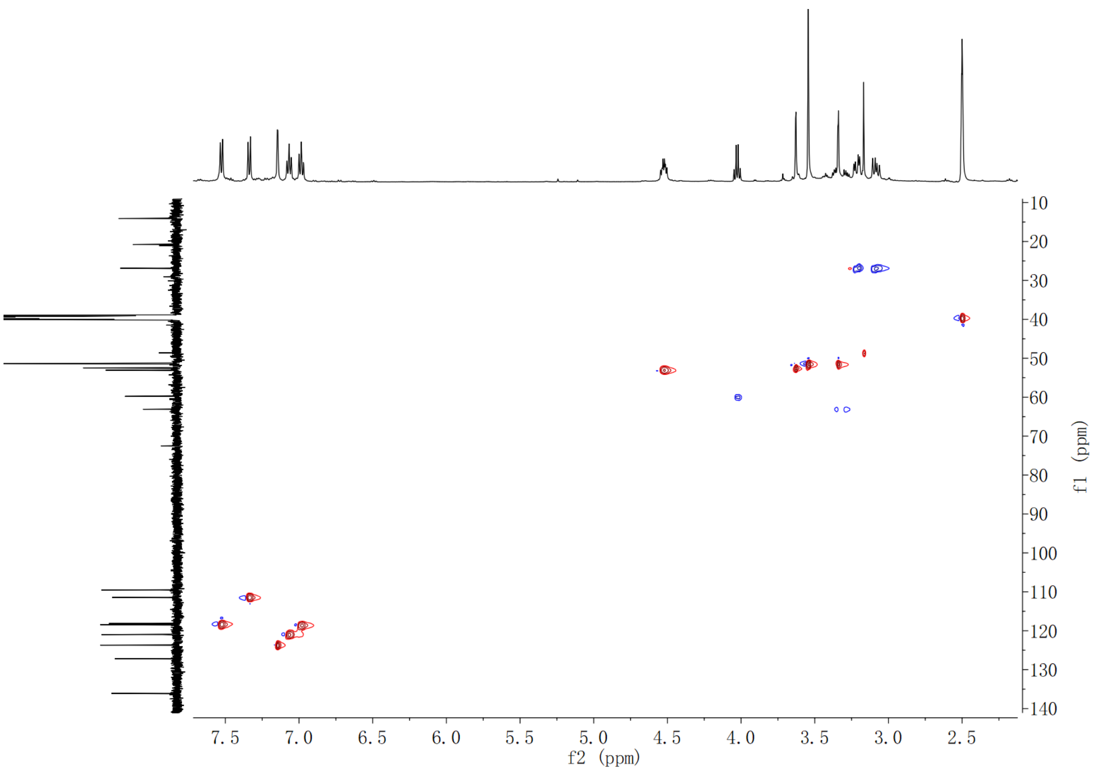

**Supplementary Fig. 121. HSQC spectrum of compound (2*S*,3*S*)-*t*-ES-Trp in DMSO-$d_6$**

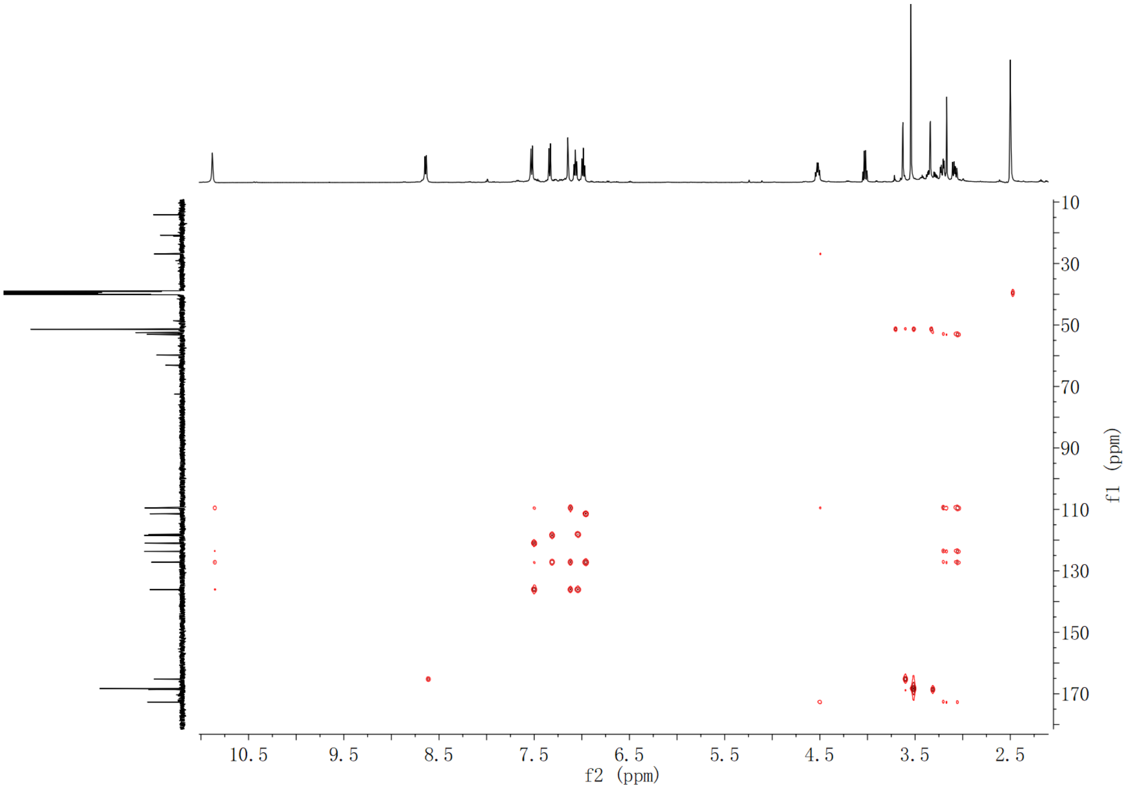

**Supplementary Fig. 122. HMBC spectrum of compound (2*S*,3*S*)-*t*-ES-Trp in DMSO-$d_6$**

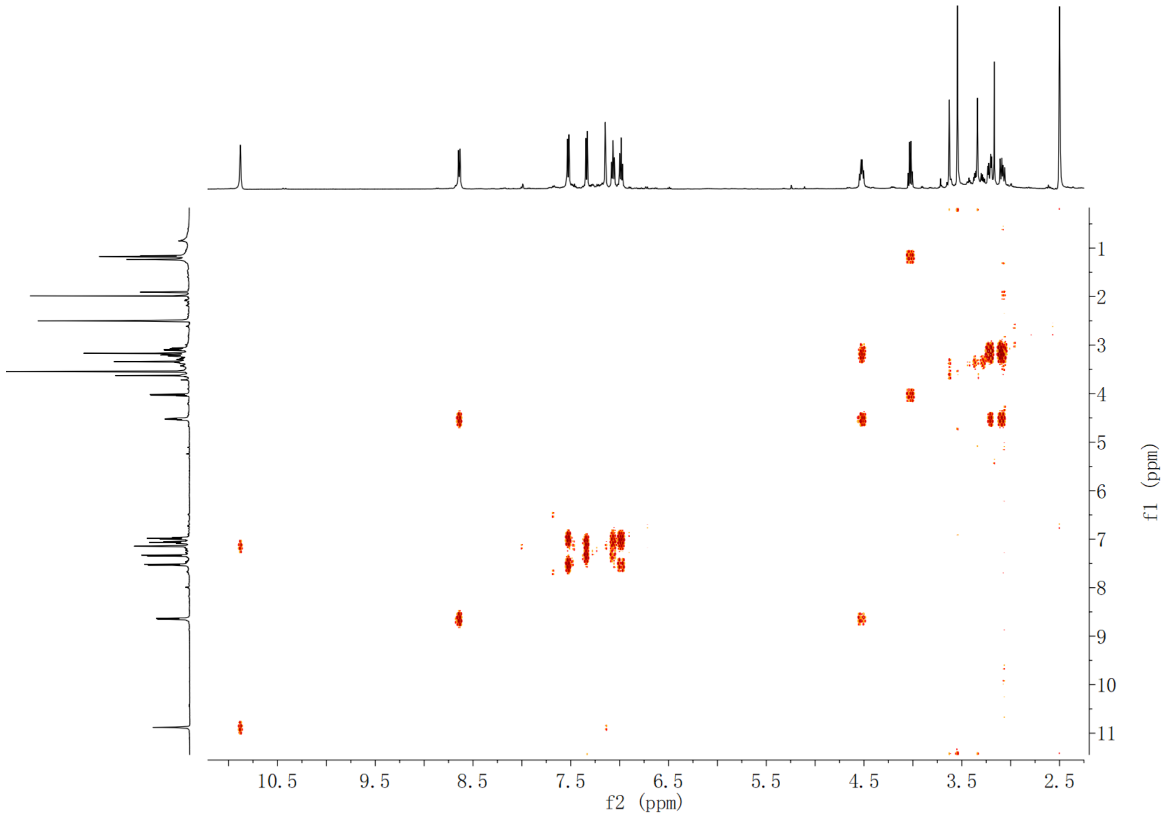

**Supplementary Fig. 123.** $^{1}H$-$^{1}H$ **COSY spectrum of compound (2*S*,3*S*)-*t*-ES-Trp in DMSO-$d_6$**

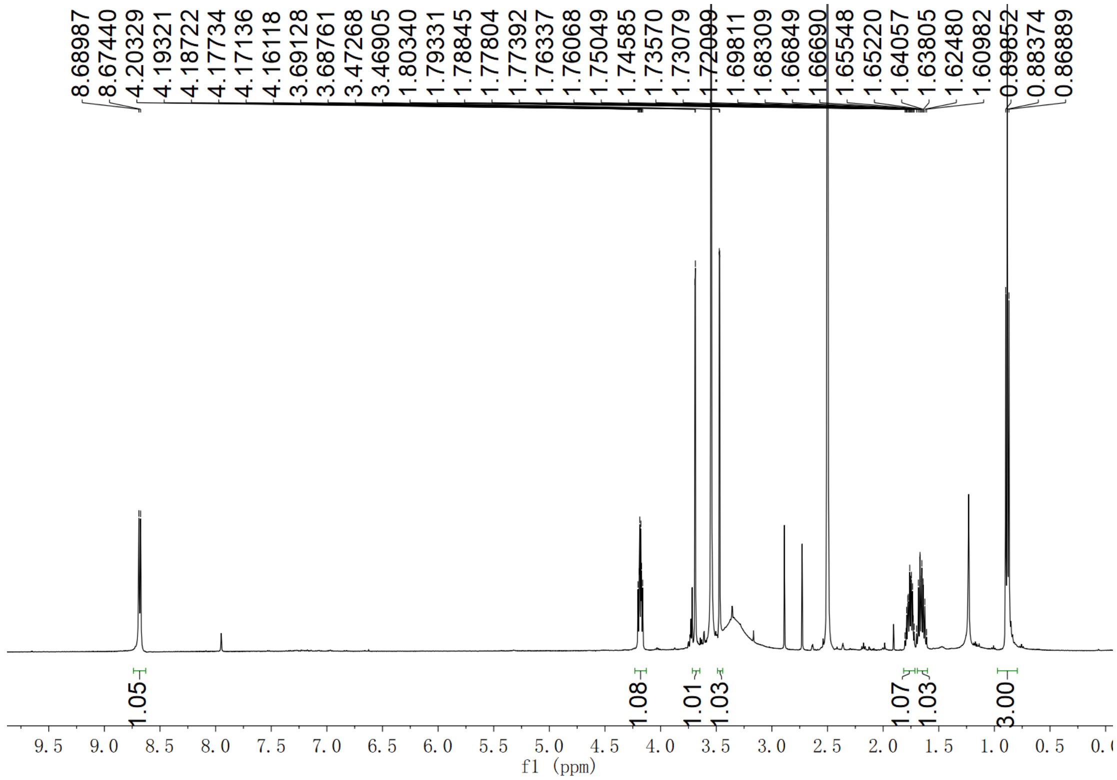

**Supplementary Fig. 124.** $^{1}H$ **NMR spectrum of compound (2*S*,3*S*)-*t*-ES-a1 in DMSO-$d_6$**

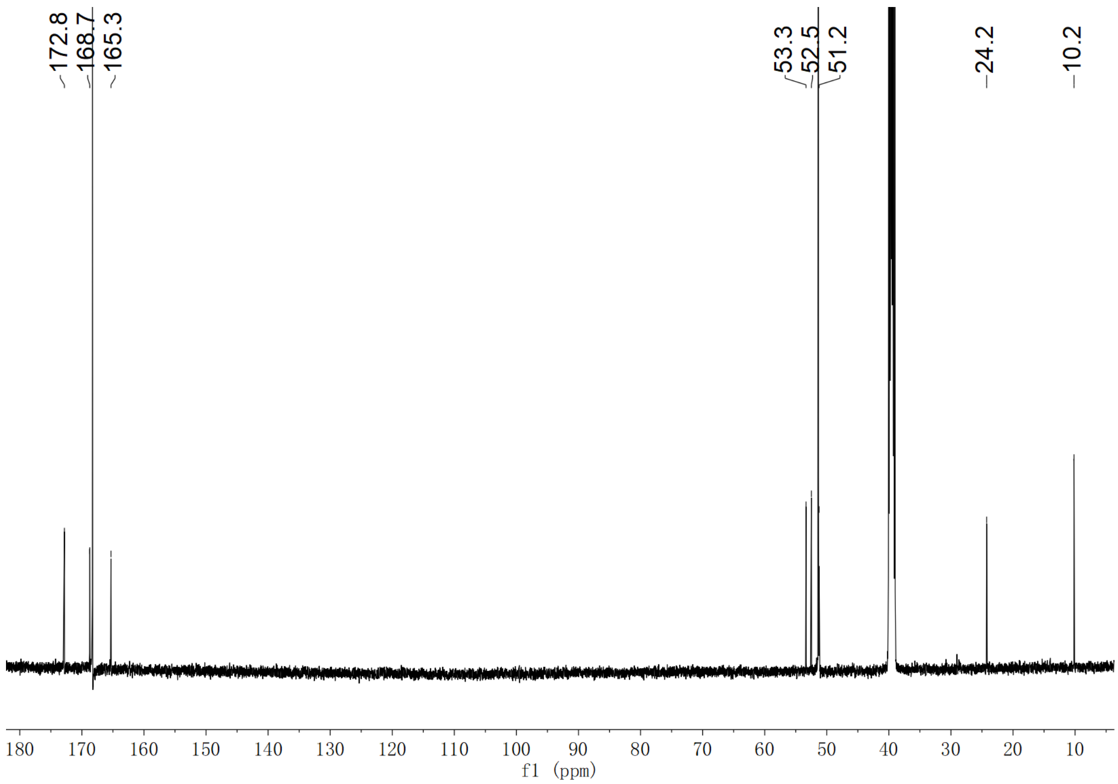

**Supplementary Fig. 125. $^{13}$C NMR spectrum of compound (2*S*,3*S*)-*t*-ES-a1 in DMSO-*d*$_6$**

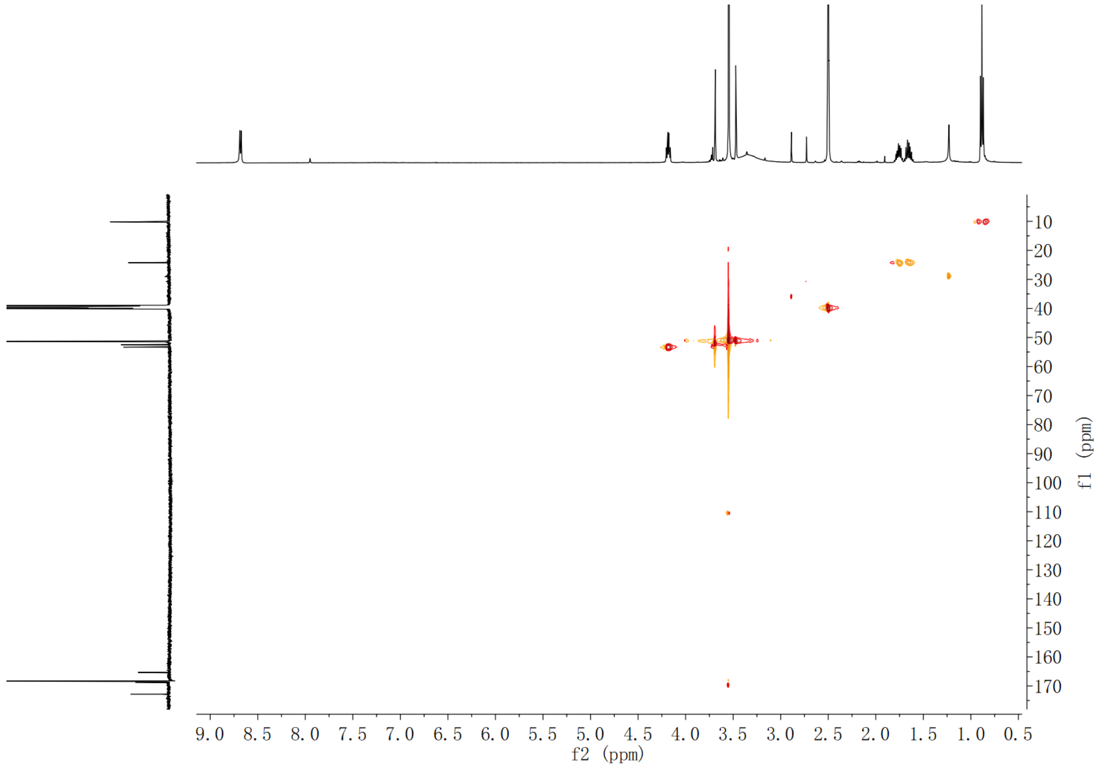

**Supplementary Fig. 126. HSQC spectrum of compound (2*S*,3*S*)-*t*-ES-a1 in DMSO-*d*$_6$**

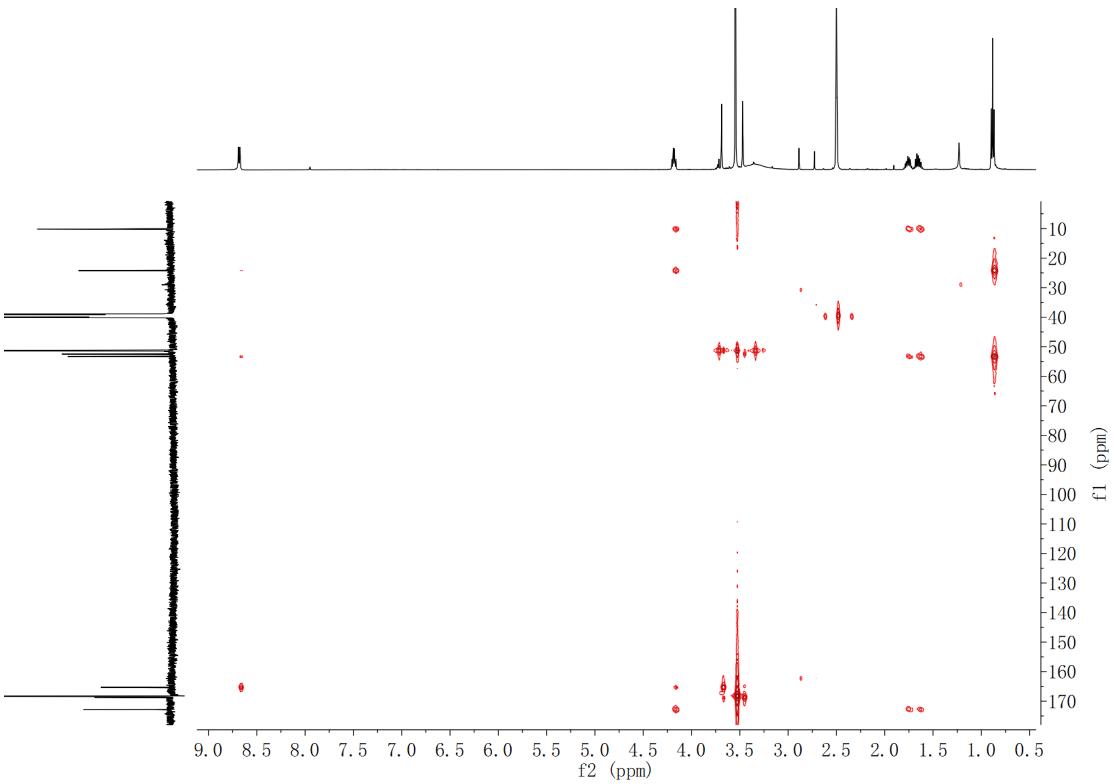

**Supplementary Fig. 127. HMBC spectrum of compound (2*S*,3*S*)-*t*-ES-a1 in DMSO-$d_6$**

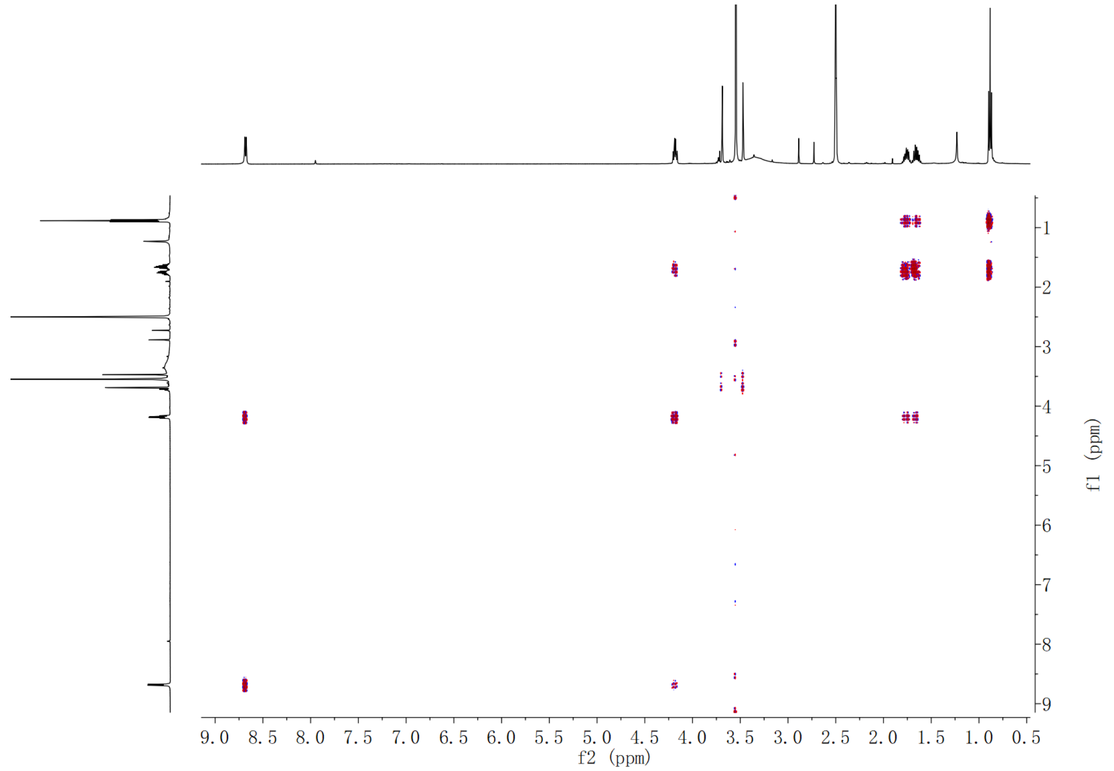

**Supplementary Fig. 128. $^{1}$H-$^{1}$H COSY spectrum of compound (2*S*,3*S*)-*t*-ES-a1 in DMSO-$d_6$**

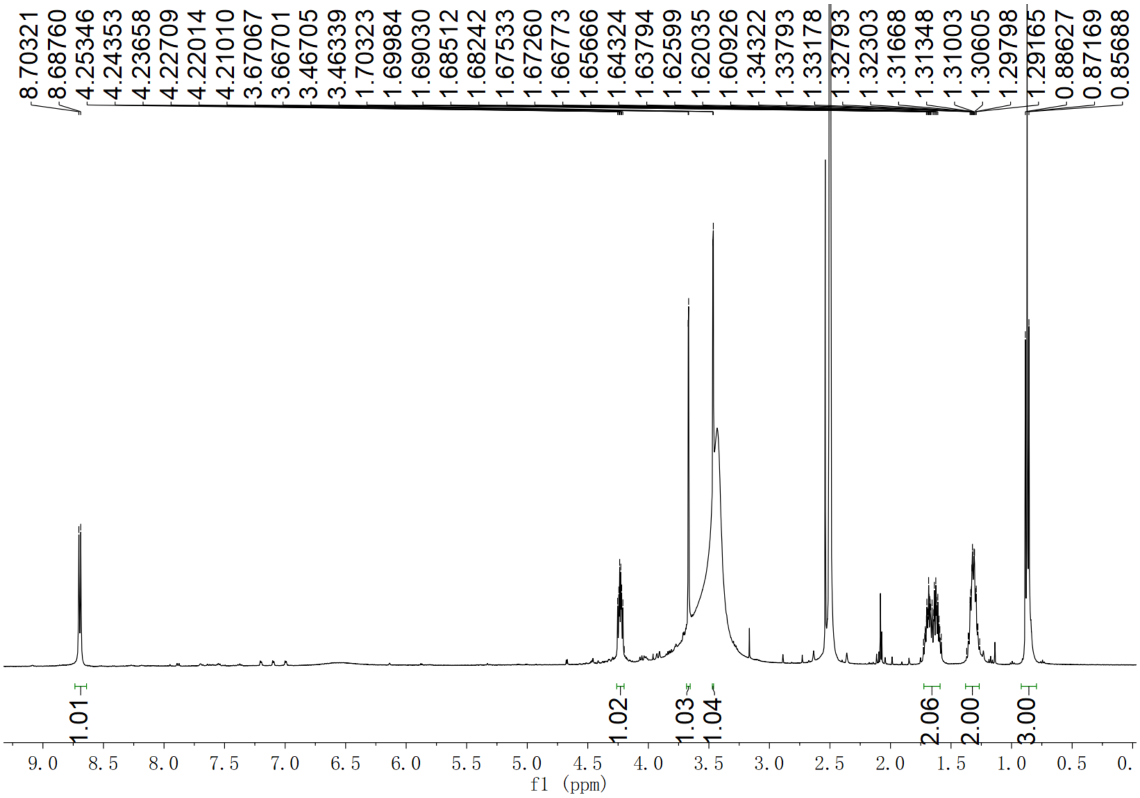

**Supplementary Fig. 129.** $^{1}$**H NMR spectrum of compound (2*S*,3*S*)-*t*-ES-a2 in DMSO-*d*$_6$**

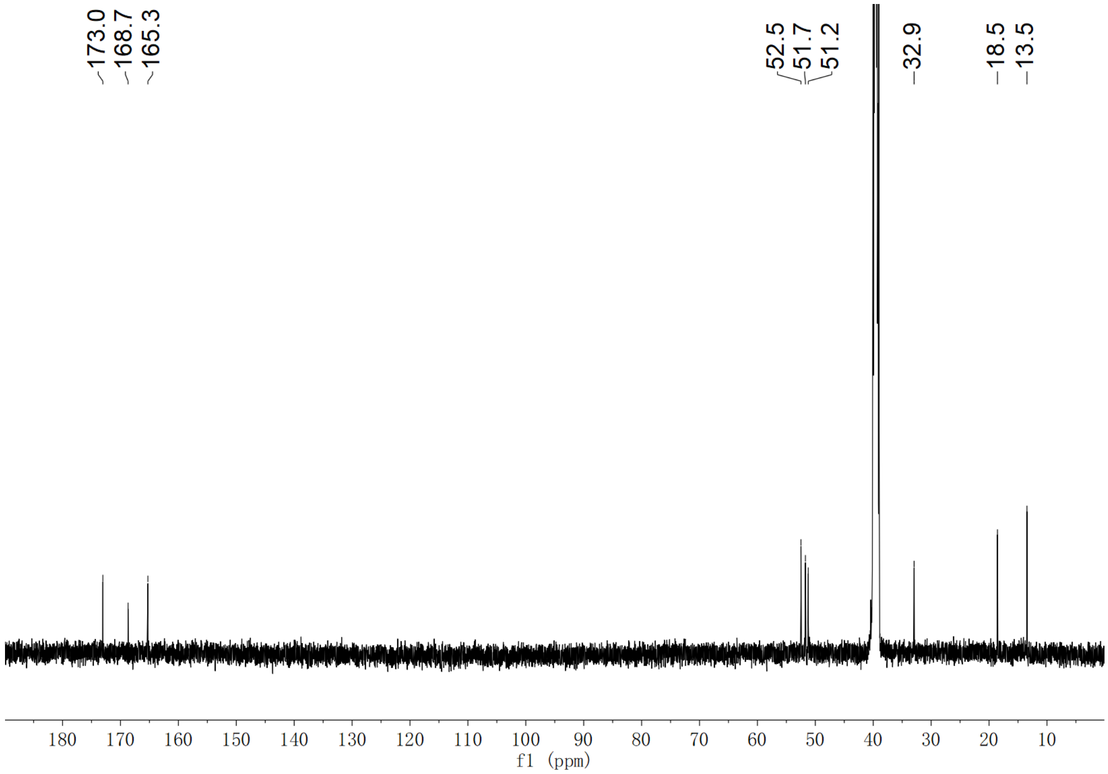

**Supplementary Fig. 130.** $^{13}$**C NMR spectrum of compound (2*S*,3*S*)-*t*-ES-a2 in DMSO-*d*$_6$**

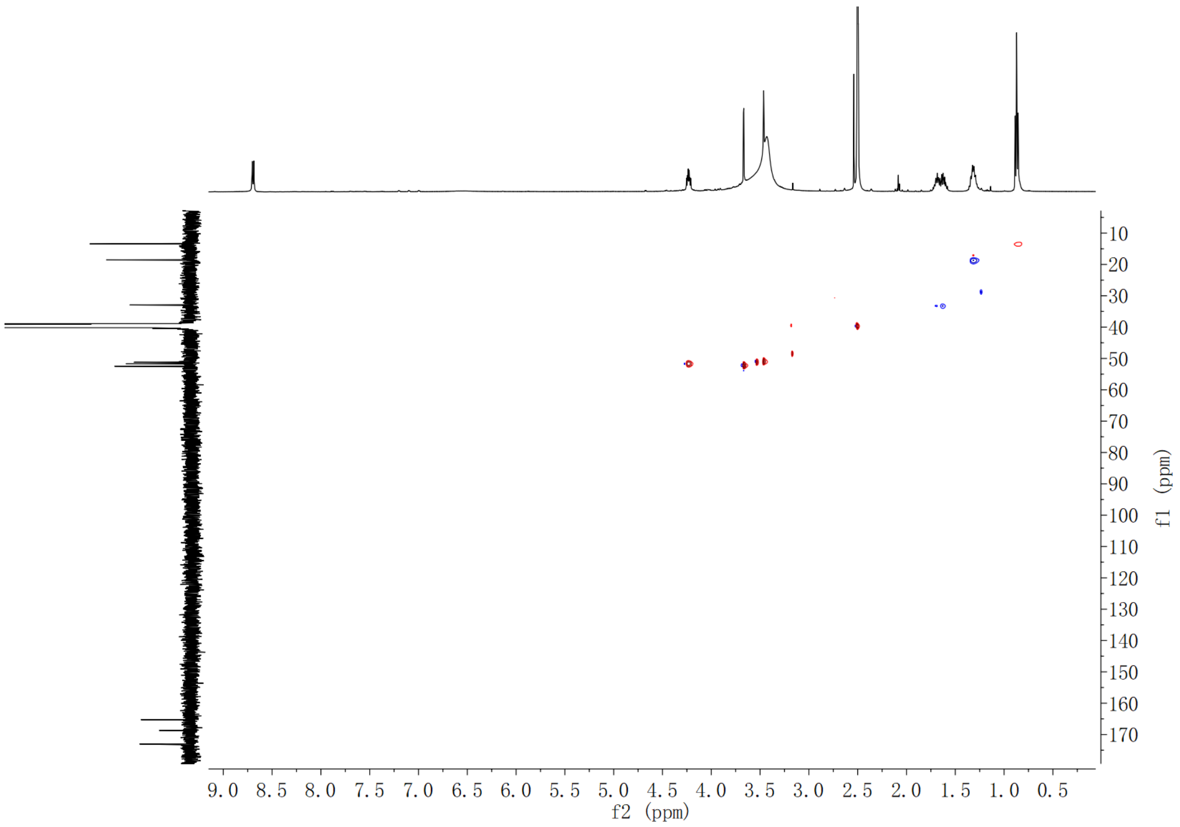

**Supplementary Fig. 131. HSQC spectrum of compound (2*S*,3*S*)-*t*-ES-a2 in DMSO-$d_6$**

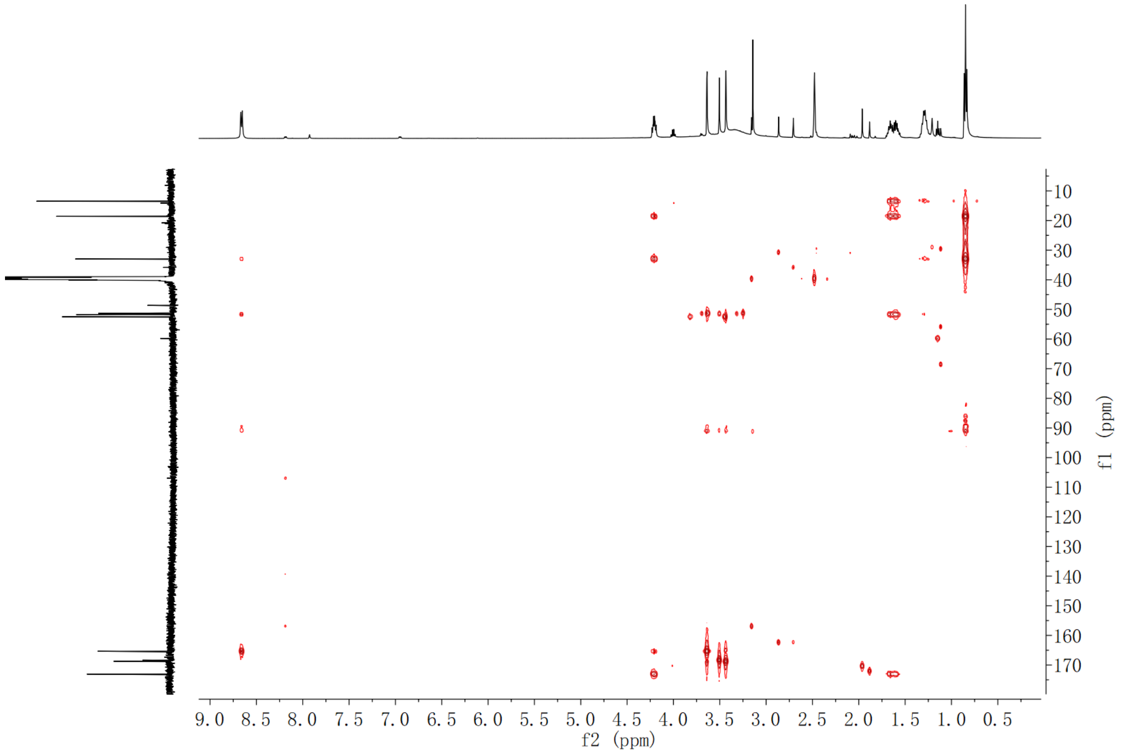

**Supplementary Fig. 132. HMBC spectrum of compound (2*S*,3*S*)-*t*-ES-a2 in DMSO-$d_6$**

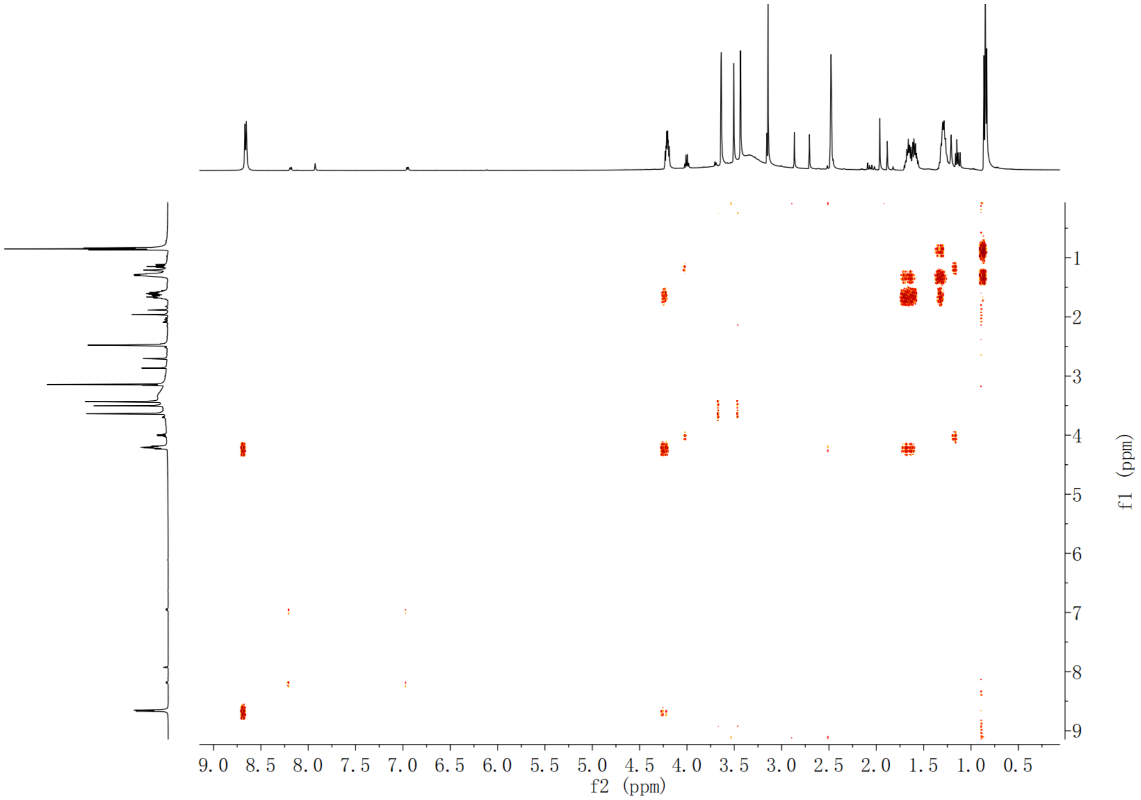

Supplementary Fig. 133. $^1$H-$^1$H COSY spectrum of compound (2*S*,3*S*)-*t*-ES-a2 in DMSO-$d_6$

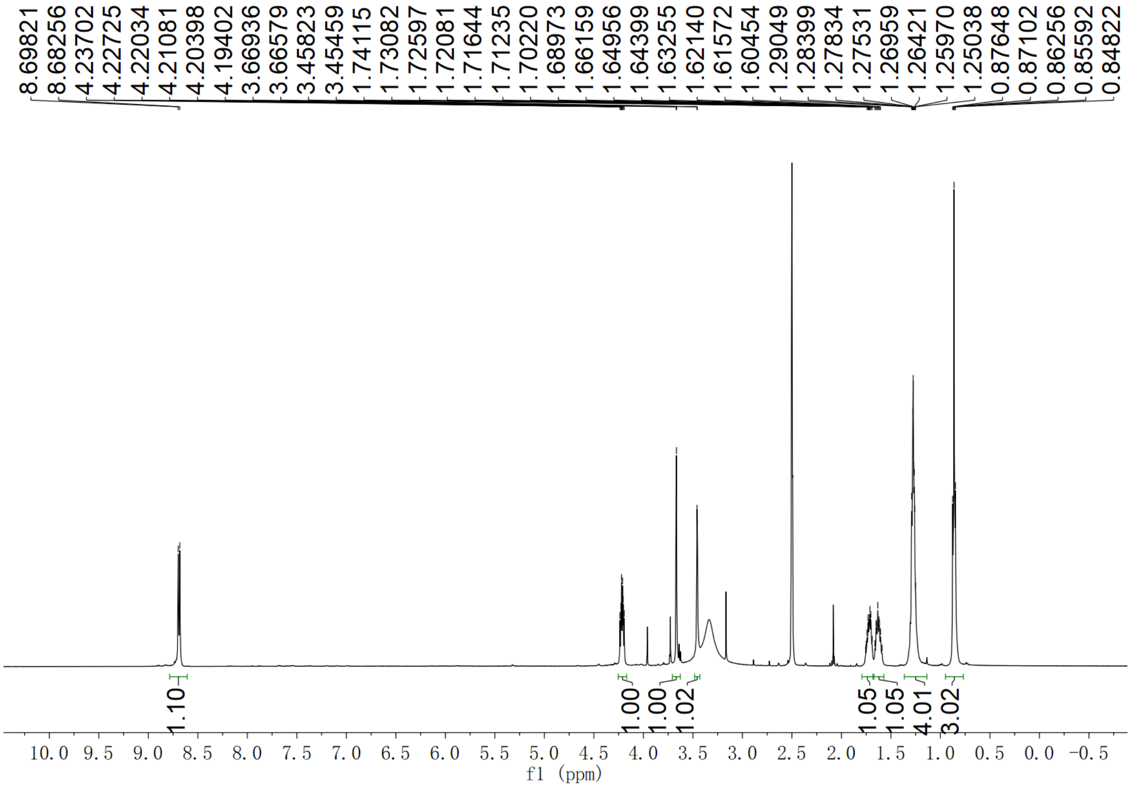

Supplementary Fig. 134. $^1$H NMR spectrum of compound (2*S*,3*S*)-*t*-ES-a3 in DMSO-$d_6$

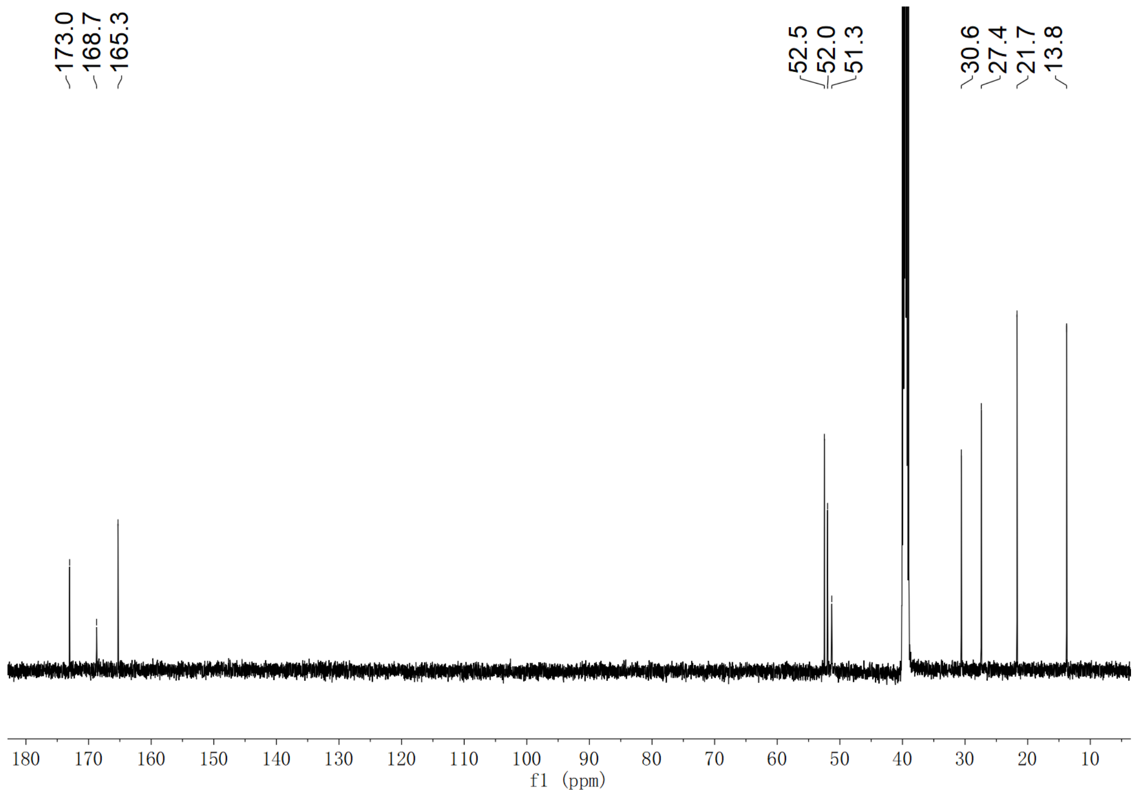

**Supplementary Fig. 135. $^{13}$C NMR spectrum of compound (2*S*,3*S*)-*t*-ES-a3 in DMSO-$d_6$**

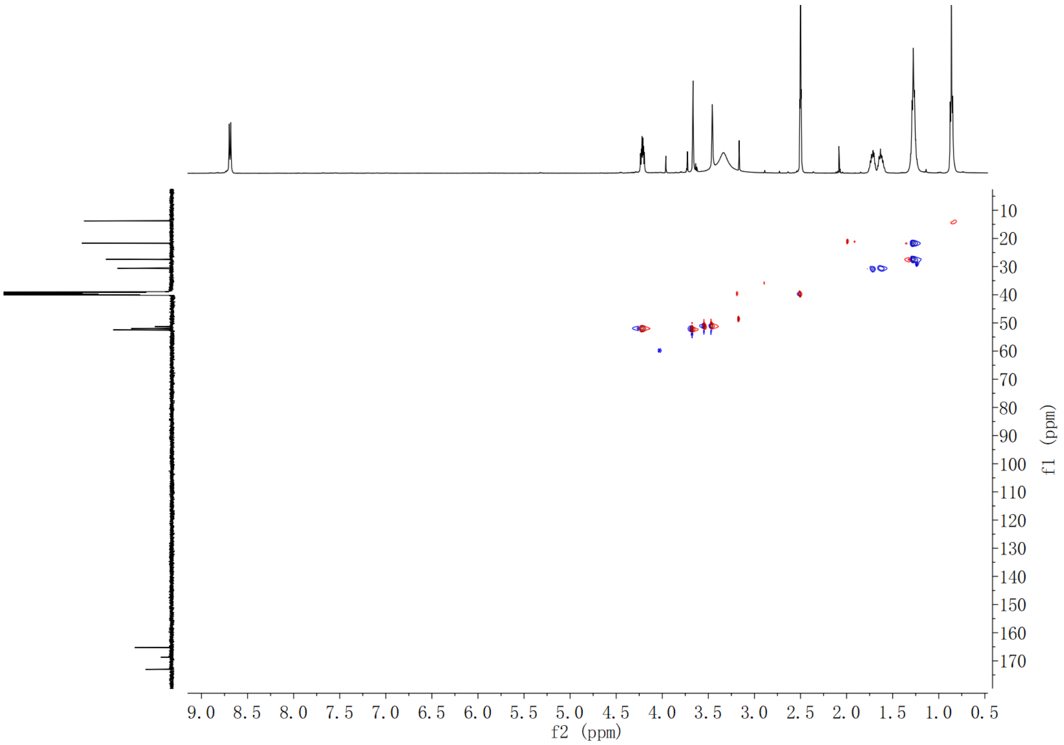

**Supplementary Fig. 136. HSQC spectrum of compound (2*S*,3*S*)-*t*-ES-a3 in DMSO-$d_6$**

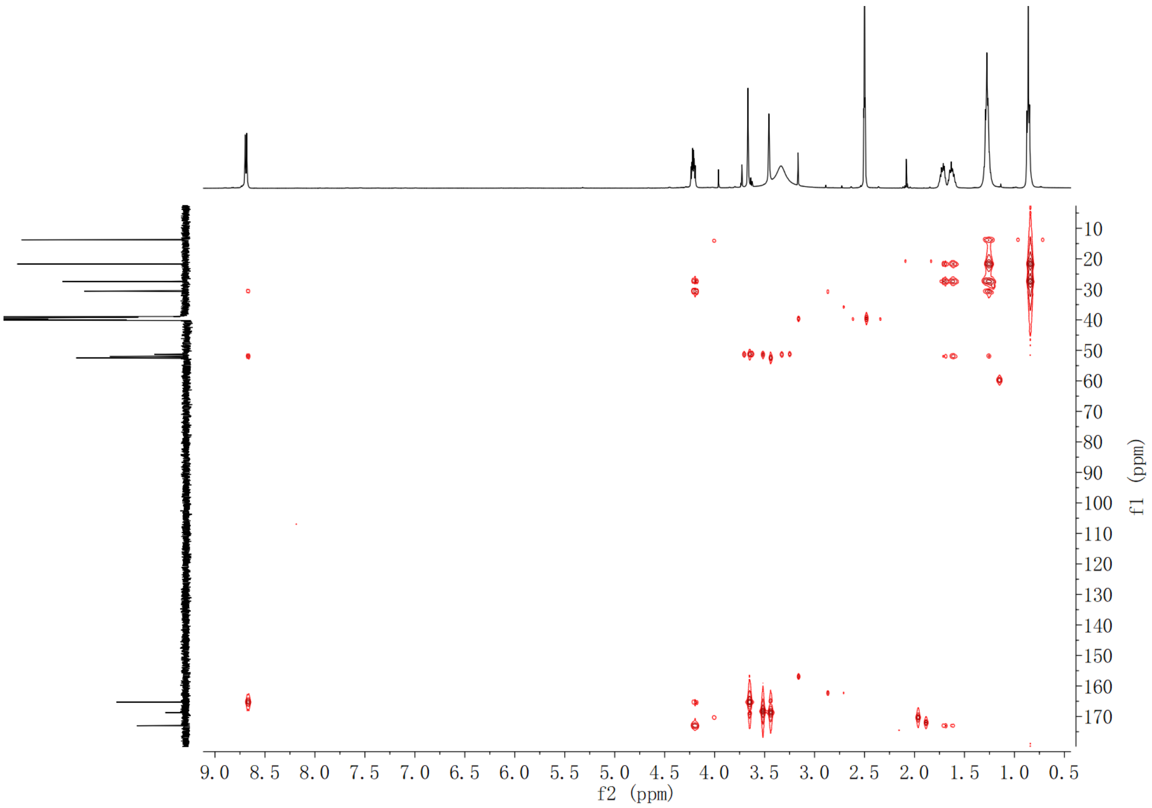

**Supplementary Fig. 137. HMBC spectrum of compound (2*S*,3*S*)-*t*-ES-a3 in DMSO-$d_6$**

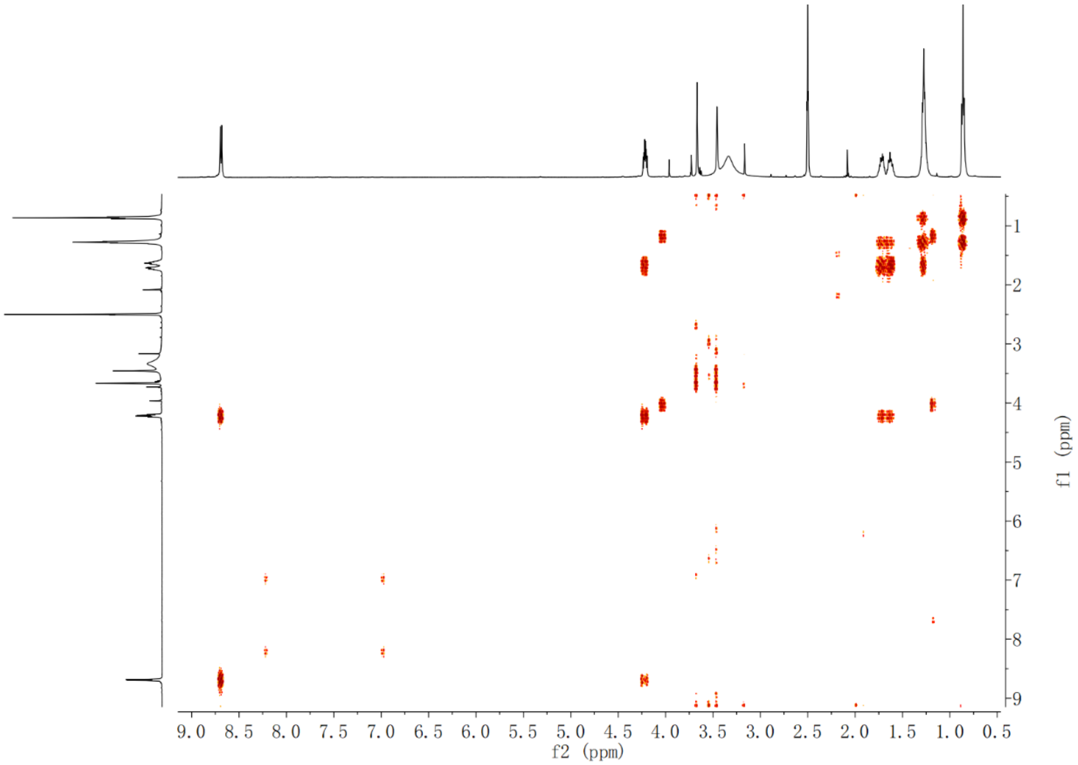

**Supplementary Fig. 138. $^1$H-$^1$H COSY spectrum of compound (2*S*,3*S*)-*t*-ES-a3 in DMSO-$d_6$**

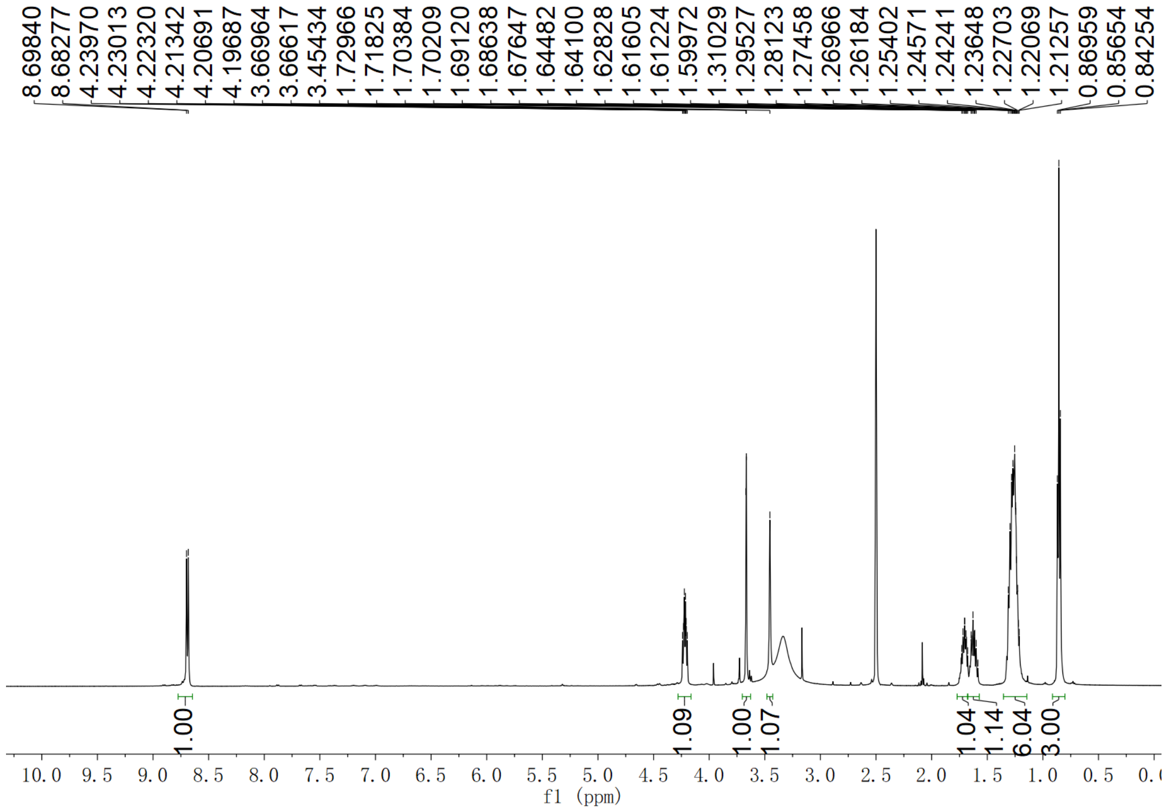

**Supplementary Fig. 139. $^{1}$H NMR spectrum of compound (2*S*,3*S*)-*t*-ES-a4 in DMSO-$d_6$**

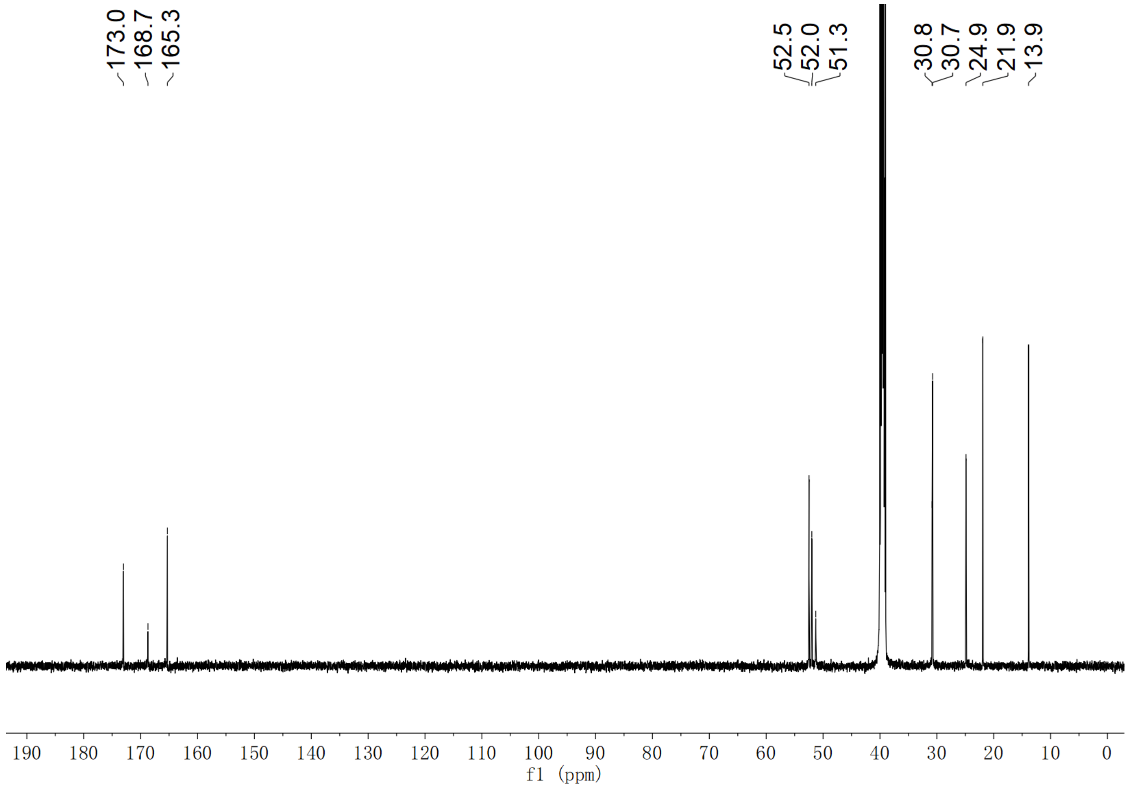

**Supplementary Fig. 140. $^{13}$C NMR spectrum of compound (2*S*,3*S*)-*t*-ES-a4 in DMSO-$d_6$**

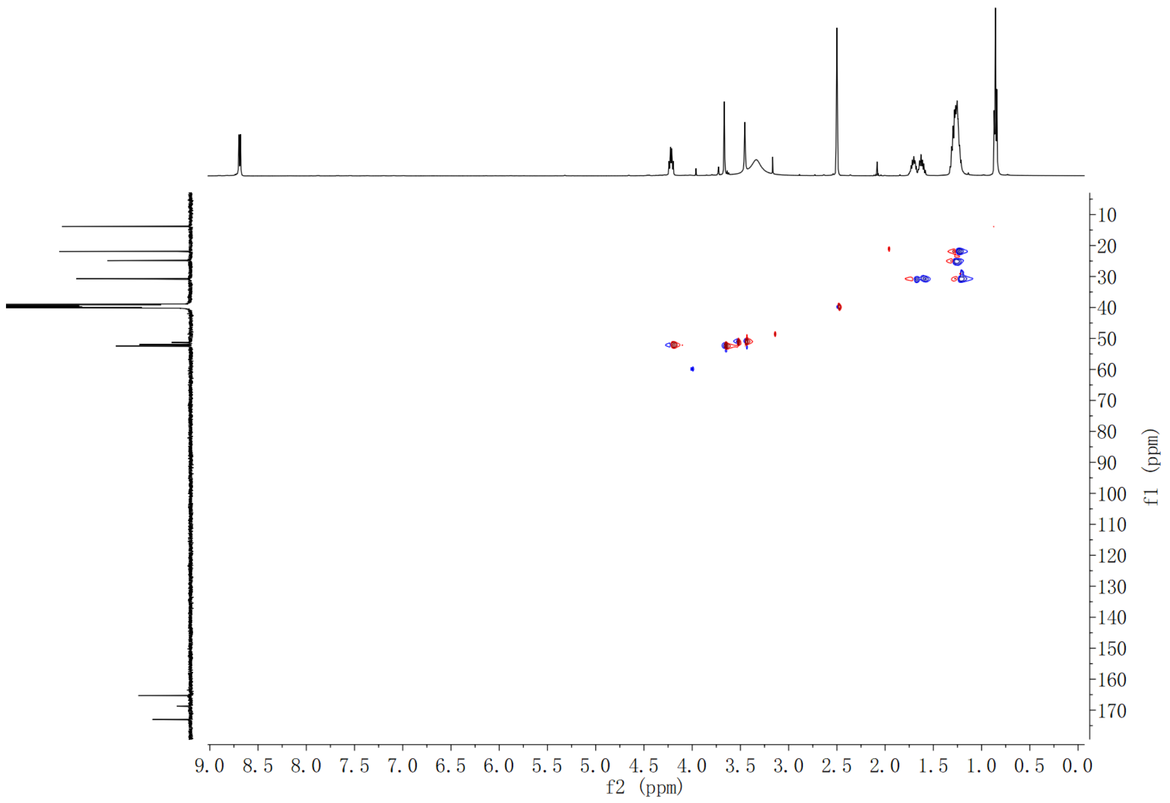

**Supplementary Fig. 141. HSQC spectrum of compound (2*S*,3*S*)-*t*-ES-a4 in DMSO-*d*$_6$**

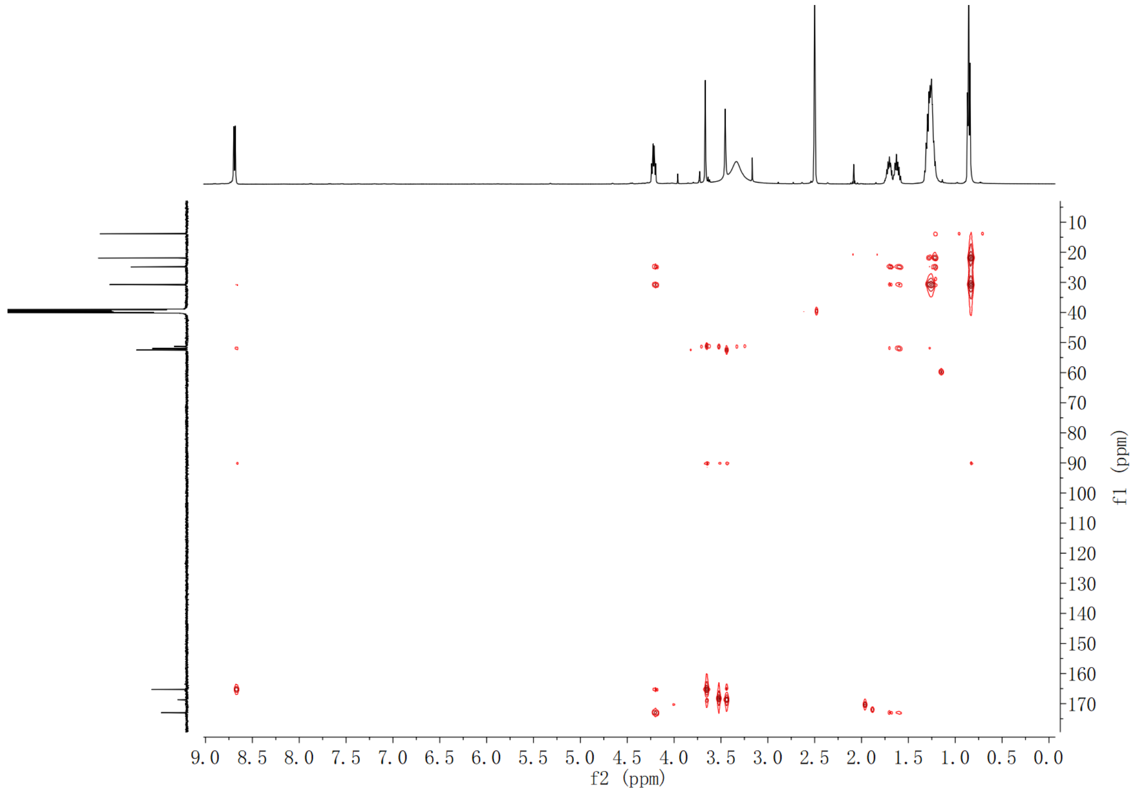

**Supplementary Fig. 142. HMBC spectrum of compound (2*S*,3*S*)-*t*-ES-a4 in DMSO-*d*$_6$**

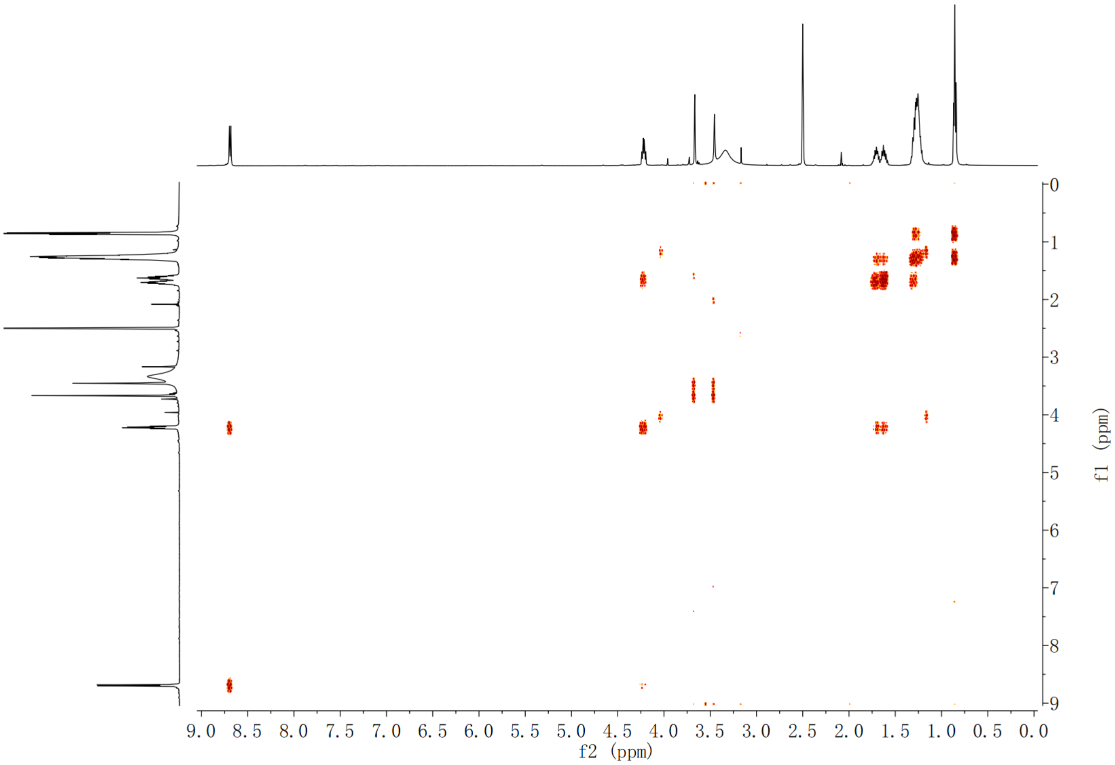

**Supplementary Fig. 143. $^{1}$H-$^{1}$H COSY spectrum of compound (2*S*,3*S*)-*t*-ES-a4 in DMSO-$d_6$**

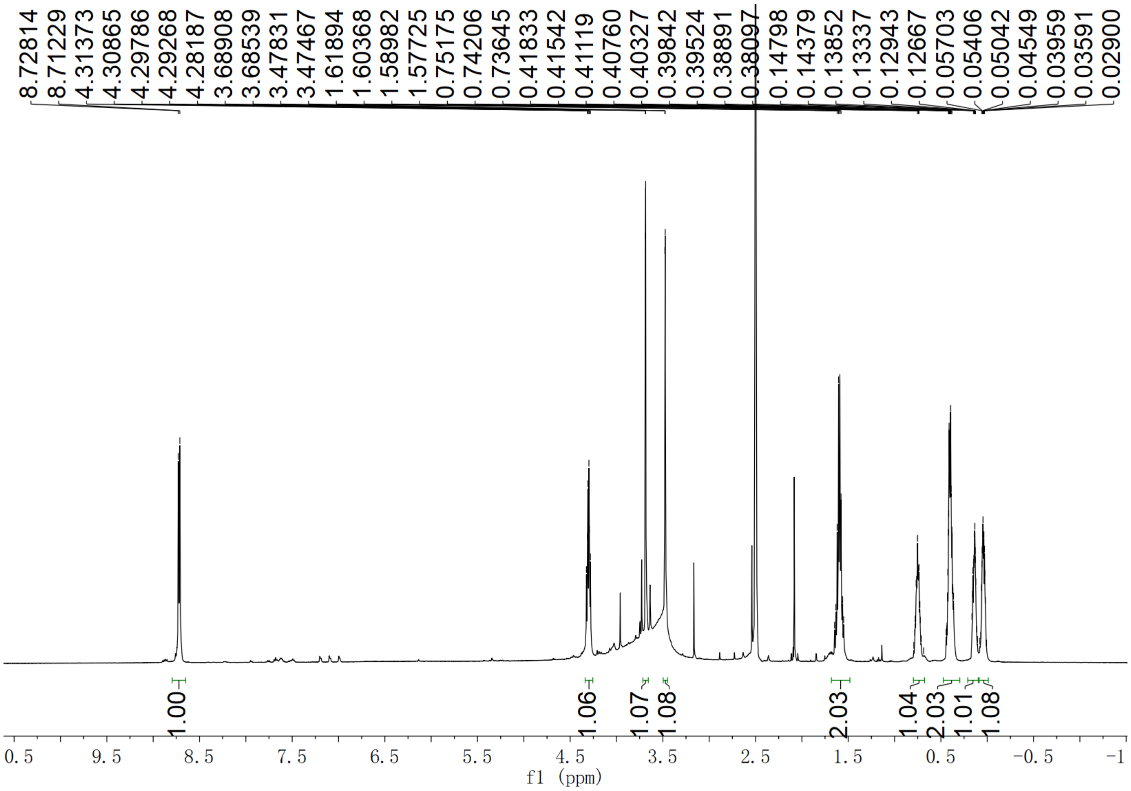

**Supplementary Fig. 144. $^{1}$H NMR spectrum of compound (2*S*,3*S*)-*t*-ES-a5 in DMSO-$d_6$**

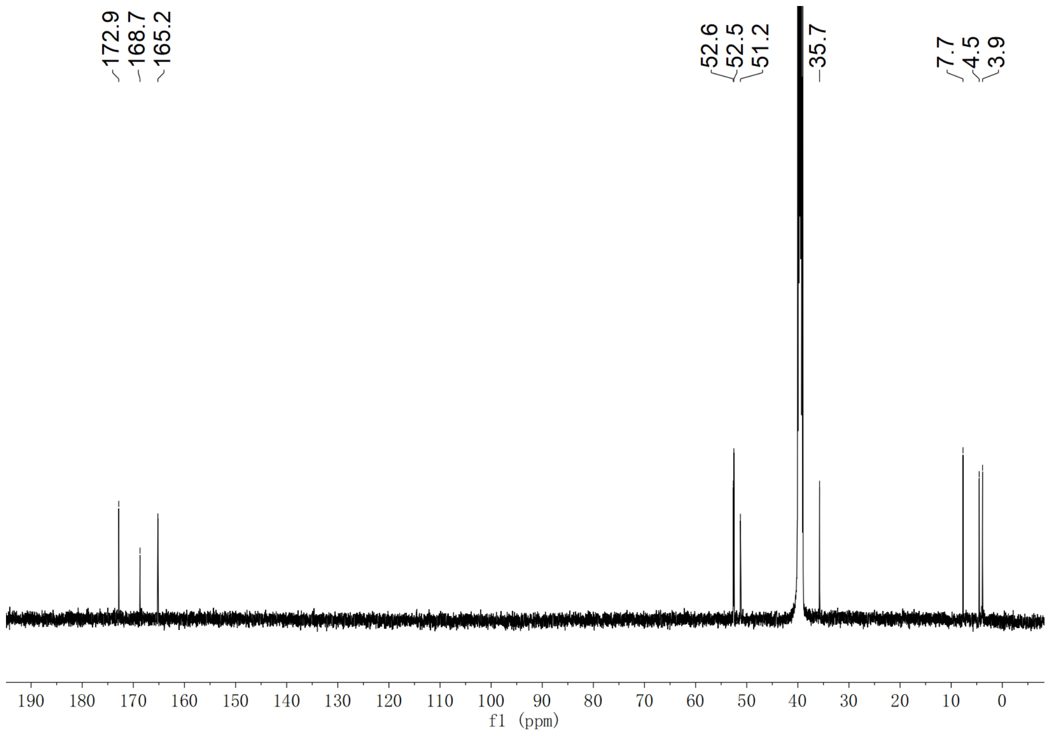

**Supplementary Fig. 145. $^{13}$C NMR spectrum of compound (2*S*,3*S*)-*t*-ES-a5 in DMSO-$d_6$**

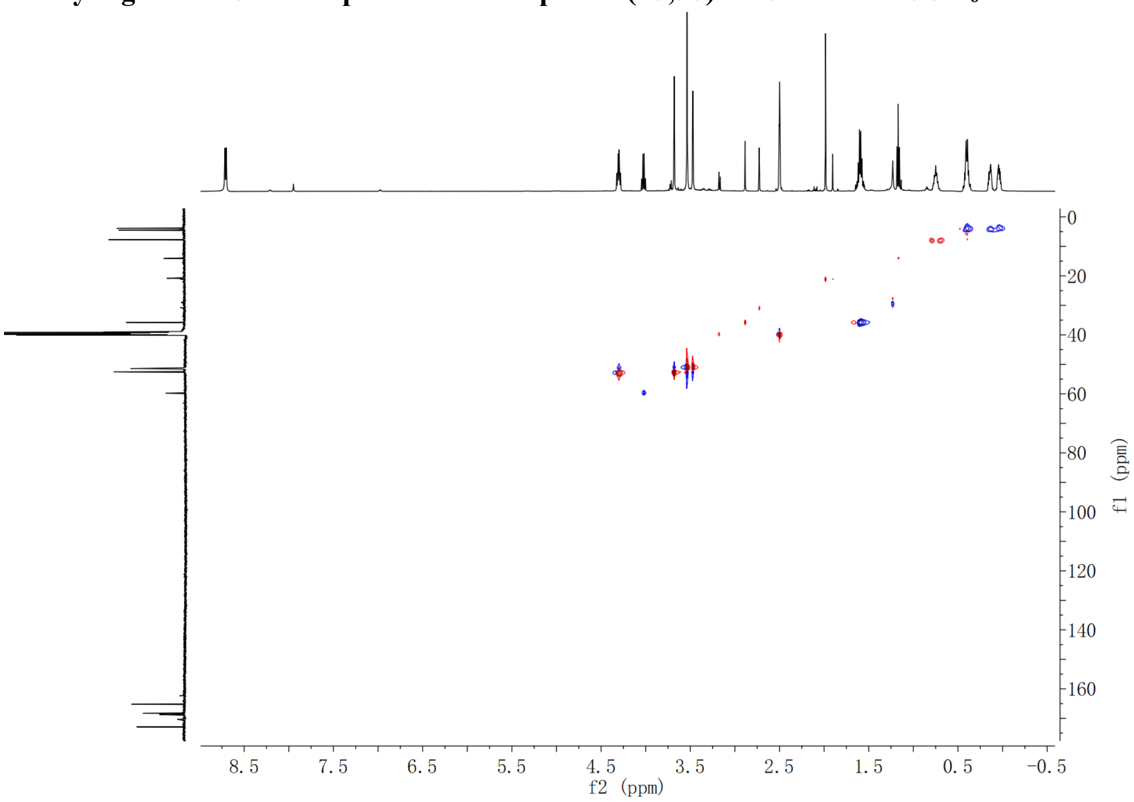

**Supplementary Fig. 146. HSQC spectrum of compound (2*S*,3*S*)-*t*-ES-a5 in DMSO-$d_6$**

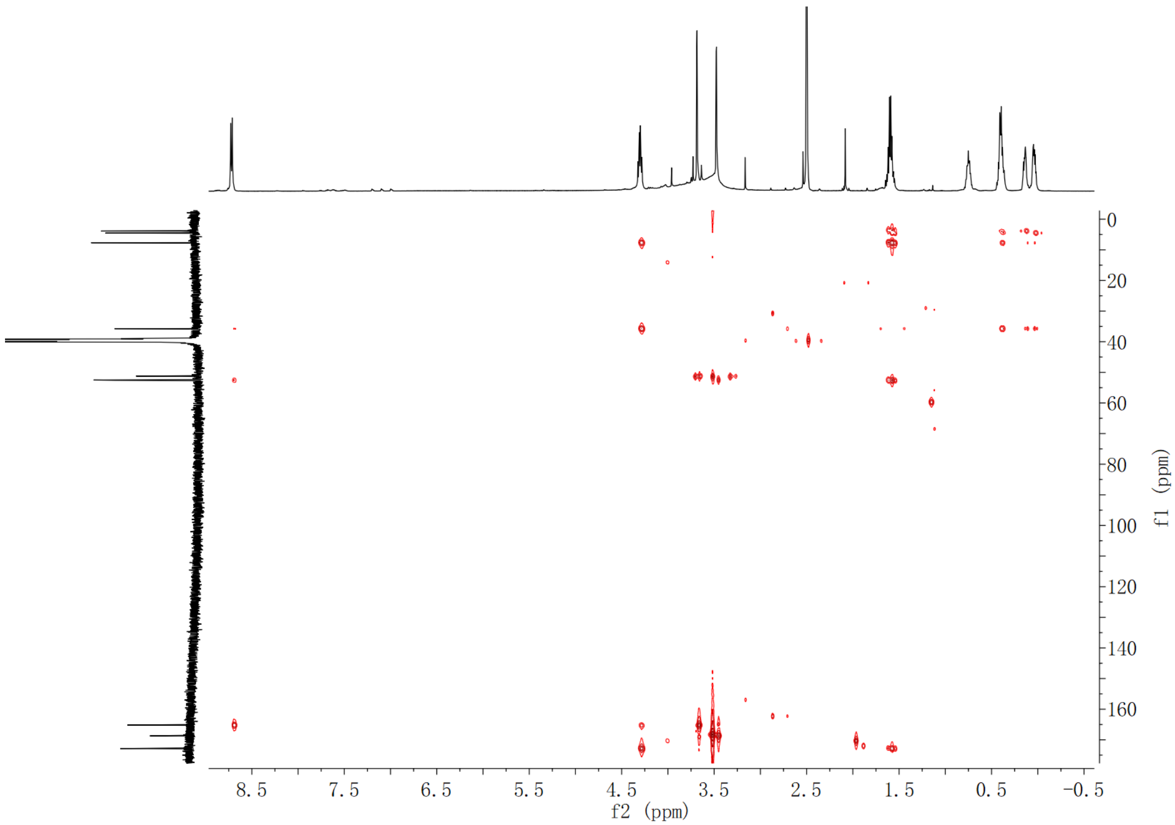

**Supplementary Fig. 147. HMBC spectrum of compound (2*S*,3*S*)-*t*-ES-a5 in DMSO-$d_6$**

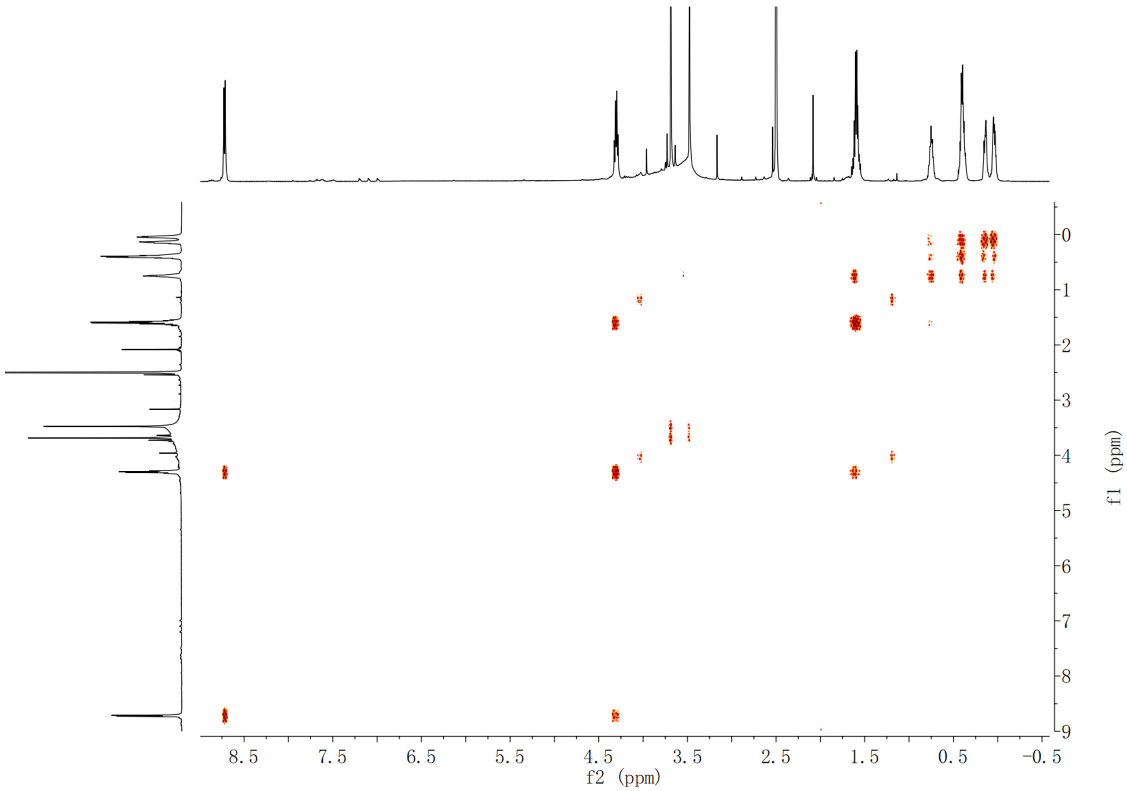

**Supplementary Fig. 148. $^{1}$H-$^{1}$H COSY spectrum of compound (2*S*,3*S*)-*t*-ES-a5 in DMSO-$d_6$**

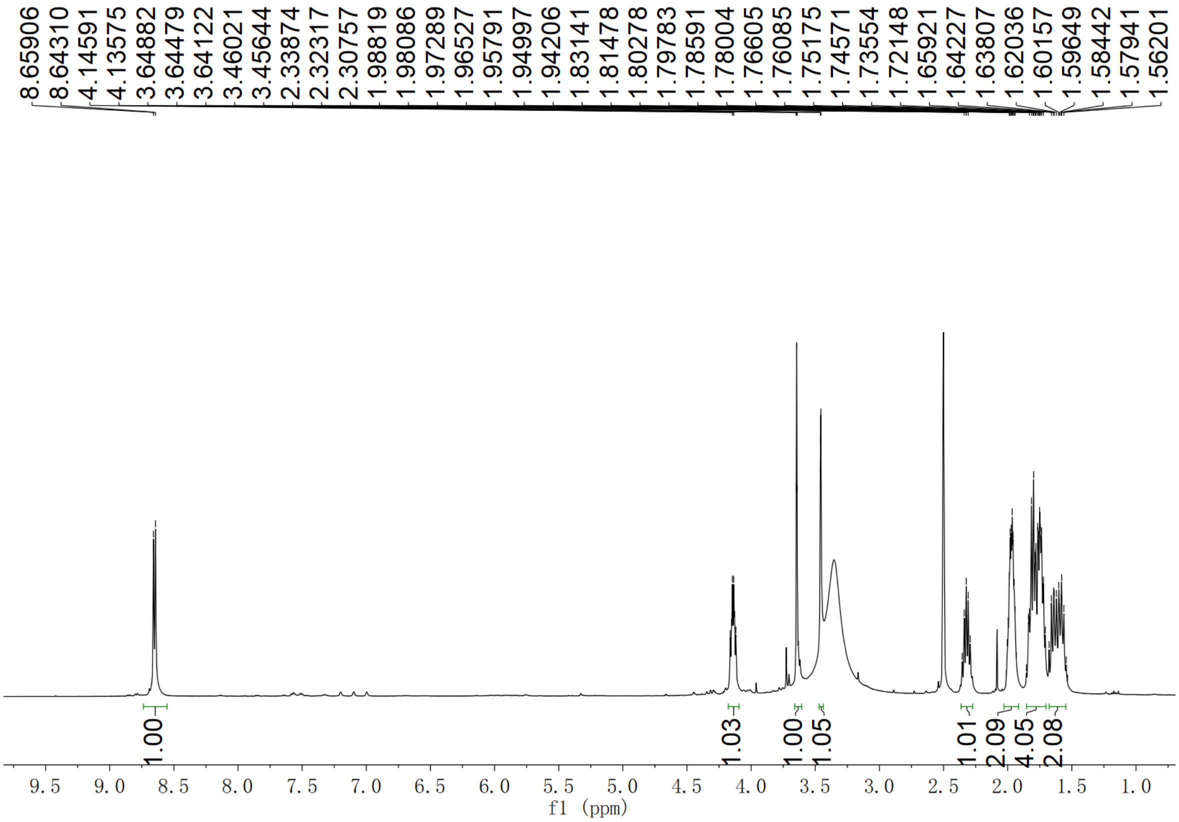

Supplementary Fig. 149. $^{1}$H NMR spectrum of compound (2*S*,3*S*)-*t*-ES-a6 in DMSO-$d_6$

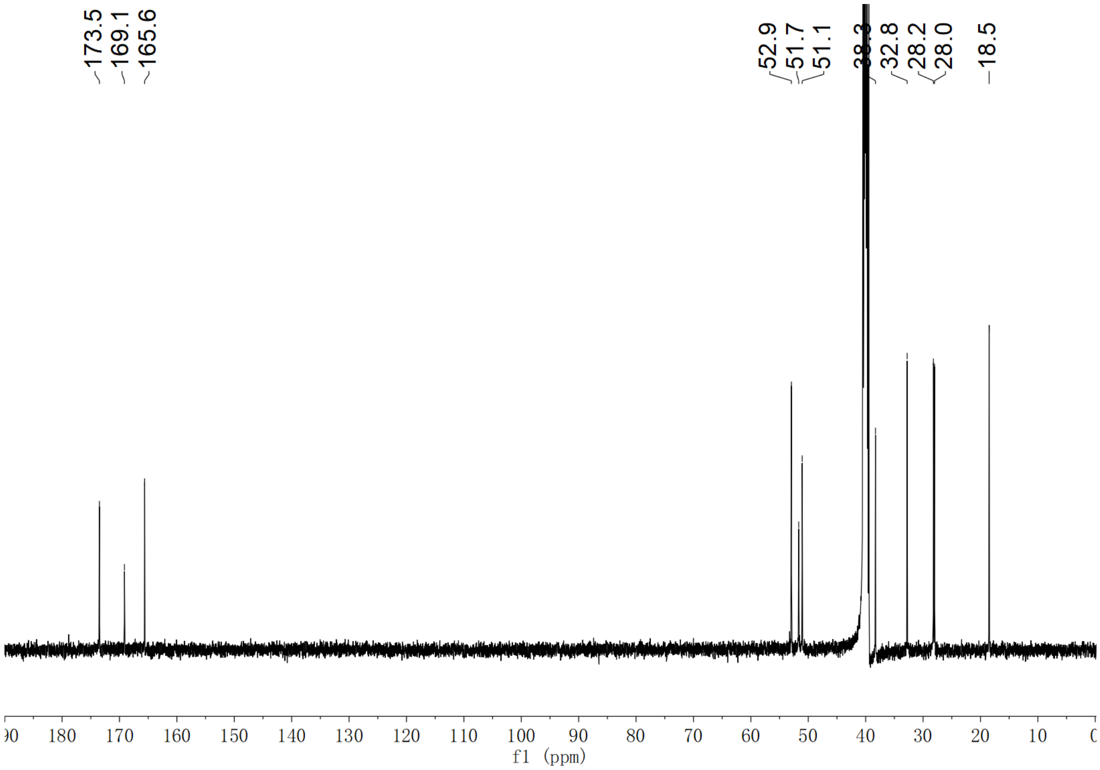

Supplementary Fig. 150. $^{13}$C NMR spectrum of compound (2*S*,3*S*)-*t*-ES-a6 in DMSO-$d_6$

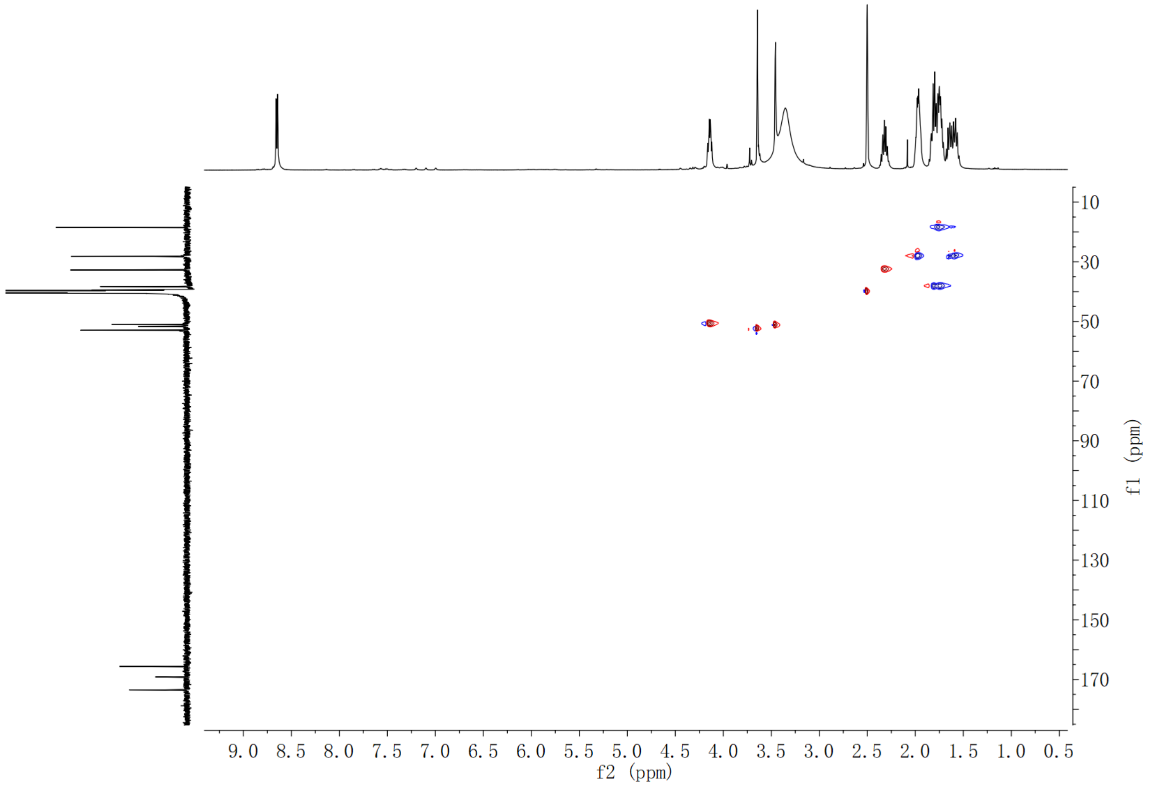

**Supplementary Fig. 151. HSQC spectrum of compound (2*S*,3*S*)-*t*-ES-a6 in DMSO-$d_6$**

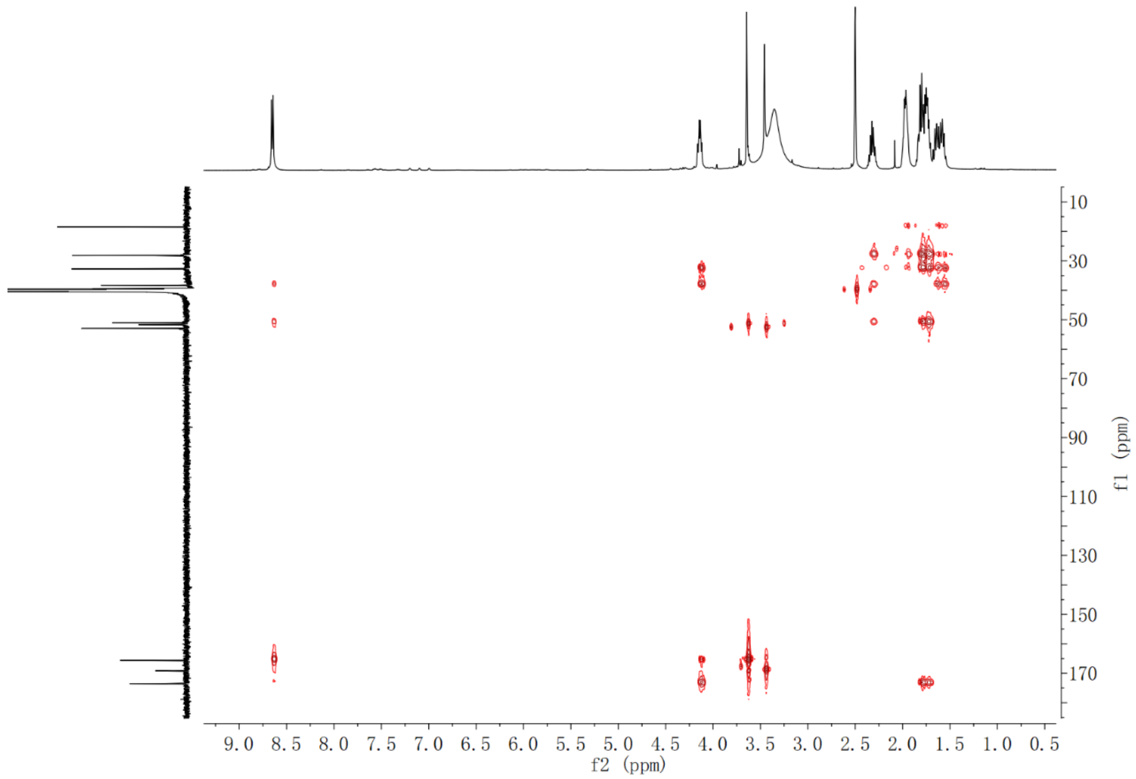

**Supplementary Fig. 152. HMBC spectrum of compound (2*S*,3*S*)-*t*-ES-a6 in DMSO-$d_6$**

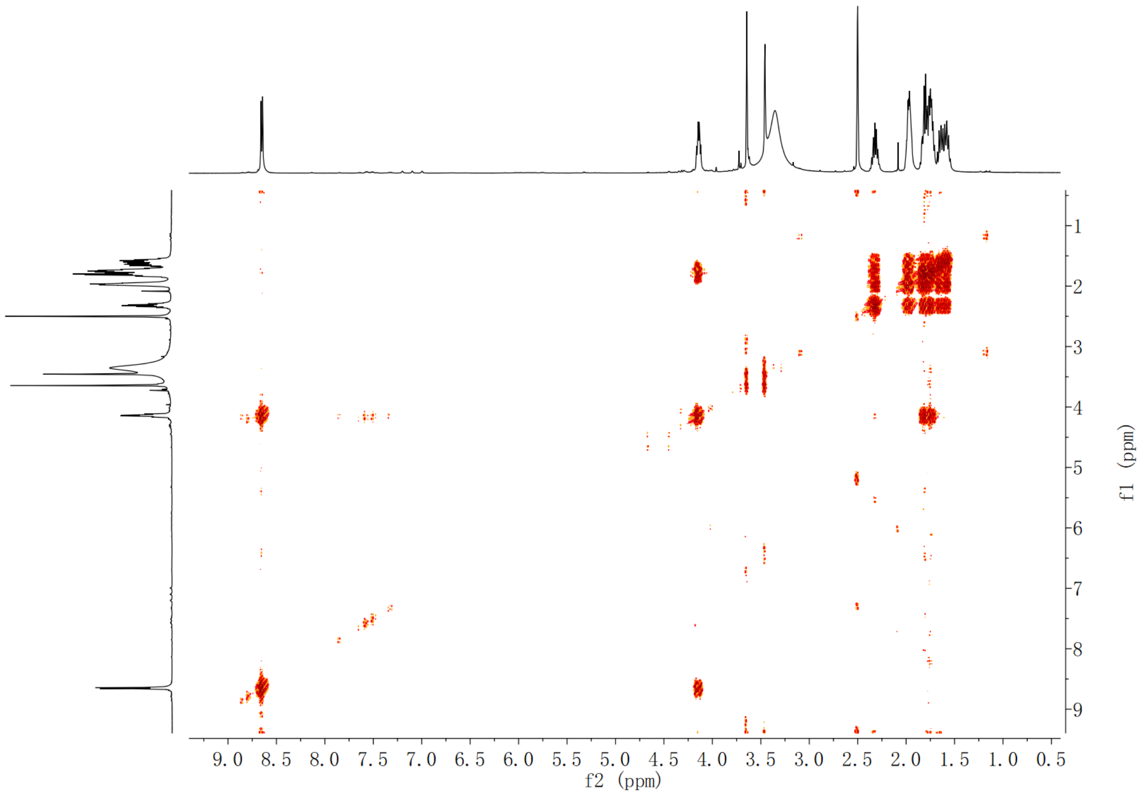

**Supplementary Fig. 153.** $^{1}$**H-**$^{1}$**H COSY spectrum of compound (2*S*,3*S*)-*t*-ES-a6 in DMSO-*d*$_6$**

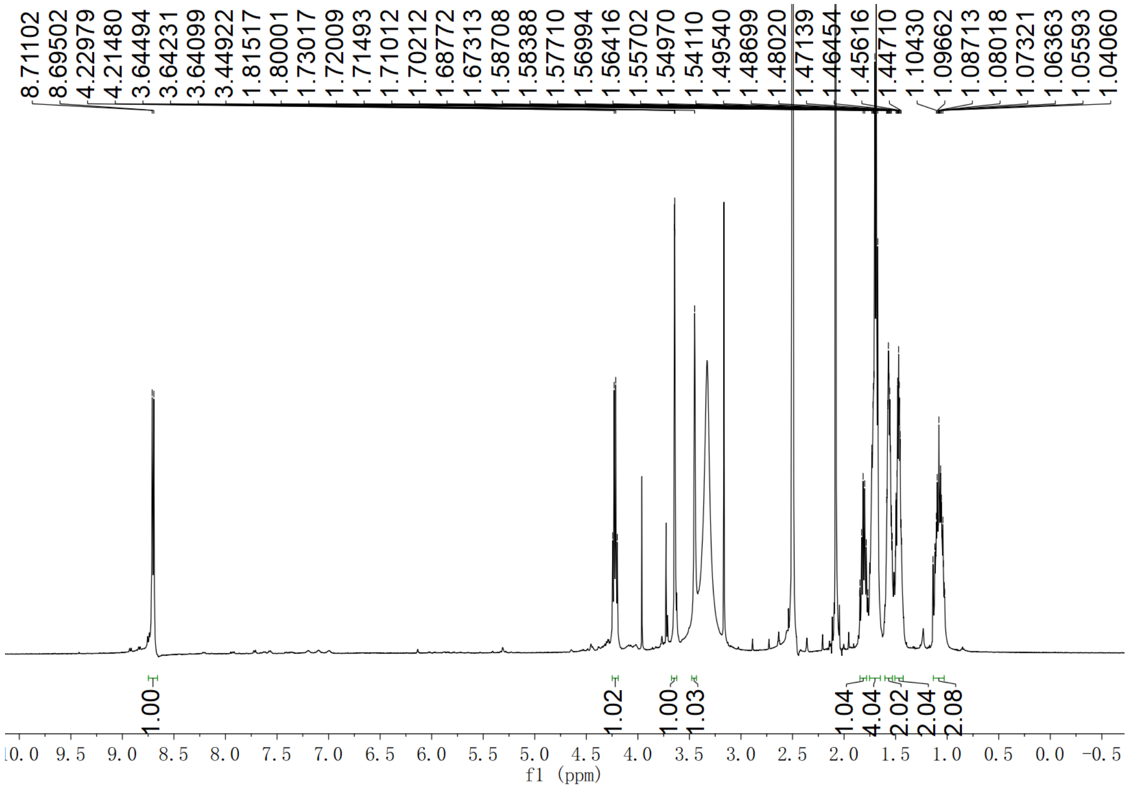

**Supplementary Fig. 154.** $^{1}$**H NMR spectrum of compound (2*S*,3*S*)-*t*-ES-a7 in DMSO-*d*$_6$**

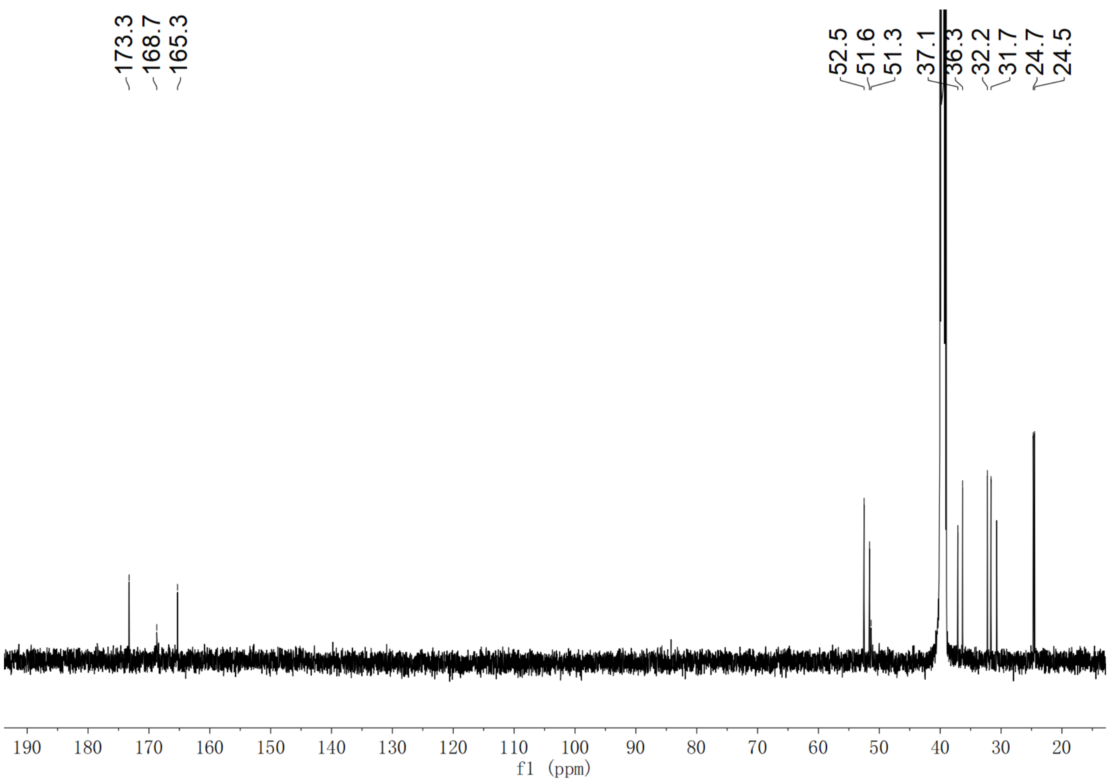

**Supplementary Fig. 155. $^{13}$C NMR spectrum of compound (2*S*,3*S*)-*t*-ES-a7 in DMSO-*d*$_6$**

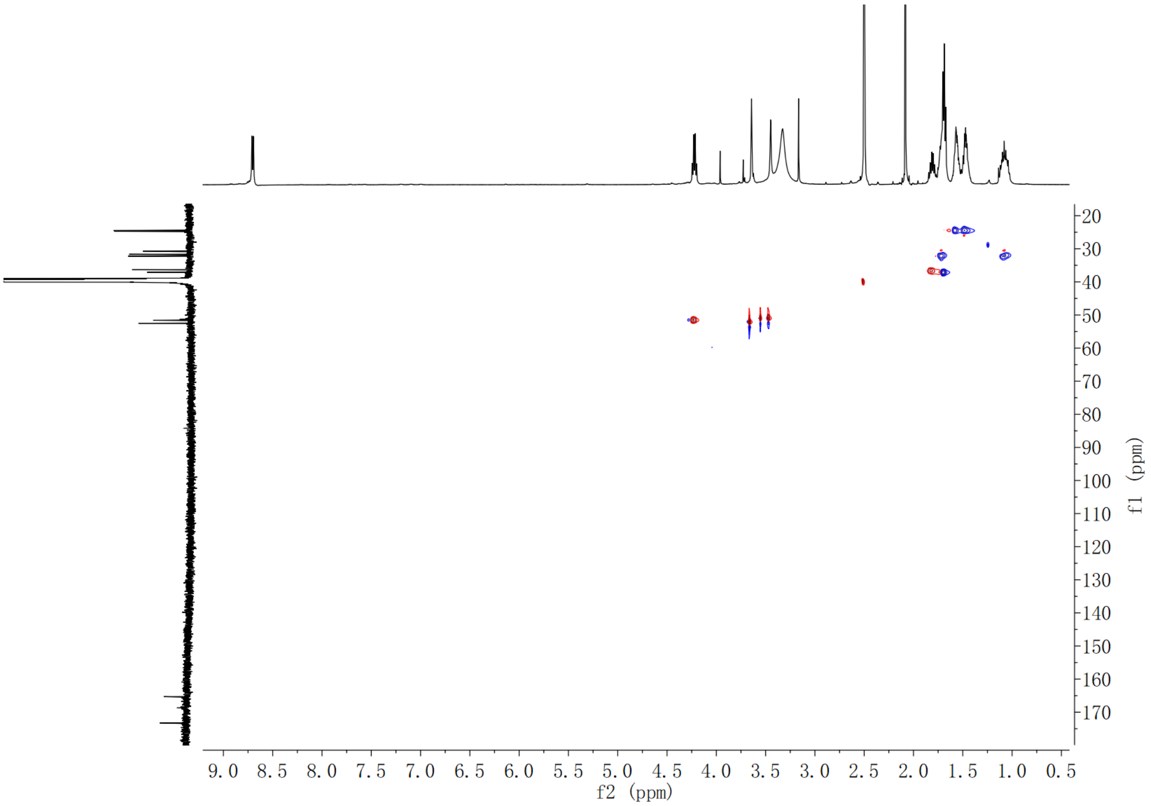

**Supplementary Fig. 156. HSQC spectrum of compound (2*S*,3*S*)-*t*-ES-a7 in DMSO-*d*$_6$**

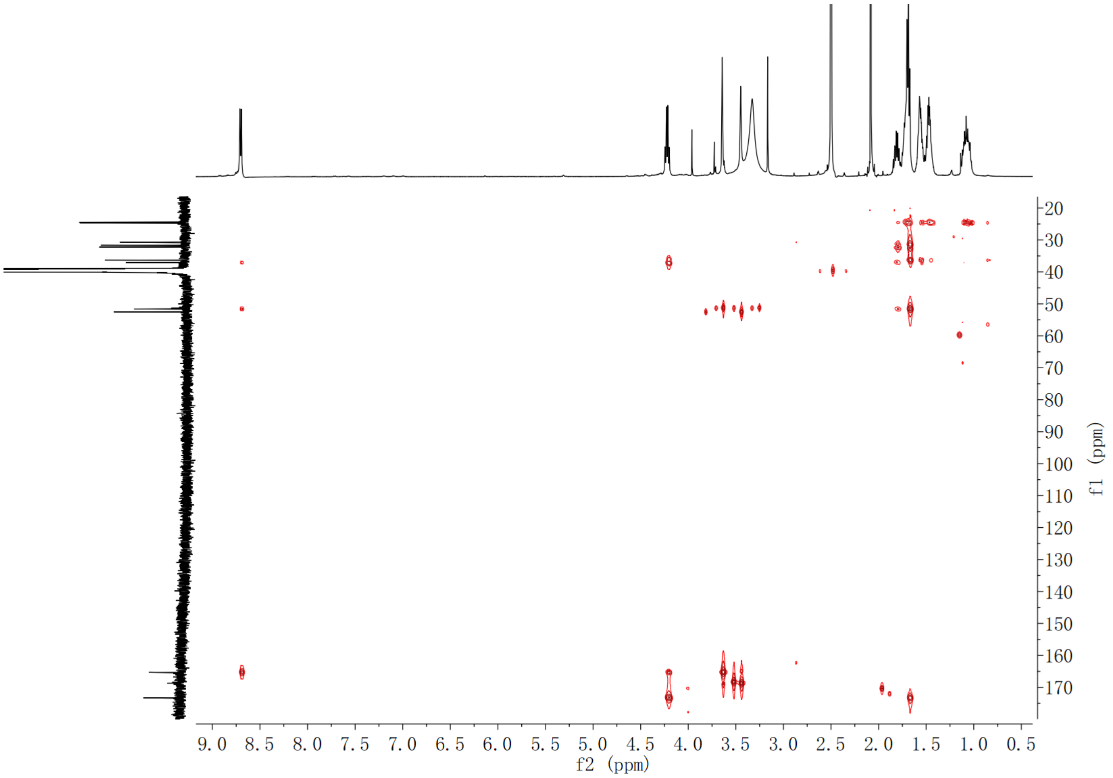

**Supplementary Fig. 157. HMBC spectrum of compound (2*S*,3*S*)-*t*-ES-a7 in DMSO-$d_6$**

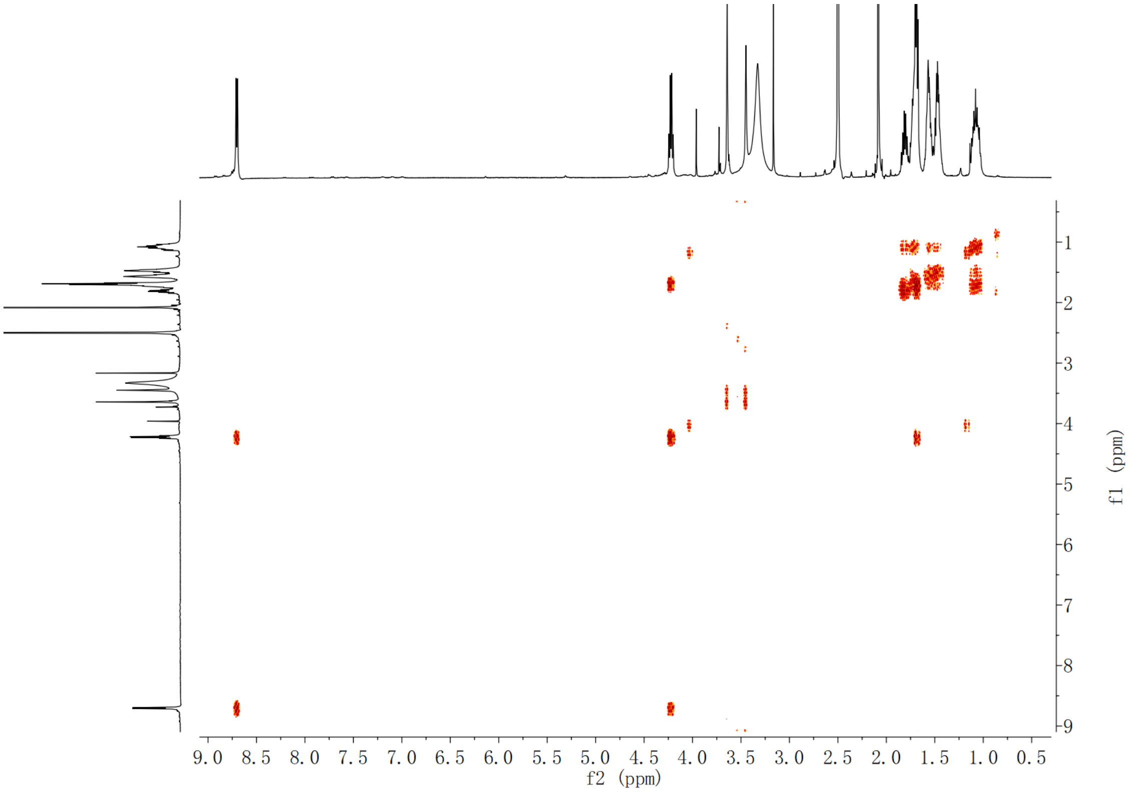

**Supplementary Fig. 158. $^1$H-$^1$H COSY spectrum of compound (2*S*,3*S*)-*t*-ES-a7 in DMSO-$d_6$**

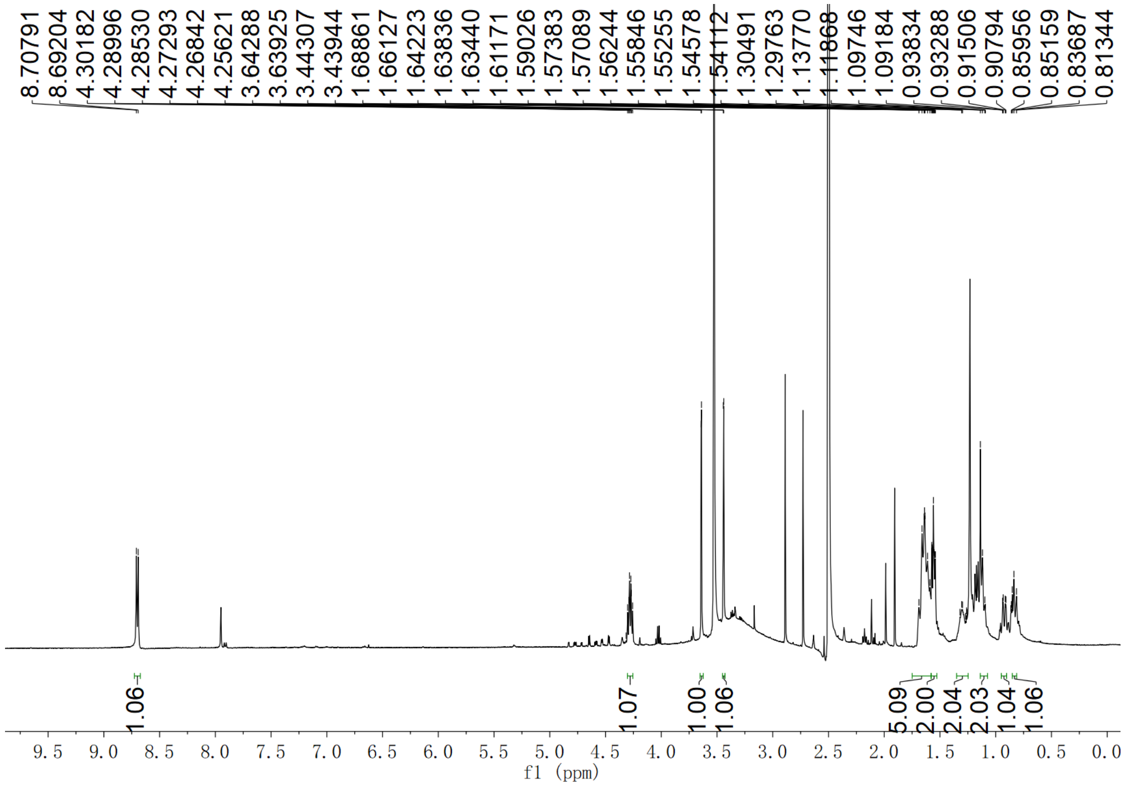

**Supplementary Fig. 159. $^{1}$H NMR spectrum of compound (2*S*,3*S*)-*t*-ES-a8 in DMSO-$d_6$**

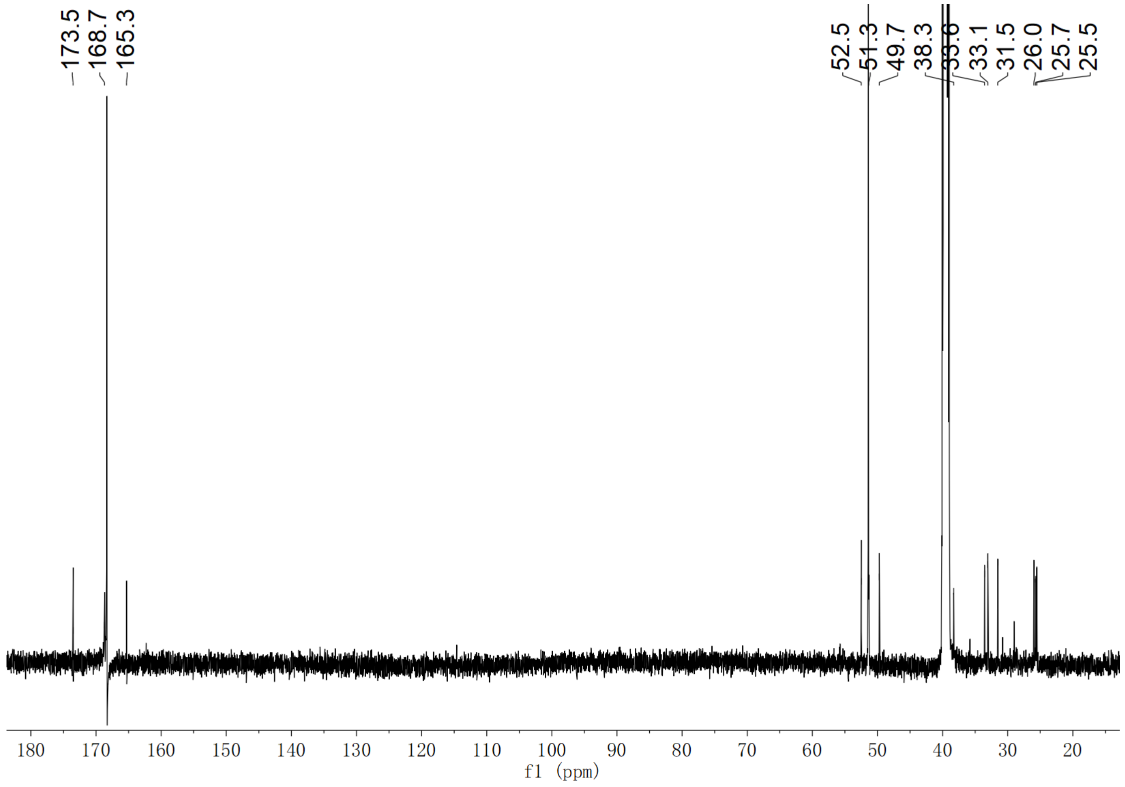

**Supplementary Fig. 160. $^{13}$C NMR spectrum of compound (2*S*,3*S*)-*t*-ES-a8 in DMSO-$d_6$**

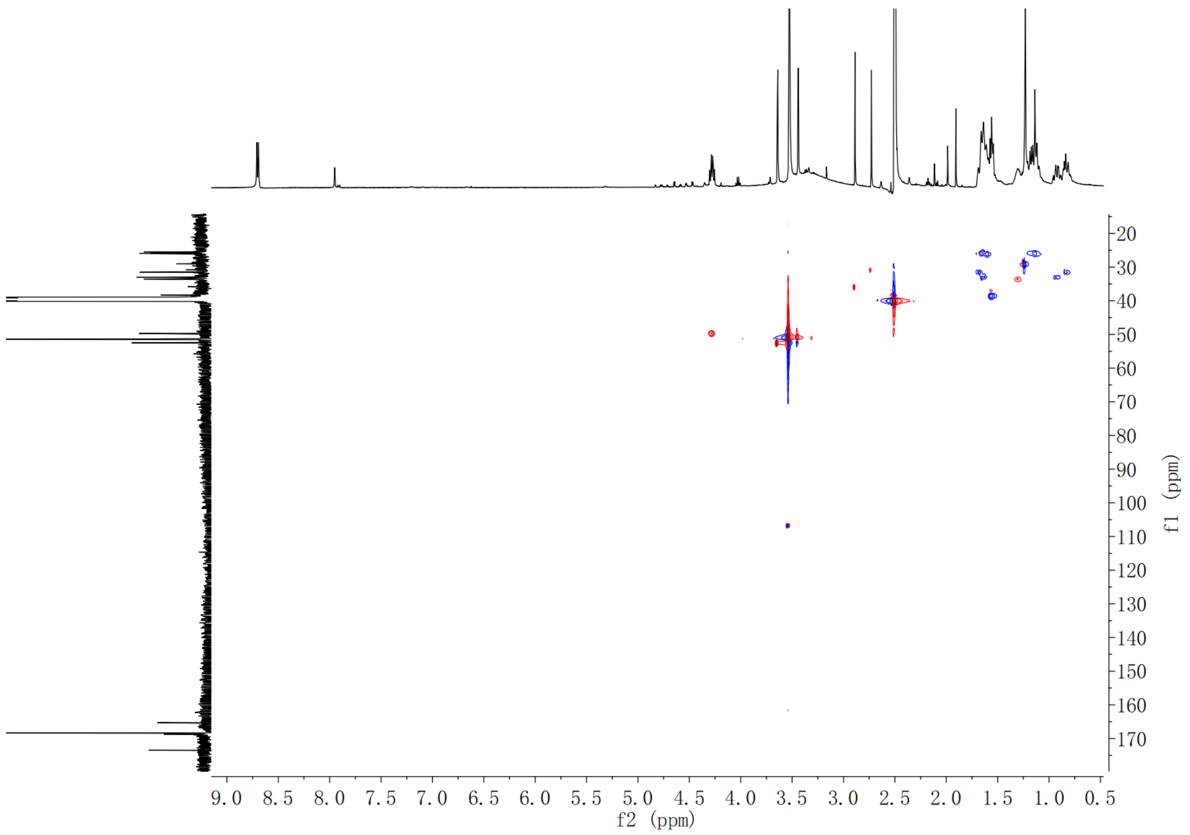

**Supplementary Fig. 161. HSQC spectrum of compound (2*S*,3*S*)-*t*-ES-a8 in DMSO-$d_6$**

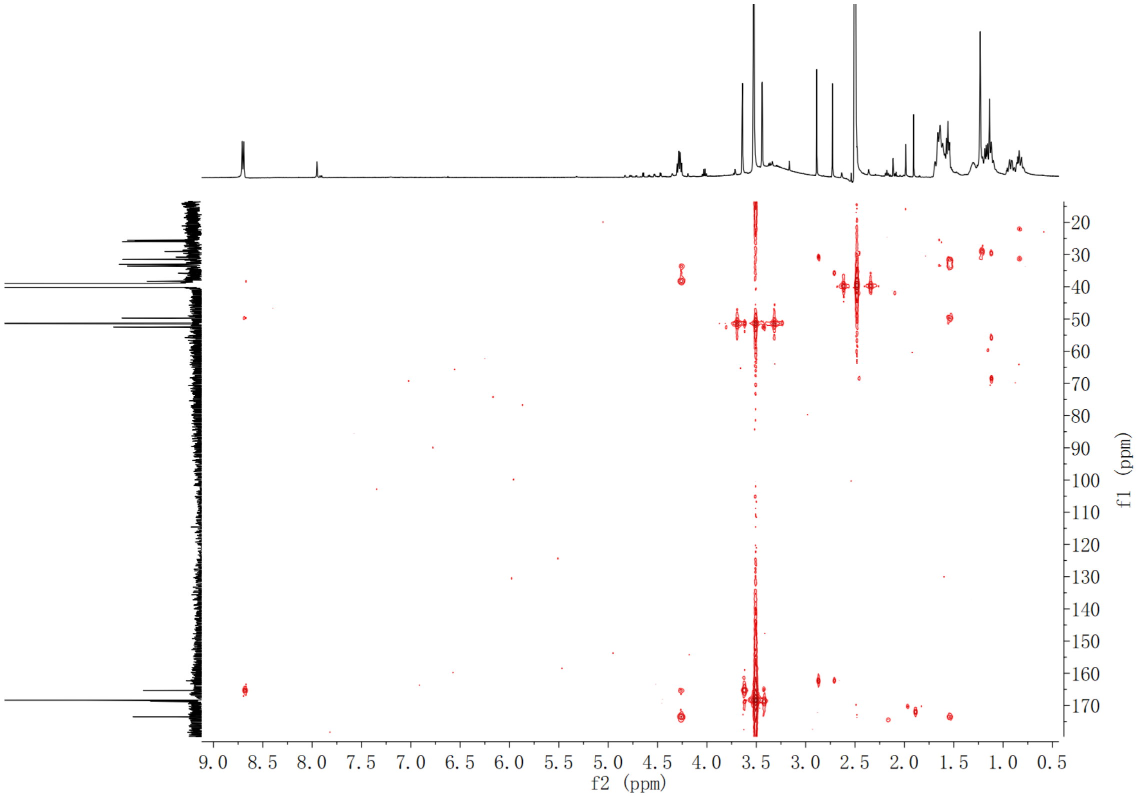

**Supplementary Fig. 162. HMBC spectrum of compound (2*S*,3*S*)-*t*-ES-a8 in DMSO-$d_6$**

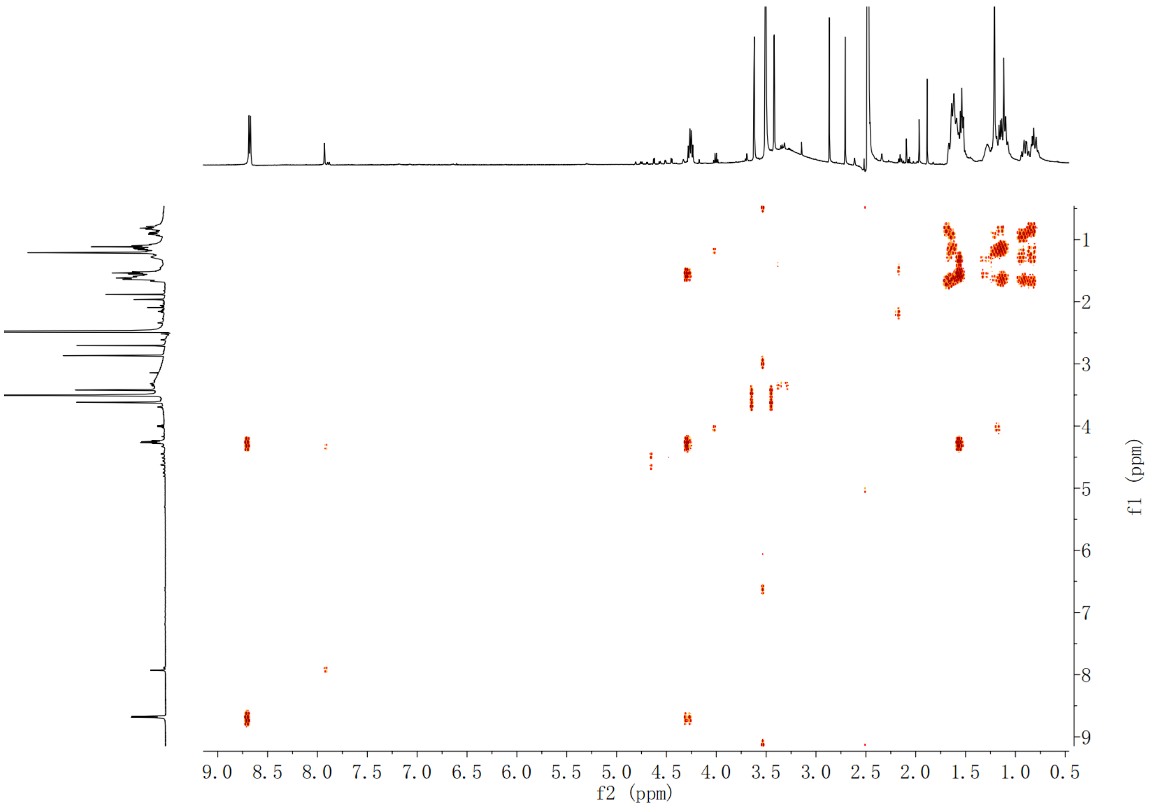

**Supplementary Fig. 163. $^{1}$H-$^{1}$H COSY spectrum of compound (2*S*,3*S*)-*t*-ES-a8 in DMSO-$d_6$**

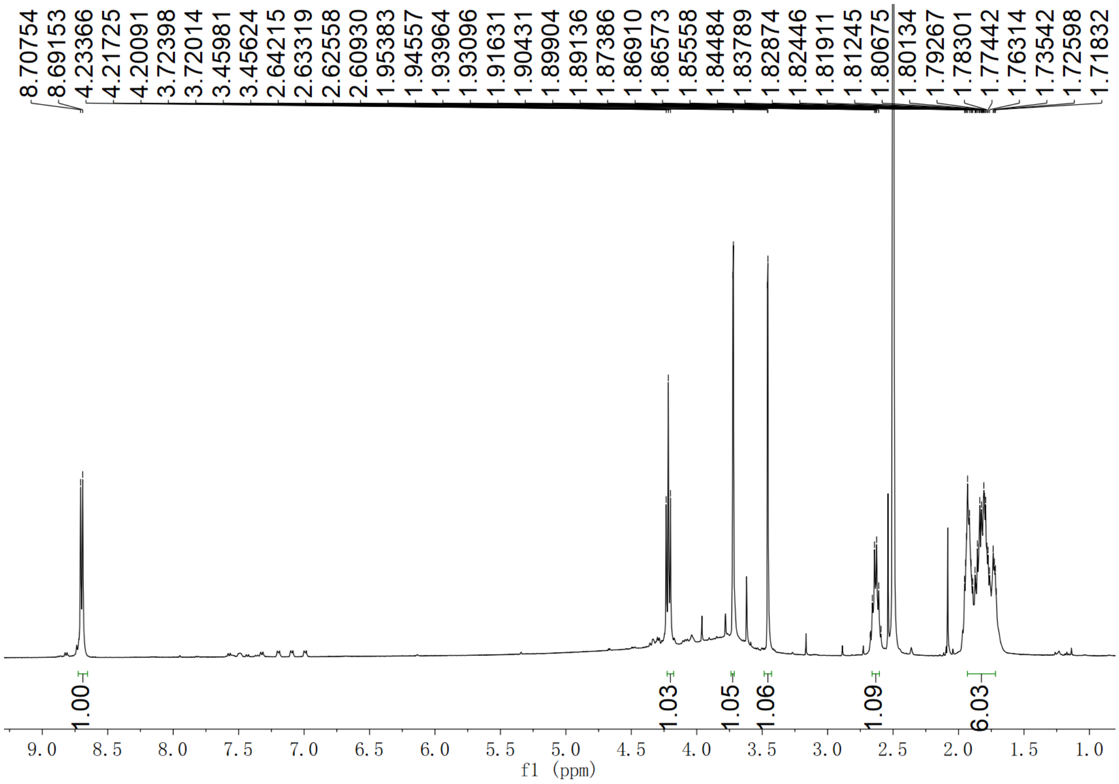

**Supplementary Fig. 164. $^{1}$H NMR spectrum of compound (2*S*,3*S*)-*t*-ES-a9 in DMSO-$d_6$**

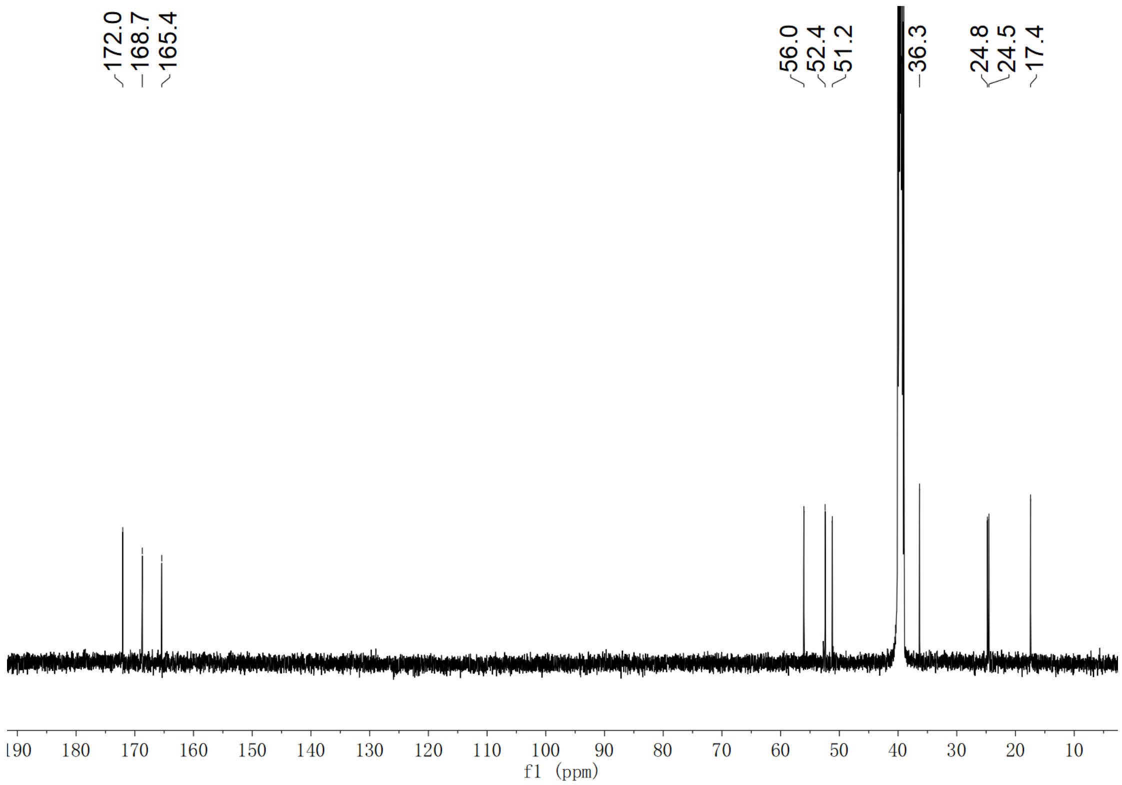

**Supplementary Fig. 165. [13]C NMR spectrum of compound (2*S*,3*S*)-*t*-ES-a9 in DMSO-*d*$_6$**

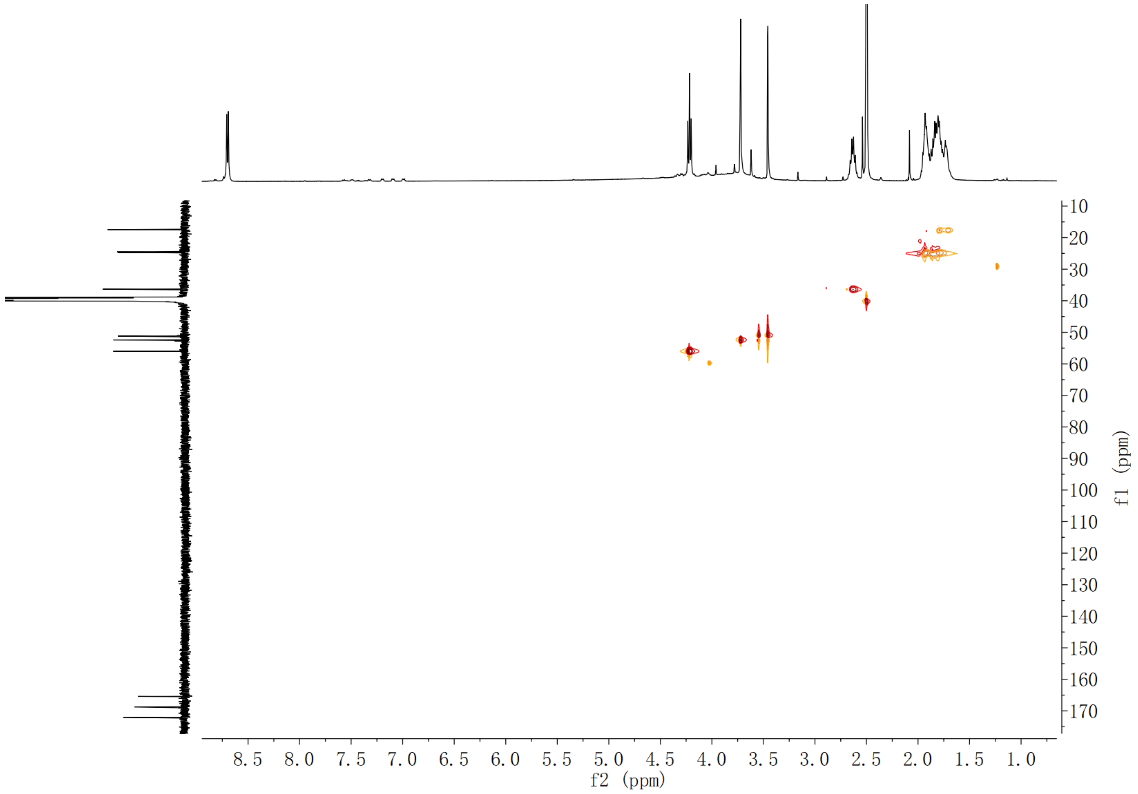

**Supplementary Fig. 166. HSQC spectrum of compound (2*S*,3*S*)-*t*-ES-a9 in DMSO-*d*$_6$**

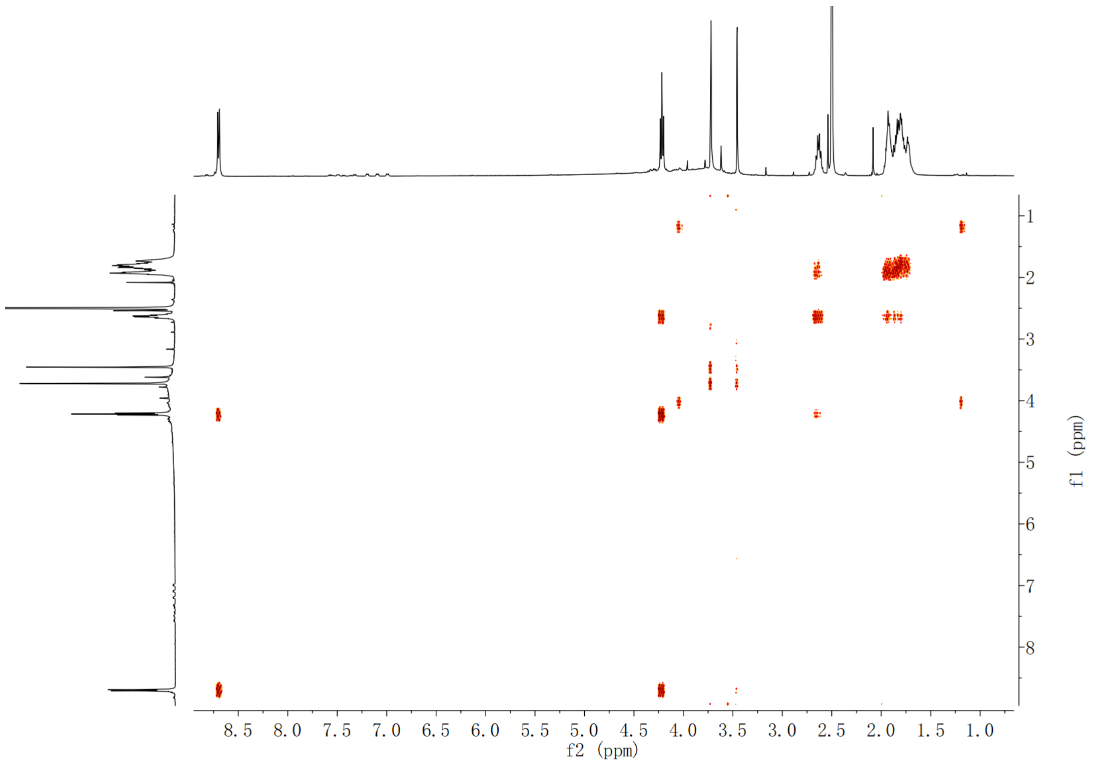

**Supplementary Fig. 167. HMBC spectrum of compound (2*S*,3*S*)-*t*-ES-a9 in DMSO-$d_6$**

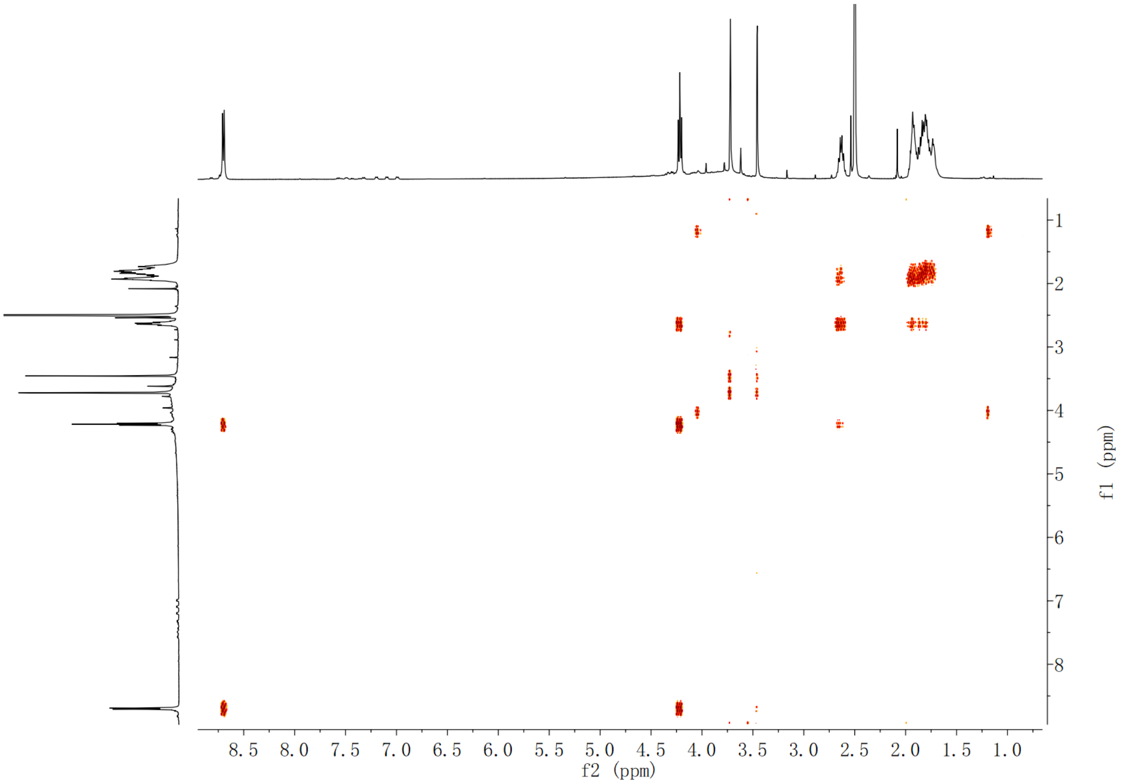

**Supplementary Fig. 168. $^{1}$H-$^{1}$H COSY spectrum of compound (2*S*,3*S*)-*t*-ES-a9 in DMSO-$d_6$**

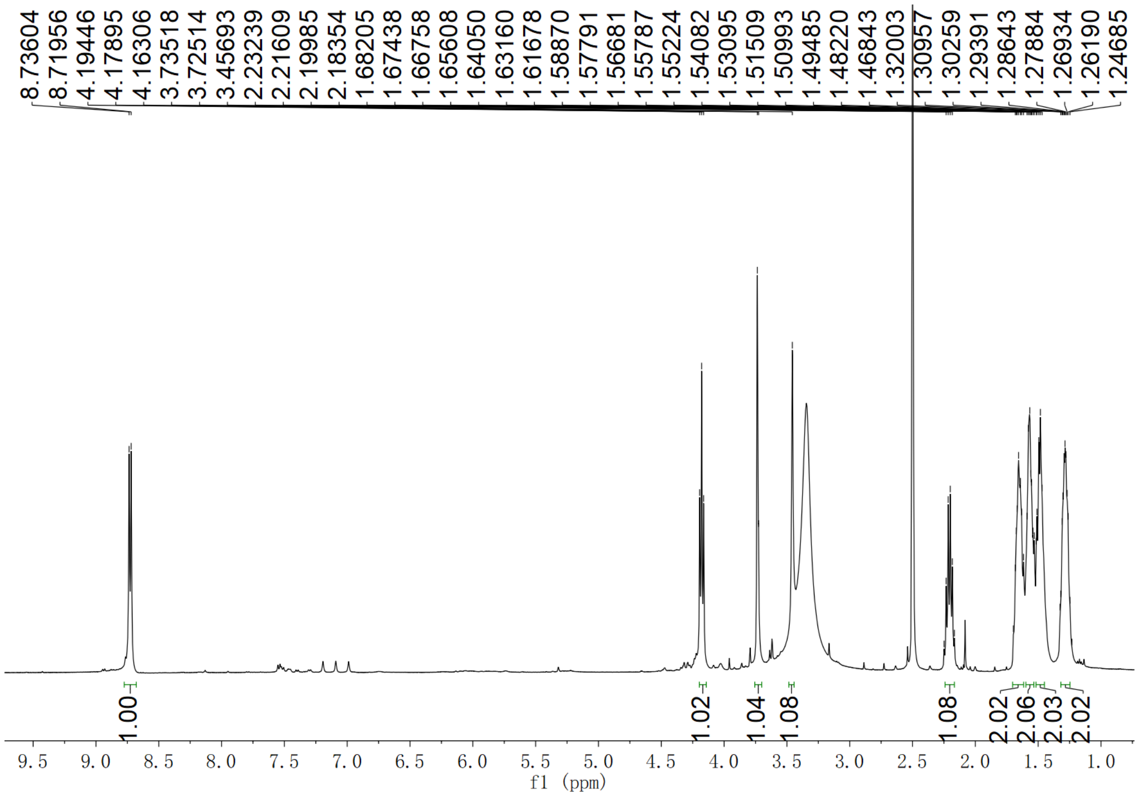

**Supplementary Fig. 169. $^{1}$H NMR spectrum of compound (2*S*,3*S*)-*t*-ES-a10 in DMSO-$d_6$**

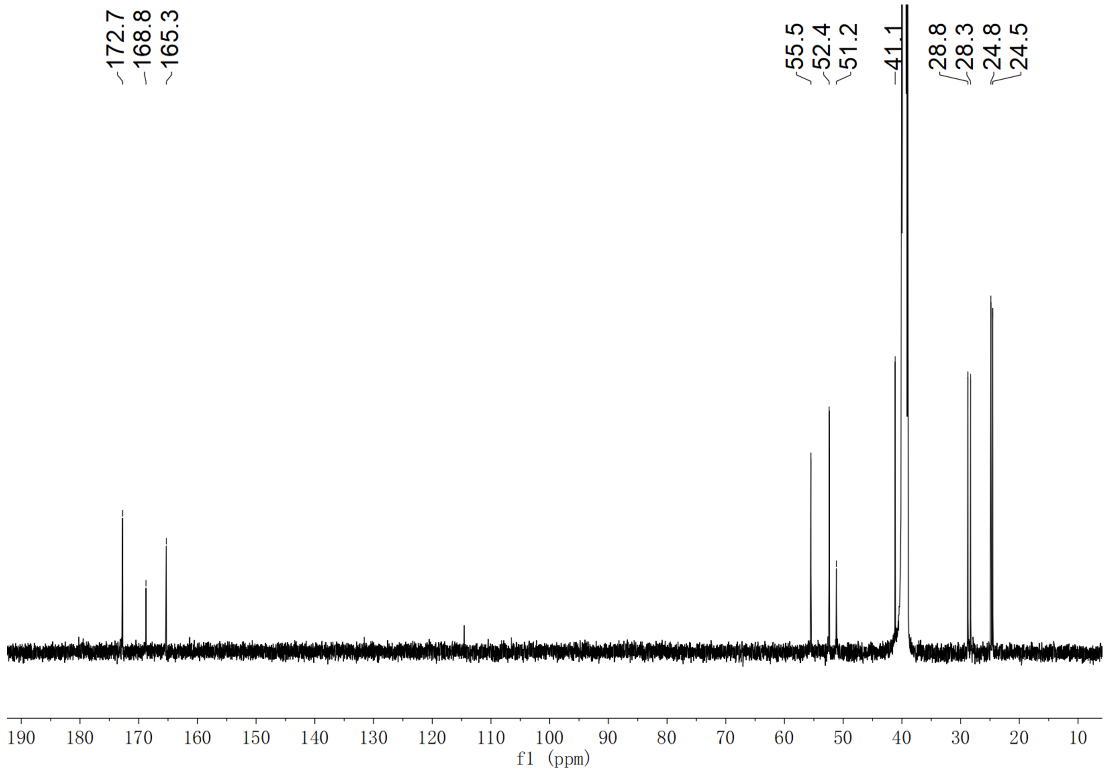

**Supplementary Fig. 170. $^{13}$C NMR spectrum of compound (2*S*,3*S*)-*t*-ES-a10 in DMSO-$d_6$**

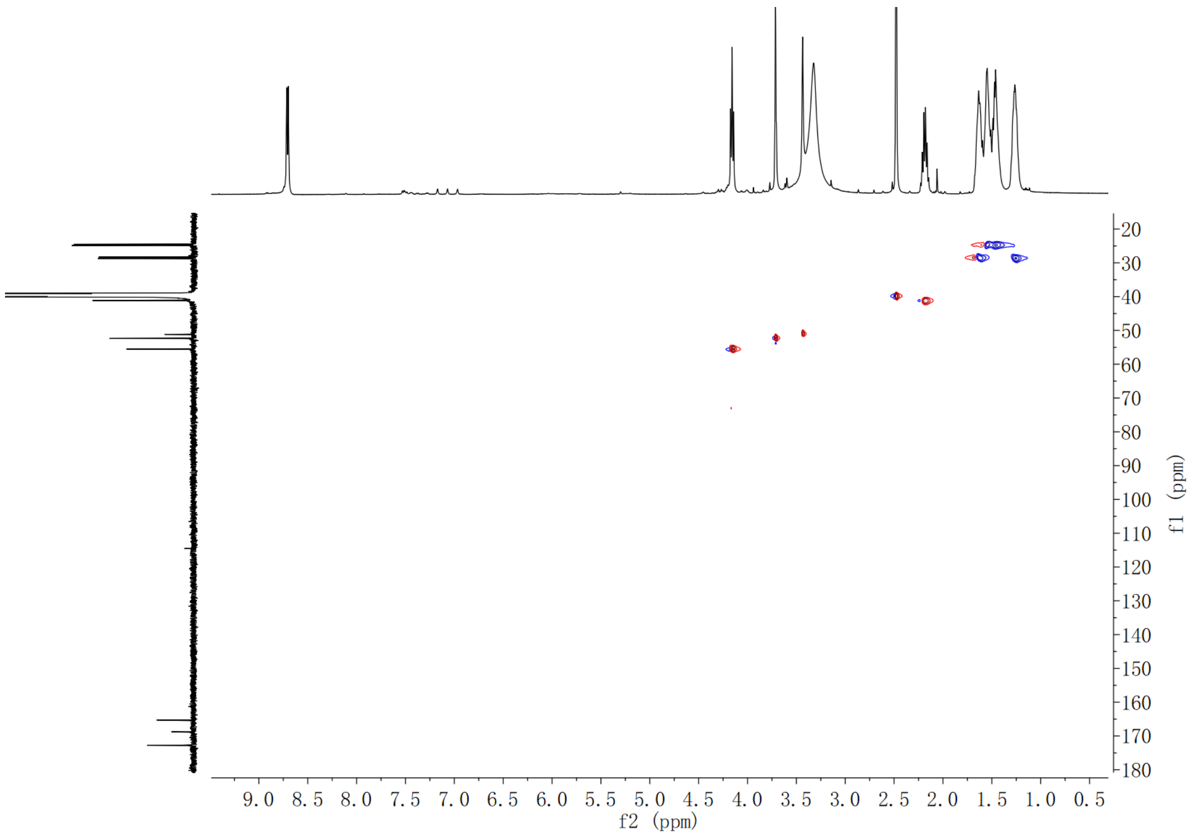

**Supplementary Fig. 171. HSQC spectrum of compound (2*S*,3*S*)-*t*-ES-a10 in DMSO-$d_6$**

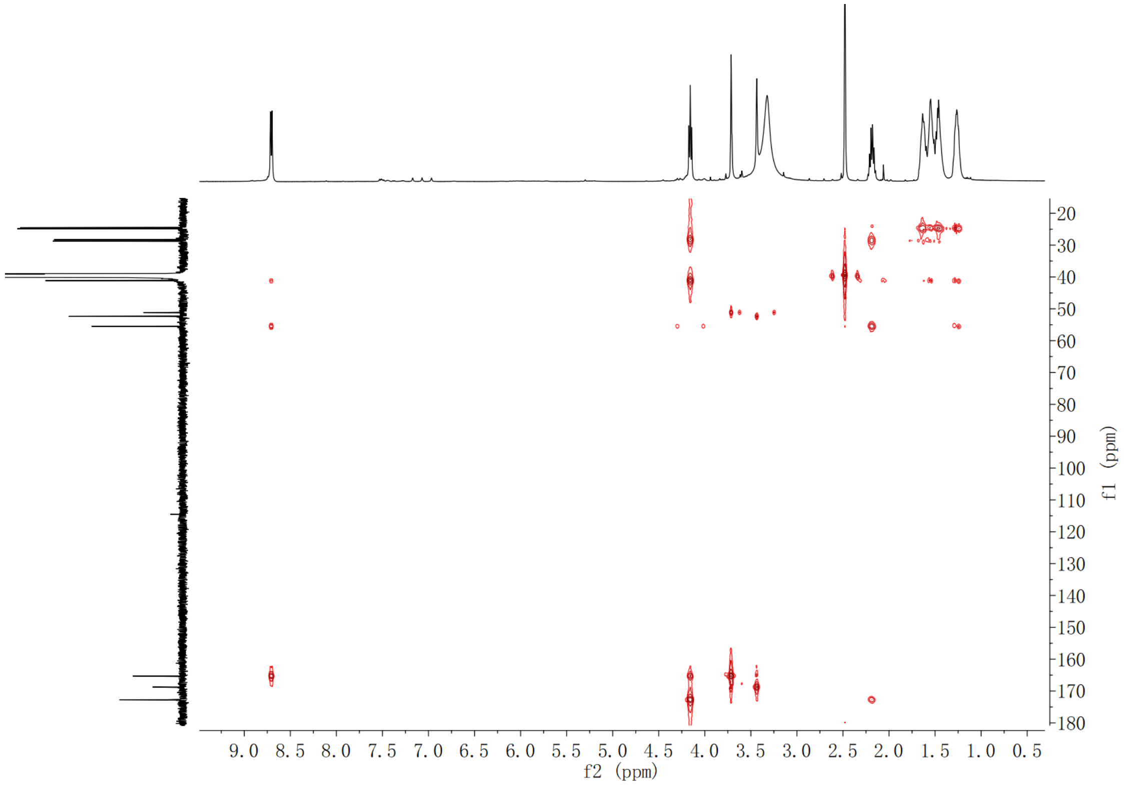

**Supplementary Fig. 172. HMBC spectrum of compound (2*S*,3*S*)-*t*-ES-a10 in DMSO-$d_6$**

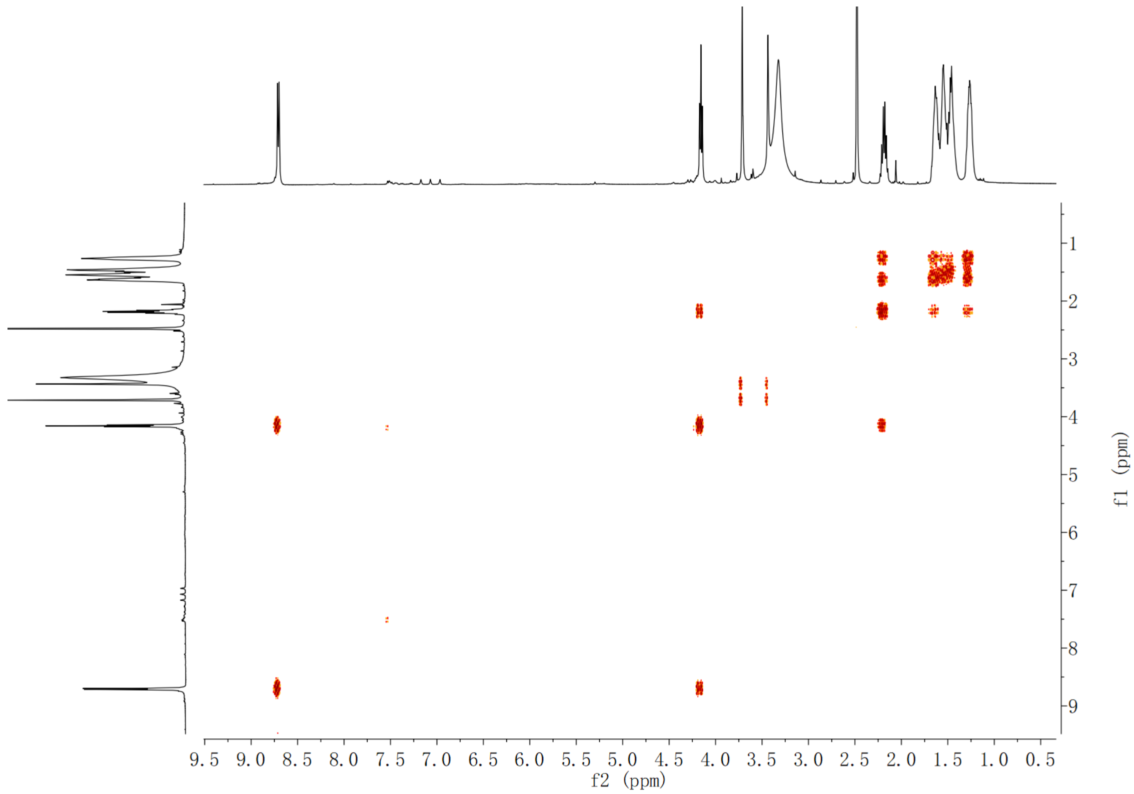

**Supplementary Fig. 173.** **¹H-¹H COSY spectrum of compound (2*S*,3*S*)-*t*-ES-a10 in DMSO-*d*₆**

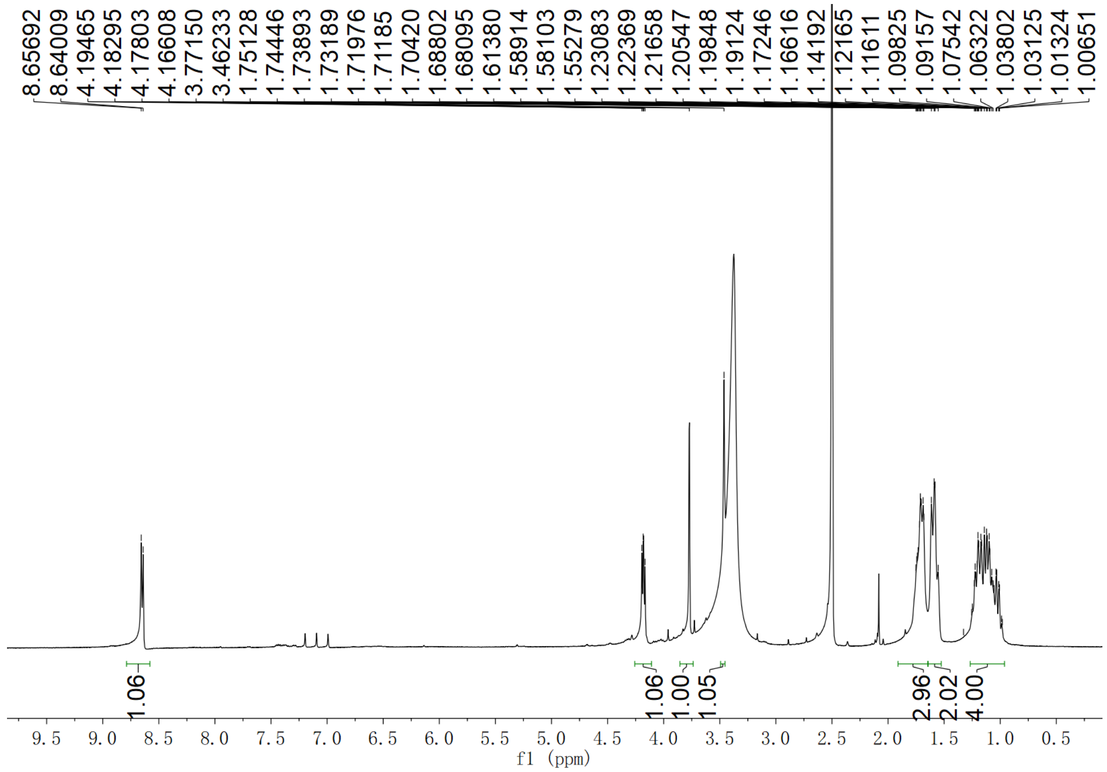

**Supplementary Fig. 174.** **¹H NMR spectrum of compound (2*S*,3*S*)-*t*-ES-a11 in DMSO-*d*₆**

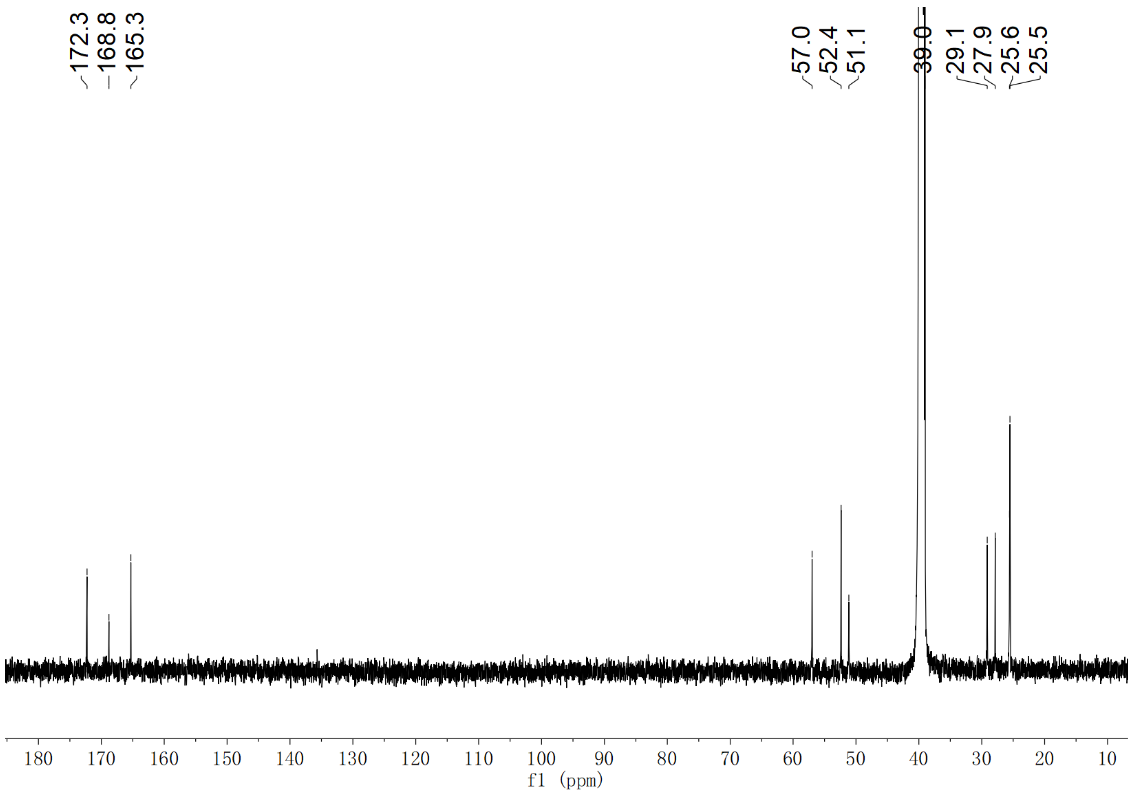

**Supplementary Fig. 175. $^{13}$C NMR spectrum of compound (2*S*,3*S*)-*t*-ES-a11 in DMSO-$d_6$**

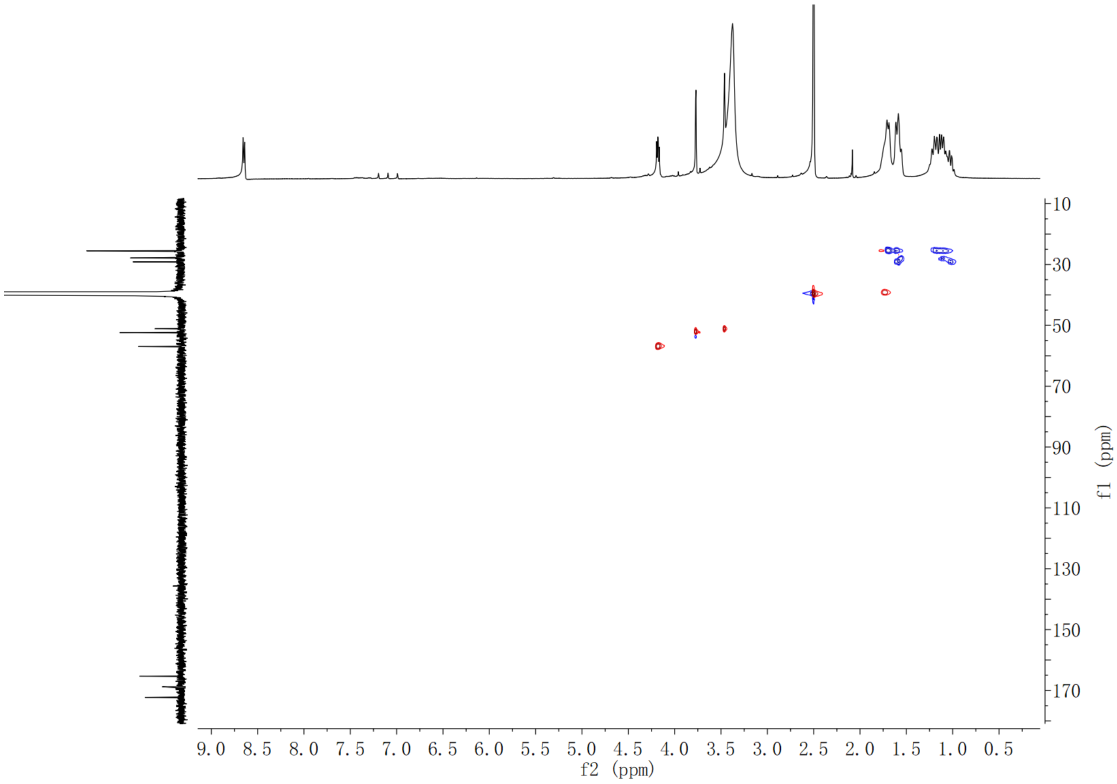

**Supplementary Fig. 176. HSQC spectrum of compound (2*S*,3*S*)-*t*-ES-a11 in DMSO-$d_6$**

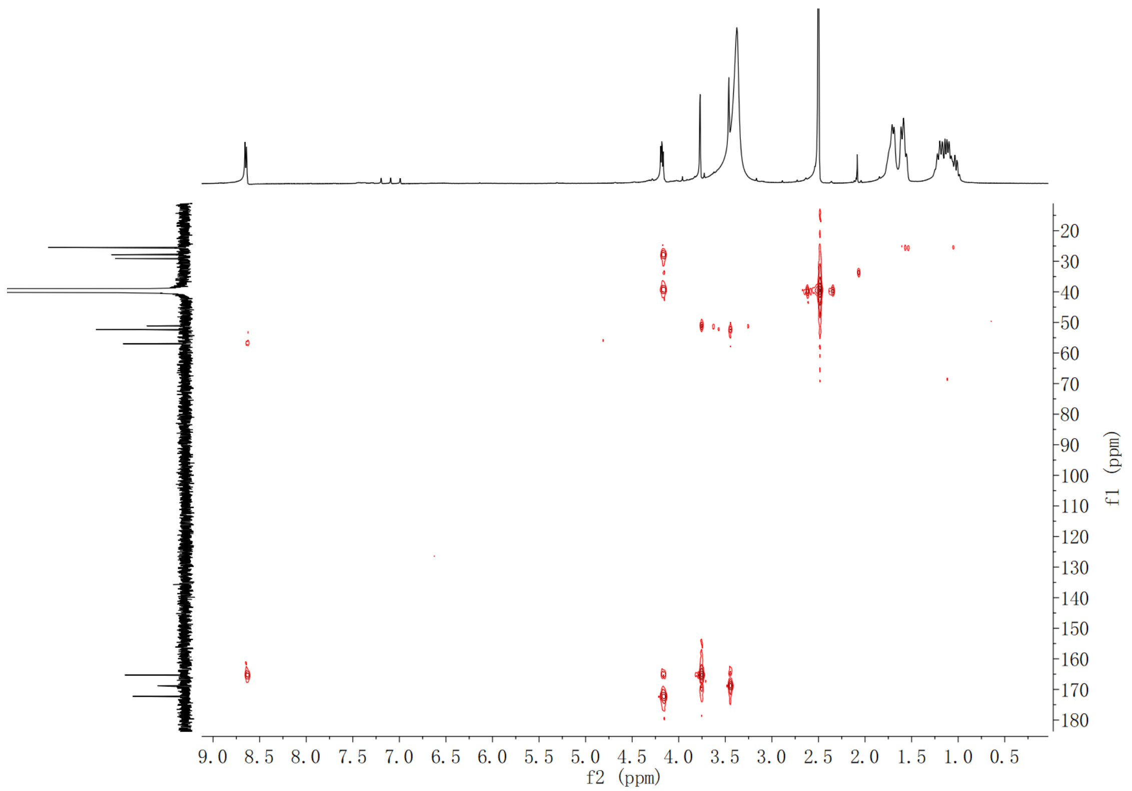

**Supplementary Fig. 177. HMBC spectrum of compound (2*S*,3*S*)-*t*-ES-a11 in DMSO-$d_6$**

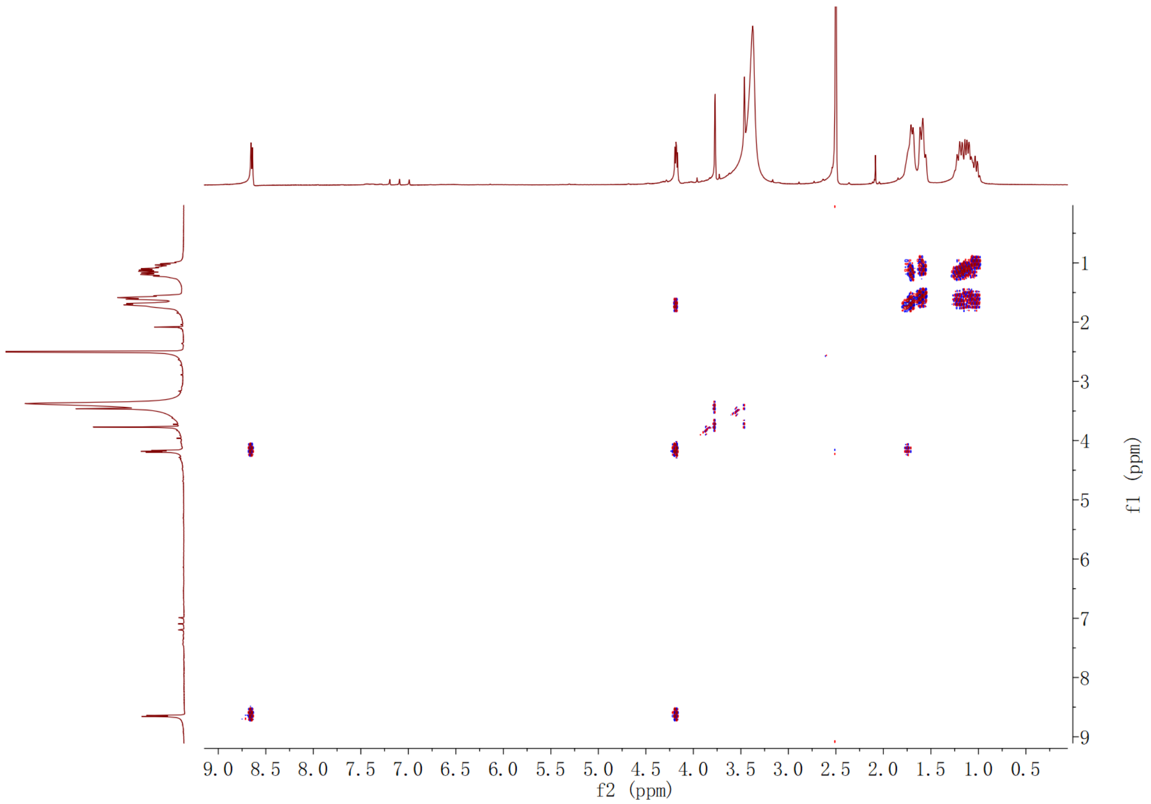

**Supplementary Fig. 178. $^1$H-$^1$H COSY spectrum of compound (2*S*,3*S*)-*t*-ES-a11 in DMSO-$d_6$**

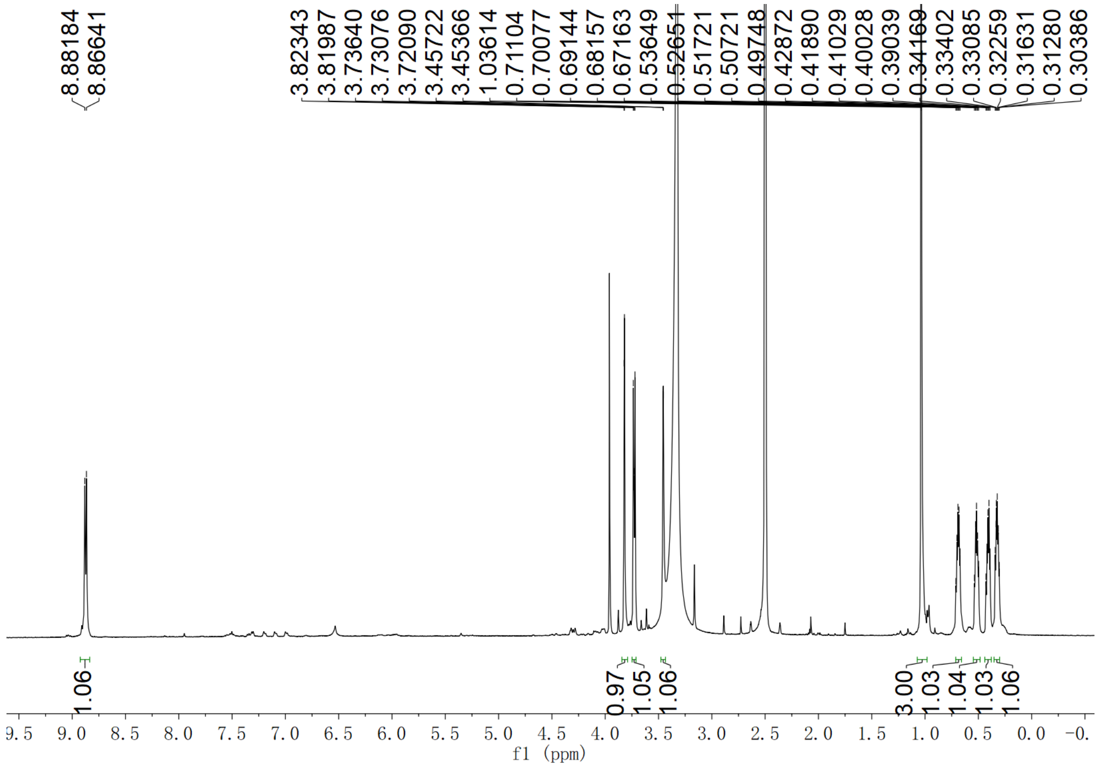

**Supplementary Fig. 179. $^{1}$H NMR spectrum of compound (2*S*,3*S*)-*t*-ES-a12 in DMSO-$d_6$**

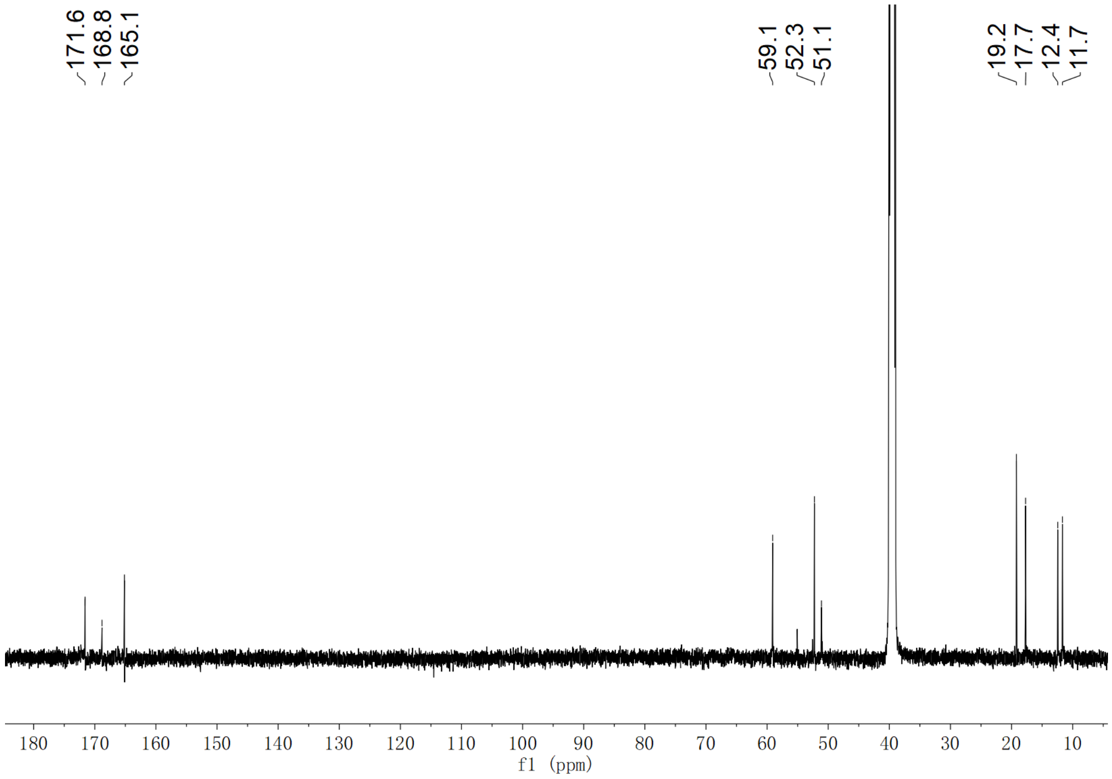

**Supplementary Fig. 180. $^{13}$C NMR spectrum of compound (2*S*,3*S*)-*t*-ES-a12 in DMSO-$d_6$**

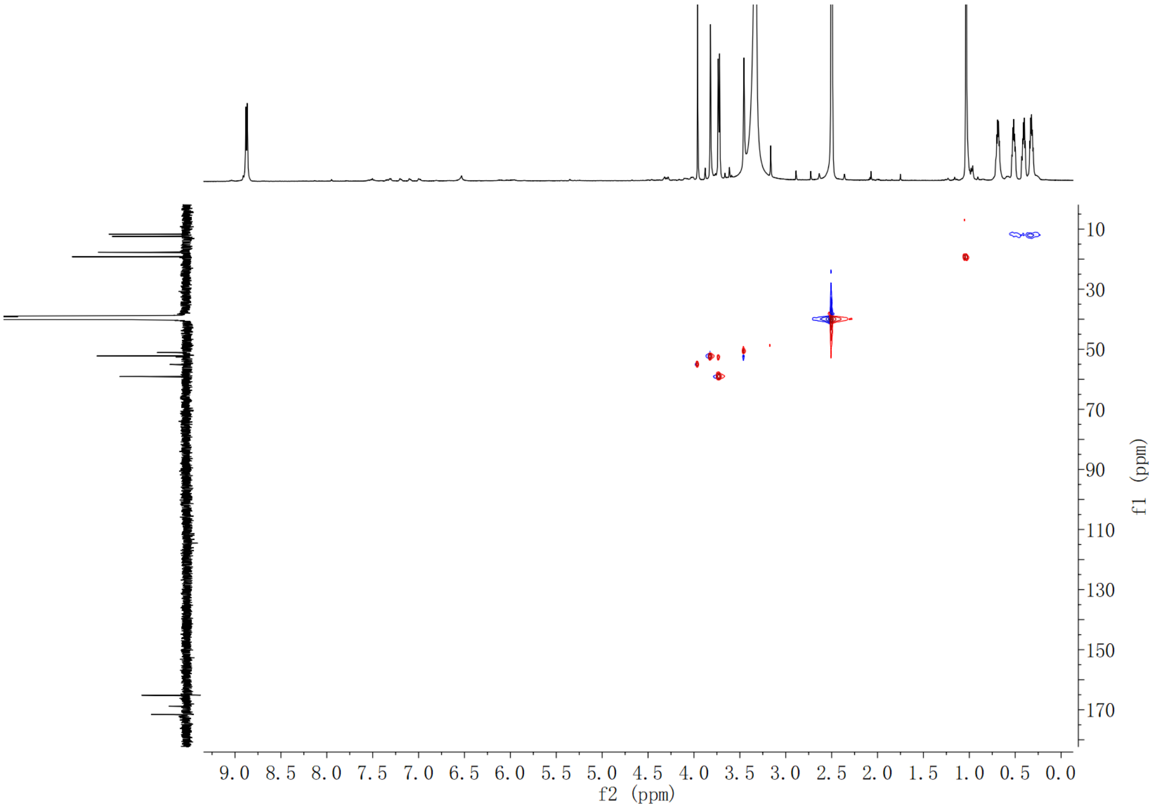

**Supplementary Fig. 181. HSQC spectrum of compound (2*S*,3*S*)-*t*-ES-a12 in DMSO-$d_6$**

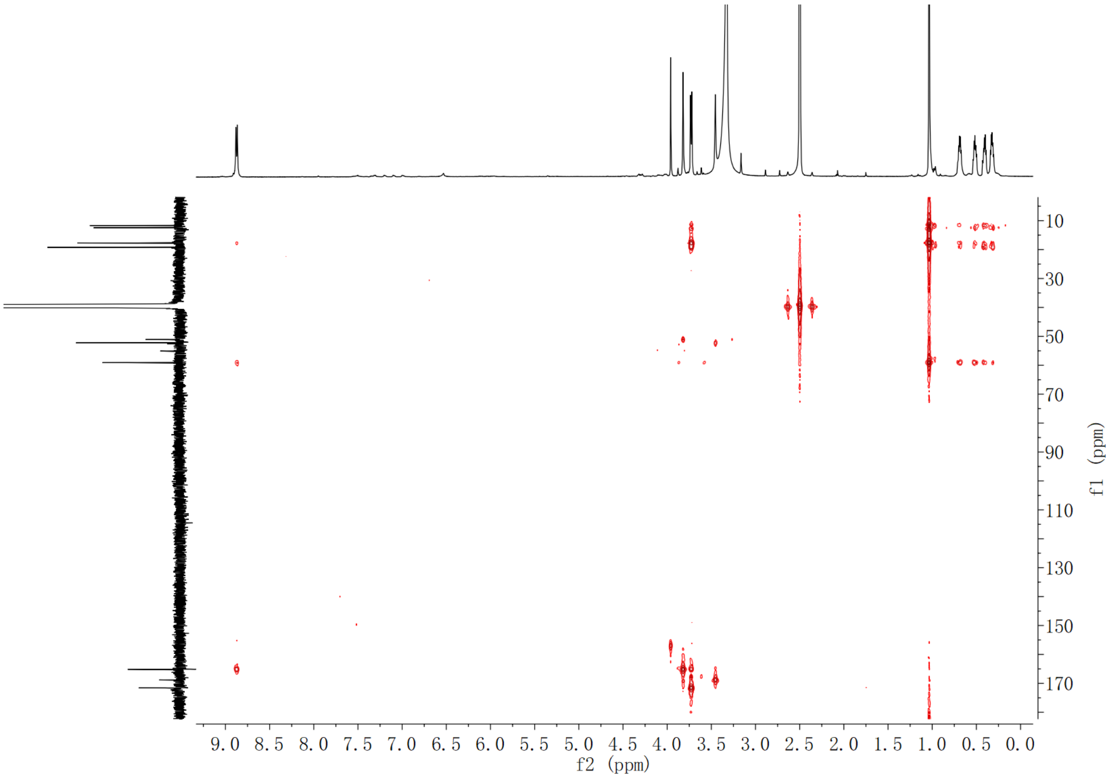

**Supplementary Fig. 182. HMBC spectrum of compound (2*S*,3*S*)-*t*-ES-a12 in DMSO-$d_6$**

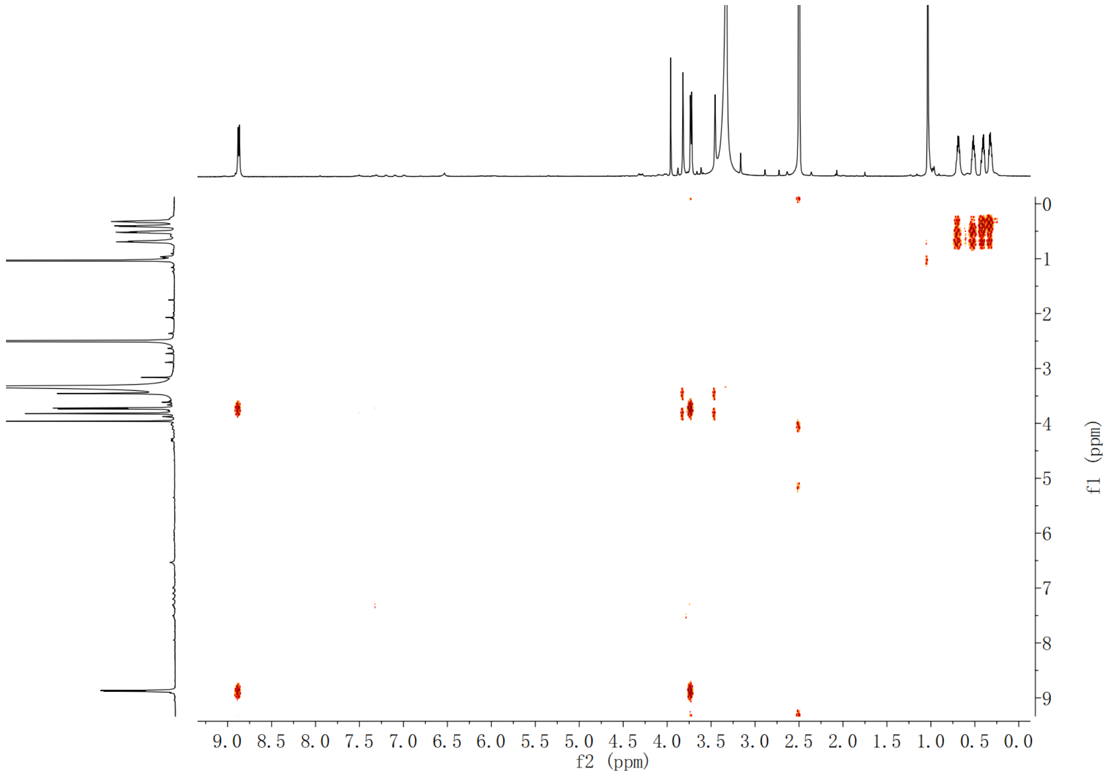

**Supplementary Fig. 183.** $^{1}$**H-**$^{1}$**H COSY spectrum of compound (2*S*,3*S*)-*t*-ES-a12 in DMSO-*d*$_6$**

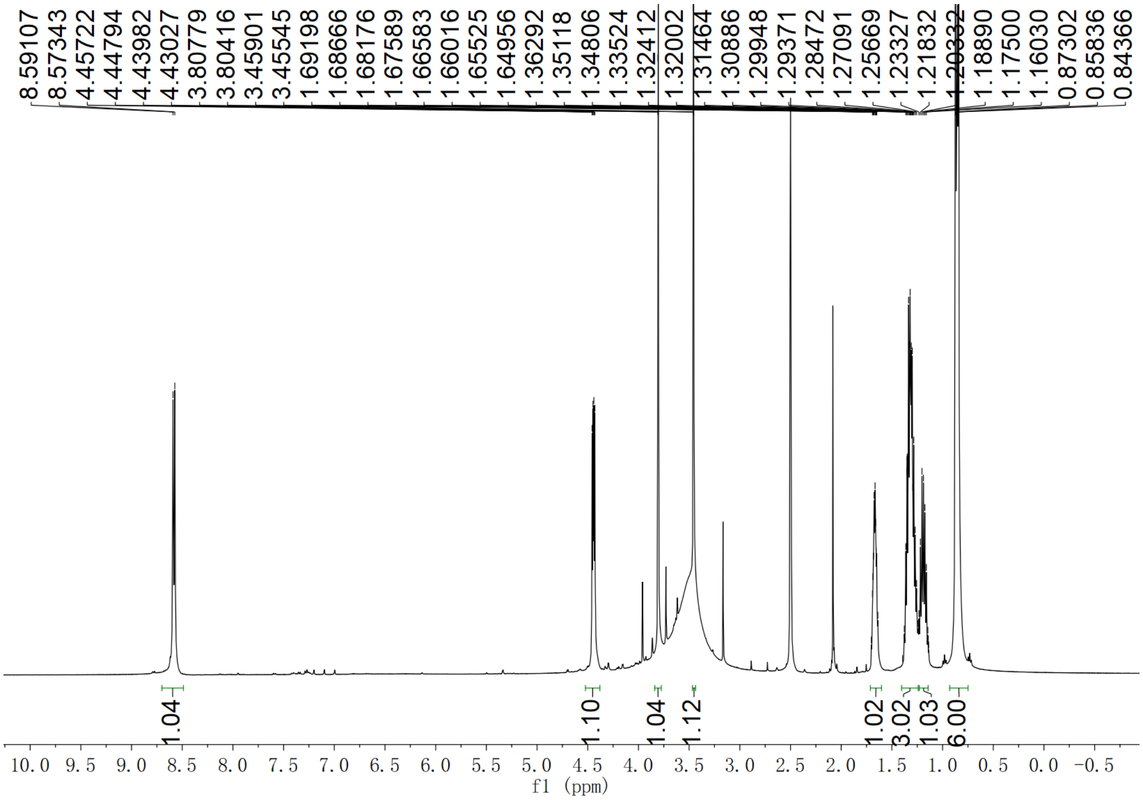

**Supplementary Fig. 184.** $^{1}$**H NMR spectrum of compound (2*S*,3*S*)-*t*-ES-a13 in DMSO-*d*$_6$**

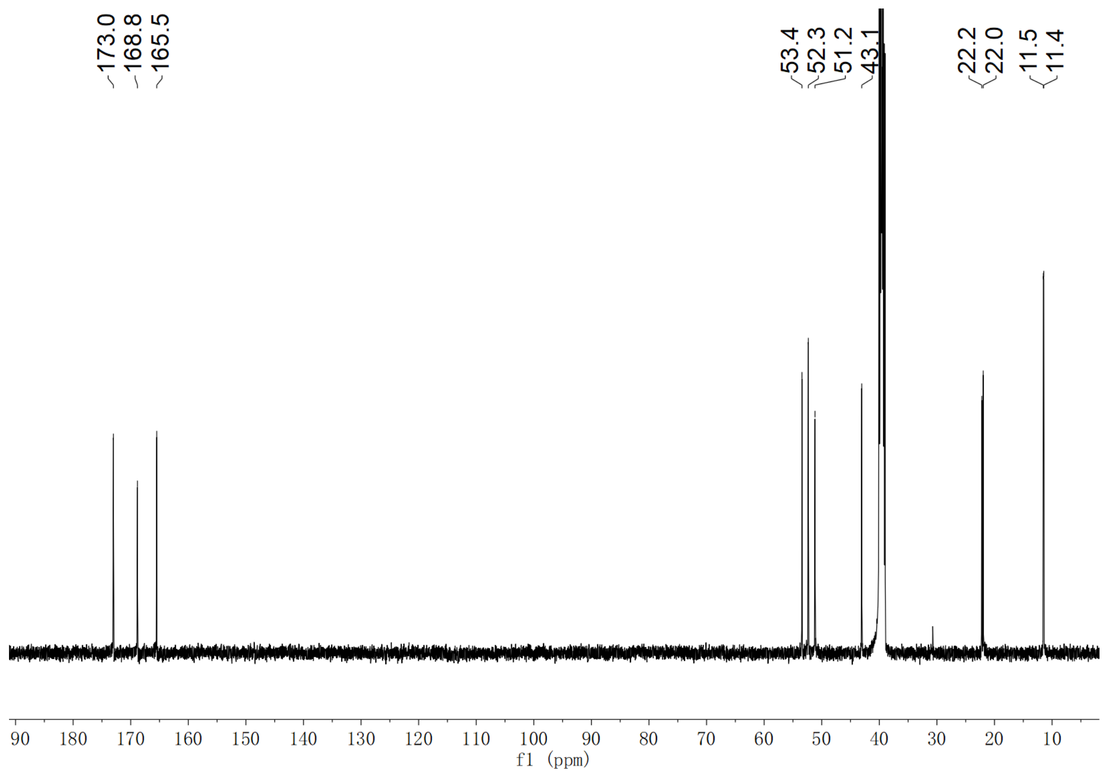

**Supplementary Fig. 185. $^{13}$C NMR spectrum of compound (2*S*,3*S*)-*t*-ES-a13 in DMSO-$d_6$**

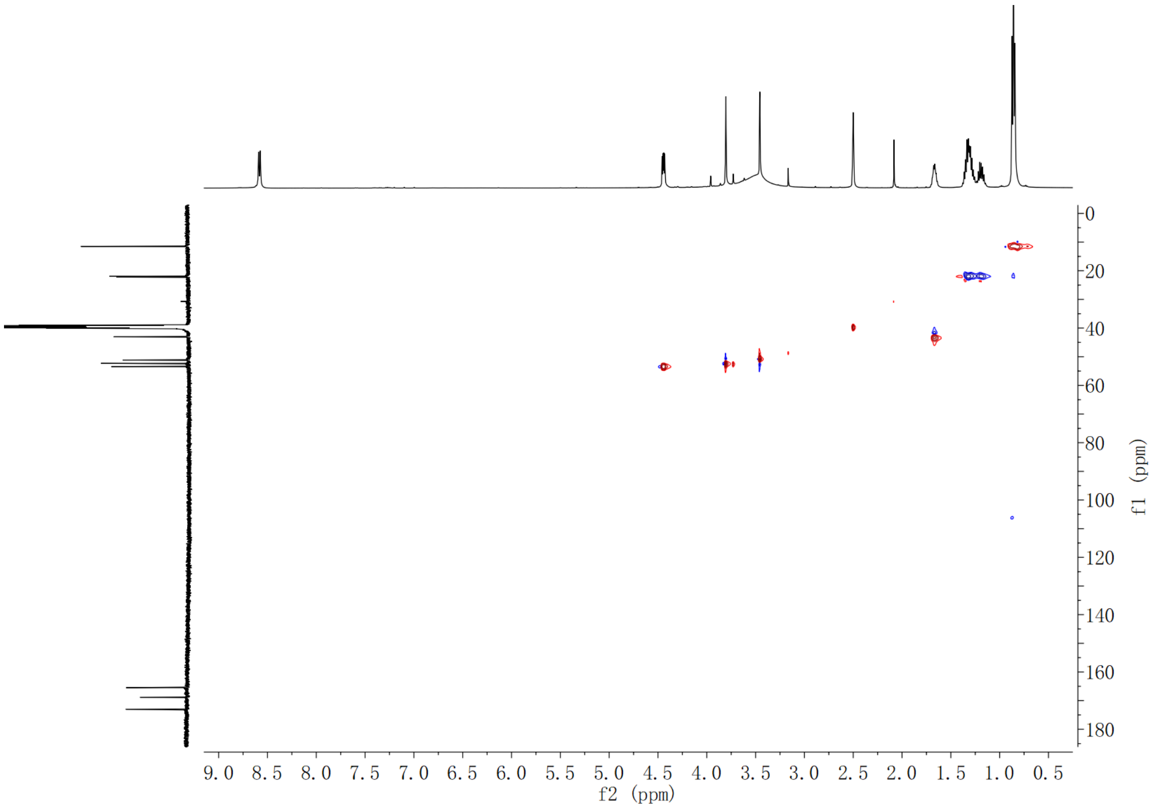

**Supplementary Fig. 186. HSQC spectrum of compound (2*S*,3*S*)-*t*-ES-a13 in DMSO-$d_6$**

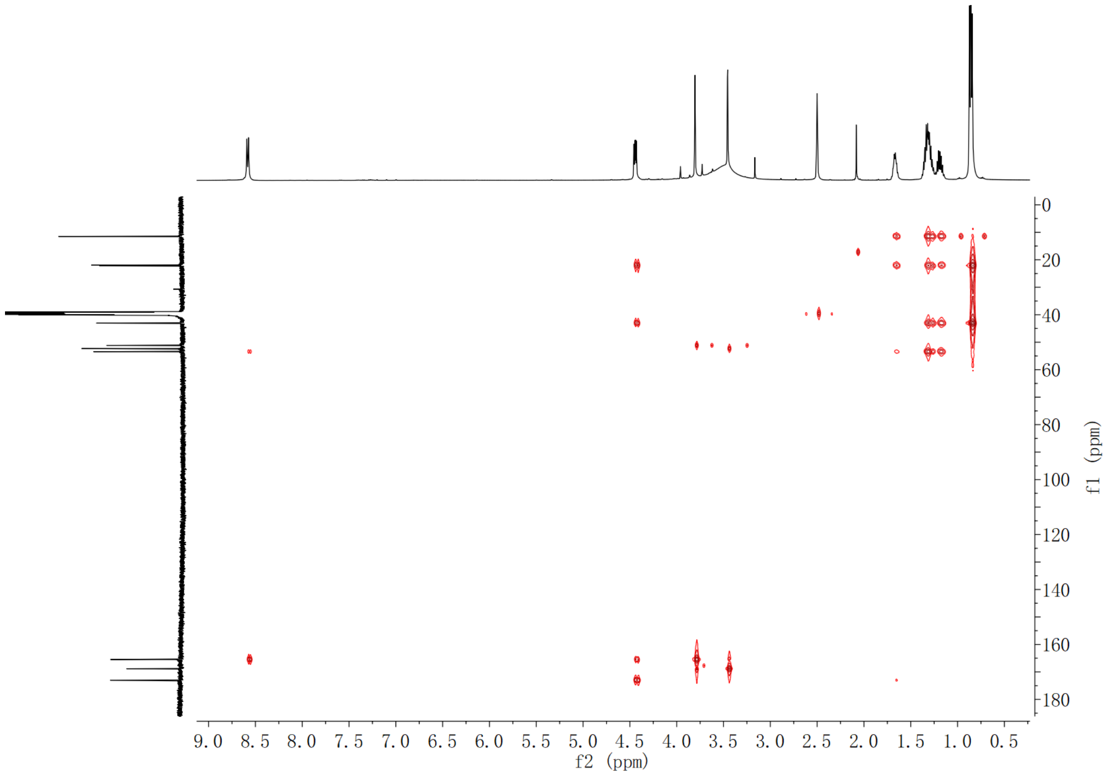

**Supplementary Fig. 187. HMBC spectrum of compound (2*S*,3*S*)-*t*-ES-a13 in DMSO-$d_6$**

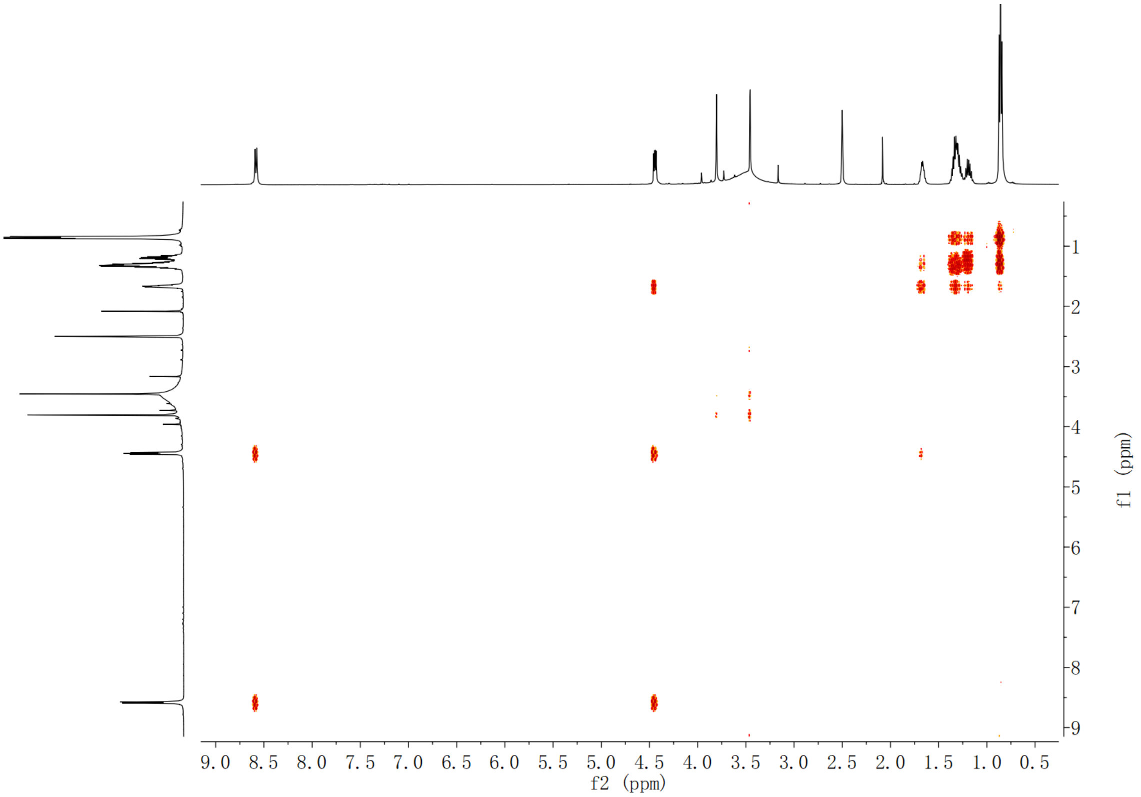

**Supplementary Fig. 188. $^{1}$H-$^{1}$H COSY spectrum of compound (2*S*,3*S*)-*t*-ES-a13 in DMSO-$d_6$**

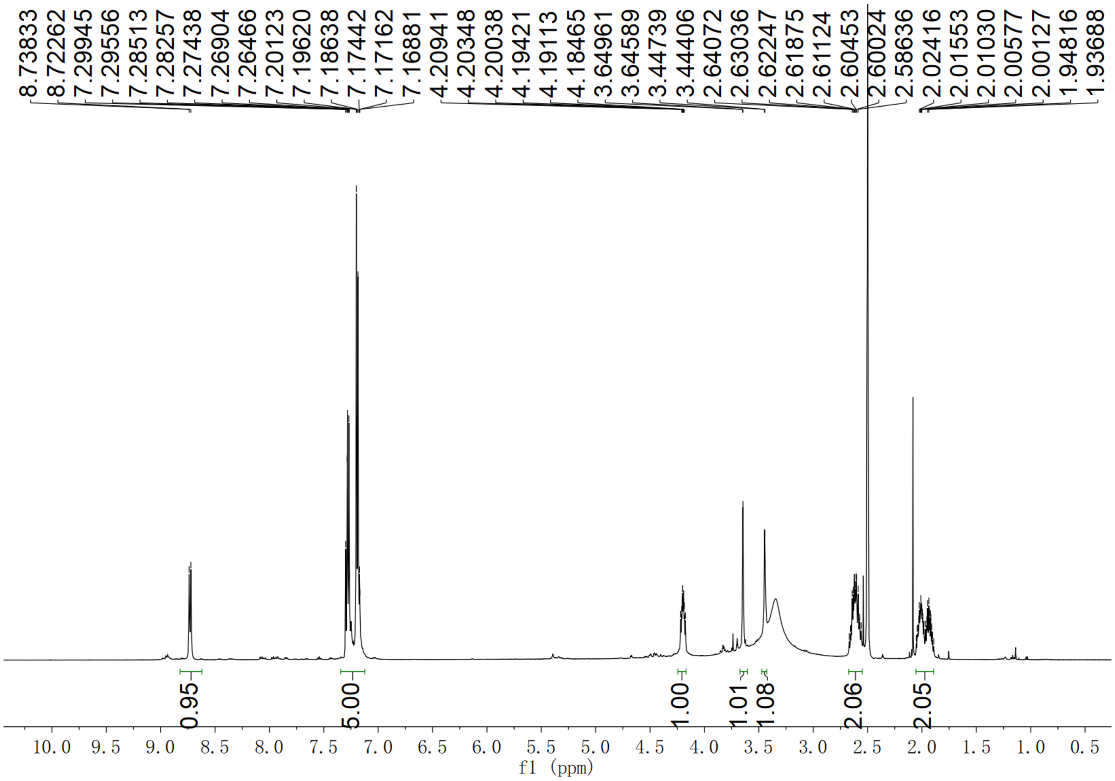

**Supplementary Fig. 189.** $^{1}$**H NMR spectrum of compound (2*S*,3*S*)-*t*-ES-a14 in DMSO-*d*$_6$**

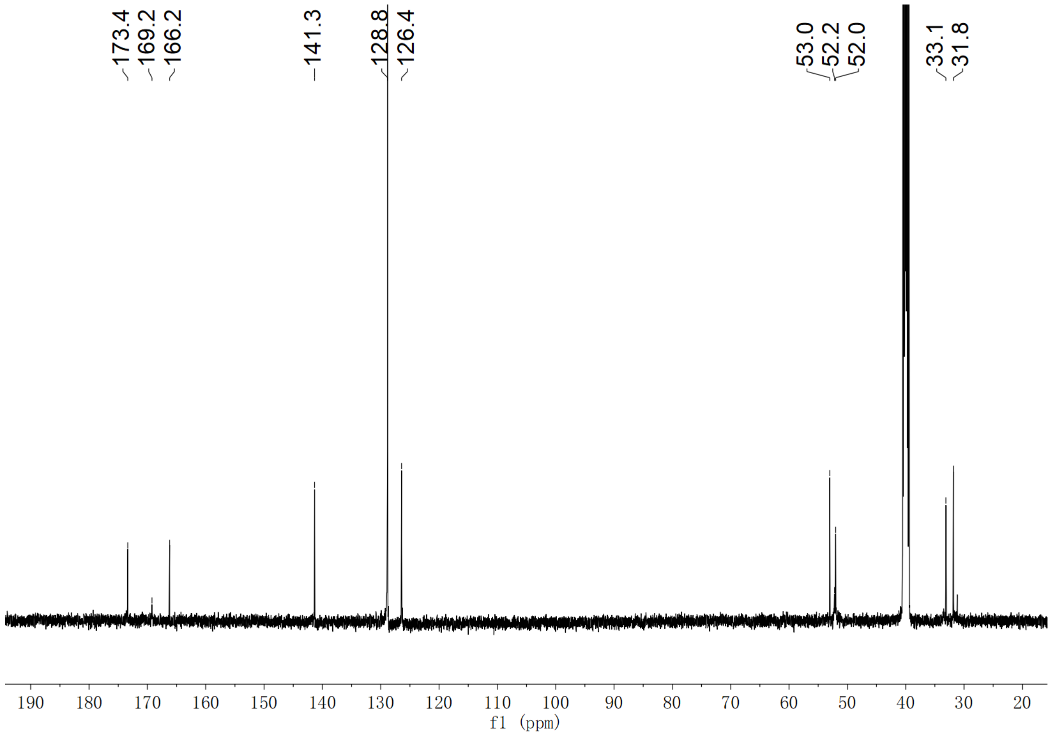

**Supplementary Fig. 190.** $^{13}$**C NMR spectrum of compound (2*S*,3*S*)-*t*-ES-a14 in DMSO-*d*$_6$**

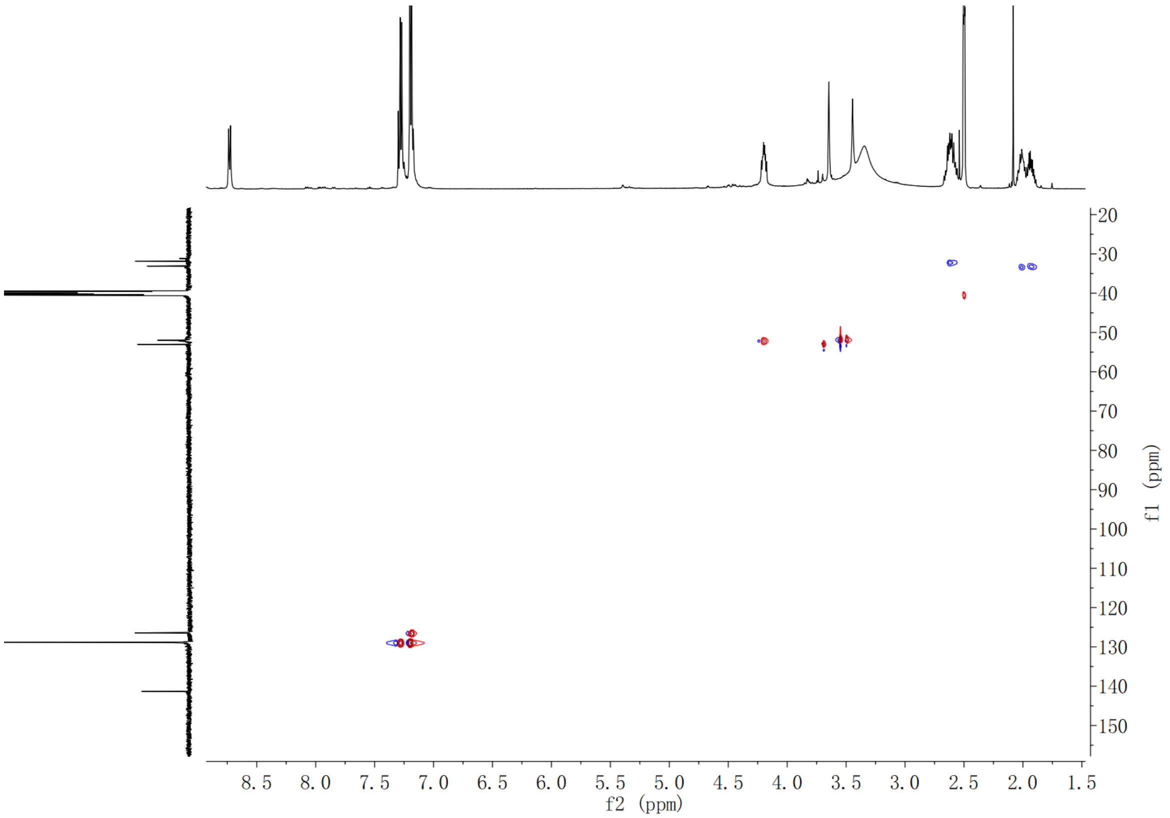

**Supplementary Fig. 191. HSQC spectrum of compound (2*S*,3*S*)-*t*-ES-a14 in DMSO-*d*$_6$**

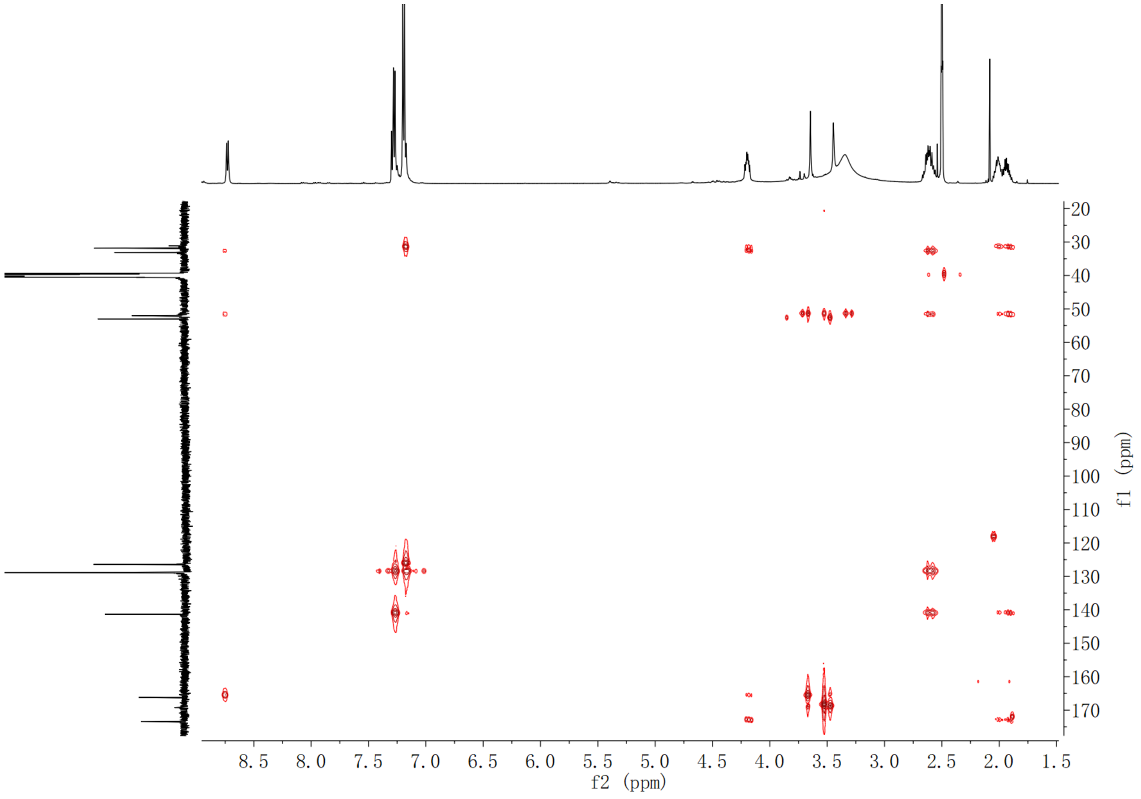

**Supplementary Fig. 192. HMBC spectrum of compound (2*S*,3*S*)-*t*-ES-a14 in DMSO-*d*$_6$**

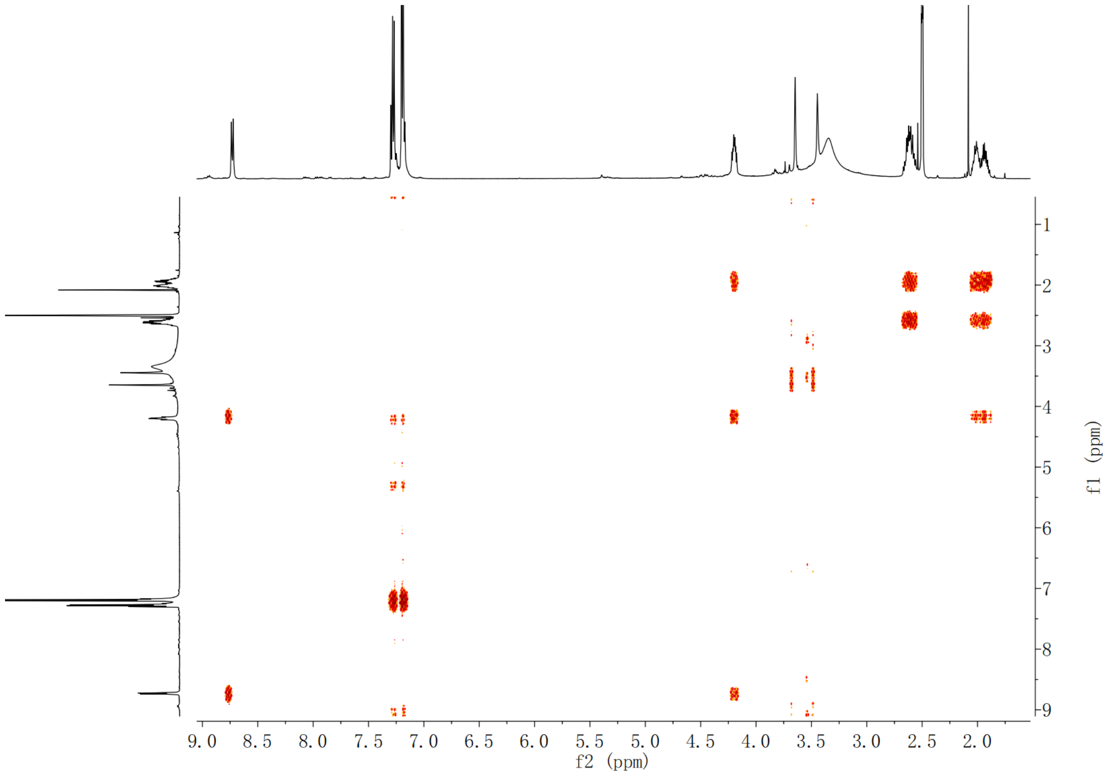

Supplementary Fig. 193. $^1$H-$^1$H COSY spectrum of compound (2*S*,3*S*)-*t*-ES-a14 in DMSO-$d_6$

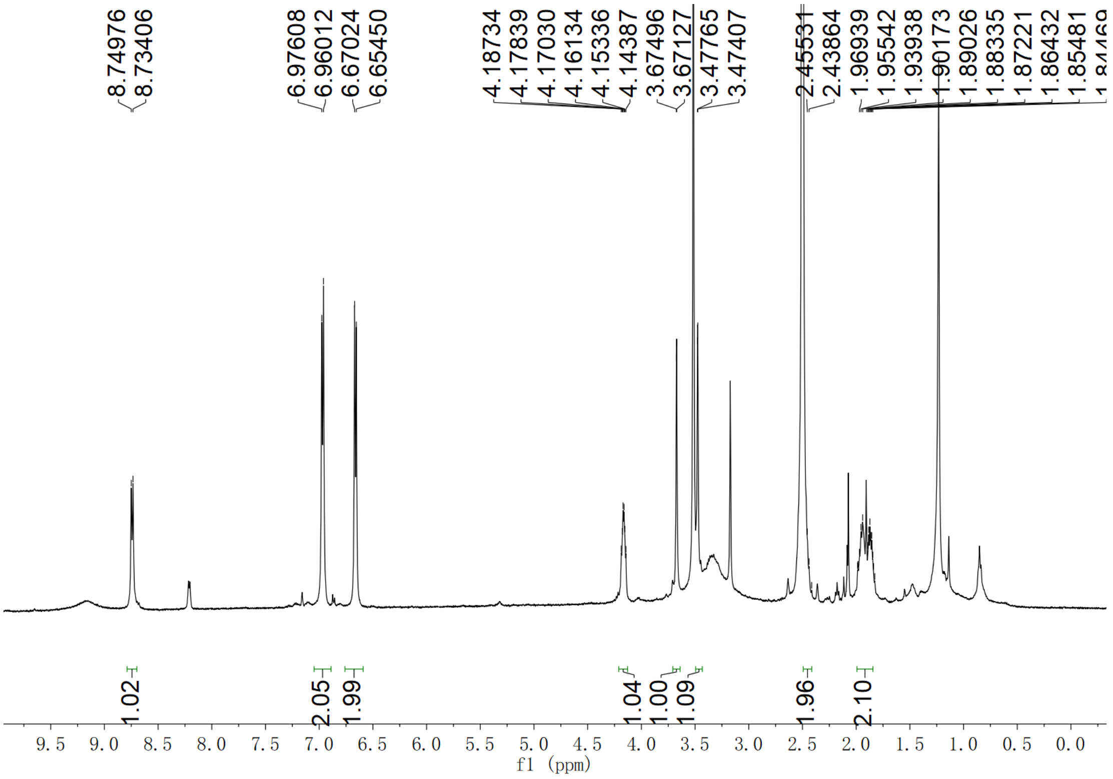

Supplementary Fig. 194. $^1$H NMR spectrum of compound (2*S*,3*S*)-*t*-ES-a15 in DMSO-$d_6$

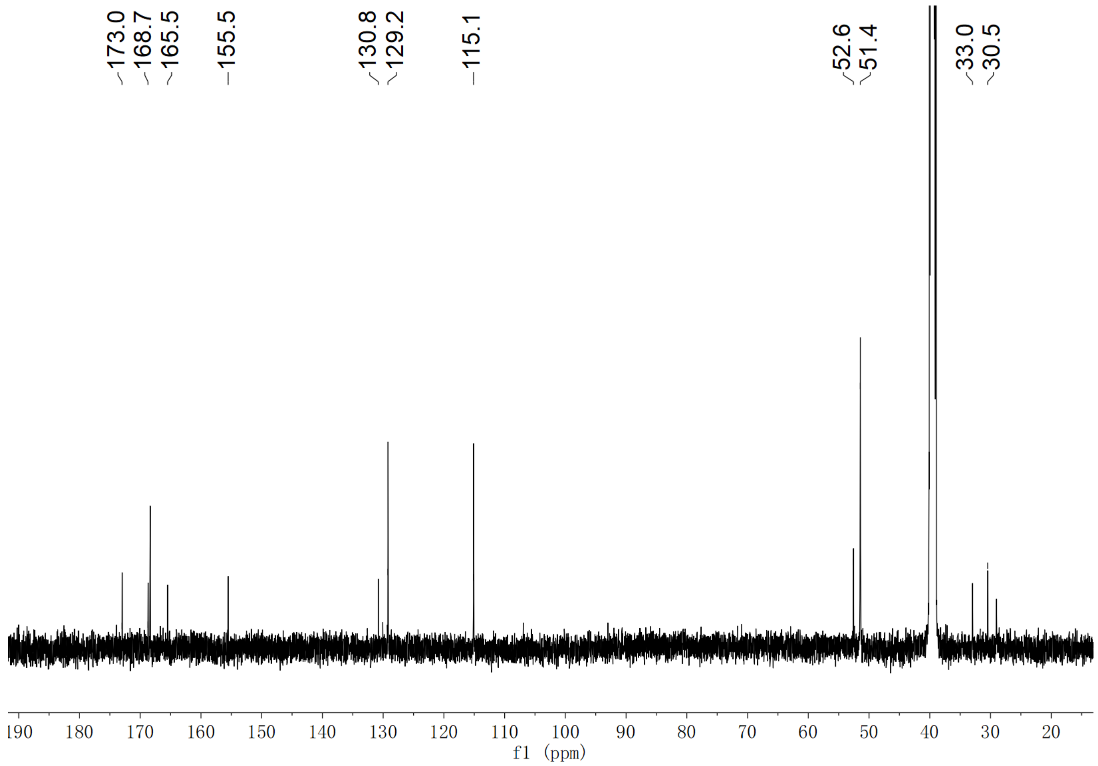

**Supplementary Fig. 195.** **$^{13}$C NMR spectrum of compound (2*S*,3*S*)-*t*-ES-a15 in DMSO-*d*$_6$**

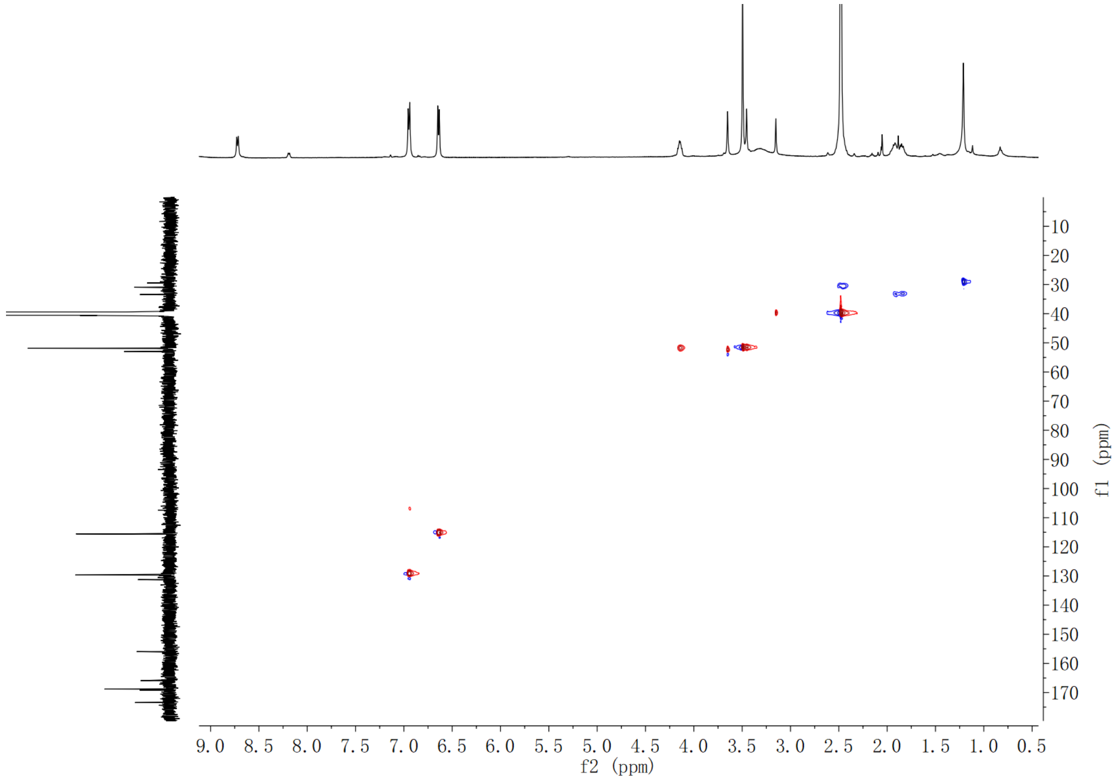

**Supplementary Fig. 196. HSQC spectrum of compound (2*S*,3*S*)-*t*-ES-a15 in DMSO-*d*$_6$**

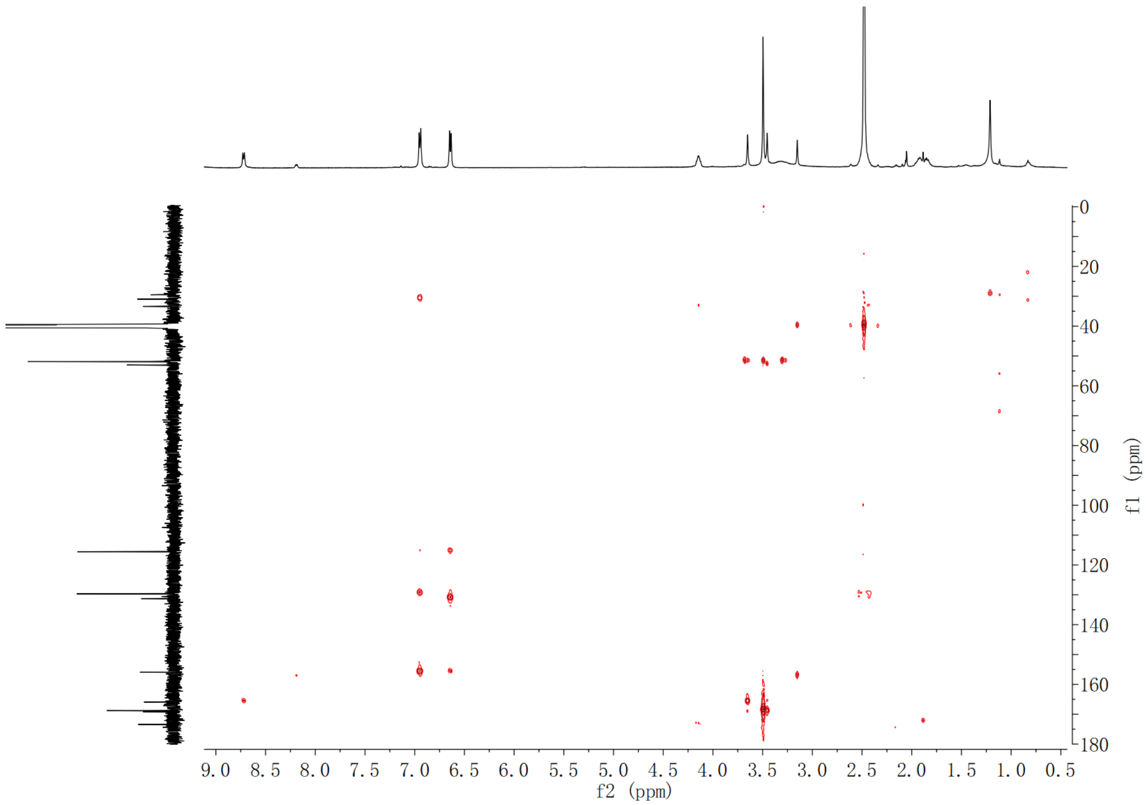

**Supplementary Fig. 197. HMBC spectrum of compound (2*S*,3*S*)-*t*-ES-a15 in DMSO-$d_6$**

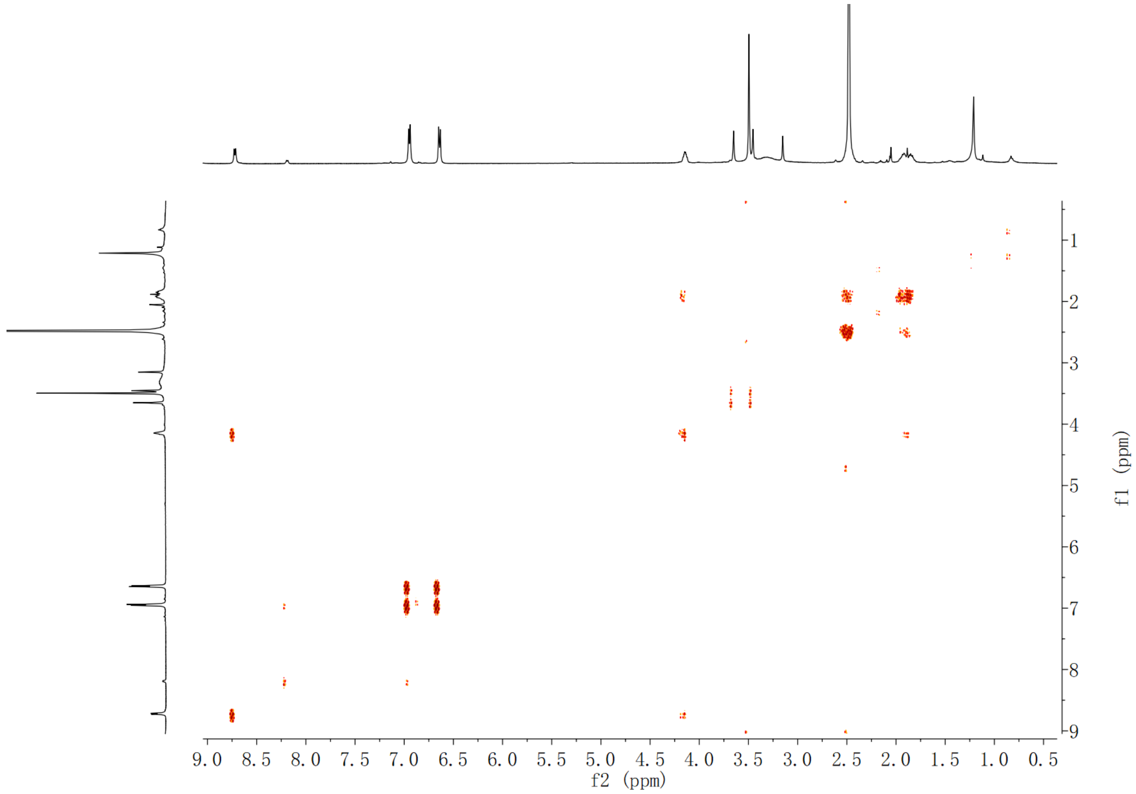

**Supplementary Fig. 198. $^1$H-$^1$H COSY spectrum of compound (2*S*,3*S*)-*t*-ES-a15 in DMSO-$d_6$**

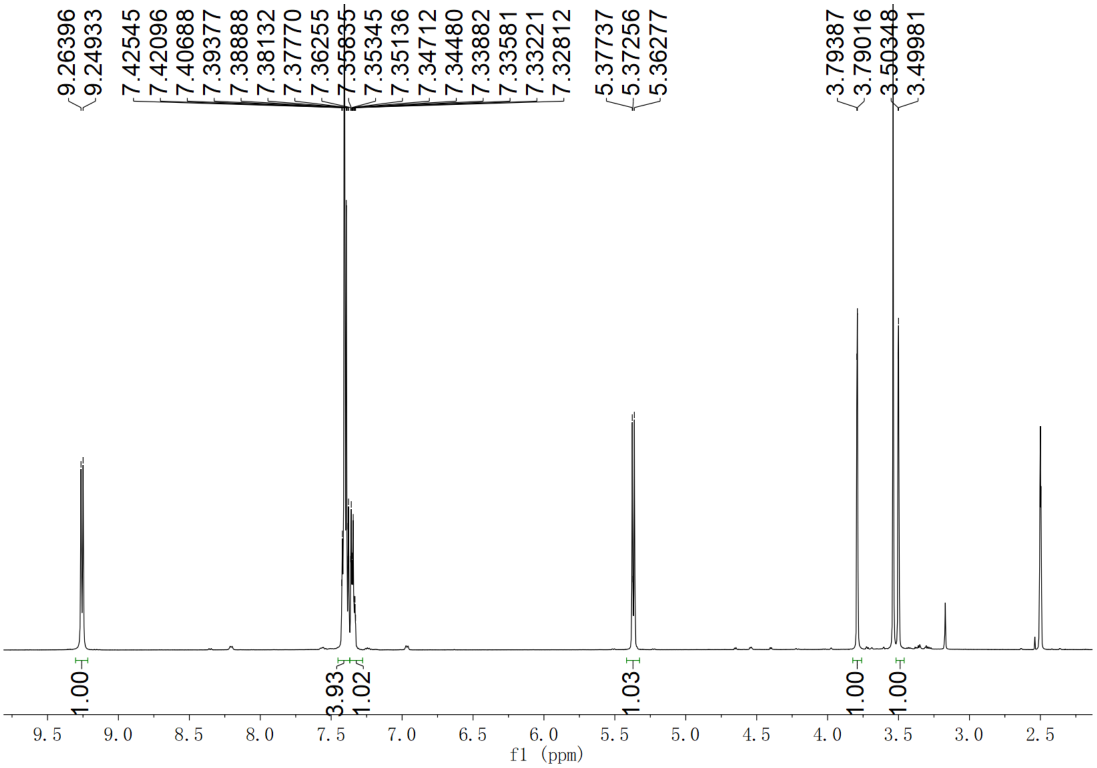

**Supplementary Fig. 199.** $^{1}$**H NMR spectrum of compound (2*S*,3*S*)-*t*-ES-a16 in DMSO-*d*$_6$**

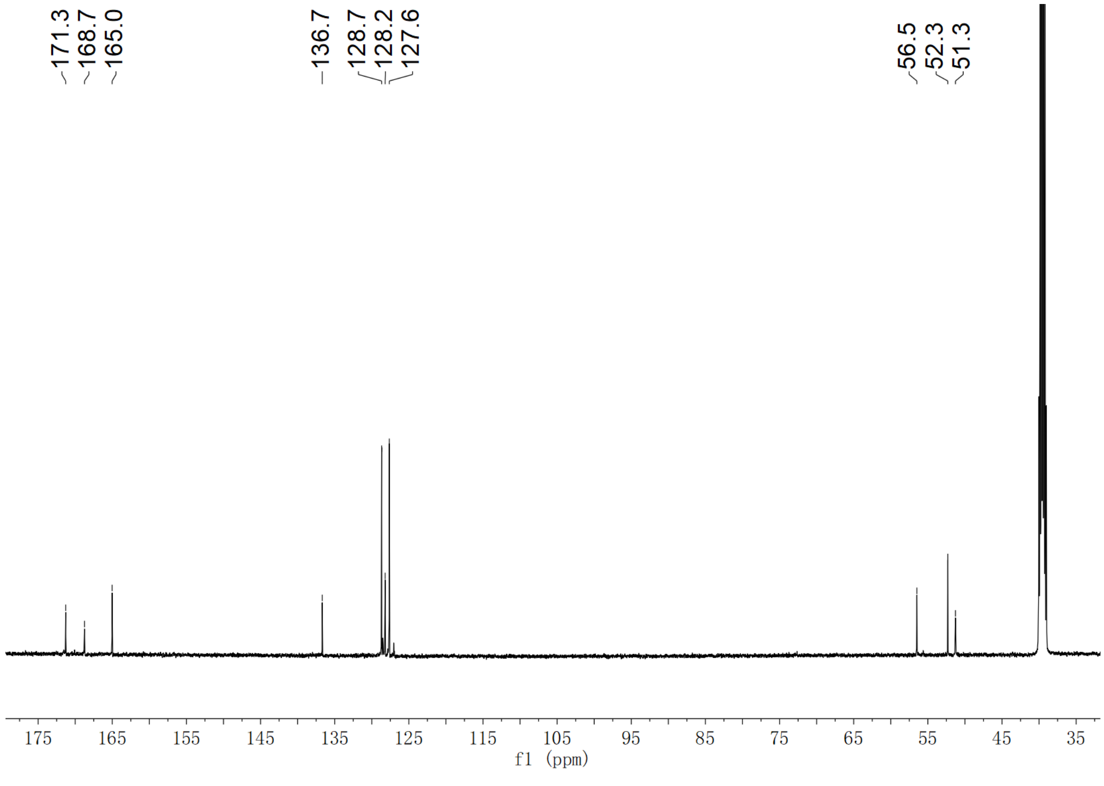

**Supplementary Fig. 200.** $^{13}$**C NMR spectrum of compound (2*S*,3*S*)-*t*-ES-a16 in DMSO-*d*$_6$**

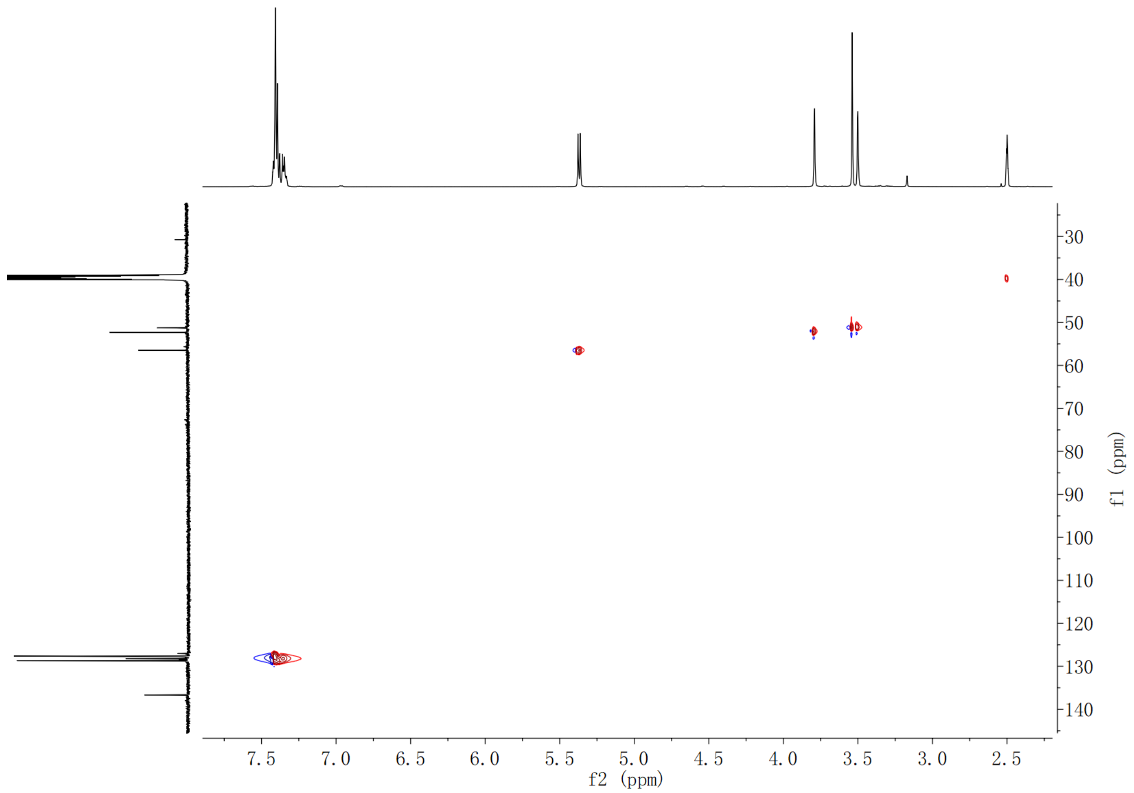

Supplementary Fig. 201. HSQC spectrum of compound (2*S*,3*S*)-*t*-ES-a16 in DMSO-$d_6$

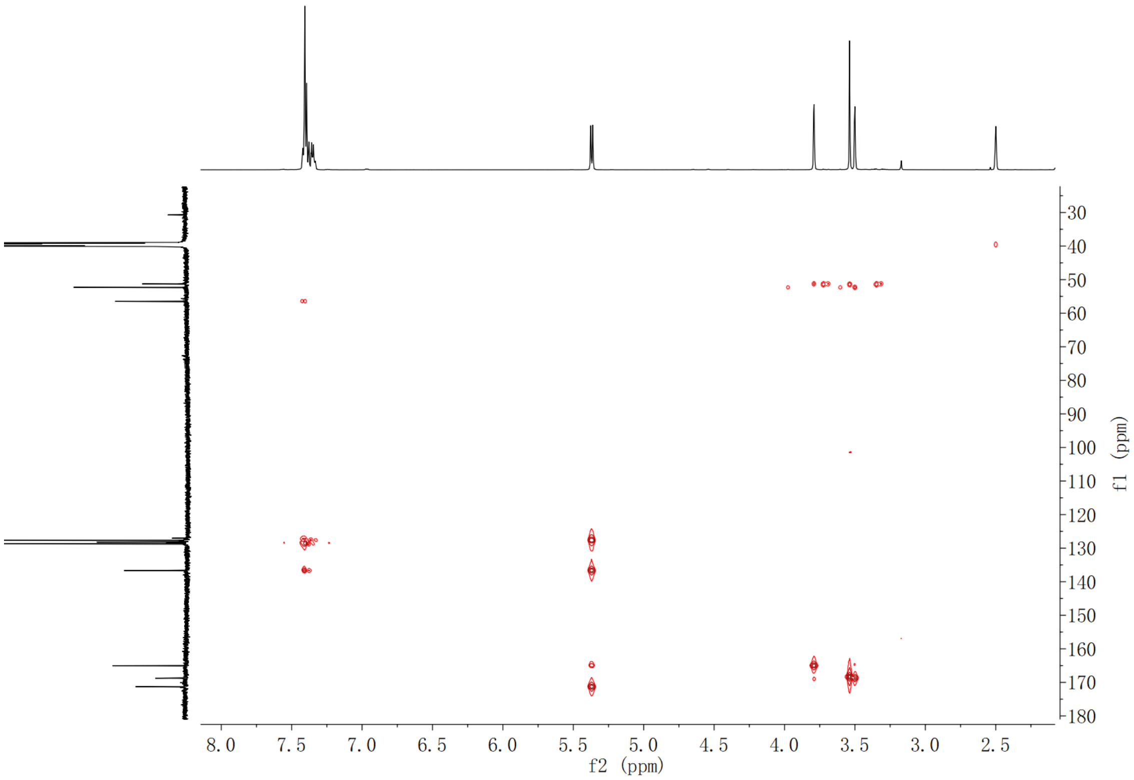

Supplementary Fig. 202. HMBC spectrum of compound (2*S*,3*S*)-*t*-ES-a16 in DMSO-$d_6$

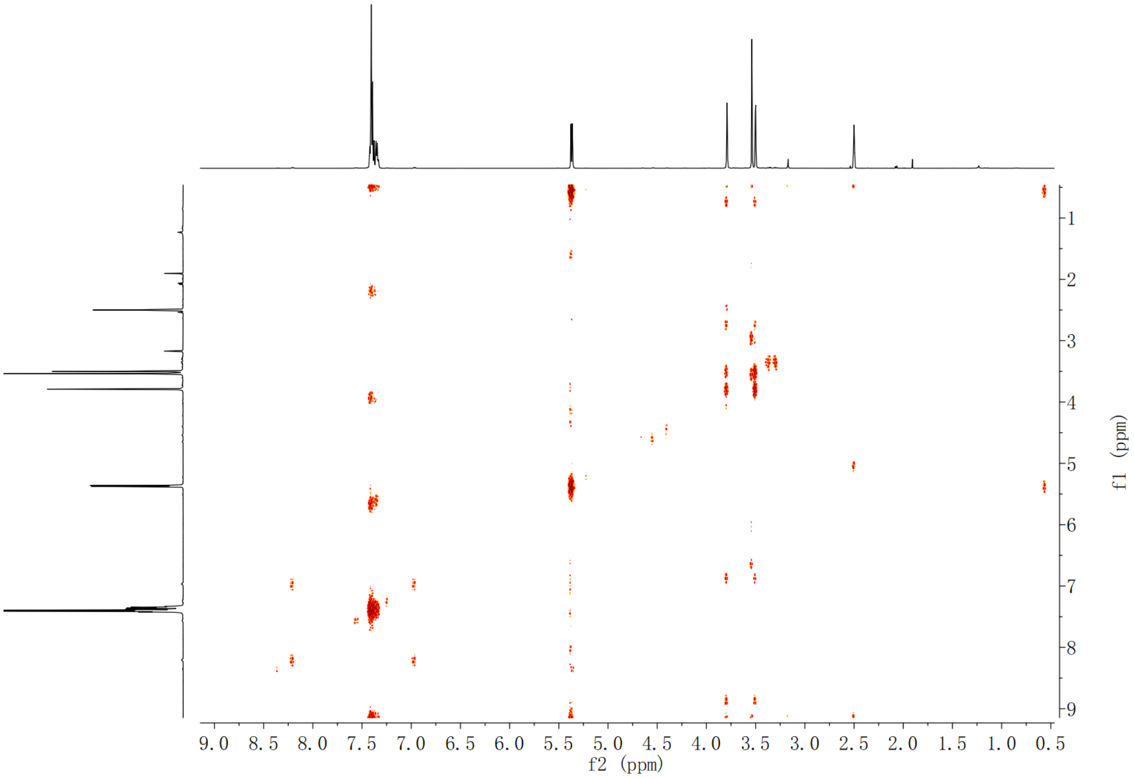

**Supplementary Fig. 203.** $^{1}$**H-**$^{1}$**H COSY spectrum of compound (2*S*,3*S*)-*t*-ES-a16 in DMSO-*d*$_6$**

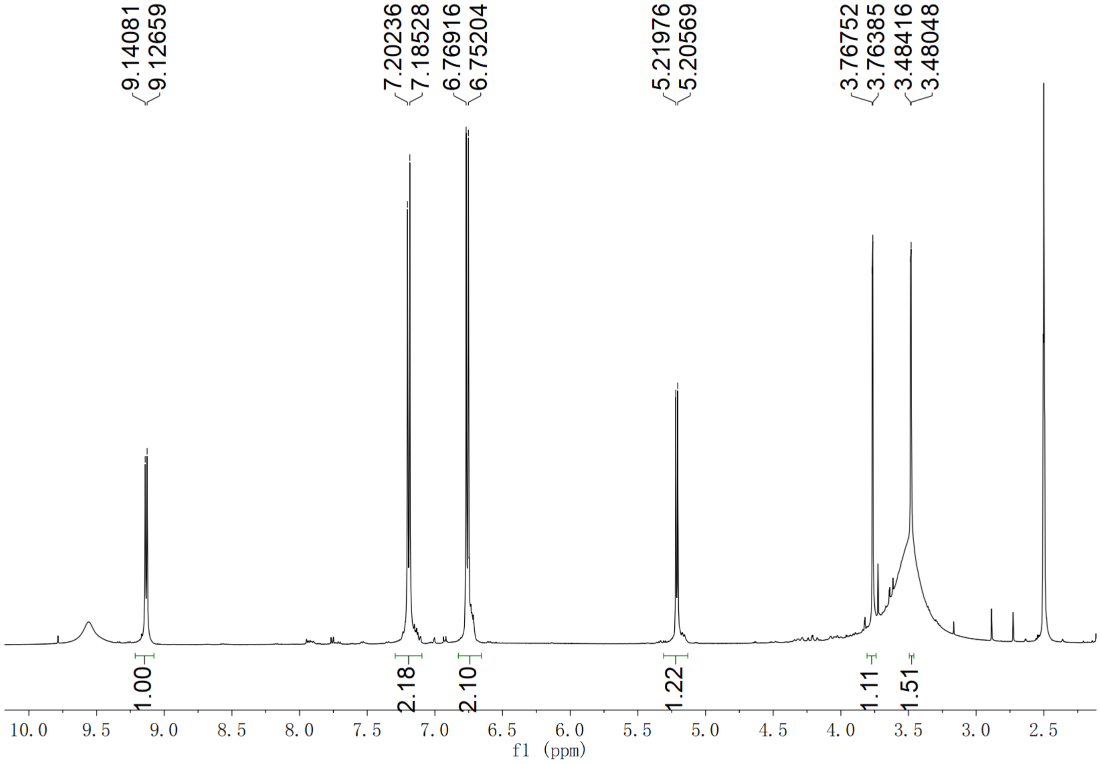

**Supplementary Fig. 204.** $^{1}$**H NMR spectrum of compound (2*S*,3*S*)-*t*-ES-a17 in DMSO-*d*$_6$**

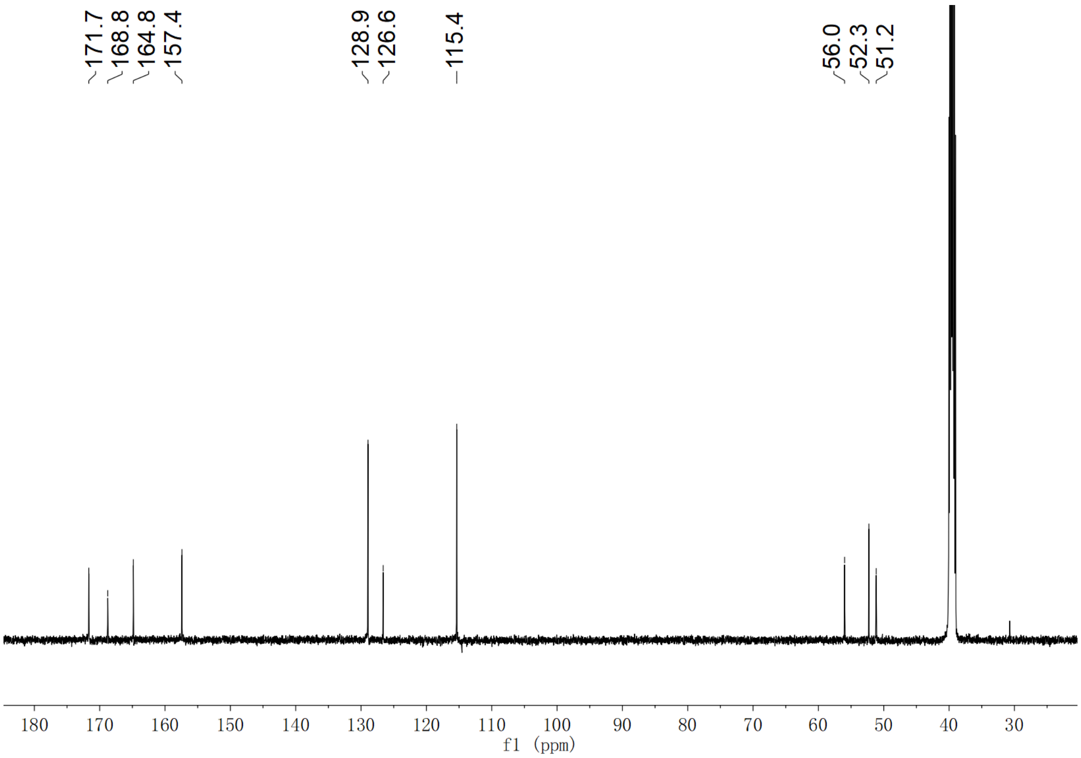

**Supplementary Fig. 205.** $^{13}$**C NMR spectrum of compound (2*S*,3*S*)-*t*-ES-a17 in DMSO-*d*$_6$**

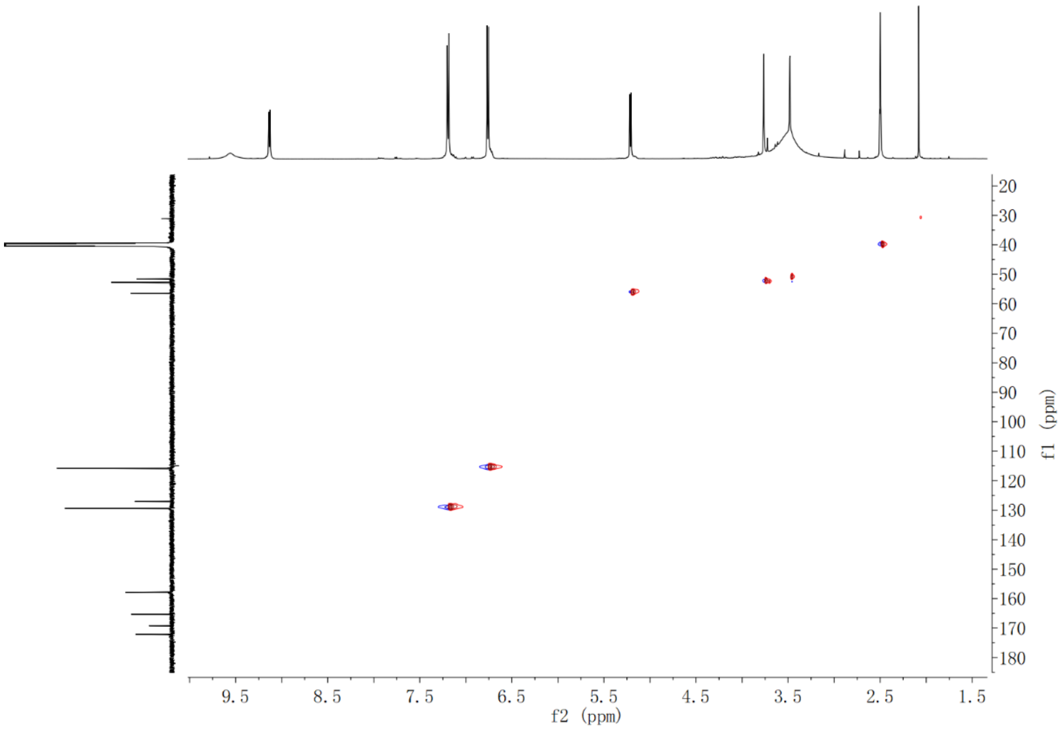

**Supplementary Fig. 206. HSQC spectrum of compound (2*S*,3*S*)-*t*-ES-a17 in DMSO-*d*$_6$**

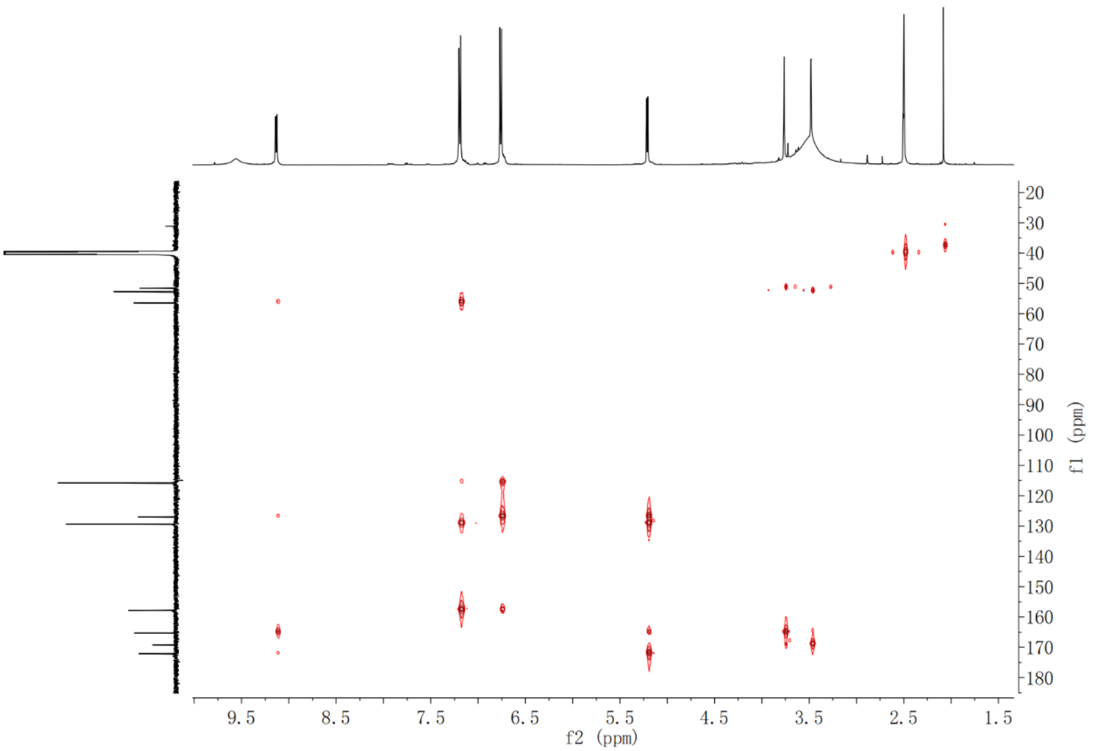

**Supplementary Fig. 207. HMBC spectrum of compound (2*S*,3*S*)-*t*-ES-a17 in DMSO-$d_6$**

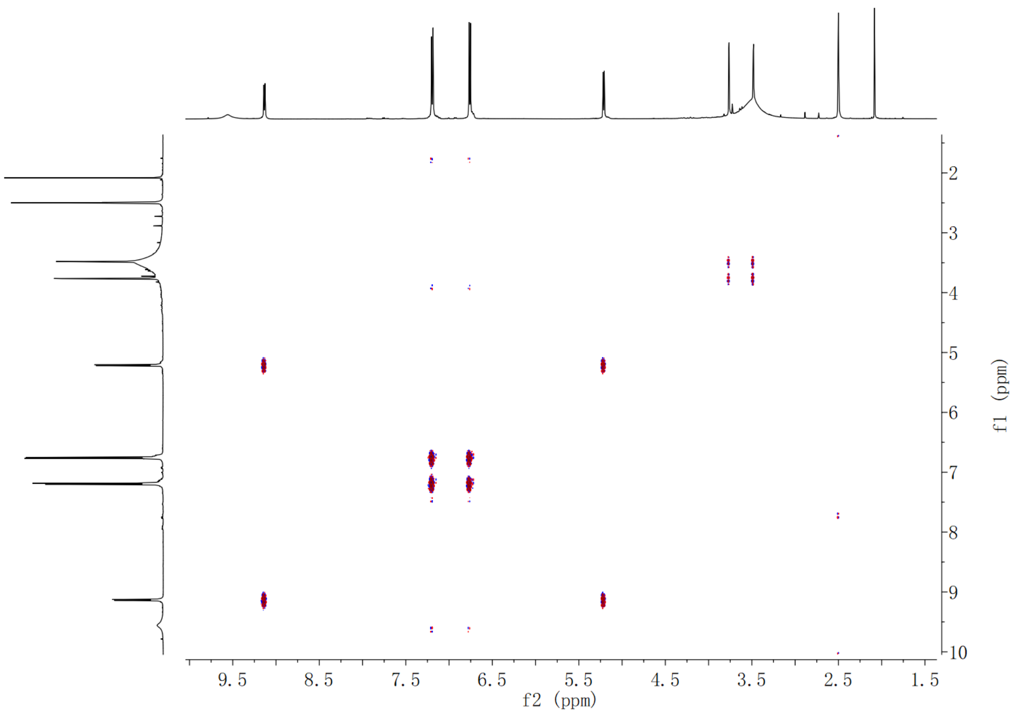

**Supplementary Fig. 208. $^1$H-$^1$H COSY spectrum of compound (2*S*,3*S*)-*t*-ES-a17 in DMSO-$d_6$**

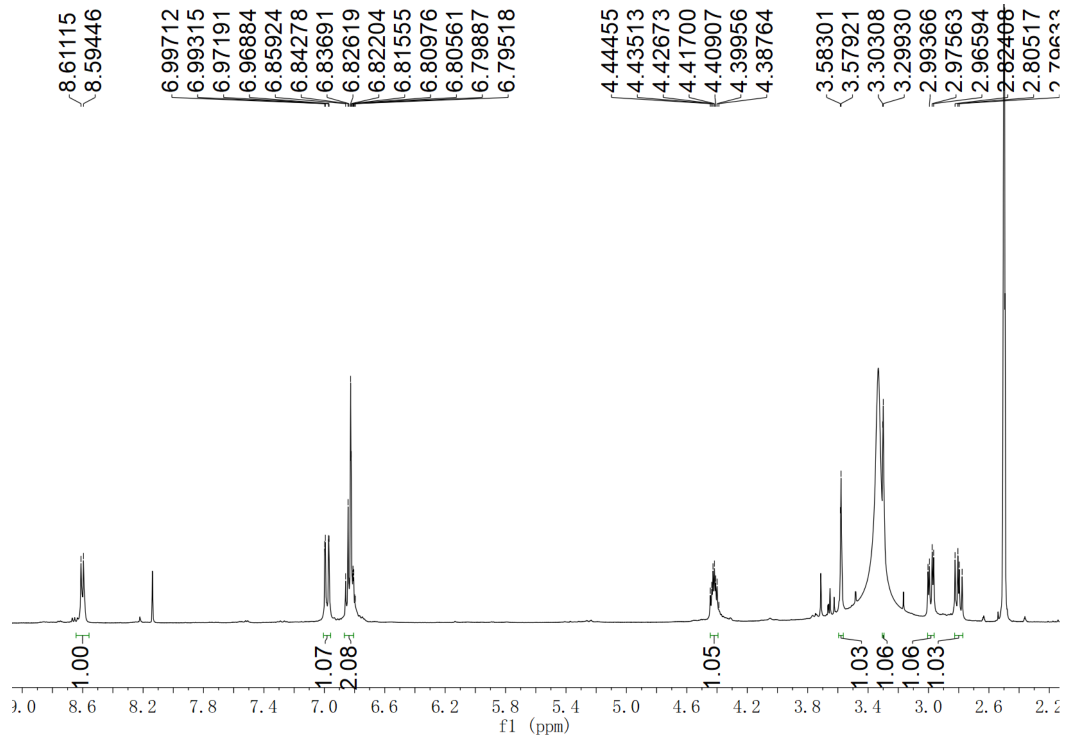

**Supplementary Fig. 209.** **[1]H NMR spectrum of compound (2*S*,3*S*)-*t*-ES-a18 in DMSO-$d_6$**

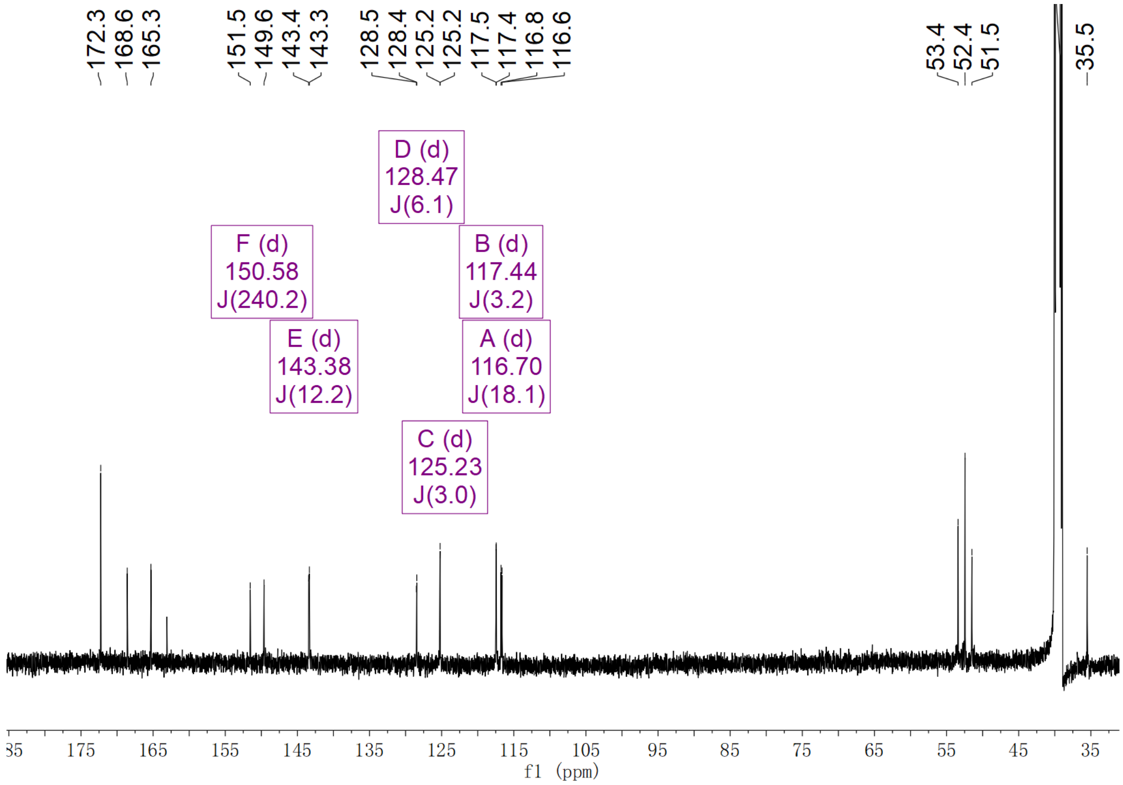

**Supplementary Fig. 210.** **[13]C NMR spectrum of compound (2*S*,3*S*)-*t*-ES-a18 in DMSO-$d_6$**

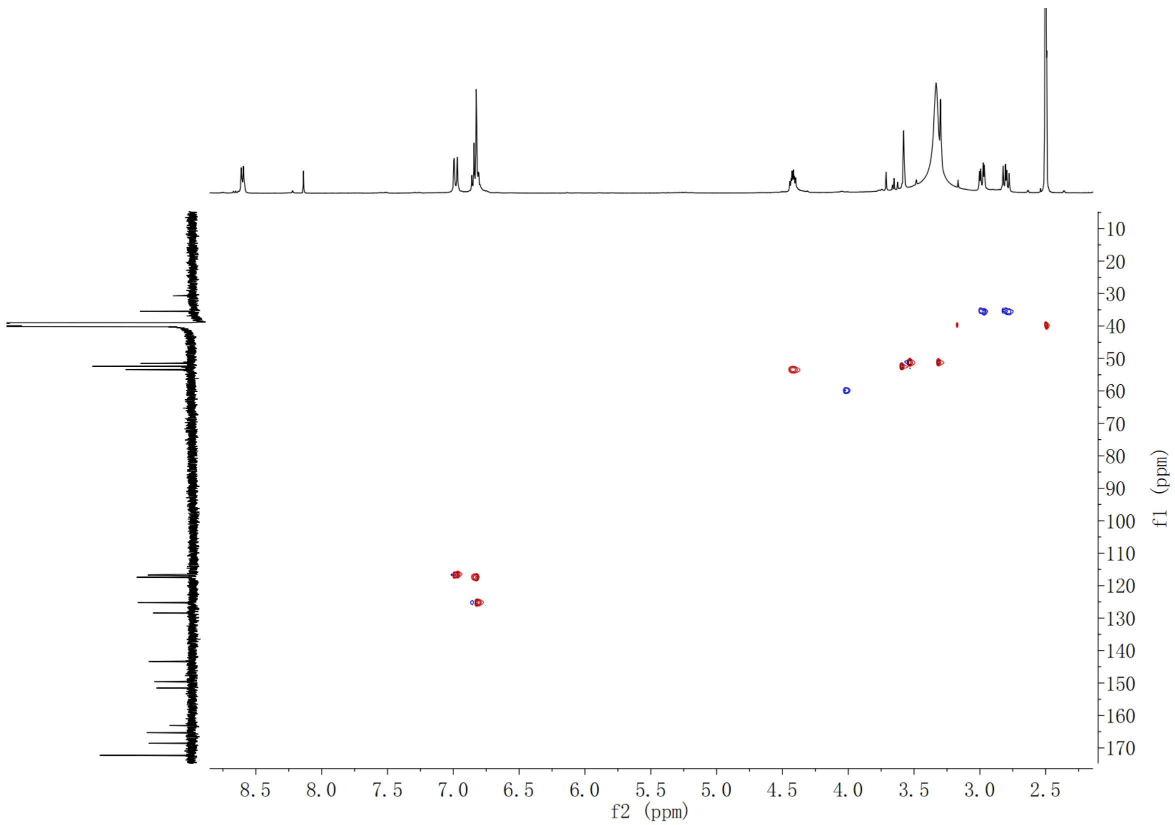

**Supplementary Fig. 211. HSQC spectrum of compound (2*S*,3*S*)-*t*-ES-a18 in DMSO-*d*$_6$**

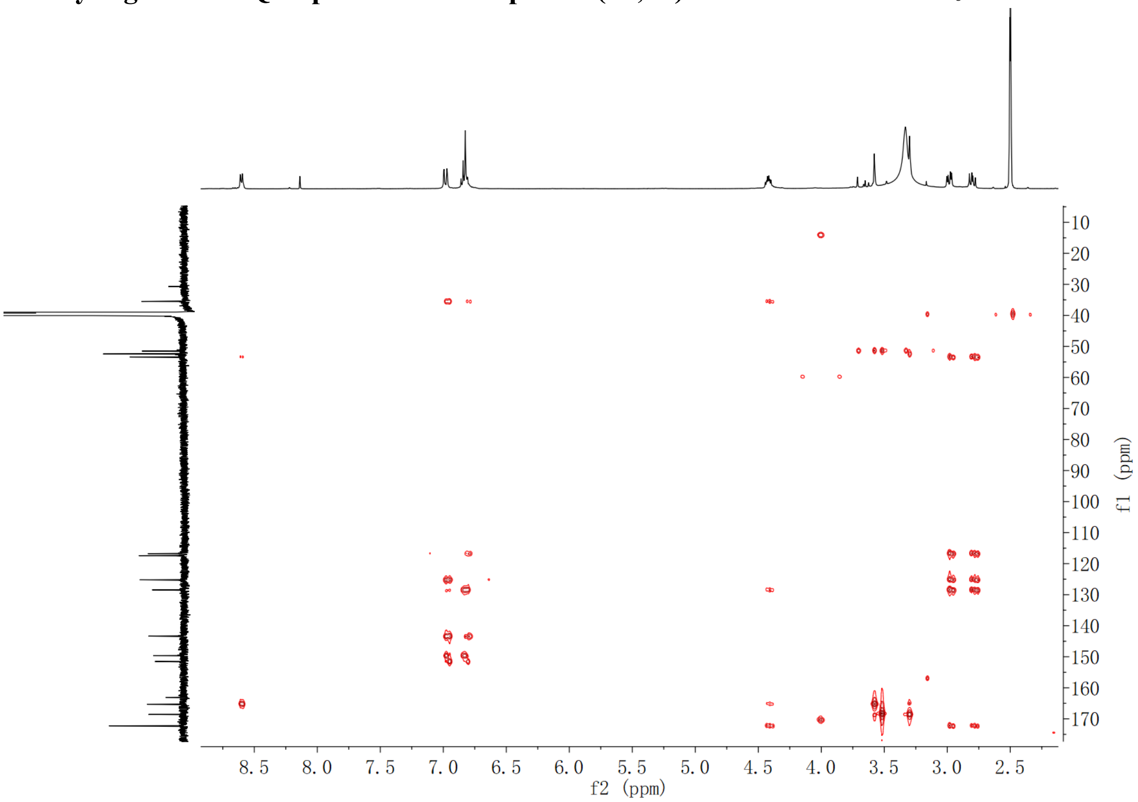

**Supplementary Fig. 212. HMBC spectrum of compound (2*S*,3*S*)-*t*-ES-a18 in DMSO-*d*$_6$**

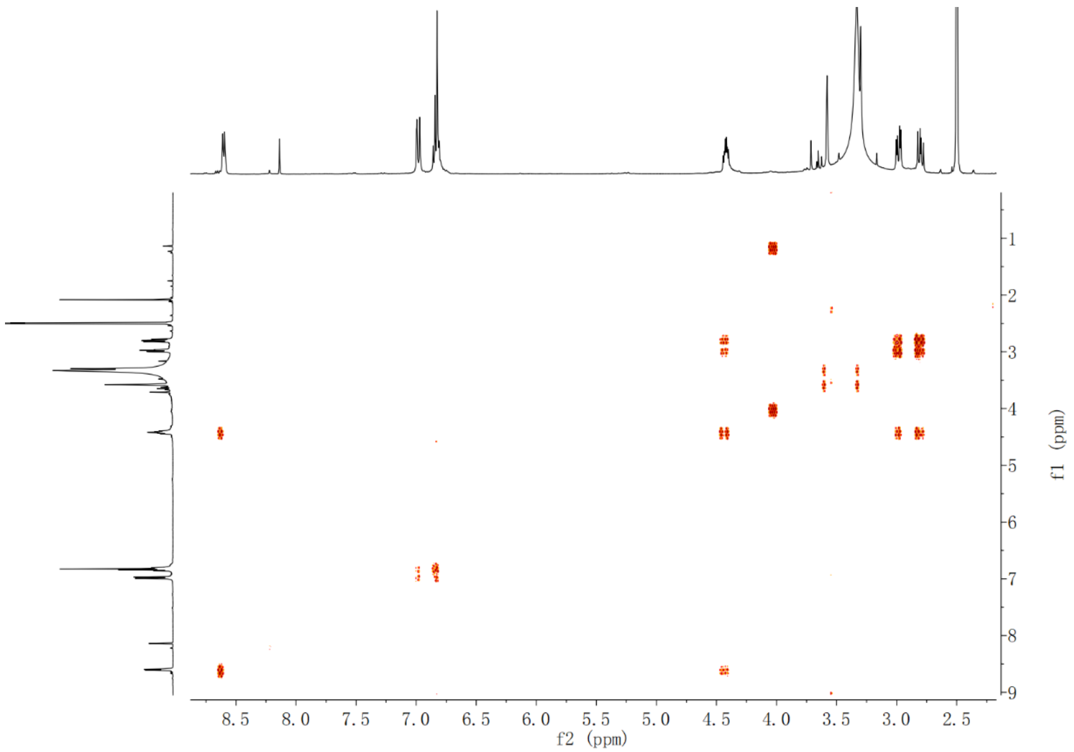

Supplementary Fig. 213. $^{1}$H-$^{1}$H COSY spectrum of compound (2*S*,3*S*)-*t*-ES-a18 in DMSO-$d_6$

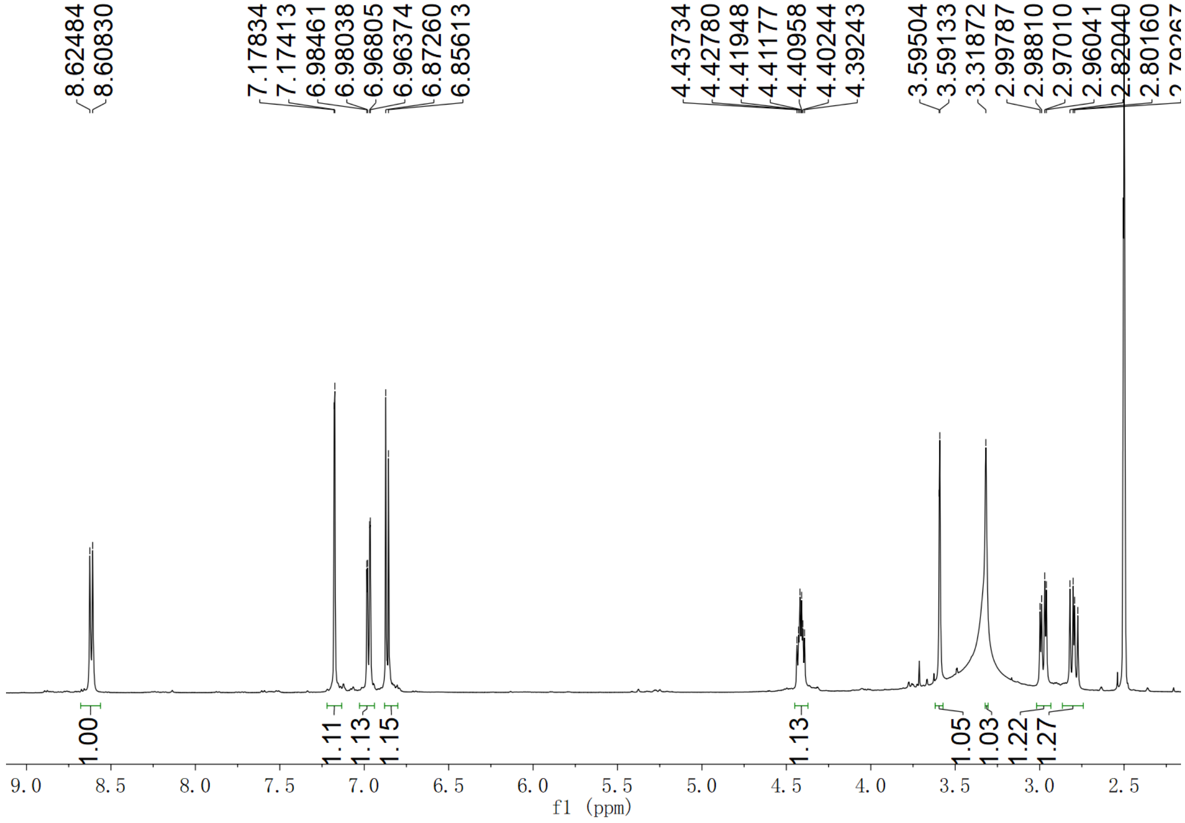

Supplementary Fig. 214. $^{1}$H NMR spectrum of compound (2*S*,3*S*)-*t*-ES-a19 in DMSO-$d_6$

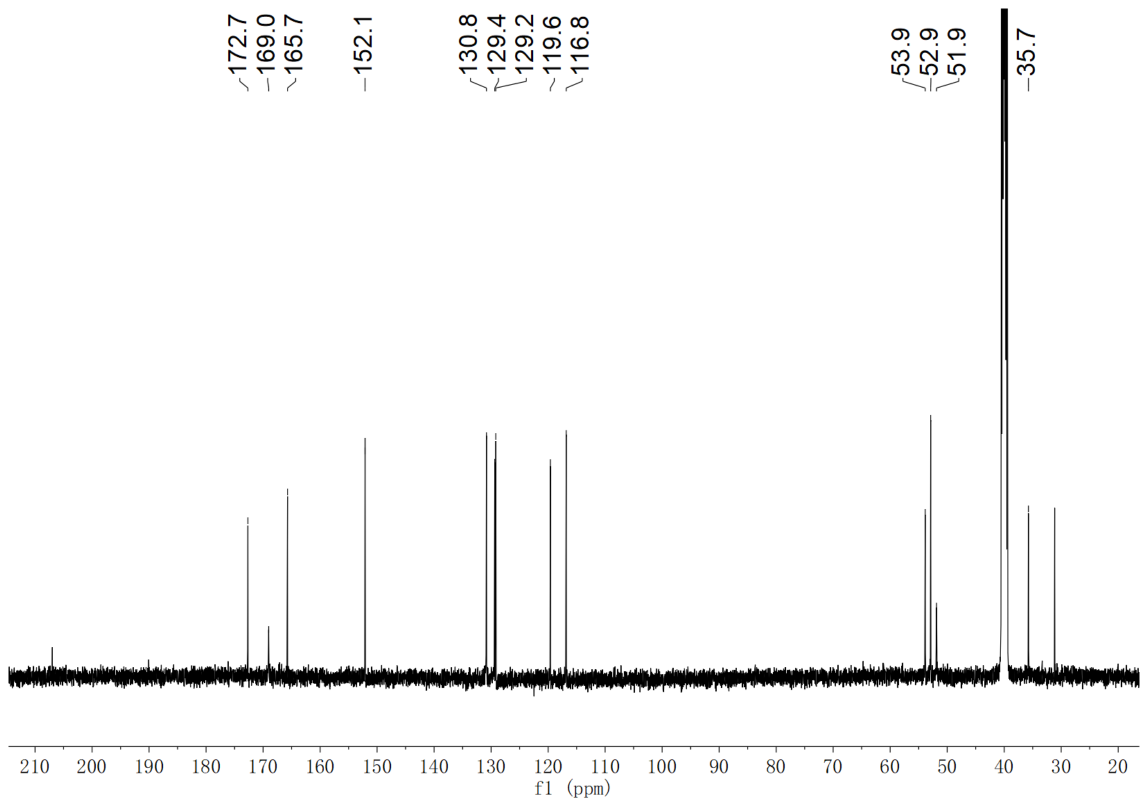

**Supplementary Fig. 215. $^{13}$C NMR spectrum of compound (2*S*,3*S*)-*t*-ES-a19 in DMSO-$d_6$**

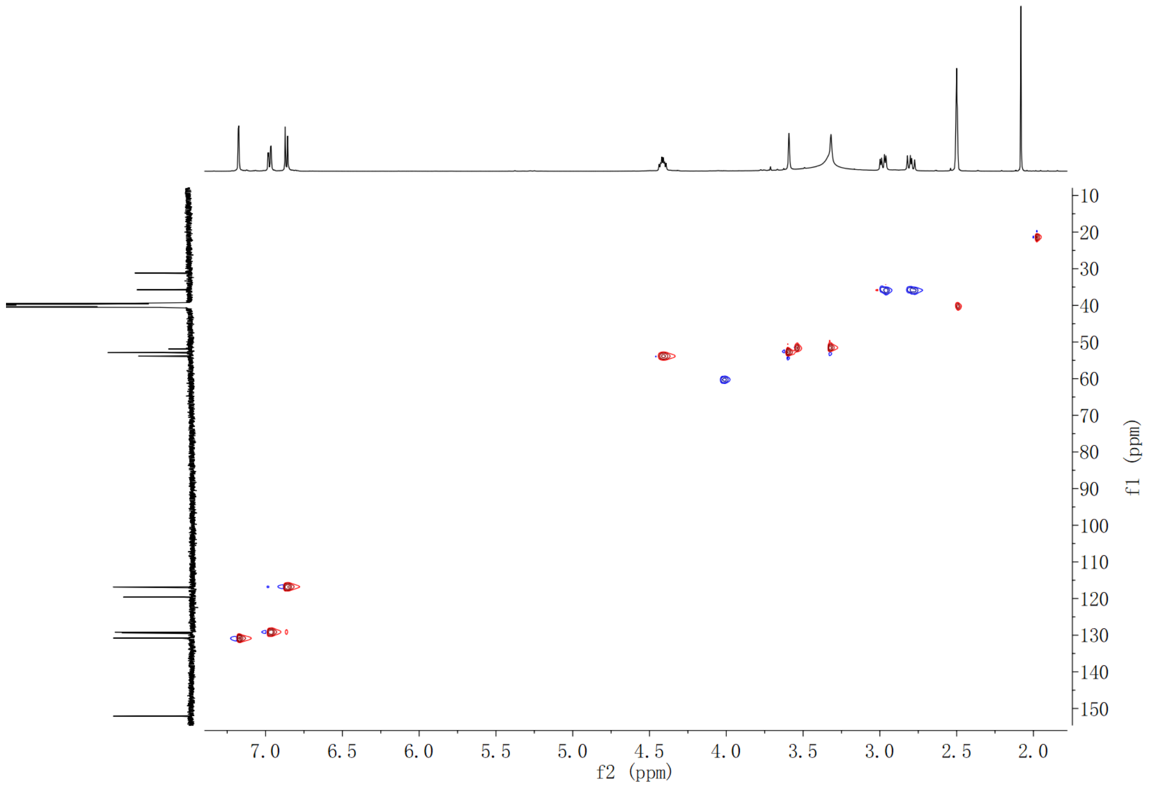

**Supplementary Fig. 216. HSQC spectrum of compound (2*S*,3*S*)-*t*-ES-a19 in DMSO-$d_6$**

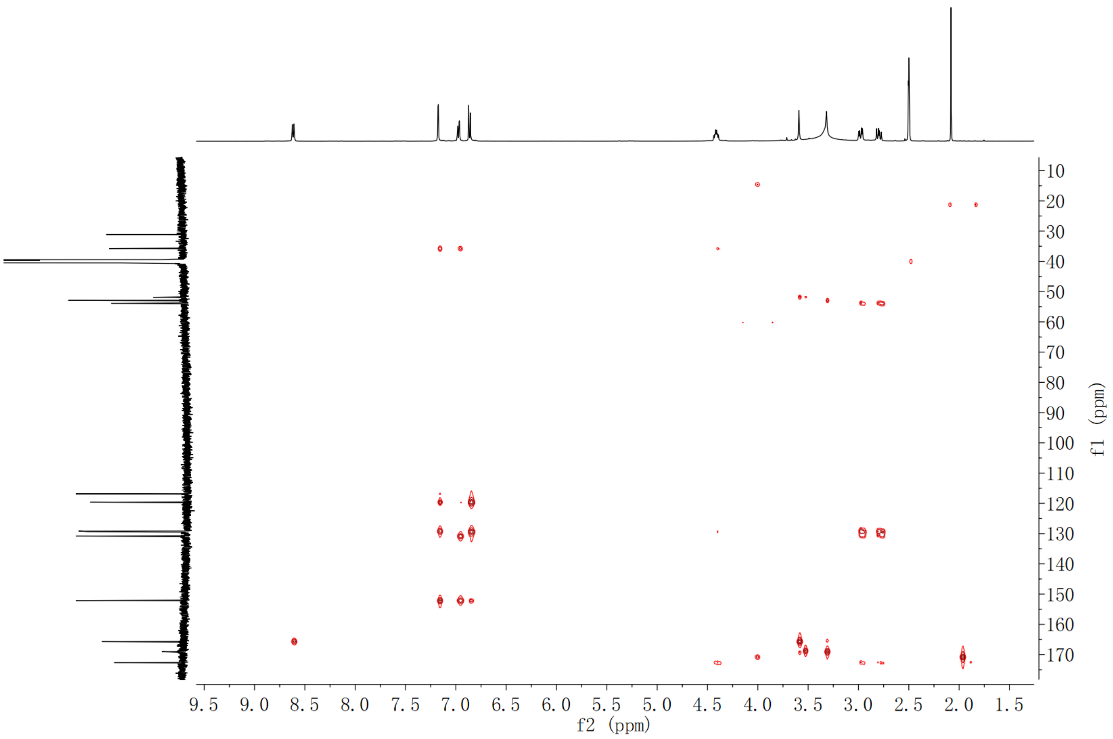

**Supplementary Fig. 217. HMBC spectrum of compound (2*S*,3*S*)-*t*-ES-a19 in DMSO-$d_6$**

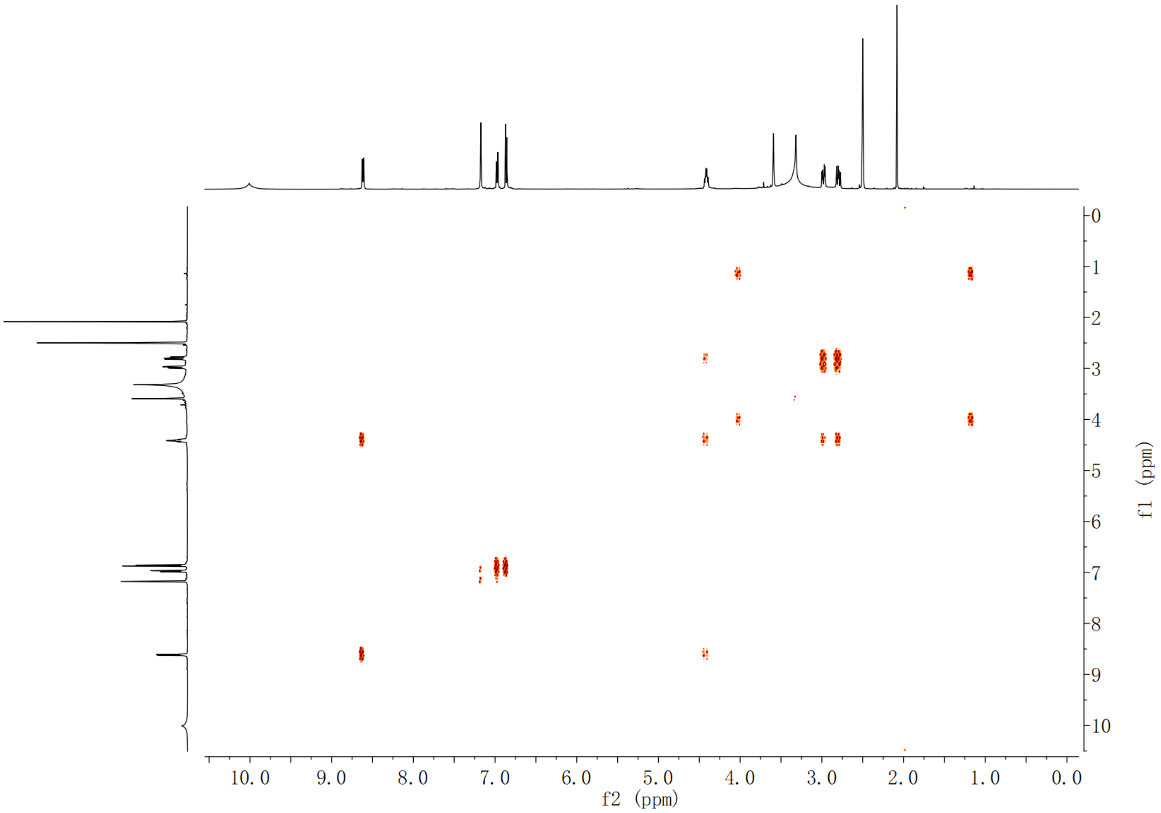

**Supplementary Fig. 218. $^1$H-$^1$H COSY spectrum of compound (2*S*,3*S*)-*t*-ES-a19 in DMSO-$d_6$**

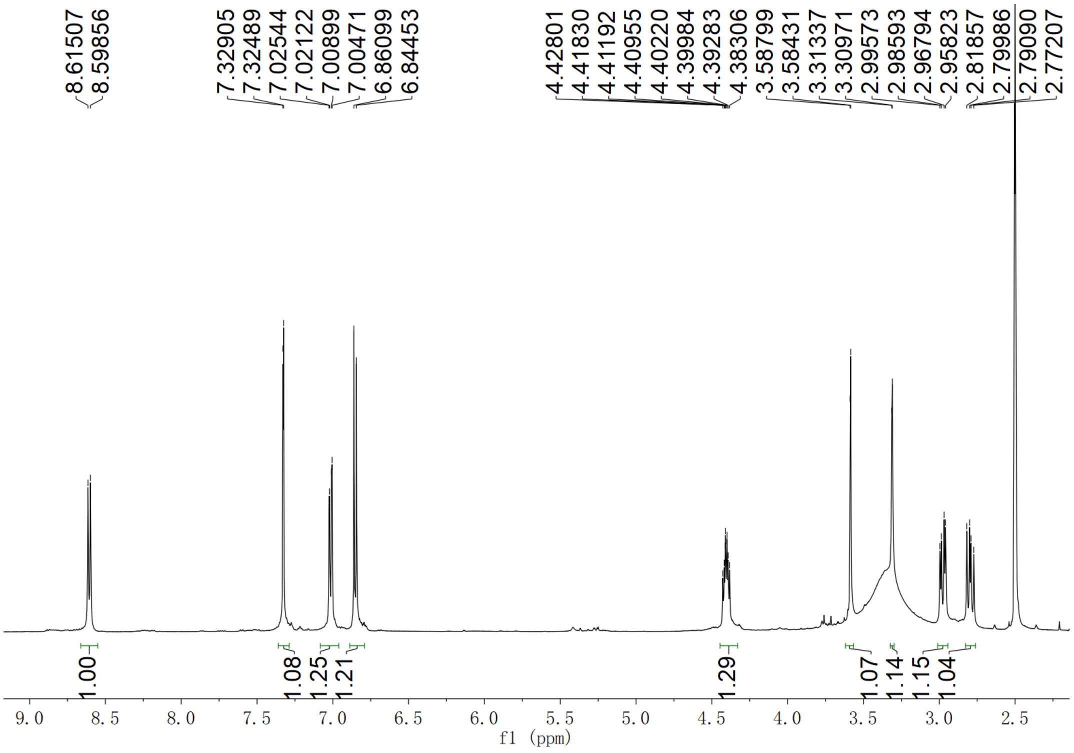

**Supplementary Fig. 219.** **$^{1}$H NMR spectrum of compound (2*S*,3*S*)-*t*-ES-a20 in DMSO-$d_6$**

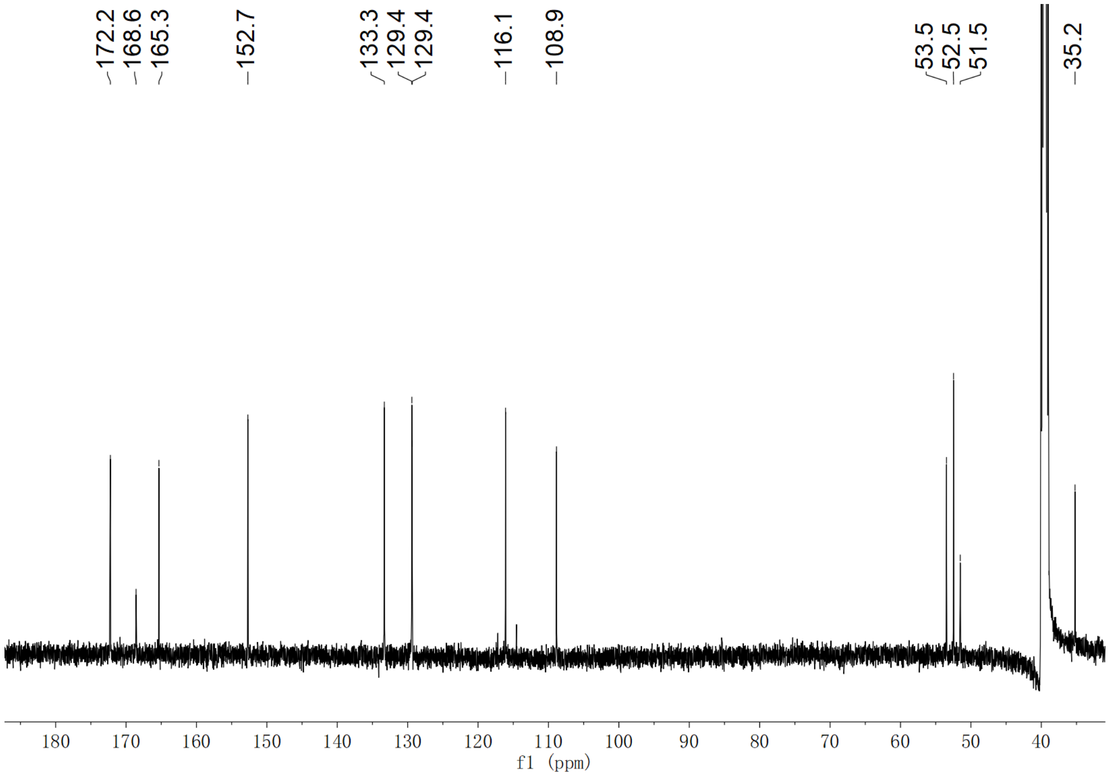

**Supplementary Fig. 220.** **$^{13}$C NMR spectrum of compound (2*S*,3*S*)-*t*-ES-a20 in DMSO-$d_6$**

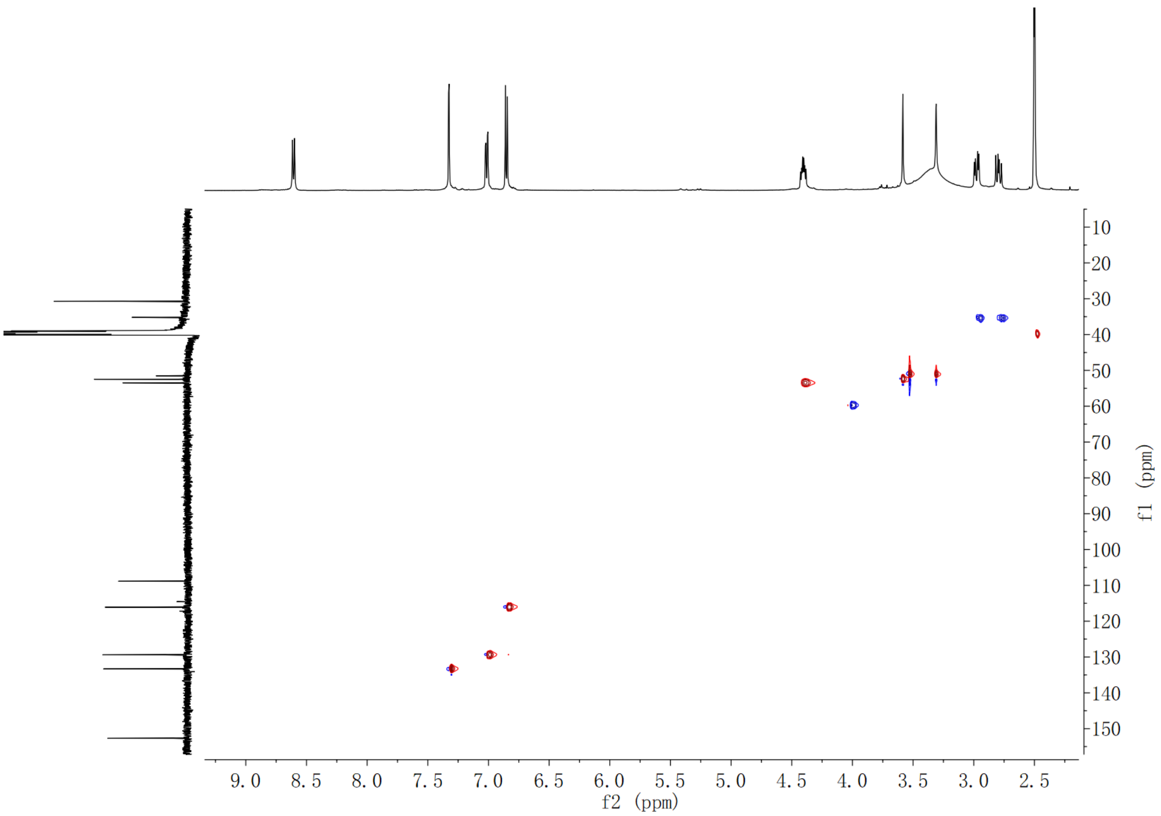

**Supplementary Fig. 221. HSQC spectrum of compound (2*S*,3*S*)-*t*-ES-a20 in DMSO-$d_6$**

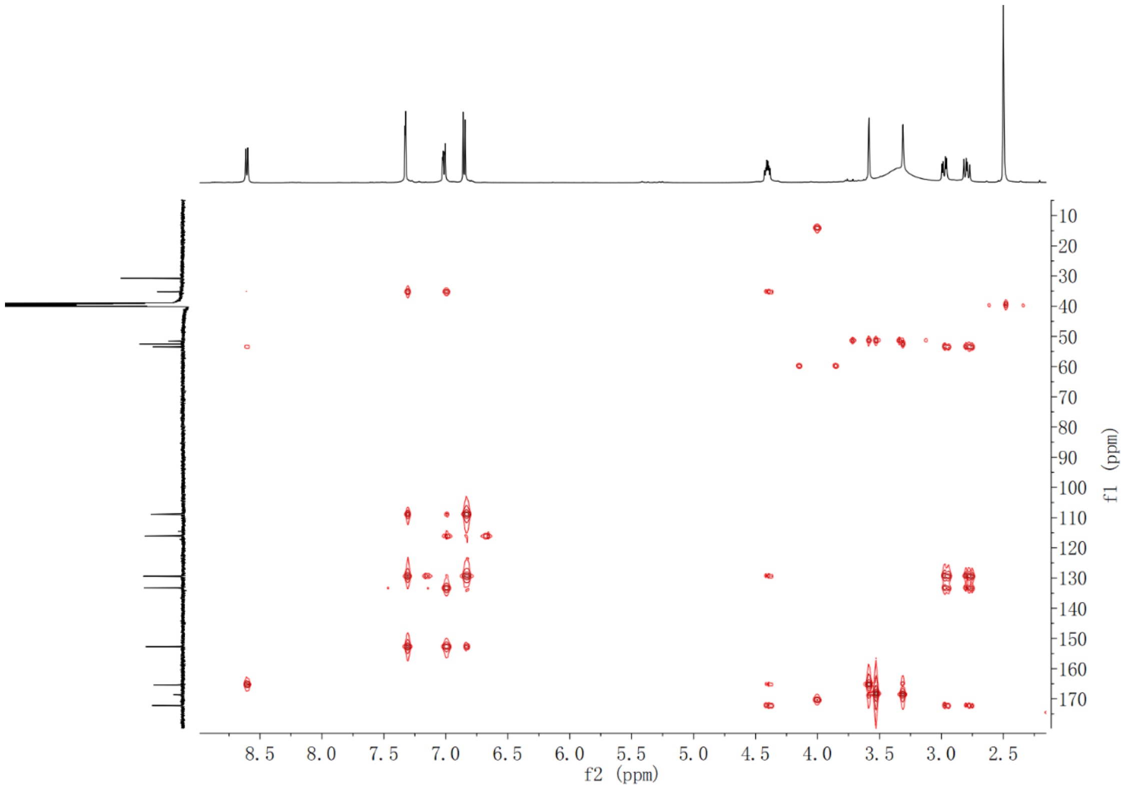

**Supplementary Fig. 222. HMBC spectrum of compound (2*S*,3*S*)-*t*-ES-a20 in DMSO-$d_6$**

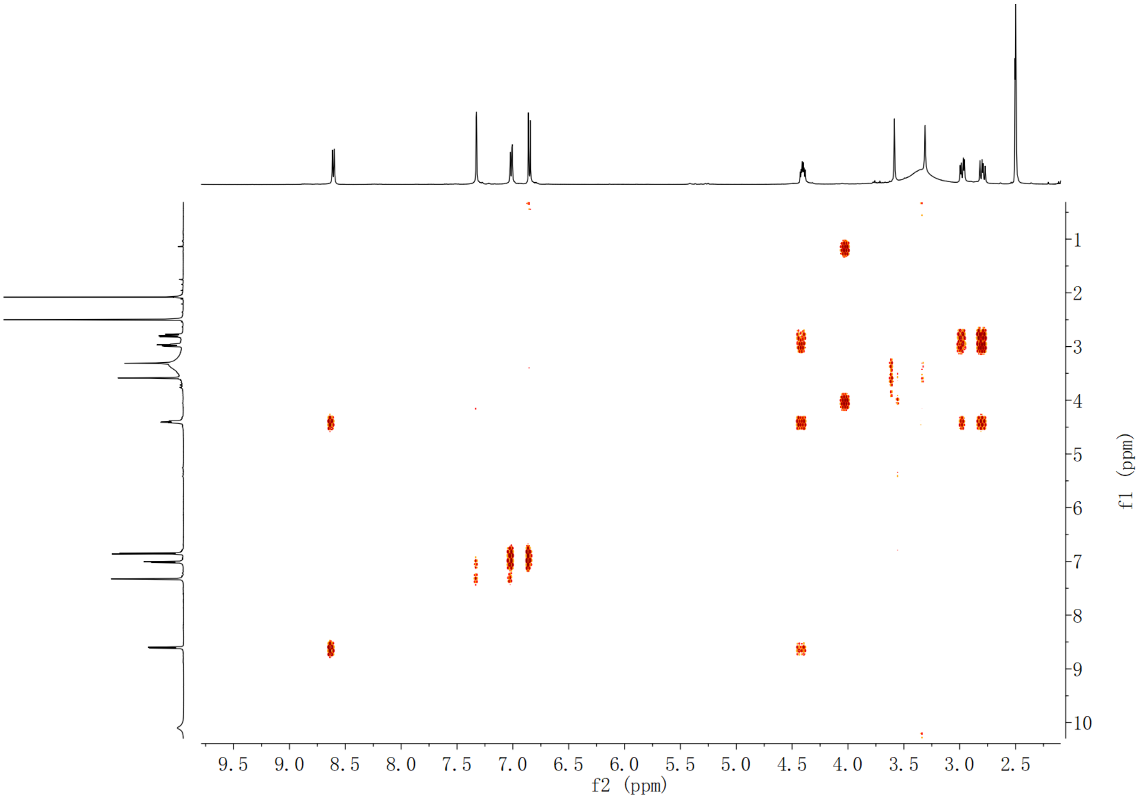

**Supplementary Fig. 223.** **[1]H-[1]H COSY spectrum of compound (2*S*,3*S*)-*t*-ES-a20 in DMSO-$d_6$**

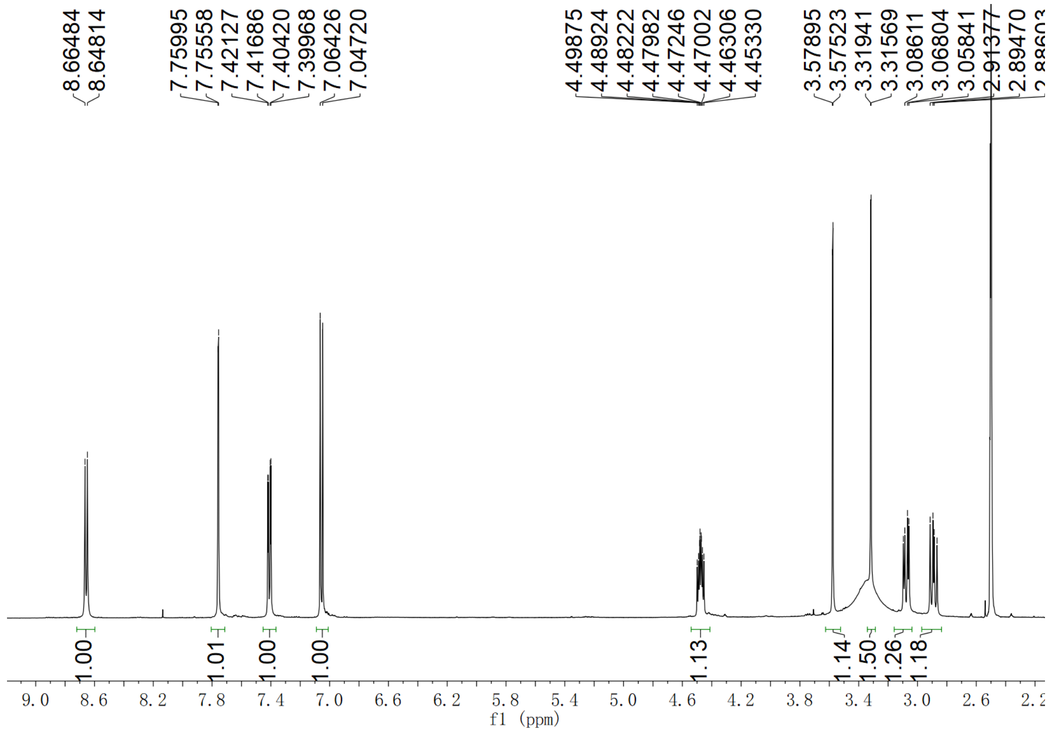

**Supplementary Fig. 224.** **[1]H NMR spectrum of compound (2*S*,3*S*)-*t*-ES-a21 in DMSO-$d_6$**

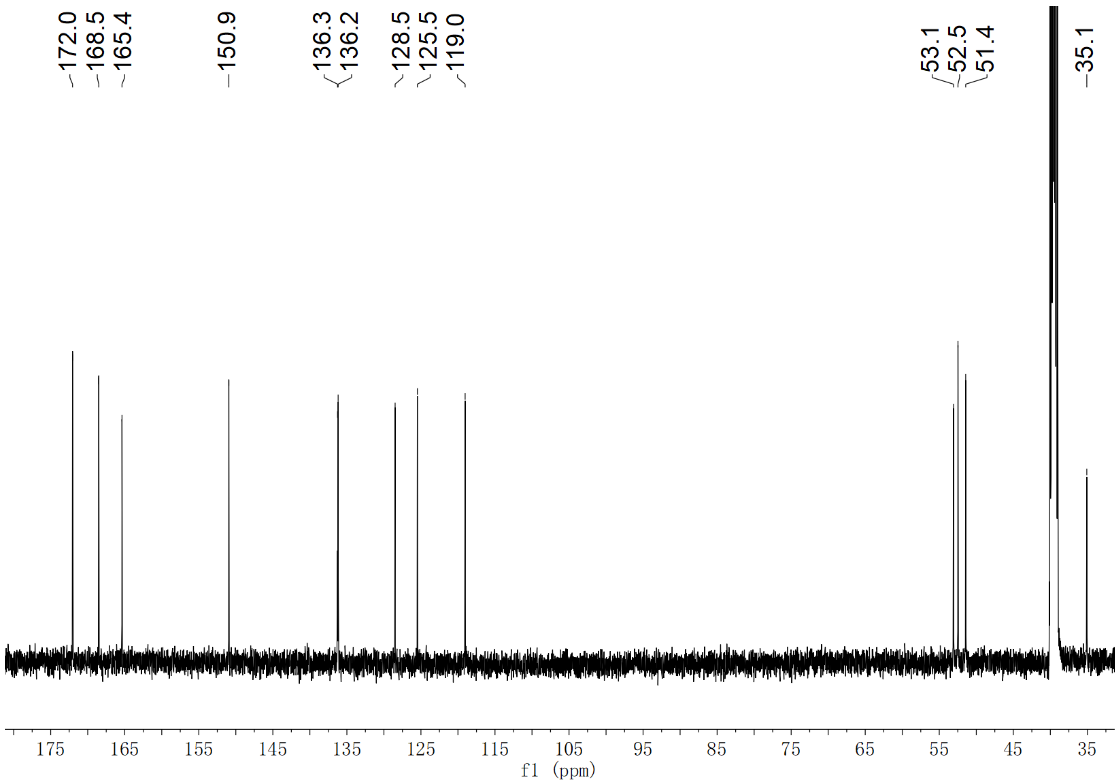

**Supplementary Fig. 225. $^{13}$C NMR spectrum of compound (2*S*,3*S*)-*t*-ES-a21 in DMSO-$d_6$**

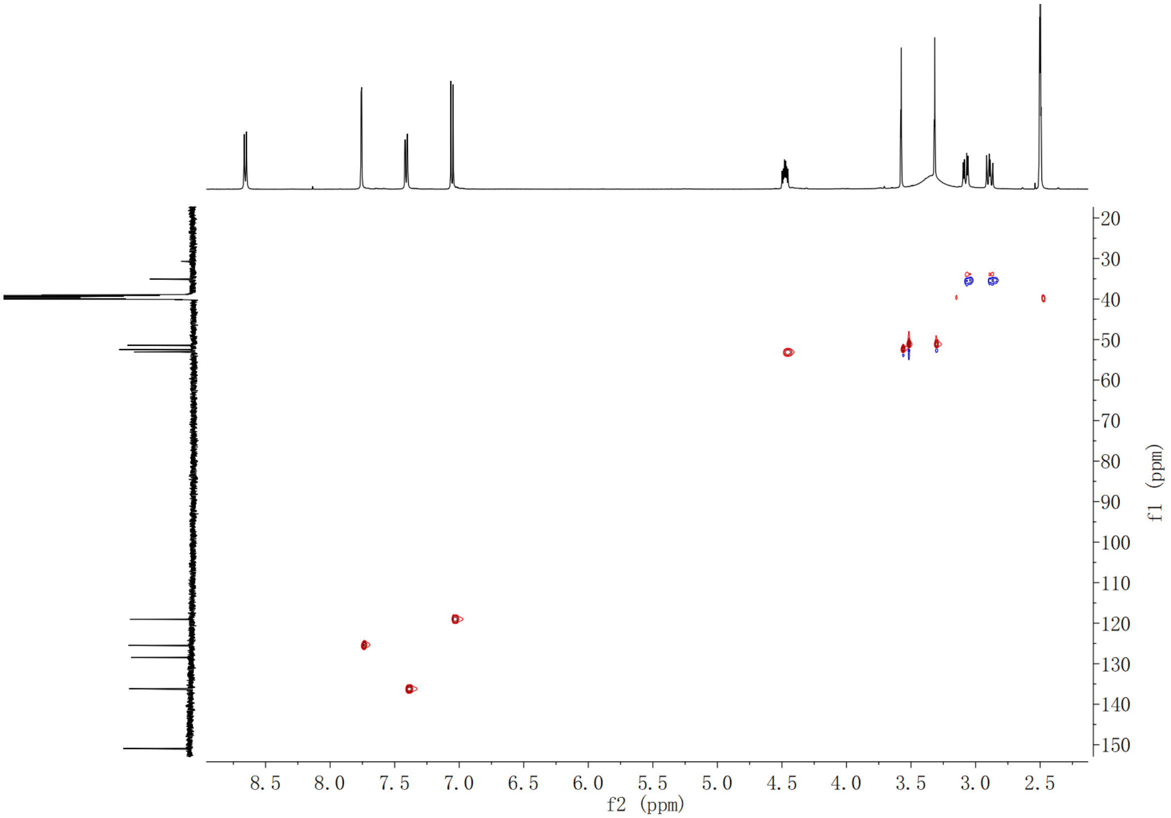

**Supplementary Fig. 226. HSQC spectrum of compound (2*S*,3*S*)-*t*-ES-a21 in DMSO-$d_6$**

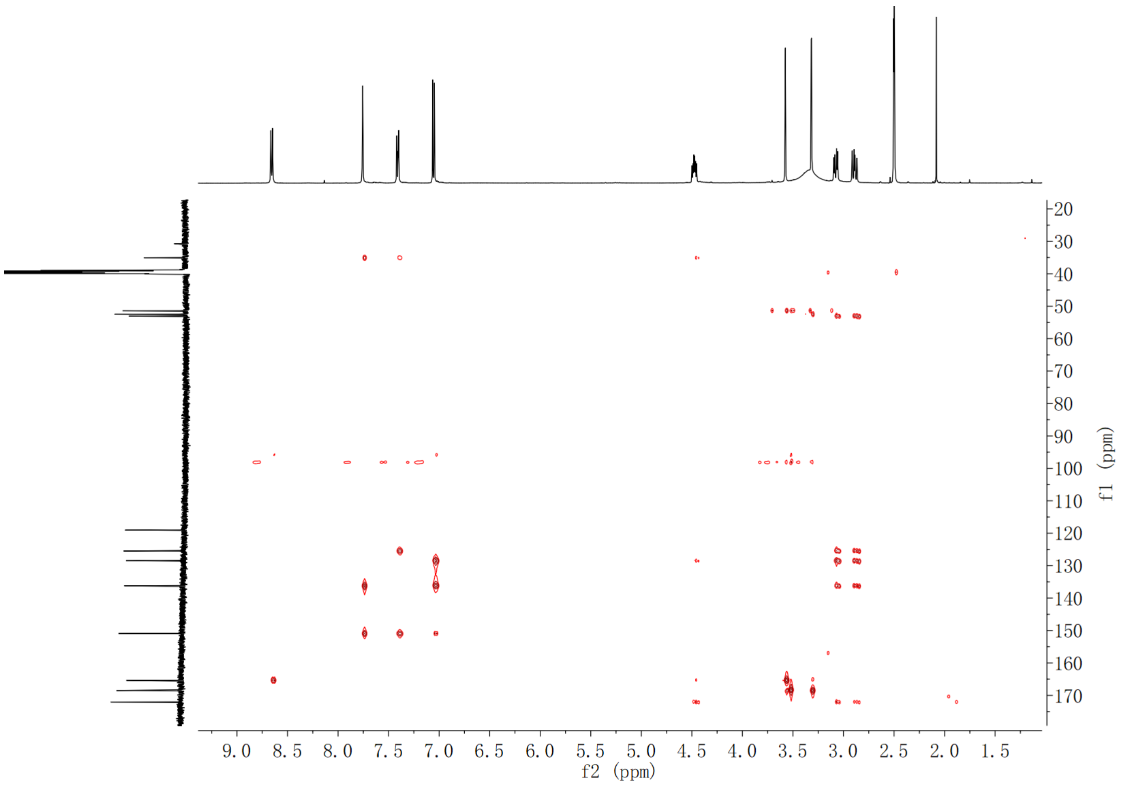

**Supplementary Fig. 227. HMBC spectrum of compound (2*S*,3*S*)-*t*-ES-a21 in DMSO-$d_6$**

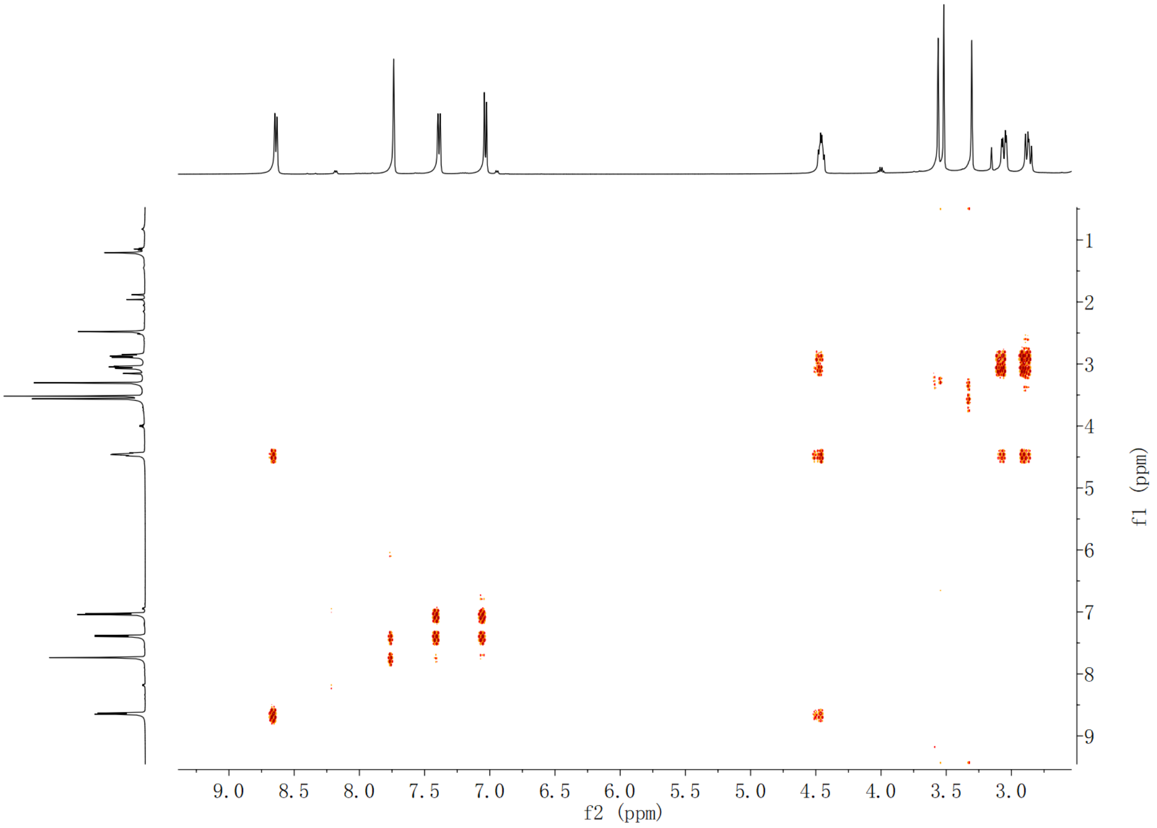

**Supplementary Fig. 228. $^1$H-$^1$H COSY spectrum of compound (2*S*,3*S*)-*t*-ES-a21 in DMSO-$d_6$**

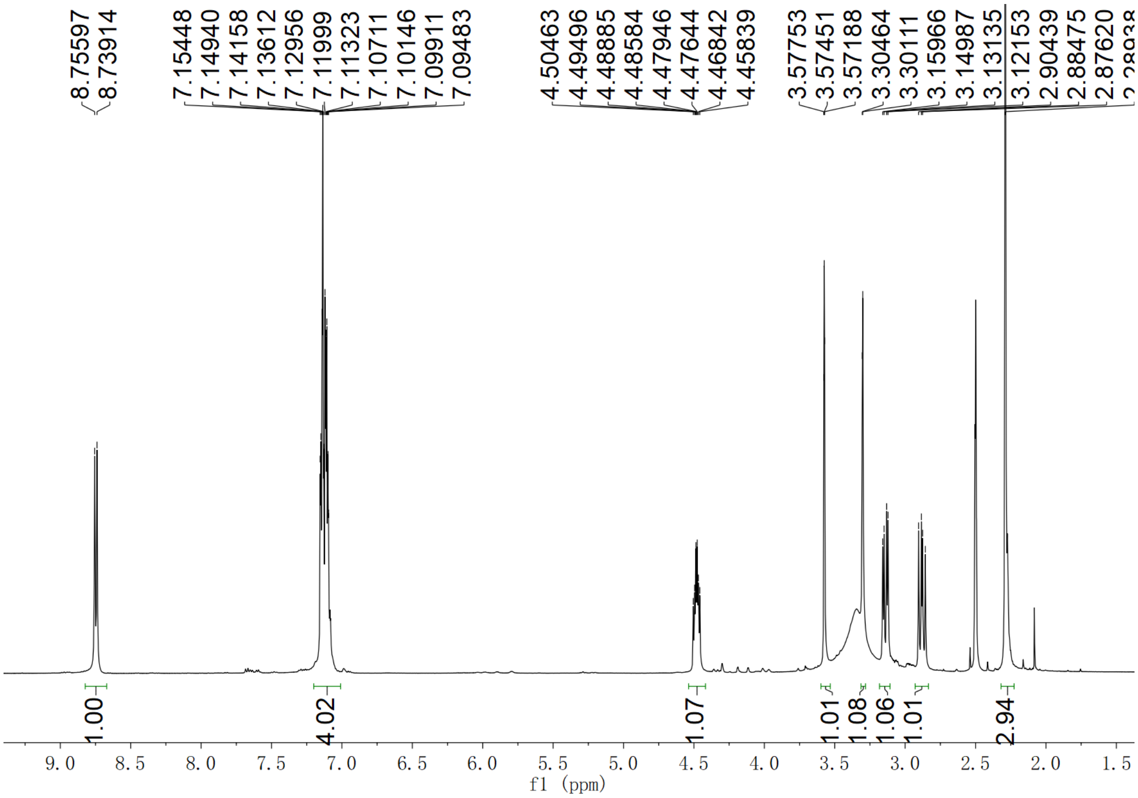

Supplementary Fig. 229. $^{1}$H NMR spectrum of compound (2*S*,3*S*)-*t*-ES-a22 in DMSO-$d_6$

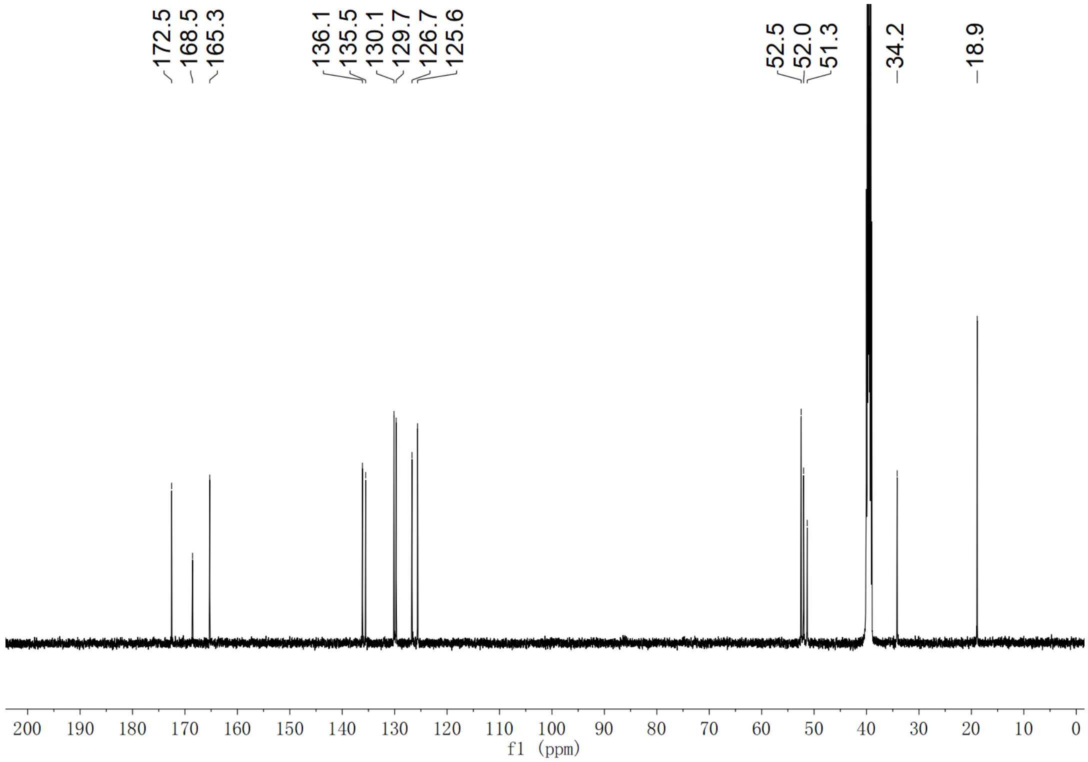

Supplementary Fig. 230. $^{13}$C NMR spectrum of compound (2*S*,3*S*)-*t*-ES-a22 in DMSO-$d_6$

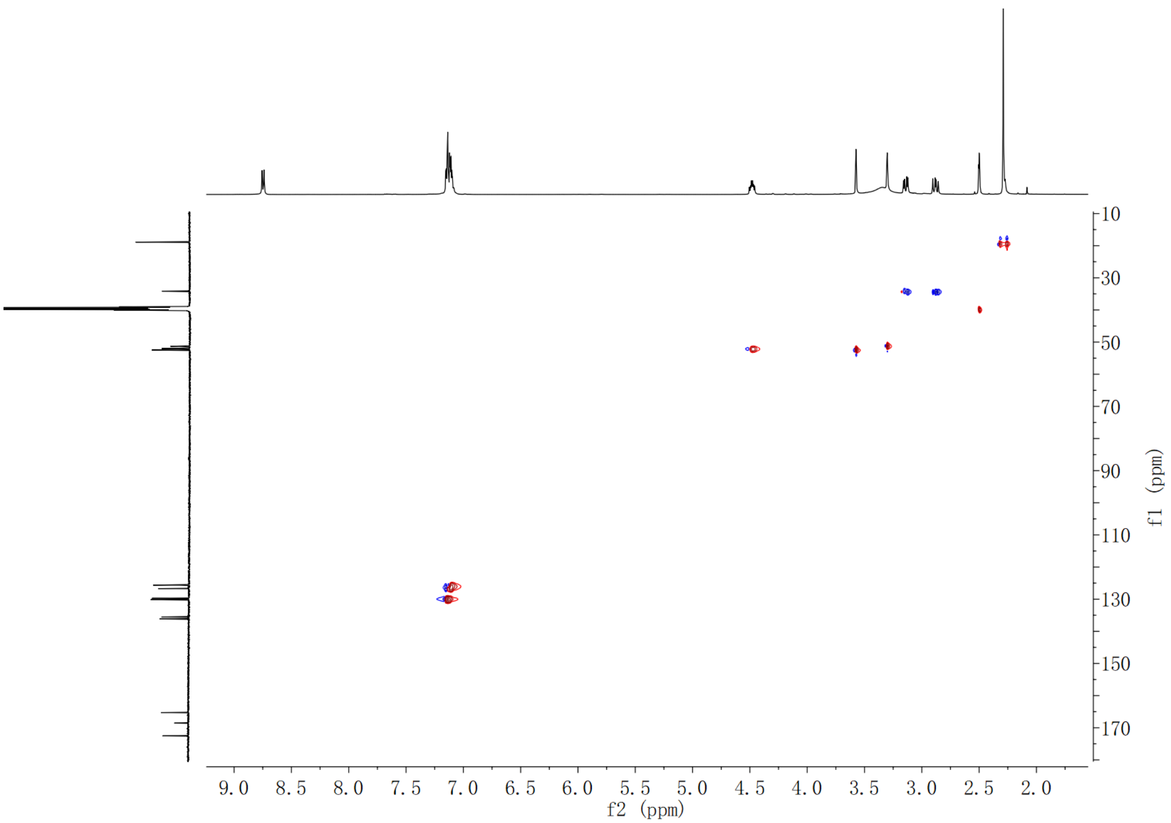

**Supplementary Fig. 231. HSQC spectrum of compound (2*S*,3*S*)-*t*-ES-a22 in DMSO-$d_6$**

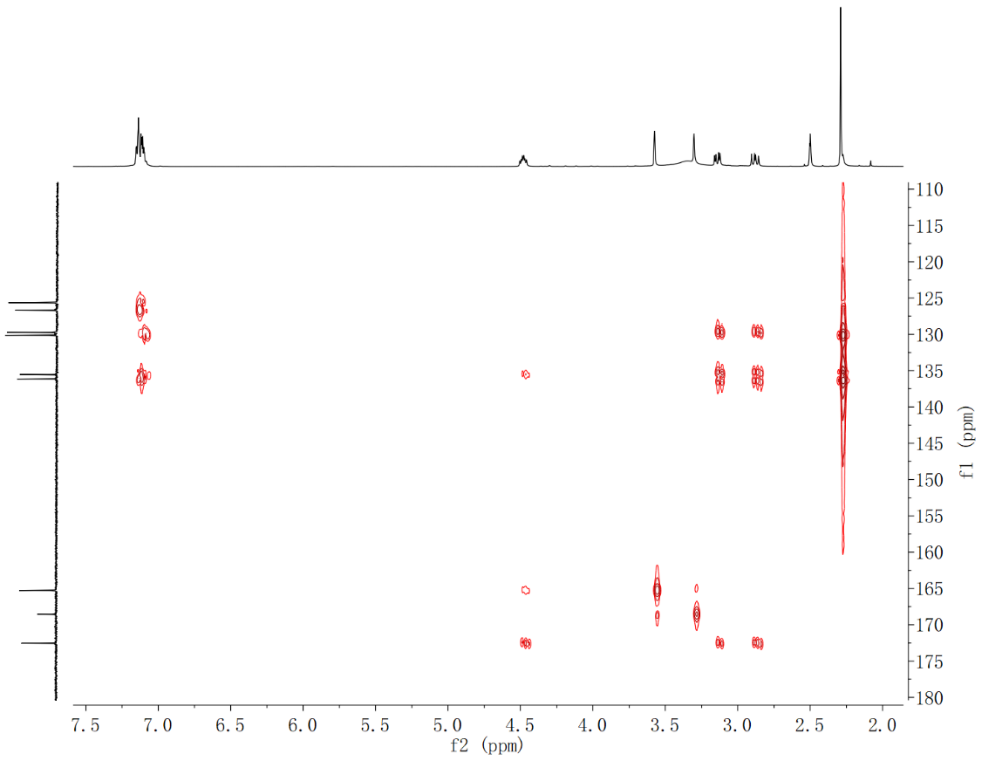

**Supplementary Fig. 232. HMBC spectrum of compound (2*S*,3*S*)-*t*-ES-a22 in DMSO-$d_6$**

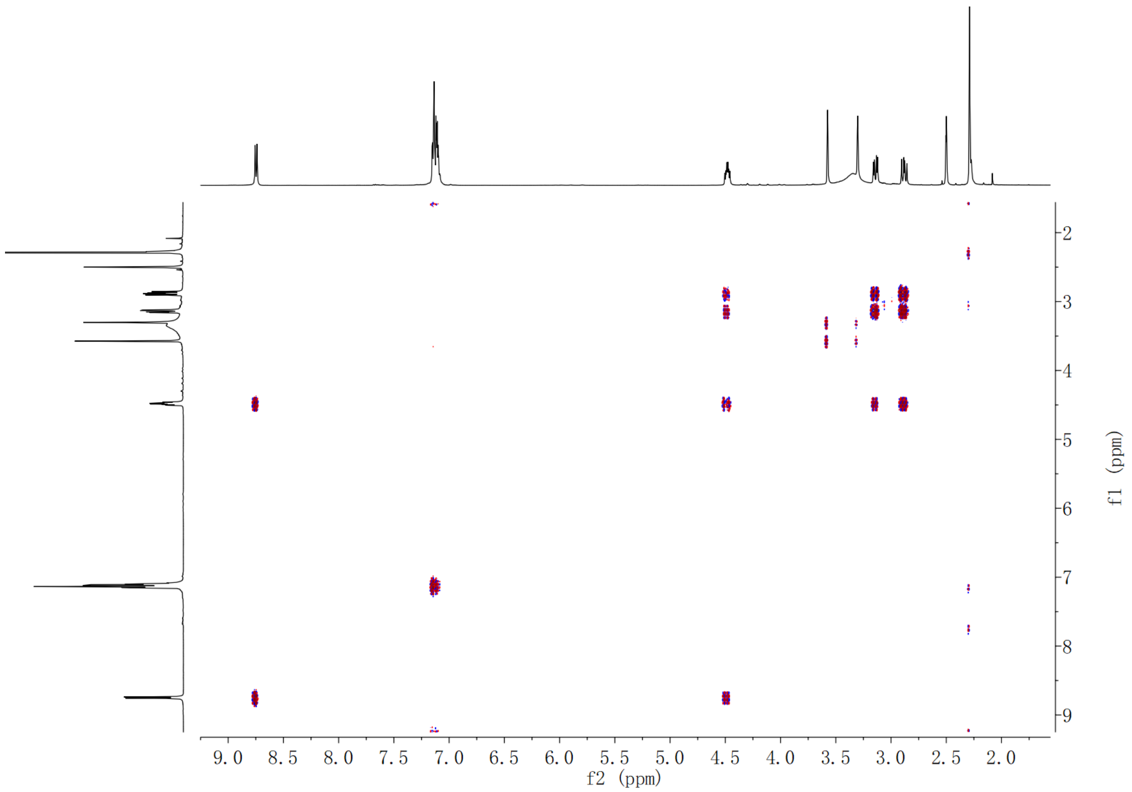

Supplementary Fig. 233. $^{1}H$-$^{1}H$ COSY spectrum of compound (2*S*,3*S*)-*t*-ES-a22 in DMSO-$d_6$

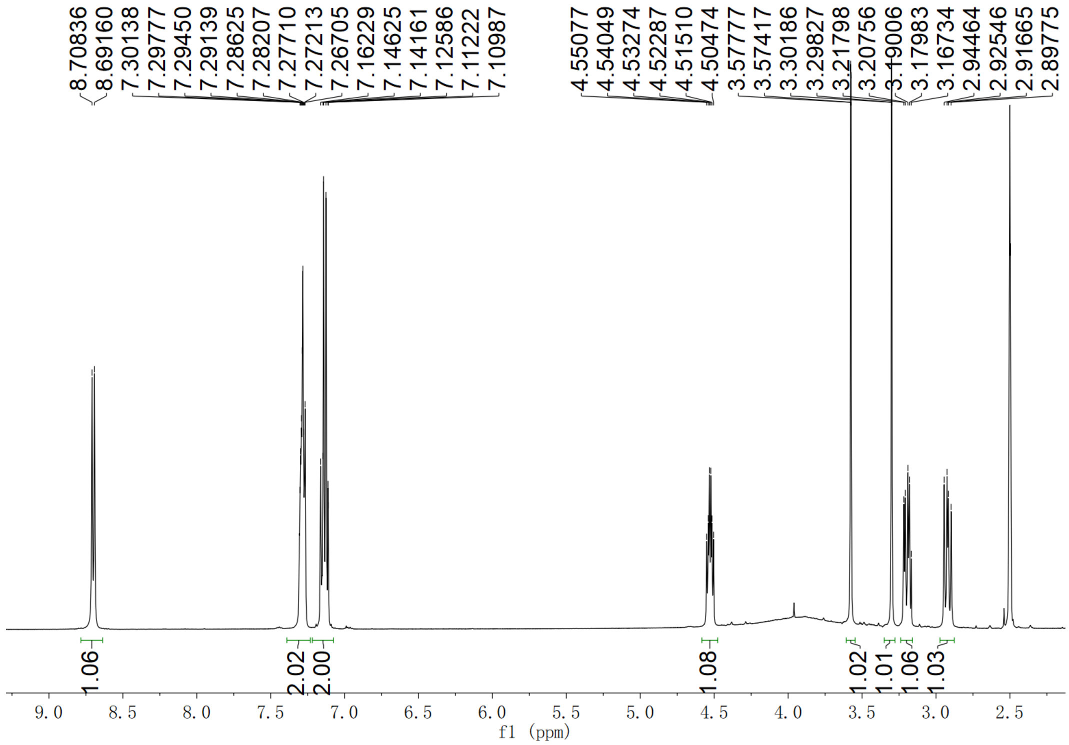

Supplementary Fig. 234. $^{1}H$ NMR spectrum of compound (2*S*,3*S*)-*t*-ES-a23 in DMSO-$d_6$

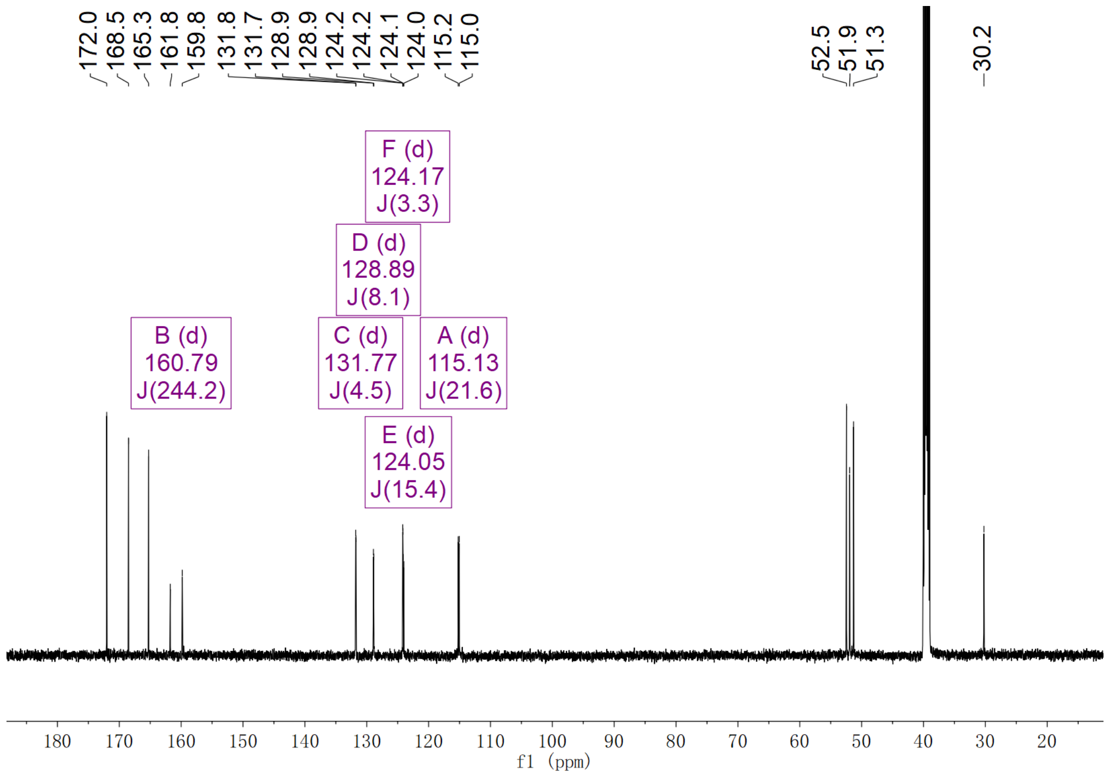

**Supplementary Fig. 235. $^{13}$C NMR spectrum of compound (2*S*,3*S*)-*t*-ES-a23 in DMSO-$d_6$**

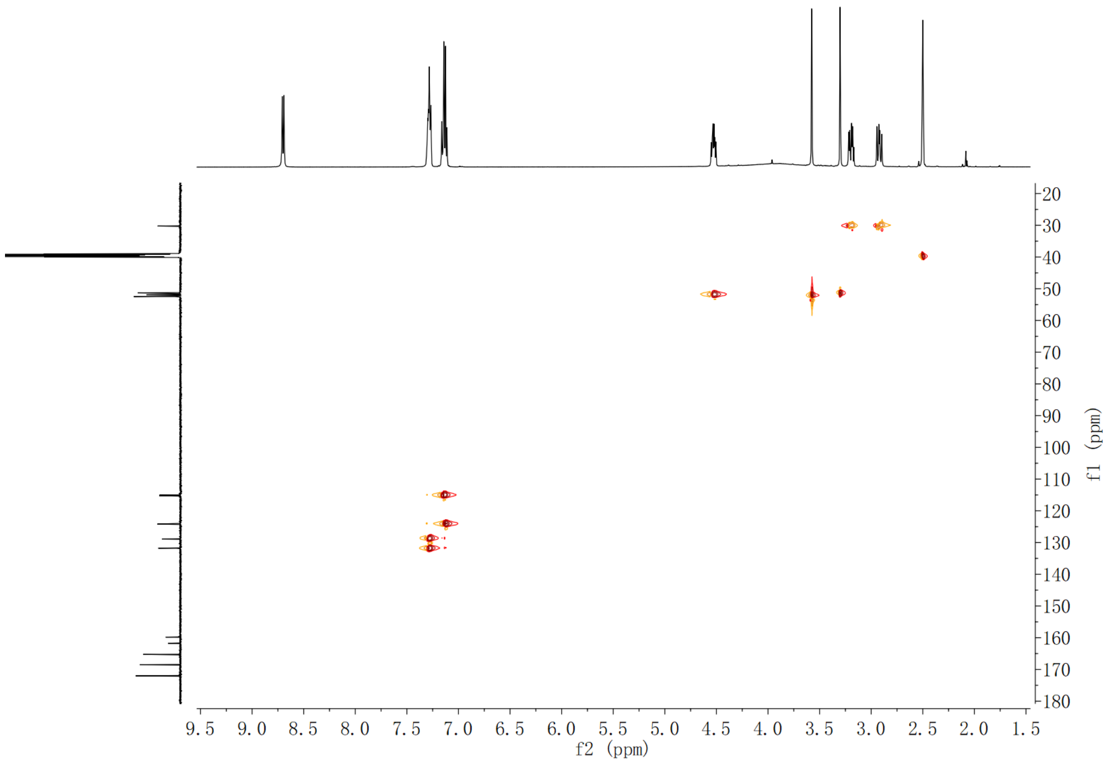

**Supplementary Fig. 236. HSQC spectrum of compound (2*S*,3*S*)-*t*-ES-a23 in DMSO-$d_6$**

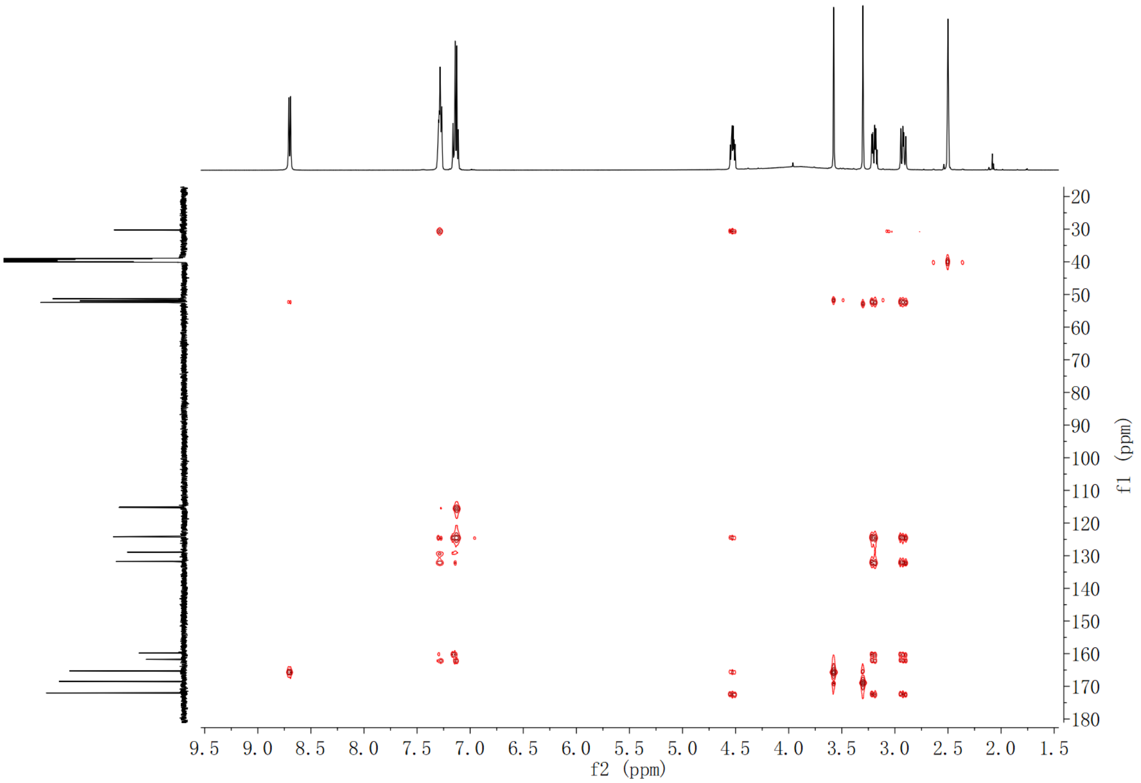

**Supplementary Fig. 237. HMBC spectrum of compound (2*S*,3*S*)-*t*-ES-a23 in DMSO-$d_6$**

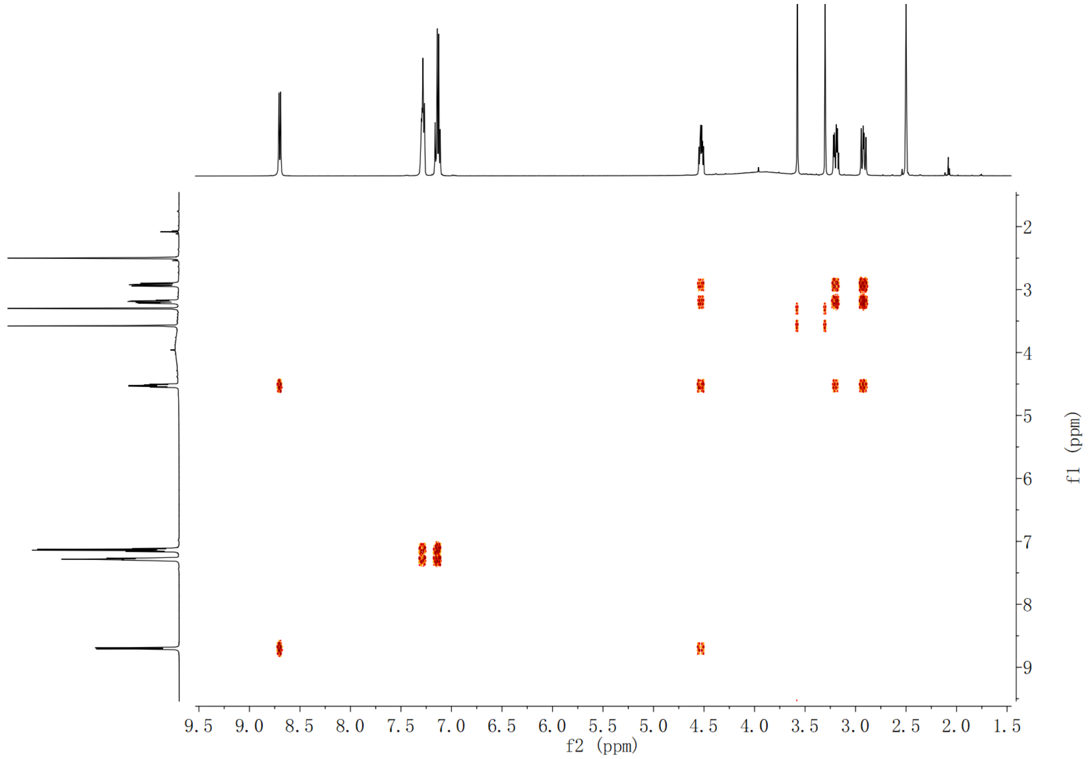

**Supplementary Fig. 238. $^1$H-$^1$H COSY spectrum of compound (2*S*,3*S*)-*t*-ES-a23 in DMSO-$d_6$**

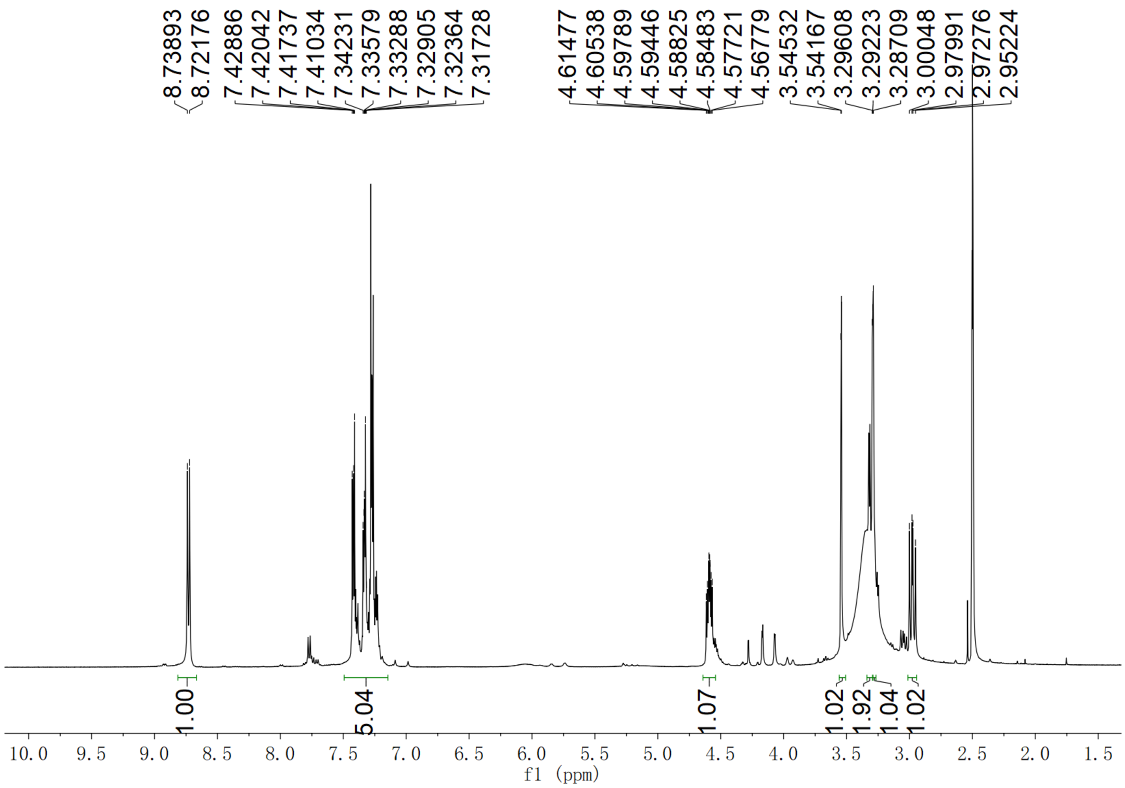

**Supplementary Fig. 239.** $^{1}$**H NMR spectrum of compound (2*S*,3*S*)-*t*-ES-a24 in DMSO-$d_6$**

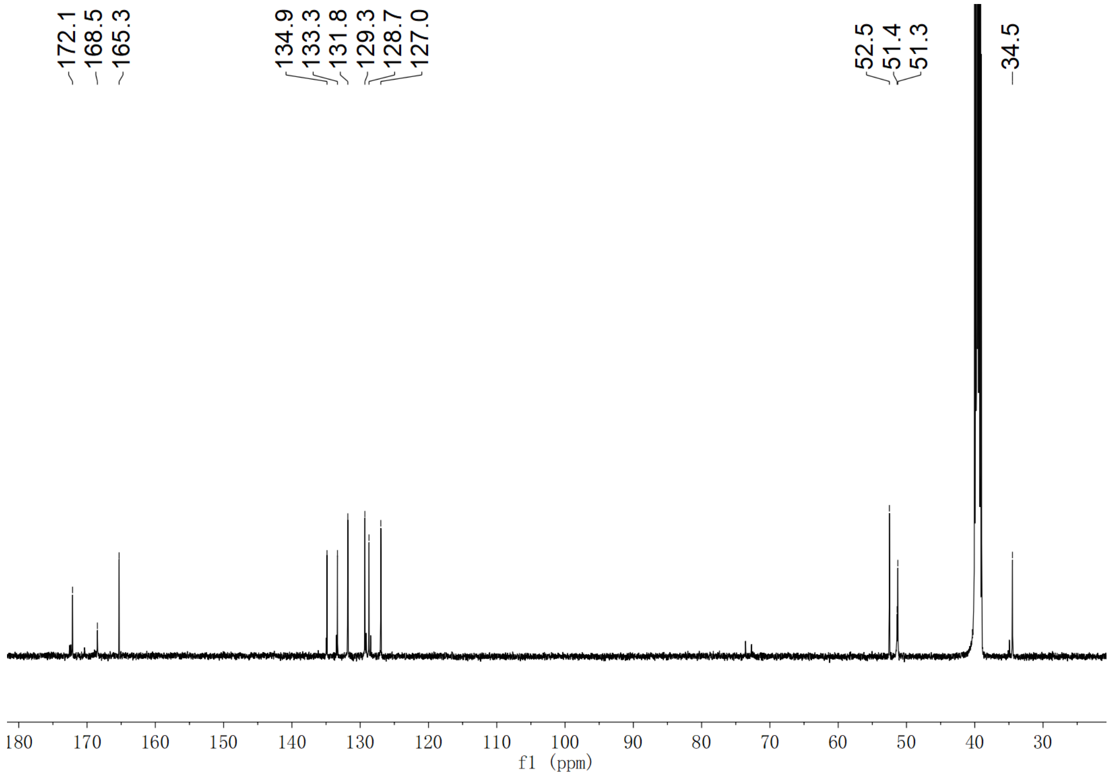

**Supplementary Fig. 240.** $^{13}$**C NMR spectrum of compound (2*S*,3*S*)-*t*-ES-a24 in DMSO-$d_6$**

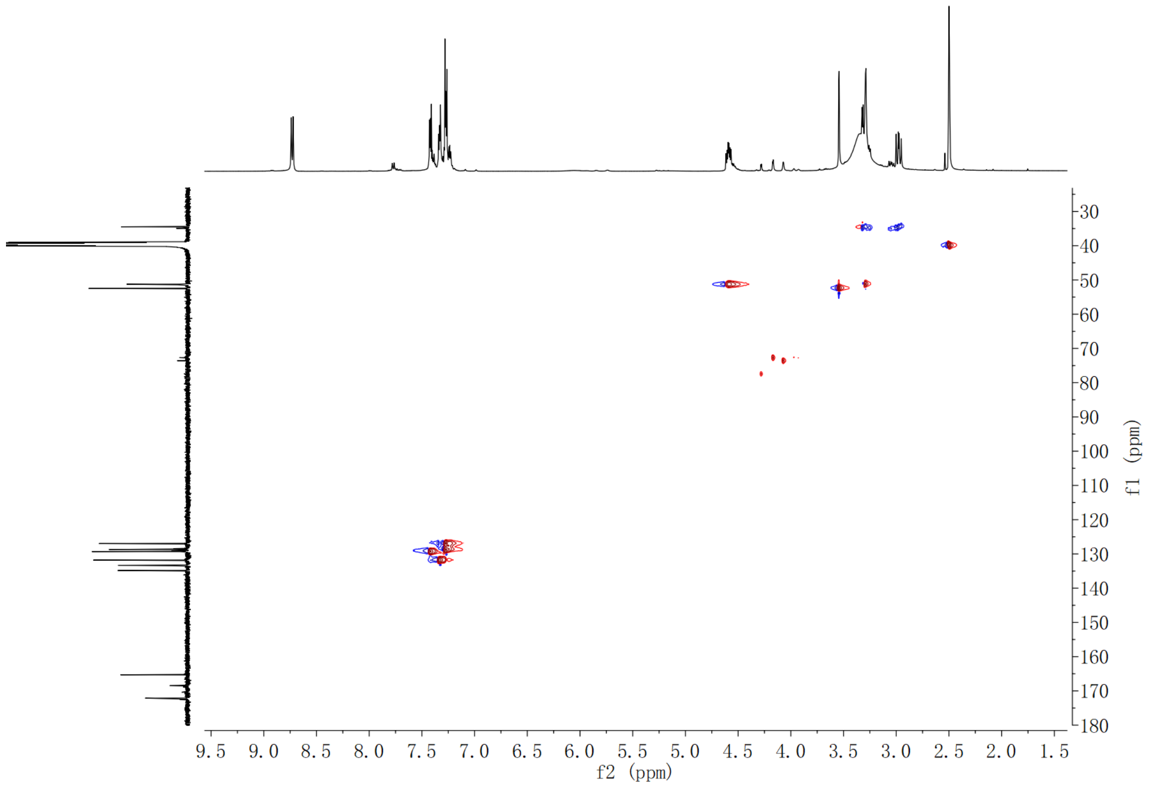

**Supplementary Fig. 241. HSQC spectrum of compound (2*S*,3*S*)-*t*-ES-a24 in DMSO-*d*$_6$**

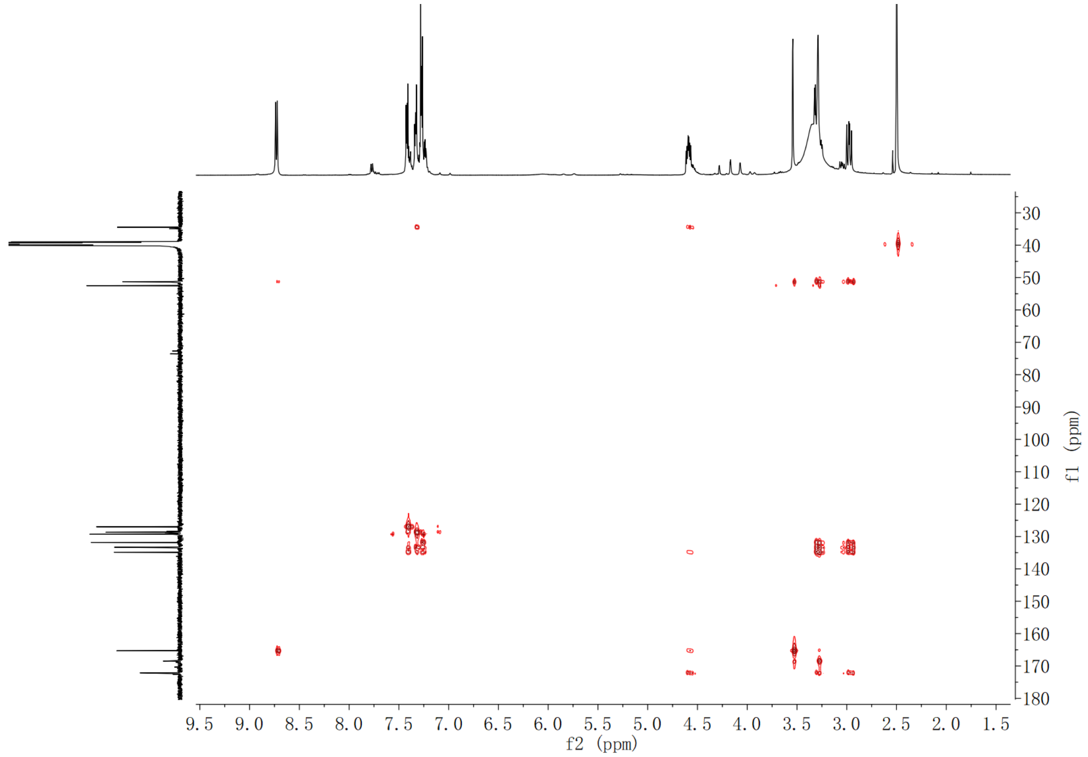

**Supplementary Fig. 242. HMBC spectrum of compound (2*S*,3*S*)-*t*-ES-a24 in DMSO-*d*$_6$**

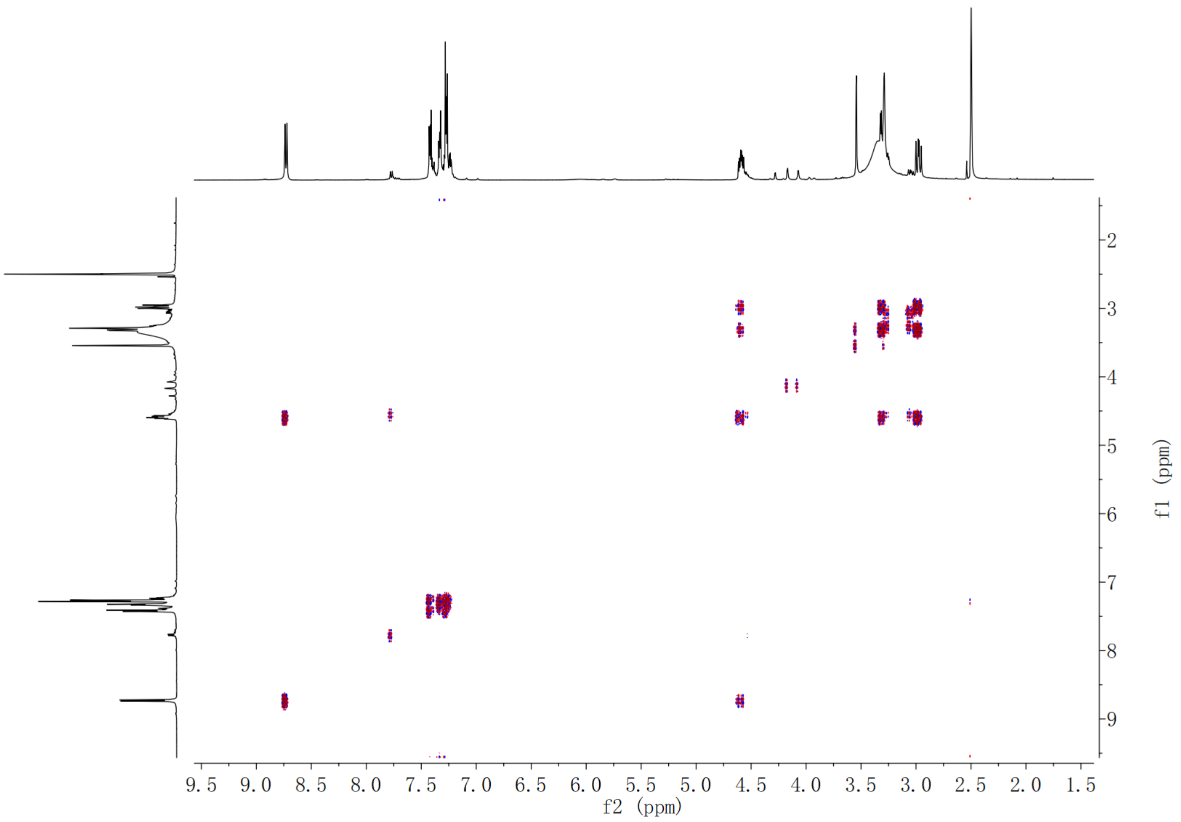

**Supplementary Fig. 243.** **$^{1}$H-$^{1}$H COSY spectrum of compound (2*S*,3*S*)-*t*-ES-a24 in DMSO-*d*$_6$**

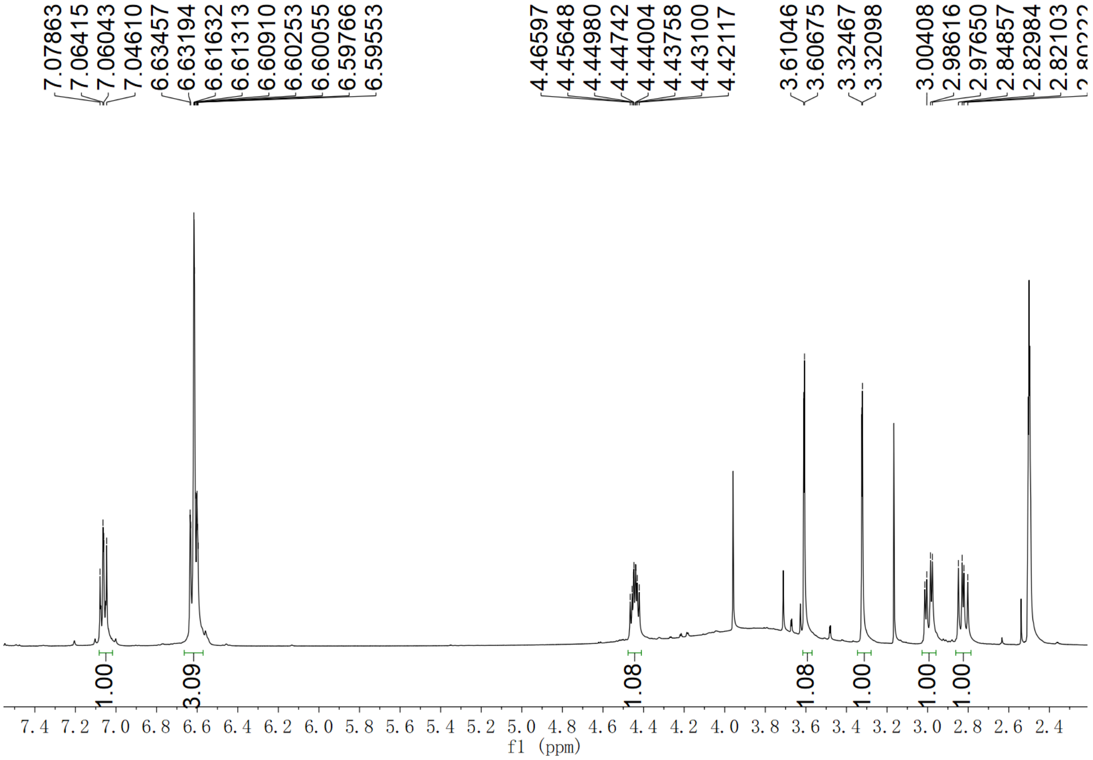

**Supplementary Fig. 244.** **$^{1}$H NMR spectrum of compound (2*S*,3*S*)-*t*-ES-a25 in DMSO-*d*$_6$**

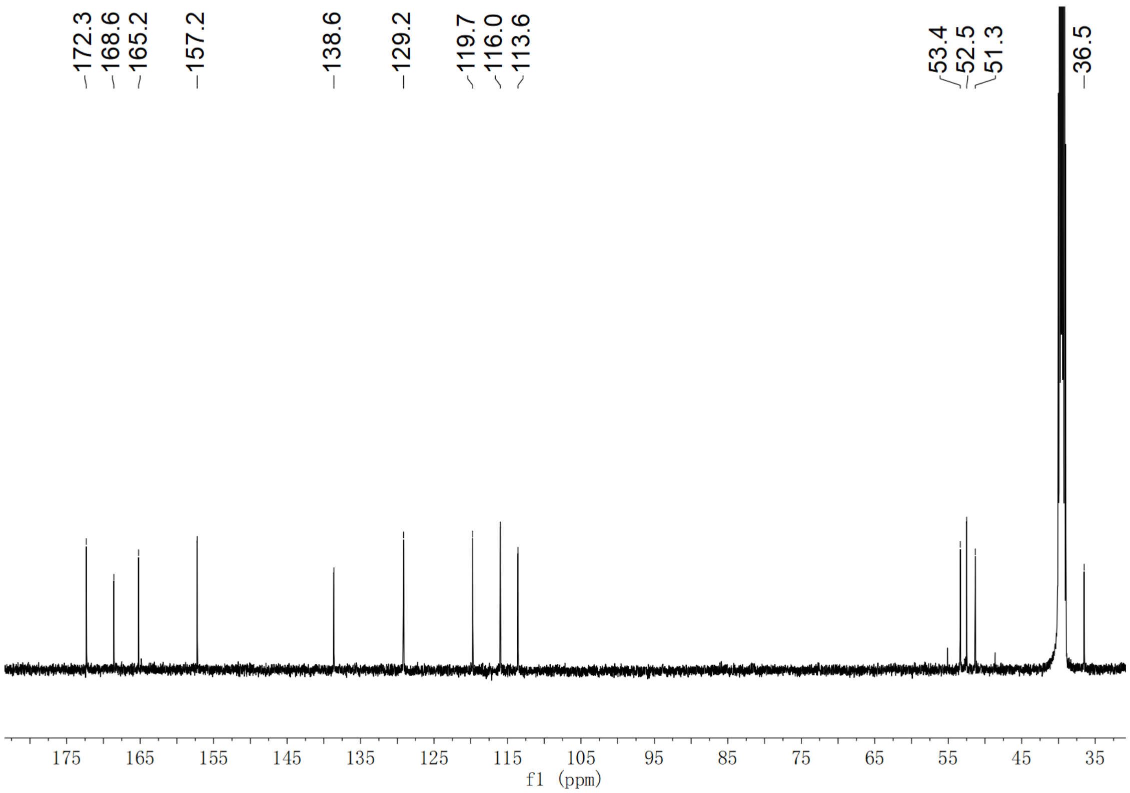

**Supplementary Fig. 245. $^{13}$C NMR spectrum of compound (2*S*,3*S*)-*t*-ES-a25 in DMSO-$d_6$**

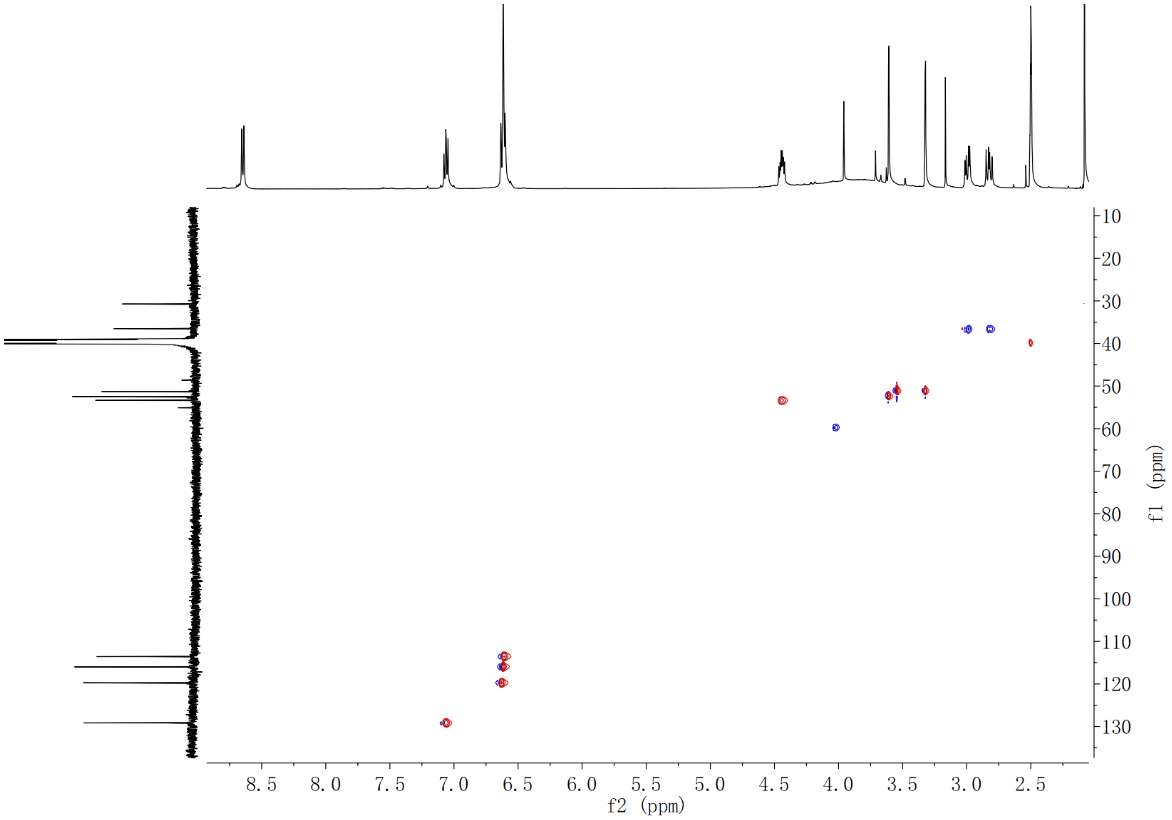

**Supplementary Fig. 246. HSQC spectrum of compound (2*S*,3*S*)-*t*-ES-a25 in DMSO-$d_6$**

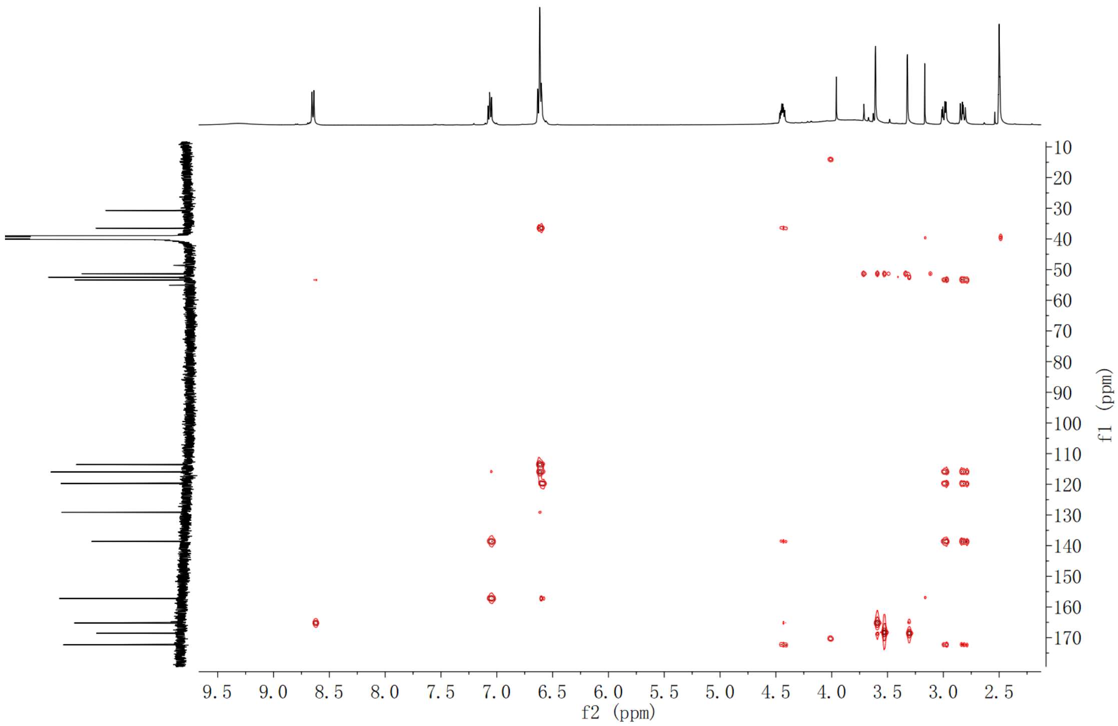

**Supplementary Fig. 247. HMBC spectrum of compound (2*S*,3*S*)-*t*-ES-a25 in DMSO-$d_6$**

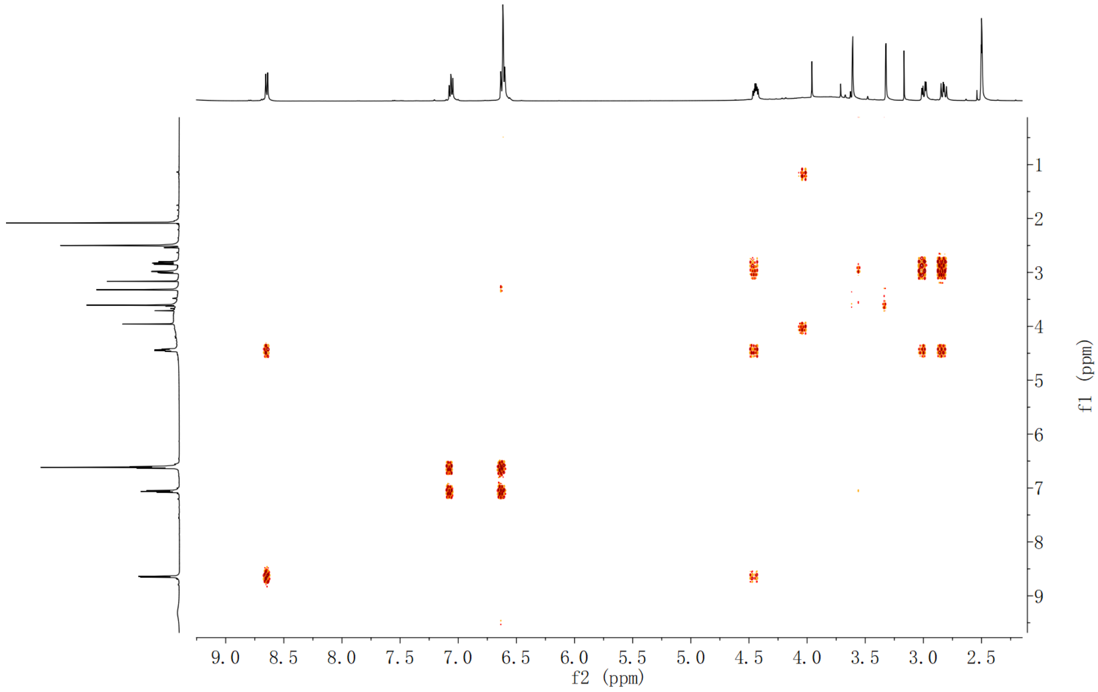

**Supplementary Fig. 248. $^1$H-$^1$H COSY spectrum of compound (2*S*,3*S*)-*t*-ES-a25 in DMSO-$d_6$**

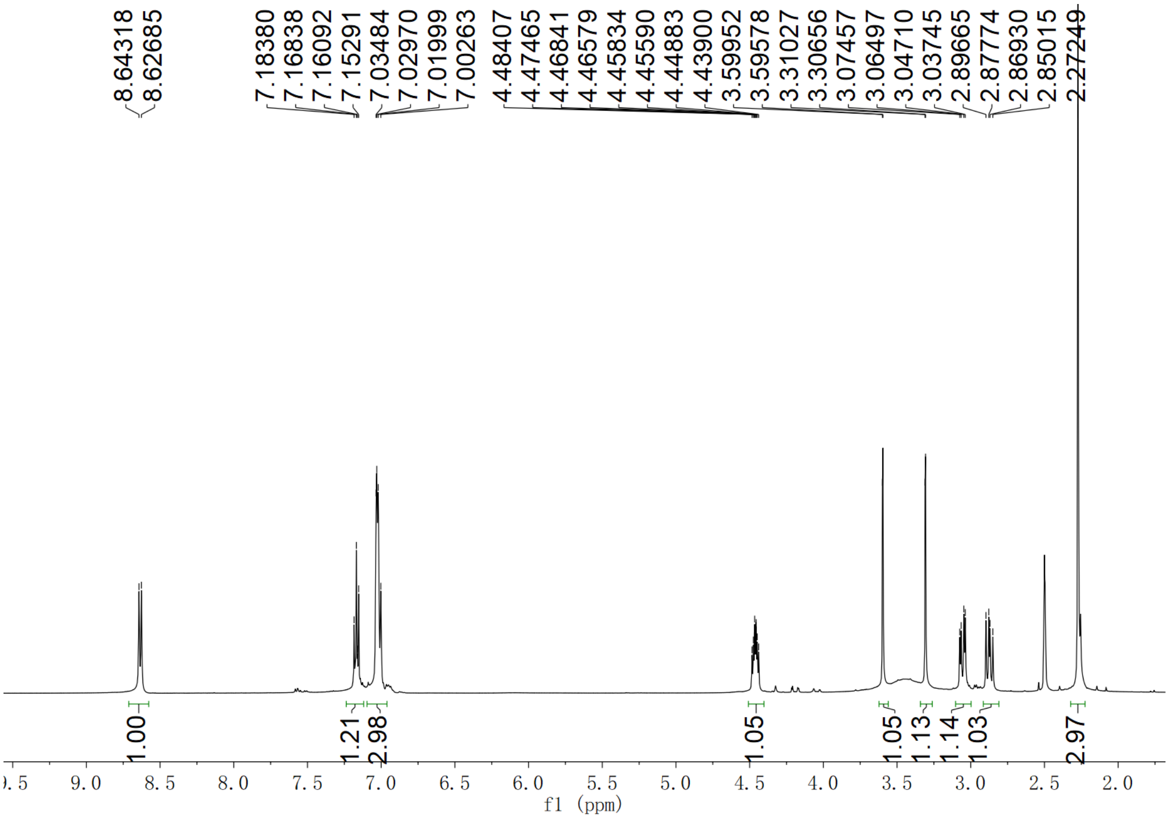

**Supplementary Fig. 249.** $^{1}$**H NMR spectrum of compound (2*S*,3*S*)-*t*-ES-a26 in DMSO-*d*$_6$**

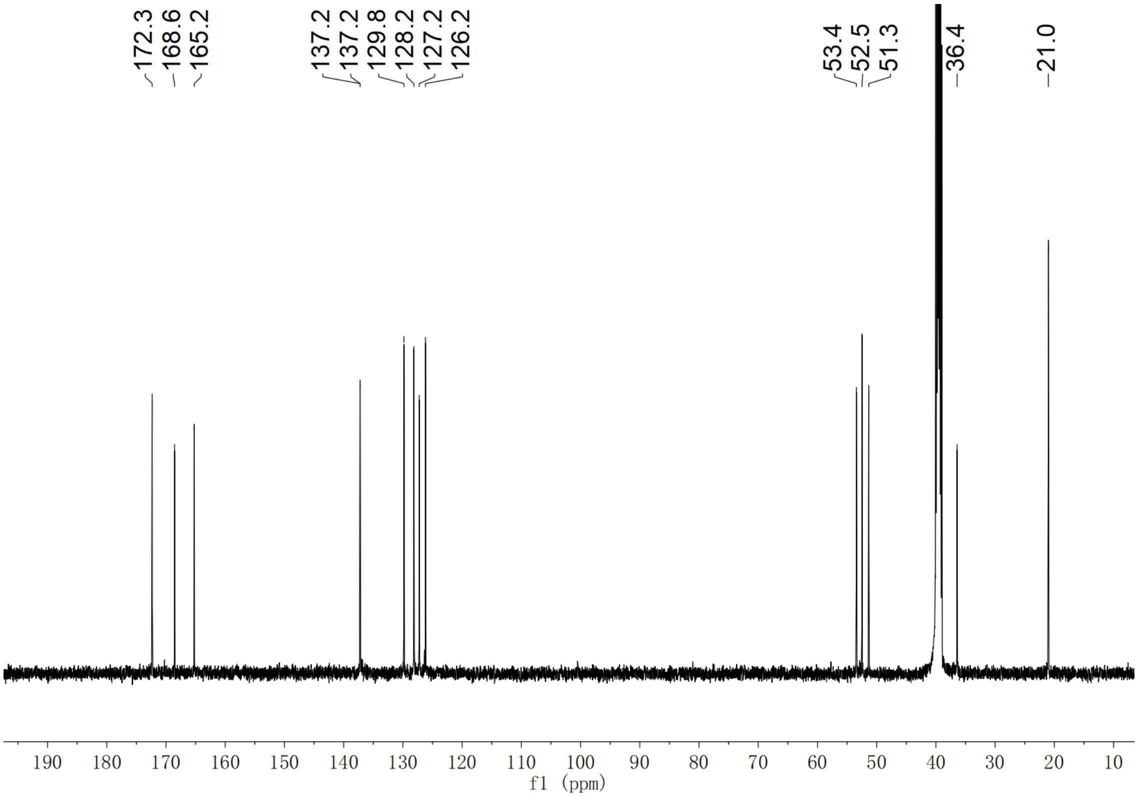

**Supplementary Fig. 250.** $^{13}$**C NMR spectrum of compound (2*S*,3*S*)-*t*-ES-a26 in DMSO-*d*$_6$**

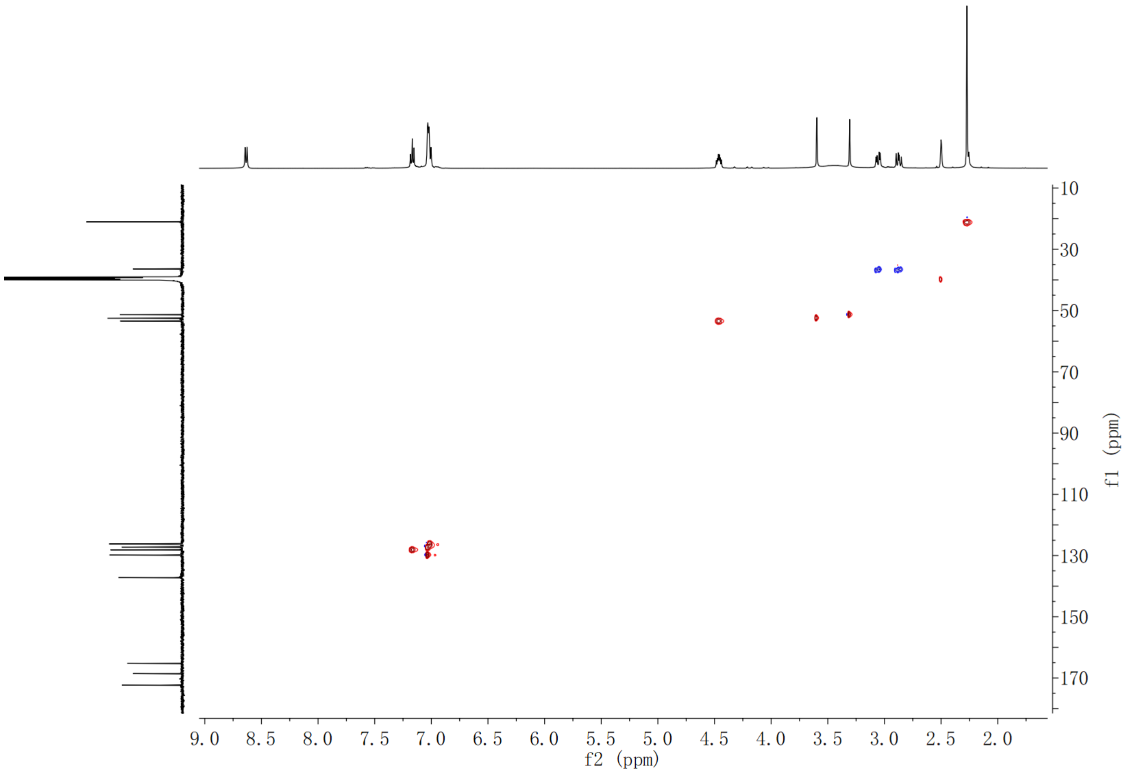

**Supplementary Fig. 251. HSQC spectrum of compound (2*S*,3*S*)-*t*-ES-a26 in DMSO-$d_6$**

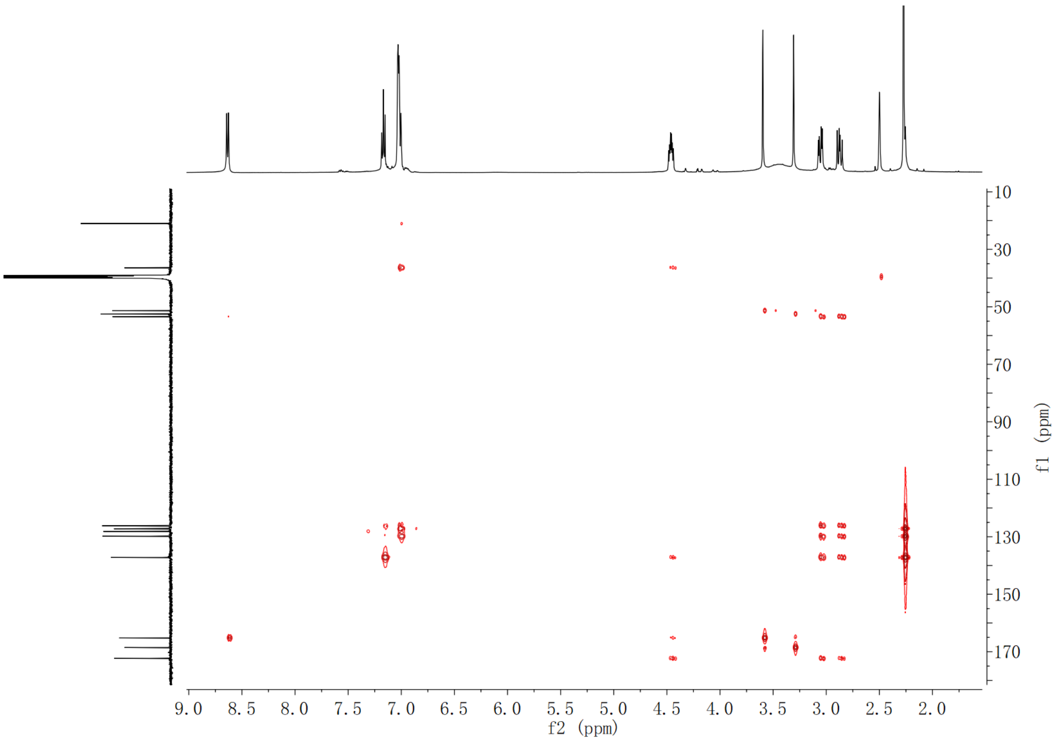

**Supplementary Fig. 252. HMBC spectrum of compound (2*S*,3*S*)-*t*-ES-a26 in DMSO-$d_6$**

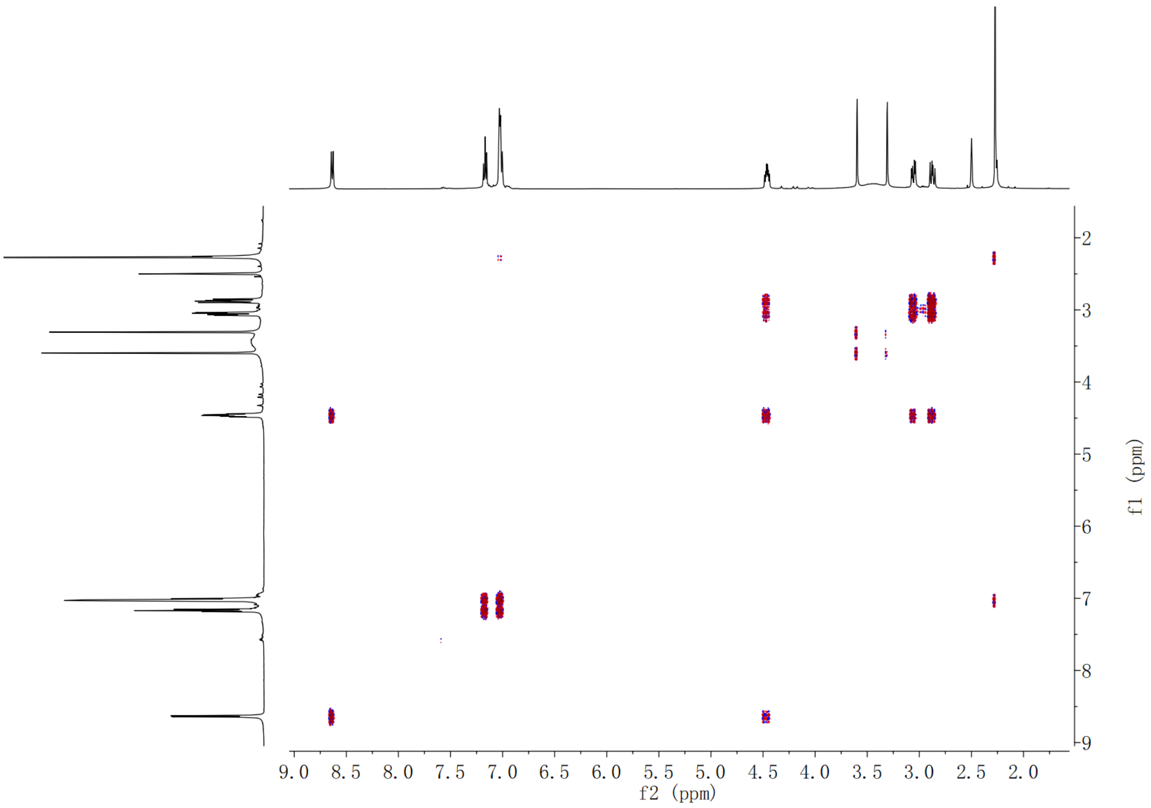

**Supplementary Fig. 253.** $^{1}$**H-**$^{1}$**H COSY spectrum of compound (2*S*,3*S*)-*t*-ES-a26 in DMSO-*d*$_6$**

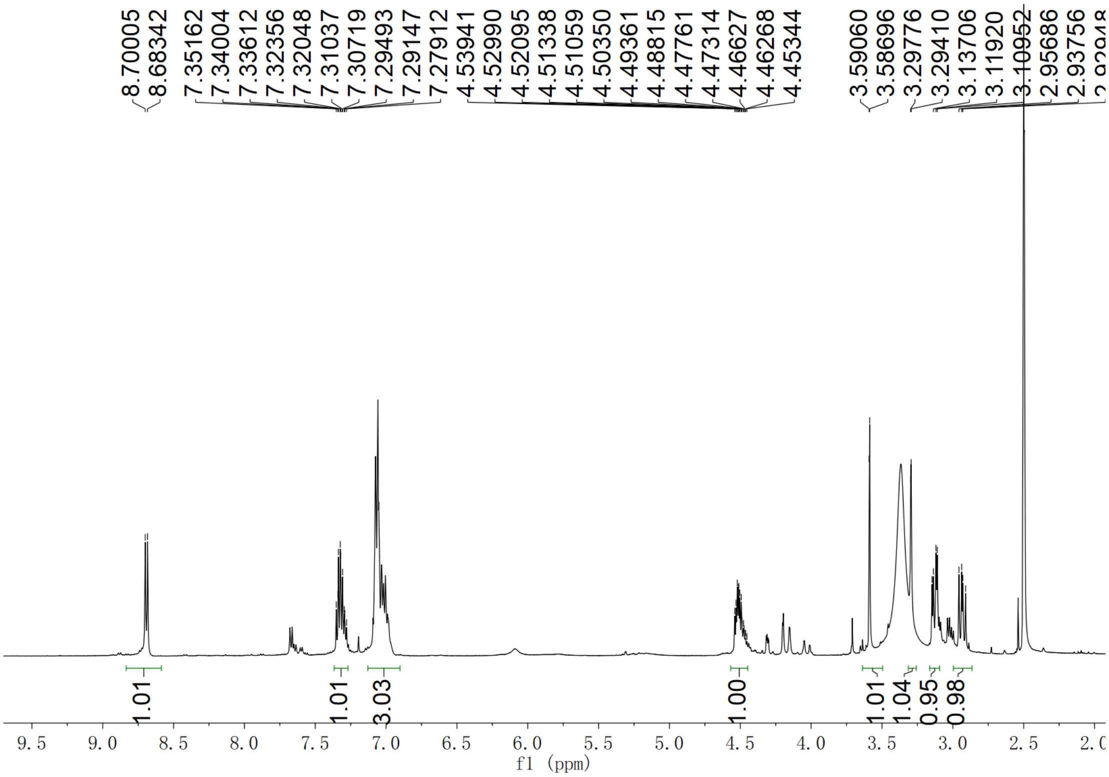

**Supplementary Fig. 254.** $^{1}$**H NMR spectrum of compound (2*S*,3*S*)-*t*-ES-a27 in DMSO-*d*$_6$**

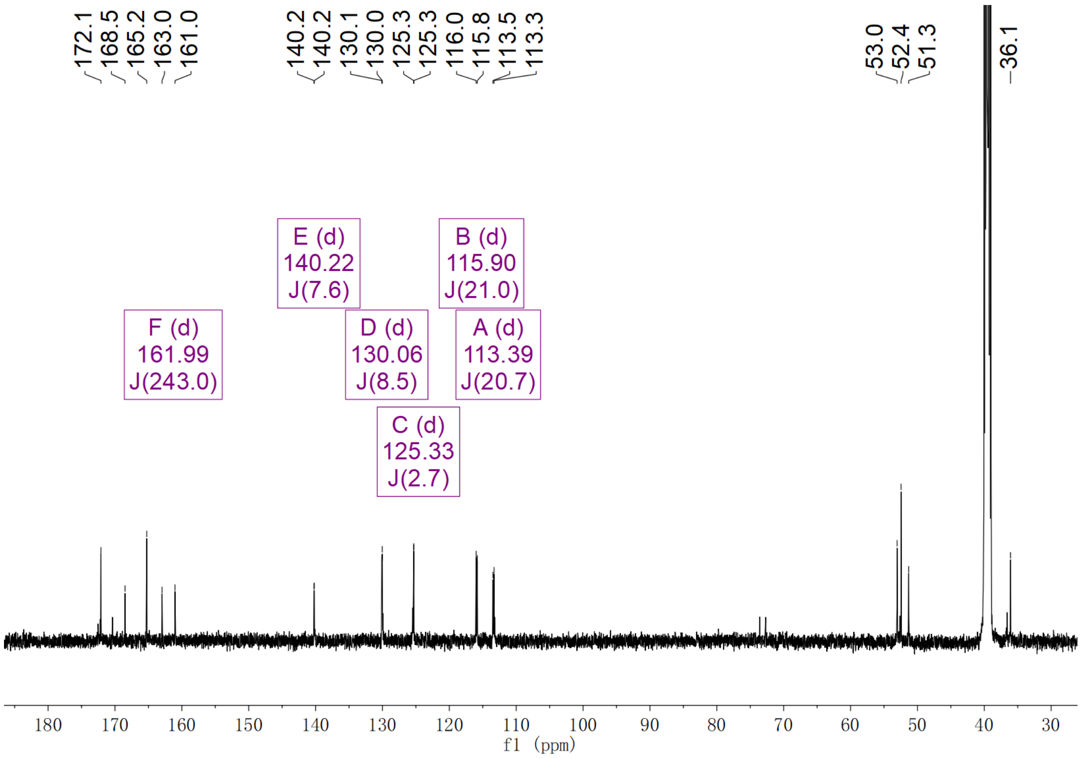

**Supplementary Fig. 255. $^{13}$C NMR spectrum of compound (2*S*,3*S*)-*t*-ES-a27 in DMSO-$d_6$**

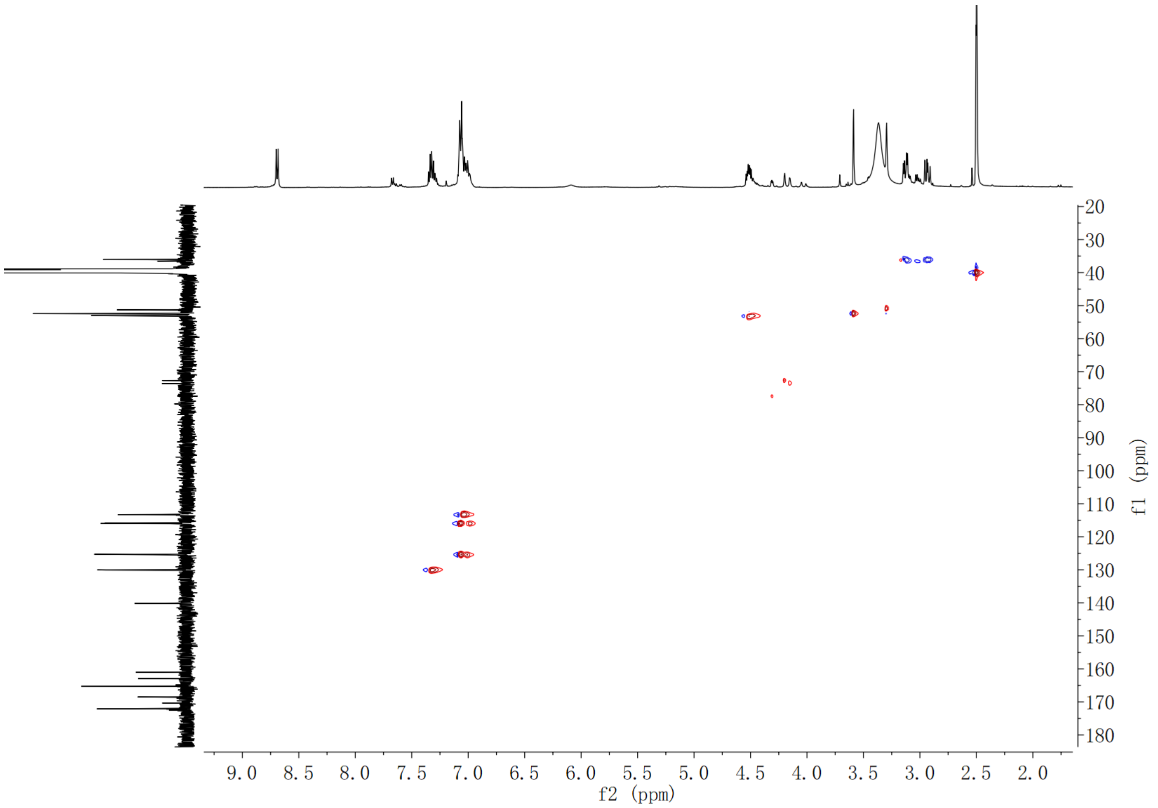

**Supplementary Fig. 256. HSQC spectrum of compound (2*S*,3*S*)-*t*-ES-a27 in DMSO-$d_6$**

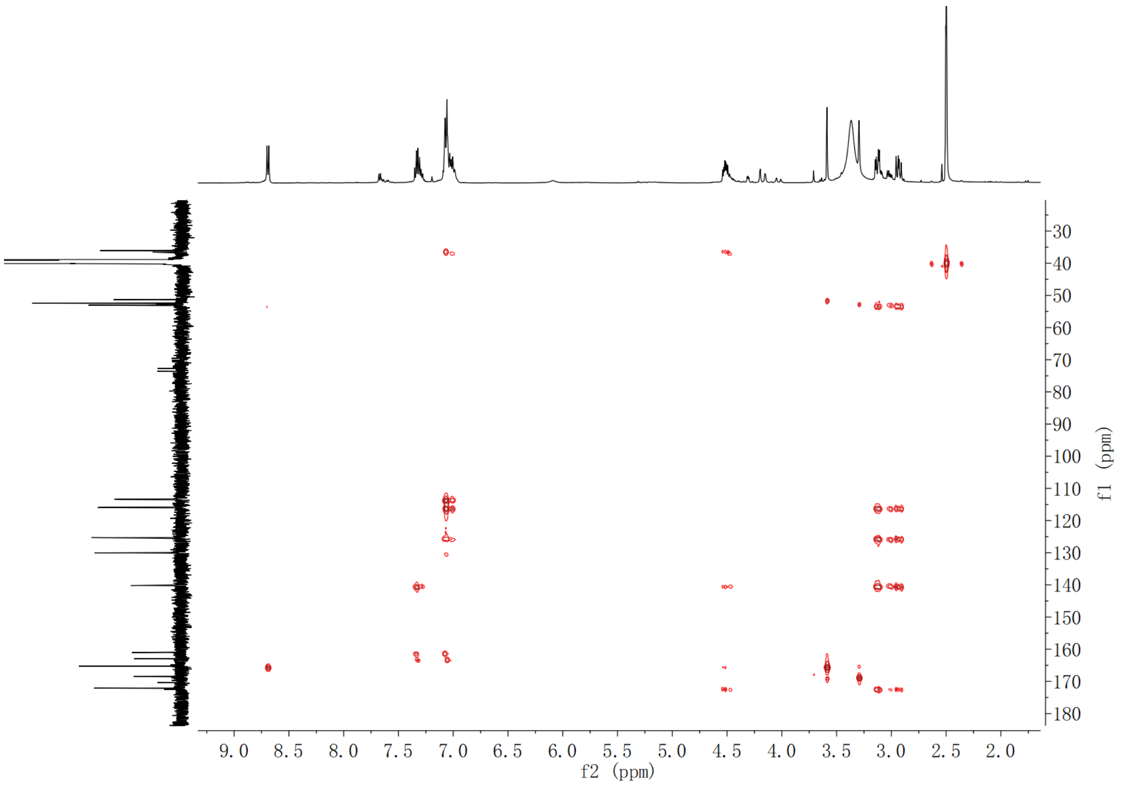

**Supplementary Fig. 257. HMBC spectrum of compound (2*S*,3*S*)-*t*-ES-a27 in DMSO-$d_6$**

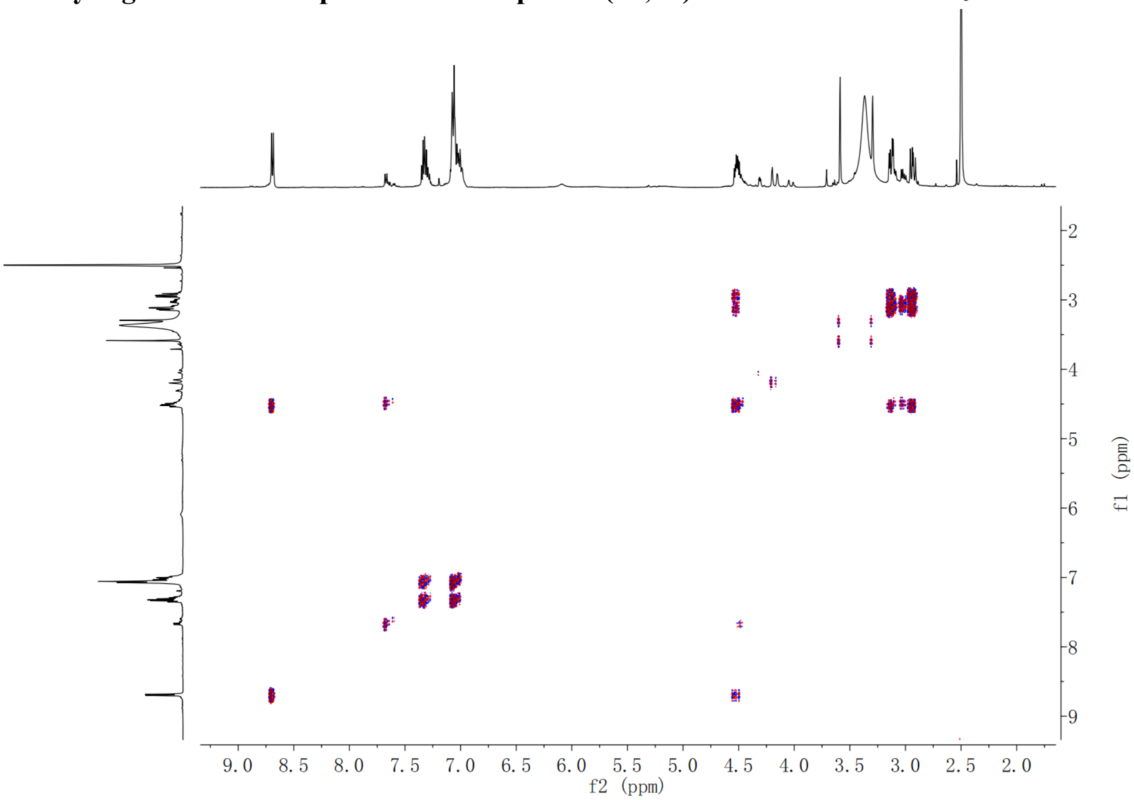

**Supplementary Fig. 258. $^1$H-$^1$H COSY spectrum of compound (2*S*,3*S*)-*t*-ES-a27 in DMSO-$d_6$**

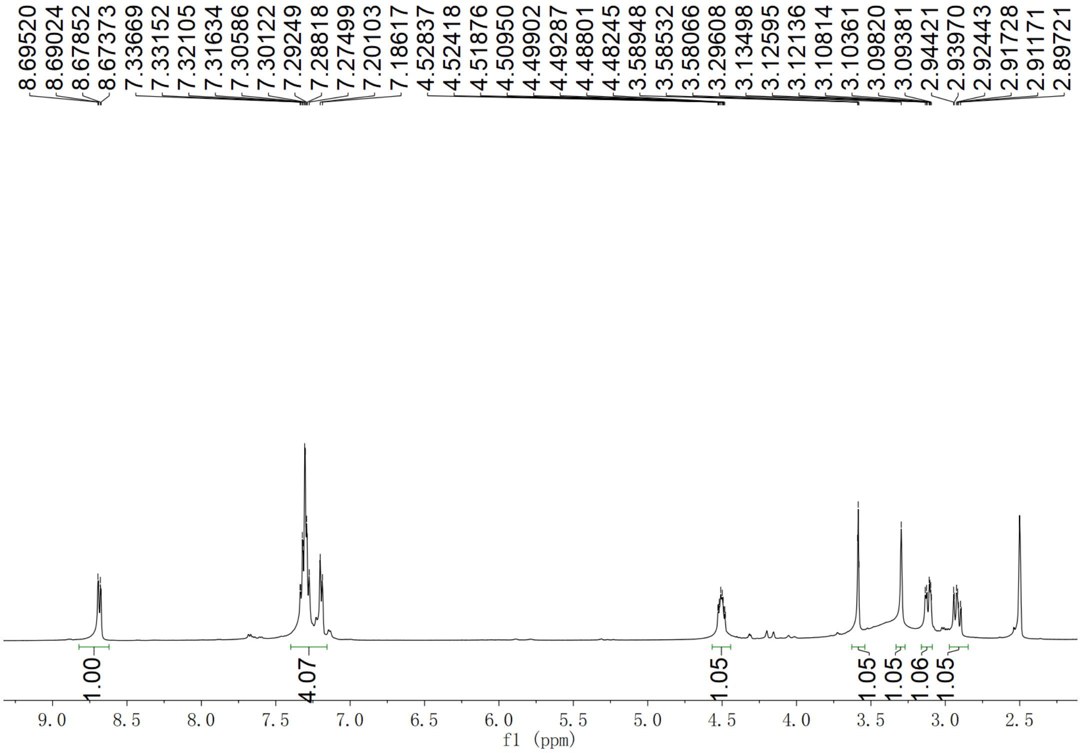

**Supplementary Fig. 259.** $^{1}$**H NMR spectrum of compound (2*S*,3*S*)-*t*-ES-a28 in DMSO-*d*$_6$**

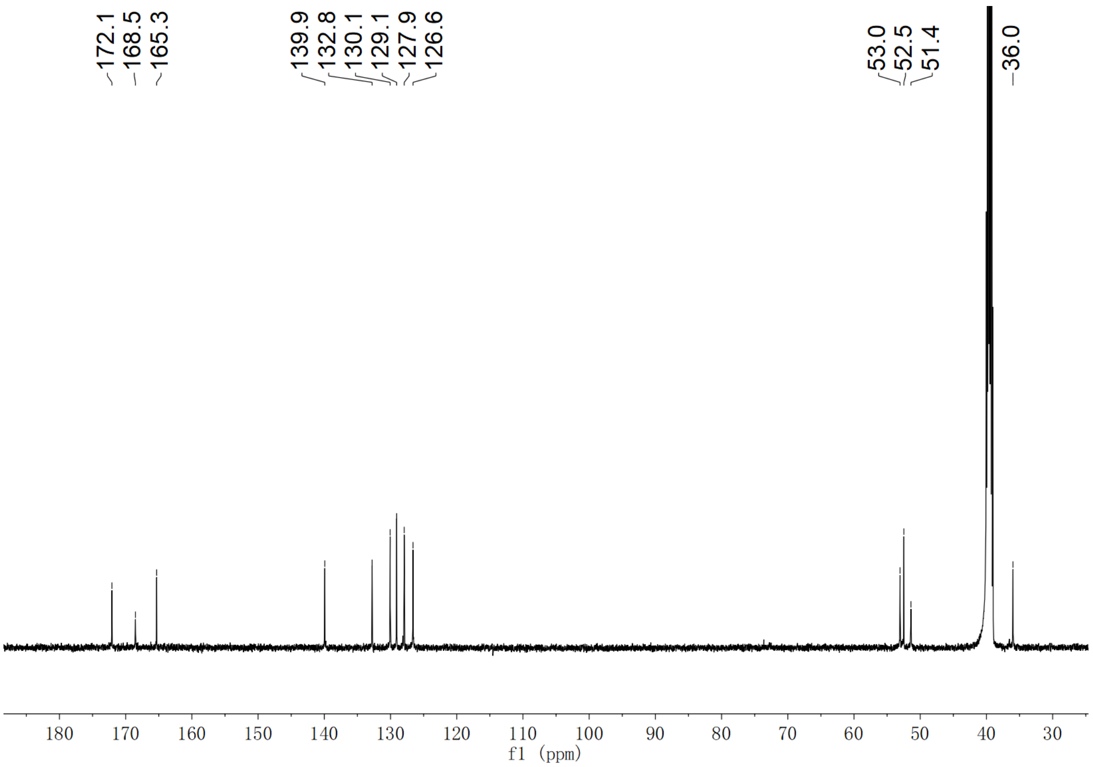

**Supplementary Fig. 260.** $^{13}$**C NMR spectrum of compound (2*S*,3*S*)-*t*-ES-a28 in DMSO-*d*$_6$**

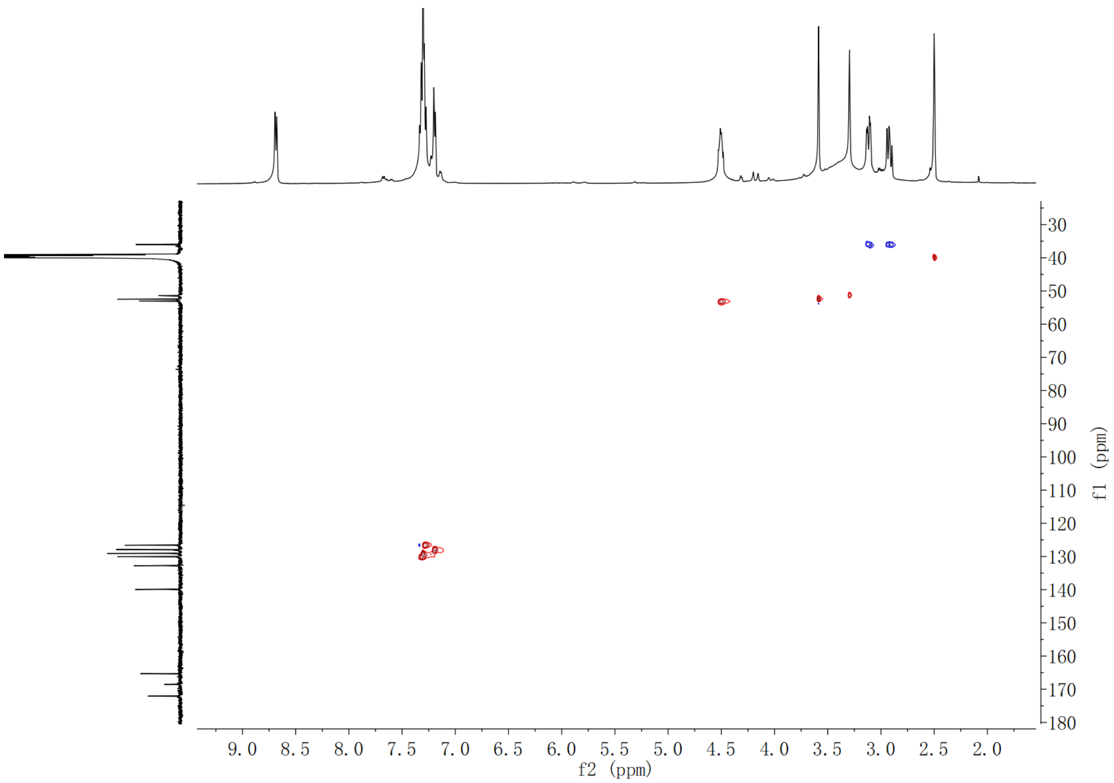

**Supplementary Fig. 261. HSQC spectrum of compound (2*S*,3*S*)-*t*-ES-a28 in DMSO-$d_6$**

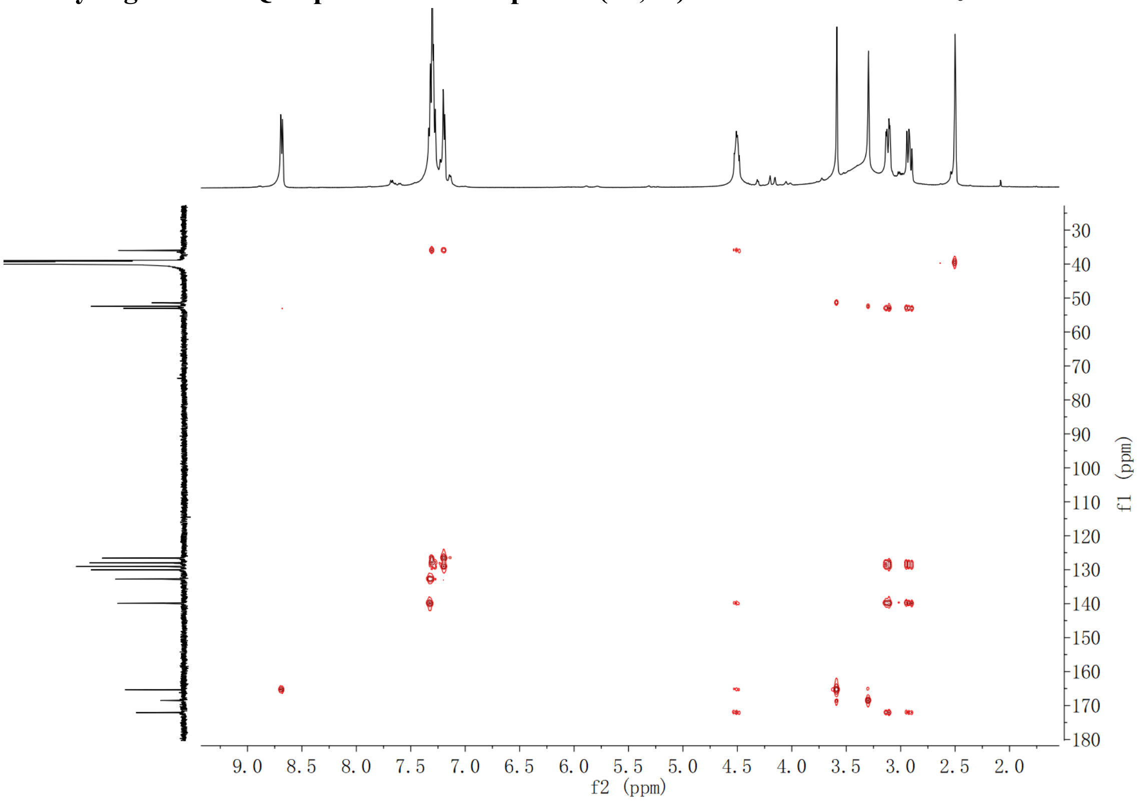

**Supplementary Fig. 262. HMBC spectrum of compound (2*S*,3*S*)-*t*-ES-a28 in DMSO-$d_6$**

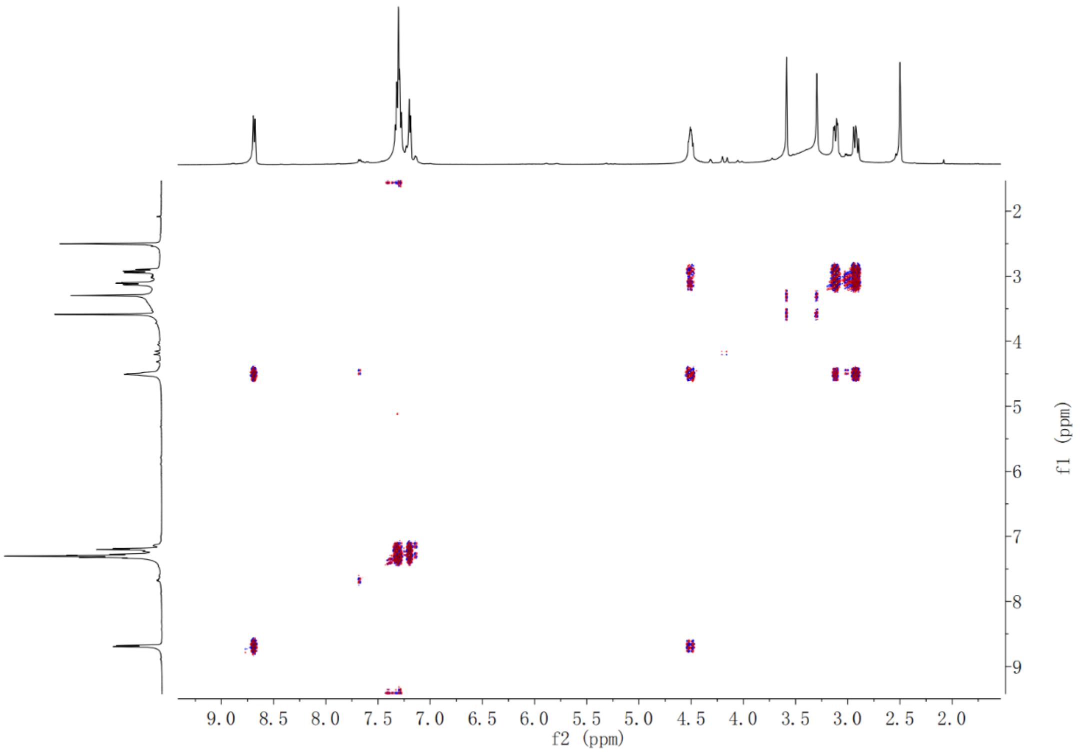

Supplementary Fig. 263. $^{1}H$-$^{1}H$ COSY spectrum of compound (2*S*,3*S*)-*t*-ES-a28 in DMSO-$d_6$

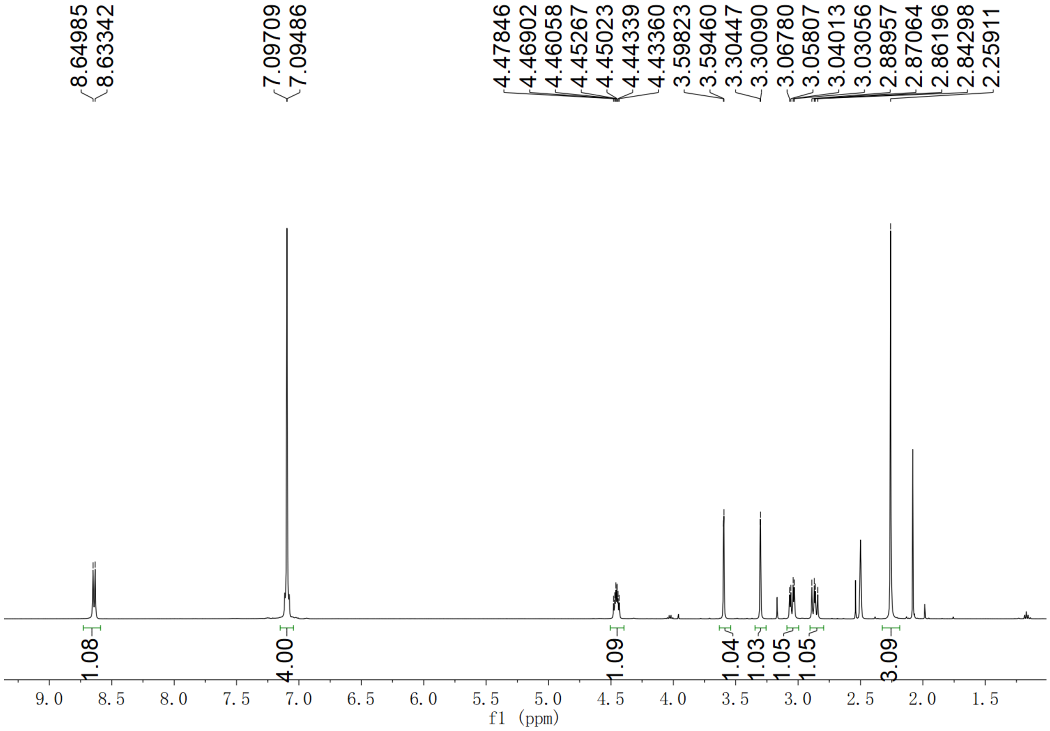

Supplementary Fig. 264. $^{1}H$ NMR spectrum of compound (2*S*,3*S*)-*t*-ES-a29 in DMSO-$d_6$

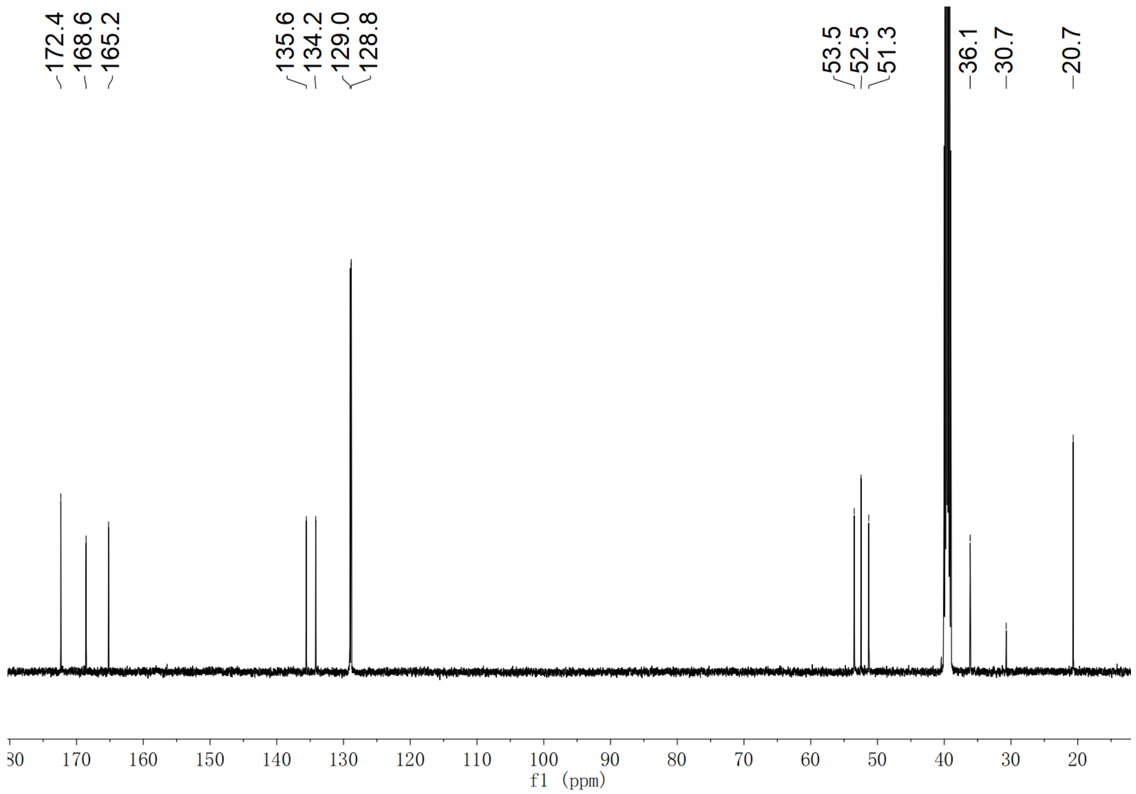

Supplementary Fig. 265. $^{13}$C NMR spectrum of compound (2*S*,3*S*)-*t*-ES-a29 in DMSO-$d_6$

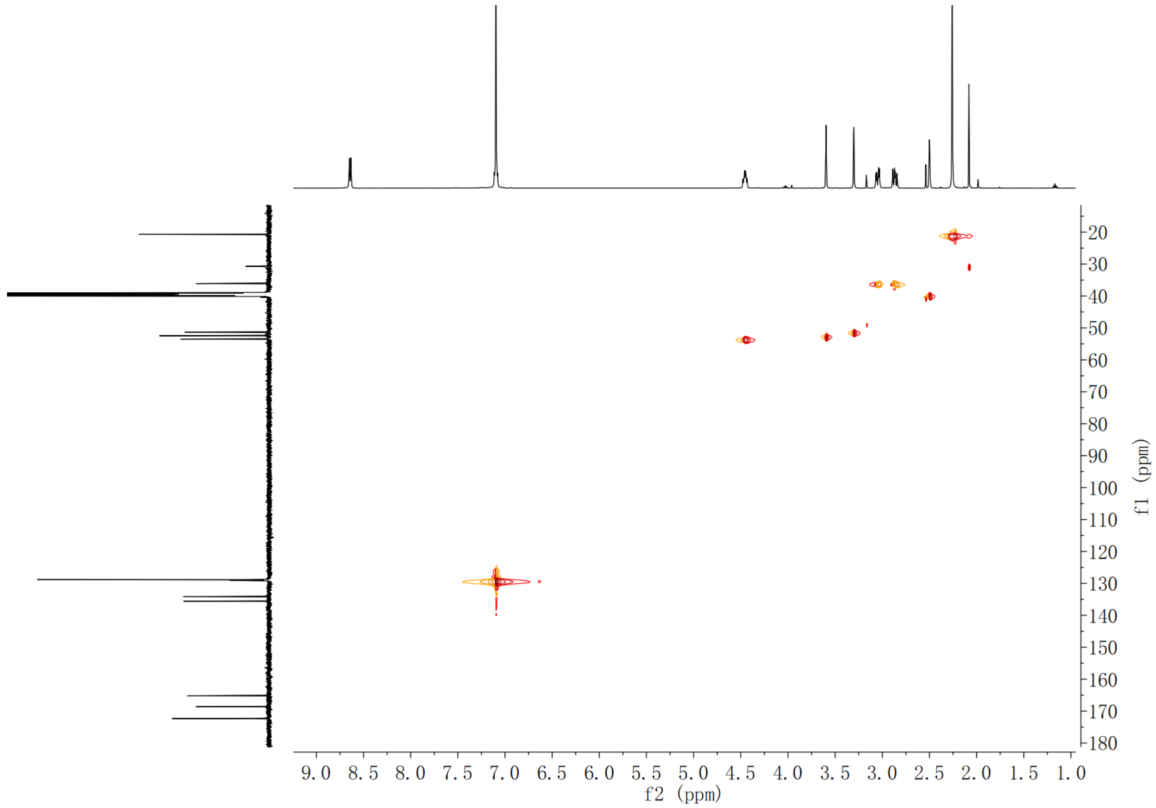

Supplementary Fig. 266. HSQC spectrum of compound (2*S*,3*S*)-*t*-ES-a29 in DMSO-$d_6$

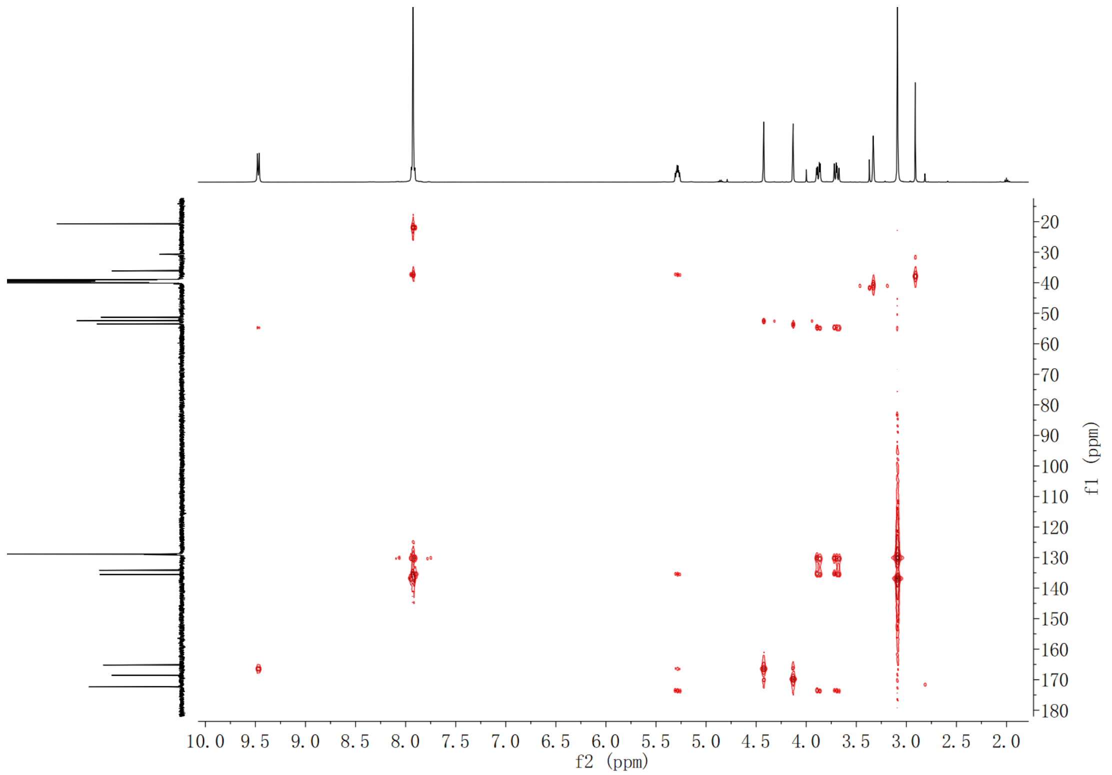

**Supplementary Fig. 267. HMBC spectrum of compound (2*S*,3*S*)-*t*-ES-a29 in DMSO-$d_6$**

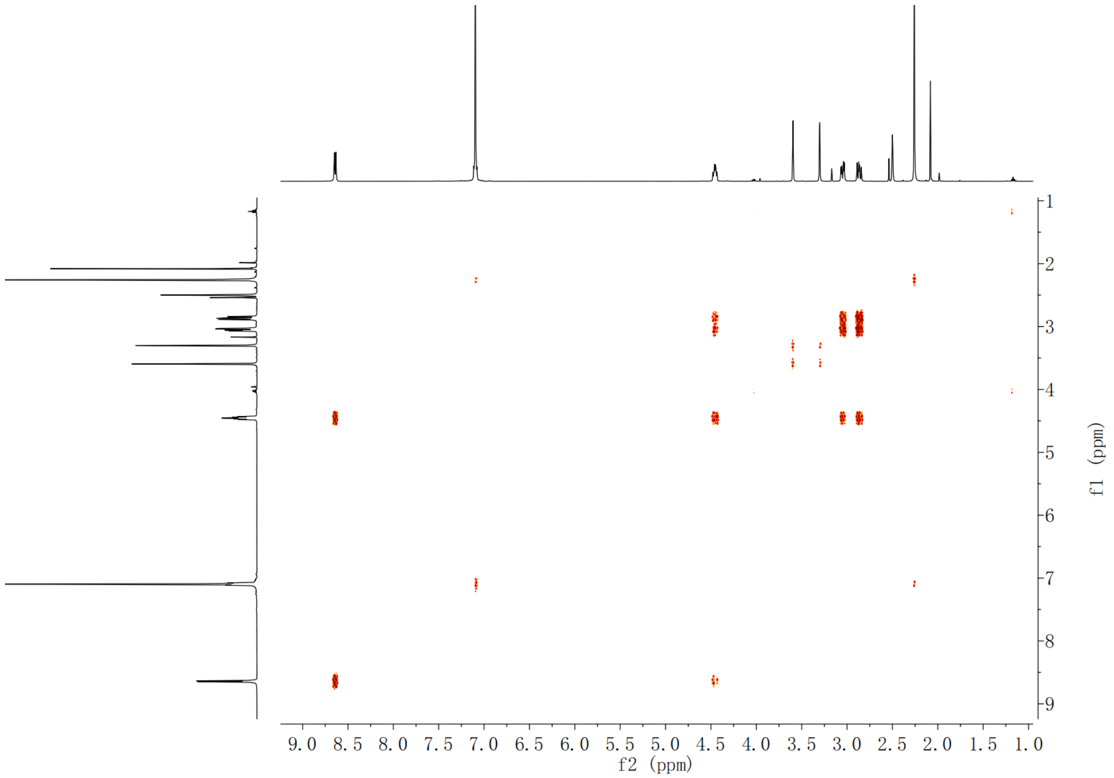

**Supplementary Fig. 268. $^1$H-$^1$H COSY spectrum of compound (2*S*,3*S*)-*t*-ES-a29 in DMSO-$d_6$**

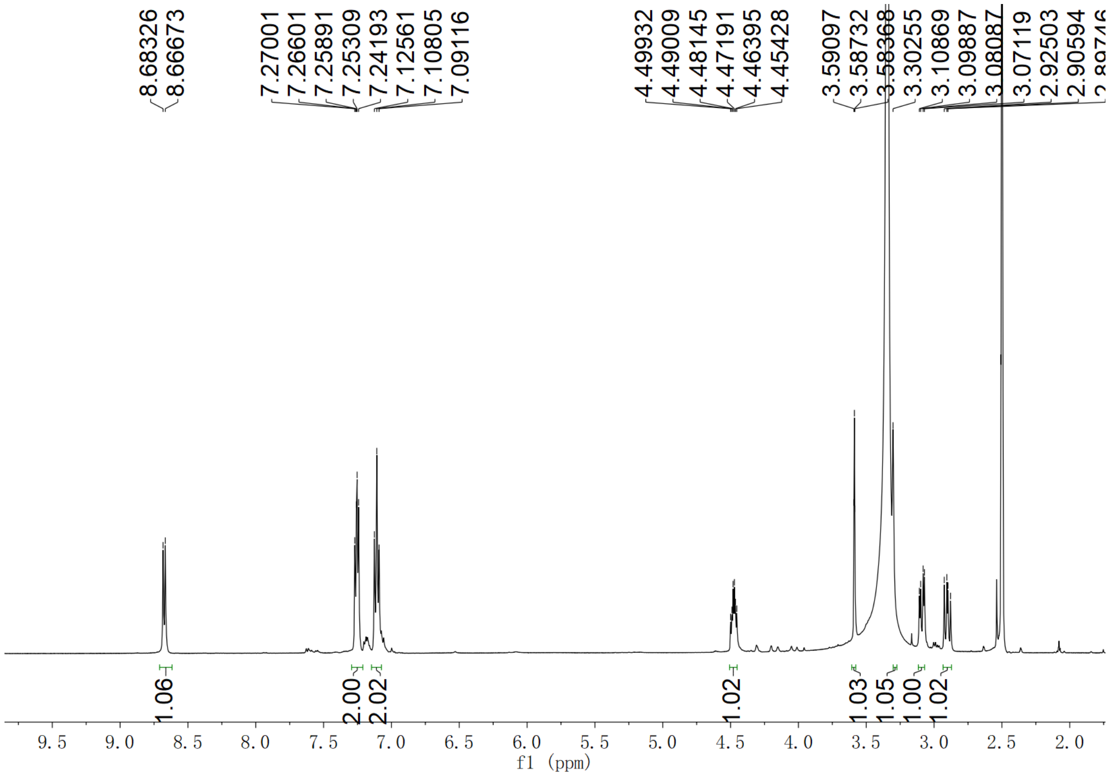

Supplementary Fig. 269. $^{1}$H NMR spectrum of compound (2*S*,3*S*)-*t*-ES-a30 in DMSO-$d_6$

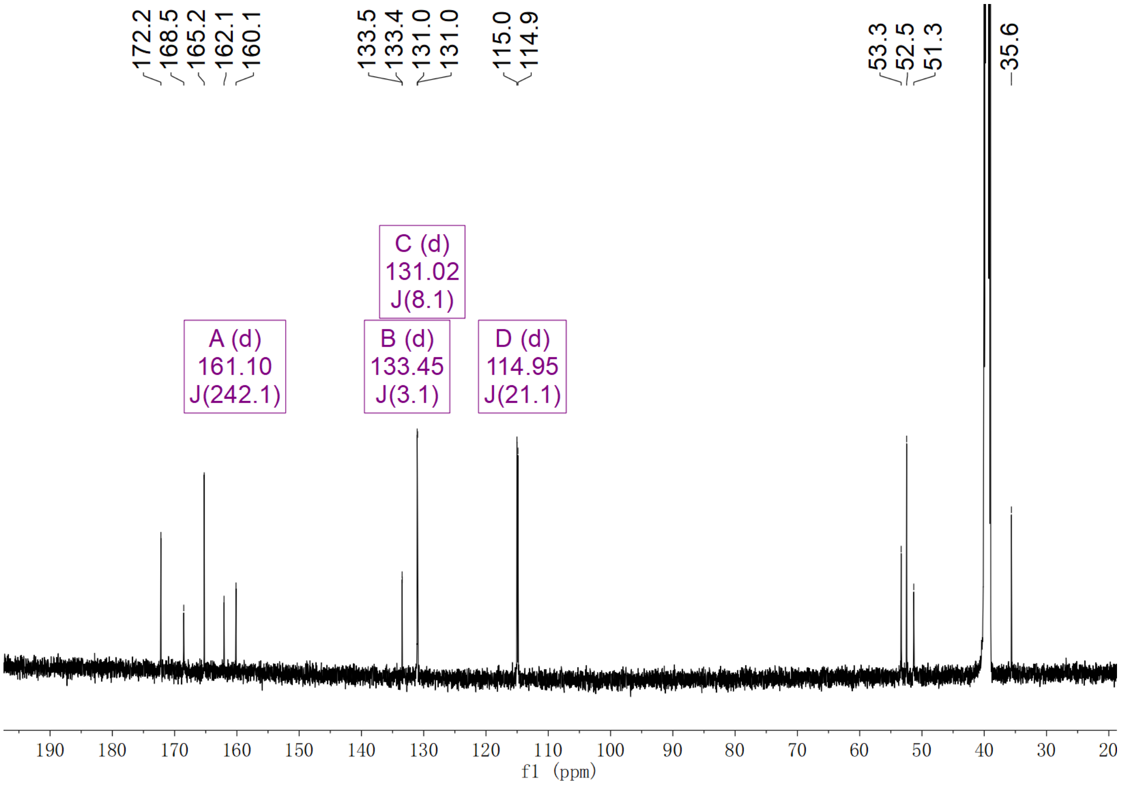

Supplementary Fig. 270. $^{13}$C NMR spectrum of compound (2*S*,3*S*)-*t*-ES-a30 in DMSO-$d_6$

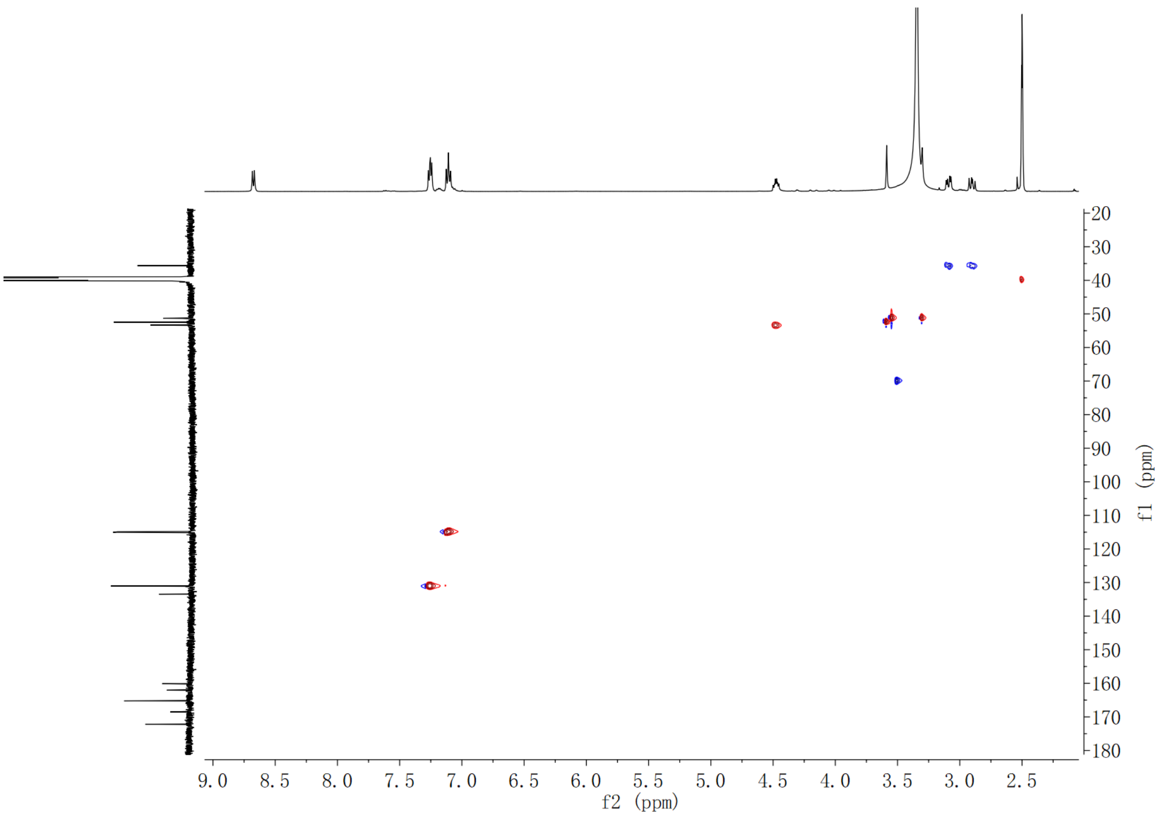

**Supplementary Fig. 271. HSQC spectrum of compound (2*S*,3*S*)-*t*-ES-a30 in DMSO-$d_6$**

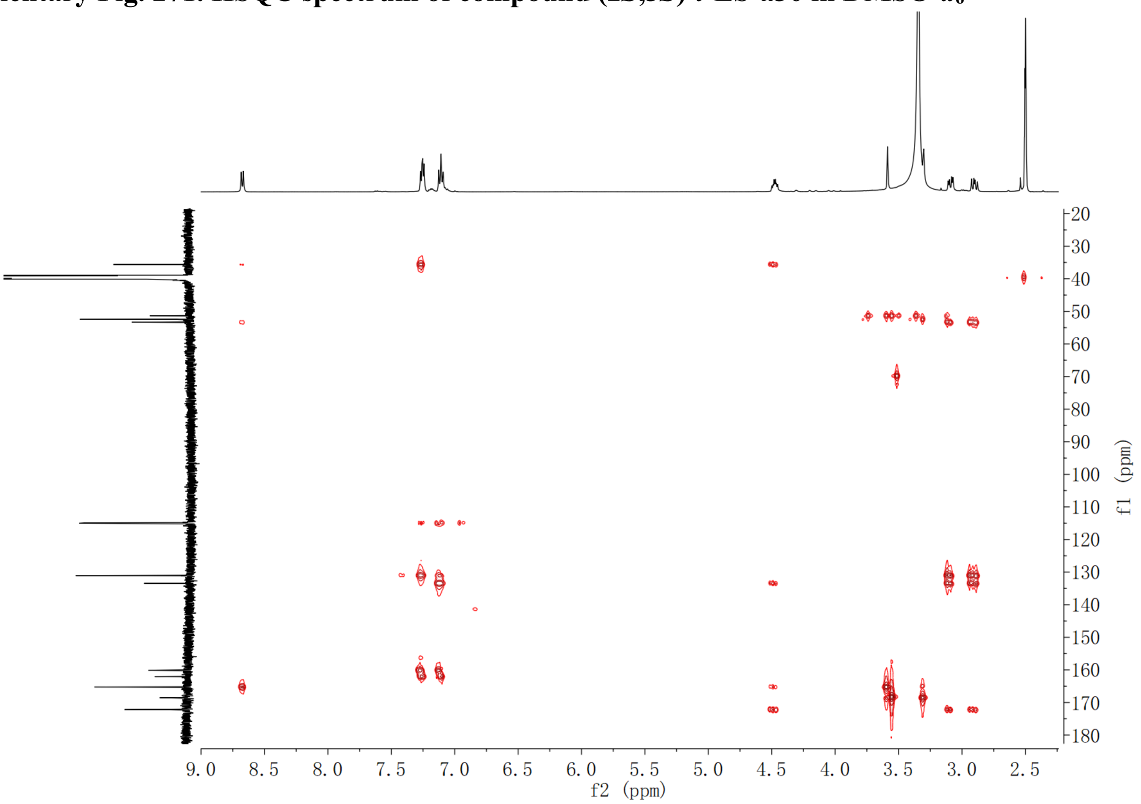

**Supplementary Fig. 272. HMBC spectrum of compound (2*S*,3*S*)-*t*-ES-a30 in DMSO-$d_6$**

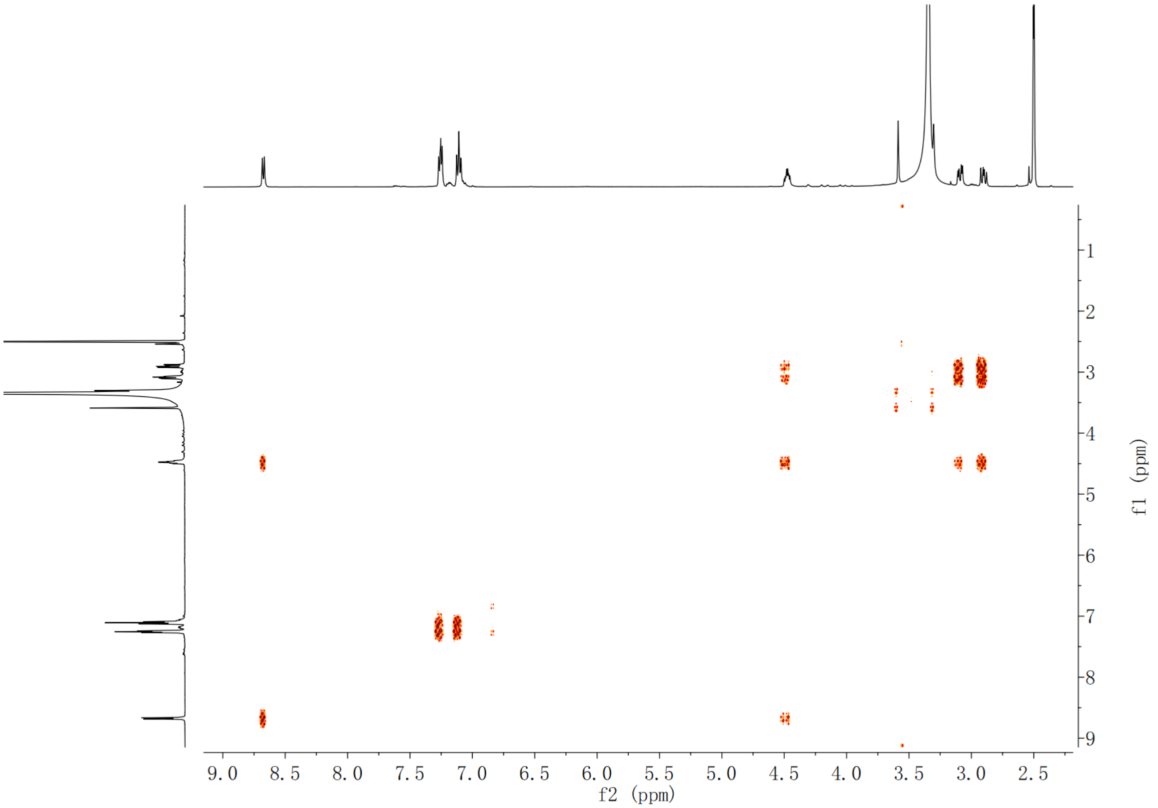

Supplementary Fig. 273. $^{1}$H-$^{1}$H COSY spectrum of compound (2*S*,3*S*)-*t*-ES-a30 in DMSO-$d_6$

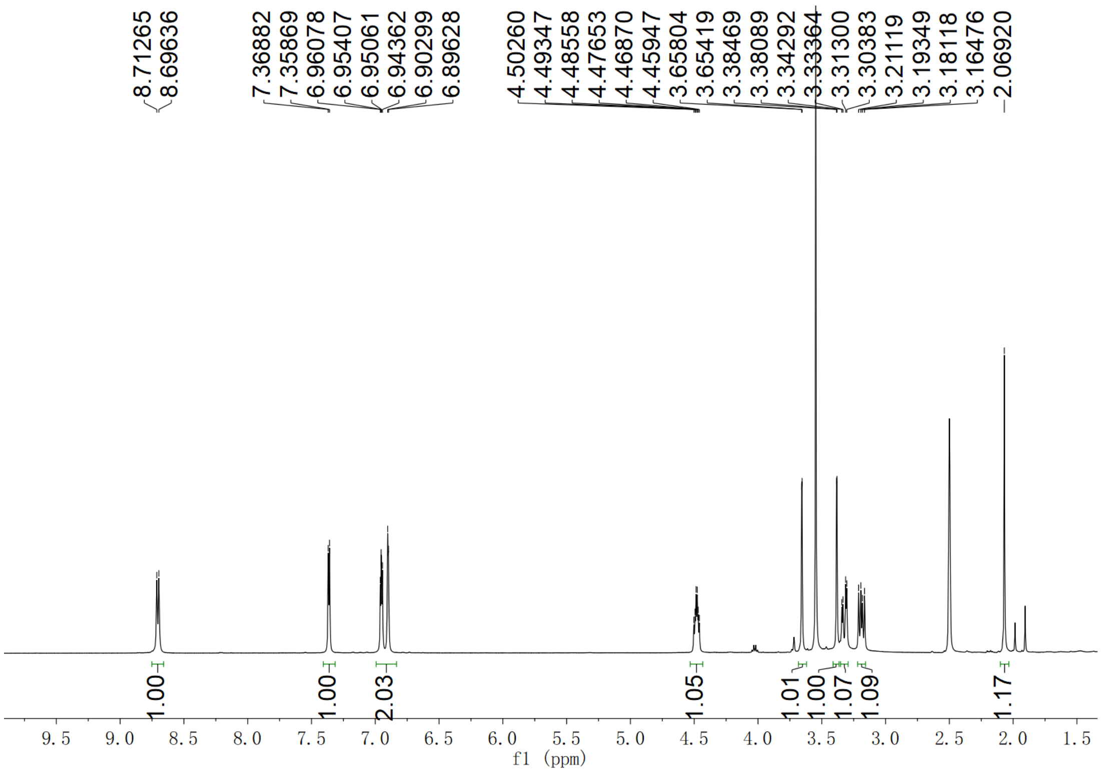

Supplementary Fig. 274. $^{1}$H NMR spectrum of compound (2*S*,3*S*)-*t*-ES-a31 in DMSO-$d_6$

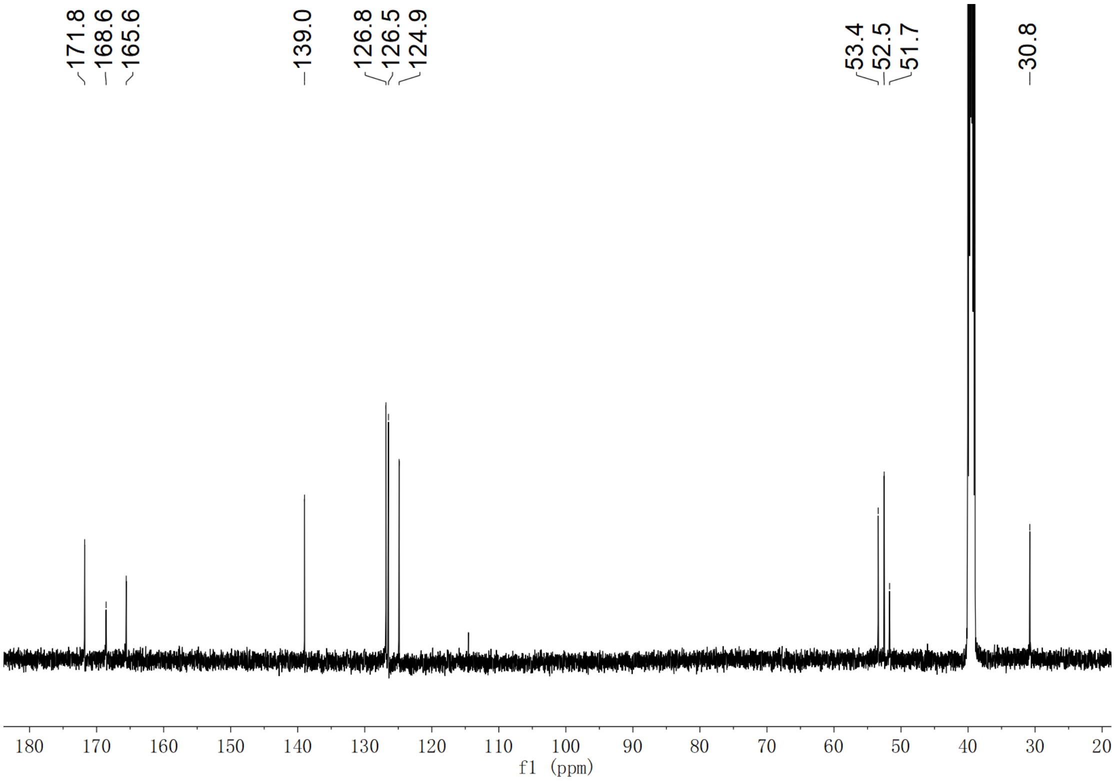

**Supplementary Fig. 275. $^{13}$C NMR spectrum of compound (2*S*,3*S*)-*t*-ES-a31 in DMSO-$d_6$**

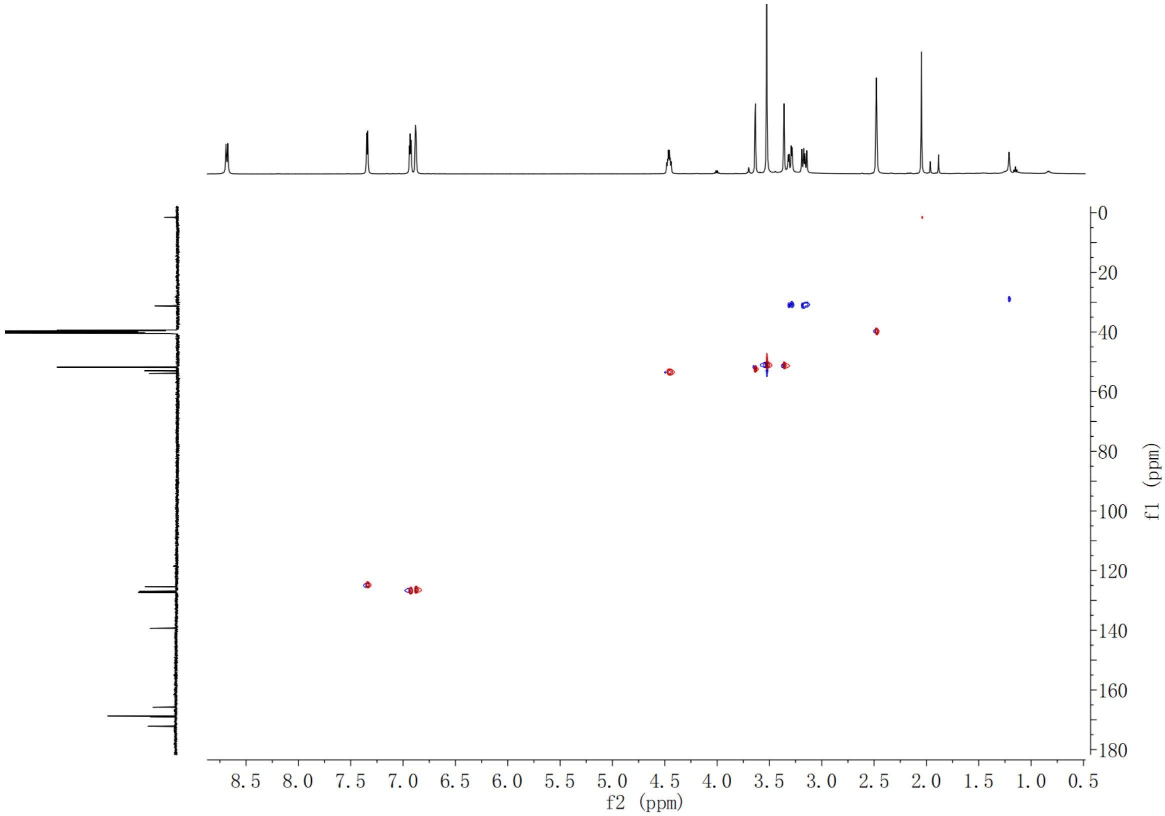

**Supplementary Fig. 276. HSQC spectrum of compound (2*S*,3*S*)-*t*-ES-a31 in DMSO-$d_6$**

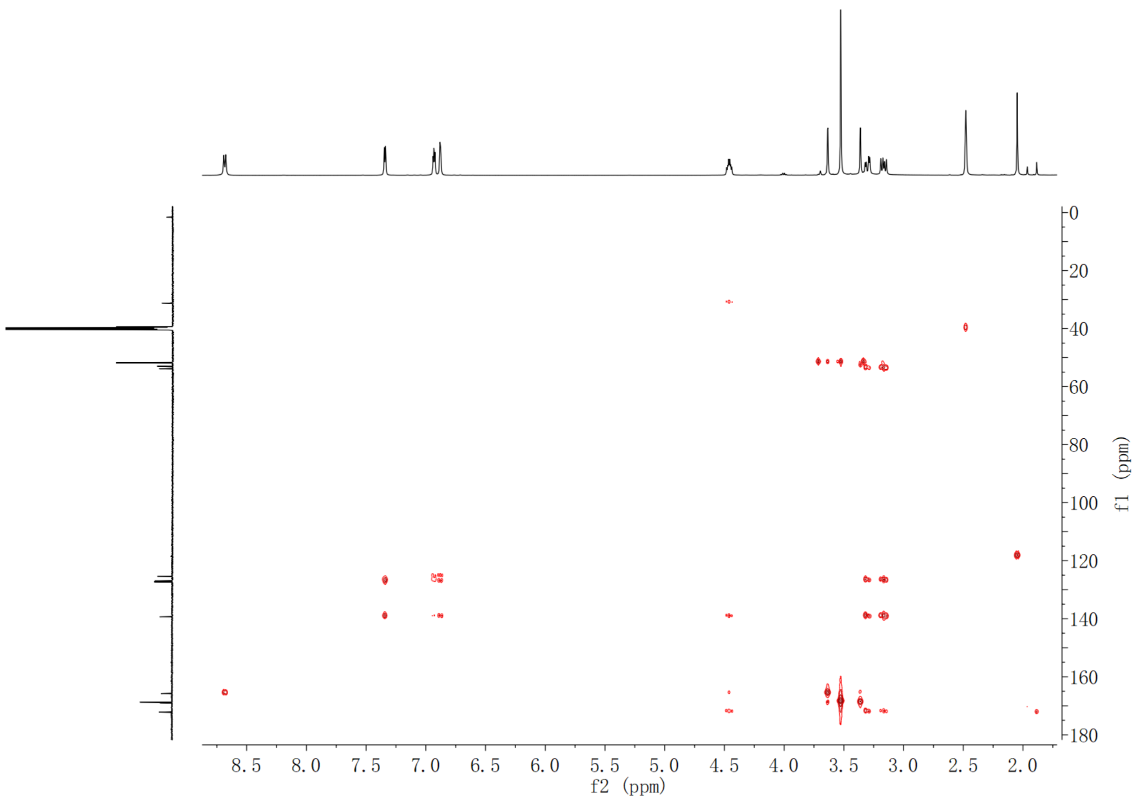

**Supplementary Fig. 277. HMBC spectrum of compound (2*S*,3*S*)-*t*-ES-a31 in DMSO-*d*$_6$**

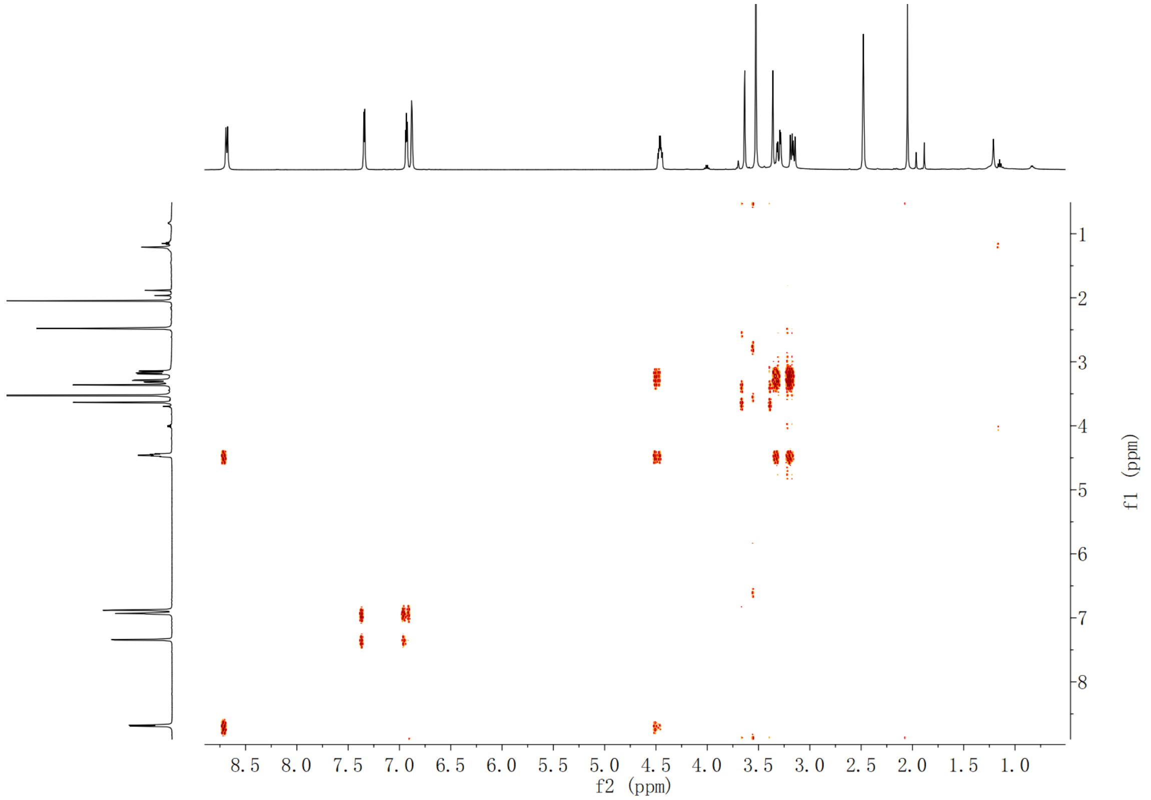

**Supplementary Fig. 278. $^1$H-$^1$H COSY spectrum of compound (2*S*,3*S*)-*t*-ES-a31 in DMSO-*d*$_6$**

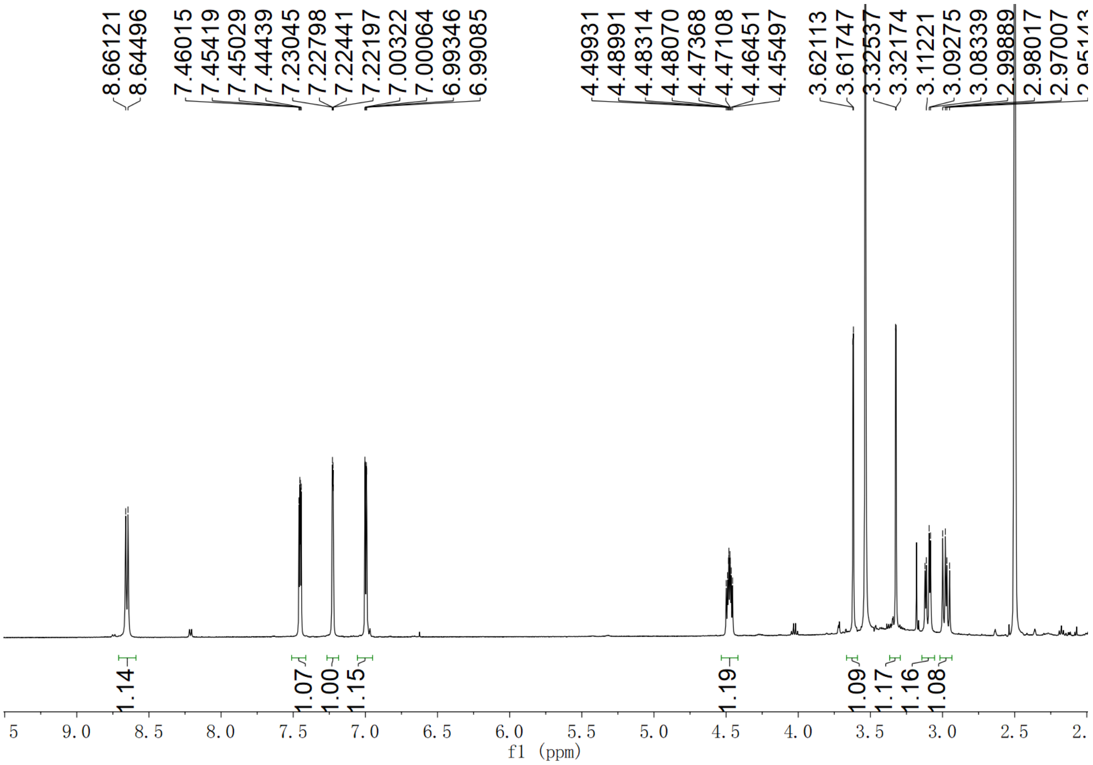

Supplementary Fig. 279. $^{1}$H NMR spectrum of compound (2*S*,3*S*)-*t*-ES-a32 in DMSO-$d_6$

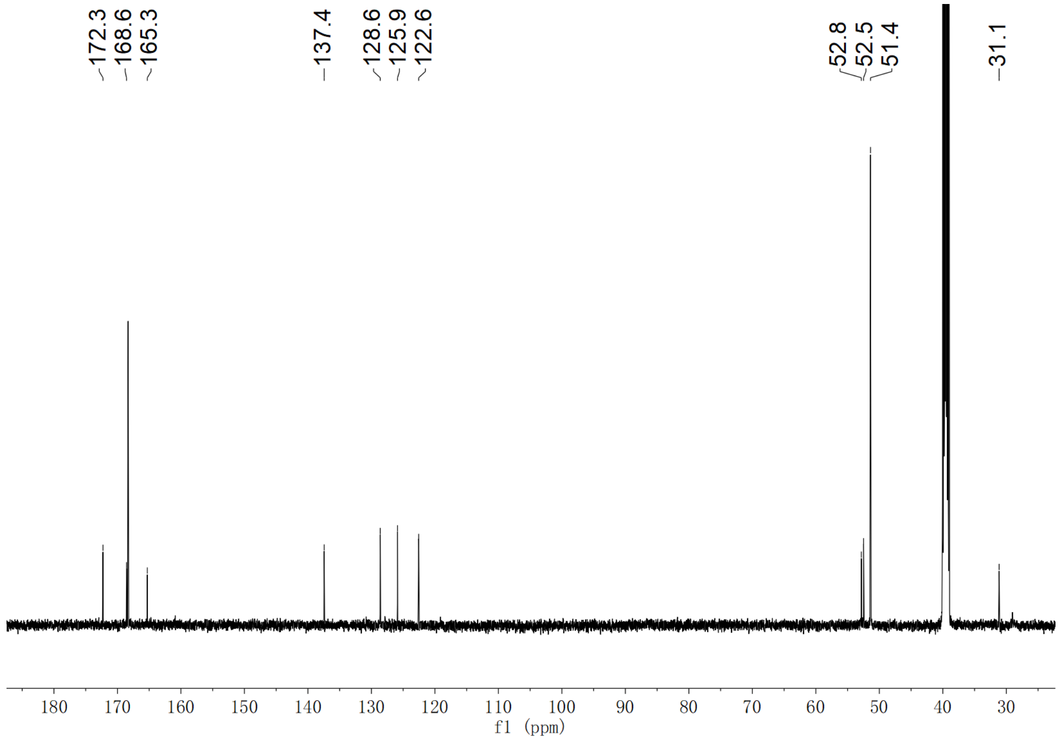

Supplementary Fig. 280. $^{13}$C NMR spectrum of compound (2*S*,3*S*)-*t*-ES-a32 in DMSO-$d_6$

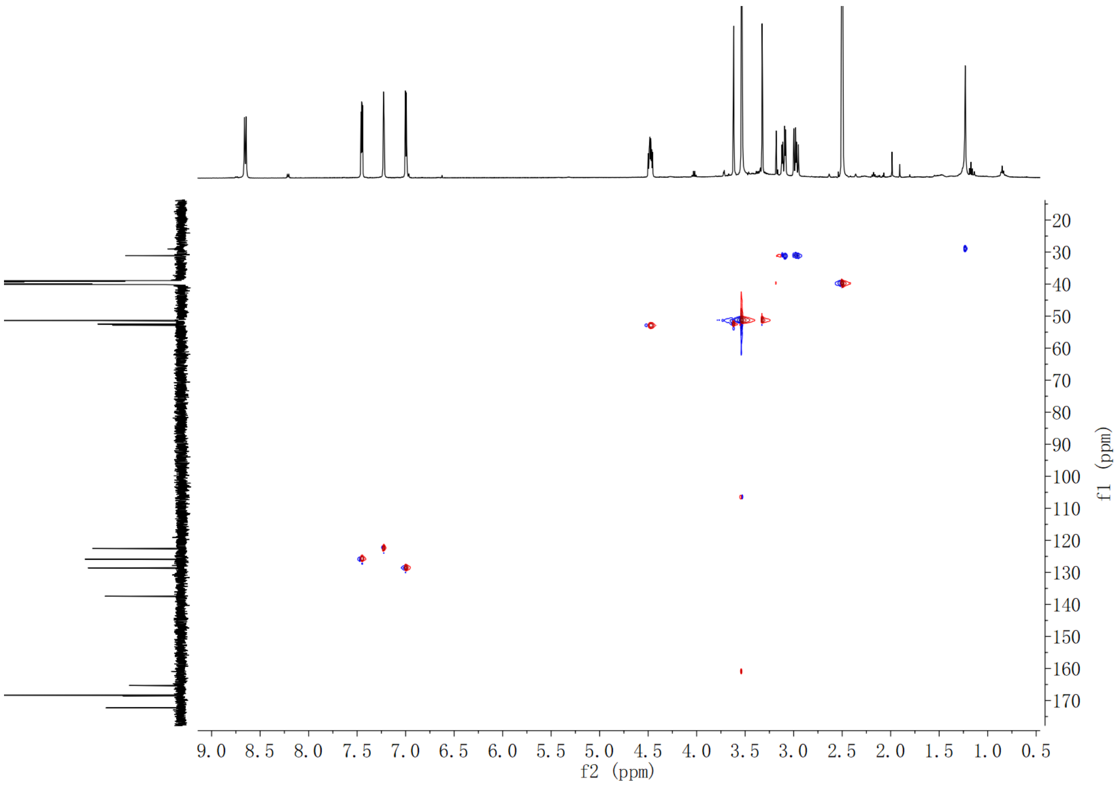

**Supplementary Fig. 281. HSQC spectrum of compound (2*S*,3*S*)-*t*-ES-a32 in DMSO-$d_6$**

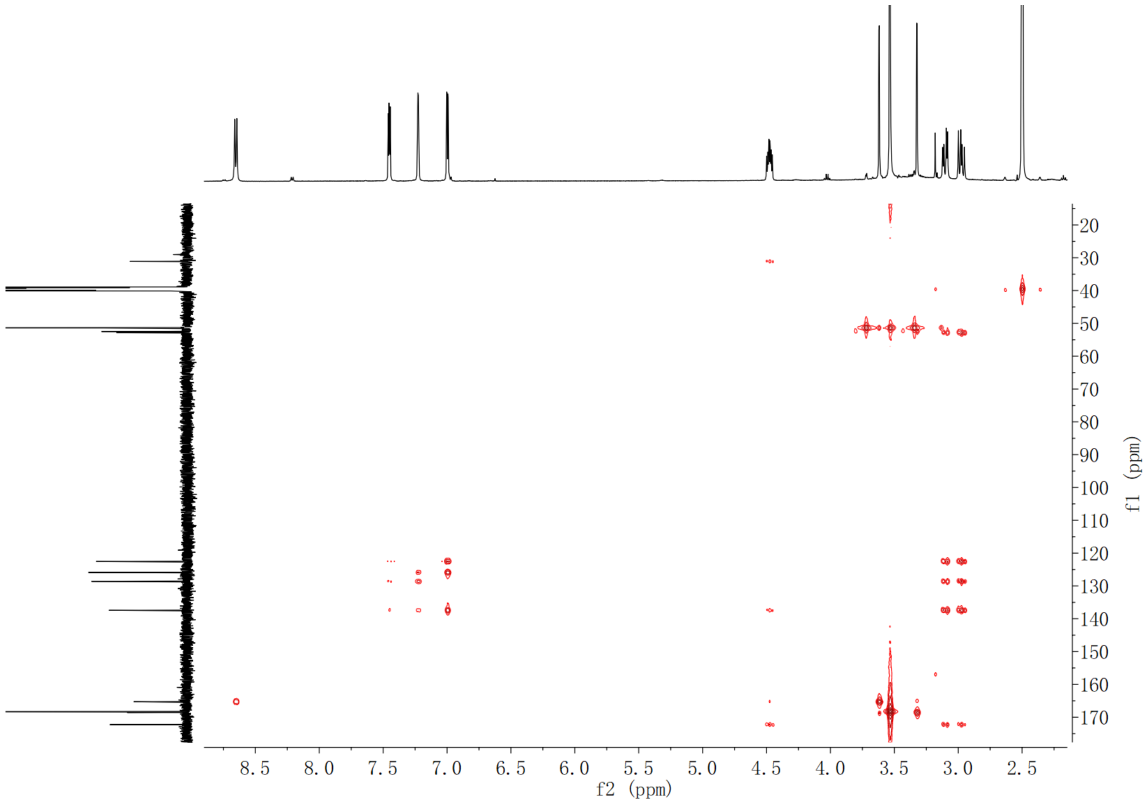

**Supplementary Fig. 282. HMBC spectrum of compound (2*S*,3*S*)-*t*-ES-a32 in DMSO-$d_6$**

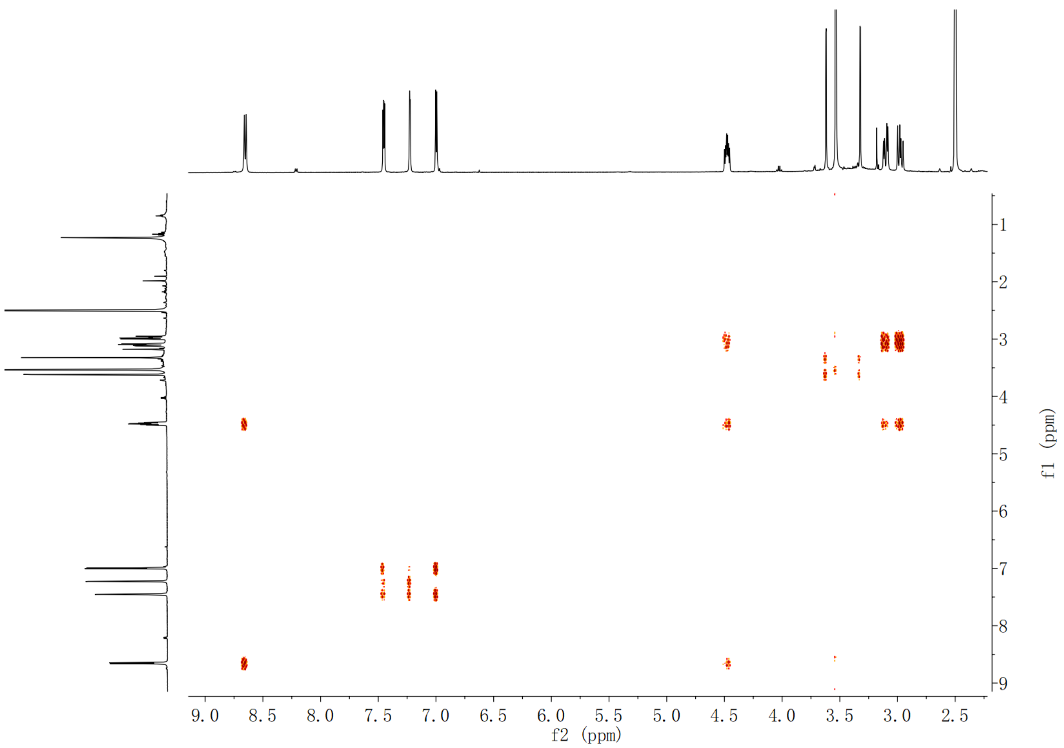

**Supplementary Fig. 283.** $^{1}H$-$^{1}H$ **COSY spectrum of compound (2*S*,3*S*)-*t*-ES-a32 in DMSO-*d*$_6$**

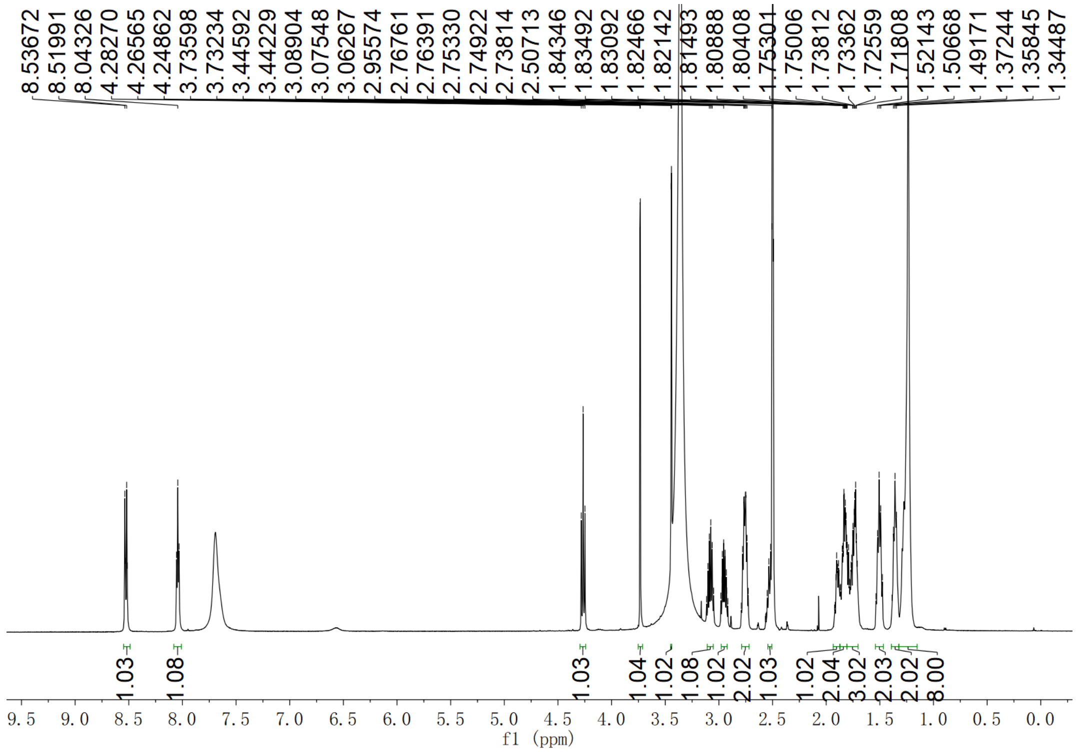

**Supplementary Fig. 284.** $^{1}H$ **NMR spectrum of compound (2*S*,3*S*)-*t*-ES-a9-b7 in DMSO-*d*$_6$**

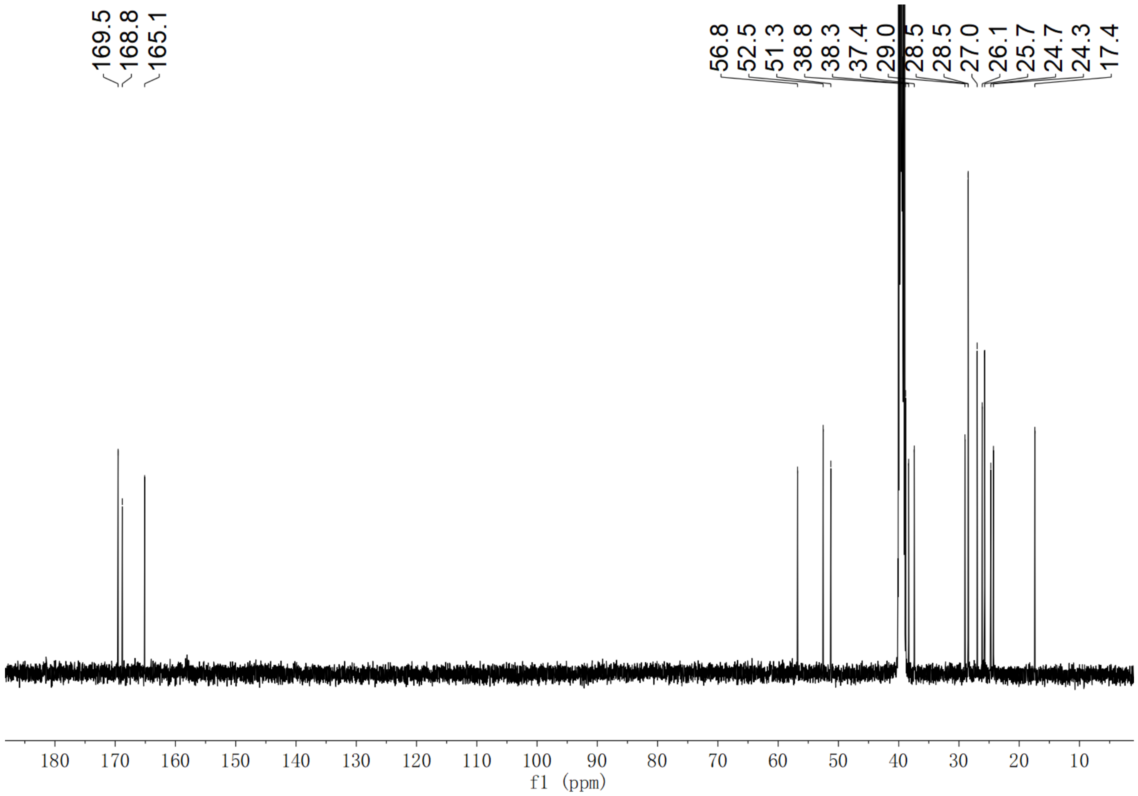

**Supplementary Fig. 285. $^{13}$C NMR spectrum of compound (2*S*,3*S*)-*t*-ES-a9-b7 in DMSO-$d_6$**

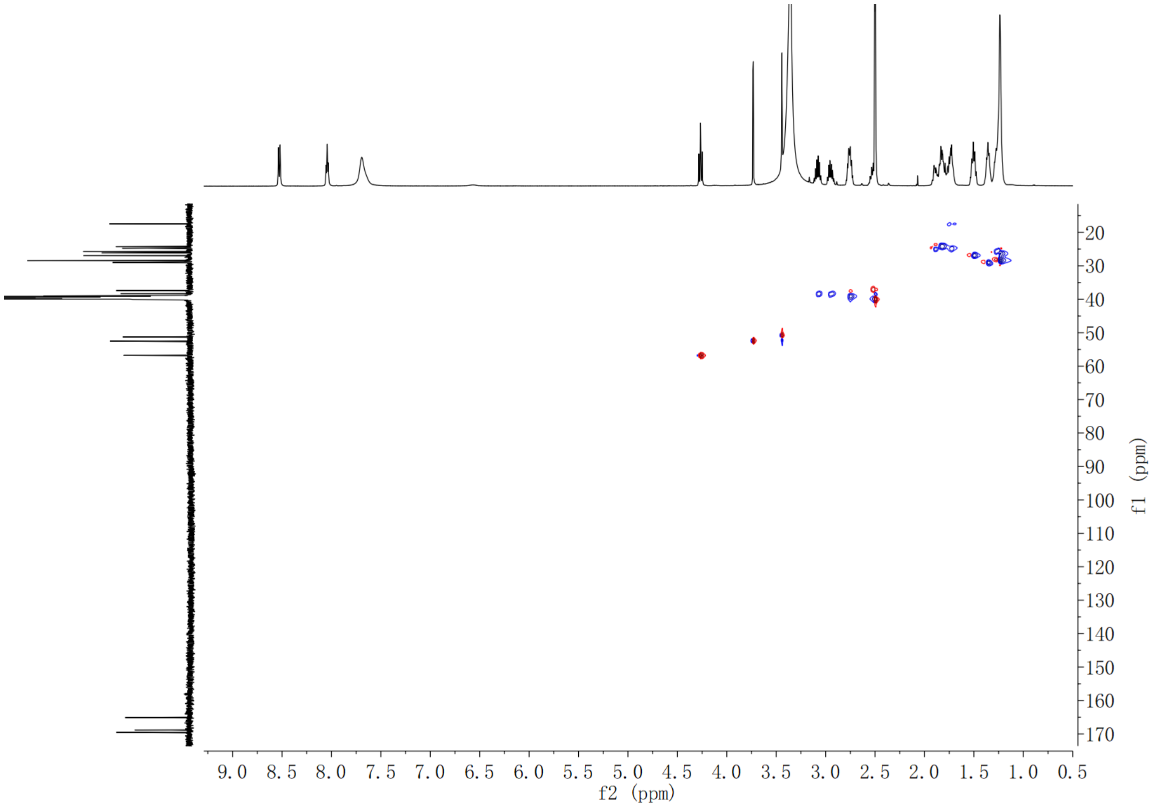

**Supplementary Fig. 286. HSQC spectrum of compound (2*S*,3*S*)-*t*-ES-a9-b7 in DMSO-$d_6$**

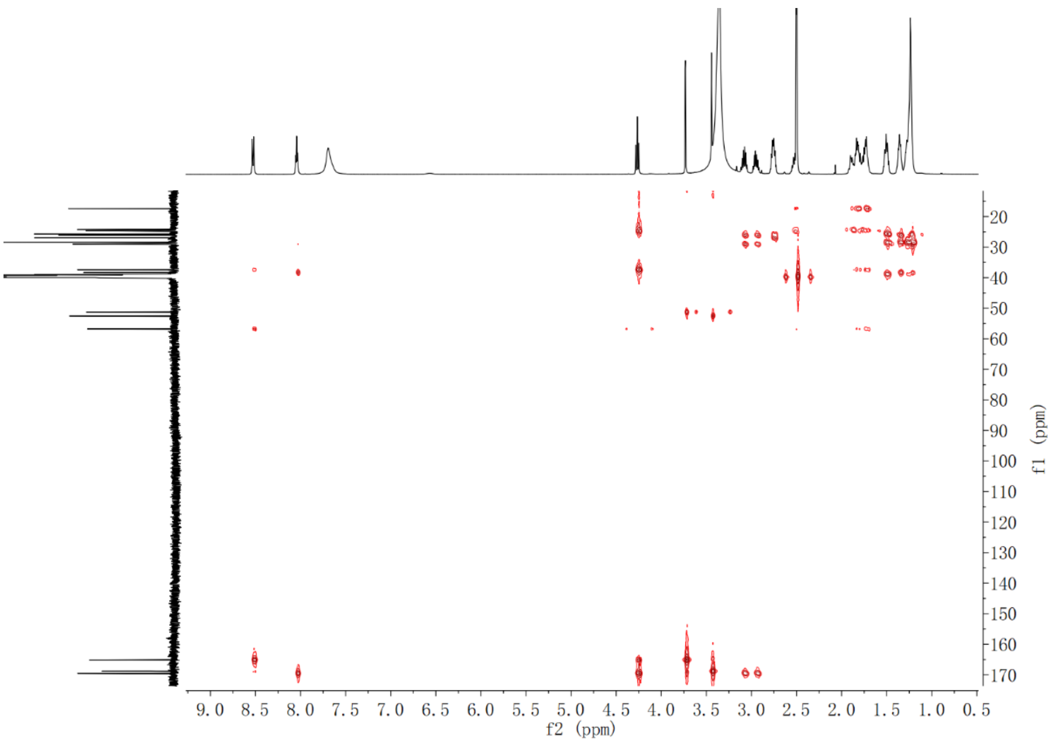

**Supplementary Fig. 287. HMBC spectrum of compound (2*S*,3*S*)-*t*-ES-a9-b7 in DMSO-$d_6$**

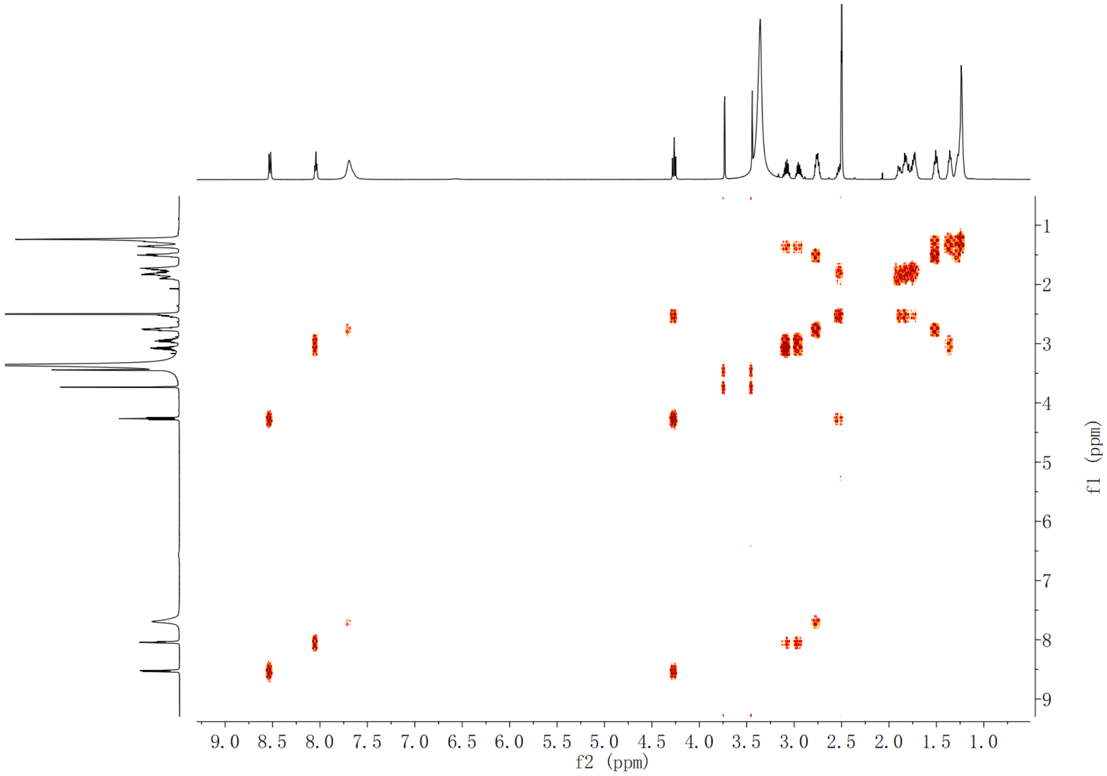

**Supplementary Fig. 288. $^1$H-$^1$H COSY spectrum of compound (2*S*,3*S*)-*t*-ES-a9-b7 in DMSO-$d_6$**

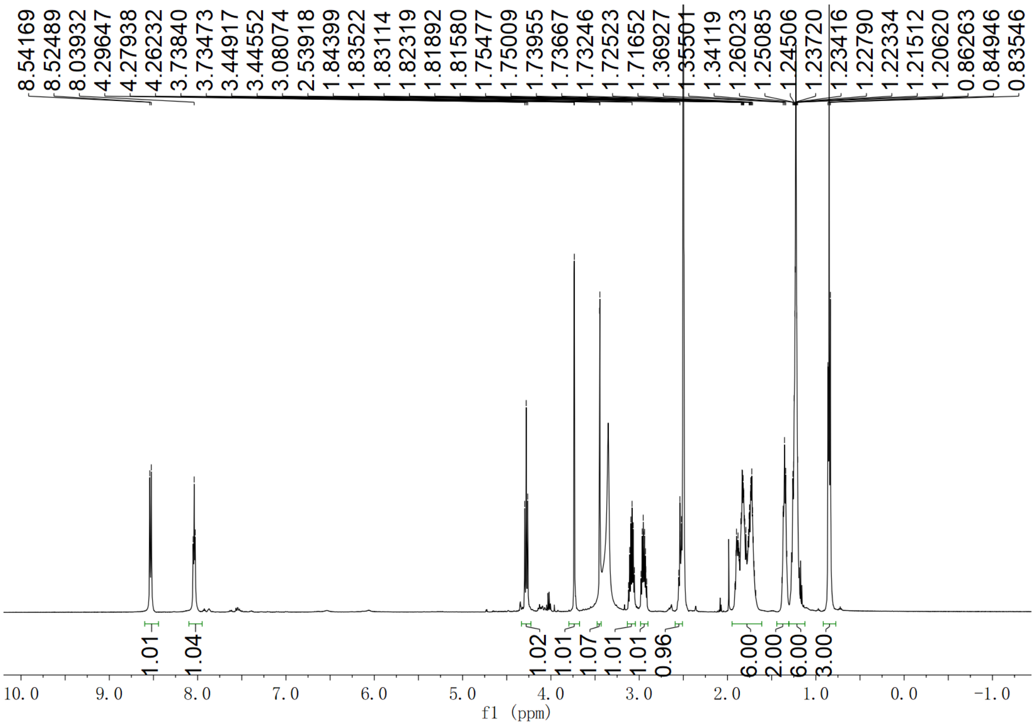

Supplementary Fig. 289. $^{1}H$ NMR spectrum of compound (2*S*,3*S*)-*t*-ES-a9-b12 in DMSO-$d_6$

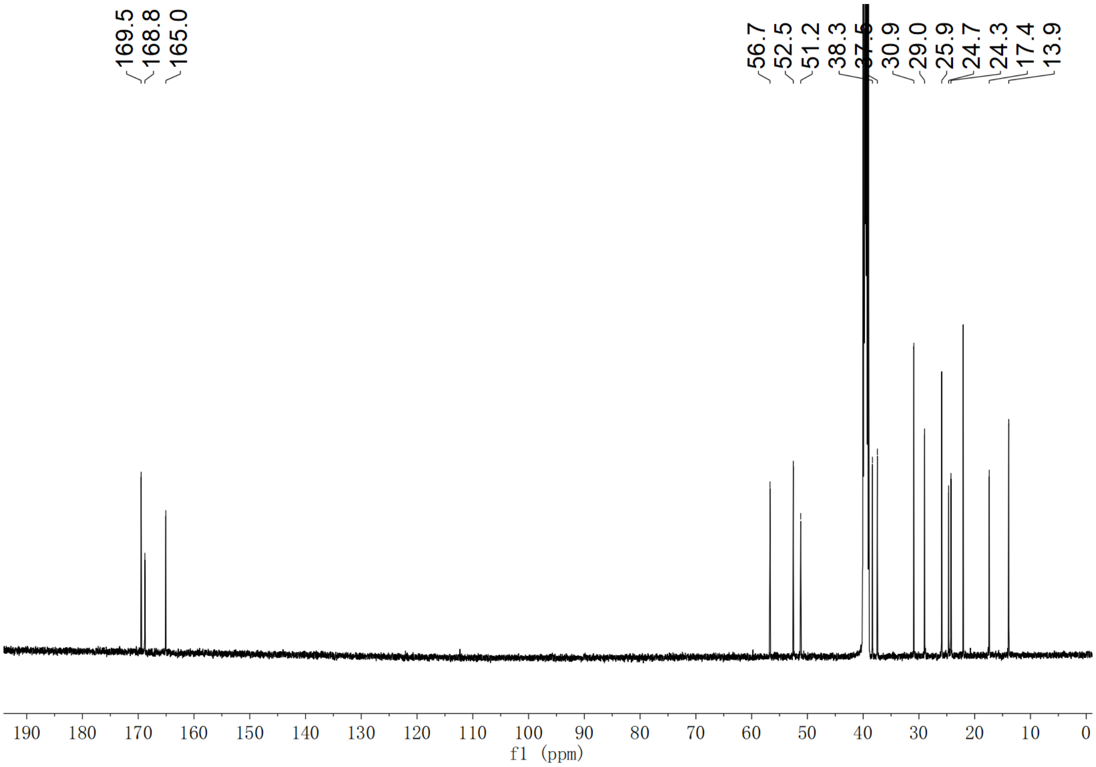

Supplementary Fig. 290. $^{13}C$ NMR spectrum of compound (2*S*,3*S*)-*t*-ES-a9-b12 in DMSO-$d_6$

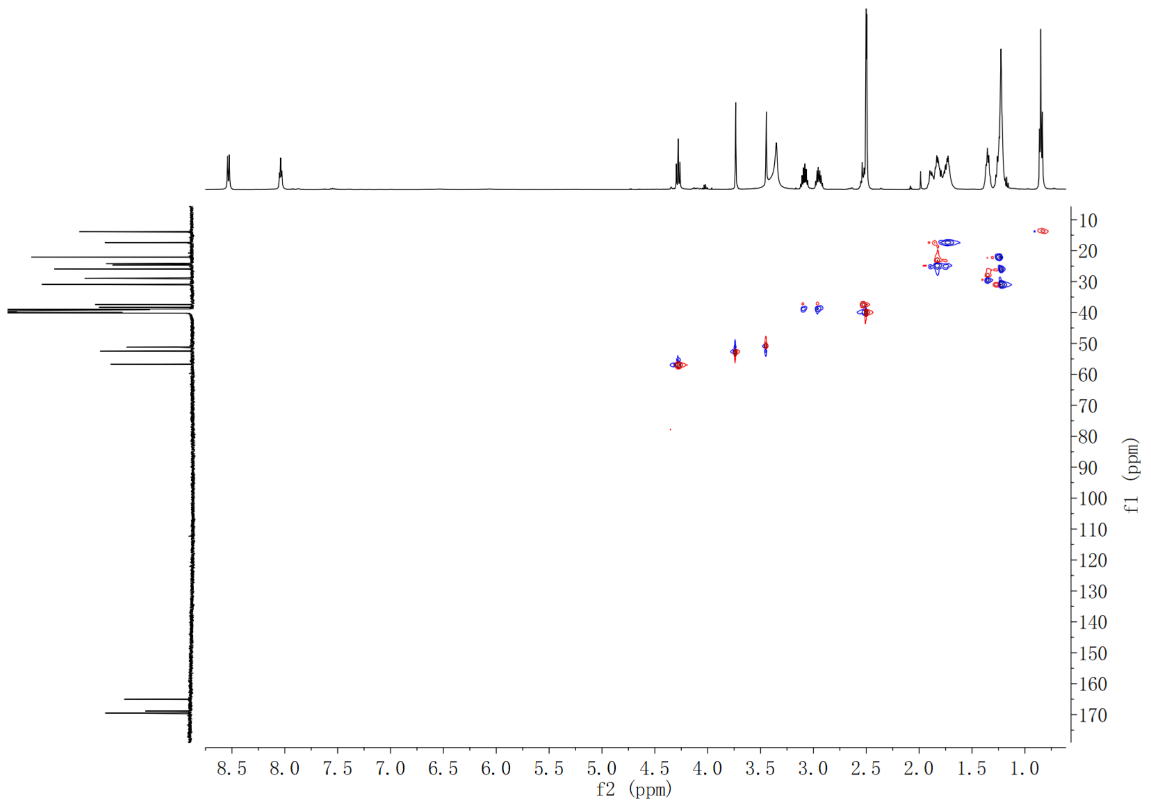

**Supplementary Fig. 291. HSQC spectrum of compound (2*S*,3*S*)-*t*-ES-a9-b12 in DMSO-$d_6$**

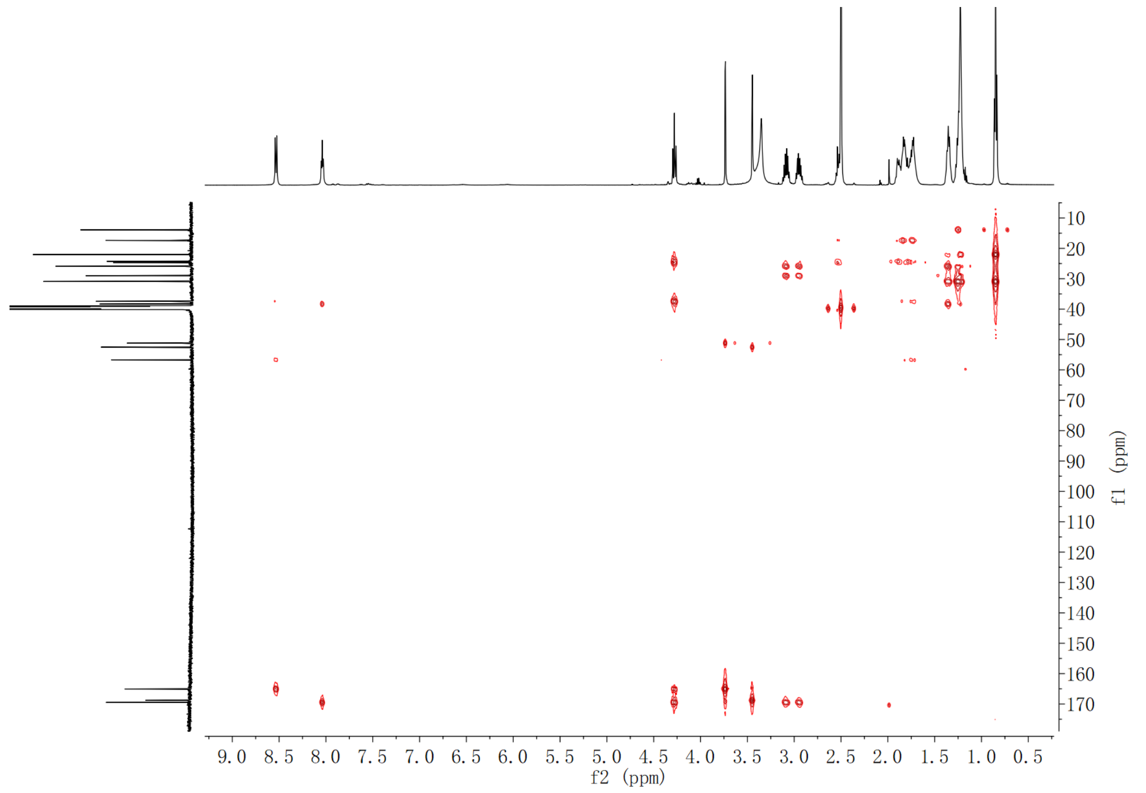

**Supplementary Fig. 292. HMBC spectrum of compound (2*S*,3*S*)-*t*-ES-a9-b12 in DMSO-$d_6$**

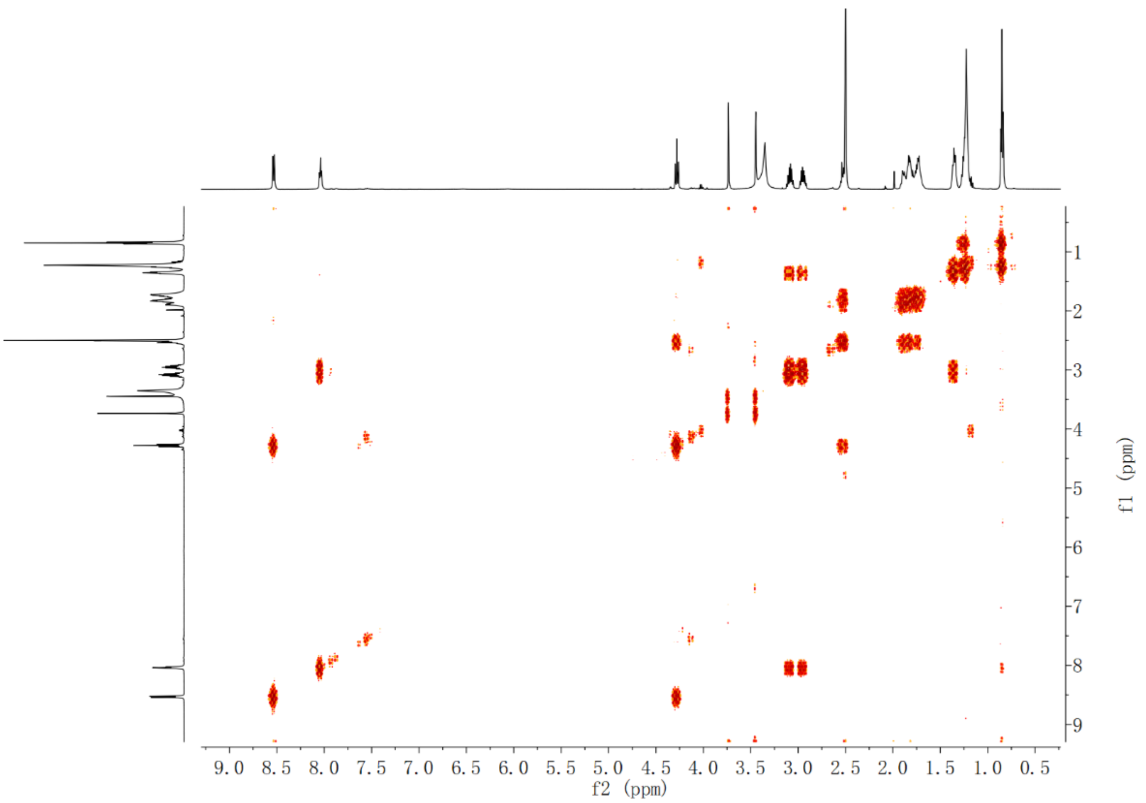

Supplementary Fig. 293. $^{1}$H-$^{1}$H COSY spectrum of compound (2*S*,3*S*)-*t*-ES-a9-b12 in DMSO-$d_6$

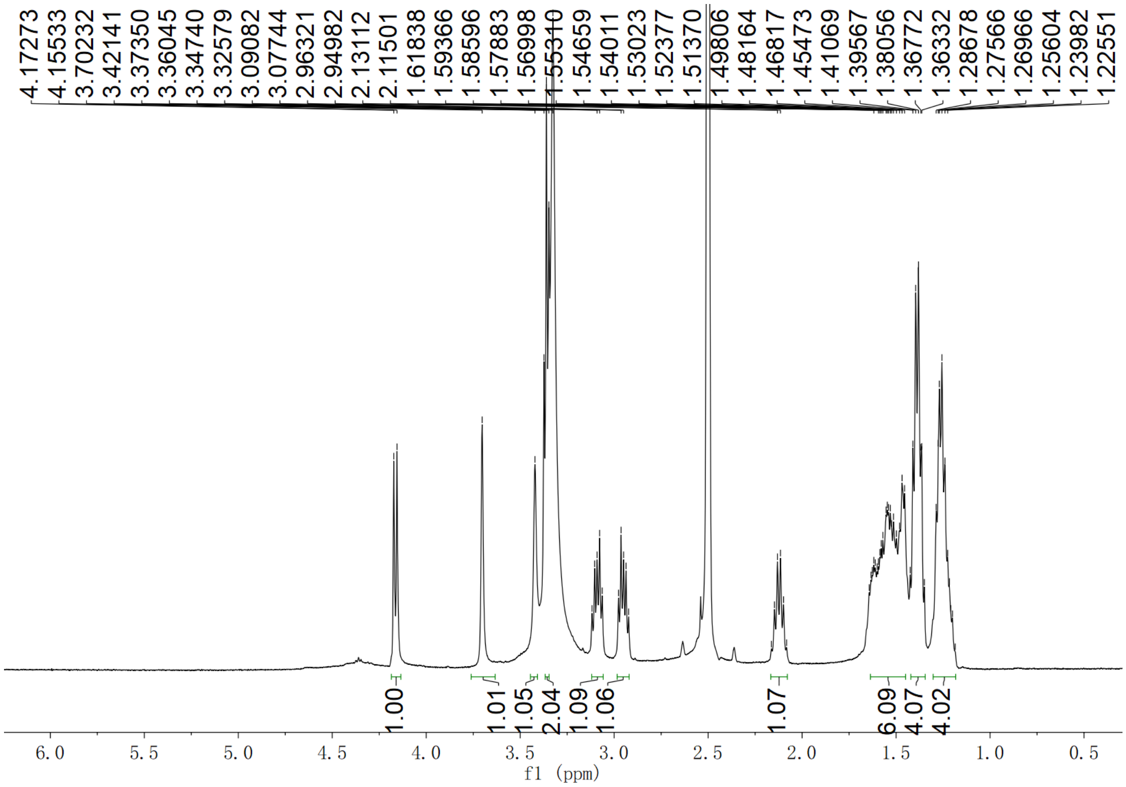

Supplementary Fig. 294. $^{1}$H NMR spectrum of compound (2*S*,3*S*)-*t*-ES-a10-b9 in DMSO-$d_6$

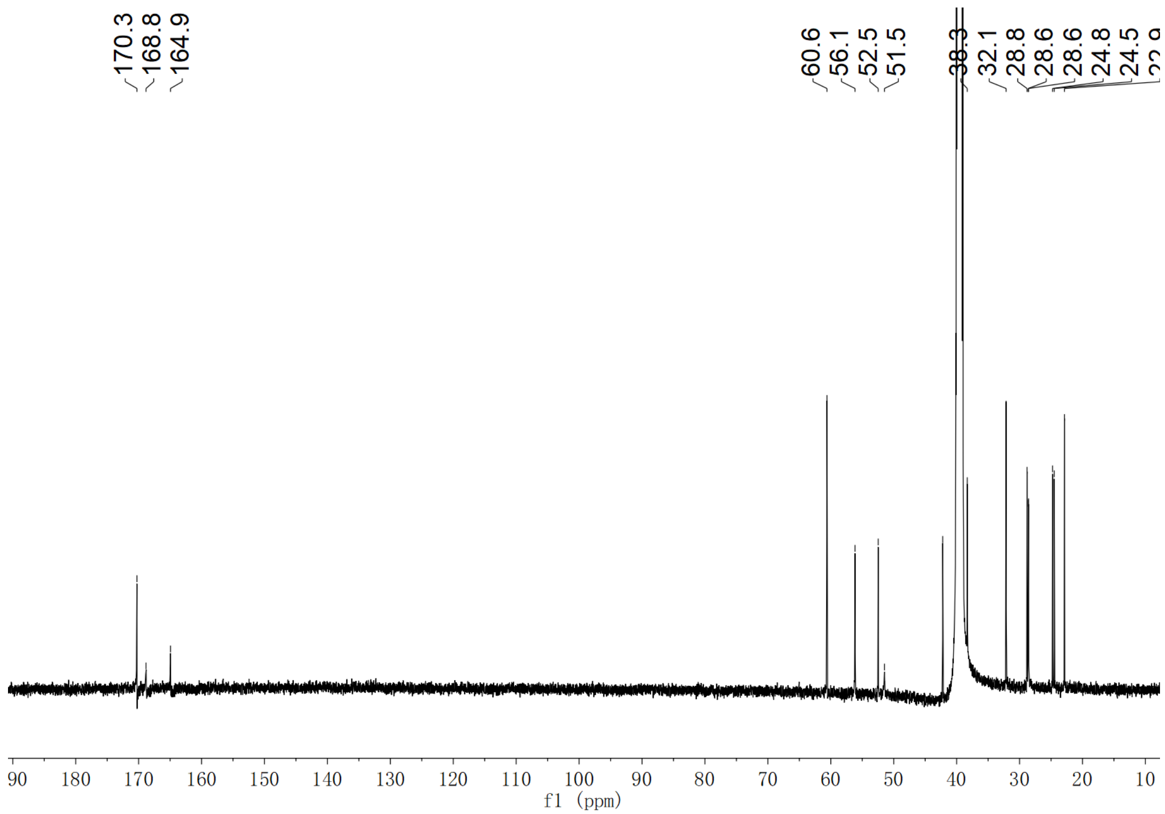

**Supplementary Fig. 295. $^{13}$C NMR spectrum of compound (2*S*,3*S*)-*t*-ES-a10-b9 in DMSO-$d_6$**

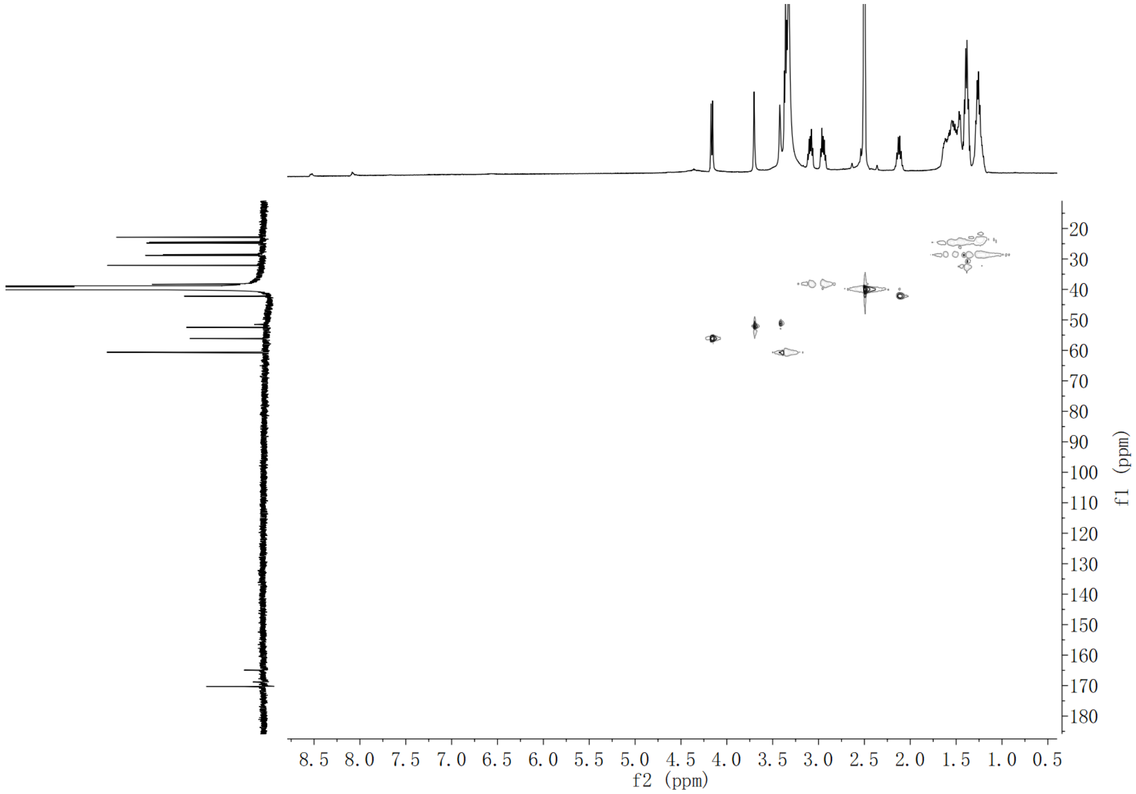

**Supplementary Fig. 296. HSQC spectrum of compound (2*S*,3*S*)-*t*-ES-a10-b9 in DMSO-$d_6$**

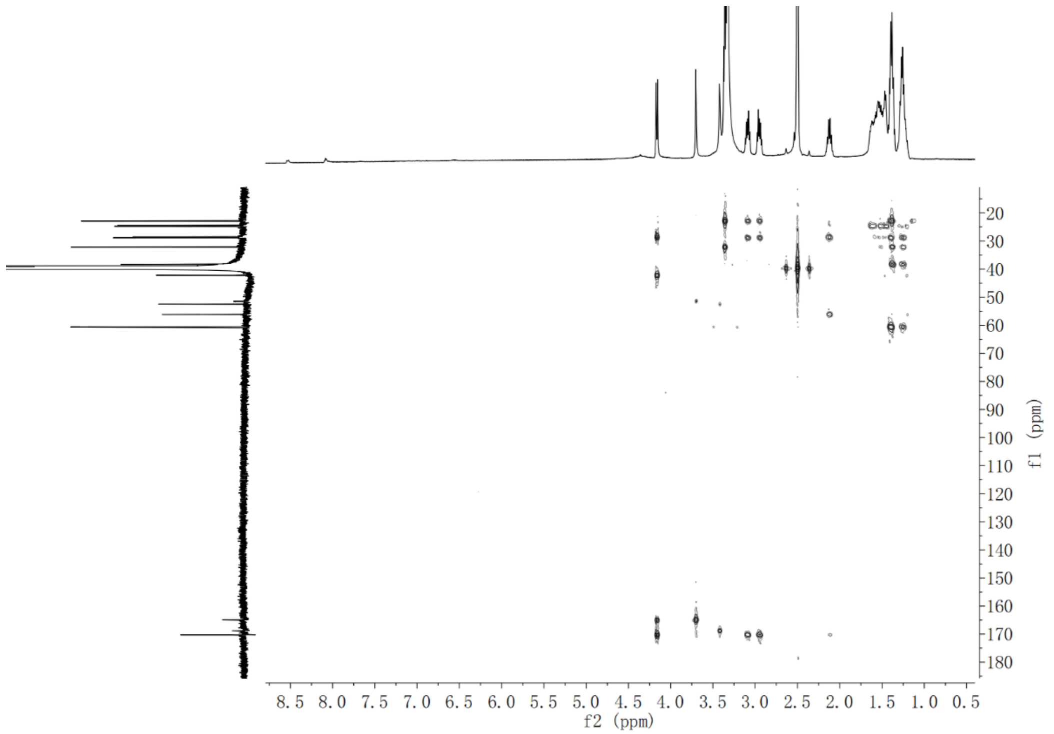

**Supplementary Fig. 297. HMBC spectrum of compound (2*S*,3*S*)-*t*-ES-a10-b9 in DMSO-*d*$_6$**

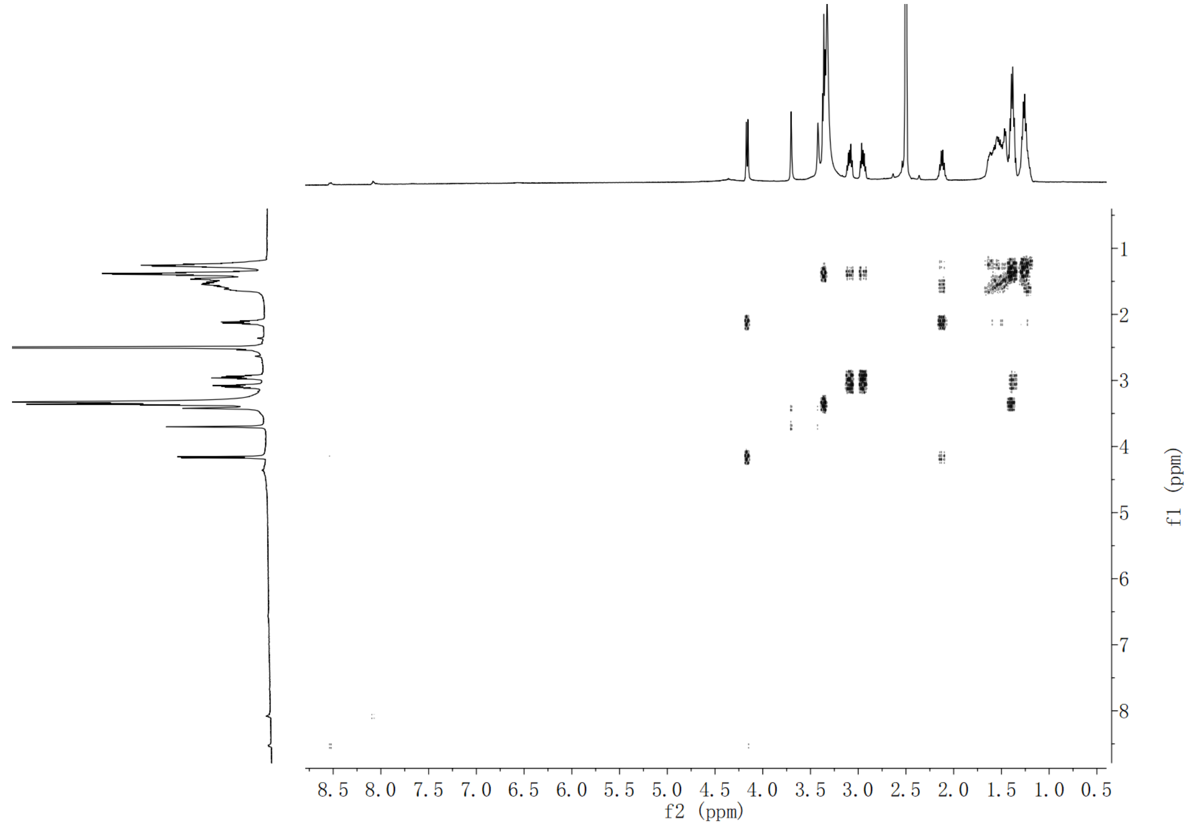

**Supplementary Fig. 298. $^1$H-$^1$H COSY spectrum of compound (2*S*,3*S*)-*t*-ES-a10-b9 in DMSO-*d*$_6$**

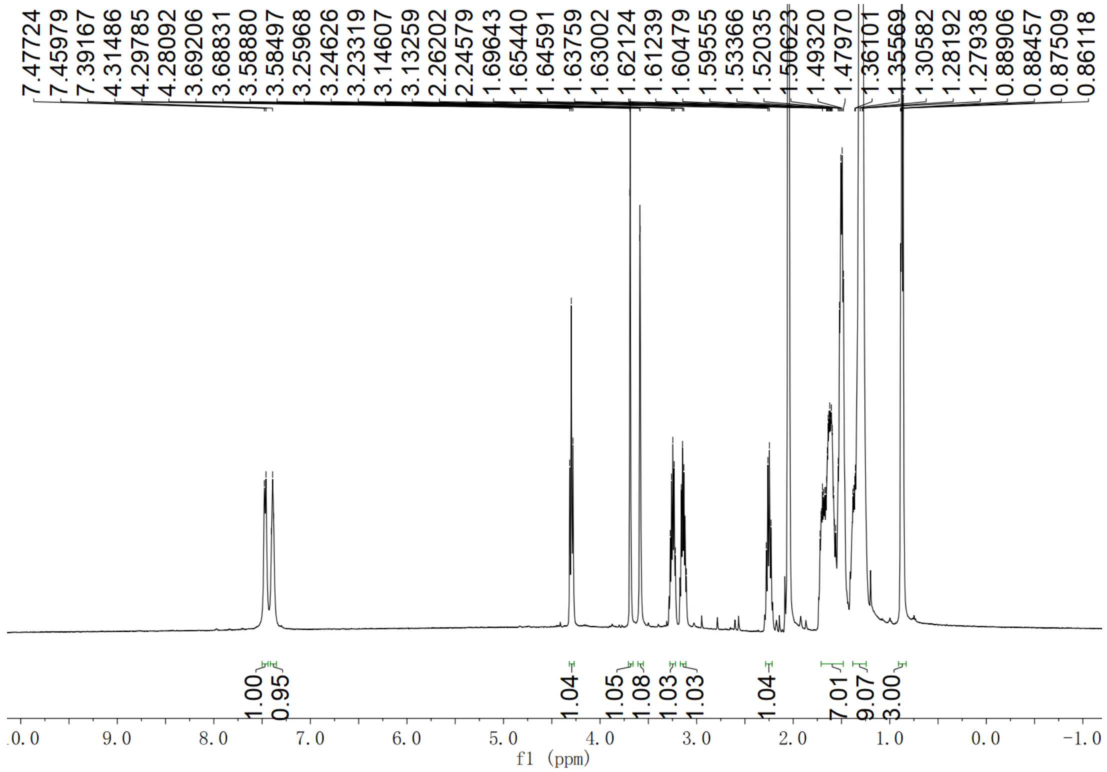

**Supplementary Fig. 299. $^{1}$H NMR spectrum of compound (2*S*,3*S*)-*t*-ES-a10-b13 in acetone-$d_6$**

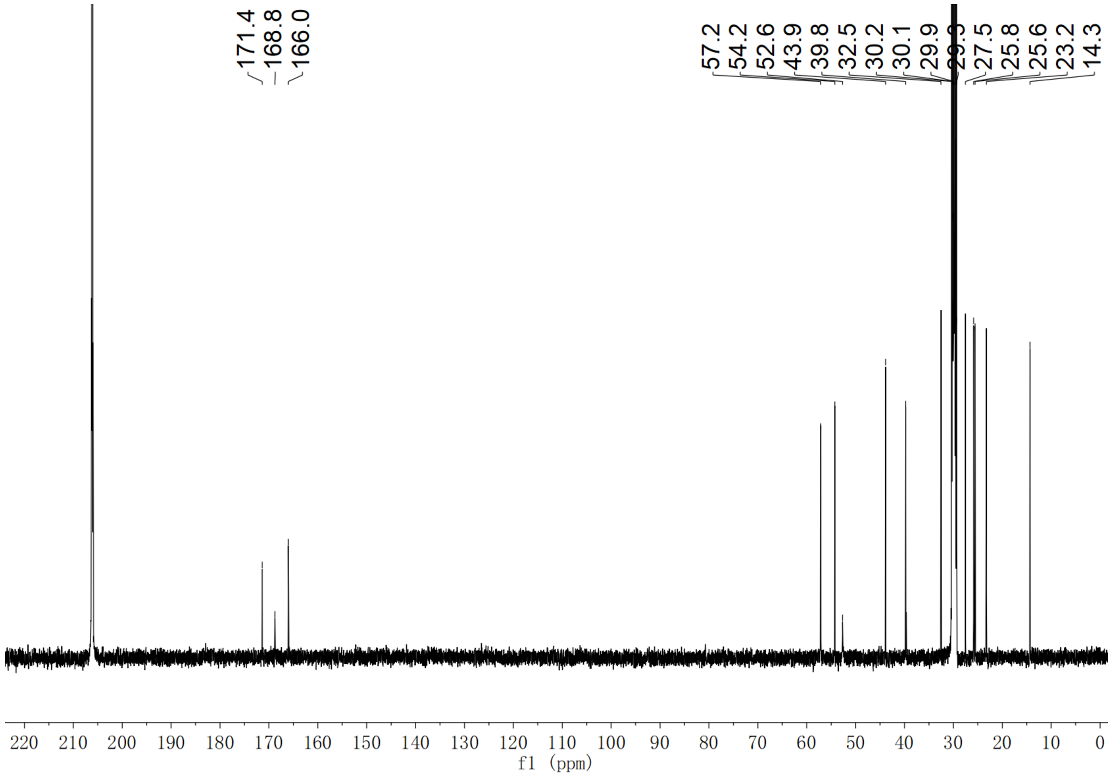

**Supplementary Fig. 300. $^{13}$C NMR spectrum of compound (2*S*,3*S*)-*t*-ES-a10-b13 in acetone-$d_6$**

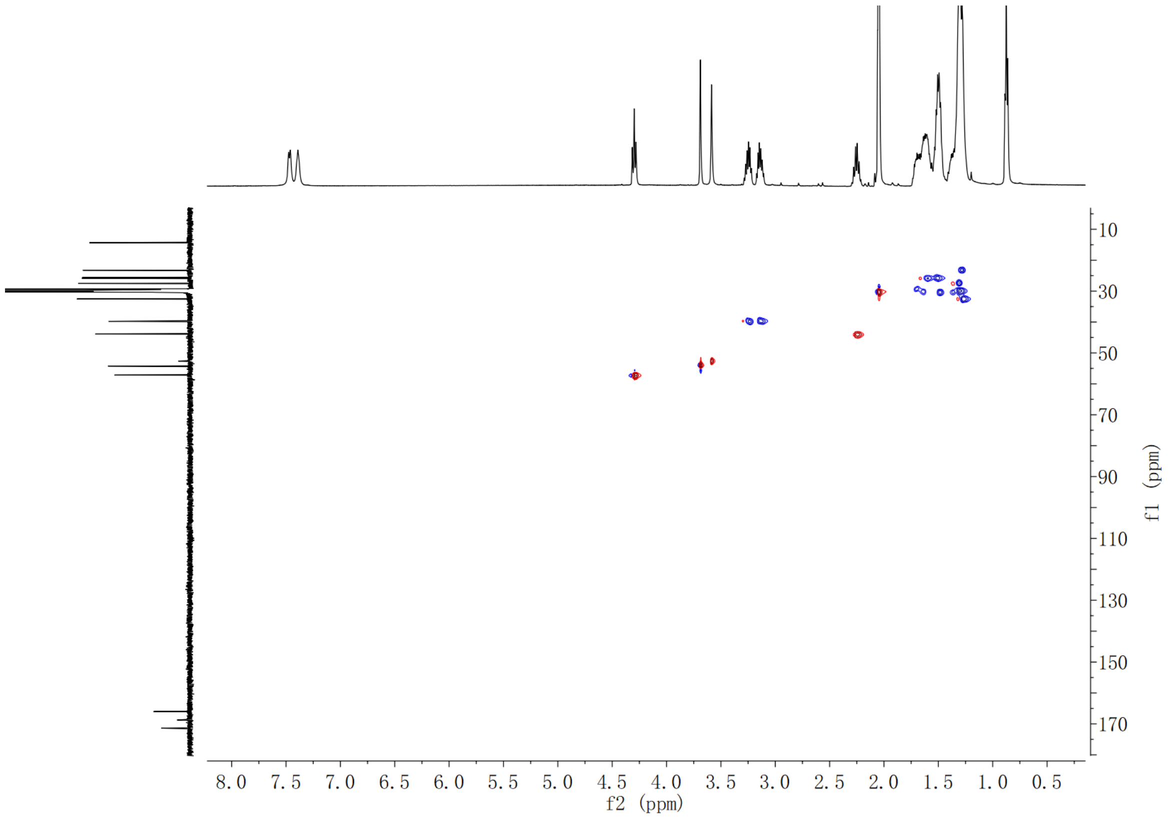

**Supplementary Fig. 301. HSQC spectrum of compound (2*S*,3*S*)-*t*-ES-a10-b13 in acetone-$d_6$**

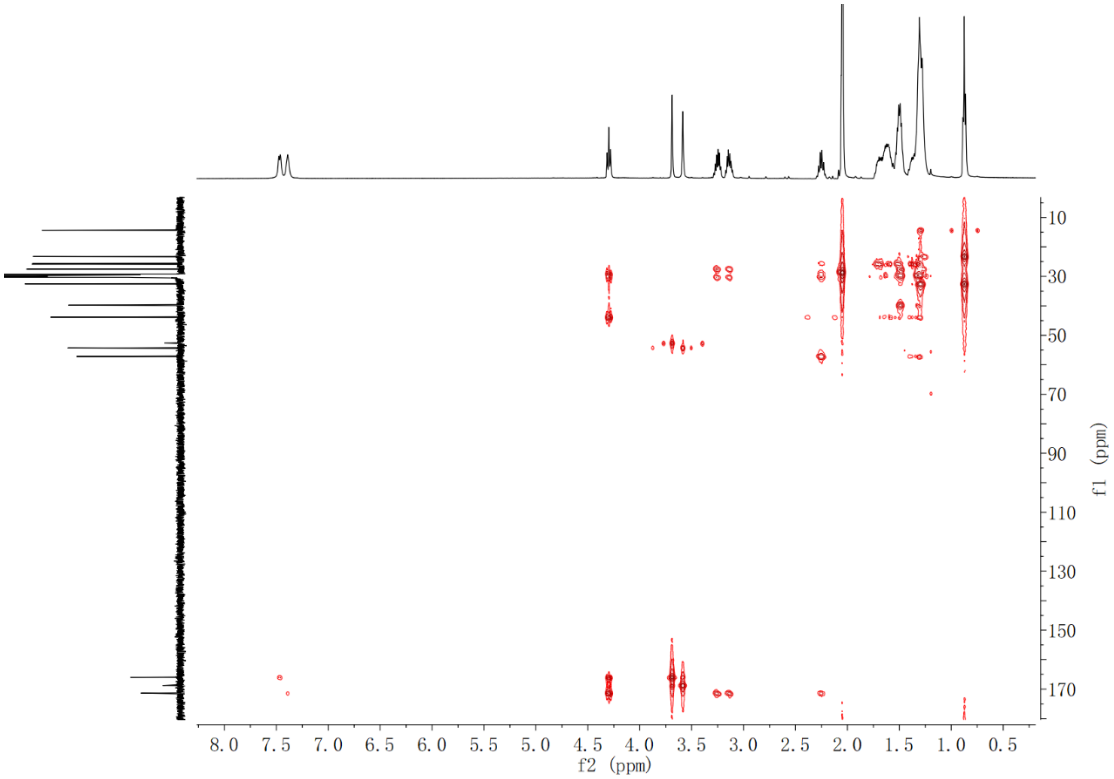

**Supplementary Fig. 302. HMBC spectrum of compound (2*S*,3*S*)-*t*-ES-a10-b13 in acetone-$d_6$**

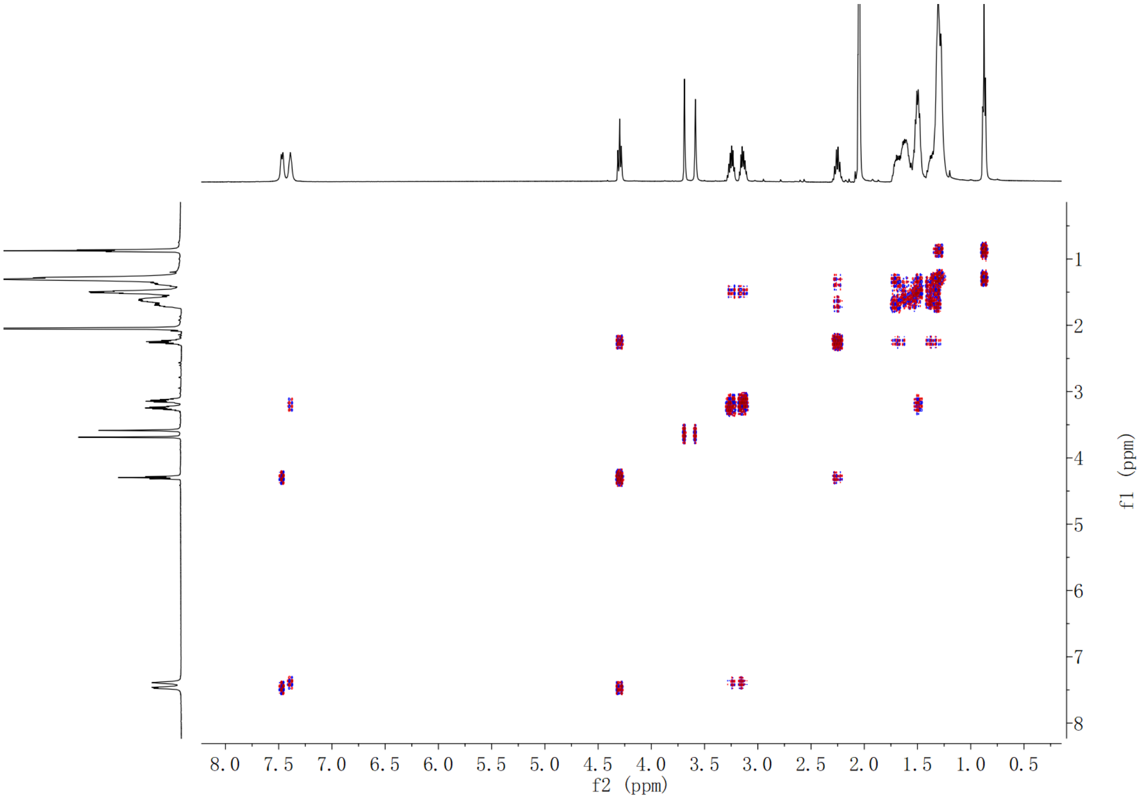

**Supplementary Fig. 303.** $^1H$-$^1H$ **COSY spectrum of compound (2*S*,3*S*)-*t*-ES-a10-b13 in acetone-$d_6$**

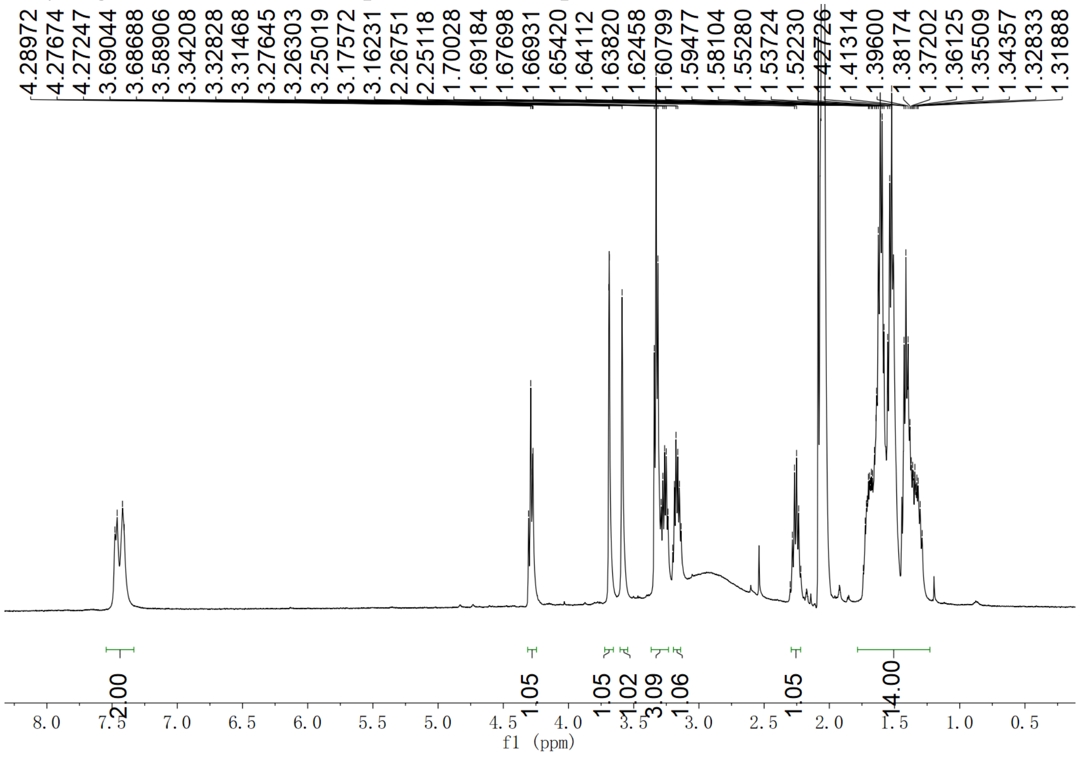

**Supplementary Fig. 304.** $^1H$ **NMR spectrum of compound (2*S*,3*S*)-*t*-ES-a10-b26 in acetone-$d_6$**

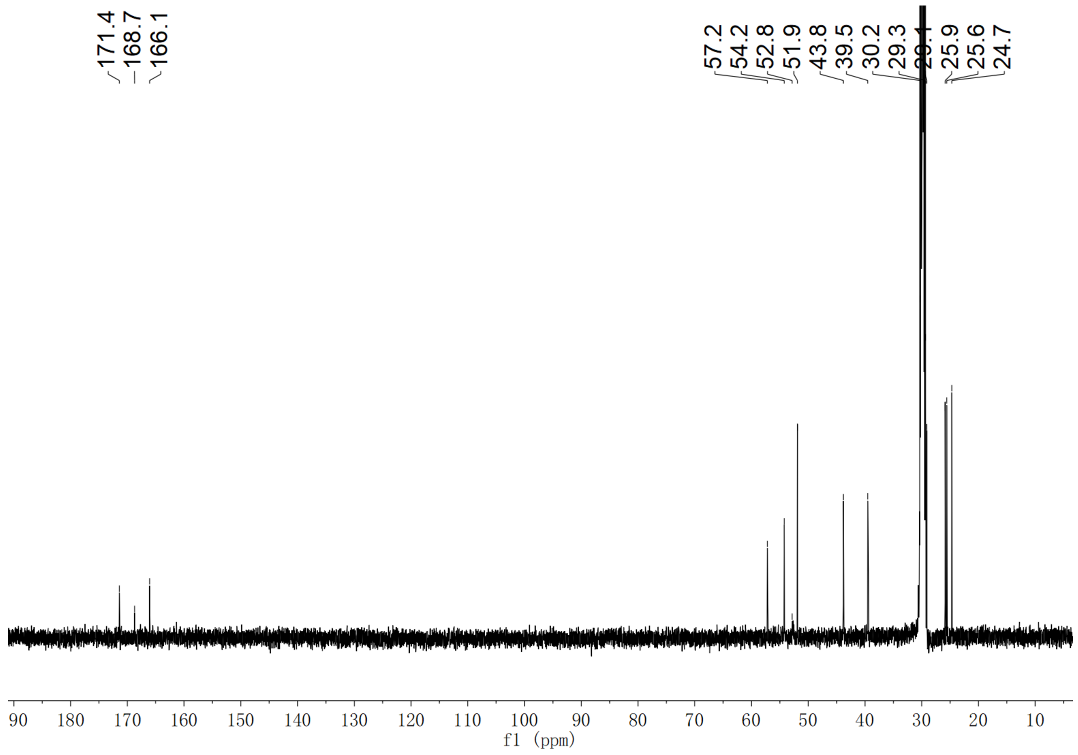

**Supplementary Fig. 305. $^{13}$C NMR spectrum of compound (2*S*,3*S*)-*t*-ES-a10-b26 in acetone-$d_6$**

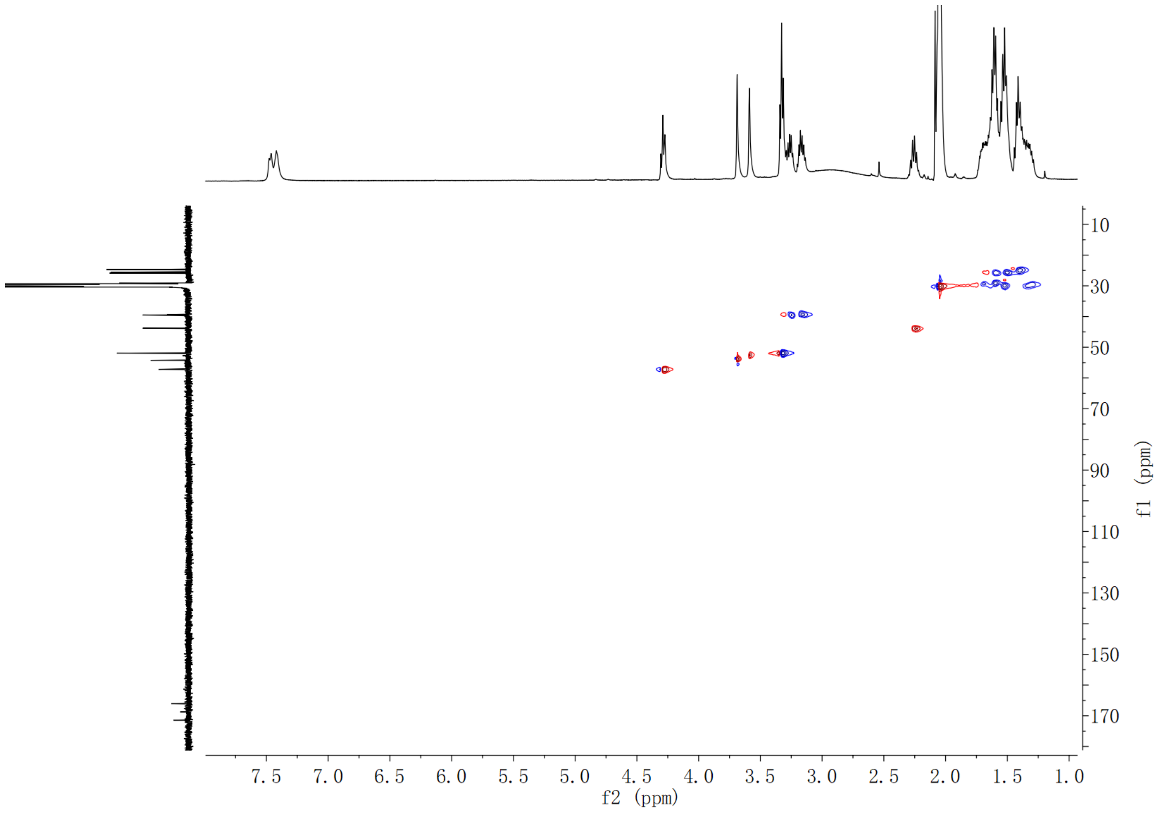

**Supplementary Fig. 306. HSQC spectrum of compound (2*S*,3*S*)-*t*-ES-a10-b26 in acetone-$d_6$**

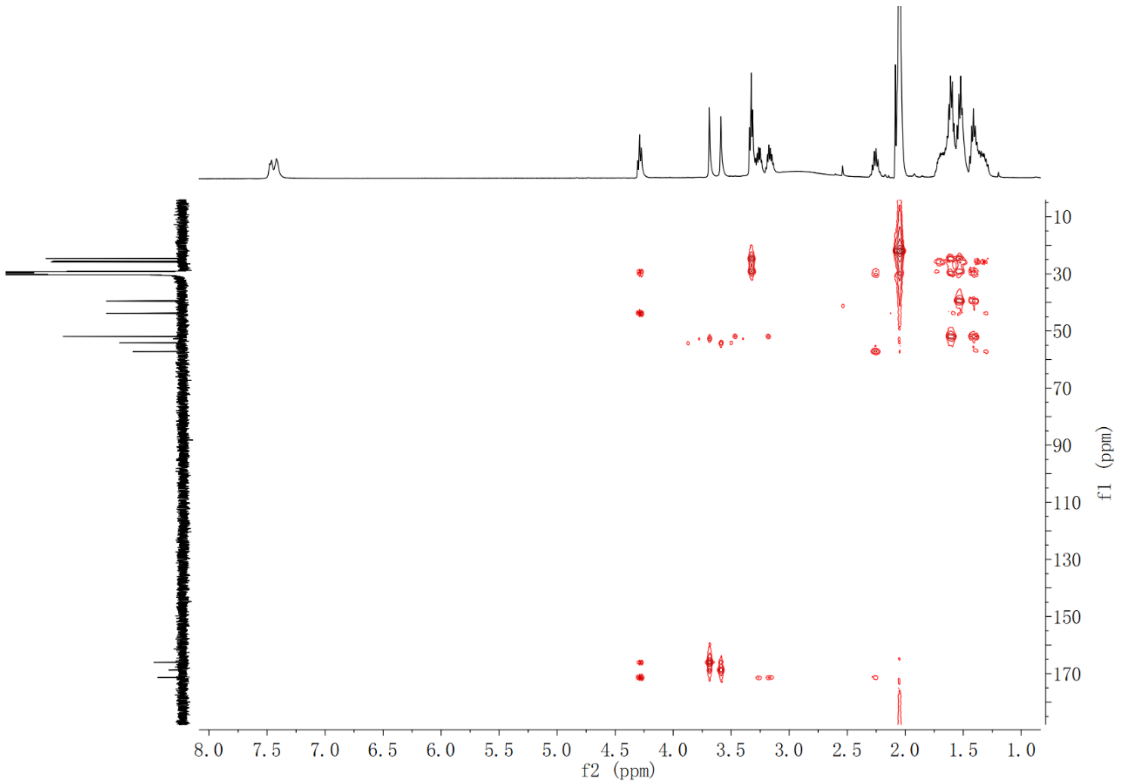

**Supplementary Fig. 307. HMBC spectrum of compound (2*S*,3*S*)-*t*-ES-a10-b26 in acetone-$d_6$**

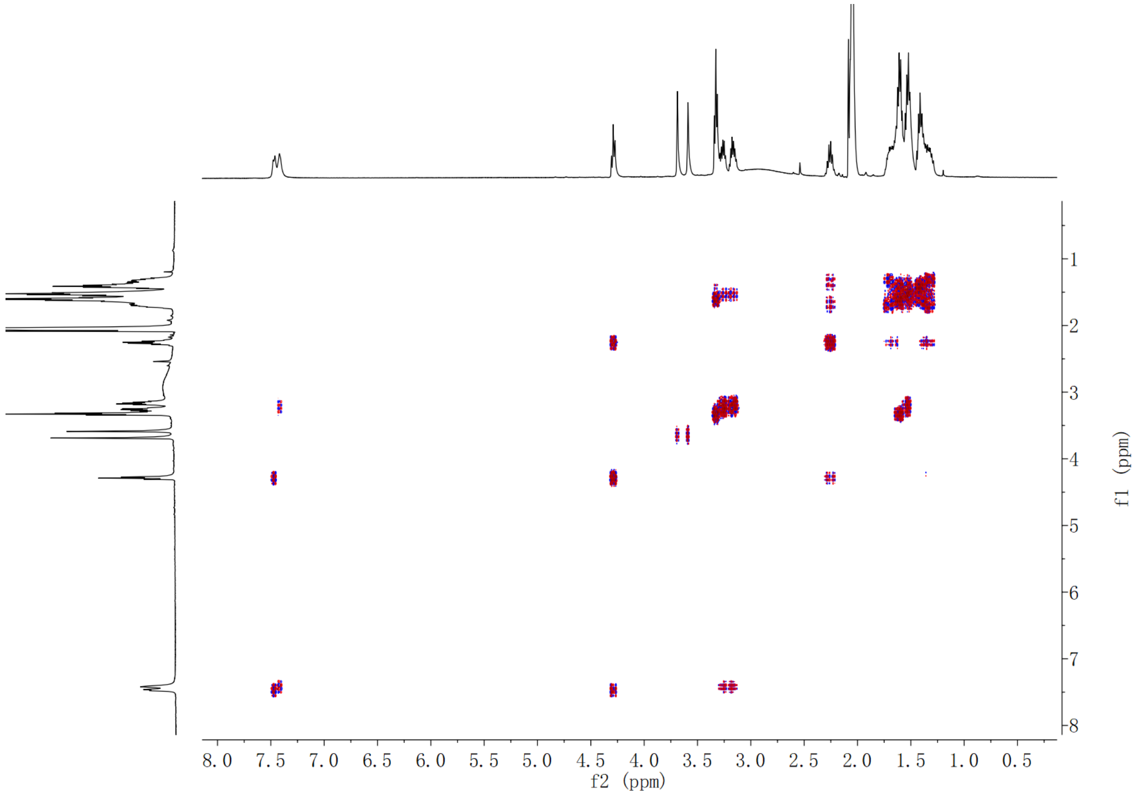

**Supplementary Fig. 308. $^1$H-$^1$H COSY spectrum of compound (2*S*,3*S*)-*t*-ES-a10-b26 in acetone-$d_6$**

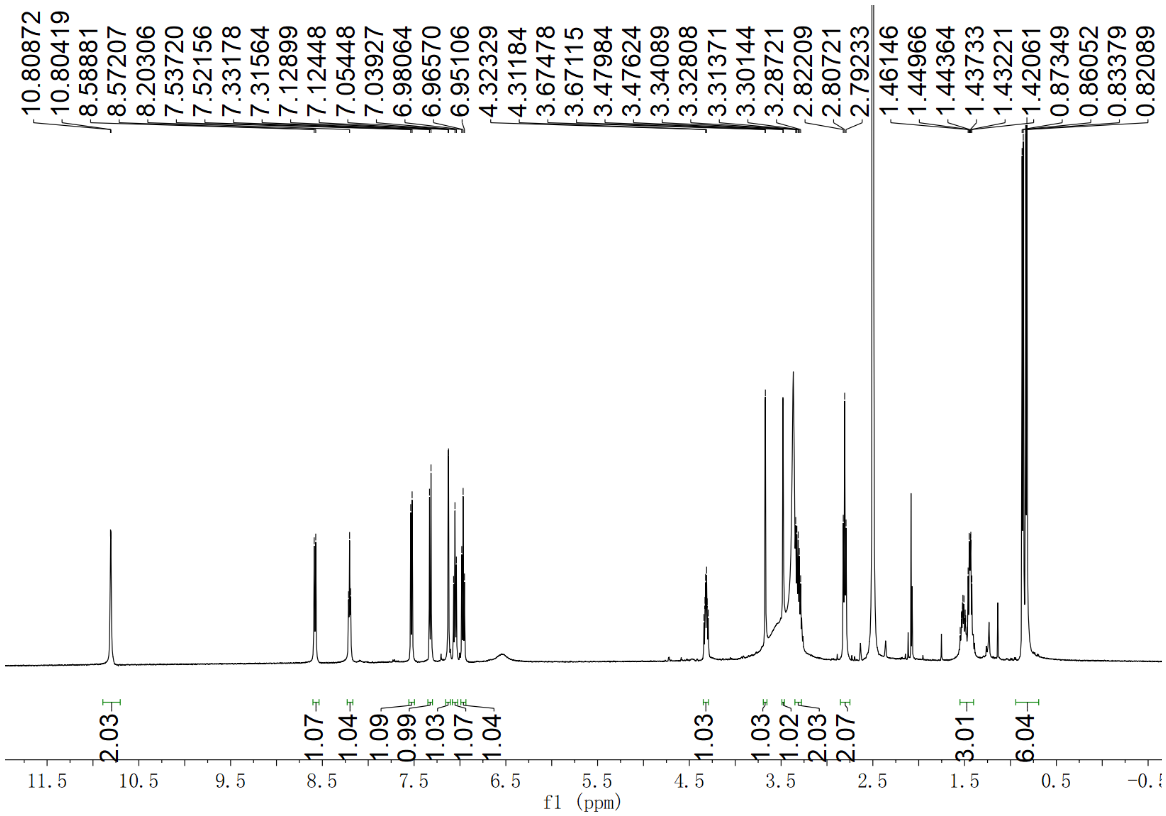

**Supplementary Fig. 309.** $^{1}$**H NMR spectrum of compound (2*S*,3*S*)-*t*-ES-Leu-b43 in DMSO-*d*$_6$**

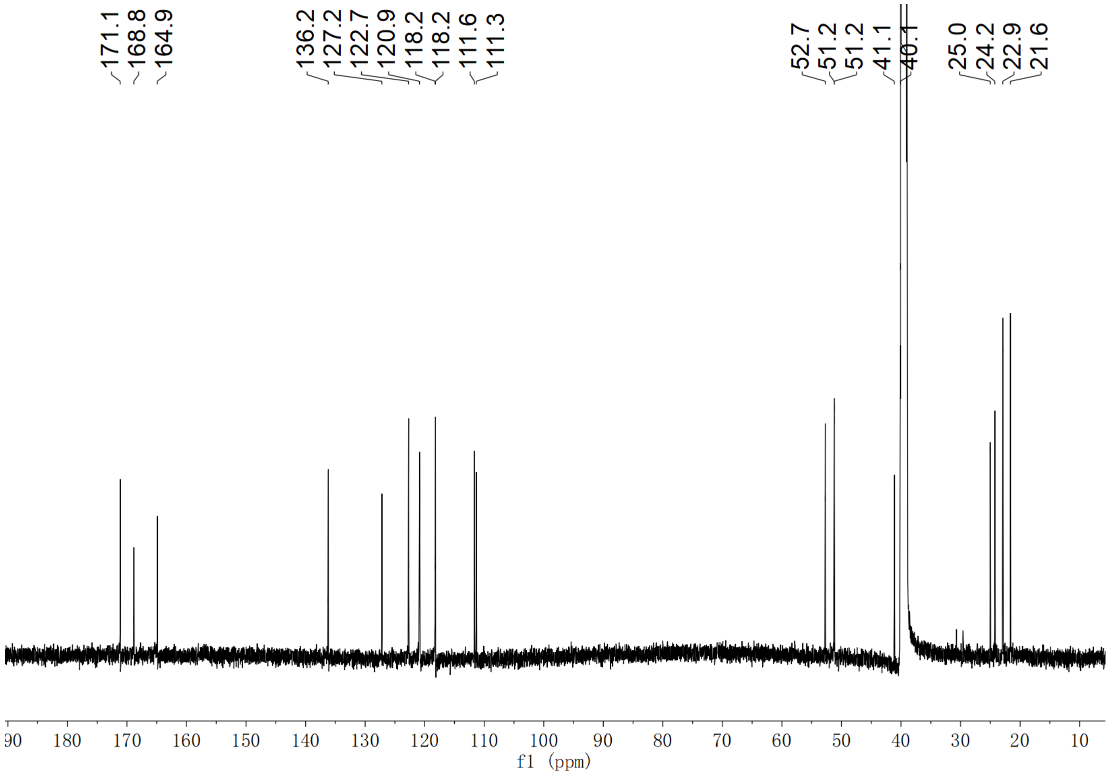

**Supplementary Fig. 310.** $^{13}$**C NMR spectrum of compound (2*S*,3*S*)-*t*-ES-Leu-b43 in DMSO-*d*$_6$**

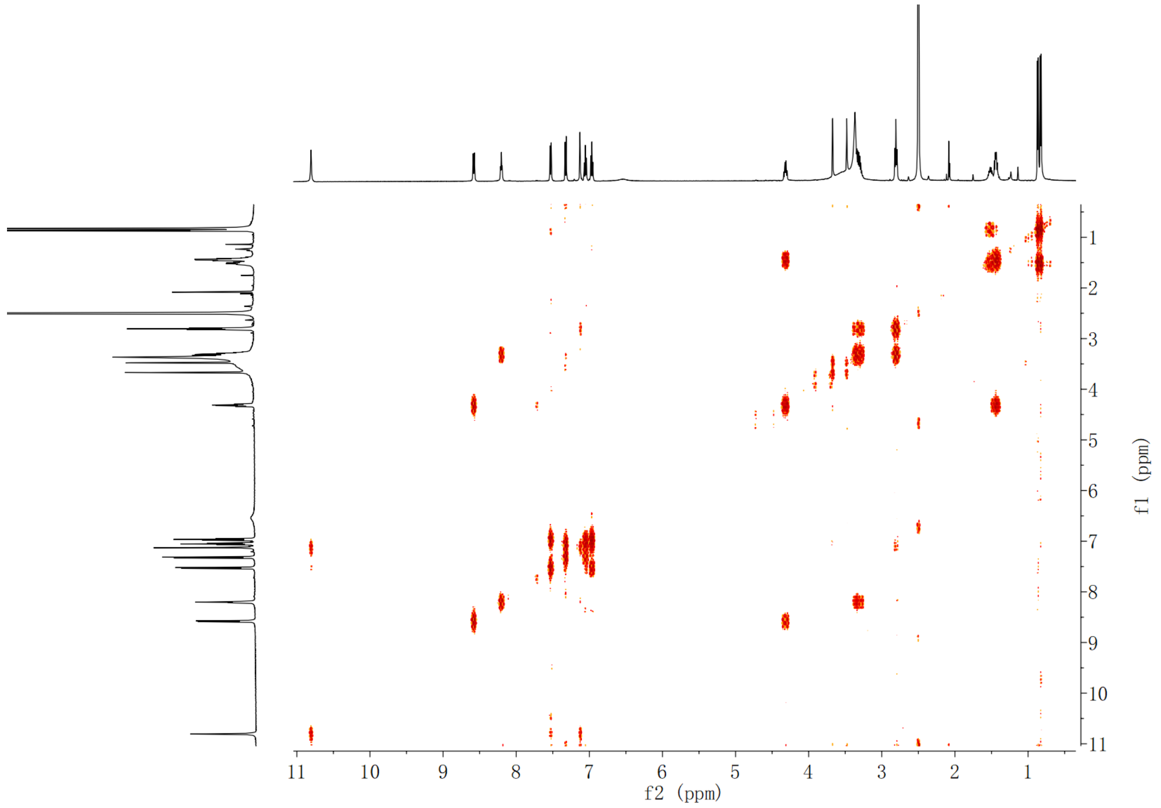

**Supplementary Fig. 311. $^1$H-$^1$H COSY spectrum of compound (2*S*,3*S*)-*t*-ES-Leu-b43 in DMSO-*d*$_6$**

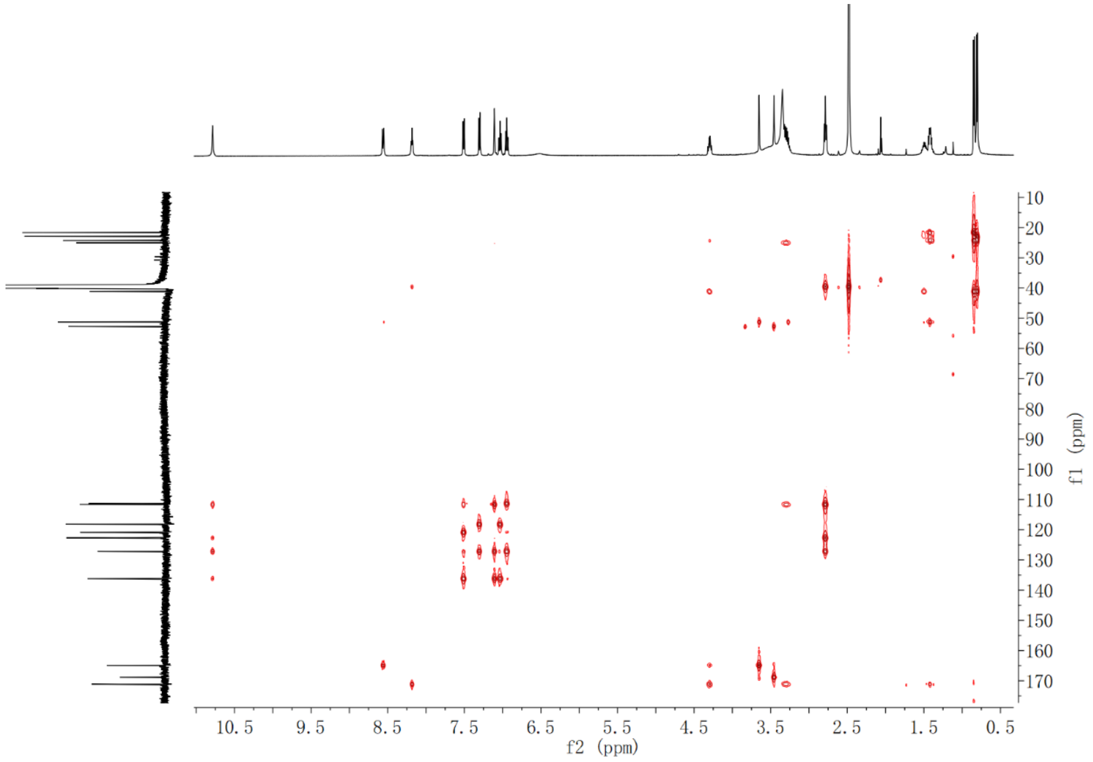

**Supplementary Fig. 312. HMBC spectrum of compound (2*S*,3*S*)-*t*-ES-Leu-b43 in DMSO-*d*$_6$**

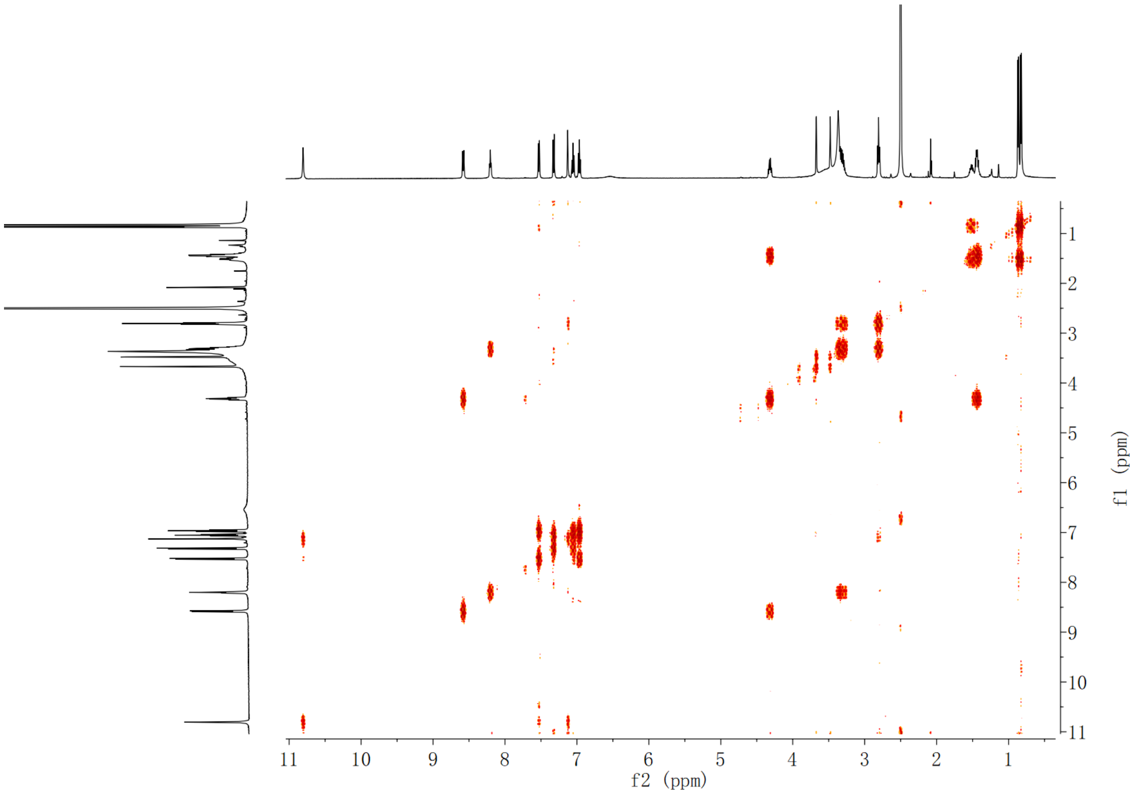

**Supplementary Fig. 313. [1]H-[1]H COSY spectrum of compound (2*S*,3*S*)-*t*-ES-Leu-b43 in DMSO-*d*$_6$**

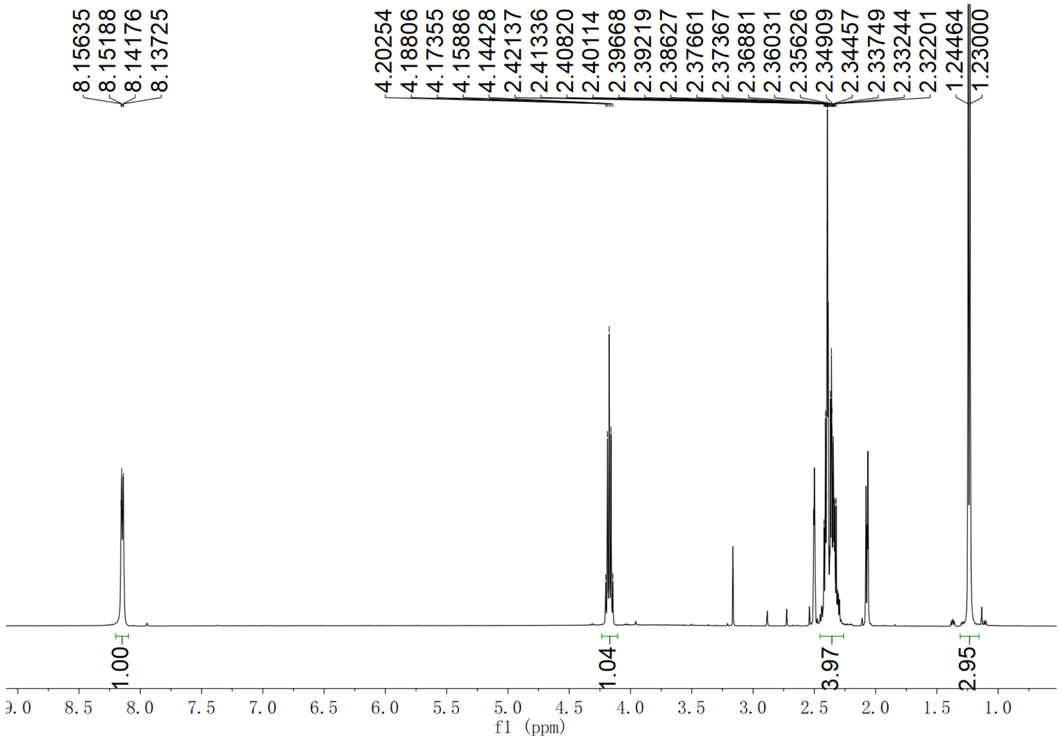

**Supplementary Fig. 314. [1]H NMR spectrum of compound *N*-succinyl-L-alanine in DMSO-*d*$_6$**

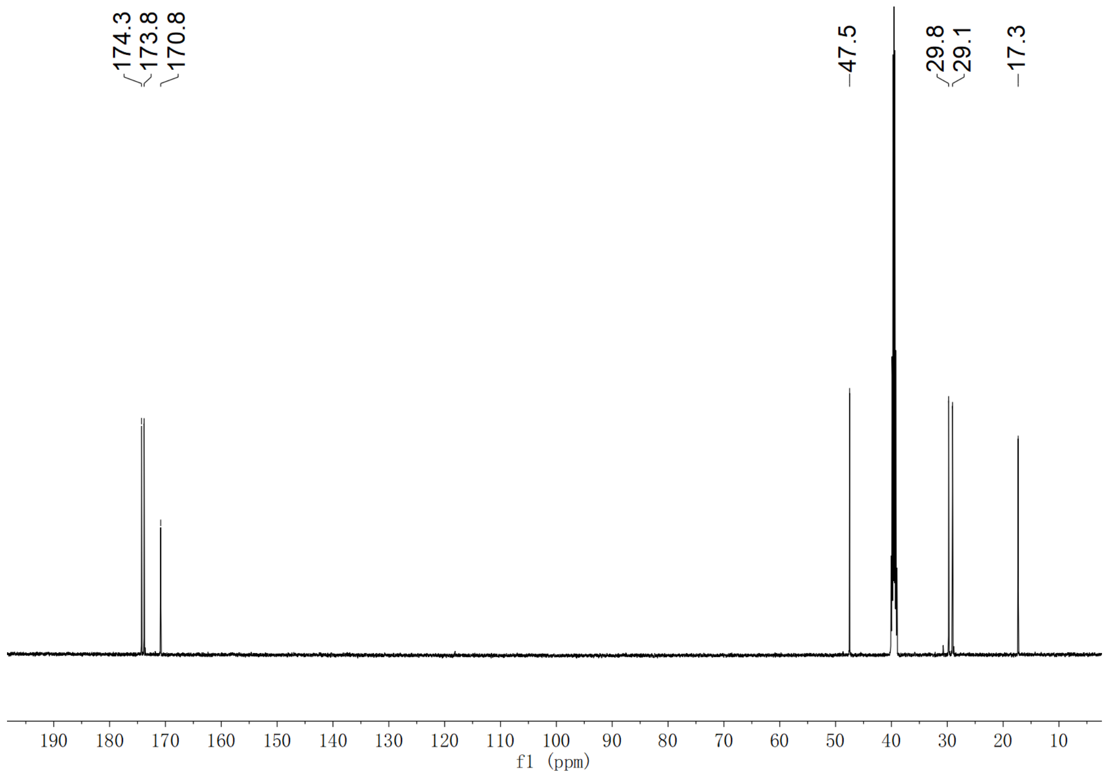

**Supplementary Fig. 315.** $^{13}$**C NMR spectrum of compound *N*-succinyl-L-alanine in DMSO-*d*$_6$**

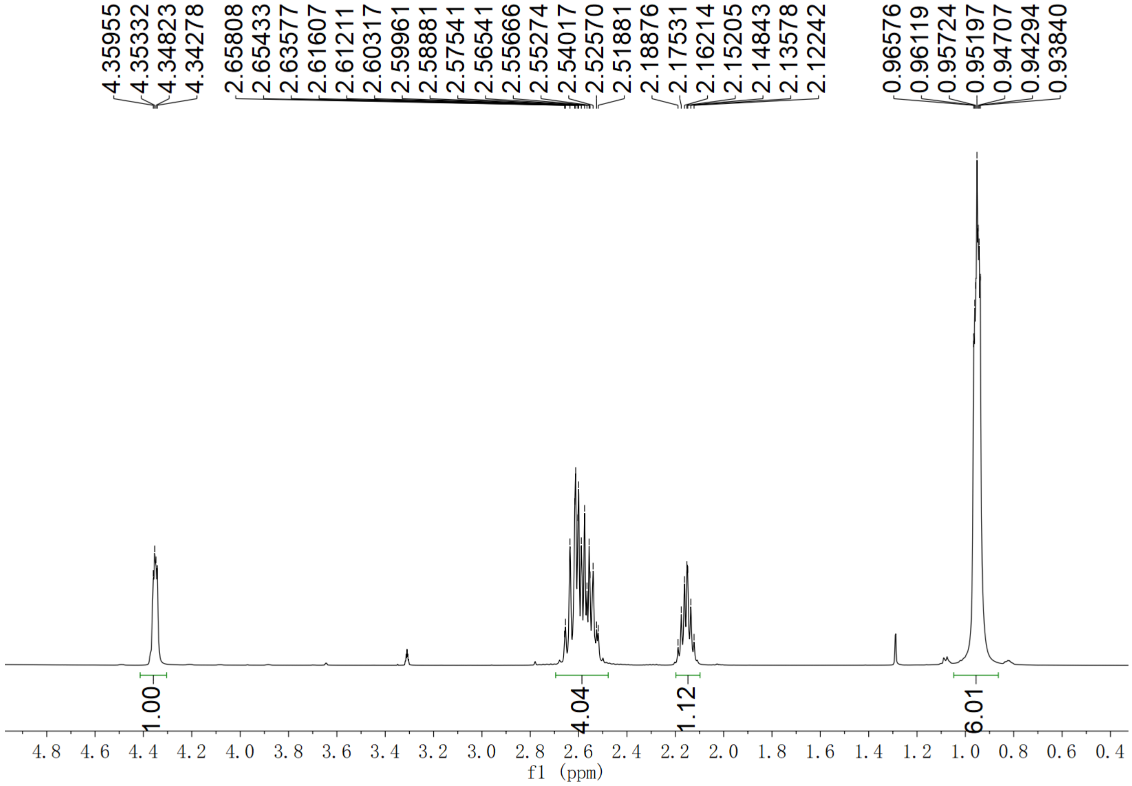

**Supplementary Fig. 316.** $^{1}$**H NMR spectrum of compound *N*-succinyl-L-valine in $CD_3OD$**

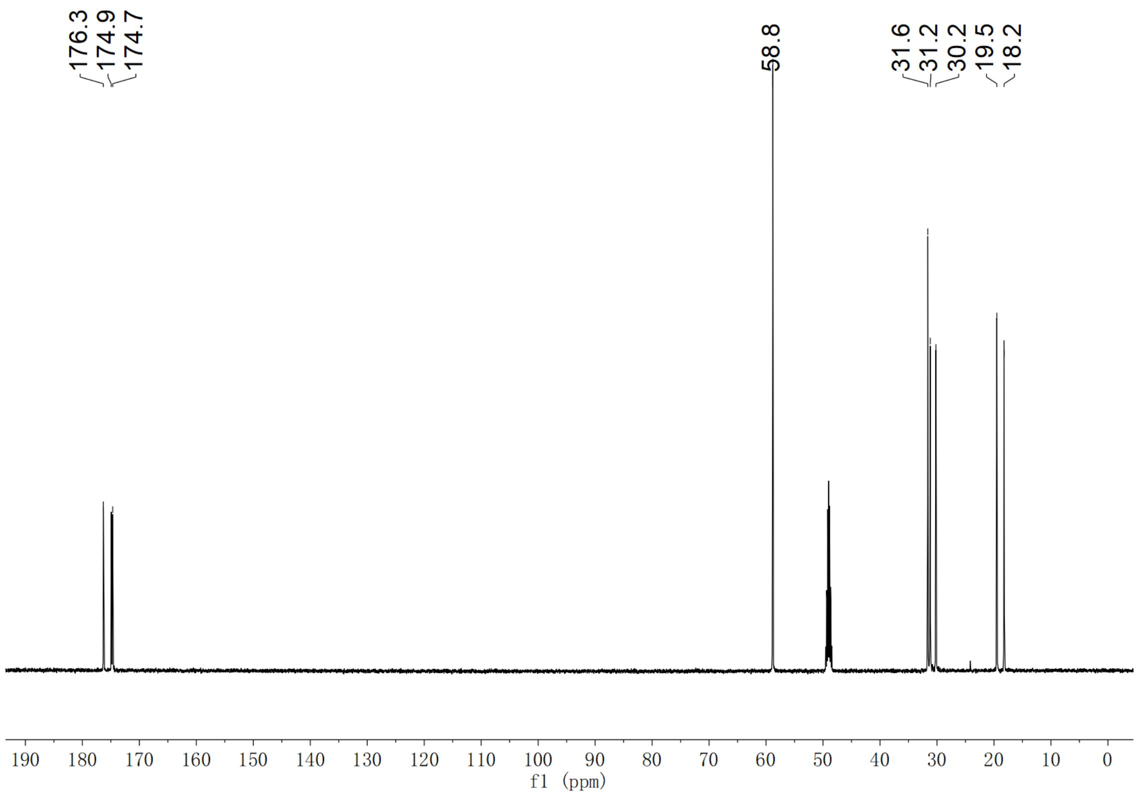

**Supplementary Fig. 317. $^{13}$C NMR spectrum of compound *N*-succinyl-L-valine in $CD_3OD$**

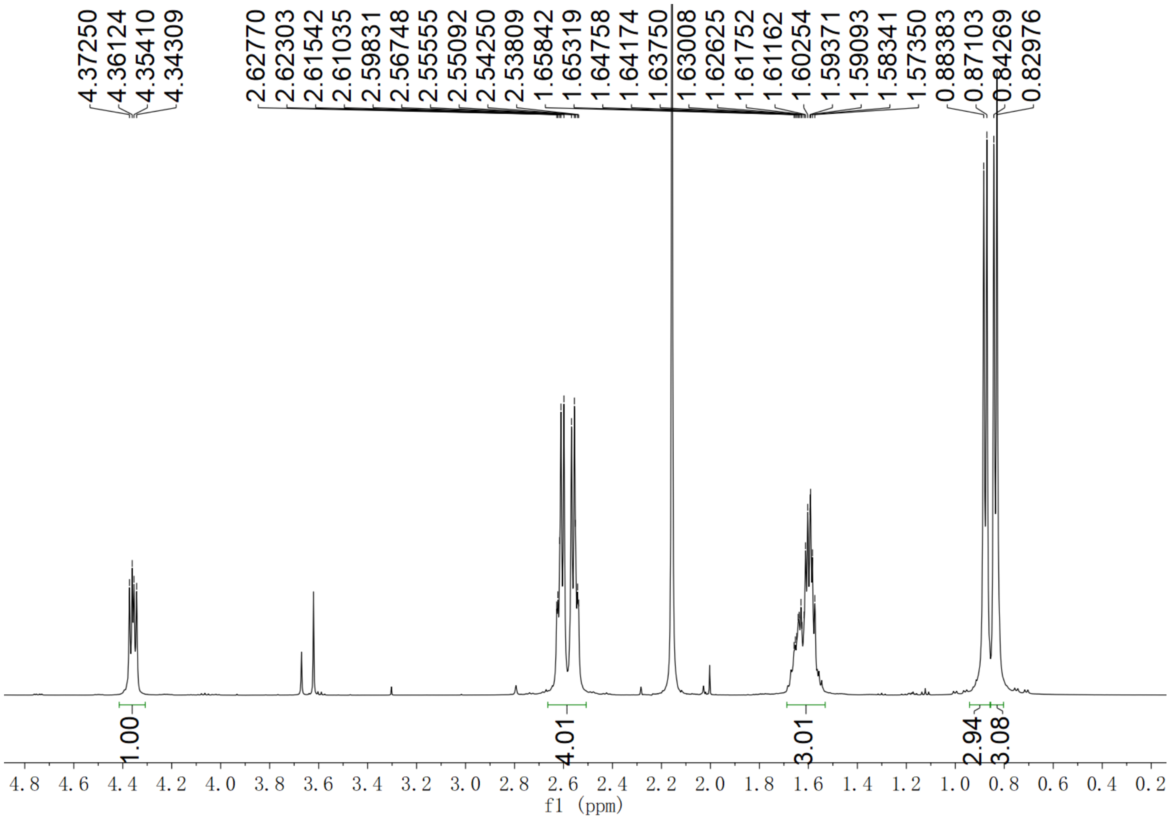

**Supplementary Fig. 318. $^{1}$H NMR spectrum of compound *N*-succinyl-L-leucine in $D_2O$**

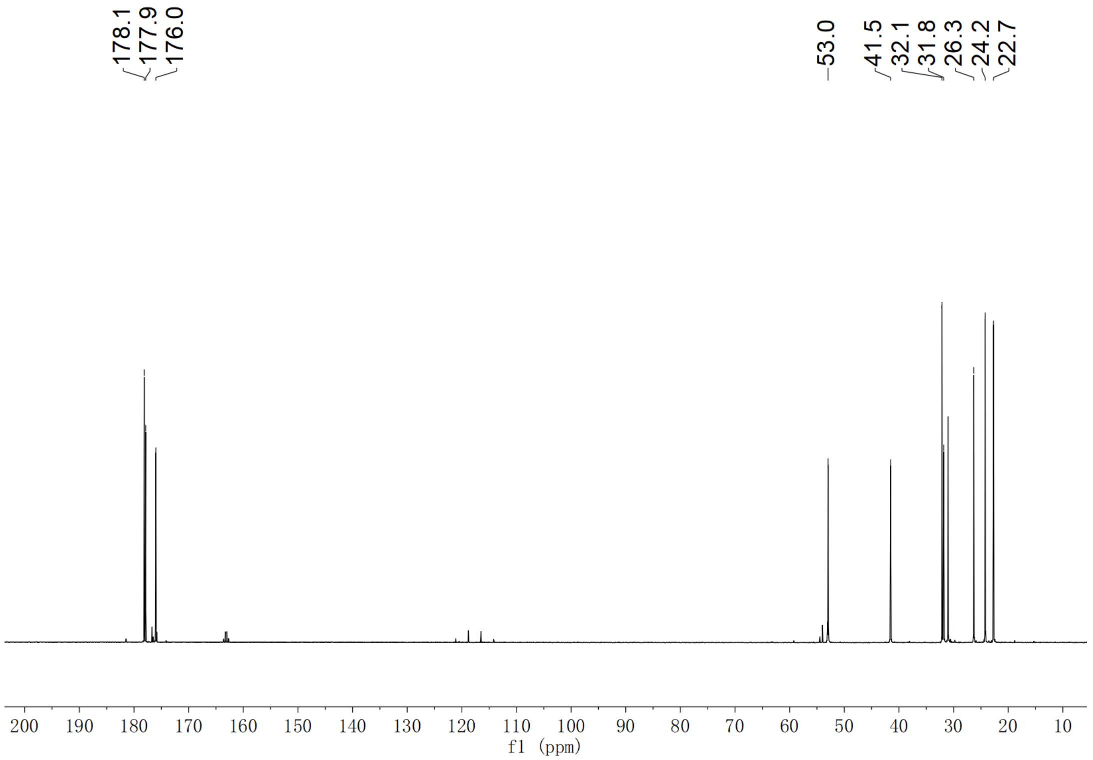

**Supplementary Fig. 319. $^{13}$C NMR spectrum of compound *N*-succinyl-L-leucine in $D_2O$**

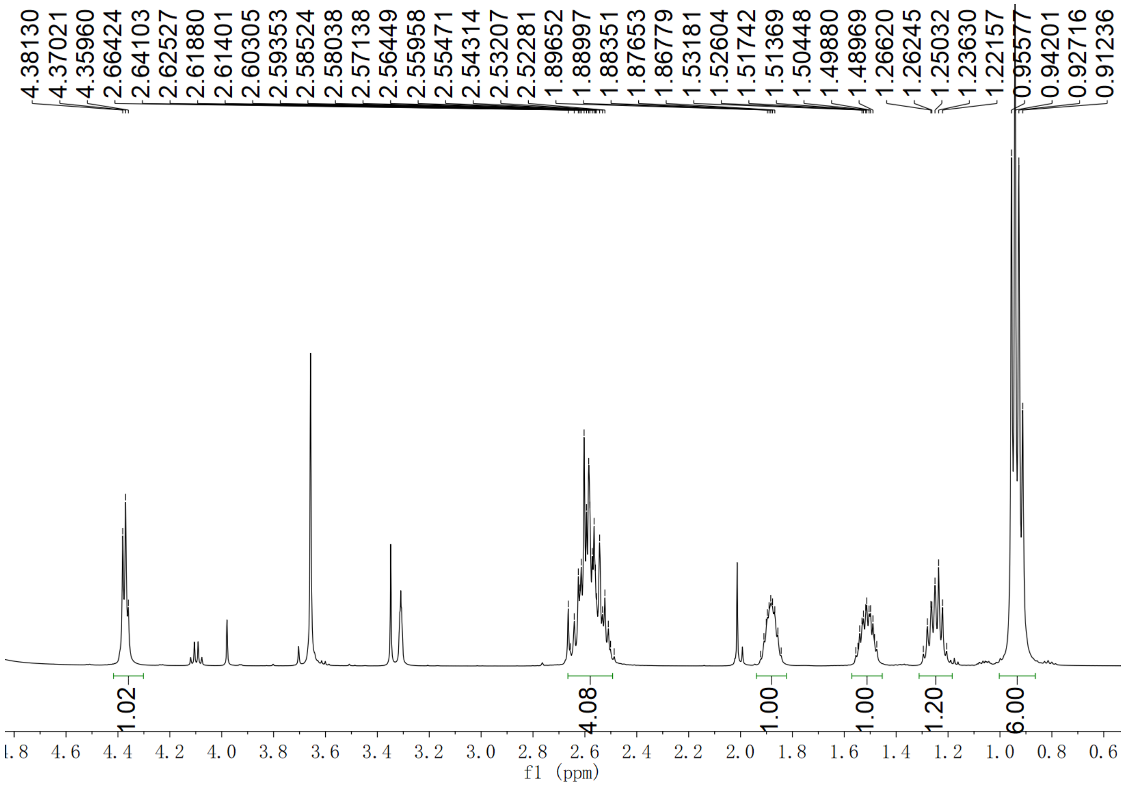

**Supplementary Fig. 320. $^{1}$H NMR spectrum of compound *N*-succinyl-L-isoleucine in $CD_3OD$**

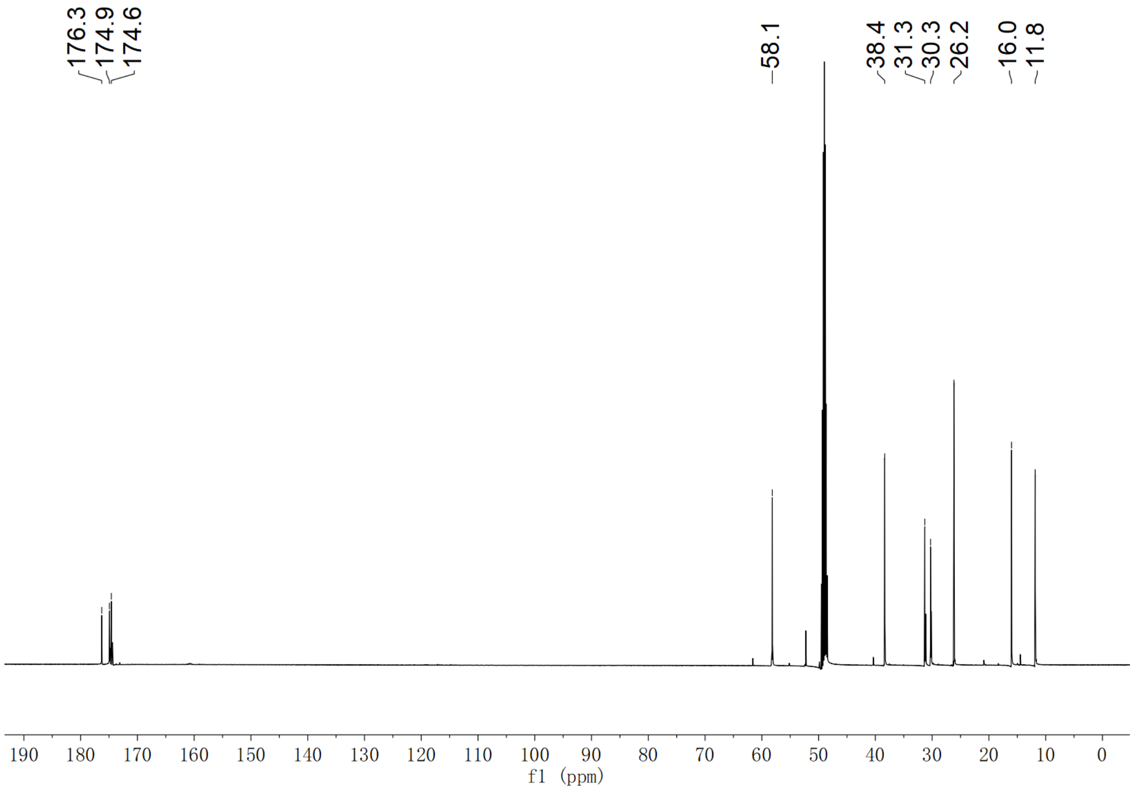

Supplementary Fig. 321. $^{13}$C NMR spectrum of compound *N*-succinyl-L-isoleucine in $CD_3OD$

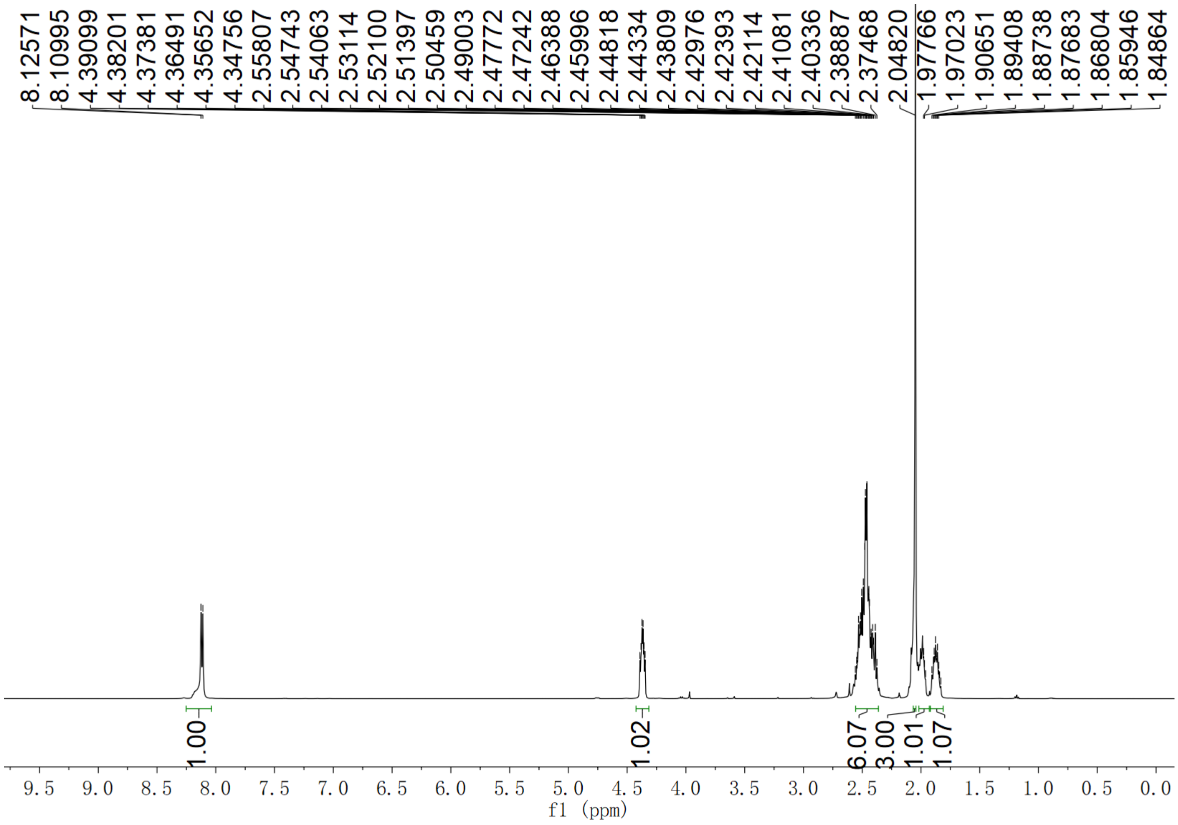

Supplementary Fig. 322. $^{1}$H NMR spectrum of compound *N*-succinyl-L-methionine in DMSO-$d_6$

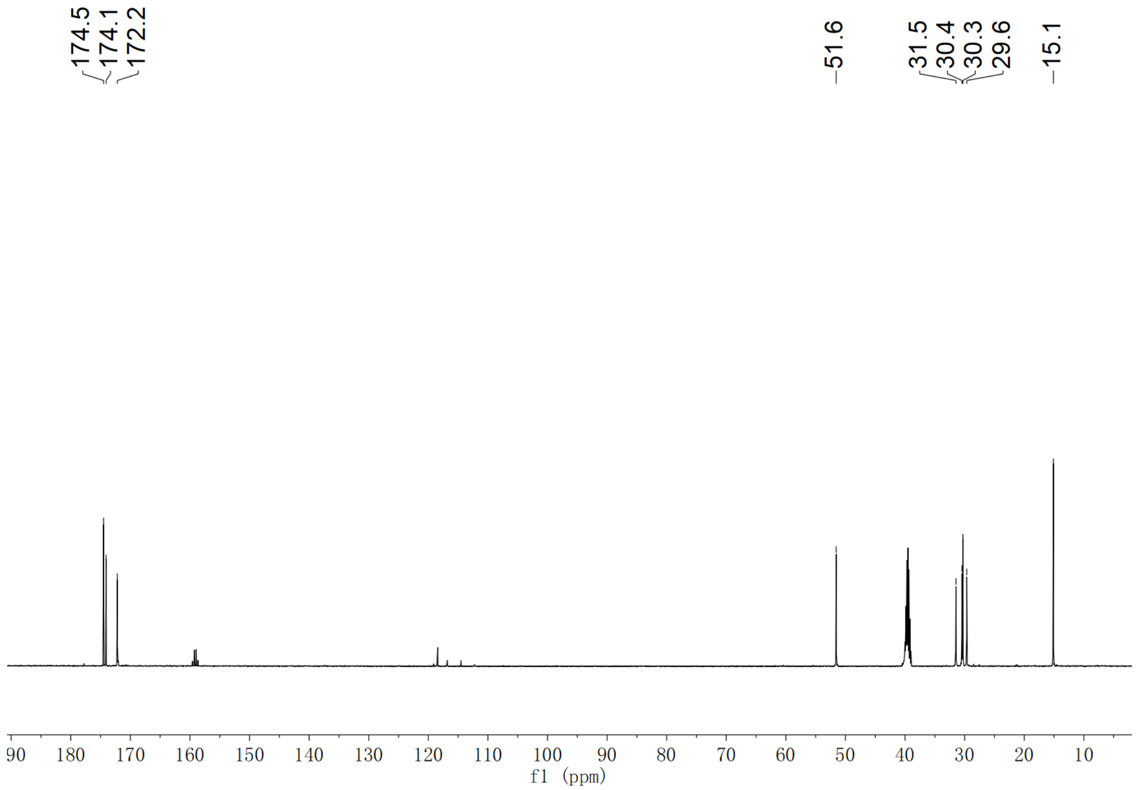

**Supplementary Fig. 323.** $^{13}$**C NMR spectrum of compound** ***N*****-succinyl-L-methionine in DMSO-*d*$_6$**

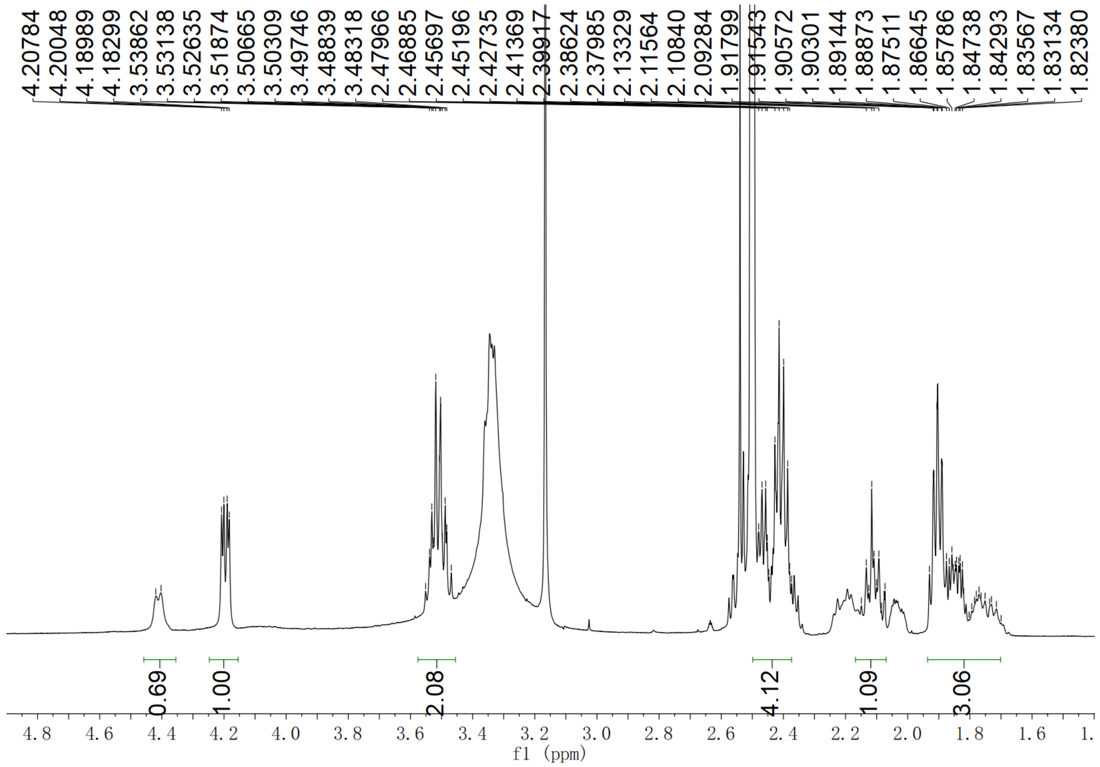

**Supplementary Fig. 324.** $^{1}$**H NMR spectrum of compound** ***N*****-succinyl-L-proline in DMSO-*d*$_6$**

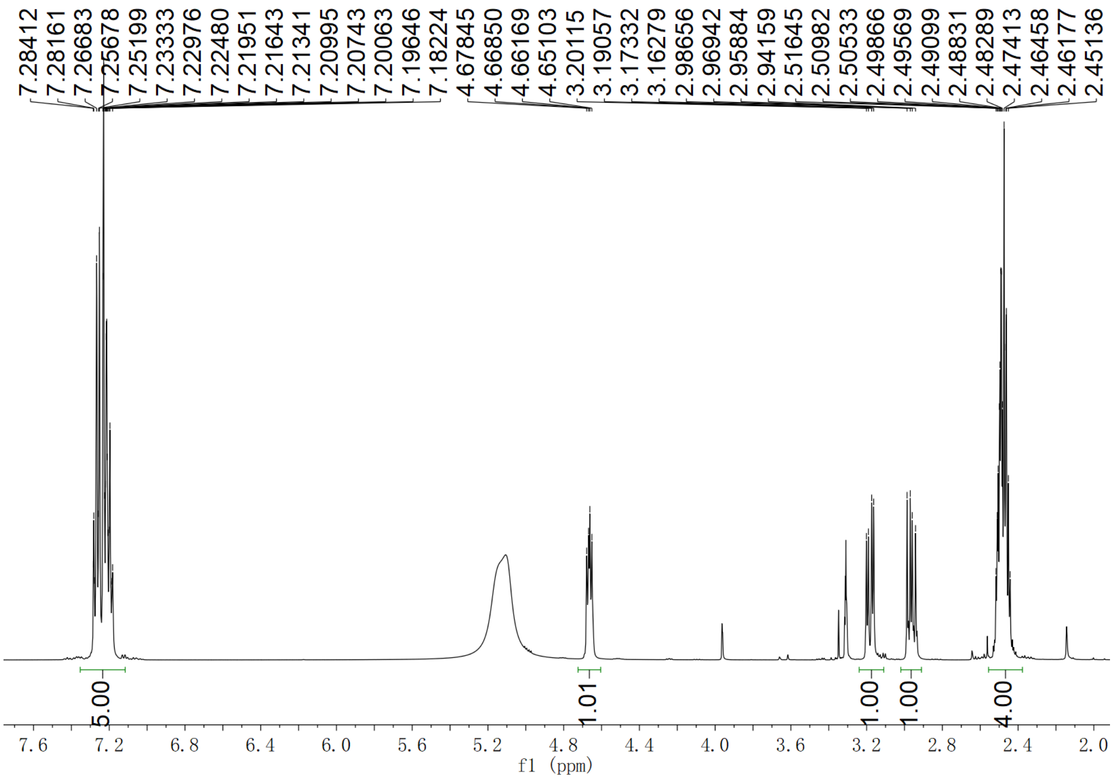

**Supplementary Fig. 325.** **$^{1}$H NMR spectrum of compound *N*-succinyl-L-phenylalanine in $CD_3OD$**

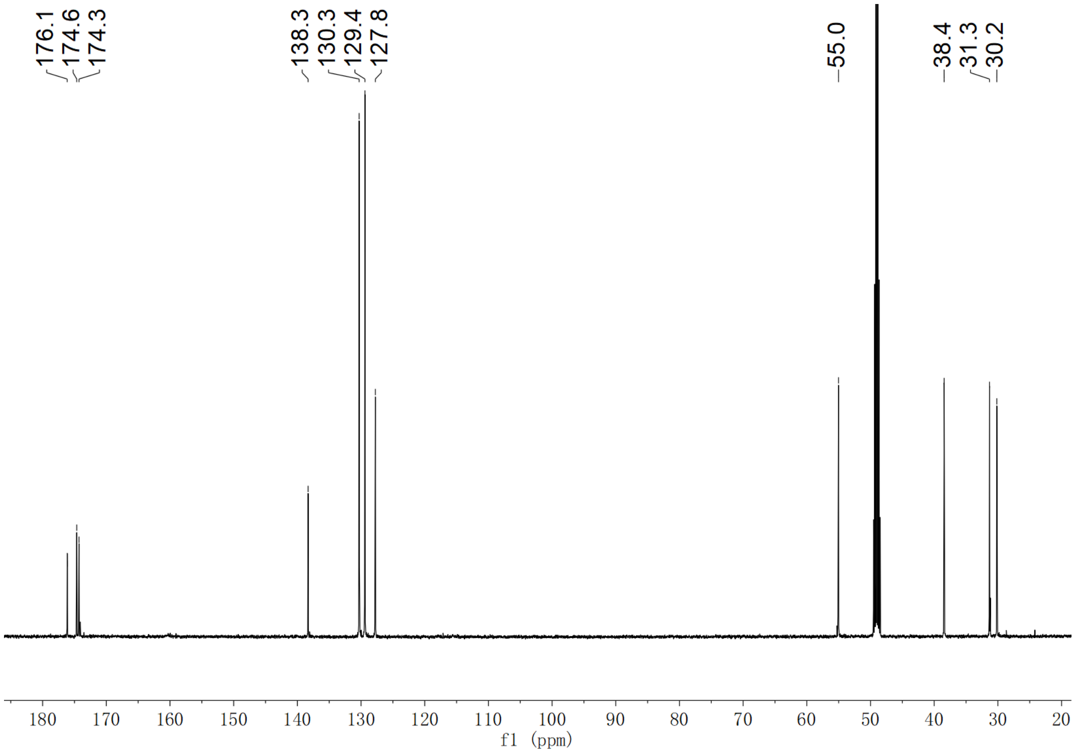

**Supplementary Fig. 326.** **$^{13}$C NMR spectrum of compound *N*-succinyl-L-phenylalanine in $CD_3OD$**

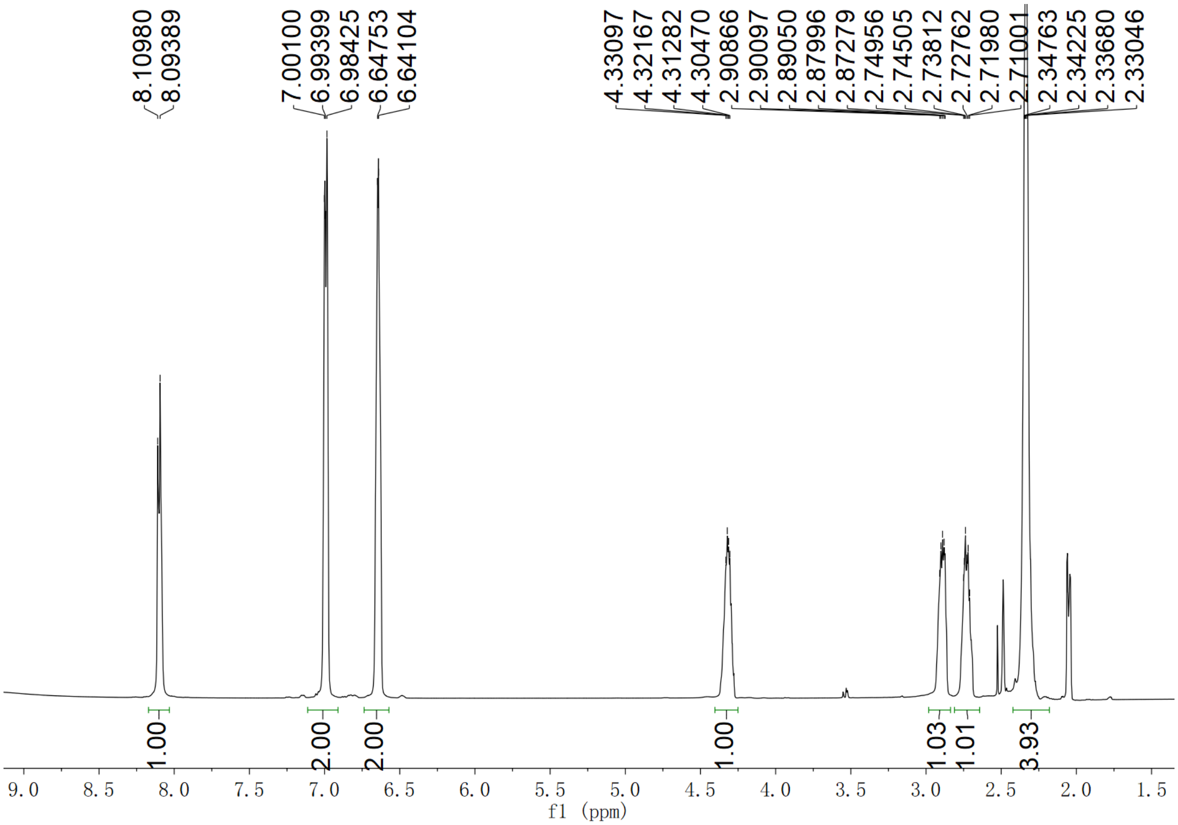

**Supplementary Fig. 327. $^{1}$H NMR spectrum of compound *N*-succinyl-L-tyrosine in DMSO-$d_6$**

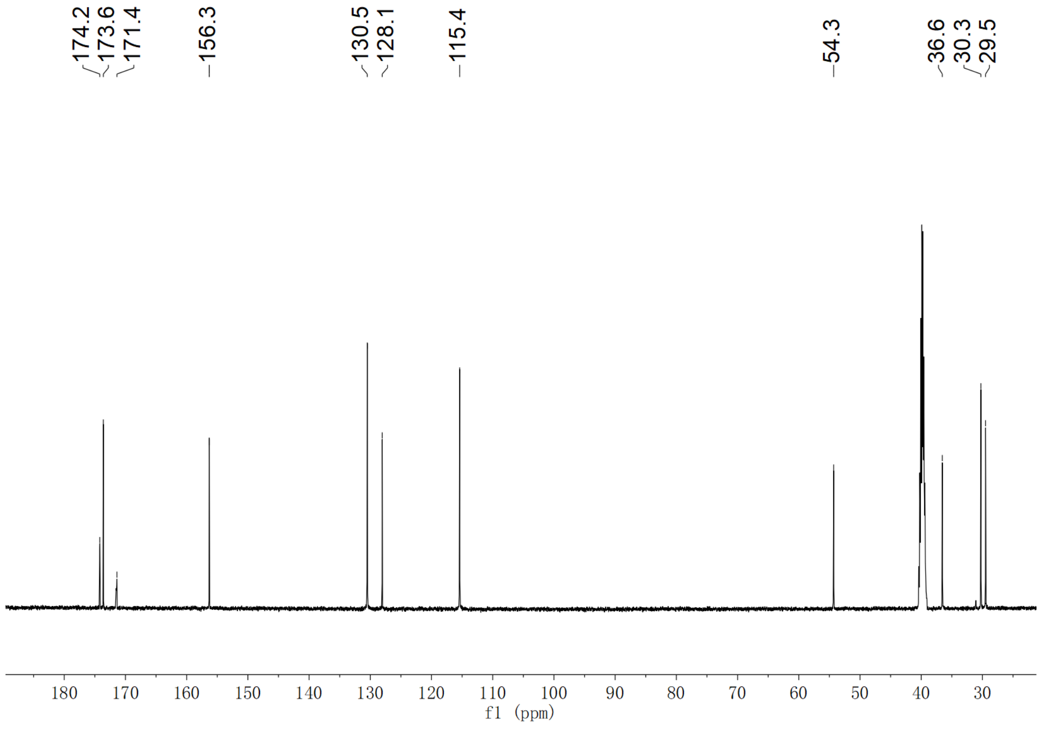

**Supplementary Fig. 328. $^{13}$C NMR spectrum of compound *N*-succinyl-L-tyrosine in DMSO-$d_6$**

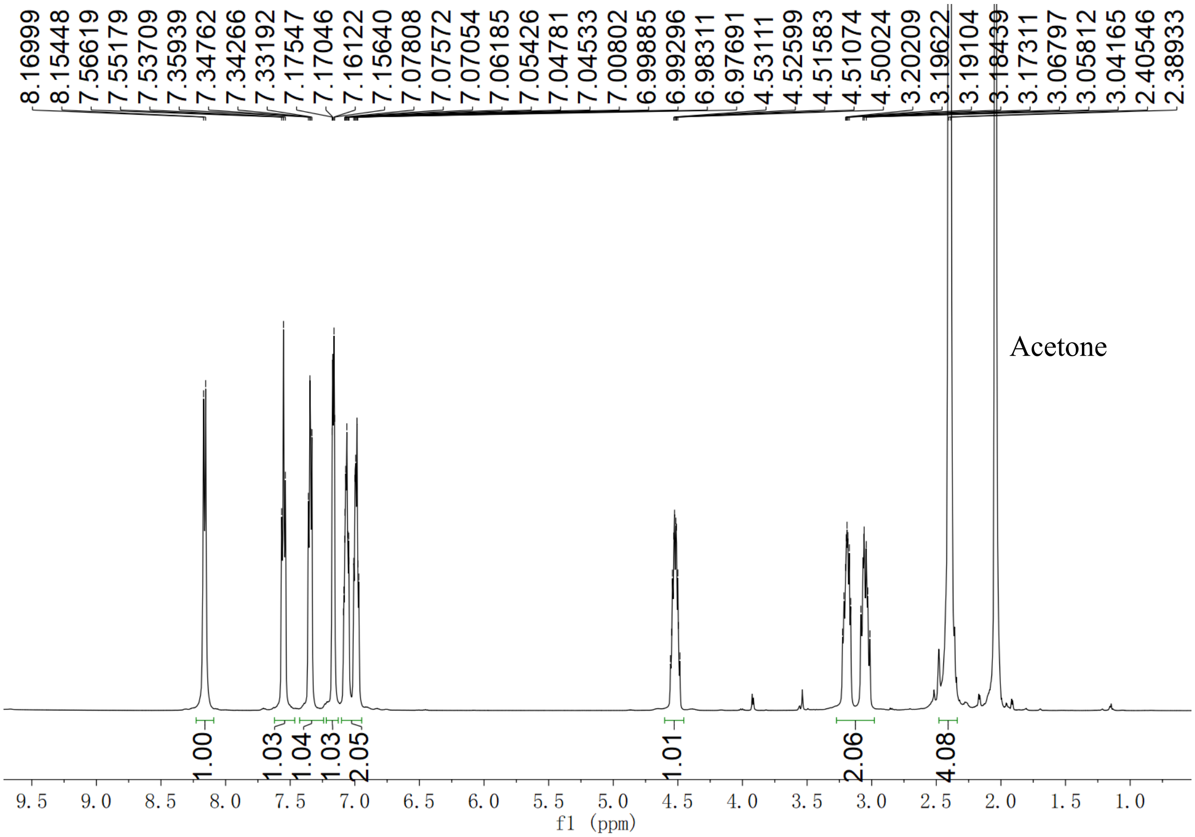

**Supplementary Fig. 329. $^{1}$H NMR spectrum of compound *N*-succinyl-L-tryptophan in DMSO-$d_6$**

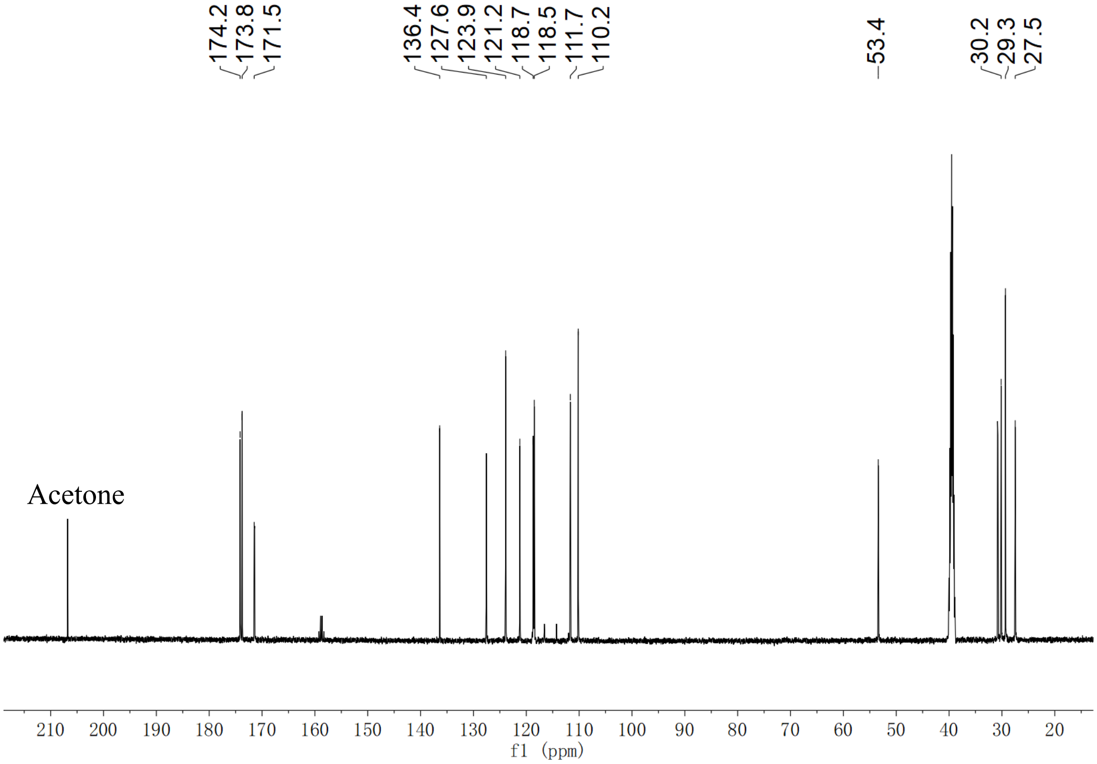

**Supplementary Fig. 330. $^{13}$C NMR spectrum of compound *N*-succinyl-L-tryptophan in DMSO-$d_6$**
